# Exome-wide analysis reveals role of *LRP1* and additional novel loci in cognition

**DOI:** 10.1101/2022.10.12.511871

**Authors:** Shreya Chakraborty, Bratati Kahali

**Affiliations:** Centre for Brain Research, Indian Institute of Science, Bangalore 560012 INDIA; Interdisciplinary Mathematical Sciences, Indian Institute of Science, Bangalore 560012 INDIA

**Author notes:** Correspondence to: Bratati Kahali, Centre for Brain Research, Indian Institute of Science, Bangalore 560012 INDIA.

**Keywords:** cognition, exome-wide association analysis, UK Biobank, metabolic risks, *APOE*, *LRP1*

## Abstract

Cognitive functioning is heritable, with metabolic risk factors known to accelerate age-associated cognitive decline. Identifying genetic underpinnings of cognition is thus crucial.

Here, we undertake single-variant and gene-based association analyses upon six neurocognitive phenotypes across six cognition domains in whole-exome sequencing data from 157,160 individuals in the UK Biobank to expound the genetic architecture of human cognition. We further identify genetic variants interacting with *APOE*, a significant genetic risk factor for cognitive decline, while controlling for lipid and glycemic risks, towards influencing cognition. Additionally, considering lipid and glycemic traits, we conduct bivariate analysis to underscore pleiotropic effects and also highlight suggestive mediation effects of metabolic risks on cognition.

We report 18 independent novel loci associated with five cognitive domains while controlling for *APOE* isoform-carrier status and metabolic risk factors. Our novel variants are mostly in genes which could also impact cognition via their functions on synaptic plasticity and connectivity, oxidative stress, neuroinflammation. Variants in or near these identified loci show genetic links to cognitive functioning in association with *APOE*, Alzheimer’s disease and related dementia phenotypes and brain morphology phenotypes, and are also eQTLs significantly controlling expression of their corresponding genes in various regions of the brain. We further report four novel pairwise interactions between exome-wide significant loci and *APOE* variants influencing episodic memory, and simple processing speed while accounting for serum lipid and serum glycemic traits. We obtain both *APOC1* and *LRP1* as significantly associated with complex processing speed and visual attention in our gene-based analysis. They also exhibit significant interaction effect with *APOE* variants in influencing visual attention. We find that variants in *APOC1* and *LRP1* act as significant eQTLs for regulating their expression in basal ganglia and cerebellar hemispheres, crucial to visual attention. Taken together, our findings suggest that *APOC1* and *LRP1* have plausible roles along pathways of amyloid-β, lipid and/or glucose metabolism in affecting visual attention and complex processing speed. Interestingly, variants in *MTFR1L, PPFIA1, PCDHB16, ATP2A1* show evidence of pleiotropy and mediation effects through serum glucose/HDL levels affecting four different cognition domains.

This is the first report from large-scale exome-wide study with evidence underscoring the effect of *LRP1* on cognition. Our research highlights a novel set of loci that augments our understanding of the genetic underpinnings of cognition during ageing, considering cooccurring metabolic conditions that can confer genetic risk to cognitive decline in addition to *APOE*, which can aid in finding causal determinants of cognitive decline.

## Introduction

Cognition refers to a plethora of mental processes that guides acquisition, transformation, storage, recovery and implementation of information and is key to good health. Understanding genetic predispositions for inter-individual differences in age-related cognitive decline is of paramount importance in healthy ageing. Genome-wide studies on cognition have shown that intelligence in humans are heritable and individual differences can be explained by genetic variations.^1–5^ Non-invasive neuropsychological cognitive assessments serve as dependable endophenotypes to assess brain functioning in healthy aging and dementia.^6,7^

The *APOE* locus, confers the highest genetic risk for Alzheimer’s dementia and is also known to be associated with nonpathological cognitive ageing.^8^ ApoE is the major apolipoprotein that plays a central role in maintaining homeostasis in the brain via transport and clearance of lipids and amyloid beta (Aβ). Several other age-associated metabolic disorders, namely, obesity, type 2 diabetes, dyslipidemia and cardiovascular disease can act as modifiable risk factors for cognitive impairment.^9^ Interplay among ApoE, lipid homeostasis, brain glucose, and Aβ trafficking in animal models of Alzheimer’s disease has been reported.^10^

In this study, we decipher the genetic underpinnings of cognitive functioning while considering the effects of putative interrelations with metabolic risk factors in the UK Biobank. We also identify variants in crucial genes that work in conjunction and interact with *APOE* in influencing cognitive functioning at a granularity of specific cognitive domains in the presence of lipid and glycemic metabolic risk factors.

## Materials and Methods

### Samples and participants

We present our analysis based on whole exomes of 200,643 individuals enrolled in UK Biobank (approved project-ID 55652).^11^

### Phenotypes

We consider six cognitive domains of simple processing speed, episodic memory, fluid intelligence, working memory, visual attention, and complex processing speed corresponding to which ‘Reaction time’, ‘Pairs’, ‘Reasoning’, ‘Digit recall’, ‘Trail making’, ‘Digit-symbol substitution’ cognitive tests were administered on the UK Biobank participants (https://biobank.ctsu.ox.ac.uk/crystal/refer.cgi?id=8481) (further details in Supplementary information).

### Genetic data and quality control

We download the UK Biobank population-level exome OQFE files for ~200k exomes in pVCF format (Field id: 23156) using the ‘gfetch’ utility. After extensive quality checks (details in Supplementary information), we retain 157,160 individuals with 211,012 variants (Supplementary Table 1).

### Heritability

Before proceeding to association analyses, we assess the heritability of the six cognition phenotypes based on unrelated individuals using LDAK^12^ model (Supplementary information). Our heritability estimates (Supplementary Table 2) show good concordance with evidence from previous family-based studies and GWAS ATLAS resource.^13^

### *APOE-carrier* status determination

Out of 157,160 samples, 93 have missing genotype information for *APOE* at either rs7412 or rs429358 or both. We determine *APOE*-carrier status, by flagging samples with at least one copy of *ε*4 allele as risk, with at least one copy of *ε*2 as protective/beneficial, *∈*1/*∈*3 and *∈*3/*∈*3 carriers as neutral, to include as a covariate in association models (Supplementary Table 3).

### Statistical Analyses

#### Single variant association

With genetic data on the resultant 157,067 samples and 211,012 variants, we perform single-variant Wald test using rvtests.^14^ Our baseline model control for age, gender, educational qualification, top 10 principal components and APOE-carrier status (Supplementary information). Further, to control for age-related metabolic conditions that can adversely affect cognition, we add lipid levels (serum total cholesterol, HDL and LDL direct cholesterol, triglycerides), glucose and HbA1c levels separately as covariates to the baseline model in models 2 and 3 (Supplementary information). We obtain the residuals and test the inverse-normalized residuals against genotype of each variant (Supplementary information). We obtain Manhattan plots (Supplementary Fig. 1-4) and QQplots (Supplementary Fig. 5-6) to visualize our results, and significant hits. To ascertain novelty, we use the LDtrait module of LDlink^15^ to check if variants which are in high LD (r^2^ > 0.8) and fall within ± 500kbp with our significant variant were previously associated with any trait listed in the EBI-GWAS catalogue.^16^

#### Gene-based association

We perform gene-based association tests in 157,067 individuals with kernel based (SKAT ^17^ and unified kernel and burden based methods (SKAT-O^18^) to detect cumulative burden in genes that work concomitantly with APOE. Figure 1 represents the possible pathways in which APOE affects neuronal dysfunction and hence cognition. Consideration of genes along these pathways ensures capturing the genetic basis of cognition in association with lipids homeostasis, glucose metabolism and amyloid-beta pathogenesis that plausibly play vital roles in modulating cognition with age.

**Figure 1.**
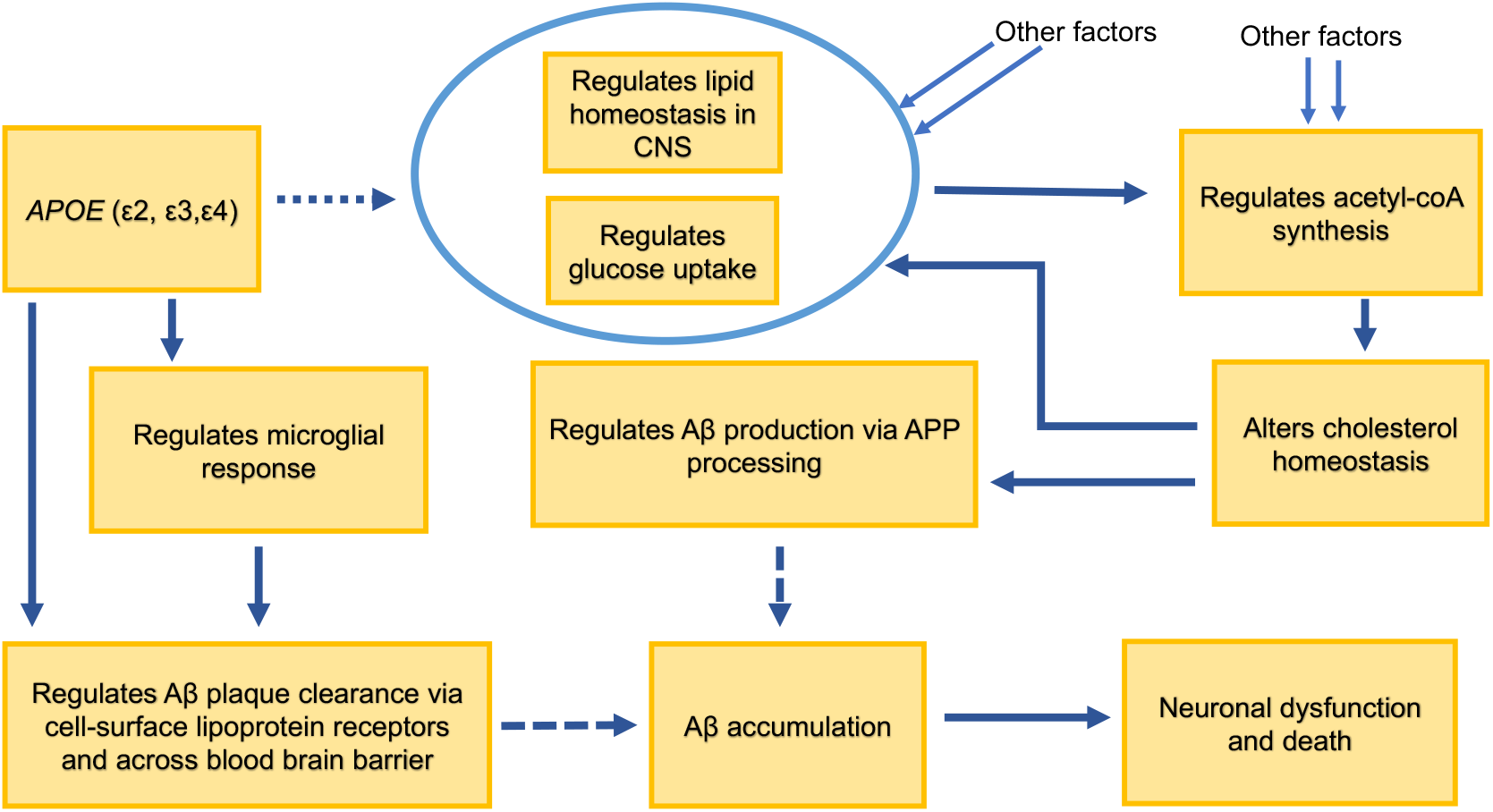
Putative pathways in which APOE isoforms regulate neuronal dysfunction.

#### Pairwise epistasis

We further uncover the interactions among the significant loci discovered by our single-variant association tests, through pairwise epistasis analysis (using plink-1.9.0 software)^19^ with each of the two *APOE* isoform-defining SNPs (rs429358 and rs7412). We control for all covariates used in models 2 and 3 except *APOE*-status and obtain the inverse-normalized residuals before testing for interaction effects.

Next, we conduct pairwise epistasis test for the variants in significant gene hits from either the SKAT or SKAT-O model (p-value < 0.0025), with *APOE* isoform-defining rs429358 and rs7412, as well as all *APOE* SNPs in two different models (Supplementary information).

#### Bivariate association tests

For each of the significant variants from our single-variant association tests, we conduct bivariate association tests for cognitive measures and lipid levels/ glycemic traits respectively, with the summary statistics obtained from the single variant Wald test results using metaMANOVA and metaUSAT^20^ (Supplementary information).

### Annotation and tissue-expression analysis

We annotate our exome-wide significant hits by mapping them to nearest genes (web resources) and calculating their deleteriousness using CADD^21^ scores, where higher scores indicate more deleteriousness. We perform functional annotation of the variants uncovered and calculate LofTool scores (web resources). Lower the LoFtool score, more intolerant is the said variant to functional changes. Also, we investigate, using GTEx,^22^ if our variants are eQTL loci or lie near eQTL loci significantly regulating expression in brain regions for the respectively annotated genes.

### Data availability

All phenotype and genotype data used in this study for analysis are available at UK BIOBANK (https://www.ukbiobank.ac.uk). We shall share the in-house scripts as required by other researchers.

## Results

### Identification of exome-wide significant variants for cognitive domains

From our exome-wide analysis on 211,012 variants in 157,160 individuals, we identify 20 independent loci associated to five different domains of cognition (summarized in Table 1, detailed results in Supplementary Table 4) with and without controlling for metabolic risks.

**Table 1:**
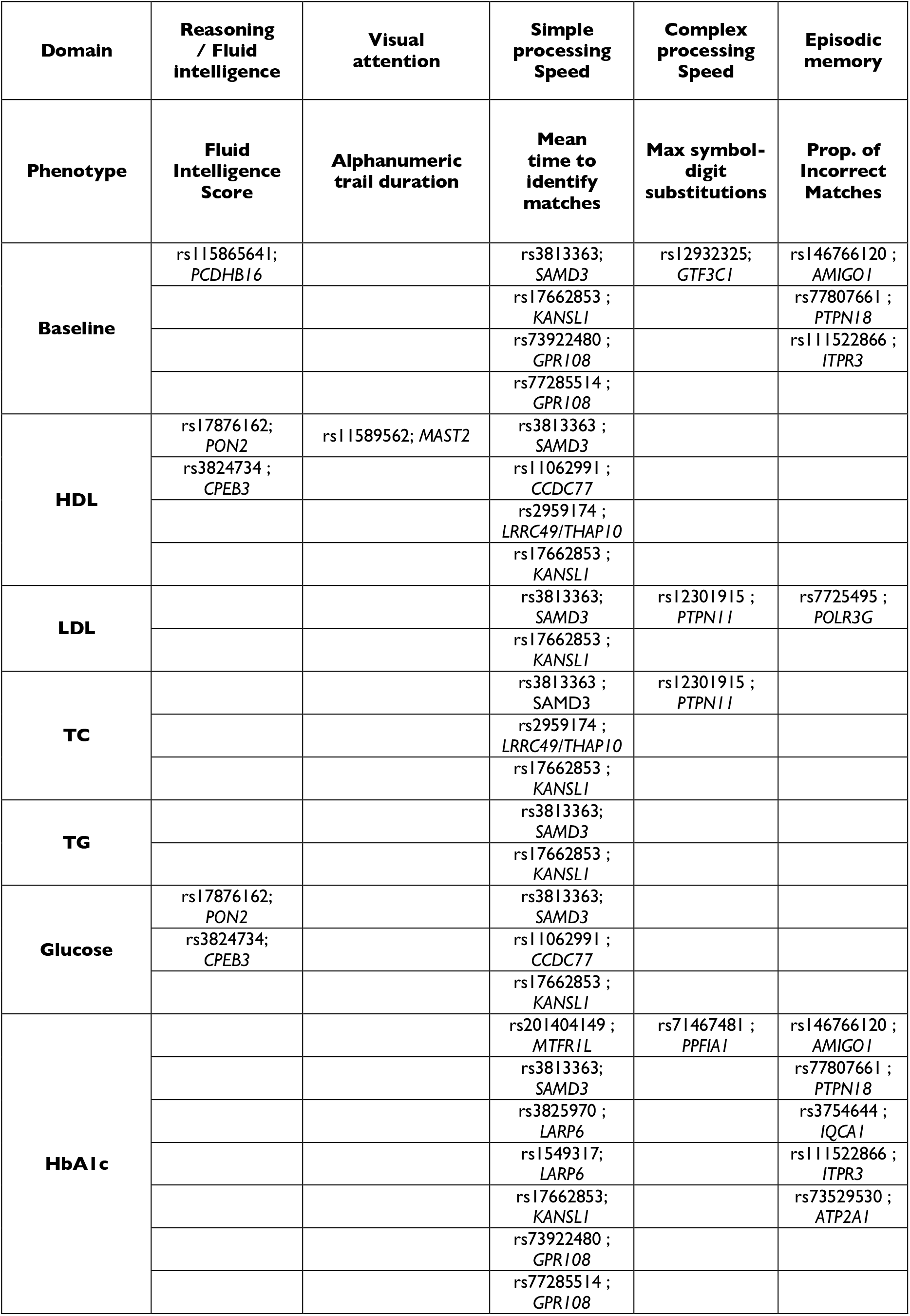
Single variant association analysis (summarized)

#### Fluid intelligence

We identify a novel rare variant rs115865641 in *PCDHB16* (3’UTR) associated with fluid intelligence (Supplementary Table 4). *PCDHB16* localizes mainly in the post-synaptic compartment and serve as a candidate gene for specification of synaptic connectivity and neuronal networks,^23^ a key element for cognition. Controlling for HDL and glucose separately, we obtain two independent significant novel hits-rs17876162 in *PON2* (intronic) and rs3824734 (synonymous) in *CPEB3* (Supplementary Table 4). *PON2* (Paraoxonase-2), a mitochondrial enzyme, has higher expression in dopaminergic regions such as striatum, striatal astrocytes, and cortical microglia,^24^ which suggest its role in protecting cells from oxidative damage and neuroinflammation.^24^ *CPEB3* is involved in synaptic protein regulation, acting as a negative regulator of AMPA receptor subunits GluA1,GluA2 to maintain long-term synaptic plasticity.^25^ Our novel synonymous rs3824734 (*CPEB3*) with a CADD score of 11.74 (Supplementary Table 4), implying that it is predicted to be among 7% of the most deleterious substitutions to the genome, could be crucial for pinpointing the role of this gene in cognition.

#### Simple processing speed

We detect rs3813363 (5’ UTR of *SAMD3*), and rs17662853, (missense variant in *KANSL1*) (CADD score 23.9) (Supplementary Table 4) to be associated with simple processing speed in the baseline as well as all models controlling for lipid and glycemic traits. rs3813363 is within 500kbp and in high LD (r^2^ > 0.8) of both rs11154580 and rs6937866, known to be associated with reaction time^5^ (Fig. 2). Highly deleterious rs17662853 is in high LD (r^2^=0.856) with intronic rs10775404 (CADD score: 1.782) previously associated with reaction time (Fig. 2)^5^, highlighting that our variant could be more impactful, and more likely to be causal. Koolen-de Vries syndrome/17q21.31 microdeletion syndrome characterized by intellectual disability has been attributed to mutations in *KANSL1*.^26^ One study showed that autophagosome accumulation at excitatory synapses in *KANSL1*-deficit neurons lead to reduced synaptic density, reduced transmission via GRIA/AMPA receptors along with impairment of neuronal network activity.^27^ We also identify two novel variants-rs73922480 and rs77285514 (synonymous and intronic *GPR108* respectively, 80 bp apart) to be associated with mean reaction time in the baseline model as well as controlling for HbA1c (Supplementary Table 4). Controlling for HbA1c, we additionally identify novel rs201404149 (synonymous *MTFR1L*), rs3825970 and rs1549317 (synonymous *LARP6*) associated with mean reaction time (Supplementary Table 4). A recent study showed that *MTFR1L* expression changed in the hippocampus and cerebral cortex in memantine-treated transgenic Alzheimer’s diseased mice.^28^ Memantine is an FDA approved prescription drug administered to improve learning and memory for moderate-to-severe Alzheimer’s cases; suggesting the importance of our identified loci in *MTFR1L* for understanding cognition in the pathophysiology of Alzheimer’s disease. We identify novel hits rs11062991 (intronic *CCDC77*) and rs2959174 (synonymous *THAP10*; intronic *LRRC49*) while controlling for serum HDL (Supplementary Table 4). rs11062991and rs2959174 are also associated with mean reaction time when we control for glucose and total cholesterol respectively. Although, immediate relevance to cognition phenotype for *LRRC49/THAP10* is unapparent, our variant in *LRRC49/THAP10* is an eQTL for *LARP6* in brain regions contextual to cognitive abilities.

**Figure 2.**
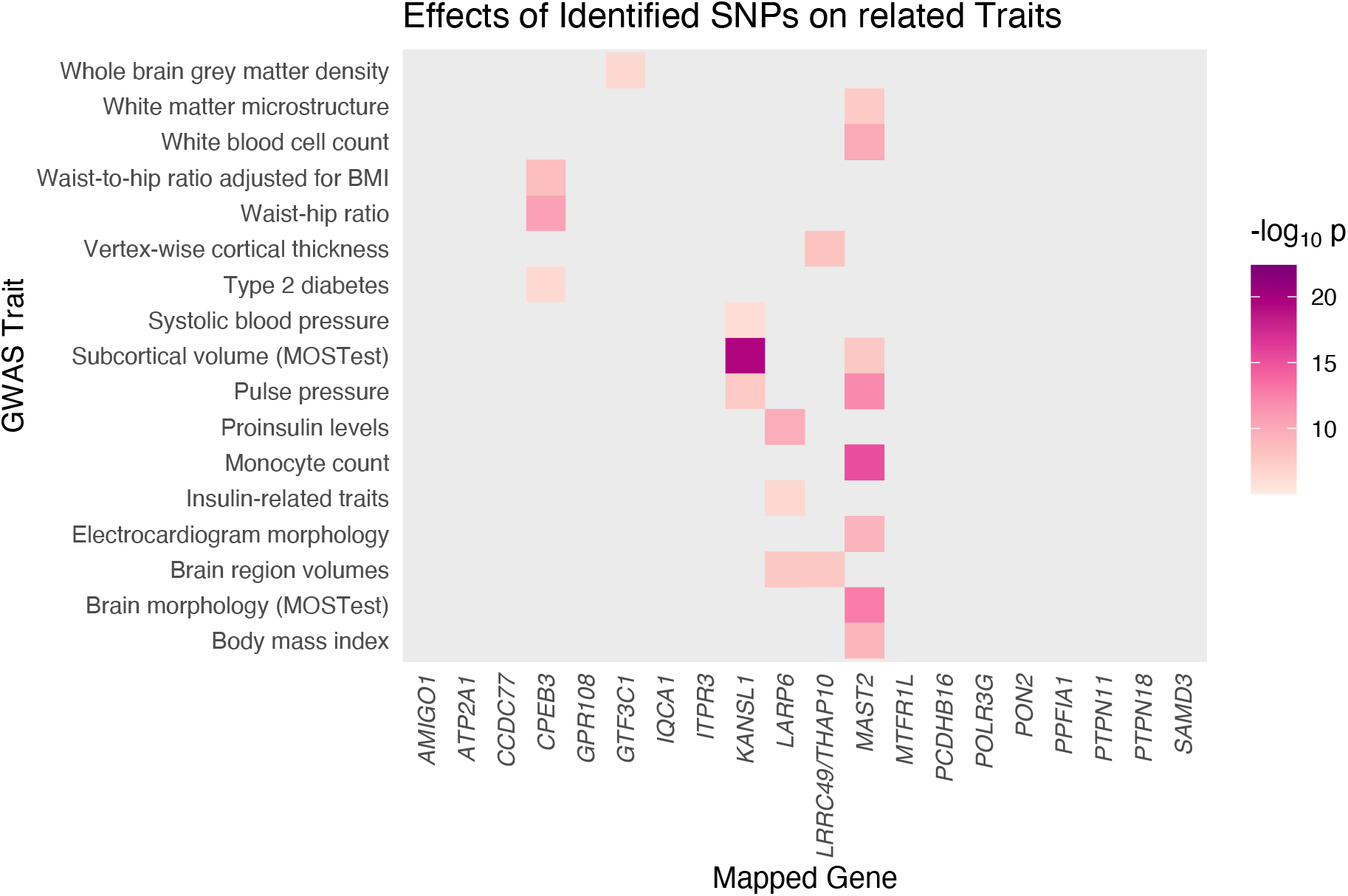
Effects of cognition-associated variants with related metabolic and brain structure traits. Previously known statistically significant effects of our exome-wide significant cognition-associated loci (mapped to nearest genes) on related metabolic and brain structure (obtained from EBI-GWAS catalogue) is highlighted in pink-purple gradient. Darker color signifies more significant association. Grey signifies no significant association.

#### Complex processing speed

We find novel rs12932325 (intronic *GTF3C1*) associated with complex processing speed from the baseline model. *GTF3C1* loci have been found to be significantly associated with entorhinal cortical thickness, an Alzheimer’s disease-related neuroimaging biomarker (Fig. 2).^29,30^ Our variant could potentially influence such related traits, and thus allude to the shared genetic mechanisms of cognition, Alzheimer’s disease and grey matter density. Controlling for LDL and total cholesterol levels independently, we detect rs12301915 (intronic *PTPN11*) as a novel hit (Supplementary Table 4). *PTPN11* is a tyrosine phosphatase that activates MAPK pathway, plays a critical role in synaptic plasticity and memory formation,^31^ and interacts with tau in Alzheimer’s patients.^32^ Mutations in *PTPN11* have been associated with numerous syndromes among which the human cognition affecting Noonan syndrome is the most common,^33^ along with cardiovascular abnormalities and congenital heart defects.^34^ Controlling for HbA1c, we identify rs71467481 (intronic *PPFIA1*) as another novel significant hit (Supplementary Table 4). *PPFIA1* encodes the neuronal scaffold protein liprin-α1 functioning in active synaptic zones and post-synaptic sites,^35^ and has been proposed as a candidate gene involved in late-onset Alzheimer’s disease etiology.^36^

#### Episodic memory

We identify three novel variants rs146766120 (missense *AMIGO1*), rs7780766 (synonymous *PTPN18*), and rs111522866 (intronic *ITPR3*) to be associated with episodic memory in the baseline as well as controlling for serum HbA1c levels (Supplementary Table 4). From the HbA1c controlled models, we additionally identify rs3754644 (missense *IQCA1*) and rs73529530 (intronic *ATP2A1*) to be associated with episodic memory (Supplementary Table 4). Upon controlling for LDL, we find another novel variant rs7725495 (intronic *POLR3G*) to be associated with episodic memory (Supplementary Table 4). rs146766120 (CADD score: 15.97) is among the top ~3% deleterious variants, and *AMIGO1* is a commonly altered marker gene in Alzheimer’s patients.^37^ *PTPN18* is a non-receptor tyrosine phosphatase expressed in neural tissues, likely influencing Alzheimer’s disease progression.^38^ *ITPR3* encodes inositol 1,4,5-trisphosphate receptor, type 3, which mediates release of intracellular calcium and facilitates crucial intra-organellar Ca^2+^ signal transmission from the endoplasmic reticulum (ER) to the mitochondria^39^ for maintaining proper cognition. *IQCA1*, found to be upregulated in the hippocampus of Alzheimer’s-like monkeys as compared to normal aged monkeys, is postulated to be associated with brain AMPKα2 activity playing a pivotal role in de-novo protein synthesis, an indispensable phenomenon for long-term synaptic plasticity and memory formation.^40^ *ATP2A1* encodes proteins associated with mitochondria-associated-ER membrane (MAM)^41^ and disruption at this locus could perturb MAM functioning which is posited to play a role in Alzheimer’s disease pathogenesis. Our results thus provide genetic insights into cognition in Alzheimer’s disease.

#### Visual attention

We detect novel rs11589562 (intronic *MAST2*) as associated with visual attention, measured by alphanumeric trail duration, when controlling for HDL levels. rs11589562 can significantly control expression of several nearby genes, including *MAST2* in different brain regions.

### Genes implicated in cognition from kernel and burden tests

We identify *APOC1* to be significantly associated with complex processing speed and visual attention in baseline model and in models controlling for LDL, total cholesterol, triglycerides and HbA1c (Supplementary Table 5). *APOC1* (~5kb downstream of *APOE*) encodes the smallest of all lipoproteins participating in lipid transport and metabolism and is known to be pleiotropically associated with serum HDL, LDL, triglyceride and cholesterol and HbA1c levels.^42^ Animal model studies have indicated the role of *APOC1*, expressed in astrocytes and endothelial cells of hippocampus, in cognitive processes in both *APOE* dependent and independent manner. ^43^ rs4420638 (*APOC1*), has been implicated in general intelligence^44^ and CSF biomarker levels.^45^ However, we report the first evidence of *APOC1* influencing two specific cognitive domains through collective burden of all variants in the gene in a human population. We could uncover this effect of *APOC1*, after filtering out the more plausible effects of *APOC1* in lipid and glycemic pathways, thereby highlighting the independent role of *APOC1* in cognition and the importance of considering appropriate co-occurring metabolic risks in genetic epidemiological studies.

Controlling for HDL and the baseline covariates, we also identify *LRP1* as a significant gene influencing visual attention (Supplementary Table 5). A few targeted studies indicate that *LRP1* SNPs and haplotypes influence cognitive performance in Chinese patients with risk of Alzheimer’s disease.^46,47^ This gene encodes the low-density lipoprotein receptor-related protein1, an endocytotic receptor with over 40 ligands including ApoE and Aβ, regulating Aβ uptake and clearance across the blood-brain barrier along with its signaling role in Alzheimerls disease pathology.^48^ Our results provide first ever evidence from large scale human whole-exome based analysis on the role of the elusive *LRP1*, in visual attention.

### Pleiotropy and mediation

Out of the 20 independent loci (Fig. 3A), 15 independent loci (*PCDHB16, PON2, MTFR1L, SAMD3, LARP6, KANSL1, GPR108, PPFIA1, PTPN11, AMIGO1, PTPN18, IQCA1, POLR3G, ITPR3, ATP2A1*) exhibit pleiotropic effects on lipid and/or glycemic phenotypes (Supplementary Tables 6-10). Interestingly, we identify suggestive mediating effects of four of these 20 loci on their respective cognitive domains. rs115865641 (*PCDHB16*), associated to fluid intelligence in our baseline model, is also found to be associated with HDL and glucose, but shows effect sizes reduced in magnitude when we control for HDL and glucose levels separately, and is also pleiotropically associated with lipid and glycemic traits (Supplementary Table 6). This suggests that serum HDL and glucose levels could partially mediate the effect of rs115865641on fluid intelligence along with its pleiotropic effect. Similar effects were observed for rs201404149 (*MTFR1L*) associated to simple processing speed controlling for HbA1c. rs201404149 is significant from the baseline model, pleiotropically associated with serum glucose levels and this variant shows reduced effect size on mean reaction time when controlling for serum glucose levels (Supplementary Table 7) indicating that the effect of this variant on reaction time could be partially mediated through its effect on serum glucose levels, providing further evidence of metabolic risk affecting cognition. Similarly, the *PPFIA1* variant rs71467481 is significant in the baseline model, and is associated with serum glucose levels but shows reduced effect size than baseline when controlling for glucose implying that the effect of this variant may be mediated through glucose homeostasis in influencing complex processing speed. This variant also shows pleiotropic association with HDL, LDL and glucose levels (Supplementary Table 9). rs73529530 in *ATP2A1* shows association with HDL and glucose levels in addition to pleiotropic association with cognition phenotype and all lipid levels and serum glucose levels. rs73529530 may also affect episodic memory by mediation through serum HDL and glucose levels as reflected by the reduction in magnitude of effect size compared to baseline when the phenotype is controlled for HDL and glucose levels (Supplementary Table 10).

**Figure 3.**
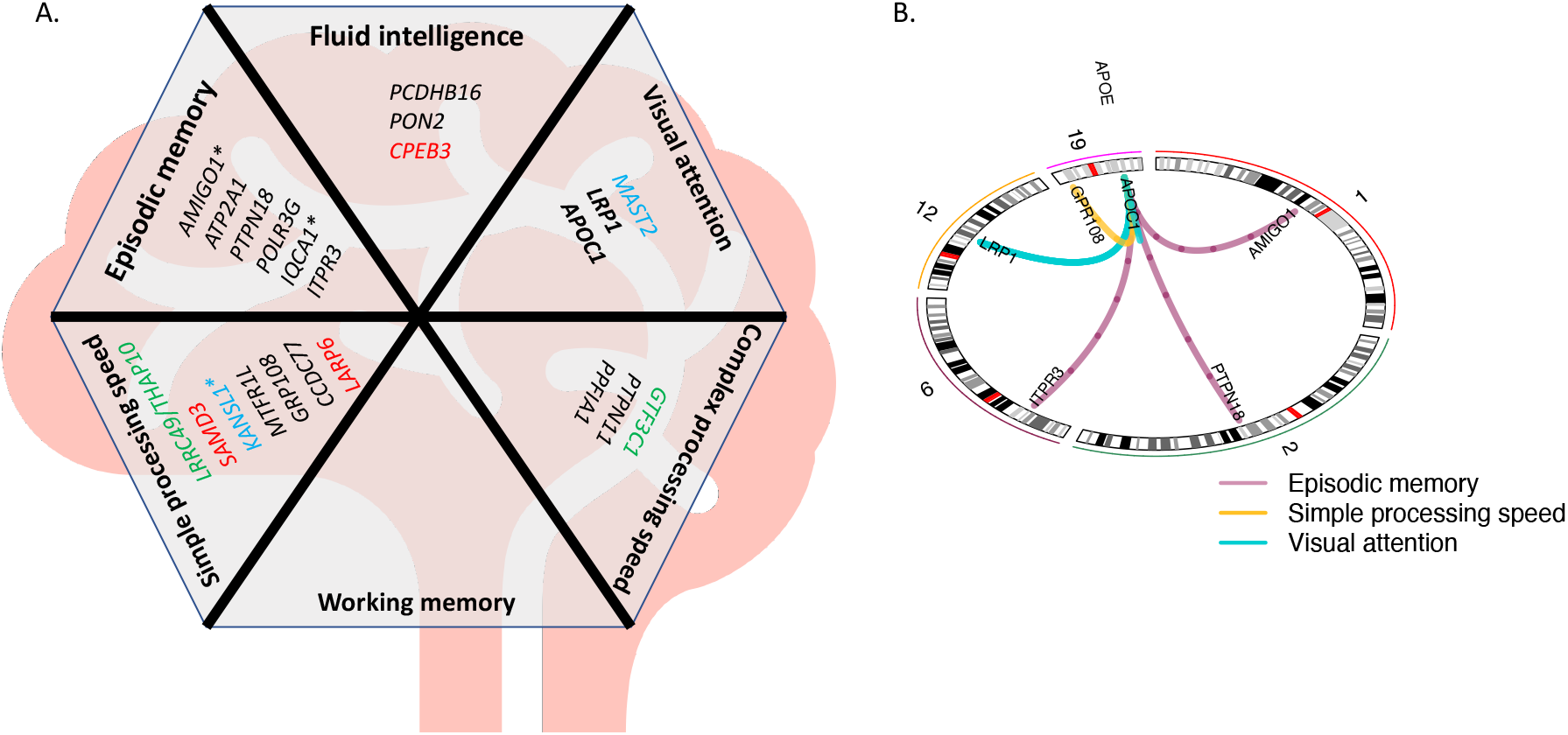
Summary figure showing association hits mapped to nearby genes corresponding to diverse cognition domains and their interactions with *APOE*. **(A)** Variants and genes we have uncovered associated to diverse cognition domains. The genes to which our single variant hits have been annotated and the genes identified from gene-based tests (given in bold) have been represented here. The variants which are eQTLs for the genes they have been mapped to have been represented in red; which are eQTLs for nearby genes have been given in green; and the ones which are eQTLs for both their annotated and nearby genes have been represented in light blue. The genes corresponding to variants which are suggestive eQTLs (because of their proximity to eQTL variants) have been shown in black. The missense variants have been represented with asterisk sign (*) beside their corresponding genes. (**B**) Circos plot showing pairwise interactions of loci with APOE influencing diverse domains of cognition. The numbers on the periphery of the circle represent the chromosome. The purple lines represent interactions influencing episodic memory, the yellow lines represent interactions influencing simple processing speed and the turquoise lines represent interactions affecting visual attention. Tables 2 and 3 contain related details.

### Expression profile analysis

#### eQTL analysis of significant loci associated with fluid intelligence

We identify novel rs115865641(3’ UTR of *PCDHB16*) associated with fluid intelligence scores from the baseline model. Controlling for HDL and glucose levels separately, we obtain two independent significant novel hits-rs17876162 in *PON2* and rs3824734 in *CPEB3*. rs3824734 is an eQTL controlling significant expression of *CPEB3* in cerebellar hemispheres (NES=0.22, p-value=3.8 x 10^-5^) (Supplementary Table 11) which is known for its role in influencing intelligence.^49^ Even though rs115865641 (rare variant) is not a significant eQTL controlling expression of *PCDHB16* as per GTEx data, we find that all eQTL variants lying within ± 500kb of our novel variant significantly control expression of *PCDHB16* in cerebrum, which contains the prefrontal cerebral cortex – the postulated seat of fluid intelligence,^49,50^ and also in cerebellar hemispheres, hippocampus and basal ganglia (Supplementary Table 12). Tissue specific expression data reveals that *PON2* is highly expressed in frontal cortex, anterior cingulate cortex, and basal ganglia (Supplementary Fig. 7) which are areas in the brain correlated with fluid intelligence.^50–52^

#### eQTL analysis of significant loci associated with simple processing speed

We find that rs3813363 in SAMD3, the association hit for mean reaction time from all models, as a significant eQTL controlling expression of *SAMD3* in the cortical regions of the brain (NES=-0.4, p-value=1.1 x 10^-5^) (Supplementary Table 11). Several studies have established that cortical regions of the brain are well correlated with reaction time phenotypes assessing the domain of simple processing speed.^53^ Similarly, rs17662853-the missense hit in *KANSL1* for reaction time, is an eQTL with significant expression for *KANSL1* in the cerebellum (NES = −0.4; p-value=2.3 x 10^-5^) and anterior cingulate cortex (NES = −0.49; p-value=2.5 x 10^-5^) (Supplementary Table 11), regions responsible for perception and motor response whose coordination is necessary for completion of a reaction time task.^53,54^ rs17662853 is also an eQTL controlling expression of *NSFP1, LRRC37A, ARL17A, ARL17B*, RP11-798G7.8, *NSF, NSFP1, FAM215B* in several brain regions including cortex and cerebellum (Supplementary Table 11), highlighting the importance of significantly associated variants obtained from exome-wide analysis, that could regulate expression of nearby genes relevant to the biology of the trait. eQTL variants in *GPR108* significantly control expression of *GPR108* in various brain tissues with the lead eQTLs within 500kbp of our lead SNP controlling *GPR108* expression significantly in the cortex (Supplementary Table 12). Our eQTL analysis reveals that loci around 500kbp of, rs11062991 (intronic *CCDC77*) most significantly regulates expression of *CCDC77* in hypothalamus and cerebellar hemispheres (Supplementary Table 12). rs2959174 (synonymous *THAP10/* intronic *LRRC49*) is a significant eQTL regulating the high expression of *LARP6* in cerebellum, cerebellar hemispheres and putamen of basal ganglia, hippocampus and cortex (Supplementary Table 11).Thus, our eQTL analysis reveals another relevant gene *LARP6* for understanding the biology of cognition, even though the identified variant itself annotates to *LRRC49* and *THAP10*, with less relevance to cognition (https://maayanlab.cloud/Harmonizome/gene_set/Cognition+Disorders/CTD+Gene-Disease+Associations).^55^ The *LARP6* loci identified from our analysis (rs3825970 and rs1549317) is also a significant eQTL controlling *LARP6* expression in cerebellum, cerebellar hemispheres and putamen of basal ganglia (Supplementary Table 11). eQTLs within 500kbp of rs201404149 (synonymous *MTFR1L*) are significant for expression of *MTFR1L* in cerebellum, cortex, frontal cortex, cerebellar hemispheres, and caudate nucleus of basal ganglia (Supplementary Table 12).

#### eQTL analysis of significant loci associated with complex processing speed

The baseline model for this domain yields one significant hit-rs12932325 in intronic region of *GTF3C1*. rs12932325 is an eQTL for *IL21R* (Supplementary Table 11) which impacts Alzheimer’s disease pathology by enhancing brain and peripheral immune and inflammatory responses and leads to increased deposition of Aβ plaques.^56^ Both the models controlling for LDL and total cholesterol levels independently yield a novel intronic variant in *PTPN11* as a novel significant hit for complex processing speed. The lead eQTL variant near ±500 kb of this variant significantly controls expression of *PTPN11* in the substantia niagra of the brain (Supplementary Table 12). Research has shown that Parkinson’s disease causes loss of dopamine producing neurons in the substantia nigra and dopaminergic processes have been shown to be involved in cognitive functions like processing speed.^57^ Controlling for HbA1c, we identify rs71467481 in intronic region of *PPFIA1* as another novel significant hit for complex processing speed. Our eQTL analysis shows that variants around 500 kbp of this SNP can significantly regulate expression of *PPFIA1* in many brain regions (Supplementary Fig. 8, Supplementary Table 12).

#### eQTL analysis of significant loci associated with episodic memory

We identify a significant novel missense variant rs146766120 in *AMIGO1* to be associated with episodic memory with and without controlling for serum HbA1c levels. *AMIGO1* is expressed in the astrocytes, hippocampus and cortical neurons and it is postulated to influence neuron survival.^58^ In our eQTL analysis too, we find that variants within 500kb of rs146766120 significantly influences expression of *AMIGO1* in the brain, especially in cortex (NES = 0.2, p-value = 8.2 x 10^-12^) and hippocampus (NES = 0.17, p-value = 1.4 x 10^-11^) (Supplementary Table 12, Supplementary Fig. 9), areas in the brain which interact among each other to encode and retrieve episodic memory,^59,60^ thus highlighting the importance of our identified hit in influencing episodic memory. Similarly we identify another novel synonymous variant rs7780766 in *PTPN18* both with and without controlling for serum HbA1c levels. Significant eQTL variants around 500 kbp of rs7780766 can regulate expression of *PTPN18* in cortex, prefrontal cortex, cerebellum, cerebellar hemispheres, caudate basal ganglia and nucleus accumbens (Supplementary Table 12 and Supplementary Fig. 10), thus pinpointing to the established crucial role of cerebellum in episodic memory via cortical-cerebellar brain networks.^61^ Studies also suggest that memory formation in hippocampus is guided by motivational significance of events whose effect on memory is thought to depend on interactions between hippocampus, ventral tegmental area and nucleus accumbens.^62^ The baseline model as well as the model controlling for HbA1c also yield rs111522866 in the intronic region of *ITPR3* as another novel significant variant for episodic memory. eQTL variants around 500kb of this variant are significantly influence expression of *ITPR3* in cerebellar hemispheres as well as in caudate basal ganglia (Supplementary Table 12). Upon controlling for LDL, we find another novel variant rs7725495 in intronic region of *POLR3G* to be associated with episodic memory. eQTL variants within 500kb of rs7725495 significantly influences expression of *POLR3G* in cerebellum, cortex, anterior cingulate cortex, hypothalamus, nucleus accumbens and putamen (Supplementary Table 12).

We identify two additional novel hits - missense rs3754644 (*IQCA1*) and rs73529530 (intronic *ATP2A1*) to be associated to episodic memory when controlled for HbA1c levels. eQTL variants around 500kb of rs3754644 also significantly control *IQCA1* expression in amygdala, cerebellar hemispheres, cerebellum, cortex, frontal cortex, anterior cingulate cortex and nucleus accumbens (Supplementary Table 12). eQTL variants around 500kb of rs73529530 significantly regulates expression of *ATP2A1* in hypothalamus (Supplementary Table 12).

#### eQTL analysis of significant loci associated with visual attention

We get an association signal of rs11589562 for visual attention when we adjust for HDL level. This variant is located in intronic region of *MAST2* gene. eQTL analyses shows that this variant controls expression for *MAST2* in cerebral cortex (NES = 0.21, p-value = 7.70E-06) and cerebellum (NES = 0.26, p-value = 4.6E-06) (Supplementary Table 12). It is known that the posterior parietal lobe of the cortex assesses the visual scene and it interacts with the frontal lobes in choosing object of interest to plan visually guided movement.^63^ This variant is also an eQTL significantly influencing expression of *CCDC163, TESK2*, and *PIK3R3* in several brain regions (Supplementary Table 11). *MAST2* is highly expressed in the hypothalamus and substantia niagra (Supplementary Fig. 11). Several studies have found oxytocin, synthesized in several nuclei of the hypothalamus, to regulate visual attention and eye movements to external sensory/social stimuli.^64^ Additionally, studies have shown dopamine producing neurons in the ventral tegmental area and substantia nigra to be related to multiple aspects of visual attention.^64^

### Interaction analyses

Our epistasis analysis conducted with significant variants from the single variant analysis reveals four pairs of significant epistatic interactions with the *APOE* isoform-defining variants (rs7412 and rs429358) for episodic memory and simple processing speed (Table 2; Fig. 3B). Each of the variants which interact with either of the two *APOE* variants exerts a significant effect on the phenotype in addition to its interaction effect. These variants are rare with large effect sizes conforming to the general consensus that rarer variants have larger effect sizes. Out of these interactions, we find the interaction between rs429358 (*APOE*) and rs14676612 (*AMIGO1*) and between rs429358 and rs77807661 (*PTPN18*) of particular interest. We see that both the variants in the *APOE-AMIGO1* and *APOE-PTPN18* interactions (baseline and HbA1c controlled) exert a significant main effect as well as an interaction effect on episodic memory even when we tease out the effect of serum HbA1c on episodic memory. *ITPR3* and *GPR108* variants also exhibit an interaction effect with *APOE* for episodic memory and simple processing speed respectively. The epistasis analysis conducted on the basis of gene-based tests reveals nine significant epistatic interactions between five *APOE* (rs440446, rs143063029, rs769449 rs429358 and rs7412) and eight *LRP1 SNPs* (Table 3) to be associated with visual attention, adjusting for HDL levels. It also reveals one significant interaction between rs7412 (*APOE*) and rs1064725 (3’UTR *APOC1*) to be associated with alphanumeric trail duration while controlling for baseline covariates and for HbA1c independently.

**Table 2:**
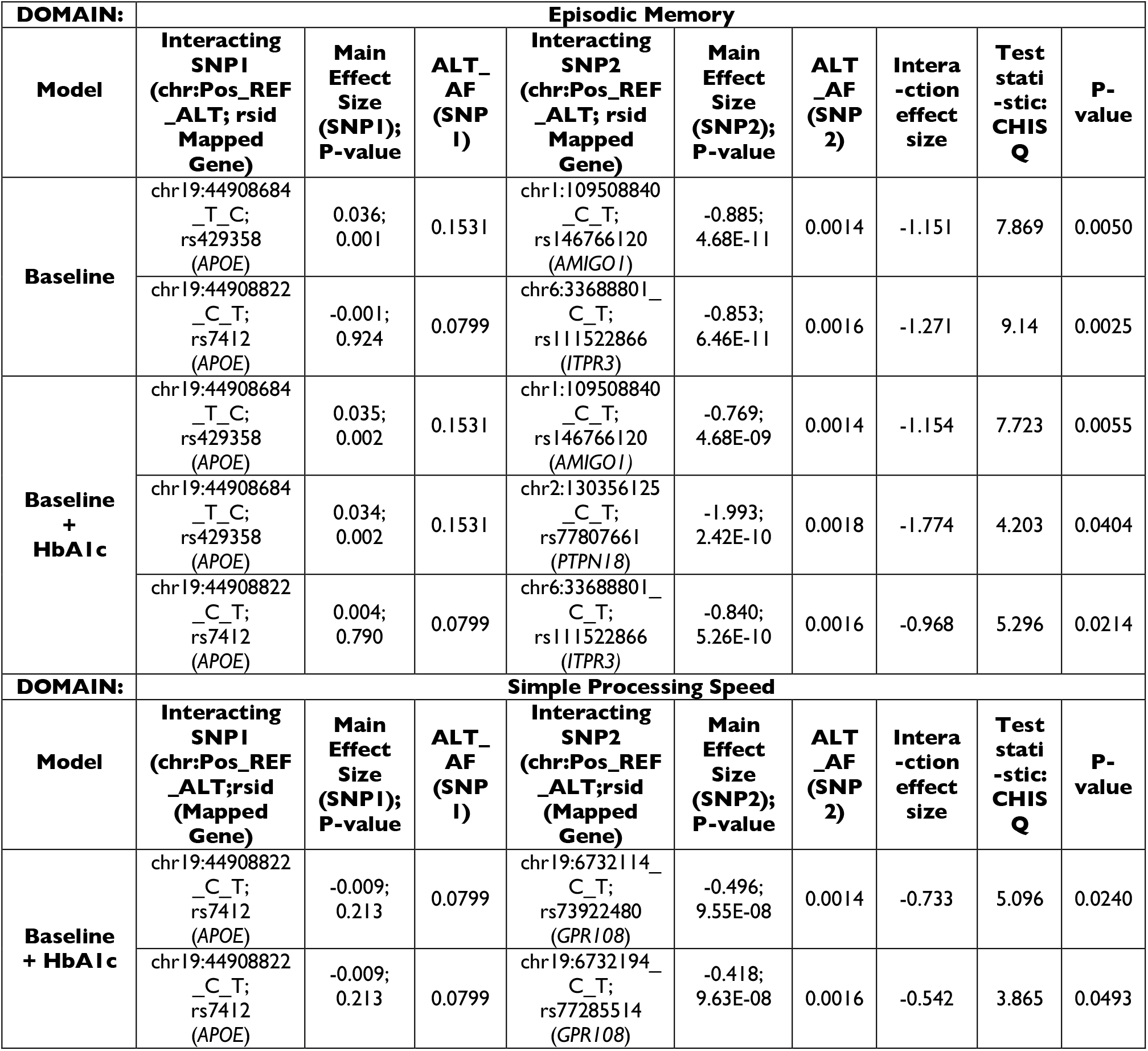
Interaction analysis with variants identified from single variant association tests.

**Table 3:**
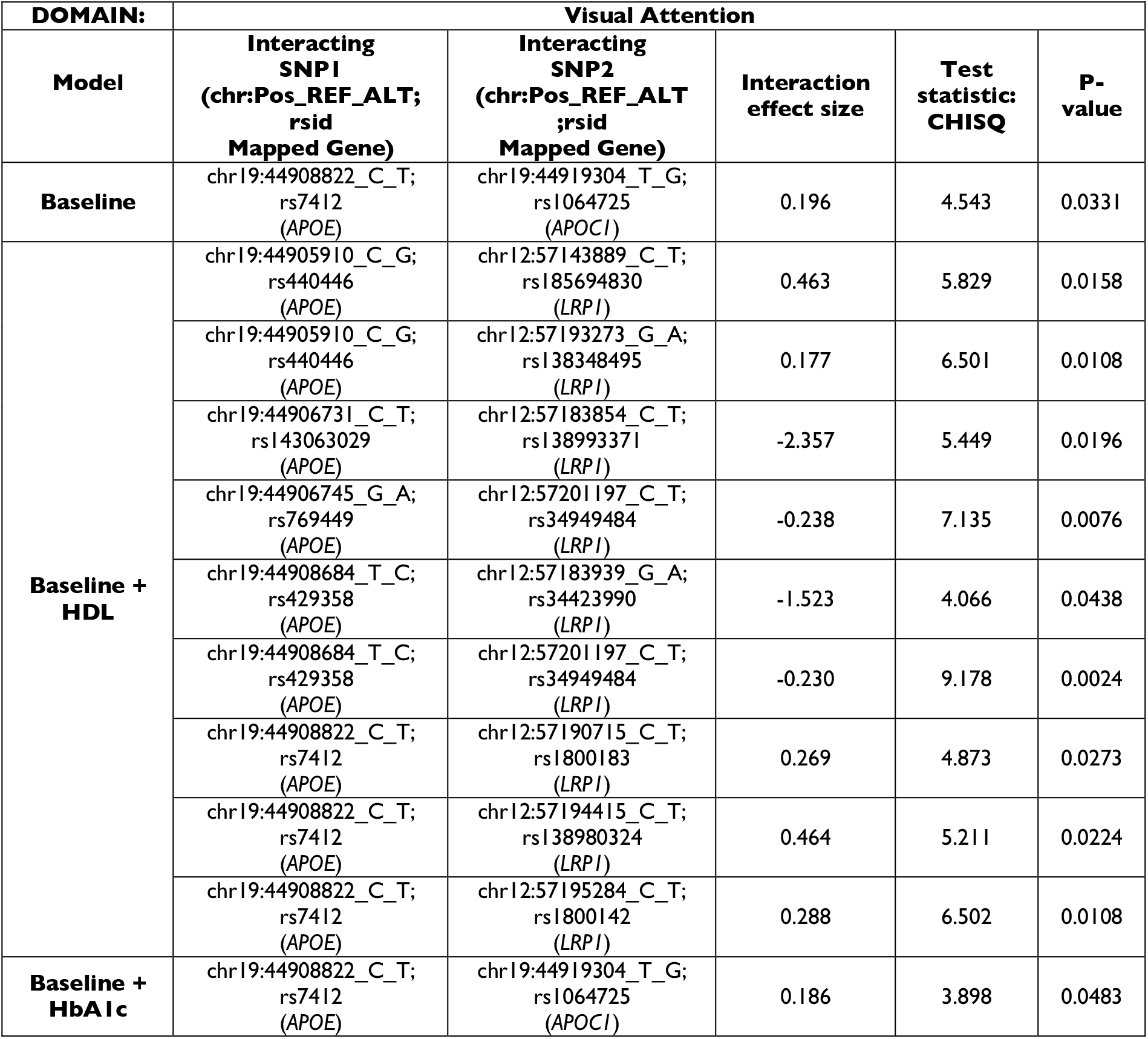
Interaction analysis of variants in genes identified from genebased association tests.

## Discussion

Our study is a comprehensive analysis to understand the genetic architecture of human cognition via single variant based, gene-based association, pairwise interaction, mediation and pleiotropy analyses (Fig. 4).

**Figure 4.**
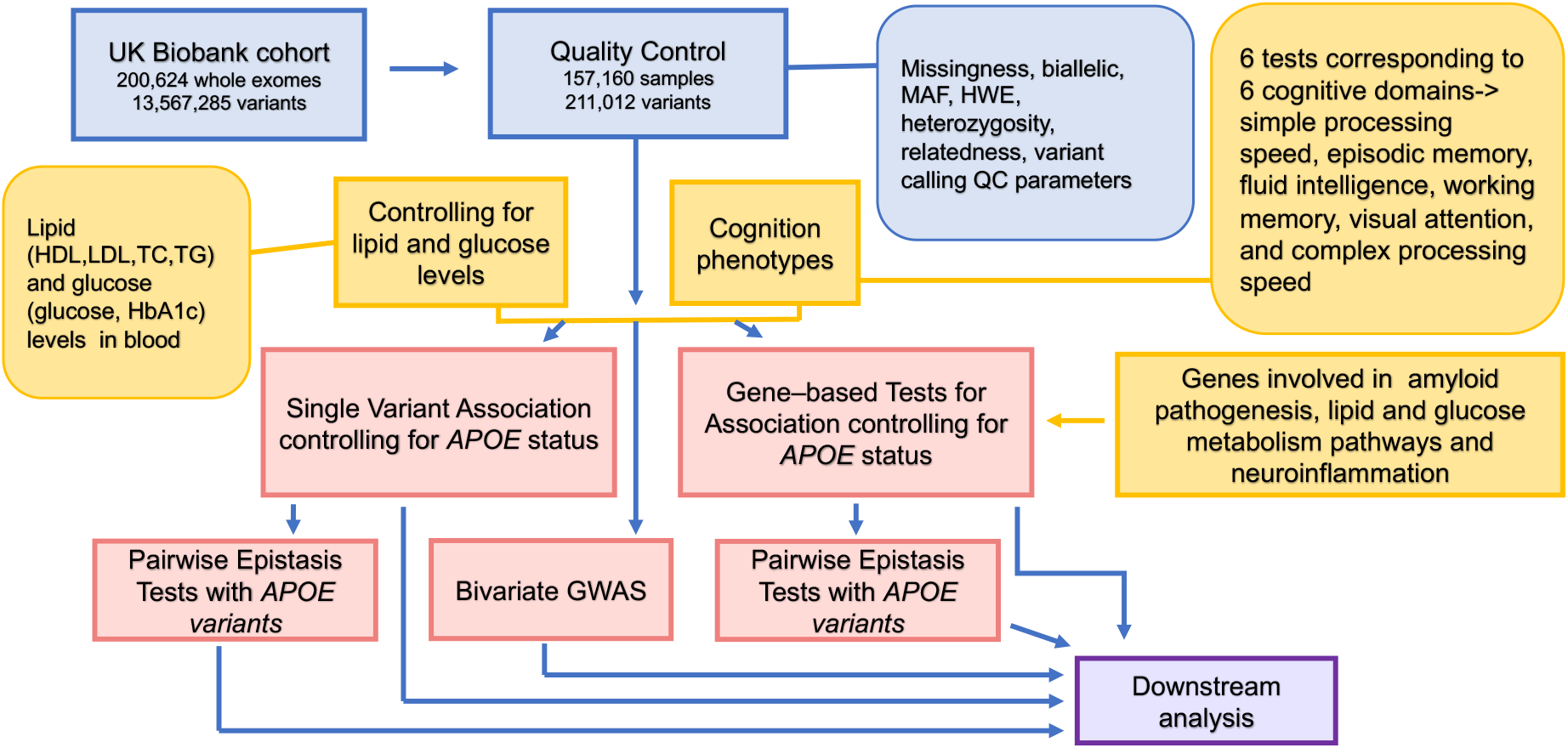
Workflow of this study. Blue boxes represent the information about the data and quality checks performed; the yellow boxes are indicative of the phenotypes and genes considered for gene-level analysis. Red boxes highlight the statistical tests performed; and the purple box indicates downstream analysis performed such as annotations, gene expression analysis and mediation analysis.

Our single-variant and gene-based association identifies novel independent loci in *PCDHB16*, *PON2*, *CPEB3*, *LRRC49/THAP10, CCDC77, LARP6, MTFR1L, GPR108, GTF3C1, PTPN11, PPFIA1, AMIGO1, ITPR3, PTPN18, IQCA1, ATP2A1, POLR3G, MAST2, APOC1, LRP1*, and previously known *KANSL1, SAMD3* as associated with diverse cognition domains (Fig. 3A) in baseline as well as while adjusting for serum lipids and glycemic levels which are postulated to be modifiable metabolic risk factors for cognition. We note that these implicated genes are known to impact Alzheimer’s disease and related dementias through their functioning in synaptic plasticity and connectivity, oxidative stress, neuroinflammation. Interestingly, all risk alleles of single variant hits affecting cognition are common in the population with allele frequency > 5%, thus highlighting the significance of our work for studying the genetic context of cognitive abilities of individuals in the general population in order to understand the risk factors for cognitive decline. We have also obtained significant hits harboured in the coding region which are in LD with genotyped variants identified by Davies et al.^5^ associated with reaction time, thus highlighting the importance of exome-based analysis in uncovering likely causal associations.

Functional annotation of the significant variants reveal that majority of them are rare and have possibly damaging effects on the gene function (LoFTool score < 0.25), explaining comparatively higher proportion of variance (Fig. 5). However, as exceptions, we note a few common and low frequency variants with possible deleterious effects, yet explaining comparatively moderate or low proportion of variation by virtue of lower effect size (Fig. 5).

**Figure 5.**
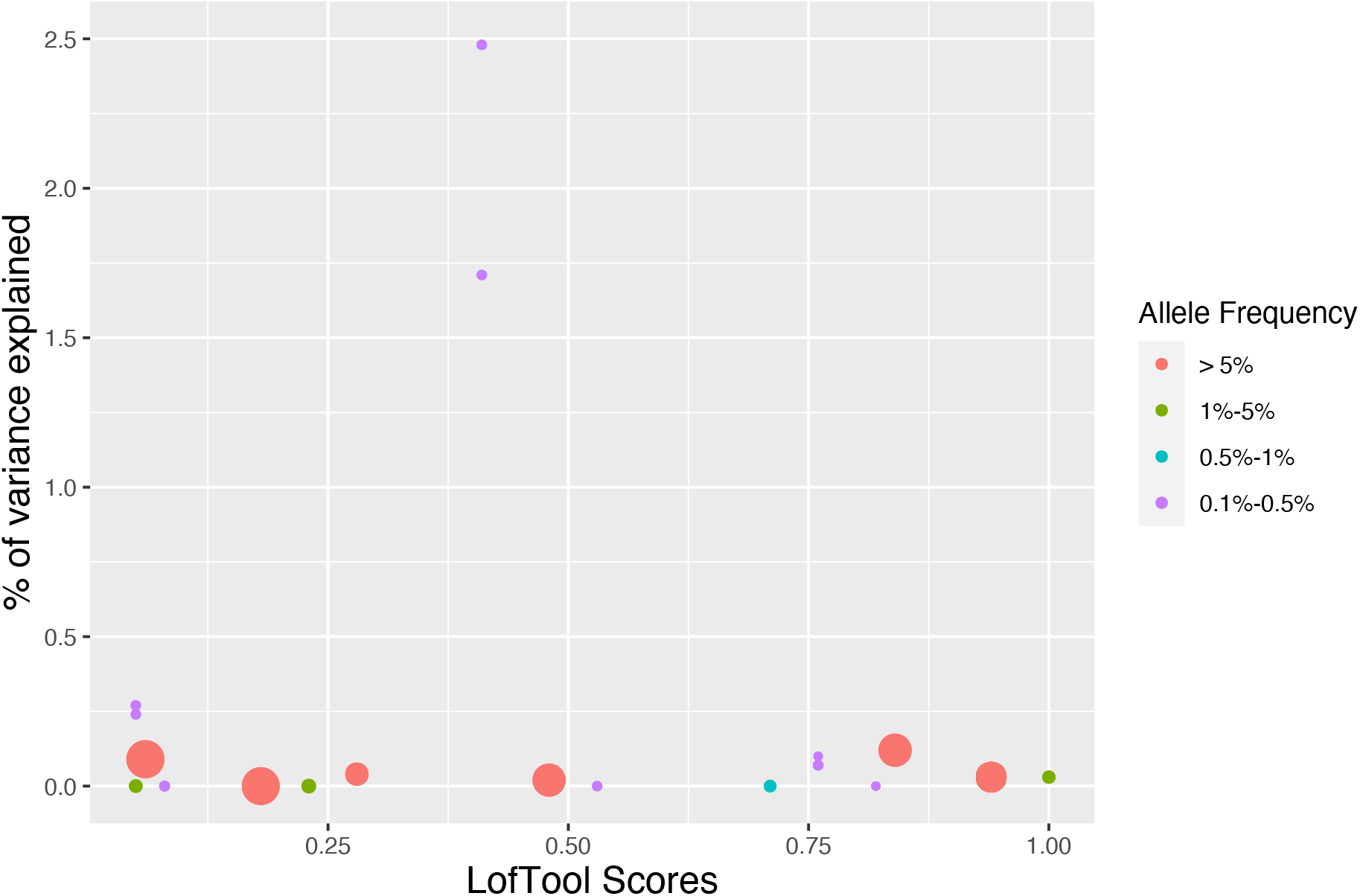
Phenotypic variance explained with respect to evolutionary constraint acting on the variants identified from single variant association tests. This figure represents the proportion of phenotypic variance explained by the exome-wide significantly associated variants for diverse cognitive domains vs their intolerance to genic functional changes. Coral and green, solid circles represent common, low variants respectively while turquoise and violet solid circles represent rare frequency variants. The sizes of the circles are proportional to the allele frequency of the corresponding variants represented by the circles

While the general consensus is that rare variants exhibit higher effects and are more likely to be deleterious, our results show that disease-associated common variants can also be intolerant to loss-of-function.

Out of the 20 independent loci (Fig. 3A), rs3824734 (*CPEB3*), rs3813363 (*SAMD3*) and rs3825970 (*LARP6*) are eQTL loci significantly controlling expression of their respective genes in cerebellum, cortex and basal ganglia. rs2959174 (*LRRC49/THAP10*) and rs12932325 (*GTF3C1*) are significant eQTL controlling expression of nearby genes such as *LARP6* and *IL21R* respectively. rs11589562 (*MAST2*) and rs17662853 (*KANSL1*) are significant eQTLs controlling expression of the respective mapped genes as well as nearby genes. For the remaining loci, we observe eQTL in the vicinity (± 500kbp) controlling the expression of their respectively annotated genes in different brain regions pertinent to cognition. Thus, our eQTL analysis of the significant exome-wide variants show that the genes mapped to these variants are highly expressed in brain regions deemed responsible for completion of neuropsychological tasks corresponding to respective cognitive domains thus providing convincing relevance for the significance of our results. We find that variants in *APOC1* and *LRP1* act as significant eQTLs for regulating their expression in basal ganglia and cerebellar hemispheres (Supplementary Table 13), crucial to visual attention.^64^

Our study is the first-ever evidence of *LRP1* association with cognition. Furthermore, we find that six out of the eight *LRP1* SNPs which interact with *APOE* are rare and remaining two are of low frequency. Targeted experiments have shown roles for *APOC1* and *LRP1^65^* in cognitive decline or neurodegeneration, however, our interaction analysis (Fig. 3B, Table 3) is the first to identify SNPs in *APOC1* and *LRP1* acting in conjunction with *APOE* in governing cognitive abilities, thus providing direct evidence for the role of *LRP1* on cognition. In total, we have identified 14 pairwise interactions relevant to episodic memory, simple processing speed, visual attention between *APOE* and our exome-wide associated hits, many of them are interestingly rare (allele frequency 0.12-4%). Our study is the first to report evidence of interactions between *APOE* and *AMIGO1, PTPN18, ITPR3. GPR108* in influencing cognition or neurodegeneration.

Despite several strengths of this study, we acknowledge the fact that the results reported herein must be considered in the light of some limitations. Firstly, even though the initial sample size is quite large (~157,000), effective sample sizes varies for each test (~27000 −121,000) is lesser because we have ensured that each participant has non-missing data on all variables of interest (phenotype and covariates) for all models. Secondly, our analyses has been based on individuals from European ancestry only. So caution must be exercised while generalizing the results for diverse ancestries.

## Abbreviations

CADD: Combined Annotation Dependent Depletion
eQTL: Expression Quantitative Trait Locus
LD: Linkage disequilibrium
GWAS: Genome-wide association studies
HDL: High-density lipoprotein
LDL: low-density lipoprotein
HbA1c: hemoglobin A1C; glycated haemoglobin
SKAT: Sequence Kernel Association Test
FDA: Food and Drug Administration

## Web resources

UK Biobank: https://www.ukbiobank.ac.uk

1000 Genomes Project: https://www.internationalgenome.org/

LDtrait: https: https://ldlink.nci.nih.gov/?tab=ldtrait

Uniprot: https://www.uniprot.org/

GWAS ATLAS: https://atlas.ctglab.nl/

VEP (LofTool): https://asia.ensembl.org/Homo_sapiens/Tools/VEP/

Harmonizome: https://maayanlab.cloud/Harmonizome/

Gtex: https://gtexportal.org/home/

NCBI: https://www.ncbi.nlm.nih.gov/ dbSNP: https://www.ncbi.nlm.nih.gov/snp/

UCSC: https://genome.ucsc.edu/

## Acknowledgements

We are extremely thankful to Dr. Balaji Jayaprakash for his support and enthusiastic encouragement towards this work. We would also like to thank Mr. Sheldon D’Silva for his help in carrying out quality control processes for the genetic data. We are also grateful to our system administrators Mr. Naveenan Srinivasan, Mr Karthik Sundaram and Mr Anand Kumar E for their support in managing technical challenges encountered while carrying out this computational work. We also thank the funding agencies-Science & Engineering Research Board (SERB), Government of India (ECR/2018/001429), Department of Biotechnology, Government of India (BT/RLF/29/2016) and National Supercomputing Mission, Government of India (DST/NSM/R&D_HPC_Applications/2021/03.12) for funding equipment and data acquisition for this study.

We conducted this research using the UK Biobank Resource under application number 55652. resource. UK Biobank’s database includes blood samples, heart and brain scans, and genetic data of the 500,000 volunteer participants and is globally accessible to approved researchers undertaking public health-related research. UK Biobank recruited 500,000 people between 40 and 69 years of age from 2006–2010 across the UK. The organization has more than 150 dedicated members of staff based in multiple locations across the UK who collected and stored detailed information about their lifestyle, physical measures, and blood, urine, and saliva samples with their consent. Since its inception in April 2012, over 20,000 researchers from 90+ countries have been approved to use this resource, and more than 2000 peer-reviewed papers that used it have now been published. This resource thus significantly contributes to advances in modern medicine and treatment, enabling better understanding of the prevention and diagnosis of a wide range of severe and life-threatening illnesses—including cancer, heart diseases, and stroke. And to run its operations, the UK Biobank receives generous support from its founding funders, the Wellcome Trust and UK Medical Research Council, the British Heart Foundation, Cancer Research UK, Department of Health, the Northwest Regional Development Agency, and the Scottish Government. We thus extend our sincere gratitude to all UK Biobank participants, researchers, clinicians, technicians, administrative staff and funding authorities who enabled curation of this enriched biomedical resource.

## Author’s contributions

B.K. conceived and designed the study. S.C, and B.K. performed the data analysis. S.C, and B.K. wrote the manuscript. S.C., and B.K. prepared the figures. Both the authors have read and approved the final manuscript.

## Supplementary material

Supplementary material is available at *Brain* online.

## References

1. Davies G, Tenesa A, Payton A, et al. Genome-wide association studies establish that human intelligence is highly heritable and polygenic. Mol Psychiatry. 2011;16(10):996–1005. doi:10.1038/mp.2011.85

2. Davies G, Armstrong N, Bis JC, et al. Genetic contributions to variation in general cognitive function: a meta-analysis of genome-wide association studies in the CHARGE consortium (N=53 949). Mol Psychiatry. 2015;20(2):183–192. doi:10.1038/mp.2014.188

3. Trampush JW, Yang MLZ, Yu J, et al. GWAS meta-analysis reveals novel loci and genetic correlates for general cognitive function: a report from the COGENT consortium. Mol Psychiatry. 2017;22(3):336–345. doi:10.1038/mp.2016.244

4. Sniekers S, Stringer S, Watanabe K, et al. Genome-wide association meta-analysis of 78,308 individuals identifies new loci and genes influencing human intelligence. Nat Genet. 2017;49(7):1107–1112. doi:10.1038/ng.3869

5. Davies G, Lam M, Harris SE, et al. Study of 300,486 individuals identifies 148 independent genetic loci influencing general cognitive function. Nat Commun. 2018;9(1):2098. doi:10.1038/s41467-018-04362-x

6. Gupta A, Kahali B. Machine learning-based cognitive impairment classification with optimal combination of neuropsychological tests. Alzheimer’s & Dementia: Translational Research & Clinical Interventions. 2020;6(1). doi:10.1002/trc2.12049

7. Homann J, Osburg T, Ohlei O, et al. Genome-Wide Association Study of Alzheimer’s Disease Brain Imaging Biomarkers and Neuropsychological Phenotypes in the European Medical Information Framework for Alzheimer’s Disease Multimodal Biomarker Discovery Dataset. Front Aging Neurosci. 2022;14. doi:10.3389/fnagi.2022.840651

8. Davies G, Harris SE, Reynolds CA, et al. A genome-wide association study implicates the APOE locus in nonpathological cognitive ageing. Mol Psychiatry. 2014;19(1):76–87. doi:10.1038/mp.2012.159

9. Raisi-Estabragh Z, M’Charrak A, McCracken C, et al. Associations of cognitive performance with cardiovascular magnetic resonance phenotypes in the UK Biobank. Eur Heart J Cardiovasc Imaging. 2022;23(5):663–672. doi:10.1093/ehjci/jeab075

10. Lane RM, Farlow MR. Lipid homeostasis and apolipoprotein E in the development and progression of Alzheimer’s disease. J Lipid Res. 2005;46(5):949–968. doi:10.1194/jlr.M400486-JLR200

11. Bycroft C, Freeman C, Petkova D, et al. The UK Biobank resource with deep phenotyping and genomic data. Nature. 2018;562(7726):203–209. doi:10.1038/s41586-018-0579-z

12. Speed D, Hemani G, Johnson MR, Balding DJ. Improved Heritability Estimation from Genome-wide SNPs. The American Journal of Human Genetics. 2012;91(6):1011–1021. doi:10.1016/j.ajhg.2012.10.010

13. Watanabe K, Stringer S, Frei O, et al. A global overview of pleiotropy and genetic architecture in complex traits. Nat Genet. 2019;51(9):1339–1348. doi:10.1038/s41588-019-0481-0

14. Zhan X, Hu Y, Li B, Abecasis GR, Liu DJ. RVTESTS: an efficient and comprehensive tool for rare variant association analysis using sequence data. Bioinformatics. 2016;32(9):1423–1426. doi:10.1093/bioinformatics/btw079

15. Machiela MJ, Chanock SJ. LDlink: a web-based application for exploring population-specific haplotype structure and linking correlated alleles of possible functional variants. Bioinformatics. 2015;31(21):3555–3557. doi:10.1093/bioinformatics/btv402

16. Buniello A, MacArthur JAL, Cerezo M, et al. The NHGRI-EBI GWAS Catalog of published genome-wide association studies, targeted arrays and summary statistics 2019. Nucleic Acids Res. 2019;47(D1):D1005-D1012. doi:10.1093/nar/gky1120

17. Wu MC, Lee S, Cai T, Li Y, Boehnke M, Lin X. Rare-variant association testing for sequencing data with the sequence kernel association test. Am J Hum Genet. 2011;89(1):82–93. doi:10.1016/j.ajhg.2011.05.029

18. Lee S, Emond MJ, Bamshad MJ, et al. Optimal unified approach for rare-variant association testing with application to small-sample case-control whole-exome sequencing studies. Am J Hum Genet. 2012;91(2):224–237. doi:10.1016/j.ajhg.2012.06.007

19. Purcell S, Neale B, Todd-Brown K, et al. PLINK: a tool set for whole-genome association and population-based linkage analyses. Am J Hum Genet. 2007;81(3):559–575. doi:10.1086/519795

20. Ray D, Boehnke M. Methods for meta-analysis of multiple traits using GWAS summary statistics. Genet Epidemiol. 2018;42(2):134–145. doi:10.1002/gepi.22105

21. Kircher M, Witten DM, Jain P, O’Roak BJ, Cooper GM, Shendure J. A general framework for estimating the relative pathogenicity of human genetic variants. Nat Genet. 2014;46(3):310–315. doi:10.1038/ng.2892

22. Lonsdale J, Thomas J, Salvatore M, et al. The Genotype-Tissue Expression (GTEx) project. Nat Genet. 2013;45(6):580–585. doi:10.1038/ng.2653

23. Junghans D, Heidenreich M, Hack I, Taylor V, Frotscher M, Kemler R. Postsynaptic and differential localization to neuronal subtypes of protocadherin β16 in the mammalian central nervous system. European Journal of Neuroscience. 2008;27(3):559–571. doi:10.1111/j.1460-9568.2008.06052.x

24. Giordano G, Cole TB, Furlong CE, Costa LG. Paraoxonase 2 (PON2) in the mouse central nervous system: A neuroprotective role? Toxicol Appl Pharmacol. 2011;256(3):369–378. doi:10.1016/j.taap.2011.02.014

25. Qu WR, Sun QH, Liu QQ, et al. Role of CPEB3 protein in learning and memory: new insights from synaptic plasticity. Aging. 2020;12(14):15169–15182. doi:10.18632/aging.103404

26. Moreno-Igoa M, Hernández-Charro B, Bengoa-Alonso A, et al. KANSL1 gene disruption associated with the full clinical spectrum of 17q21.31 microdeletion syndrome. BMC Med Genet. 2015;16(1):68. doi:10.1186/s12881-015-0211-0

27. Linda K, Lewerissa EI, Verboven AHA, et al. Imbalanced autophagy causes synaptic deficits in a human model for neurodevelopmental disorders. Autophagy. 2022;18(2):423–442. doi:10.1080/15548627.2021.1936777

28. Zhou X, Wang L, Xiao W, et al. Memantine Improves Cognitive Function and Alters Hippocampal and Cortical Proteome in Triple Transgenic Mouse Model of Alzheimer’s Disease. Exp Neurobiol. 2019;28(3):390–403. doi:10.5607/en.2019.28.3.390

29. Miller JE, Shivakumar MK, Risacher SL, et al. Codon bias among synonymous rare variants is associated with Alzheimer’s disease imaging biomarker. Pac Symp Biocomput. 2018;23:365–376.

30. Velayudhan L, Proitsi P, Westman E, et al. Entorhinal cortex thickness predicts cognitive decline in Alzheimer’s disease. J Alzheimers Dis. 2013;33(3):755–766. doi:10.3233/JAD-2012-121408

31. Kusakari S, Saitow F, Ago Y, et al. Shp2 in Forebrain Neurons Regulates Synaptic Plasticity, Locomotion, and Memory Formation in Mice. Mol Cell Biol. 2015;35(9):1557–1572. doi:10.1128/MCB.01339-14

32. Kim Y, Liu G, Leugers CJ, et al. Tau interacts with SHP2 in neuronal systems and in Alzheimer’s disease. J Cell Sci. Published online January 1, 2019. doi:10.1242/jcs.229054

33. Johnson EM, Ishak AD, Naylor PE, Stevenson DA, Reiss AL, Green T. PTPN11 Gain-of-Function Mutations Affect the Developing Human Brain, Memory, and Attention. Cerebral Cortex. 2019;29(7):2915–2923. doi:10.1093/cercor/bhy158

34. Linglart L, Gelb BD. Congenital heart defects in Noonan syndrome: Diagnosis, management, and treatment. Am J Med Genet C Semin Med Genet. 2020;184(1):73–80. doi:10.1002/ajmg.c.31765

35. Xie X, Liang M, Yu C, Wei Z. Liprin-α-Mediated Assemblies and Their Roles in Synapse Formation. Front Cell Dev Biol. 2021;9. doi:10.3389/fcell.2021.653381

36. Scholz CJ, Weber H, Jungwirth S, et al. Explorative results from multistep screening for potential genetic risk loci of Alzheimer’s disease in the longitudinal VITA study cohort. J Neural Transm. 2018;125(1):77–87. doi:10.1007/s00702-017-1796-6

37. Bayraktar A, Lam S, Altay O, et al. Revealing the Molecular Mechanisms of Alzheimer’s Disease Based on Network Analysis. Int J Mol Sci. 2021;22(21):11556. doi:10.3390/ijms222111556

38. Stewart Alexandre FR, Chen HH. Activation of tyrosine phosphatases in the progression of Alzheimer’s disease. Neural Regen Res. 2020;15(12):2245. doi:10.4103/1673-5374.284986

39. Bartok A, Weaver D, Golenár T, et al. IP3 receptor isoforms differently regulate ER-mitochondrial contacts and local calcium transfer. Nat Commun. 2019;10(1):3726. doi:10.1038/s41467-019-11646-3

40. Wang X, Zhou X, Uberseder B, et al. Isoform-specific dysregulation of AMP-activated protein kinase signaling in a non-human primate model of Alzheimer’s disease. Neurobiol Dis. 2021;158:105463. doi:10.1016/j.nbd.2021.105463

41. Schon EA, Area-Gomez E. Mitochondria-associated ER membranes in Alzheimer disease. Molecular and Cellular Neuroscience. 2013;55:26–36. doi:10.1016/j.mcn.2012.07.011

42. Ligthart S, de Vries PS, Uitterlinden AG, et al. Pleiotropy among Common Genetic Loci Identified for Cardiometabolic Disorders and C-Reactive Protein. PLoS One. 2015;10(3):e0118859. doi:10.1371/journal.pone.0118859

43. Fuior E v., Gafencu A v. Apolipoprotein C1: Its Pleiotropic Effects in Lipid Metabolism and Beyond. Int J Mol Sci. 2019;20(23):5939. doi:10.3390/ijms20235939

44. de la Fuente J, Davies G, Grotzinger AD, Tucker-Drob EM, Deary IJ. A general dimension of genetic sharing across diverse cognitive traits inferred from molecular data. Nat Hum Behav. 2021;5(1):49–58. doi:10.1038/s41562-020-00936-2

45. Liu C, Chyr J, Zhao W, et al. Genome-Wide Association and Mechanistic Studies Indicate That Immune Response Contributes to Alzheimer’s Disease Development. Front Genet. 2018;9. doi:10.3389/fgene.2018.00410

46. Cao W, Tian S, Zhang H, et al. Association of Low-Density Lipoprotein Receptor-Related Protein 1 and Its rs1799986 Polymorphism With Mild Cognitive Impairment in Chinese Patients With Type 2 Diabetes. Front Neurosci. 2020;14. doi:10.3389/fnins.2020.00743

47. Shi YM, Zhou H, Zhang ZJ, et al. Association of the LRP1 gene and cognitive performance with amnestic mild cognitive impairment in elderly Chinese. Int Psychogeriatr. 2009;21(6):1072–1080. doi:10.1017/S104161020999072X

48. Shinohara M, Tachibana M, Kanekiyo T, Bu G. Role of LRP1 in the pathogenesis of Alzheimer’s disease: evidence from clinical and preclinical studies. J Lipid Res. 2017;58(7): 1267-1281. doi:10.1194/jlr.R075796

49. Yoon YB, Shin WG, Lee TY, et al. Brain Structural Networks Associated with Intelligence and Visuomotor Ability. Sci Rep. 2017;7(1):2177. doi:10.1038/s41598-017-02304-z

50. Chen PY, Chen CL, Hsu YC, Tseng WYI. Fluid intelligence is associated with cortical volume and white matter tract integrity within multiple-demand system across adult lifespan. Neuroimage. 2020;212:116576. doi:10.1016/j.neuroimage.2020.116576

51. Burgaleta M, MacDonald PA, Martínez K, et al. Subcortical regional morphology correlates with fluid and spatial intelligence. Hum Brain Mapp. 2014;35(5):1957–1968. doi:10.1002/hbm.22305

52. Rhein C, Mühle C, Richter-Schmidinger T, Alexopoulos P, Doerfler A, Kornhuber J. Neuroanatomical Correlates of Intelligence in Healthy Young Adults: The Role of Basal Ganglia Volume. PLoS One. 2014;9(4):e93623. doi:10.1371/journal.pone.0093623

53. Paraskevopoulou SE, Coon WG, Brunner P, Miller KJ, Schalk G. Within-subject reaction time variability: Role of cortical networks and underlying neurophysiological mechanisms. Neuroimage. 2021;237:118127. doi:10.1016/j.neuroimage.2021.118127

54. Coon WG, Gunduz A, Brunner P, Ritaccio AL, Pesaran B, Schalk G. Oscillatory phase modulates the timing of neuronal activations and resulting behavior. Neuroimage. 2016;133:294–301. doi:10.1016/j.neuroimage.2016.02.080

55. Rouillard AD, Gundersen GW, Fernandez NF, et al. The harmonizome: a collection of processed datasets gathered to serve and mine knowledge about genes and proteins. Database. 2016;2016:baw100. doi:10.1093/database/baw100

56. Agrawal S, Baulch JE, Madan S, et al. Impact of IL-21-associated peripheral and brain crosstalk on the Alzheimer’s disease neuropathology. Cellular and Molecular Life Sciences. 2022;79(6):331. doi:10.1007/s00018-022-04347-6

57. Nguyen H, Hall B, Higginson CI, Sigvardt KA, Zweig R, Disbrow EA. Theory of Cognitive Aging in Parkinson Disease. J Alzheimers Dis Parkinsonism. 2017;7(5). doi:10.4172/2161-0460.1000369

58. Chen Y, Hor HH, Tang BL. AMIGO is expressed in multiple brain cell types and may regulate dendritic growth and neuronal survival. J Cell Physiol. 2012;227(5):2217–2229. doi:10.1002/jcp.22958

59. Dickerson BC, Eichenbaum H. The Episodic Memory System: Neurocircuitry and Disorders. Neuropsychopharmacology. 2010;35(1):86–104. doi:10.1038/npp.2009.126

60. Meenakshi P, Balaji J. Neural Circuits of Memory Consolidation and Generalisation. J Indian Inst Sci. 2017;97(4):487–495. doi:10.1007/s41745-017-0042-4

61. Andreasen NC, O’Leary DS, Paradiso S, et al. The cerebellum plays a role in conscious episodic memory retrieval. Hum Brain Mapp. 1999;8(4):226–234. doi:10.1002/(SICI)1097-0193(1999)8:4<226::AID-HBM6>3.0.CO;2-4

62. Kahn I, Shohamy D. Intrinsic connectivity between the hippocampus, nucleus accumbens, and ventral tegmental area in humans. Hippocampus. 2013;23(3):187–192. doi:10.1002/hipo.22077

63. Das M, Bennett DM, Dutton GN. Visual attention as an important visual function: an outline of manifestations, diagnosis and management of impaired visual attention. British Journal of Ophthalmology. 2007;91(11):1556–1560. doi:10.1136/bjo.2006.104844

64. Lockhofen DEL, Mulert C. Neurochemistry of Visual Attention. Front Neurosci. 2021; 15. doi:10.3389/fnins.2021.643597

65. Zang F, Zhu Y, Zhang Q, Tan C, Wang Q, Xie C. APOE genotype moderates the relationship between LRP1 polymorphism and cognition across the Alzheimer’s disease spectrum via disturbing default mode network. CNS Neurosci Ther. 2021;27(11):1385–1395. doi:10.1111/cns.13716

