## Supplementary table 11 for "Exome-wide analysis reveals role of *LRP1* and additional novel loci in cognition"

**Suuplementary Table 11: Single variant association hits which are eQTLs controlling expression of relevant genes in brain tissues**

**Header Descriptions**

Mapped Gene: Genes to which our single variant hits have been annotated
SNP Id: rsid of significant eQTLs
NES: Normalized effect size of association of the significant eQTLs with tissue expression
P-value: p-value of the association

| **Cognition Domain** | **Cognition-associated SNPs** | **Novelty for cognition** | **Mapped Gene** | **Gene Symbol** | **SNP ID** | **P-Value** | **NES** | **Tissue** |
|---|---|---|---|---|---|---|---|---|
| Fluid Intelligence/ Reasoning | rs3824734 | Novel | *CPEB3* | *CPEB3* | rs3824734 | 3.80E-05 | 0.22 | Brain Cerebellar Hemisphere |
| Visual Attention | rs11589562 | Novel | *MAST2* | *CCDC163* | rs11589562 | 2.10E-11 | -0.69 | Brain-Cerebellum |
| Visual Attention | rs11589562 | Novel | *MAST2* | *CCDC163* | rs11589562 | 4.20E-11 | -0.55 | Brain-Cortex |
| Visual Attention | rs11589562 | Novel | *MAST2* | *CCDC163* | rs11589562 | 1.40E-10 | -0.55 | Brain-Caudate (basal ganglia) |
| Visual Attention | rs11589562 | Novel | *MAST2* | *CCDC163* | rs11589562 | 8.10E-10 | -0.56 | Brain-Anterior cingulate cortex (BA24) |
| Visual Attention | rs11589562 | Novel | *MAST2* | *CCDC163* | rs11589562 | 3.10E-09 | -0.52 | Brain-Frontal Cortex (BA9) |
| Visual Attention | rs11589562 | Novel | *MAST2* | *CCDC163* | rs11589562 | 3.80E-09 | -0.73 | Brain-Cerebellar Hemisphere |
| Visual Attention | rs11589562 | Novel | *MAST2* | *CCDC163* | rs11589562 | 7.30E-09 | -0.59 | Brain-Nucleus accumbens (basal ganglia) |
| Visual Attention | rs11589562 | Novel | *MAST2* | *CCDC163* | rs11589562 | 1.50E-08 | -0.63 | Brain-Putamen (basal ganglia) |
| Visual Attention | rs11589562 | Novel | *MAST2* | *TESK2* | rs11589562 | 3.40E-06 | -0.27 | Brain-Caudate (basal ganglia) |
| Visual Attention | rs11589562 | Novel | *MAST2* | *MAST2* | rs11589562 | 4.60E-06 | 0.26 | Brain-Cerebellum |
| Visual Attention | rs11589562 | Novel | *MAST2* | *CCDC163* | rs11589562 | 6.60E-06 | -0.36 | Brain-Hippocampus |
| Visual Attention | rs11589562 | Novel | *MAST2* | *MAST2* | rs11589562 | 7.70E-06 | 0.21 | Brain-Cortex |
| Visual Attention | rs11589562 | Novel | *MAST2* | *PIK3R3* | rs11589562 | 6.60E-05 | 0.28 | Brain-Cortex |
| Simple Processing Speed | rs3813363 | - | *SAMD3* | *SAMD3* | rs3813363 | 1.10E-05 | -0.4 | Brain-Cortex |
| Simple Processing Speed | rs17662853 | - | *KANSL1* | *NSFP1* | rs17662853 | 3.90E-20 | 1.2 | Brain- Cortex |
| Simple Processing Speed | rs17662853 | - | *KANSL1* | *NSFP1* | rs17662853 | 1.00E-19 | 1.2 | Brain- Cerebellum |
| Simple Processing Speed | rs17662853 | - | *KANSL1* | *NSFP1* | rs17662853 | 6.60E-19 | 1.1 | Brain- Nucleus accumbens (basal ganglia) |
| Simple Processing Speed | rs17662853 | - | *KANSL1* | *NSFP1* | rs17662853 | 2.90E-16 | 1.2 | Brain- Caudate (basal ganglia) |
| Simple Processing Speed | rs17662853 | - | *KANSL1* | *NSFP1* | rs17662853 | 6.90E-15 | 1.2 | Brain- Cerebellar Hemisphere |
| Simple Processing Speed | rs17662853 | - | *KANSL1* | *NSFP1* | rs17662853 | 7.30E-15 | 1.2 | Brain- Putamen (basal ganglia) |

| | | | | | | | | |
|---|---|---|---|---|---|---|---|---|
| Simple Processing Speed | rs17662853 | - | *KANSL1* | *NSFP1* | rs17662853 | 1.20E-14 | 1.1 | Brain- Anterior cingulate cortex (BA24) |
| Simple Processing Speed | rs17662853 | - | *KANSL1* | *NSFP1* | rs17662853 | 1.40E-13 | 1.3 | Brain- Amygdala |
| Simple Processing Speed | rs17662853 | - | *KANSL1* | *NSFP1* | rs17662853 | 1.50E-13 | 1.1 | Brain- Hypothalamus |
| Simple Processing Speed | rs17662853 | - | *KANSL1* | *NSFP1* | rs17662853 | 5.90E-13 | 1.1 | Brain- Frontal Cortex (BA9) |
| Simple Processing Speed | rs17662853 | - | *KANSL1* | *NSFP1* | rs17662853 | 1.00E-11 | 1.1 | Brain- Hippocampus |
| Simple Processing Speed | rs17662853 | - | *KANSL1* | *ARL17B* | rs17662853 | 3.50E-11 | 1.1 | Brain- Caudate (basal ganglia) |
| Simple Processing Speed | rs17662853 | - | *KANSL1* | *LRRC37A* | rs17662853 | 7.90E-11 | 0.87 | Brain- Cortex |
| Simple Processing Speed | rs17662853 | - | *KANSL1* | *ARL17B* | rs17662853 | 9.40E-11 | 0.92 | Brain- Nucleus accumbens (basal ganglia) |
| Simple Processing Speed | rs17662853 | - | *KANSL1* | *ARL17B* | rs17662853 | 9.80E-11 | 0.93 | Brain- Cortex |
| Simple Processing Speed | rs17662853 | - | *KANSL1* | *ARL17B* | rs17662853 | 1.10E-10 | 0.95 | Brain- Putamen (basal ganglia) |
| Simple Processing Speed | rs17662853 | - | *KANSL1* | *NSFP1* | rs17662853 | 1.10E-10 | 1 | Brain- Spinal cord (cervical c-1) |
| Simple Processing Speed | rs17662853 | - | *KANSL1* | *LRRC37A* | rs17662853 | 5.60E-09 | 0.85 | Brain- Nucleus accumbens (basal ganglia) |
| Simple Processing Speed | rs17662853 | - | *KANSL1* | *LRRC37A* | rs17662853 | 1.80E-08 | 0.91 | Brain- Putamen (basal ganglia) |
| Simple Processing Speed | rs17662853 | - | *KANSL1* | *LRRC37A* | rs17662853 | 4.40E-08 | 0.88 | Brain- Anterior cingulate cortex (BA24) |
| Simple Processing Speed | rs17662853 | - | *KANSL1* | *ARL17B* | rs17662853 | 8.70E-08 | 0.78 | Brain- Cerebellum |
| Simple Processing Speed | rs17662853 | - | *KANSL1* | *LRRC37A* | rs17662853 | 1.10E-07 | 0.85 | Brain- Caudate (basal ganglia) |
| Simple Processing Speed | rs17662853 | - | *KANSL1* | *ARL17B* | rs17662853 | 6.10E-07 | 0.77 | Brain- Anterior cingulate cortex (BA24) |
| Simple Processing Speed | rs17662853 | - | *KANSL1* | *ARL17B* | rs17662853 | 8.60E-07 | 0.82 | Brain- Hypothalamus |
| Simple Processing Speed | rs17662853 | - | *KANSL1* | *RP11-798G7.*[illegible] | rs17662853 | 8.80E-07 | -0.62 | Brain- Cerebellum |
| Simple Processing Speed | rs17662853 | - | *KANSL1* | *LRRC37A* | rs17662853 | 9.70E-07 | 0.83 | Brain- Hypothalamus |
| Simple Processing Speed | rs17662853 | - | *KANSL1* | *KANSL1-AS1* | rs17662853 | 1.50E-06 | -0.6 | Brain- Nucleus accumbens (basal ganglia) |
| Simple Processing Speed | rs17662853 | - | *KANSL1* | *RP11-798G7.*[illegible] | rs17662853 | 2.00E-06 | -0.7 | Brain- Putamen (basal ganglia) |
| Simple Processing Speed | rs17662853 | - | *KANSL1* | *RP11-798G7.*[illegible] | rs17662853 | 4.50E-06 | -0.59 | Brain- Cortex |
| Simple Processing Speed | rs17662853 | - | *KANSL1* | *ARL17B* | rs17662853 | 6.60E-06 | 0.78 | Brain- Frontal Cortex (BA9) |
| Simple Processing Speed | rs17662853 | - | *KANSL1* | *LRRC37A* | rs17662853 | 7.90E-06 | 0.8 | Brain- Frontal Cortex (BA9) |
| Simple Processing Speed | rs17662853 | - | *KANSL1* | *NSFP1* | rs17662853 | 1.10E-05 | 0.84 | Brain- Substantia nigra |
| Simple Processing Speed | rs17662853 | - | *KANSL1* | *RP11-798G7.*[illegible] | rs17662853 | 1.20E-05 | -0.6 | Brain- Nucleus accumbens (basal ganglia) |
| Simple Processing Speed | rs17662853 | - | *KANSL1* | *ARL17B* | rs17662853 | 1.40E-05 | 0.86 | Brain- Amygdala |
| Simple Processing Speed | rs17662853 | - | *KANSL1* | *KANSL1-AS1* | rs17662853 | 1.50E-05 | -0.58 | Brain- Cerebellum |
| Simple Processing Speed | rs17662853 | - | *KANSL1* | *ARL17B* | rs17662853 | 1.70E-05 | 0.76 | Brain- Hippocampus |
| Simple Processing Speed | rs17662853 | - | *KANSL1* | *ARL17A* | rs17662853 | 1.90E-05 | -0.57 | Brain- Cerebellum |
| Simple Processing Speed | rs17662853 | - | *KANSL1* | *KANSL1* | rs17662853 | 2.30E-05 | -0.4 | Brain- Cerebellum |

| | | | | | | | | |
|---|---|---|---|---|---|---|---|---|
| Simple Processing Speed | rs17662853 | - | *KANSL1* | *KANSL1* | rs17662853 | 2.50E-05 | -0.49 | Brain- Anterior cingulate cortex (BA24) |
| Simple Processing Speed | rs17662853 | - | *KANSL1* | *NSF* | rs17662853 | 3.30E-05 | 0.2 | Brain- Cerebellum |
| Simple Processing Speed | rs17662853 | - | *KANSL1* | *NSF* | rs17662853 | 3.70E-05 | 0.23 | Brain- Cerebellar Hemisphere |
| Simple Processing Speed | rs17662853 | - | *KANSL1* | *RP11-798G7.* | rs17662853 | 4.70E-05 | -0.65 | Brain- Cerebellar Hemisphere |
| Simple Processing Speed | rs17662853 | - | *KANSL1* | *RP11-798G7.* | rs17662853 | 4.90E-05 | -0.63 | Brain- Caudate (basal ganglia) |
| Simple Processing Speed | rs17662853 | - | *KANSL1* | *KANSL1-AS1* | rs17662853 | 9.20E-05 | -0.51 | Brain- Cortex |
| Simple Processing Speed | rs17662853 | - | *KANSL1* | *ARL17B* | rs17662853 | 1.10E-04 | 0.62 | Brain- Cerebellar Hemisphere |
| Simple Processing Speed | rs17662853 | - | *KANSL1* | *FAM215B* | rs17662853 | 1.40E-04 | -0.47 | Brain- Cerebellum |
| Simple Processing Speed | rs2959174 | Novel | *LRRC49* | *LARP6* | rs2959174 | 8.90E-18 | 0.49 | Brain-Cerebellum |
| Simple Processing Speed | rs2959174 | Novel | *LRRC49* | *LARP6* | rs2959174 | 4.10E-10 | 0.39 | Brain-Cerebellar Hemisphere |
| Simple Processing Speed | rs2959174 | Novel | *LRRC49* | *LARP6* | rs2959174 | 5.30E-10 | 0.2 | Brain-Putamen (basal ganglia) |
| Simple Processing Speed | rs2959174 | Novel | *LRRC49* | *LARP6* | rs2959174 | 2.20E-08 | 0.22 | Brain-Hippocampus |
| Simple Processing Speed | rs2959174 | Novel | *LRRC49* | *LARP6* | rs2959174 | 1.60E-06 | 0.35 | Brain-Spinal cord (cervical c-1) |
| Simple Processing Speed | rs2959174 | Novel | *LRRC49* | *LARP6* | rs2959174 | 1.10E-05 | 0.13 | Brain-Cortex |
| Simple Processing Speed | rs3825970 | Novel | *LARP6* | *LARP6* | rs3825970 | 2.60E-22 | -0.52 | Brain- Cerebellum |
| Simple Processing Speed | rs3825970 | Novel | *LARP6* | *LARP6* | rs3825970 | 7.80E-16 | -0.46 | Brain- Cerebellar Hemisphere |
| Simple Processing Speed | rs3825970 | Novel | *LARP6* | *LARP6* | rs3825970 | 2.70E-06 | -0.15 | Brain-Putamen (basal ganglia) |
| Complex Processing Speed | rs12932325 | Novel | *GTF3C1* | *IL21R-AS1* | rs12932325 | 9.30E-46 | -1.4 | Brain-Cerebellum |
| Complex Processing Speed | rs12932325 | Novel | *GTF3C1* | *IL21R-AS1* | rs12932325 | 2.20E-41 | -1.3 | Brain-Cortex |
| Complex Processing Speed | rs12932325 | Novel | *GTF3C1* | *IL21R-AS1* | rs12932325 | 3.40E-36 | -0.97 | Brain-Nucleus accumbens (basal ganglia) |
| Complex Processing Speed | rs12932325 | Novel | *GTF3C1* | *IL21R-AS1* | rs12932325 | 1.60E-33 | -1.1 | Brain-Caudate (basal ganglia) |
| Complex Processing Speed | rs12932325 | Novel | *GTF3C1* | *IL21R-AS1* | rs12932325 | 5.00E-32 | -1.2 | Brain-Cerebellar Hemisphere |
| Complex Processing Speed | rs12932325 | Novel | *GTF3C1* | *IL21R-AS1* | rs12932325 | 4.20E-31 | -1.1 | Brain-Frontal Cortex (BA9) |
| Complex Processing Speed | rs12932325 | Novel | *GTF3C1* | *IL21R-AS1* | rs12932325 | 4.50E-27 | -1 | Brain-Hypothalamus |
| Complex Processing Speed | rs12932325 | Novel | *GTF3C1* | *IL21R* | rs12932325 | 2.20E-25 | -1.1 | Brain-Cerebellum |
| Complex Processing Speed | rs12932325 | Novel | *GTF3C1* | *IL21R-AS1* | rs12932325 | 6.50E-23 | -1.1 | Brain-Hippocampus |
| Complex Processing Speed | rs12932325 | Novel | *GTF3C1* | *IL21R-AS1* | rs12932325 | 9.80E-23 | -1.2 | Brain-Anterior cingulate cortex (BA24) |
| Complex Processing Speed | rs12932325 | Novel | *GTF3C1* | *IL21R-AS1* | rs12932325 | 4.70E-19 | -0.96 | Brain-Putamen (basal ganglia) |
| Complex Processing Speed | rs12932325 | Novel | *GTF3C1* | *IL21R-AS1* | rs12932325 | 2.00E-17 | -0.93 | Brain-Amygdala |
| Complex Processing Speed | rs12932325 | Novel | *GTF3C1* | *IL21R* | rs12932325 | 2.00E-17 | -1.2 | Brain-Cerebellar Hemisphere |
| Complex Processing Speed | rs12932325 | Novel | *GTF3C1* | *IL21R-AS1* | rs12932325 | 1.90E-14 | -1.1 | Brain-Substantia nigra |
| Complex Processing Speed | rs12932325 | Novel | *GTF3C1* | *IL21R-AS1* | rs12932325 | 6.10E-13 | -0.95 | Brain-Spinal cord (cervical c-1) |

| | | | | | | | | |
|---|---|---|---|---|---|---|---|---|
| Complex Processing Speed | rs12932325 | Novel | *GTF3C1* | *IL21R* | rs12932325 | 4.10E-11 | -0.64 | Brain-Nucleus accumbens (basal ganglia) |
| Complex Processing Speed | rs12932325 | Novel | *GTF3C1* | *IL21R* | rs12932325 | 1.00E-10 | -0.73 | Brain-Cortex |
| Complex Processing Speed | rs12932325 | Novel | *GTF3C1* | *IL21R* | rs12932325 | 1.40E-07 | -0.59 | Brain-Frontal Cortex (BA9) |
| Complex Processing Speed | rs12932325 | Novel | *GTF3C1* | *IL21R* | rs12932325 | 2.00E-07 | -0.64 | Brain-Hippocampus |
| Complex Processing Speed | rs12932325 | Novel | *GTF3C1* | *IL21R* | rs12932325 | 1.50E-05 | -0.45 | Brain-Caudate (basal ganglia) |
