## Supplementary table 12 for "Exome-wide analysis reveals role of *LRP1* and additional novel loci in cognition"

**Suuplementary Table 12: eQTLs in genes mapped to novel variants discovered in our analysis**

**Header Descriptions**

Gencode ID, Gene Symbol for mapped gene: Gene ID and Symbol of genes to which our novel variants were mapped.
SNP Id: rsid of significant eQTLs
NES: Normalized effect size of association of the significant eQTLs with tissue expression
P-value: p-value of the association
Query Pos:
Query Start, Query End: -+500kbp from novel variants identified in our study
Within-Yes/No

| Gencode Id | Gene Symbol for Mapped Gene | SNP Id | P-Value | NES | Tissue | Query Pos | Query Start | Query End | Within-Yes/No |
|---|---|---|---|---|---|---|---|---|---|
| ENSG00000272674.3 | PCDHB16 | rs13171154 | 2.60E-05 | -0.45 | Brain - Cerebellar Hemisphere | 141185287 | 140685287 | 141685287 | No |
| ENSG00000272674.3 | PCDHB16 | rs71592123 | 2.60E-05 | -0.45 | Brain - Cerebellar Hemisphere | 141185287 | 140685287 | 141685287 | No |
| ENSG00000272674.3 | PCDHB16 | rs12523459 | 4.60E-05 | -1.1 | Brain - Cerebellar Hemisphere | 141185287 | 140685287 | 141685287 | No |
| ENSG00000272674.3 | PCDHB16 | rs13180584 | 2.50E-05 | -0.44 | Brain - Cerebellar Hemisphere | 141185287 | 140685287 | 141685287 | No |
| ENSG00000272674.3 | PCDHB16 | rs10463297 | 2.50E-05 | -0.44 | Brain - Cerebellar Hemisphere | 141185287 | 140685287 | 141685287 | No |
| ENSG00000272674.3 | PCDHB16 | rs11167532 | 8.90E-06 | -0.45 | Brain - Cerebellar Hemisphere | 141185287 | 140685287 | 141685287 | No |
| ENSG00000272674.3 | PCDHB16 | rs34517992 | 6.60E-06 | -0.46 | Brain - Cerebellar Hemisphere | 141185287 | 140685287 | 141685287 | No |
| ENSG00000272674.3 | PCDHB16 | rs2915863 | 9.30E-06 | 0.45 | Brain - Cerebellar Hemisphere | 141185287 | 140685287 | 141685287 | No |
| ENSG00000272674.3 | PCDHB16 | rs251350 | 5.60E-05 | 0.33 | Brain - Cerebellar Hemisphere | 141185287 | 140685287 | 141685287 | Yes |
| ENSG00000272674.3 | PCDHB16 | rs251351 | 7.90E-05 | 0.35 | Brain - Cerebellar Hemisphere | 141185287 | 140685287 | 141685287 | Yes |
| ENSG00000272674.3 | PCDHB16 | rs251352 | 8.90E-05 | 0.33 | Brain - Cerebellar Hemisphere | 141185287 | 140685287 | 141685287 | Yes |
| ENSG00000272674.3 | PCDHB16 | rs251353 | 4.10E-05 | 0.34 | Brain - Cerebellar Hemisphere | 141185287 | 140685287 | 141685287 | Yes |
| ENSG00000272674.3 | PCDHB16 | rs251354 | 5.50E-05 | 0.36 | Brain - Cerebellar Hemisphere | 141185287 | 140685287 | 141685287 | Yes |
| ENSG00000272674.3 | PCDHB16 | rs251355 | 2.60E-05 | 0.35 | Brain - Cerebellar Hemisphere | 141185287 | 140685287 | 141685287 | Yes |
| ENSG00000272674.3 | PCDHB16 | rs369636 | 5.50E-05 | 0.36 | Brain - Cerebellar Hemisphere | 141185287 | 140685287 | 141685287 | Yes |
| ENSG00000272674.3 | PCDHB16 | rs369639 | 5.50E-05 | 0.36 | Brain - Cerebellar Hemisphere | 141185287 | 140685287 | 141685287 | Yes |
| ENSG00000272674.3 | PCDHB16 | rs577069 | 1.30E-05 | 0.36 | Brain - Cerebellar Hemisphere | 141185287 | 140685287 | 141685287 | Yes |
| ENSG00000272674.3 | PCDHB16 | rs251358 | 4.50E-06 | 0.37 | Brain - Cerebellar Hemisphere | 141185287 | 140685287 | 141685287 | Yes |

| | | | | | | | | | |
|---|---|---|---|---|---|---|---|---|---|
| ENSG00000272674.3 | PCDHB16 | rs251359 | 1.70E-05 | 0.35 | Brain - Cerebellar Hemisphere | 141185287 | 140685287 | 141685287 | Yes |
| ENSG00000272674.3 | PCDHB16 | rs251360 | 3.80E-05 | 0.36 | Brain - Cerebellar Hemisphere | 141185287 | 140685287 | 141685287 | Yes |
| ENSG00000272674.3 | PCDHB16 | rs251361 | 6.80E-06 | 0.37 | Brain - Cerebellar Hemisphere | 141185287 | 140685287 | 141685287 | Yes |
| ENSG00000272674.3 | PCDHB16 | rs251362 | 6.80E-06 | 0.37 | Brain - Cerebellar Hemisphere | 141185287 | 140685287 | 141685287 | Yes |
| ENSG00000272674.3 | PCDHB16 | rs251363 | 3.80E-05 | 0.36 | Brain - Cerebellar Hemisphere | 141185287 | 140685287 | 141685287 | Yes |
| ENSG00000272674.3 | PCDHB16 | rs251364 | 3.80E-05 | 0.36 | Brain - Cerebellar Hemisphere | 141185287 | 140685287 | 141685287 | Yes |
| ENSG00000272674.3 | PCDHB16 | rs630162 | 3.80E-05 | 0.36 | Brain - Cerebellar Hemisphere | 141185287 | 140685287 | 141685287 | Yes |
| ENSG00000272674.3 | PCDHB16 | rs251365 | 2.10E-05 | 0.41 | Brain - Cerebellar Hemisphere | 141185287 | 140685287 | 141685287 | Yes |
| ENSG00000272674.3 | PCDHB16 | rs251367 | 2.10E-05 | 0.41 | Brain - Cerebellar Hemisphere | 141185287 | 140685287 | 141685287 | Yes |
| ENSG00000272674.3 | PCDHB16 | rs3834243 | 2.10E-05 | 0.41 | Brain - Cerebellar Hemisphere | 141185287 | 140685287 | 141685287 | Yes |
| ENSG00000272674.3 | PCDHB16 | rs11321479 | 2.10E-05 | 0.41 | Brain - Cerebellar Hemisphere | 141185287 | 140685287 | 141685287 | Yes |
| ENSG00000272674.3 | PCDHB16 | rs251369 | 2.10E-05 | 0.41 | Brain - Cerebellar Hemisphere | 141185287 | 140685287 | 141685287 | Yes |
| ENSG00000272674.3 | PCDHB16 | rs251371 | 2.10E-05 | 0.41 | Brain - Cerebellar Hemisphere | 141185287 | 140685287 | 141685287 | Yes |
| ENSG00000272674.3 | PCDHB16 | rs251374 | 1.80E-05 | 0.42 | Brain - Cerebellar Hemisphere | 141185287 | 140685287 | 141685287 | Yes |
| ENSG00000272674.3 | PCDHB16 | rs251378 | 2.10E-05 | 0.41 | Brain - Cerebellar Hemisphere | 141185287 | 140685287 | 141685287 | Yes |
| ENSG00000272674.3 | PCDHB16 | rs251379 | 3.50E-05 | 0.4 | Brain - Cerebellar Hemisphere | 141185287 | 140685287 | 141685287 | Yes |
| ENSG00000272674.3 | PCDHB16 | rs59479 | 2.10E-05 | 0.41 | Brain - Cerebellar Hemisphere | 141185287 | 140685287 | 141685287 | Yes |
| ENSG00000272674.3 | PCDHB16 | rs251380 | 2.10E-05 | 0.41 | Brain - Cerebellar Hemisphere | 141185287 | 140685287 | 141685287 | Yes |
| ENSG00000272674.3 | PCDHB16 | rs251381 | 2.10E-05 | 0.41 | Brain - Cerebellar Hemisphere | 141185287 | 140685287 | 141685287 | Yes |
| ENSG00000272674.3 | PCDHB16 | rs35160890 | 1.80E-05 | -0.42 | Brain - Cerebellar Hemisphere | 141185287 | 140685287 | 141685287 | Yes |
| ENSG00000272674.3 | PCDHB16 | rs13174119 | 4.10E-05 | -0.39 | Brain - Cerebellar Hemisphere | 141185287 | 140685287 | 141685287 | Yes |
| ENSG00000272674.3 | PCDHB16 | rs11167645 | 4.10E-05 | -0.39 | Brain - Cerebellar Hemisphere | 141185287 | 140685287 | 141685287 | Yes |
| ENSG00000272674.3 | PCDHB16 | rs199581699 | 2.00E-05 | -0.42 | Brain - Cerebellar Hemisphere | 141185287 | 140685287 | 141685287 | Yes |
| ENSG00000272674.3 | PCDHB16 | rs62384484 | 4.10E-05 | -0.39 | Brain - Cerebellar Hemisphere | 141185287 | 140685287 | 141685287 | Yes |
| ENSG00000272674.3 | PCDHB16 | rs6874218 | 4.10E-05 | -0.39 | Brain - Cerebellar Hemisphere | 141185287 | 140685287 | 141685287 | Yes |
| ENSG00000272674.3 | PCDHB16 | rs34807894 | 4.10E-05 | -0.39 | Brain - Cerebellar Hemisphere | 141185287 | 140685287 | 141685287 | Yes |
| ENSG00000272674.3 | PCDHB16 | rs62384488 | 4.10E-05 | -0.39 | Brain - Cerebellar Hemisphere | 141185287 | 140685287 | 141685287 | Yes |
| ENSG00000272674.3 | PCDHB16 | rs13182228 | 4.10E-05 | -0.39 | Brain - Cerebellar Hemisphere | 141185287 | 140685287 | 141685287 | Yes |
| ENSG00000272674.3 | PCDHB16 | rs7704223 | 4.10E-05 | -0.39 | Brain - Cerebellar Hemisphere | 141185287 | 140685287 | 141685287 | Yes |
| ENSG00000272674.3 | PCDHB16 | rs1606122 | 2.00E-05 | -0.42 | Brain - Cerebellar Hemisphere | 141185287 | 140685287 | 141685287 | Yes |
| ENSG00000272674.3 | PCDHB16 | rs1849054 | 4.10E-05 | -0.39 | Brain - Cerebellar Hemisphere | 141185287 | 140685287 | 141685287 | Yes |
| ENSG00000272674.3 | PCDHB16 | rs1858296 | 4.10E-05 | -0.39 | Brain - Cerebellar Hemisphere | 141185287 | 140685287 | 141685287 | Yes |

| | | | | | | | | | |
|---|---|---|---|---|---|---|---|---|---|
| ENSG00000272674.3 | PCDHB16 | rs11167655 | 4.10E-05 | -0.39 | Brain - Cerebellar Hemisphere | 141185287 | 140685287 | 141685287 | Yes |
| ENSG00000272674.3 | PCDHB16 | rs155804 | 4.10E-05 | -0.39 | Brain - Cerebellar Hemisphere | 141185287 | 140685287 | 141685287 | Yes |
| ENSG00000272674.3 | PCDHB16 | rs155803 | 2.70E-05 | -0.4 | Brain - Cerebellar Hemisphere | 141185287 | 140685287 | 141685287 | Yes |
| ENSG00000272674.3 | PCDHB16 | rs155801 | 4.10E-05 | -0.39 | Brain - Cerebellar Hemisphere | 141185287 | 140685287 | 141685287 | Yes |
| ENSG00000272674.3 | PCDHB16 | rs155800 | 4.10E-05 | -0.39 | Brain - Cerebellar Hemisphere | 141185287 | 140685287 | 141685287 | Yes |
| ENSG00000272674.3 | PCDHB16 | rs56003 | 4.10E-05 | -0.39 | Brain - Cerebellar Hemisphere | 141185287 | 140685287 | 141685287 | Yes |
| ENSG00000272674.3 | PCDHB16 | rs56002 | 4.10E-05 | -0.39 | Brain - Cerebellar Hemisphere | 141185287 | 140685287 | 141685287 | Yes |
| ENSG00000272674.3 | PCDHB16 | rs35005979 | 4.10E-05 | -0.39 | Brain - Cerebellar Hemisphere | 141185287 | 140685287 | 141685287 | Yes |
| ENSG00000272674.3 | PCDHB16 | rs155817 | 4.10E-05 | -0.39 | Brain - Cerebellar Hemisphere | 141185287 | 140685287 | 141685287 | Yes |
| ENSG00000272674.3 | PCDHB16 | rs155818 | 4.10E-05 | -0.39 | Brain - Cerebellar Hemisphere | 141185287 | 140685287 | 141685287 | Yes |
| ENSG00000272674.3 | PCDHB16 | rs122183103 | 4.10E-05 | -0.39 | Brain - Cerebellar Hemisphere | 141185287 | 140685287 | 141685287 | Yes |
| ENSG00000272674.3 | PCDHB16 | rs155820 | 4.10E-05 | -0.39 | Brain - Cerebellar Hemisphere | 141185287 | 140685287 | 141685287 | Yes |
| ENSG00000272674.3 | PCDHB16 | rs155821 | 1.40E-05 | -0.42 | Brain - Cerebellar Hemisphere | 141185287 | 140685287 | 141685287 | Yes |
| ENSG00000272674.3 | PCDHB16 | rs3836748 | 4.10E-05 | -0.39 | Brain - Cerebellar Hemisphere | 141185287 | 140685287 | 141685287 | Yes |
| ENSG00000272674.3 | PCDHB16 | rs140472192 | 4.10E-05 | -0.39 | Brain - Cerebellar Hemisphere | 141185287 | 140685287 | 141685287 | Yes |
| ENSG00000272674.3 | PCDHB16 | rs155823 | 4.10E-05 | -0.39 | Brain - Cerebellar Hemisphere | 141185287 | 140685287 | 141685287 | Yes |
| ENSG00000272674.3 | PCDHB16 | rs155824 | 4.10E-05 | -0.39 | Brain - Cerebellar Hemisphere | 141185287 | 140685287 | 141685287 | Yes |
| ENSG00000272674.3 | PCDHB16 | rs155825 | 4.10E-05 | -0.39 | Brain - Cerebellar Hemisphere | 141185287 | 140685287 | 141685287 | Yes |
| ENSG00000272674.3 | PCDHB16 | rs246073 | 1.70E-05 | -0.41 | Brain - Cerebellar Hemisphere | 141185287 | 140685287 | 141685287 | Yes |
| ENSG00000272674.3 | PCDHB16 | rs149003334 | 4.10E-05 | -0.39 | Brain - Cerebellar Hemisphere | 141185287 | 140685287 | 141685287 | Yes |
| ENSG00000272674.3 | PCDHB16 | rs155816 | 4.10E-05 | -0.39 | Brain - Cerebellar Hemisphere | 141185287 | 140685287 | 141685287 | Yes |
| ENSG00000272674.3 | PCDHB16 | rs155815 | 4.10E-05 | -0.39 | Brain - Cerebellar Hemisphere | 141185287 | 140685287 | 141685287 | Yes |
| ENSG00000272674.3 | PCDHB16 | rs246068 | 4.10E-05 | -0.39 | Brain - Cerebellar Hemisphere | 141185287 | 140685287 | 141685287 | Yes |
| ENSG00000272674.3 | PCDHB16 | rs155814 | 1.40E-05 | -0.42 | Brain - Cerebellar Hemisphere | 141185287 | 140685287 | 141685287 | Yes |
| ENSG00000272674.3 | PCDHB16 | rs155813 | 4.10E-05 | -0.39 | Brain - Cerebellar Hemisphere | 141185287 | 140685287 | 141685287 | Yes |
| ENSG00000272674.3 | PCDHB16 | rs269553 | 4.10E-05 | -0.39 | Brain - Cerebellar Hemisphere | 141185287 | 140685287 | 141685287 | Yes |
| ENSG00000272674.3 | PCDHB16 | rs145130356 | 2.50E-05 | -0.42 | Brain - Cerebellar Hemisphere | 141185287 | 140685287 | 141685287 | Yes |
| ENSG00000272674.3 | PCDHB16 | rs246040 | 4.10E-05 | -0.39 | Brain - Cerebellar Hemisphere | 141185287 | 140685287 | 141685287 | Yes |
| ENSG00000272674.3 | PCDHB16 | rs246039 | 2.60E-05 | -0.37 | Brain - Cerebellar Hemisphere | 141185287 | 140685287 | 141685287 | Yes |
| ENSG00000272674.3 | PCDHB16 | rs246015 | 4.10E-05 | -0.39 | Brain - Cerebellar Hemisphere | 141185287 | 140685287 | 141685287 | Yes |
| ENSG00000272674.3 | PCDHB16 | rs155799 | 2.00E-05 | -0.42 | Brain - Cerebellar Hemisphere | 141185287 | 140685287 | 141685287 | Yes |
| ENSG00000272674.3 | PCDHB16 | rs135310774 | 4.10E-05 | -0.39 | Brain - Cerebellar Hemisphere | 141185287 | 140685287 | 141685287 | Yes |

| | | | | | | | | | |
|---|---|---|---|---|---|---|---|---|---|
| ENSG00000272674.3 | PCDHB16 | rs5871756 | 1.40E-05 | -0.42 | Brain - Cerebellar Hemisphere | 141185287 | 140685287 | 141685287 | Yes |
| ENSG00000272674.3 | PCDHB16 | rs246004 | 2.00E-05 | -0.41 | Brain - Cerebellar Hemisphere | 141185287 | 140685287 | 141685287 | Yes |
| ENSG00000272674.3 | PCDHB16 | rs151988 | 1.90E-05 | -0.41 | Brain - Cerebellar Hemisphere | 141185287 | 140685287 | 141685287 | Yes |
| ENSG00000272674.3 | PCDHB16 | rs155812 | 2.00E-05 | -0.41 | Brain - Cerebellar Hemisphere | 141185287 | 140685287 | 141685287 | Yes |
| ENSG00000272674.3 | PCDHB16 | rs155811 | 7.70E-06 | -0.42 | Brain - Cerebellar Hemisphere | 141185287 | 140685287 | 141685287 | Yes |
| ENSG00000272674.3 | PCDHB16 | rs155808 | 2.30E-05 | -0.4 | Brain - Cerebellar Hemisphere | 141185287 | 140685287 | 141685287 | Yes |
| ENSG00000272674.3 | PCDHB16 | rs155807 | 4.10E-05 | -0.39 | Brain - Cerebellar Hemisphere | 141185287 | 140685287 | 141685287 | Yes |
| ENSG00000272674.3 | PCDHB16 | rs3776114 | 2.10E-07 | -0.53 | Brain - Cerebellar Hemisphere | 141185287 | 140685287 | 141685287 | Yes |
| ENSG00000272674.3 | PCDHB16 | rs4912736 | 8.30E-07 | -0.5 | Brain - Cerebellar Hemisphere | 141185287 | 140685287 | 141685287 | Yes |
| ENSG00000272674.3 | PCDHB16 | rs3756325 | 5.00E-07 | -0.51 | Brain - Cerebellar Hemisphere | 141185287 | 140685287 | 141685287 | Yes |
| ENSG00000272674.3 | PCDHB16 | rs31874 | 1.20E-05 | -0.47 | Brain - Cerebellar Hemisphere | 141185287 | 140685287 | 141685287 | Yes |
| ENSG00000272674.3 | PCDHB16 | rs3756324 | 5.20E-07 | -0.51 | Brain - Cerebellar Hemisphere | 141185287 | 140685287 | 141685287 | Yes |
| ENSG00000272674.3 | PCDHB16 | rs31872 | 1.50E-05 | -0.43 | Brain - Cerebellar Hemisphere | 141185287 | 140685287 | 141685287 | Yes |
| ENSG00000272674.3 | PCDHB16 | rs17119346 | 5.00E-07 | -0.51 | Brain - Cerebellar Hemisphere | 141185287 | 140685287 | 141685287 | Yes |
| ENSG00000272674.3 | PCDHB16 | rs17119367 | 3.10E-08 | -0.61 | Brain - Cerebellar Hemisphere | 141185287 | 140685287 | 141685287 | Yes |
| ENSG00000272674.3 | PCDHB16 | rs17119367 | 3.10E-06 | -0.46 | Brain - Cerebellum | 141185287 | 140685287 | 141685287 | Yes |
| ENSG00000272674.3 | PCDHB16 | rs11742520 | 2.50E-08 | -0.63 | Brain - Cerebellar Hemisphere | 141185287 | 140685287 | 141685287 | Yes |
| ENSG00000272674.3 | PCDHB16 | rs11742520 | 9.80E-07 | -0.5 | Brain - Cerebellum | 141185287 | 140685287 | 141685287 | Yes |
| ENSG00000272674.3 | PCDHB16 | rs31746 | 7.50E-05 | -0.4 | Brain - Cerebellar Hemisphere | 141185287 | 140685287 | 141685287 | Yes |
| ENSG00000272674.3 | PCDHB16 | rs3776105 | 3.10E-08 | -0.61 | Brain - Cerebellar Hemisphere | 141185287 | 140685287 | 141685287 | Yes |
| ENSG00000272674.3 | PCDHB16 | rs3776105 | 3.10E-06 | -0.46 | Brain - Cerebellum | 141185287 | 140685287 | 141685287 | Yes |
| ENSG00000272674.3 | PCDHB16 | rs151331262 | 1.20E-08 | -0.64 | Brain - Cerebellar Hemisphere | 141185287 | 140685287 | 141685287 | Yes |
| ENSG00000272674.3 | PCDHB16 | rs151331262 | 1.00E-06 | -0.5 | Brain - Cerebellum | 141185287 | 140685287 | 141685287 | Yes |
| ENSG00000272674.3 | PCDHB16 | rs74855082 | 1.20E-08 | -0.64 | Brain - Cerebellar Hemisphere | 141185287 | 140685287 | 141685287 | Yes |
| ENSG00000272674.3 | PCDHB16 | rs74855082 | 1.00E-06 | -0.5 | Brain - Cerebellum | 141185287 | 140685287 | 141685287 | Yes |
| ENSG00000272674.3 | PCDHB16 | rs702390 | 2.40E-06 | 0.37 | Brain - Cerebellum | 141185287 | 140685287 | 141685287 | Yes |
| ENSG00000272674.3 | PCDHB16 | rs702390 | 6.10E-06 | 0.46 | Brain - Cerebellar Hemisphere | 141185287 | 140685287 | 141685287 | Yes |
| ENSG00000272674.3 | PCDHB16 | rs6580163 | 3.10E-08 | -0.61 | Brain - Cerebellar Hemisphere | 141185287 | 140685287 | 141685287 | Yes |
| ENSG00000272674.3 | PCDHB16 | rs6580163 | 3.10E-06 | -0.46 | Brain - Cerebellum | 141185287 | 140685287 | 141685287 | Yes |
| ENSG00000272674.3 | PCDHB16 | rs62378059 | 3.10E-08 | -0.61 | Brain - Cerebellar Hemisphere | 141185287 | 140685287 | 141685287 | Yes |
| ENSG00000272674.3 | PCDHB16 | rs62378059 | 3.10E-06 | -0.46 | Brain - Cerebellum | 141185287 | 140685287 | 141685287 | Yes |
| ENSG00000272674.3 | PCDHB16 | rs31801 | 2.90E-05 | -0.33 | Brain - Cerebellum | 141185287 | 140685287 | 141685287 | Yes |

| | | | | | | | | | |
|---|---|---|---|---|---|---|---|---|---|
| ENSG00000272674.3 | PCDHB16 | rs77437352 | 3.10E-08 | -0.61 | Brain - Cerebellar Hemisphere | 141185287 | 140685287 | 141685287 | Yes |
| ENSG00000272674.3 | PCDHB16 | rs77437352 | 3.10E-06 | -0.46 | Brain - Cerebellum | 141185287 | 140685287 | 141685287 | Yes |
| ENSG00000272674.3 | PCDHB16 | rs31797 | 4.10E-05 | -0.33 | Brain - Cerebellum | 141185287 | 140685287 | 141685287 | Yes |
| ENSG00000272674.3 | PCDHB16 | rs11746036 | 2.50E-08 | -0.63 | Brain - Cerebellar Hemisphere | 141185287 | 140685287 | 141685287 | Yes |
| ENSG00000272674.3 | PCDHB16 | rs11746036 | 9.80E-07 | -0.5 | Brain - Cerebellum | 141185287 | 140685287 | 141685287 | Yes |
| ENSG00000272674.3 | PCDHB16 | rs2059127 | 3.10E-08 | -0.61 | Brain - Cerebellar Hemisphere | 141185287 | 140685287 | 141685287 | Yes |
| ENSG00000272674.3 | PCDHB16 | rs2059127 | 3.10E-06 | -0.46 | Brain - Cerebellum | 141185287 | 140685287 | 141685287 | Yes |
| ENSG00000272674.3 | PCDHB16 | rs11746543 | 3.10E-08 | -0.61 | Brain - Cerebellar Hemisphere | 141185287 | 140685287 | 141685287 | Yes |
| ENSG00000272674.3 | PCDHB16 | rs11746543 | 3.10E-06 | -0.46 | Brain - Cerebellum | 141185287 | 140685287 | 141685287 | Yes |
| ENSG00000272674.3 | PCDHB16 | rs62378862 | 2.50E-08 | -0.63 | Brain - Cerebellar Hemisphere | 141185287 | 140685287 | 141685287 | Yes |
| ENSG00000272674.3 | PCDHB16 | rs62378862 | 9.80E-07 | -0.5 | Brain - Cerebellum | 141185287 | 140685287 | 141685287 | Yes |
| ENSG00000272674.3 | PCDHB16 | rs4912598 | 5.90E-08 | -0.6 | Brain - Cerebellar Hemisphere | 141185287 | 140685287 | 141685287 | Yes |
| ENSG00000272674.3 | PCDHB16 | rs4912598 | 2.60E-06 | -0.46 | Brain - Cerebellum | 141185287 | 140685287 | 141685287 | Yes |
| ENSG00000272674.3 | PCDHB16 | rs17844376 | 1.40E-07 | -0.59 | Brain - Cerebellar Hemisphere | 141185287 | 140685287 | 141685287 | Yes |
| ENSG00000272674.3 | PCDHB16 | rs17844376 | 1.30E-06 | -0.48 | Brain - Cerebellum | 141185287 | 140685287 | 141685287 | Yes |
| ENSG00000272674.3 | PCDHB16 | rs31850 | 4.10E-05 | -0.33 | Brain - Cerebellum | 141185287 | 140685287 | 141685287 | Yes |
| ENSG00000272674.3 | PCDHB16 | rs31848 | 3.30E-06 | -0.37 | Brain - Cerebellum | 141185287 | 140685287 | 141685287 | Yes |
| ENSG00000272674.3 | PCDHB16 | rs31848 | 7.50E-06 | -0.46 | Brain - Cerebellar Hemisphere | 141185287 | 140685287 | 141685287 | Yes |
| ENSG00000272674.3 | PCDHB16 | rs7722330 | 3.10E-08 | -0.61 | Brain - Cerebellar Hemisphere | 141185287 | 140685287 | 141685287 | Yes |
| ENSG00000272674.3 | PCDHB16 | rs7722330 | 3.10E-06 | -0.46 | Brain - Cerebellum | 141185287 | 140685287 | 141685287 | Yes |
| ENSG00000272674.3 | PCDHB16 | rs10463345 | 2.50E-08 | -0.63 | Brain - Cerebellar Hemisphere | 141185287 | 140685287 | 141685287 | Yes |
| ENSG00000272674.3 | PCDHB16 | rs10463345 | 9.80E-07 | -0.5 | Brain - Cerebellum | 141185287 | 140685287 | 141685287 | Yes |
| ENSG00000272674.3 | PCDHB16 | rs6887594 | 3.10E-08 | -0.61 | Brain - Cerebellar Hemisphere | 141185287 | 140685287 | 141685287 | Yes |
| ENSG00000272674.3 | PCDHB16 | rs6887594 | 3.10E-06 | -0.46 | Brain - Cerebellum | 141185287 | 140685287 | 141685287 | Yes |
| ENSG00000272674.3 | PCDHB16 | rs3776100 | 2.50E-08 | -0.63 | Brain - Cerebellar Hemisphere | 141185287 | 140685287 | 141685287 | Yes |
| ENSG00000272674.3 | PCDHB16 | rs3776100 | 9.80E-07 | -0.5 | Brain - Cerebellum | 141185287 | 140685287 | 141685287 | Yes |
| ENSG00000272674.3 | PCDHB16 | rs3088331 | 2.50E-08 | -0.63 | Brain - Cerebellar Hemisphere | 141185287 | 140685287 | 141685287 | Yes |
| ENSG00000272674.3 | PCDHB16 | rs3088331 | 9.80E-07 | -0.5 | Brain - Cerebellum | 141185287 | 140685287 | 141685287 | Yes |
| ENSG00000272674.3 | PCDHB16 | rs55902608 | 1.40E-08 | -0.64 | Brain - Cerebellar Hemisphere | 141185287 | 140685287 | 141685287 | Yes |
| ENSG00000272674.3 | PCDHB16 | rs55902608 | 1.20E-06 | -0.49 | Brain - Cerebellum | 141185287 | 140685287 | 141685287 | Yes |
| ENSG00000272674.3 | PCDHB16 | rs2338528 | 6.00E-06 | -0.39 | Brain - Cerebellum | 141185287 | 140685287 | 141685287 | Yes |
| ENSG00000272674.3 | PCDHB16 | rs2338528 | 7.50E-06 | -0.44 | Brain - Cerebellar Hemisphere | 141185287 | 140685287 | 141685287 | Yes |

| | | | | | | | | | |
|---|---|---|---|---|---|---|---|---|---|
| ENSG00000272674.3 | PCDHB16 | rs6864827 | 6.80E-05 | -0.32 | Brain - Cerebellum | 141185287 | 140685287 | 141685287 | Yes |
| ENSG00000272674.3 | PCDHB16 | rs246723 | 1.30E-06 | -0.48 | Brain - Cerebellar Hemisphere | 141185287 | 140685287 | 141685287 | Yes |
| ENSG00000272674.3 | PCDHB16 | rs246723 | 2.60E-06 | -0.4 | Brain - Cerebellum | 141185287 | 140685287 | 141685287 | Yes |
| ENSG00000272674.3 | PCDHB16 | rs7730410 | 6.00E-06 | -0.39 | Brain - Cerebellum | 141185287 | 140685287 | 141685287 | Yes |
| ENSG00000272674.3 | PCDHB16 | rs7730410 | 7.50E-06 | -0.44 | Brain - Cerebellar Hemisphere | 141185287 | 140685287 | 141685287 | Yes |
| ENSG00000272674.3 | PCDHB16 | rs246722 | 1.20E-08 | -0.52 | Brain - Cerebellar Hemisphere | 141185287 | 140685287 | 141685287 | Yes |
| ENSG00000272674.3 | PCDHB16 | rs246722 | 1.90E-08 | -0.47 | Brain - Cerebellum | 141185287 | 140685287 | 141685287 | Yes |
| ENSG00000272674.3 | PCDHB16 | rs62378868 | 2.70E-11 | -0.77 | Brain - Cerebellar Hemisphere | 141185287 | 140685287 | 141685287 | Yes |
| ENSG00000272674.3 | PCDHB16 | rs62378868 | 8.60E-10 | -0.65 | Brain - Cerebellum | 141185287 | 140685287 | 141685287 | Yes |
| ENSG00000272674.3 | PCDHB16 | rs246718 | 7.80E-08 | -0.49 | Brain - Cerebellar Hemisphere | 141185287 | 140685287 | 141685287 | Yes |
| ENSG00000272674.3 | PCDHB16 | rs246718 | 7.90E-08 | -0.44 | Brain - Cerebellum | 141185287 | 140685287 | 141685287 | Yes |
| ENSG00000272674.3 | PCDHB16 | rs246717 | 4.70E-08 | -0.44 | Brain - Cerebellum | 141185287 | 140685287 | 141685287 | Yes |
| ENSG00000272674.3 | PCDHB16 | rs246717 | 3.20E-06 | -0.46 | Brain - Cerebellar Hemisphere | 141185287 | 140685287 | 141685287 | Yes |
| ENSG00000272674.3 | PCDHB16 | rs246716 | 7.80E-08 | -0.49 | Brain - Cerebellar Hemisphere | 141185287 | 140685287 | 141685287 | Yes |
| ENSG00000272674.3 | PCDHB16 | rs246716 | 7.90E-08 | -0.44 | Brain - Cerebellum | 141185287 | 140685287 | 141685287 | Yes |
| ENSG00000272674.3 | PCDHB16 | rs12654066 | 2.70E-11 | -0.77 | Brain - Cerebellar Hemisphere | 141185287 | 140685287 | 141685287 | Yes |
| ENSG00000272674.3 | PCDHB16 | rs12654066 | 3.10E-09 | -0.63 | Brain - Cerebellum | 141185287 | 140685287 | 141685287 | Yes |
| ENSG00000272674.3 | PCDHB16 | rs35397499 | 1.20E-08 | -0.52 | Brain - Cerebellar Hemisphere | 141185287 | 140685287 | 141685287 | Yes |
| ENSG00000272674.3 | PCDHB16 | rs35397499 | 1.90E-08 | -0.47 | Brain - Cerebellum | 141185287 | 140685287 | 141685287 | Yes |
| ENSG00000272674.3 | PCDHB16 | rs246714 | 1.20E-08 | -0.52 | Brain - Cerebellar Hemisphere | 141185287 | 140685287 | 141685287 | Yes |
| ENSG00000272674.3 | PCDHB16 | rs246714 | 1.90E-08 | -0.47 | Brain - Cerebellum | 141185287 | 140685287 | 141685287 | Yes |
| ENSG00000272674.3 | PCDHB16 | rs62378869 | 2.70E-11 | -0.77 | Brain - Cerebellar Hemisphere | 141185287 | 140685287 | 141685287 | Yes |
| ENSG00000272674.3 | PCDHB16 | rs62378869 | 8.60E-10 | -0.65 | Brain - Cerebellum | 141185287 | 140685287 | 141685287 | Yes |
| ENSG00000272674.3 | PCDHB16 | rs55634432 | 2.70E-11 | -0.77 | Brain - Cerebellar Hemisphere | 141185287 | 140685287 | 141685287 | Yes |
| ENSG00000272674.3 | PCDHB16 | rs55634432 | 8.60E-10 | -0.65 | Brain - Cerebellum | 141185287 | 140685287 | 141685287 | Yes |
| ENSG00000272674.3 | PCDHB16 | rs35627549 | 2.70E-11 | -0.77 | Brain - Cerebellar Hemisphere | 141185287 | 140685287 | 141685287 | Yes |
| ENSG00000272674.3 | PCDHB16 | rs35627549 | 8.60E-10 | -0.65 | Brain - Cerebellum | 141185287 | 140685287 | 141685287 | Yes |
| ENSG00000272674.3 | PCDHB16 | rs34762107 | 2.70E-11 | -0.77 | Brain - Cerebellar Hemisphere | 141185287 | 140685287 | 141685287 | Yes |
| ENSG00000272674.3 | PCDHB16 | rs34762107 | 8.60E-10 | -0.65 | Brain - Cerebellum | 141185287 | 140685287 | 141685287 | Yes |
| ENSG00000272674.3 | PCDHB16 | rs62378893 | 2.70E-11 | -0.77 | Brain - Cerebellar Hemisphere | 141185287 | 140685287 | 141685287 | Yes |
| ENSG00000272674.3 | PCDHB16 | rs62378893 | 8.60E-10 | -0.65 | Brain - Cerebellum | 141185287 | 140685287 | 141685287 | Yes |
| ENSG00000272674.3 | PCDHB16 | rs246711 | 1.20E-08 | -0.52 | Brain - Cerebellar Hemisphere | 141185287 | 140685287 | 141685287 | Yes |

| | | | | | | | | | |
|---|---|---|---|---|---|---|---|---|---|
| ENSG00000272674.3 | PCDHB16 | rs246711 | 1.90E-08 | -0.47 | Brain - Cerebellum | 141185287 | 140685287 | 141685287 | Yes |
| ENSG00000272674.3 | PCDHB16 | rs17629011 | 6.80E-05 | -0.91 | Brain - Cerebellar Hemisphere | 141185287 | 140685287 | 141685287 | Yes |
| ENSG00000272674.3 | PCDHB16 | rs246710 | 1.20E-08 | -0.52 | Brain - Cerebellar Hemisphere | 141185287 | 140685287 | 141685287 | Yes |
| ENSG00000272674.3 | PCDHB16 | rs246710 | 1.90E-08 | -0.47 | Brain - Cerebellum | 141185287 | 140685287 | 141685287 | Yes |
| ENSG00000272674.3 | PCDHB16 | rs246709 | 1.20E-08 | -0.52 | Brain - Cerebellar Hemisphere | 141185287 | 140685287 | 141685287 | Yes |
| ENSG00000272674.3 | PCDHB16 | rs246709 | 1.90E-08 | -0.47 | Brain - Cerebellum | 141185287 | 140685287 | 141685287 | Yes |
| ENSG00000272674.3 | PCDHB16 | rs12653888 | 2.20E-11 | -0.77 | Brain - Cerebellar Hemisphere | 141185287 | 140685287 | 141685287 | Yes |
| ENSG00000272674.3 | PCDHB16 | rs12653888 | 7.60E-10 | -0.66 | Brain - Cerebellum | 141185287 | 140685287 | 141685287 | Yes |
| ENSG00000272674.3 | PCDHB16 | rs4151693 | 2.20E-11 | -0.77 | Brain - Cerebellar Hemisphere | 141185287 | 140685287 | 141685287 | Yes |
| ENSG00000272674.3 | PCDHB16 | rs4151693 | 7.60E-10 | -0.66 | Brain - Cerebellum | 141185287 | 140685287 | 141685287 | Yes |
| ENSG00000272674.3 | PCDHB16 | rs4151694 | 2.20E-11 | -0.77 | Brain - Cerebellar Hemisphere | 141185287 | 140685287 | 141685287 | Yes |
| ENSG00000272674.3 | PCDHB16 | rs4151694 | 7.60E-10 | -0.66 | Brain - Cerebellum | 141185287 | 140685287 | 141685287 | Yes |
| ENSG00000272674.3 | PCDHB16 | rs3776096 | 2.20E-11 | -0.77 | Brain - Cerebellar Hemisphere | 141185287 | 140685287 | 141685287 | Yes |
| ENSG00000272674.3 | PCDHB16 | rs3776096 | 7.60E-10 | -0.66 | Brain - Cerebellum | 141185287 | 140685287 | 141685287 | Yes |
| ENSG00000272674.3 | PCDHB16 | rs17844439 | 2.20E-11 | -0.77 | Brain - Cerebellar Hemisphere | 141185287 | 140685287 | 141685287 | Yes |
| ENSG00000272674.3 | PCDHB16 | rs17844439 | 7.60E-10 | -0.66 | Brain - Cerebellum | 141185287 | 140685287 | 141685287 | Yes |
| ENSG00000272674.3 | PCDHB16 | rs246703 | 4.20E-08 | -0.44 | Brain - Cerebellum | 141185287 | 140685287 | 141685287 | Yes |
| ENSG00000272674.3 | PCDHB16 | rs246703 | 6.90E-06 | -0.45 | Brain - Cerebellar Hemisphere | 141185287 | 140685287 | 141685287 | Yes |
| ENSG00000272674.3 | PCDHB16 | rs12658810 | 2.20E-11 | -0.77 | Brain - Cerebellar Hemisphere | 141185287 | 140685287 | 141685287 | Yes |
| ENSG00000272674.3 | PCDHB16 | rs12658810 | 7.60E-10 | -0.66 | Brain - Cerebellum | 141185287 | 140685287 | 141685287 | Yes |
| ENSG00000272674.3 | PCDHB16 | rs246702 | 1.20E-08 | -0.52 | Brain - Cerebellar Hemisphere | 141185287 | 140685287 | 141685287 | Yes |
| ENSG00000272674.3 | PCDHB16 | rs246702 | 1.90E-08 | -0.47 | Brain - Cerebellum | 141185287 | 140685287 | 141685287 | Yes |
| ENSG00000272674.3 | PCDHB16 | rs17096929 | 2.20E-11 | -0.77 | Brain - Cerebellar Hemisphere | 141185287 | 140685287 | 141685287 | Yes |
| ENSG00000272674.3 | PCDHB16 | rs17096929 | 7.60E-10 | -0.66 | Brain - Cerebellum | 141185287 | 140685287 | 141685287 | Yes |
| ENSG00000272674.3 | PCDHB16 | rs246701 | 1.20E-08 | -0.52 | Brain - Cerebellar Hemisphere | 141185287 | 140685287 | 141685287 | Yes |
| ENSG00000272674.3 | PCDHB16 | rs246701 | 1.90E-08 | -0.47 | Brain - Cerebellum | 141185287 | 140685287 | 141685287 | Yes |
| ENSG00000272674.3 | PCDHB16 | rs246700 | 1.20E-08 | -0.52 | Brain - Cerebellar Hemisphere | 141185287 | 140685287 | 141685287 | Yes |
| ENSG00000272674.3 | PCDHB16 | rs246700 | 1.90E-08 | -0.47 | Brain - Cerebellum | 141185287 | 140685287 | 141685287 | Yes |
| ENSG00000272674.3 | PCDHB16 | rs62378894 | 2.20E-11 | -0.77 | Brain - Cerebellar Hemisphere | 141185287 | 140685287 | 141685287 | Yes |
| ENSG00000272674.3 | PCDHB16 | rs62378894 | 7.60E-10 | -0.66 | Brain - Cerebellum | 141185287 | 140685287 | 141685287 | Yes |
| ENSG00000272674.3 | PCDHB16 | rs246694 | 1.30E-09 | -0.54 | Brain - Cerebellar Hemisphere | 141185287 | 140685287 | 141685287 | Yes |
| ENSG00000272674.3 | PCDHB16 | rs246694 | 1.40E-08 | -0.46 | Brain - Cerebellum | 141185287 | 140685287 | 141685287 | Yes |

| | | | | | | | | | |
|---|---|---|---|---|---|---|---|---|---|
| ENSG00000272674.3 | PCDHB16 | rs33978132 | 2.80E-08 | -0.44 | Brain - Cerebellum | 141185287 | 140685287 | 141685287 | Yes |
| ENSG00000272674.3 | PCDHB16 | rs33978132 | 1.10E-06 | -0.47 | Brain - Cerebellar Hemisphere | 141185287 | 140685287 | 141685287 | Yes |
| ENSG00000272674.3 | PCDHB16 | rs246693 | 8.60E-11 | -0.49 | Brain - Cerebellum | 141185287 | 140685287 | 141685287 | Yes |
| ENSG00000272674.3 | PCDHB16 | rs246693 | 9.40E-07 | -0.5 | Brain - Cerebellar Hemisphere | 141185287 | 140685287 | 141685287 | Yes |
| ENSG00000272674.3 | PCDHB16 | rs10701498 | 1.30E-09 | -0.54 | Brain - Cerebellar Hemisphere | 141185287 | 140685287 | 141685287 | Yes |
| ENSG00000272674.3 | PCDHB16 | rs10701498 | 1.40E-08 | -0.46 | Brain - Cerebellum | 141185287 | 140685287 | 141685287 | Yes |
| ENSG00000272674.3 | PCDHB16 | rs11748310 | 6.30E-13 | -0.8 | Brain - Cerebellar Hemisphere | 141185287 | 140685287 | 141685287 | Yes |
| ENSG00000272674.3 | PCDHB16 | rs11748310 | 3.10E-10 | -0.66 | Brain - Cerebellum | 141185287 | 140685287 | 141685287 | Yes |
| ENSG00000272674.3 | PCDHB16 | rs11270142 | 1.30E-09 | -0.54 | Brain - Cerebellar Hemisphere | 141185287 | 140685287 | 141685287 | Yes |
| ENSG00000272674.3 | PCDHB16 | rs11270142 | 1.40E-08 | -0.46 | Brain - Cerebellum | 141185287 | 140685287 | 141685287 | Yes |
| ENSG00000272674.3 | PCDHB16 | rs4912600 | 6.30E-13 | -0.8 | Brain - Cerebellar Hemisphere | 141185287 | 140685287 | 141685287 | Yes |
| ENSG00000272674.3 | PCDHB16 | rs4912600 | 3.10E-10 | -0.66 | Brain - Cerebellum | 141185287 | 140685287 | 141685287 | Yes |
| ENSG00000272674.3 | PCDHB16 | rs56233997 | 6.30E-13 | -0.8 | Brain - Cerebellar Hemisphere | 141185287 | 140685287 | 141685287 | Yes |
| ENSG00000272674.3 | PCDHB16 | rs56233997 | 3.10E-10 | -0.66 | Brain - Cerebellum | 141185287 | 140685287 | 141685287 | Yes |
| ENSG00000272674.3 | PCDHB16 | rs145354 | 1.30E-09 | -0.54 | Brain - Cerebellar Hemisphere | 141185287 | 140685287 | 141685287 | Yes |
| ENSG00000272674.3 | PCDHB16 | rs145354 | 1.40E-08 | -0.46 | Brain - Cerebellum | 141185287 | 140685287 | 141685287 | Yes |
| ENSG00000272674.3 | PCDHB16 | rs62378897 | 6.30E-13 | -0.8 | Brain - Cerebellar Hemisphere | 141185287 | 140685287 | 141685287 | Yes |
| ENSG00000272674.3 | PCDHB16 | rs62378897 | 3.10E-10 | -0.66 | Brain - Cerebellum | 141185287 | 140685287 | 141685287 | Yes |
| ENSG00000272674.3 | PCDHB16 | rs246690 | 2.80E-08 | -0.44 | Brain - Cerebellum | 141185287 | 140685287 | 141685287 | Yes |
| ENSG00000272674.3 | PCDHB16 | rs246690 | 1.10E-06 | -0.47 | Brain - Cerebellar Hemisphere | 141185287 | 140685287 | 141685287 | Yes |
| ENSG00000272674.3 | PCDHB16 | rs11321809 | 6.30E-13 | -0.8 | Brain - Cerebellar Hemisphere | 141185287 | 140685287 | 141685287 | Yes |
| ENSG00000272674.3 | PCDHB16 | rs11321809 | 3.10E-10 | -0.66 | Brain - Cerebellum | 141185287 | 140685287 | 141685287 | Yes |
| ENSG00000272674.3 | PCDHB16 | rs246689 | 1.30E-09 | -0.54 | Brain - Cerebellar Hemisphere | 141185287 | 140685287 | 141685287 | Yes |
| ENSG00000272674.3 | PCDHB16 | rs246689 | 1.40E-08 | -0.46 | Brain - Cerebellum | 141185287 | 140685287 | 141685287 | Yes |
| ENSG00000272674.3 | PCDHB16 | rs246688 | 2.80E-08 | -0.44 | Brain - Cerebellum | 141185287 | 140685287 | 141685287 | Yes |
| ENSG00000272674.3 | PCDHB16 | rs246688 | 1.10E-06 | -0.47 | Brain - Cerebellar Hemisphere | 141185287 | 140685287 | 141685287 | Yes |
| ENSG00000272674.3 | PCDHB16 | rs246686 | 1.30E-09 | -0.54 | Brain - Cerebellar Hemisphere | 141185287 | 140685287 | 141685287 | Yes |
| ENSG00000272674.3 | PCDHB16 | rs246686 | 1.40E-08 | -0.46 | Brain - Cerebellum | 141185287 | 140685287 | 141685287 | Yes |
| ENSG00000272674.3 | PCDHB16 | rs55805572 | 6.30E-13 | -0.8 | Brain - Cerebellar Hemisphere | 141185287 | 140685287 | 141685287 | Yes |
| ENSG00000272674.3 | PCDHB16 | rs55805572 | 3.10E-10 | -0.66 | Brain - Cerebellum | 141185287 | 140685287 | 141685287 | Yes |
| ENSG00000272674.3 | PCDHB16 | rs246683 | 1.30E-09 | -0.54 | Brain - Cerebellar Hemisphere | 141185287 | 140685287 | 141685287 | Yes |
| ENSG00000272674.3 | PCDHB16 | rs246683 | 1.40E-08 | -0.46 | Brain - Cerebellum | 141185287 | 140685287 | 141685287 | Yes |

| | | | | | | | | | |
|---|---|---|---|---|---|---|---|---|---|
| ENSG00000272674.3 | PCDHB16 | rs246682 | 1.30E-09 | -0.54 | Brain - Cerebellar Hemisphere | 141185287 | 140685287 | 141685287 | Yes |
| ENSG00000272674.3 | PCDHB16 | rs246682 | 1.40E-08 | -0.46 | Brain - Cerebellum | 141185287 | 140685287 | 141685287 | Yes |
| ENSG00000272674.3 | PCDHB16 | rs246681 | 1.30E-09 | -0.54 | Brain - Cerebellar Hemisphere | 141185287 | 140685287 | 141685287 | Yes |
| ENSG00000272674.3 | PCDHB16 | rs246681 | 1.40E-08 | -0.46 | Brain - Cerebellum | 141185287 | 140685287 | 141685287 | Yes |
| ENSG00000272674.3 | PCDHB16 | rs246680 | 1.30E-09 | -0.54 | Brain - Cerebellar Hemisphere | 141185287 | 140685287 | 141685287 | Yes |
| ENSG00000272674.3 | PCDHB16 | rs246680 | 1.40E-08 | -0.46 | Brain - Cerebellum | 141185287 | 140685287 | 141685287 | Yes |
| ENSG00000272674.3 | PCDHB16 | rs6884127 | 1.30E-09 | -0.54 | Brain - Cerebellar Hemisphere | 141185287 | 140685287 | 141685287 | Yes |
| ENSG00000272674.3 | PCDHB16 | rs6884127 | 1.40E-08 | -0.46 | Brain - Cerebellum | 141185287 | 140685287 | 141685287 | Yes |
| ENSG00000272674.3 | PCDHB16 | rs1528587 | 1.40E-12 | -0.79 | Brain - Cerebellar Hemisphere | 141185287 | 140685287 | 141685287 | Yes |
| ENSG00000272674.3 | PCDHB16 | rs1528587 | 1.30E-10 | -0.67 | Brain - Cerebellum | 141185287 | 140685287 | 141685287 | Yes |
| ENSG00000272674.3 | PCDHB16 | rs2910313 | 2.80E-08 | -0.44 | Brain - Cerebellum | 141185287 | 140685287 | 141685287 | Yes |
| ENSG00000272674.3 | PCDHB16 | rs2910313 | 1.10E-06 | -0.47 | Brain - Cerebellar Hemisphere | 141185287 | 140685287 | 141685287 | Yes |
| ENSG00000272674.3 | PCDHB16 | rs2907330 | 2.80E-08 | -0.44 | Brain - Cerebellum | 141185287 | 140685287 | 141685287 | Yes |
| ENSG00000272674.3 | PCDHB16 | rs2907330 | 1.10E-06 | -0.47 | Brain - Cerebellar Hemisphere | 141185287 | 140685287 | 141685287 | Yes |
| ENSG00000272674.3 | PCDHB16 | rs2907329 | 2.80E-08 | -0.44 | Brain - Cerebellum | 141185287 | 140685287 | 141685287 | Yes |
| ENSG00000272674.3 | PCDHB16 | rs2907329 | 1.10E-06 | -0.47 | Brain - Cerebellar Hemisphere | 141185287 | 140685287 | 141685287 | Yes |
| ENSG00000272674.3 | PCDHB16 | rs4912601 | 5.40E-09 | -0.51 | Brain - Cerebellar Hemisphere | 141185287 | 140685287 | 141685287 | Yes |
| ENSG00000272674.3 | PCDHB16 | rs4912601 | 1.00E-07 | -0.43 | Brain - Cerebellum | 141185287 | 140685287 | 141685287 | Yes |
| ENSG00000272674.3 | PCDHB16 | rs1811237 | 1.10E-09 | -0.63 | Brain - Cerebellar Hemisphere | 141185287 | 140685287 | 141685287 | Yes |
| ENSG00000272674.3 | PCDHB16 | rs1811237 | 5.10E-08 | -0.54 | Brain - Cerebellum | 141185287 | 140685287 | 141685287 | Yes |
| ENSG00000272674.3 | PCDHB16 | rs11741863 | 1.40E-12 | -0.79 | Brain - Cerebellar Hemisphere | 141185287 | 140685287 | 141685287 | Yes |
| ENSG00000272674.3 | PCDHB16 | rs11741863 | 1.30E-10 | -0.67 | Brain - Cerebellum | 141185287 | 140685287 | 141685287 | Yes |
| ENSG00000272674.3 | PCDHB16 | rs6892507 | 1.30E-09 | -0.54 | Brain - Cerebellar Hemisphere | 141185287 | 140685287 | 141685287 | Yes |
| ENSG00000272674.3 | PCDHB16 | rs6892507 | 1.40E-08 | -0.46 | Brain - Cerebellum | 141185287 | 140685287 | 141685287 | Yes |
| ENSG00000272674.3 | PCDHB16 | rs2910315 | 8.60E-13 | -0.78 | Brain - Cerebellar Hemisphere | 141185287 | 140685287 | 141685287 | Yes |
| ENSG00000272674.3 | PCDHB16 | rs2910315 | 2.50E-10 | -0.64 | Brain - Cerebellum | 141185287 | 140685287 | 141685287 | Yes |
| ENSG00000272674.3 | PCDHB16 | rs17844498 | 1.30E-09 | -0.54 | Brain - Cerebellar Hemisphere | 141185287 | 140685287 | 141685287 | Yes |
| ENSG00000272674.3 | PCDHB16 | rs17844498 | 1.40E-08 | -0.46 | Brain - Cerebellum | 141185287 | 140685287 | 141685287 | Yes |
| ENSG00000272674.3 | PCDHB16 | rs2740582 | 1.30E-09 | -0.54 | Brain - Cerebellar Hemisphere | 141185287 | 140685287 | 141685287 | Yes |
| ENSG00000272674.3 | PCDHB16 | rs2740582 | 1.40E-08 | -0.46 | Brain - Cerebellum | 141185287 | 140685287 | 141685287 | Yes |
| ENSG00000272674.3 | PCDHB16 | rs2740583 | 2.50E-11 | -0.53 | Brain - Cerebellum | 141185287 | 140685287 | 141685287 | Yes |
| ENSG00000272674.3 | PCDHB16 | rs2740583 | 3.10E-07 | -0.53 | Brain - Cerebellar Hemisphere | 141185287 | 140685287 | 141685287 | Yes |

| | | | | | | | | | |
|---|---|---|---|---|---|---|---|---|---|
| ENSG00000272674.3 | PCDHB16 | rs2697540 | 1.30E-09 | -0.54 | Brain - Cerebellar Hemisphere | 141185287 | 140685287 | 141685287 | Yes |
| ENSG00000272674.3 | PCDHB16 | rs2697540 | 1.40E-08 | -0.46 | Brain - Cerebellum | 141185287 | 140685287 | 141685287 | Yes |
| ENSG00000272674.3 | PCDHB16 | rs7730978 | 1.10E-09 | -0.63 | Brain - Cerebellar Hemisphere | 141185287 | 140685287 | 141685287 | Yes |
| ENSG00000272674.3 | PCDHB16 | rs7730978 | 5.10E-08 | -0.54 | Brain - Cerebellum | 141185287 | 140685287 | 141685287 | Yes |
| ENSG00000272674.3 | PCDHB16 | rs2740584 | 1.10E-09 | -0.63 | Brain - Cerebellar Hemisphere | 141185287 | 140685287 | 141685287 | Yes |
| ENSG00000272674.3 | PCDHB16 | rs2740584 | 5.10E-08 | -0.54 | Brain - Cerebellum | 141185287 | 140685287 | 141685287 | Yes |
| ENSG00000272674.3 | PCDHB16 | rs2204704 | 4.70E-08 | -0.43 | Brain - Cerebellum | 141185287 | 140685287 | 141685287 | Yes |
| ENSG00000272674.3 | PCDHB16 | rs2204704 | 5.20E-07 | -0.49 | Brain - Cerebellar Hemisphere | 141185287 | 140685287 | 141685287 | Yes |
| ENSG00000272674.3 | PCDHB16 | rs2697534 | 6.30E-08 | -0.43 | Brain - Cerebellum | 141185287 | 140685287 | 141685287 | Yes |
| ENSG00000272674.3 | PCDHB16 | rs2697534 | 3.70E-07 | -0.49 | Brain - Cerebellar Hemisphere | 141185287 | 140685287 | 141685287 | Yes |
| ENSG00000272674.3 | PCDHB16 | rs17844646 | 9.80E-13 | -0.8 | Brain - Cerebellar Hemisphere | 141185287 | 140685287 | 141685287 | Yes |
| ENSG00000272674.3 | PCDHB16 | rs17844646 | 2.30E-10 | -0.67 | Brain - Cerebellum | 141185287 | 140685287 | 141685287 | Yes |
| ENSG00000272674.3 | PCDHB16 | rs17844664 | 6.30E-13 | -0.8 | Brain - Cerebellar Hemisphere | 141185287 | 140685287 | 141685287 | Yes |
| ENSG00000272674.3 | PCDHB16 | rs17844664 | 3.10E-10 | -0.66 | Brain - Cerebellum | 141185287 | 140685287 | 141685287 | Yes |
| ENSG00000272674.3 | PCDHB16 | rs13176519 | 1.40E-12 | -0.79 | Brain - Cerebellar Hemisphere | 141185287 | 140685287 | 141685287 | Yes |
| ENSG00000272674.3 | PCDHB16 | rs13176519 | 1.30E-10 | -0.67 | Brain - Cerebellum | 141185287 | 140685287 | 141685287 | Yes |
| ENSG00000272674.3 | PCDHB16 | rs17844666 | 1.40E-12 | -0.79 | Brain - Cerebellar Hemisphere | 141185287 | 140685287 | 141685287 | Yes |
| ENSG00000272674.3 | PCDHB16 | rs17844666 | 1.30E-10 | -0.67 | Brain - Cerebellum | 141185287 | 140685287 | 141685287 | Yes |
| ENSG00000272674.3 | PCDHB16 | rs2980409 | 8.90E-12 | -0.53 | Brain - Cerebellum | 141185287 | 140685287 | 141685287 | Yes |
| ENSG00000272674.3 | PCDHB16 | rs2980409 | 1.50E-07 | -0.55 | Brain - Cerebellar Hemisphere | 141185287 | 140685287 | 141685287 | Yes |
| ENSG00000272674.3 | PCDHB16 | rs2950846 | 6.30E-08 | -0.43 | Brain - Cerebellum | 141185287 | 140685287 | 141685287 | Yes |
| ENSG00000272674.3 | PCDHB16 | rs2950846 | 3.70E-07 | -0.49 | Brain - Cerebellar Hemisphere | 141185287 | 140685287 | 141685287 | Yes |
| ENSG00000272674.3 | PCDHB16 | rs6580182 | 6.30E-08 | -0.43 | Brain - Cerebellum | 141185287 | 140685287 | 141685287 | Yes |
| ENSG00000272674.3 | PCDHB16 | rs6580182 | 3.70E-07 | -0.49 | Brain - Cerebellar Hemisphere | 141185287 | 140685287 | 141685287 | Yes |
| ENSG00000272674.3 | PCDHB16 | rs59617728 | 1.40E-12 | -0.79 | Brain - Cerebellar Hemisphere | 141185287 | 140685287 | 141685287 | Yes |
| ENSG00000272674.3 | PCDHB16 | rs59617728 | 1.30E-10 | -0.67 | Brain - Cerebellum | 141185287 | 140685287 | 141685287 | Yes |
| ENSG00000272674.3 | PCDHB16 | rs2740588 | 1.90E-12 | -0.79 | Brain - Cerebellar Hemisphere | 141185287 | 140685287 | 141685287 | Yes |
| ENSG00000272674.3 | PCDHB16 | rs2740588 | 3.10E-10 | -0.66 | Brain - Cerebellum | 141185287 | 140685287 | 141685287 | Yes |
| ENSG00000272674.3 | PCDHB16 | rs17844512 | 4.20E-12 | -0.77 | Brain - Cerebellar Hemisphere | 141185287 | 140685287 | 141685287 | Yes |
| ENSG00000272674.3 | PCDHB16 | rs17844512 | 1.30E-10 | -0.67 | Brain - Cerebellum | 141185287 | 140685287 | 141685287 | Yes |
| ENSG00000272674.3 | PCDHB16 | rs11167743 | 6.30E-08 | -0.43 | Brain - Cerebellum | 141185287 | 140685287 | 141685287 | Yes |
| ENSG00000272674.3 | PCDHB16 | rs11167743 | 3.70E-07 | -0.49 | Brain - Cerebellar Hemisphere | 141185287 | 140685287 | 141685287 | Yes |

| | | | | | | | | | |
|---|---|---|---|---|---|---|---|---|---|
| ENSG00000272674.3 | PCDHB16 | rs2697531 | 4.20E-12 | -0.77 | Brain - Cerebellar Hemisphere | 141185287 | 140685287 | 141685287 | Yes |
| ENSG00000272674.3 | PCDHB16 | rs2697531 | 1.30E-10 | -0.67 | Brain - Cerebellum | 141185287 | 140685287 | 141685287 | Yes |
| ENSG00000272674.3 | PCDHB16 | rs3733691 | 1.90E-12 | -0.79 | Brain - Cerebellar Hemisphere | 141185287 | 140685287 | 141685287 | Yes |
| ENSG00000272674.3 | PCDHB16 | rs3733691 | 3.10E-10 | -0.66 | Brain - Cerebellum | 141185287 | 140685287 | 141685287 | Yes |
| ENSG00000272674.3 | PCDHB16 | rs3733690 | 5.50E-10 | -0.56 | Brain - Cerebellar Hemisphere | 141185287 | 140685287 | 141685287 | Yes |
| ENSG00000272674.3 | PCDHB16 | rs3733690 | 4.00E-08 | -0.45 | Brain - Cerebellum | 141185287 | 140685287 | 141685287 | Yes |
| ENSG00000272674.3 | PCDHB16 | rs2907326 | 6.30E-08 | -0.43 | Brain - Cerebellum | 141185287 | 140685287 | 141685287 | Yes |
| ENSG00000272674.3 | PCDHB16 | rs2907326 | 3.70E-07 | -0.49 | Brain - Cerebellar Hemisphere | 141185287 | 140685287 | 141685287 | Yes |
| ENSG00000272674.3 | PCDHB16 | rs2910324 | 1.40E-07 | -0.42 | Brain - Cerebellum | 141185287 | 140685287 | 141685287 | Yes |
| ENSG00000272674.3 | PCDHB16 | rs2910324 | 2.60E-06 | -0.45 | Brain - Cerebellar Hemisphere | 141185287 | 140685287 | 141685287 | Yes |
| ENSG00000272674.3 | PCDHB16 | rs7735365 | 5.50E-10 | -0.56 | Brain - Cerebellar Hemisphere | 141185287 | 140685287 | 141685287 | Yes |
| ENSG00000272674.3 | PCDHB16 | rs7735365 | 4.00E-08 | -0.45 | Brain - Cerebellum | 141185287 | 140685287 | 141685287 | Yes |
| ENSG00000272674.3 | PCDHB16 | rs4912740 | 5.50E-10 | -0.56 | Brain - Cerebellar Hemisphere | 141185287 | 140685287 | 141685287 | Yes |
| ENSG00000272674.3 | PCDHB16 | rs4912740 | 4.00E-08 | -0.45 | Brain - Cerebellum | 141185287 | 140685287 | 141685287 | Yes |
| ENSG00000272674.3 | PCDHB16 | rs12654953 | 1.30E-11 | -0.62 | Brain - Cerebellar Hemisphere | 141185287 | 140685287 | 141685287 | Yes |
| ENSG00000272674.3 | PCDHB16 | rs12654953 | 4.40E-09 | -0.47 | Brain - Cerebellum | 141185287 | 140685287 | 141685287 | Yes |
| ENSG00000272674.3 | PCDHB16 | rs12657056 | 5.50E-10 | -0.56 | Brain - Cerebellar Hemisphere | 141185287 | 140685287 | 141685287 | Yes |
| ENSG00000272674.3 | PCDHB16 | rs12657056 | 4.00E-08 | -0.45 | Brain - Cerebellum | 141185287 | 140685287 | 141685287 | Yes |
| ENSG00000272674.3 | PCDHB16 | rs4912741 | 5.50E-10 | -0.56 | Brain - Cerebellar Hemisphere | 141185287 | 140685287 | 141685287 | Yes |
| ENSG00000272674.3 | PCDHB16 | rs4912741 | 4.00E-08 | -0.45 | Brain - Cerebellum | 141185287 | 140685287 | 141685287 | Yes |
| ENSG00000272674.3 | PCDHB16 | rs4912742 | 5.50E-10 | -0.56 | Brain - Cerebellar Hemisphere | 141185287 | 140685287 | 141685287 | Yes |
| ENSG00000272674.3 | PCDHB16 | rs4912742 | 4.00E-08 | -0.45 | Brain - Cerebellum | 141185287 | 140685287 | 141685287 | Yes |
| ENSG00000272674.3 | PCDHB16 | rs597064 | 5.50E-10 | -0.56 | Brain - Cerebellar Hemisphere | 141185287 | 140685287 | 141685287 | Yes |
| ENSG00000272674.3 | PCDHB16 | rs597064 | 4.00E-08 | -0.45 | Brain - Cerebellum | 141185287 | 140685287 | 141685287 | Yes |
| ENSG00000272674.3 | PCDHB16 | rs610836 | 5.50E-10 | -0.56 | Brain - Cerebellar Hemisphere | 141185287 | 140685287 | 141685287 | Yes |
| ENSG00000272674.3 | PCDHB16 | rs610836 | 4.00E-08 | -0.45 | Brain - Cerebellum | 141185287 | 140685287 | 141685287 | Yes |
| ENSG00000272674.3 | PCDHB16 | rs622424 | 5.50E-10 | -0.56 | Brain - Cerebellar Hemisphere | 141185287 | 140685287 | 141685287 | Yes |
| ENSG00000272674.3 | PCDHB16 | rs622424 | 4.00E-08 | -0.45 | Brain - Cerebellum | 141185287 | 140685287 | 141685287 | Yes |
| ENSG00000272674.3 | PCDHB16 | rs702386 | 6.30E-08 | -0.43 | Brain - Cerebellum | 141185287 | 140685287 | 141685287 | Yes |
| ENSG00000272674.3 | PCDHB16 | rs702386 | 3.70E-07 | -0.49 | Brain - Cerebellar Hemisphere | 141185287 | 140685287 | 141685287 | Yes |
| ENSG00000272674.3 | PCDHB16 | rs17844596 | 2.50E-11 | -0.77 | Brain - Cerebellar Hemisphere | 141185287 | 140685287 | 141685287 | Yes |
| ENSG00000272674.3 | PCDHB16 | rs17844596 | 2.10E-09 | -0.64 | Brain - Cerebellum | 141185287 | 140685287 | 141685287 | Yes |

| | | | | | | | | | |
|---|---|---|---|---|---|---|---|---|---|
| ENSG00000272674.3 | PCDHB16 | rs111745700 | 1.40E-06 | -0.43 | Brain - Cerebellar Hemisphere | 141185287 | 140685287 | 141685287 | Yes |
| ENSG00000272674.3 | PCDHB16 | rs111745700 | 2.00E-06 | -0.38 | Brain - Cerebellum | 141185287 | 140685287 | 141685287 | Yes |
| ENSG00000272674.3 | PCDHB16 | rs629168 | 6.30E-08 | -0.43 | Brain - Cerebellum | 141185287 | 140685287 | 141685287 | Yes |
| ENSG00000272674.3 | PCDHB16 | rs629168 | 3.70E-07 | -0.49 | Brain - Cerebellar Hemisphere | 141185287 | 140685287 | 141685287 | Yes |
| ENSG00000272674.3 | PCDHB16 | rs640835 | 6.30E-08 | -0.43 | Brain - Cerebellum | 141185287 | 140685287 | 141685287 | Yes |
| ENSG00000272674.3 | PCDHB16 | rs640835 | 3.70E-07 | -0.49 | Brain - Cerebellar Hemisphere | 141185287 | 140685287 | 141685287 | Yes |
| ENSG00000272674.3 | PCDHB16 | rs590042 | 4.20E-12 | -0.77 | Brain - Cerebellar Hemisphere | 141185287 | 140685287 | 141685287 | Yes |
| ENSG00000272674.3 | PCDHB16 | rs590042 | 1.30E-10 | -0.67 | Brain - Cerebellum | 141185287 | 140685287 | 141685287 | Yes |
| ENSG00000272674.3 | PCDHB16 | rs643864 | 5.50E-10 | -0.56 | Brain - Cerebellar Hemisphere | 141185287 | 140685287 | 141685287 | Yes |
| ENSG00000272674.3 | PCDHB16 | rs643864 | 4.00E-08 | -0.45 | Brain - Cerebellum | 141185287 | 140685287 | 141685287 | Yes |
| ENSG00000272674.3 | PCDHB16 | rs643918 | 4.20E-12 | -0.77 | Brain - Cerebellar Hemisphere | 141185287 | 140685287 | 141685287 | Yes |
| ENSG00000272674.3 | PCDHB16 | rs643918 | 1.30E-10 | -0.67 | Brain - Cerebellum | 141185287 | 140685287 | 141685287 | Yes |
| ENSG00000272674.3 | PCDHB16 | rs799834 | 2.10E-09 | -0.54 | Brain - Cerebellar Hemisphere | 141185287 | 140685287 | 141685287 | Yes |
| ENSG00000272674.3 | PCDHB16 | rs799834 | 8.30E-08 | -0.43 | Brain - Cerebellum | 141185287 | 140685287 | 141685287 | Yes |
| ENSG00000272674.3 | PCDHB16 | rs799833 | 1.90E-12 | -0.79 | Brain - Cerebellar Hemisphere | 141185287 | 140685287 | 141685287 | Yes |
| ENSG00000272674.3 | PCDHB16 | rs799833 | 2.10E-10 | -0.68 | Brain - Cerebellum | 141185287 | 140685287 | 141685287 | Yes |
| ENSG00000272674.3 | PCDHB16 | rs638054 | 1.90E-12 | -0.79 | Brain - Cerebellar Hemisphere | 141185287 | 140685287 | 141685287 | Yes |
| ENSG00000272674.3 | PCDHB16 | rs638054 | 2.50E-10 | -0.66 | Brain - Cerebellum | 141185287 | 140685287 | 141685287 | Yes |
| ENSG00000272674.3 | PCDHB16 | rs592195 | 5.50E-10 | -0.56 | Brain - Cerebellar Hemisphere | 141185287 | 140685287 | 141685287 | Yes |
| ENSG00000272674.3 | PCDHB16 | rs592195 | 3.30E-08 | -0.45 | Brain - Cerebellum | 141185287 | 140685287 | 141685287 | Yes |
| ENSG00000272674.3 | PCDHB16 | rs782703560 | 3.00E-09 | -0.77 | Brain - Cerebellar Hemisphere | 141185287 | 140685287 | 141685287 | Yes |
| ENSG00000272674.3 | PCDHB16 | rs782703560 | 4.00E-09 | -0.67 | Brain - Cerebellum | 141185287 | 140685287 | 141685287 | Yes |
| ENSG00000272674.3 | PCDHB16 | rs672062 | 4.70E-08 | -0.43 | Brain - Cerebellum | 141185287 | 140685287 | 141685287 | Yes |
| ENSG00000272674.3 | PCDHB16 | rs672062 | 3.70E-07 | -0.49 | Brain - Cerebellar Hemisphere | 141185287 | 140685287 | 141685287 | Yes |
| ENSG00000272674.3 | PCDHB16 | rs676177 | 1.40E-09 | -0.54 | Brain - Cerebellar Hemisphere | 141185287 | 140685287 | 141685287 | Yes |
| ENSG00000272674.3 | PCDHB16 | rs676177 | 1.70E-07 | -0.42 | Brain - Cerebellum | 141185287 | 140685287 | 141685287 | Yes |
| ENSG00000272674.3 | PCDHB16 | rs643502 | 5.50E-10 | -0.56 | Brain - Cerebellar Hemisphere | 141185287 | 140685287 | 141685287 | Yes |
| ENSG00000272674.3 | PCDHB16 | rs643502 | 3.30E-08 | -0.45 | Brain - Cerebellum | 141185287 | 140685287 | 141685287 | Yes |
| ENSG00000272674.3 | PCDHB16 | rs652435 | 1.30E-11 | -0.53 | Brain - Cerebellum | 141185287 | 140685287 | 141685287 | Yes |
| ENSG00000272674.3 | PCDHB16 | rs652435 | 1.50E-07 | -0.55 | Brain - Cerebellar Hemisphere | 141185287 | 140685287 | 141685287 | Yes |
| ENSG00000272674.3 | PCDHB16 | rs651056 | 1.90E-12 | -0.79 | Brain - Cerebellar Hemisphere | 141185287 | 140685287 | 141685287 | Yes |
| ENSG00000272674.3 | PCDHB16 | rs651056 | 2.50E-10 | -0.66 | Brain - Cerebellum | 141185287 | 140685287 | 141685287 | Yes |

| | | | | | | | | | |
|---|---|---|---|---|---|---|---|---|---|
| ENSG00000272674.3 | PCDHB16 | rs628301 | 4.00E-08 | -0.44 | Brain - Cerebellum | 141185287 | 140685287 | 141685287 | Yes |
| ENSG00000272674.3 | PCDHB16 | rs628301 | 2.70E-07 | -0.5 | Brain - Cerebellar Hemisphere | 141185287 | 140685287 | 141685287 | Yes |
| ENSG00000272674.3 | PCDHB16 | rs702384 | 1.90E-12 | -0.79 | Brain - Cerebellar Hemisphere | 141185287 | 140685287 | 141685287 | Yes |
| ENSG00000272674.3 | PCDHB16 | rs702384 | 2.50E-10 | -0.66 | Brain - Cerebellum | 141185287 | 140685287 | 141685287 | Yes |
| ENSG00000272674.3 | PCDHB16 | rs584204 | 4.00E-08 | -0.44 | Brain - Cerebellum | 141185287 | 140685287 | 141685287 | Yes |
| ENSG00000272674.3 | PCDHB16 | rs584204 | 2.70E-07 | -0.5 | Brain - Cerebellar Hemisphere | 141185287 | 140685287 | 141685287 | Yes |
| ENSG00000272674.3 | PCDHB16 | rs671959 | 4.60E-13 | -0.82 | Brain - Cerebellar Hemisphere | 141185287 | 140685287 | 141685287 | Yes |
| ENSG00000272674.3 | PCDHB16 | rs671959 | 2.50E-10 | -0.66 | Brain - Cerebellum | 141185287 | 140685287 | 141685287 | Yes |
| ENSG00000272674.3 | PCDHB16 | rs2907315 | 5.00E-08 | -0.43 | Brain - Cerebellum | 141185287 | 140685287 | 141685287 | Yes |
| ENSG00000272674.3 | PCDHB16 | rs2907315 | 7.70E-08 | -0.52 | Brain - Cerebellar Hemisphere | 141185287 | 140685287 | 141685287 | Yes |
| ENSG00000272674.3 | PCDHB16 | rs631696 | 1.90E-12 | -0.79 | Brain - Cerebellar Hemisphere | 141185287 | 140685287 | 141685287 | Yes |
| ENSG00000272674.3 | PCDHB16 | rs631696 | 2.50E-10 | -0.66 | Brain - Cerebellum | 141185287 | 140685287 | 141685287 | Yes |
| ENSG00000272674.3 | PCDHB16 | rs587342 | 1.80E-08 | -0.44 | Brain - Cerebellum | 141185287 | 140685287 | 141685287 | Yes |
| ENSG00000272674.3 | PCDHB16 | rs587342 | 8.20E-07 | -0.48 | Brain - Cerebellar Hemisphere | 141185287 | 140685287 | 141685287 | Yes |
| ENSG00000272674.3 | PCDHB16 | rs660082 | 1.90E-12 | -0.79 | Brain - Cerebellar Hemisphere | 141185287 | 140685287 | 141685287 | Yes |
| ENSG00000272674.3 | PCDHB16 | rs660082 | 2.50E-10 | -0.66 | Brain - Cerebellum | 141185287 | 140685287 | 141685287 | Yes |
| ENSG00000272674.3 | PCDHB16 | rs618506 | 4.60E-13 | -0.82 | Brain - Cerebellar Hemisphere | 141185287 | 140685287 | 141685287 | Yes |
| ENSG00000272674.3 | PCDHB16 | rs618506 | 2.50E-10 | -0.66 | Brain - Cerebellum | 141185287 | 140685287 | 141685287 | Yes |
| ENSG00000272674.3 | PCDHB16 | rs652104 | 4.60E-13 | -0.82 | Brain - Cerebellar Hemisphere | 141185287 | 140685287 | 141685287 | Yes |
| ENSG00000272674.3 | PCDHB16 | rs652104 | 2.50E-10 | -0.66 | Brain - Cerebellum | 141185287 | 140685287 | 141685287 | Yes |
| ENSG00000272674.3 | PCDHB16 | rs618096 | 9.80E-09 | -0.45 | Brain - Cerebellum | 141185287 | 140685287 | 141685287 | Yes |
| ENSG00000272674.3 | PCDHB16 | rs618096 | 5.90E-07 | -0.48 | Brain - Cerebellar Hemisphere | 141185287 | 140685287 | 141685287 | Yes |
| ENSG00000272674.3 | PCDHB16 | rs10516156 | 2.80E-12 | -0.79 | Brain - Cerebellar Hemisphere | 141185287 | 140685287 | 141685287 | Yes |
| ENSG00000272674.3 | PCDHB16 | rs10516156 | 4.00E-10 | -0.65 | Brain - Cerebellum | 141185287 | 140685287 | 141685287 | Yes |
| ENSG00000272674.3 | PCDHB16 | rs1638537 | 1.50E-08 | -0.44 | Brain - Cerebellum | 141185287 | 140685287 | 141685287 | Yes |
| ENSG00000272674.3 | PCDHB16 | rs1638537 | 2.10E-06 | -0.46 | Brain - Cerebellar Hemisphere | 141185287 | 140685287 | 141685287 | Yes |
| ENSG00000272674.3 | PCDHB16 | rs643844 | 1.20E-08 | -0.45 | Brain - Cerebellum | 141185287 | 140685287 | 141685287 | Yes |
| ENSG00000272674.3 | PCDHB16 | rs643844 | 2.10E-06 | -0.46 | Brain - Cerebellar Hemisphere | 141185287 | 140685287 | 141685287 | Yes |
| ENSG00000272674.3 | PCDHB16 | rs640163 | 2.90E-08 | -0.5 | Brain - Cerebellum | 141185287 | 140685287 | 141685287 | Yes |
| ENSG00000272674.3 | PCDHB16 | rs640163 | 4.30E-05 | -0.43 | Brain - Cerebellar Hemisphere | 141185287 | 140685287 | 141685287 | Yes |
| ENSG00000272674.3 | PCDHB16 | rs147767080 | 8.20E-05 | 1.4 | Brain - Cerebellar Hemisphere | 141185287 | 140685287 | 141685287 | Yes |
| ENSG00000272674.3 | PCDHB16 | rs200817766 | 8.20E-05 | 1.4 | Brain - Cerebellar Hemisphere | 141185287 | 140685287 | 141685287 | Yes |

| | | | | | | | | | |
|---|---|---|---|---|---|---|---|---|---|
| ENSG00000272674.3 | PCDHB16 | rs726684 | 7.40E-05 | 0.4 | Brain - Putamen (basal ganglia) | 141185287 | 140685287 | 141685287 | Yes |
| ENSG00000272674.3 | PCDHB16 | rs56926516 | 4.50E-05 | 0.4 | Brain - Putamen (basal ganglia) | 141185287 | 140685287 | 141685287 | Yes |
| ENSG00000272674.3 | PCDHB16 | rs4912750 | 7.20E-05 | 0.69 | Brain - Cerebellar Hemisphere | 141185287 | 140685287 | 141685287 | Yes |
| ENSG00000272674.3 | PCDHB16 | rs17208404 | 4.50E-05 | 0.4 | Brain - Putamen (basal ganglia) | 141185287 | 140685287 | 141685287 | Yes |
| ENSG00000272674.3 | PCDHB16 | rs3805697 | 2.50E-05 | 0.39 | Brain - Putamen (basal ganglia) | 141185287 | 140685287 | 141685287 | Yes |
| ENSG00000272674.3 | PCDHB16 | rs62379167 | 1.50E-05 | 0.4 | Brain - Putamen (basal ganglia) | 141185287 | 140685287 | 141685287 | Yes |
| ENSG00000272674.3 | PCDHB16 | rs2879228 | 2.10E-05 | 0.39 | Brain - Putamen (basal ganglia) | 141185287 | 140685287 | 141685287 | Yes |
| ENSG00000272674.3 | PCDHB16 | rs6874378 | 2.60E-05 | 0.41 | Brain - Putamen (basal ganglia) | 141185287 | 140685287 | 141685287 | Yes |
| ENSG00000272674.3 | PCDHB16 | rs11953841 | 2.60E-05 | 0.41 | Brain - Putamen (basal ganglia) | 141185287 | 140685287 | 141685287 | Yes |
| ENSG00000272674.3 | PCDHB16 | rs62379198 | 1.50E-05 | 0.4 | Brain - Putamen (basal ganglia) | 141185287 | 140685287 | 141685287 | Yes |
| ENSG00000272674.3 | PCDHB16 | rs34870721 | 5.90E-05 | 0.38 | Brain - Putamen (basal ganglia) | 141185287 | 140685287 | 141685287 | Yes |
| ENSG00000272674.3 | PCDHB16 | rs2233598 | 3.30E-05 | 0.39 | Brain - Putamen (basal ganglia) | 141185287 | 140685287 | 141685287 | Yes |
| ENSG00000272674.3 | PCDHB16 | rs2074912 | 1.20E-05 | 0.43 | Brain - Putamen (basal ganglia) | 141185287 | 140685287 | 141685287 | Yes |
| ENSG00000272674.3 | PCDHB16 | rs918558 | 1.20E-05 | 0.43 | Brain - Putamen (basal ganglia) | 141185287 | 140685287 | 141685287 | Yes |
| ENSG00000272674.3 | PCDHB16 | rs17208425 | 1.20E-05 | 0.43 | Brain - Putamen (basal ganglia) | 141185287 | 140685287 | 141685287 | Yes |
| ENSG00000272674.3 | PCDHB16 | rs62379205 | 1.20E-05 | 0.43 | Brain - Putamen (basal ganglia) | 141185287 | 140685287 | 141685287 | Yes |
| ENSG00000272674.3 | PCDHB16 | rs62379206 | 2.90E-05 | 0.41 | Brain - Putamen (basal ganglia) | 141185287 | 140685287 | 141685287 | Yes |
| ENSG00000272674.3 | PCDHB16 | rs11953270 | 2.90E-05 | 0.41 | Brain - Putamen (basal ganglia) | 141185287 | 140685287 | 141685287 | Yes |
| ENSG00000272674.3 | PCDHB16 | rs11350413 | 4.70E-05 | 0.41 | Brain - Putamen (basal ganglia) | 141185287 | 140685287 | 141685287 | Yes |
| ENSG00000272674.3 | PCDHB16 | rs970069 | 7.30E-05 | 0.39 | Brain - Putamen (basal ganglia) | 141185287 | 140685287 | 141685287 | Yes |
| ENSG00000272674.3 | PCDHB16 | rs13159206 | 2.30E-05 | 0.41 | Brain - Putamen (basal ganglia) | 141185287 | 140685287 | 141685287 | Yes |
| ENSG00000272674.3 | PCDHB16 | rs12522562 | 3.60E-05 | 0.41 | Brain - Putamen (basal ganglia) | 141185287 | 140685287 | 141685287 | Yes |
| ENSG00000272674.3 | PCDHB16 | rs12520873 | 6.90E-05 | 0.39 | Brain - Putamen (basal ganglia) | 141185287 | 140685287 | 141685287 | Yes |
| ENSG00000272674.3 | PCDHB16 | rs17208474 | 2.10E-05 | 0.41 | Brain - Putamen (basal ganglia) | 141185287 | 140685287 | 141685287 | Yes |
| ENSG00000272674.3 | PCDHB16 | rs34577963 | 2.10E-05 | 0.42 | Brain - Putamen (basal ganglia) | 141185287 | 140685287 | 141685287 | Yes |
| ENSG00000272674.3 | PCDHB16 | rs11954658 | 7.30E-05 | 0.39 | Brain - Putamen (basal ganglia) | 141185287 | 140685287 | 141685287 | Yes |
| ENSG00000272674.3 | PCDHB16 | rs143296685 | 7.30E-05 | 0.39 | Brain - Putamen (basal ganglia) | 141185287 | 140685287 | 141685287 | Yes |
| ENSG00000272674.3 | PCDHB16 | rs6881581 | 7.80E-05 | 0.33 | Brain - Putamen (basal ganglia) | 141185287 | 140685287 | 141685287 | Yes |
| ENSG00000272674.3 | PCDHB16 | rs11954998 | 7.30E-05 | 0.39 | Brain - Putamen (basal ganglia) | 141185287 | 140685287 | 141685287 | Yes |
| ENSG00000272674.3 | PCDHB16 | rs3217029 | 7.10E-05 | 0.39 | Brain - Putamen (basal ganglia) | 141185287 | 140685287 | 141685287 | Yes |
| ENSG00000272674.3 | PCDHB16 | rs11951636 | 7.10E-05 | 0.39 | Brain - Putamen (basal ganglia) | 141185287 | 140685287 | 141685287 | Yes |
| ENSG00000272674.3 | PCDHB16 | rs10066420 | 7.60E-05 | 0.34 | Brain - Putamen (basal ganglia) | 141185287 | 140685287 | 141685287 | Yes |

| | | | | | | | | | |
|---|---|---|---|---|---|---|---|---|---|
| ENSG00000272674.3 | PCDHB16 | rs35791104 | 7.10E-05 | 0.39 | Brain - Putamen (basal ganglia) | 141185287 | 140685287 | 141685287 | Yes |
| ENSG00000272674.3 | PCDHB16 | rs62380760 | 7.10E-05 | 0.73 | Brain - Cerebellar Hemisphere | 141185287 | 140685287 | 141685287 | Yes |
| ENSG00000272674.3 | PCDHB16 | rs17286989 | 2.70E-05 | 0.87 | Brain - Cerebellar Hemisphere | 141185287 | 140685287 | 141685287 | Yes |
| ENSG00000272674.3 | PCDHB16 | rs17287002 | 2.70E-05 | 0.87 | Brain - Cerebellar Hemisphere | 141185287 | 140685287 | 141685287 | Yes |
| ENSG00000272674.3 | PCDHB16 | rs61749636 | 2.70E-05 | 0.87 | Brain - Cerebellar Hemisphere | 141185287 | 140685287 | 141685287 | Yes |
| ENSG00000272674.3 | PCDHB16 | rs79862920 | 2.70E-05 | 0.87 | Brain - Cerebellar Hemisphere | 141185287 | 140685287 | 141685287 | Yes |
| ENSG00000272674.3 | PCDHB16 | rs17208502 | 2.70E-05 | 0.87 | Brain - Cerebellar Hemisphere | 141185287 | 140685287 | 141685287 | Yes |
| ENSG00000272674.3 | PCDHB16 | rs17287009 | 2.70E-05 | 0.87 | Brain - Cerebellar Hemisphere | 141185287 | 140685287 | 141685287 | Yes |
| ENSG00000272674.3 | PCDHB16 | rs62380786 | 2.70E-05 | 0.87 | Brain - Cerebellar Hemisphere | 141185287 | 140685287 | 141685287 | Yes |
| ENSG00000272674.3 | PCDHB16 | rs4912765 | 2.70E-05 | 0.87 | Brain - Cerebellar Hemisphere | 141185287 | 140685287 | 141685287 | Yes |
| ENSG00000272674.3 | PCDHB16 | rs62380787 | 2.70E-05 | 0.87 | Brain - Cerebellar Hemisphere | 141185287 | 140685287 | 141685287 | Yes |
| ENSG00000272674.3 | PCDHB16 | rs62380789 | 6.00E-05 | 0.82 | Brain - Cerebellar Hemisphere | 141185287 | 140685287 | 141685287 | Yes |
| ENSG00000272674.3 | PCDHB16 | rs1031904 | 1.20E-05 | -0.45 | Brain - Hippocampus | 141185287 | 140685287 | 141685287 | Yes |
| ENSG00000272674.3 | PCDHB16 | rs1421895 | 5.20E-05 | -0.48 | Brain - Cerebellar Hemisphere | 141185287 | 140685287 | 141685287 | No |
| ENSG00000272674.3 | PCDHB16 | rs1816049 | 4.20E-05 | -0.48 | Brain - Cerebellar Hemisphere | 141185287 | 140685287 | 141685287 | No |
| ENSG00000272674.3 | PCDHB16 | rs59177246 | 4.20E-05 | -0.48 | Brain - Cerebellar Hemisphere | 141185287 | 140685287 | 141685287 | No |
| ENSG00000272674.3 | PCDHB16 | rs72794023 | 5.80E-05 | -0.44 | Brain - Putamen (basal ganglia) | 141185287 | 140685287 | 141685287 | No |
| | PON2 | - | - | - | - | 95406923 | 94906923 | 95906923 | No |
| ENSG00000179295.17 | PTPN11 | rs80291072 | 6.10E-06 | 0.68 | Brain - Substantia nigra | 112482065 | 111982065 | 112982065 | Yes |
| ENSG00000131626.16 | PPFIA1 | rs1788974 | 2.00E-33 | ##### | Brain - Cerebellum | 70324575 | 69824575 | 70824575 | Yes |
| ENSG00000131626.16 | PPFIA1 | rs11356872 | 2.20E-33 | ##### | Brain - Cerebellum | 70324575 | 69824575 | 70824575 | Yes |
| ENSG00000131626.16 | PPFIA1 | rs36037711 | 2.70E-33 | ##### | Brain - Cerebellum | 70324575 | 69824575 | 70824575 | Yes |
| ENSG00000131626.16 | PPFIA1 | rs658590 | 6.20E-33 | ##### | Brain - Cerebellum | 70324575 | 69824575 | 70824575 | Yes |
| ENSG00000131626.16 | PPFIA1 | rs7942766 | 1.90E-32 | ##### | Brain - Cerebellum | 70324575 | 69824575 | 70824575 | Yes |
| ENSG00000131626.16 | PPFIA1 | rs576588 | 1.90E-32 | ##### | Brain - Cerebellum | 70324575 | 69824575 | 70824575 | Yes |
| ENSG00000131626.16 | PPFIA1 | rs488746 | 1.90E-32 | ##### | Brain - Cerebellum | 70324575 | 69824575 | 70824575 | Yes |
| ENSG00000131626.16 | PPFIA1 | rs546724 | 1.90E-32 | ##### | Brain - Cerebellum | 70324575 | 69824575 | 70824575 | Yes |
| ENSG00000131626.16 | PPFIA1 | rs61887401 | 1.90E-32 | ##### | Brain - Cerebellum | 70324575 | 69824575 | 70824575 | Yes |
| ENSG00000131626.16 | PPFIA1 | rs113180277 | 1.90E-32 | ##### | Brain - Cerebellum | 70324575 | 69824575 | 70824575 | Yes |
| ENSG00000131626.16 | PPFIA1 | rs693315 | 1.90E-32 | ##### | Brain - Cerebellum | 70324575 | 69824575 | 70824575 | Yes |
| ENSG00000131626.16 | PPFIA1 | rs531498 | 1.90E-32 | ##### | Brain - Cerebellum | 70324575 | 69824575 | 70824575 | Yes |
| ENSG00000131626.16 | PPFIA1 | rs670310 | 1.90E-32 | ##### | Brain - Cerebellum | 70324575 | 69824575 | 70824575 | Yes |

| | | | | | | | | | |
|---|---|---|---|---|---|---|---|---|---|
| ENSG00000131626.16 | PPFIA1 | rs524333 | 1.90E-32 | ##### | Brain - Cerebellum | 70324575 | 69824575 | 70824575 | Yes |
| ENSG00000131626.16 | PPFIA1 | rs559176 | 1.90E-32 | ##### | Brain - Cerebellum | 70324575 | 69824575 | 70824575 | Yes |
| ENSG00000131626.16 | PPFIA1 | rs687660 | 3.60E-32 | ##### | Brain - Cerebellum | 70324575 | 69824575 | 70824575 | Yes |
| ENSG00000131626.16 | PPFIA1 | rs3133177 | 4.40E-32 | ##### | Brain - Cerebellum | 70324575 | 69824575 | 70824575 | Yes |
| ENSG00000131626.16 | PPFIA1 | rs621749 | 4.40E-32 | ##### | Brain - Cerebellum | 70324575 | 69824575 | 70824575 | Yes |
| ENSG00000131626.16 | PPFIA1 | rs11235935 | 4.40E-32 | ##### | Brain - Cerebellum | 70324575 | 69824575 | 70824575 | Yes |
| ENSG00000131626.16 | PPFIA1 | rs4980597 | 4.40E-32 | ##### | Brain - Cerebellum | 70324575 | 69824575 | 70824575 | Yes |
| ENSG00000131626.16 | PPFIA1 | rs531917 | 4.40E-32 | ##### | Brain - Cerebellum | 70324575 | 69824575 | 70824575 | Yes |
| ENSG00000131626.16 | PPFIA1 | rs665890 | 8.20E-32 | ##### | Brain - Cerebellum | 70324575 | 69824575 | 70824575 | Yes |
| ENSG00000131626.16 | PPFIA1 | rs571554 | 1.50E-31 | ##### | Brain - Cerebellum | 70324575 | 69824575 | 70824575 | Yes |
| ENSG00000131626.16 | PPFIA1 | rs670012 | 3.00E-31 | ##### | Brain - Cerebellum | 70324575 | 69824575 | 70824575 | Yes |
| ENSG00000131626.16 | PPFIA1 | rs4980779 | 3.00E-31 | ##### | Brain - Cerebellum | 70324575 | 69824575 | 70824575 | Yes |
| ENSG00000131626.16 | PPFIA1 | rs7130178 | 3.00E-31 | ##### | Brain - Cerebellum | 70324575 | 69824575 | 70824575 | Yes |
| ENSG00000131626.16 | PPFIA1 | rs7117443 | 5.50E-31 | ##### | Brain - Cerebellum | 70324575 | 69824575 | 70824575 | Yes |
| ENSG00000131626.16 | PPFIA1 | rs625294 | 6.60E-31 | ##### | Brain - Cerebellum | 70324575 | 69824575 | 70824575 | Yes |
| ENSG00000131626.16 | PPFIA1 | rs12295241 | 6.60E-31 | ##### | Brain - Cerebellum | 70324575 | 69824575 | 70824575 | Yes |
| ENSG00000131626.16 | PPFIA1 | rs7109643 | 6.60E-31 | ##### | Brain - Cerebellum | 70324575 | 69824575 | 70824575 | Yes |
| ENSG00000131626.16 | PPFIA1 | rs12283131 | 9.50E-29 | ##### | Brain - Cerebellum | 70324575 | 69824575 | 70824575 | Yes |
| ENSG00000131626.16 | PPFIA1 | rs2282481 | 1.90E-28 | ##### | Brain - Cerebellum | 70324575 | 69824575 | 70824575 | Yes |
| ENSG00000131626.16 | PPFIA1 | rs1788973 | 3.50E-28 | ##### | Brain - Cerebellum | 70324575 | 69824575 | 70824575 | Yes |
| ENSG00000131626.16 | PPFIA1 | rs10683183 | 3.50E-28 | ##### | Brain - Cerebellum | 70324575 | 69824575 | 70824575 | Yes |
| ENSG00000131626.16 | PPFIA1 | rs6592542 | 3.50E-28 | ##### | Brain - Cerebellum | 70324575 | 69824575 | 70824575 | Yes |
| ENSG00000131626.16 | PPFIA1 | rs6592546 | 3.50E-28 | ##### | Brain - Cerebellum | 70324575 | 69824575 | 70824575 | Yes |
| ENSG00000131626.16 | PPFIA1 | rs583347 | 6.10E-28 | ##### | Brain - Cerebellum | 70324575 | 69824575 | 70824575 | Yes |
| ENSG00000131626.16 | PPFIA1 | rs552282 | 6.10E-28 | ##### | Brain - Cerebellum | 70324575 | 69824575 | 70824575 | Yes |
| ENSG00000131626.16 | PPFIA1 | rs674374 | 6.30E-27 | ##### | Brain - Cerebellum | 70324575 | 69824575 | 70824575 | Yes |
| ENSG00000131626.16 | PPFIA1 | rs141709344 | 4.20E-26 | ##### | Brain - Cerebellum | 70324575 | 69824575 | 70824575 | Yes |
| ENSG00000131626.16 | PPFIA1 | rs111368954 | 1.80E-24 | ##### | Brain - Cerebellum | 70324575 | 69824575 | 70824575 | Yes |
| ENSG00000131626.16 | PPFIA1 | rs4980782 | 1.60E-23 | ##### | Brain - Cerebellum | 70324575 | 69824575 | 70824575 | Yes |
| ENSG00000131626.16 | PPFIA1 | rs11236048 | 4.40E-23 | ##### | Brain - Cerebellum | 70324575 | 69824575 | 70824575 | Yes |
| ENSG00000131626.16 | PPFIA1 | rs561050 | 5.00E-23 | ##### | Brain - Cerebellum | 70324575 | 69824575 | 70824575 | Yes |
| ENSG00000131626.16 | PPFIA1 | rs546502 | 4.20E-22 | ##### | Brain - Cerebellum | 70324575 | 69824575 | 70824575 | Yes |

| | | | | | | | | | |
|---|---|---|---|---|---|---|---|---|---|
| ENSG00000131626.16 | PPFIA1 | rs624765 | 4.20E-22 | ##### | Brain - Cerebellum | 70324575 | 69824575 | 70824575 | Yes |
| ENSG00000131626.16 | PPFIA1 | rs3892505 | 4.20E-22 | ##### | Brain - Cerebellum | 70324575 | 69824575 | 70824575 | Yes |
| ENSG00000131626.16 | PPFIA1 | rs35795372 | 4.20E-22 | ##### | Brain - Cerebellum | 70324575 | 69824575 | 70824575 | Yes |
| ENSG00000131626.16 | PPFIA1 | rs664130 | 4.20E-22 | ##### | Brain - Cerebellum | 70324575 | 69824575 | 70824575 | Yes |
| ENSG00000131626.16 | PPFIA1 | rs10591164 | 4.50E-22 | ##### | Brain - Cerebellum | 70324575 | 69824575 | 70824575 | Yes |
| ENSG00000131626.16 | PPFIA1 | rs34141396 | 4.70E-22 | ##### | Brain - Cerebellum | 70324575 | 69824575 | 70824575 | Yes |
| ENSG00000131626.16 | PPFIA1 | rs76294966 | 4.70E-22 | ##### | Brain - Cerebellum | 70324575 | 69824575 | 70824575 | Yes |
| ENSG00000131626.16 | PPFIA1 | rs4980780 | 4.70E-22 | ##### | Brain - Cerebellum | 70324575 | 69824575 | 70824575 | Yes |
| ENSG00000131626.16 | PPFIA1 | rs7129635 | 4.70E-22 | ##### | Brain - Cerebellum | 70324575 | 69824575 | 70824575 | Yes |
| ENSG00000131626.16 | PPFIA1 | rs594763 | 4.70E-22 | ##### | Brain - Cerebellum | 70324575 | 69824575 | 70824575 | Yes |
| ENSG00000131626.16 | PPFIA1 | rs3132421 | 4.70E-22 | ##### | Brain - Cerebellum | 70324575 | 69824575 | 70824575 | Yes |
| ENSG00000131626.16 | PPFIA1 | rs3132438 | 5.40E-22 | ##### | Brain - Cerebellum | 70324575 | 69824575 | 70824575 | Yes |
| ENSG00000131626.16 | PPFIA1 | rs499484 | 2.60E-21 | ##### | Brain - Cerebellum | 70324575 | 69824575 | 70824575 | Yes |
| ENSG00000131626.16 | PPFIA1 | rs670012 | 1.40E-20 | ##### | Brain - Cerebellar Hemisphere | 70324575 | 69824575 | 70824575 | Yes |
| ENSG00000131626.16 | PPFIA1 | rs4980779 | 1.40E-20 | ##### | Brain - Cerebellar Hemisphere | 70324575 | 69824575 | 70824575 | Yes |
| ENSG00000131626.16 | PPFIA1 | rs7130178 | 1.40E-20 | ##### | Brain - Cerebellar Hemisphere | 70324575 | 69824575 | 70824575 | Yes |
| ENSG00000131626.16 | PPFIA1 | rs488746 | 3.50E-20 | ##### | Brain - Cerebellar Hemisphere | 70324575 | 69824575 | 70824575 | Yes |
| ENSG00000131626.16 | PPFIA1 | rs546724 | 3.50E-20 | ##### | Brain - Cerebellar Hemisphere | 70324575 | 69824575 | 70824575 | Yes |
| ENSG00000131626.16 | PPFIA1 | rs61887401 | 3.50E-20 | ##### | Brain - Cerebellar Hemisphere | 70324575 | 69824575 | 70824575 | Yes |
| ENSG00000131626.16 | PPFIA1 | rs113180277 | 3.50E-20 | ##### | Brain - Cerebellar Hemisphere | 70324575 | 69824575 | 70824575 | Yes |
| ENSG00000131626.16 | PPFIA1 | rs670310 | 3.50E-20 | ##### | Brain - Cerebellar Hemisphere | 70324575 | 69824575 | 70824575 | Yes |
| ENSG00000131626.16 | PPFIA1 | rs693315 | 6.40E-20 | ##### | Brain - Cerebellar Hemisphere | 70324575 | 69824575 | 70824575 | Yes |
| ENSG00000131626.16 | PPFIA1 | rs531498 | 6.40E-20 | ##### | Brain - Cerebellar Hemisphere | 70324575 | 69824575 | 70824575 | Yes |
| ENSG00000131626.16 | PPFIA1 | rs524333 | 6.40E-20 | ##### | Brain - Cerebellar Hemisphere | 70324575 | 69824575 | 70824575 | Yes |
| ENSG00000131626.16 | PPFIA1 | rs559176 | 6.40E-20 | ##### | Brain - Cerebellar Hemisphere | 70324575 | 69824575 | 70824575 | Yes |
| ENSG00000131626.16 | PPFIA1 | rs576588 | 9.40E-20 | ##### | Brain - Cerebellar Hemisphere | 70324575 | 69824575 | 70824575 | Yes |
| ENSG00000131626.16 | PPFIA1 | rs571554 | 9.70E-20 | ##### | Brain - Cerebellar Hemisphere | 70324575 | 69824575 | 70824575 | Yes |
| ENSG00000131626.16 | PPFIA1 | rs569732 | 1.50E-19 | ##### | Brain - Cerebellum | 70324575 | 69824575 | 70824575 | Yes |
| ENSG00000131626.16 | PPFIA1 | rs513487 | 1.50E-19 | ##### | Brain - Cerebellum | 70324575 | 69824575 | 70824575 | Yes |
| ENSG00000131626.16 | PPFIA1 | rs679455 | 1.50E-19 | ##### | Brain - Cerebellum | 70324575 | 69824575 | 70824575 | Yes |
| ENSG00000131626.16 | PPFIA1 | rs676889 | 1.50E-19 | ##### | Brain - Cerebellum | 70324575 | 69824575 | 70824575 | Yes |
| ENSG00000131626.16 | PPFIA1 | rs583397 | 1.70E-19 | ##### | Brain - Cerebellum | 70324575 | 69824575 | 70824575 | Yes |

| | | | | | | | | | |
|---|---|---|---|---|---|---|---|---|---|
| ENSG00000131626.16 | PPFIA1 | rs3781649 | 2.10E-19 | ##### | Brain - Cerebellum | 70324575 | 69824575 | 70824575 | Yes |
| ENSG00000131626.16 | PPFIA1 | rs3781646 | 2.10E-19 | ##### | Brain - Cerebellum | 70324575 | 69824575 | 70824575 | Yes |
| ENSG00000131626.16 | PPFIA1 | rs151079983 | 2.10E-19 | ##### | Brain - Cerebellum | 70324575 | 69824575 | 70824575 | Yes |
| ENSG00000131626.16 | PPFIA1 | rs7942766 | 5.00E-19 | ##### | Brain - Cerebellar Hemisphere | 70324575 | 69824575 | 70824575 | Yes |
| ENSG00000131626.16 | PPFIA1 | rs142385955 | 7.10E-19 | ##### | Brain - Cerebellum | 70324575 | 69824575 | 70824575 | Yes |
| ENSG00000131626.16 | PPFIA1 | rs1198234 | 7.10E-19 | ##### | Brain - Cerebellum | 70324575 | 69824575 | 70824575 | Yes |
| ENSG00000131626.16 | PPFIA1 | rs674374 | 7.10E-19 | ##### | Brain - Cerebellar Hemisphere | 70324575 | 69824575 | 70824575 | Yes |
| ENSG00000131626.16 | PPFIA1 | rs1788974 | 7.50E-19 | ##### | Brain - Cerebellar Hemisphere | 70324575 | 69824575 | 70824575 | Yes |
| ENSG00000131626.16 | PPFIA1 | rs1788973 | 7.50E-19 | ##### | Brain - Cerebellar Hemisphere | 70324575 | 69824575 | 70824575 | Yes |
| ENSG00000131626.16 | PPFIA1 | rs10683183 | 7.50E-19 | ##### | Brain - Cerebellar Hemisphere | 70324575 | 69824575 | 70824575 | Yes |
| ENSG00000131626.16 | PPFIA1 | rs6592542 | 7.50E-19 | ##### | Brain - Cerebellar Hemisphere | 70324575 | 69824575 | 70824575 | Yes |
| ENSG00000131626.16 | PPFIA1 | rs6592546 | 7.50E-19 | ##### | Brain - Cerebellar Hemisphere | 70324575 | 69824575 | 70824575 | Yes |
| ENSG00000131626.16 | PPFIA1 | rs638510 | 8.60E-19 | ##### | Brain - Cerebellum | 70324575 | 69824575 | 70824575 | Yes |
| ENSG00000131626.16 | PPFIA1 | rs611216 | 9.00E-19 | ##### | Brain - Cerebellum | 70324575 | 69824575 | 70824575 | Yes |
| ENSG00000131626.16 | PPFIA1 | rs482438 | 9.00E-19 | ##### | Brain - Cerebellum | 70324575 | 69824575 | 70824575 | Yes |
| ENSG00000131626.16 | PPFIA1 | rs643301 | 9.00E-19 | ##### | Brain - Cerebellum | 70324575 | 69824575 | 70824575 | Yes |
| ENSG00000131626.16 | PPFIA1 | rs3802780 | 9.00E-19 | ##### | Brain - Cerebellum | 70324575 | 69824575 | 70824575 | Yes |
| ENSG00000131626.16 | PPFIA1 | rs662018 | 1.00E-18 | ##### | Brain - Cerebellum | 70324575 | 69824575 | 70824575 | Yes |
| ENSG00000131626.16 | PPFIA1 | rs36037711 | 1.00E-18 | ##### | Brain - Cerebellar Hemisphere | 70324575 | 69824575 | 70824575 | Yes |
| ENSG00000131626.16 | PPFIA1 | rs11236275 | 1.20E-18 | ##### | Brain - Cerebellum | 70324575 | 69824575 | 70824575 | Yes |
| ENSG00000131626.16 | PPFIA1 | rs3132424 | 1.60E-18 | ##### | Brain - Cerebellum | 70324575 | 69824575 | 70824575 | Yes |
| ENSG00000131626.16 | PPFIA1 | rs619650 | 1.60E-18 | ##### | Brain - Cerebellum | 70324575 | 69824575 | 70824575 | Yes |
| ENSG00000131626.16 | PPFIA1 | rs625294 | 2.00E-18 | ##### | Brain - Cerebellar Hemisphere | 70324575 | 69824575 | 70824575 | Yes |
| ENSG00000131626.16 | PPFIA1 | rs7117443 | 3.40E-18 | ##### | Brain - Cerebellar Hemisphere | 70324575 | 69824575 | 70824575 | Yes |
| ENSG00000131626.16 | PPFIA1 | rs592501 | 3.50E-18 | ##### | Brain - Cerebellum | 70324575 | 69824575 | 70824575 | Yes |
| ENSG00000131626.16 | PPFIA1 | rs12295241 | 3.70E-18 | ##### | Brain - Cerebellar Hemisphere | 70324575 | 69824575 | 70824575 | Yes |
| ENSG00000131626.16 | PPFIA1 | rs7109643 | 3.70E-18 | ##### | Brain - Cerebellar Hemisphere | 70324575 | 69824575 | 70824575 | Yes |
| ENSG00000131626.16 | PPFIA1 | rs548687 | 3.90E-18 | ##### | Brain - Cerebellum | 70324575 | 69824575 | 70824575 | Yes |
| ENSG00000131626.16 | PPFIA1 | rs1198236 | 3.90E-18 | ##### | Brain - Cerebellum | 70324575 | 69824575 | 70824575 | Yes |
| ENSG00000131626.16 | PPFIA1 | rs3133177 | 4.00E-18 | ##### | Brain - Cerebellar Hemisphere | 70324575 | 69824575 | 70824575 | Yes |
| ENSG00000131626.16 | PPFIA1 | rs621749 | 4.00E-18 | ##### | Brain - Cerebellar Hemisphere | 70324575 | 69824575 | 70824575 | Yes |
| ENSG00000131626.16 | PPFIA1 | rs11235935 | 7.40E-18 | ##### | Brain - Cerebellar Hemisphere | 70324575 | 69824575 | 70824575 | Yes |

| | | | | | | | | | |
|---|---|---|---|---|---|---|---|---|---|
| ENSG00000131626.16 | PPFIA1 | rs4980597 | 7.40E-18 | ##### | Brain - Cerebellar Hemisphere | 70324575 | 69824575 | 70824575 | Yes |
| ENSG00000131626.16 | PPFIA1 | rs531917 | 7.40E-18 | ##### | Brain - Cerebellar Hemisphere | 70324575 | 69824575 | 70824575 | Yes |
| ENSG00000131626.16 | PPFIA1 | rs670141 | 1.20E-17 | ##### | Brain - Cerebellum | 70324575 | 69824575 | 70824575 | Yes |
| ENSG00000131626.16 | PPFIA1 | rs665890 | 1.60E-17 | ##### | Brain - Cerebellar Hemisphere | 70324575 | 69824575 | 70824575 | Yes |
| ENSG00000131626.16 | PPFIA1 | rs12283131 | 1.70E-17 | ##### | Brain - Cerebellar Hemisphere | 70324575 | 69824575 | 70824575 | Yes |
| ENSG00000131626.16 | PPFIA1 | rs3132436 | 2.10E-17 | ##### | Brain - Cerebellum | 70324575 | 69824575 | 70824575 | Yes |
| ENSG00000131626.16 | PPFIA1 | rs644396 | 4.50E-17 | ##### | Brain - Cerebellum | 70324575 | 69824575 | 70824575 | Yes |
| ENSG00000131626.16 | PPFIA1 | rs2353299 | 5.40E-17 | ##### | Brain - Cerebellum | 70324575 | 69824575 | 70824575 | Yes |
| ENSG00000131626.16 | PPFIA1 | rs12284490 | 7.50E-17 | ##### | Brain - Cerebellum | 70324575 | 69824575 | 70824575 | Yes |
| ENSG00000131626.16 | PPFIA1 | rs658590 | 1.40E-16 | ##### | Brain - Cerebellar Hemisphere | 70324575 | 69824575 | 70824575 | Yes |
| ENSG00000131626.16 | PPFIA1 | rs583347 | 1.40E-16 | ##### | Brain - Cerebellar Hemisphere | 70324575 | 69824575 | 70824575 | Yes |
| ENSG00000131626.16 | PPFIA1 | rs552282 | 1.40E-16 | ##### | Brain - Cerebellar Hemisphere | 70324575 | 69824575 | 70824575 | Yes |
| ENSG00000131626.16 | PPFIA1 | rs2282481 | 3.80E-16 | ##### | Brain - Cerebellar Hemisphere | 70324575 | 69824575 | 70824575 | Yes |
| ENSG00000131626.16 | PPFIA1 | rs687660 | 4.00E-16 | ##### | Brain - Cerebellar Hemisphere | 70324575 | 69824575 | 70824575 | Yes |
| ENSG00000131626.16 | PPFIA1 | rs11356872 | 6.00E-16 | ##### | Brain - Cerebellar Hemisphere | 70324575 | 69824575 | 70824575 | Yes |
| ENSG00000131626.16 | PPFIA1 | rs141709344 | 7.20E-16 | ##### | Brain - Cerebellar Hemisphere | 70324575 | 69824575 | 70824575 | Yes |
| ENSG00000131626.16 | PPFIA1 | rs111368954 | 2.20E-14 | ##### | Brain - Cerebellar Hemisphere | 70324575 | 69824575 | 70824575 | Yes |
| ENSG00000131626.16 | PPFIA1 | rs670141 | 2.10E-13 | ##### | Brain - Cerebellar Hemisphere | 70324575 | 69824575 | 70824575 | Yes |
| ENSG00000131626.16 | PPFIA1 | rs644396 | 5.20E-13 | ##### | Brain - Cerebellar Hemisphere | 70324575 | 69824575 | 70824575 | Yes |
| ENSG00000131626.16 | PPFIA1 | rs638510 | 7.90E-13 | ##### | Brain - Cerebellar Hemisphere | 70324575 | 69824575 | 70824575 | Yes |
| ENSG00000131626.16 | PPFIA1 | rs76294966 | 8.80E-13 | ##### | Brain - Cerebellar Hemisphere | 70324575 | 69824575 | 70824575 | Yes |
| ENSG00000131626.16 | PPFIA1 | rs4980780 | 8.80E-13 | ##### | Brain - Cerebellar Hemisphere | 70324575 | 69824575 | 70824575 | Yes |
| ENSG00000131626.16 | PPFIA1 | rs7129635 | 8.80E-13 | ##### | Brain - Cerebellar Hemisphere | 70324575 | 69824575 | 70824575 | Yes |
| ENSG00000131626.16 | PPFIA1 | rs594763 | 8.80E-13 | ##### | Brain - Cerebellar Hemisphere | 70324575 | 69824575 | 70824575 | Yes |
| ENSG00000131626.16 | PPFIA1 | rs34141396 | 1.40E-12 | ##### | Brain - Cerebellar Hemisphere | 70324575 | 69824575 | 70824575 | Yes |
| ENSG00000131626.16 | PPFIA1 | rs3132421 | 1.40E-12 | ##### | Brain - Cerebellar Hemisphere | 70324575 | 69824575 | 70824575 | Yes |
| ENSG00000131626.16 | PPFIA1 | rs12283131 | 2.00E-12 | ##### | Brain - Cortex | 70324575 | 69824575 | 70824575 | Yes |
| ENSG00000131626.16 | PPFIA1 | rs3132438 | 2.50E-12 | ##### | Brain - Cerebellar Hemisphere | 70324575 | 69824575 | 70824575 | Yes |
| ENSG00000131626.16 | PPFIA1 | rs7942766 | 3.10E-12 | ##### | Brain - Cortex | 70324575 | 69824575 | 70824575 | Yes |
| ENSG00000131626.16 | PPFIA1 | rs576588 | 3.10E-12 | ##### | Brain - Cortex | 70324575 | 69824575 | 70824575 | Yes |
| ENSG00000131626.16 | PPFIA1 | rs561050 | 3.30E-12 | ##### | Brain - Cerebellar Hemisphere | 70324575 | 69824575 | 70824575 | Yes |
| ENSG00000131626.16 | PPFIA1 | rs488746 | 6.90E-12 | ##### | Brain - Cortex | 70324575 | 69824575 | 70824575 | Yes |

| | | | | | | | | | |
|---|---|---|---|---|---|---|---|---|---|
| ENSG00000131626.16 | PPFIA1 | rs546724 | 6.90E-12 | ##### | Brain - Cortex | 70324575 | 69824575 | 70824575 | Yes |
| ENSG00000131626.16 | PPFIA1 | rs61887401 | 6.90E-12 | ##### | Brain - Cortex | 70324575 | 69824575 | 70824575 | Yes |
| ENSG00000131626.16 | PPFIA1 | rs113180277 | 6.90E-12 | ##### | Brain - Cortex | 70324575 | 69824575 | 70824575 | Yes |
| ENSG00000131626.16 | PPFIA1 | rs670310 | 6.90E-12 | ##### | Brain - Cortex | 70324575 | 69824575 | 70824575 | Yes |
| ENSG00000131626.16 | PPFIA1 | rs569732 | 7.70E-12 | ##### | Brain - Cerebellar Hemisphere | 70324575 | 69824575 | 70824575 | Yes |
| ENSG00000131626.16 | PPFIA1 | rs513487 | 7.70E-12 | ##### | Brain - Cerebellar Hemisphere | 70324575 | 69824575 | 70824575 | Yes |
| ENSG00000131626.16 | PPFIA1 | rs679455 | 7.70E-12 | ##### | Brain - Cerebellar Hemisphere | 70324575 | 69824575 | 70824575 | Yes |
| ENSG00000131626.16 | PPFIA1 | rs676889 | 7.70E-12 | ##### | Brain - Cerebellar Hemisphere | 70324575 | 69824575 | 70824575 | Yes |
| ENSG00000131626.16 | PPFIA1 | rs11236275 | 7.70E-12 | ##### | Brain - Cerebellar Hemisphere | 70324575 | 69824575 | 70824575 | Yes |
| ENSG00000131626.16 | PPFIA1 | rs2353299 | 8.40E-12 | ##### | Brain - Cerebellar Hemisphere | 70324575 | 69824575 | 70824575 | Yes |
| ENSG00000131626.16 | PPFIA1 | rs670012 | 8.70E-12 | ##### | Brain - Cortex | 70324575 | 69824575 | 70824575 | Yes |
| ENSG00000131626.16 | PPFIA1 | rs4980779 | 8.70E-12 | ##### | Brain - Cortex | 70324575 | 69824575 | 70824575 | Yes |
| ENSG00000131626.16 | PPFIA1 | rs7130178 | 8.70E-12 | ##### | Brain - Cortex | 70324575 | 69824575 | 70824575 | Yes |
| ENSG00000131626.16 | PPFIA1 | rs3133177 | 9.40E-12 | ##### | Brain - Cortex | 70324575 | 69824575 | 70824575 | Yes |
| ENSG00000131626.16 | PPFIA1 | rs621749 | 9.40E-12 | ##### | Brain - Cortex | 70324575 | 69824575 | 70824575 | Yes |
| ENSG00000131626.16 | PPFIA1 | rs693315 | 1.10E-11 | ##### | Brain - Cortex | 70324575 | 69824575 | 70824575 | Yes |
| ENSG00000131626.16 | PPFIA1 | rs531498 | 1.10E-11 | ##### | Brain - Cortex | 70324575 | 69824575 | 70824575 | Yes |
| ENSG00000131626.16 | PPFIA1 | rs524333 | 1.10E-11 | ##### | Brain - Cortex | 70324575 | 69824575 | 70824575 | Yes |
| ENSG00000131626.16 | PPFIA1 | rs559176 | 1.10E-11 | ##### | Brain - Cortex | 70324575 | 69824575 | 70824575 | Yes |
| ENSG00000131626.16 | PPFIA1 | rs625294 | 1.20E-11 | ##### | Brain - Cortex | 70324575 | 69824575 | 70824575 | Yes |
| ENSG00000131626.16 | PPFIA1 | rs665890 | 1.40E-11 | ##### | Brain - Cortex | 70324575 | 69824575 | 70824575 | Yes |
| ENSG00000131626.16 | PPFIA1 | rs11235935 | 1.60E-11 | ##### | Brain - Cortex | 70324575 | 69824575 | 70824575 | Yes |
| ENSG00000131626.16 | PPFIA1 | rs4980597 | 1.60E-11 | ##### | Brain - Cortex | 70324575 | 69824575 | 70824575 | Yes |
| ENSG00000131626.16 | PPFIA1 | rs531917 | 1.60E-11 | ##### | Brain - Cortex | 70324575 | 69824575 | 70824575 | Yes |
| ENSG00000131626.16 | PPFIA1 | rs546502 | 1.60E-11 | ##### | Brain - Cerebellar Hemisphere | 70324575 | 69824575 | 70824575 | Yes |
| ENSG00000131626.16 | PPFIA1 | rs624765 | 1.60E-11 | ##### | Brain - Cerebellar Hemisphere | 70324575 | 69824575 | 70824575 | Yes |
| ENSG00000131626.16 | PPFIA1 | rs3892505 | 1.60E-11 | ##### | Brain - Cerebellar Hemisphere | 70324575 | 69824575 | 70824575 | Yes |
| ENSG00000131626.16 | PPFIA1 | rs12295241 | 1.90E-11 | ##### | Brain - Cortex | 70324575 | 69824575 | 70824575 | Yes |
| ENSG00000131626.16 | PPFIA1 | rs7109643 | 1.90E-11 | ##### | Brain - Cortex | 70324575 | 69824575 | 70824575 | Yes |
| ENSG00000131626.16 | PPFIA1 | rs7117443 | 2.20E-11 | ##### | Brain - Cortex | 70324575 | 69824575 | 70824575 | Yes |
| ENSG00000131626.16 | PPFIA1 | rs10591164 | 2.50E-11 | ##### | Brain - Cerebellar Hemisphere | 70324575 | 69824575 | 70824575 | Yes |
| ENSG00000131626.16 | PPFIA1 | rs35795372 | 2.50E-11 | ##### | Brain - Cerebellar Hemisphere | 70324575 | 69824575 | 70824575 | Yes |

| | | | | | | | | | |
|---|---|---|---|---|---|---|---|---|---|
| ENSG00000131626.16 | PPFIA1 | rs664130 | 2.50E-11 | ##### | Brain - Cerebellar Hemisphere | 70324575 | 69824575 | 70824575 | Yes |
| ENSG00000131626.16 | PPFIA1 | rs11236048 | 2.60E-11 | ##### | Brain - Cerebellar Hemisphere | 70324575 | 69824575 | 70824575 | Yes |
| ENSG00000131626.16 | PPFIA1 | rs499484 | 3.60E-11 | ##### | Brain - Cerebellar Hemisphere | 70324575 | 69824575 | 70824575 | Yes |
| ENSG00000131626.16 | PPFIA1 | rs571554 | 4.00E-11 | ##### | Brain - Cortex | 70324575 | 69824575 | 70824575 | Yes |
| ENSG00000131626.16 | PPFIA1 | rs4980782 | 4.30E-11 | ##### | Brain - Cerebellar Hemisphere | 70324575 | 69824575 | 70824575 | Yes |
| ENSG00000131626.16 | PPFIA1 | rs7948535 | 5.30E-11 | ##### | Brain - Cerebellum | 70324575 | 69824575 | 70824575 | Yes |
| ENSG00000131626.16 | PPFIA1 | rs619650 | 9.70E-11 | ##### | Brain - Cerebellar Hemisphere | 70324575 | 69824575 | 70824575 | Yes |
| ENSG00000131626.16 | PPFIA1 | rs3132424 | 9.70E-11 | ##### | Brain - Cerebellar Hemisphere | 70324575 | 69824575 | 70824575 | Yes |
| ENSG00000131626.16 | PPFIA1 | rs12284490 | 1.90E-10 | ##### | Brain - Cerebellar Hemisphere | 70324575 | 69824575 | 70824575 | Yes |
| ENSG00000131626.16 | PPFIA1 | rs583397 | 2.10E-10 | ##### | Brain - Cerebellar Hemisphere | 70324575 | 69824575 | 70824575 | Yes |
| ENSG00000131626.16 | PPFIA1 | rs141709344 | 2.30E-10 | ##### | Brain - Cortex | 70324575 | 69824575 | 70824575 | Yes |
| ENSG00000131626.16 | PPFIA1 | rs36037711 | 2.50E-10 | ##### | Brain - Cortex | 70324575 | 69824575 | 70824575 | Yes |
| ENSG00000131626.16 | PPFIA1 | rs1788974 | 2.90E-10 | ##### | Brain - Cortex | 70324575 | 69824575 | 70824575 | Yes |
| ENSG00000131626.16 | PPFIA1 | rs662018 | 3.20E-10 | ##### | Brain - Cerebellar Hemisphere | 70324575 | 69824575 | 70824575 | Yes |
| ENSG00000131626.16 | PPFIA1 | rs658590 | 5.40E-10 | ##### | Brain - Cortex | 70324575 | 69824575 | 70824575 | Yes |
| ENSG00000131626.16 | PPFIA1 | rs2282481 | 6.50E-10 | ##### | Brain - Cortex | 70324575 | 69824575 | 70824575 | Yes |
| ENSG00000131626.16 | PPFIA1 | rs1788973 | 7.50E-10 | ##### | Brain - Cortex | 70324575 | 69824575 | 70824575 | Yes |
| ENSG00000131626.16 | PPFIA1 | rs10683183 | 7.50E-10 | ##### | Brain - Cortex | 70324575 | 69824575 | 70824575 | Yes |
| ENSG00000131626.16 | PPFIA1 | rs6592542 | 7.50E-10 | ##### | Brain - Cortex | 70324575 | 69824575 | 70824575 | Yes |
| ENSG00000131626.16 | PPFIA1 | rs6592546 | 7.50E-10 | ##### | Brain - Cortex | 70324575 | 69824575 | 70824575 | Yes |
| ENSG00000131626.16 | PPFIA1 | rs11356872 | 9.40E-10 | ##### | Brain - Cortex | 70324575 | 69824575 | 70824575 | Yes |
| ENSG00000131626.16 | PPFIA1 | rs687660 | 1.10E-09 | ##### | Brain - Cortex | 70324575 | 69824575 | 70824575 | Yes |
| ENSG00000131626.16 | PPFIA1 | rs674374 | 1.30E-09 | ##### | Brain - Cortex | 70324575 | 69824575 | 70824575 | Yes |
| ENSG00000131626.16 | PPFIA1 | rs583347 | 1.50E-09 | ##### | Brain - Cortex | 70324575 | 69824575 | 70824575 | Yes |
| ENSG00000131626.16 | PPFIA1 | rs552282 | 1.50E-09 | ##### | Brain - Cortex | 70324575 | 69824575 | 70824575 | Yes |
| ENSG00000131626.16 | PPFIA1 | rs142385955 | 2.70E-09 | ##### | Brain - Cerebellar Hemisphere | 70324575 | 69824575 | 70824575 | Yes |
| ENSG00000131626.16 | PPFIA1 | rs1198234 | 2.70E-09 | ##### | Brain - Cerebellar Hemisphere | 70324575 | 69824575 | 70824575 | Yes |
| ENSG00000131626.16 | PPFIA1 | rs592501 | 2.70E-09 | ##### | Brain - Cerebellar Hemisphere | 70324575 | 69824575 | 70824575 | Yes |
| ENSG00000131626.16 | PPFIA1 | rs111368954 | 3.90E-09 | ##### | Brain - Cortex | 70324575 | 69824575 | 70824575 | Yes |
| ENSG00000131626.16 | PPFIA1 | rs576588 | 4.70E-09 | ##### | Brain - Amygdala | 70324575 | 69824575 | 70824575 | Yes |
| ENSG00000131626.16 | PPFIA1 | rs611216 | 5.90E-09 | ##### | Brain - Cerebellar Hemisphere | 70324575 | 69824575 | 70824575 | Yes |
| ENSG00000131626.16 | PPFIA1 | rs548687 | 5.90E-09 | ##### | Brain - Cerebellar Hemisphere | 70324575 | 69824575 | 70824575 | Yes |

| | | | | | | | | | |
|---|---|---|---|---|---|---|---|---|---|
| ENSG00000131626.16 | PPFIA1 | rs1198236 | 5.90E-09 | ##### | Brain - Cerebellar Hemisphere | 70324575 | 69824575 | 70824575 | Yes |
| ENSG00000131626.16 | PPFIA1 | rs482438 | 5.90E-09 | ##### | Brain - Cerebellar Hemisphere | 70324575 | 69824575 | 70824575 | Yes |
| ENSG00000131626.16 | PPFIA1 | rs643301 | 5.90E-09 | ##### | Brain - Cerebellar Hemisphere | 70324575 | 69824575 | 70824575 | Yes |
| ENSG00000131626.16 | PPFIA1 | rs3802780 | 5.90E-09 | ##### | Brain - Cerebellar Hemisphere | 70324575 | 69824575 | 70824575 | Yes |
| ENSG00000131626.16 | PPFIA1 | rs3781649 | 6.00E-09 | ##### | Brain - Cerebellar Hemisphere | 70324575 | 69824575 | 70824575 | Yes |
| ENSG00000131626.16 | PPFIA1 | rs3781646 | 6.00E-09 | ##### | Brain - Cerebellar Hemisphere | 70324575 | 69824575 | 70824575 | Yes |
| ENSG00000131626.16 | PPFIA1 | rs151079983 | 6.00E-09 | ##### | Brain - Cerebellar Hemisphere | 70324575 | 69824575 | 70824575 | Yes |
| ENSG00000131626.16 | PPFIA1 | rs693315 | 6.50E-09 | ##### | Brain - Amygdala | 70324575 | 69824575 | 70824575 | Yes |
| ENSG00000131626.16 | PPFIA1 | rs531498 | 6.50E-09 | ##### | Brain - Amygdala | 70324575 | 69824575 | 70824575 | Yes |
| ENSG00000131626.16 | PPFIA1 | rs524333 | 6.50E-09 | ##### | Brain - Amygdala | 70324575 | 69824575 | 70824575 | Yes |
| ENSG00000131626.16 | PPFIA1 | rs559176 | 6.50E-09 | ##### | Brain - Amygdala | 70324575 | 69824575 | 70824575 | Yes |
| ENSG00000131626.16 | PPFIA1 | rs7942766 | 1.10E-08 | ##### | Brain - Amygdala | 70324575 | 69824575 | 70824575 | Yes |
| ENSG00000131626.16 | PPFIA1 | rs488746 | 1.30E-08 | ##### | Brain - Amygdala | 70324575 | 69824575 | 70824575 | Yes |
| ENSG00000131626.16 | PPFIA1 | rs546724 | 1.30E-08 | ##### | Brain - Amygdala | 70324575 | 69824575 | 70824575 | Yes |
| ENSG00000131626.16 | PPFIA1 | rs61887401 | 1.30E-08 | ##### | Brain - Amygdala | 70324575 | 69824575 | 70824575 | Yes |
| ENSG00000131626.16 | PPFIA1 | rs113180277 | 1.30E-08 | ##### | Brain - Amygdala | 70324575 | 69824575 | 70824575 | Yes |
| ENSG00000131626.16 | PPFIA1 | rs670310 | 1.30E-08 | ##### | Brain - Amygdala | 70324575 | 69824575 | 70824575 | Yes |
| ENSG00000131626.16 | PPFIA1 | rs638510 | 1.40E-08 | ##### | Brain - Cortex | 70324575 | 69824575 | 70824575 | Yes |
| ENSG00000131626.16 | PPFIA1 | rs571554 | 1.40E-08 | ##### | Brain - Amygdala | 70324575 | 69824575 | 70824575 | Yes |
| ENSG00000131626.16 | PPFIA1 | rs670012 | 1.50E-08 | ##### | Brain - Amygdala | 70324575 | 69824575 | 70824575 | Yes |
| ENSG00000131626.16 | PPFIA1 | rs4980779 | 1.50E-08 | ##### | Brain - Amygdala | 70324575 | 69824575 | 70824575 | Yes |
| ENSG00000131626.16 | PPFIA1 | rs7130178 | 1.50E-08 | ##### | Brain - Amygdala | 70324575 | 69824575 | 70824575 | Yes |
| ENSG00000131626.16 | PPFIA1 | rs3132436 | 1.50E-08 | ##### | Brain - Cerebellar Hemisphere | 70324575 | 69824575 | 70824575 | Yes |
| ENSG00000131626.16 | PPFIA1 | rs576588 | 1.60E-08 | ##### | Brain - Putamen (basal gangli | 70324575 | 69824575 | 70824575 | Yes |
| ENSG00000131626.16 | PPFIA1 | rs7942766 | 1.70E-08 | ##### | Brain - Frontal Cortex (BA9) | 70324575 | 69824575 | 70824575 | Yes |
| ENSG00000131626.16 | PPFIA1 | rs7942766 | 1.80E-08 | ##### | Brain - Putamen (basal gangli | 70324575 | 69824575 | 70824575 | Yes |
| ENSG00000131626.16 | PPFIA1 | rs111368954 | 2.10E-08 | ##### | Brain - Amygdala | 70324575 | 69824575 | 70824575 | Yes |
| ENSG00000131626.16 | PPFIA1 | rs488746 | 2.40E-08 | ##### | Brain - Putamen (basal gangli | 70324575 | 69824575 | 70824575 | Yes |
| ENSG00000131626.16 | PPFIA1 | rs546724 | 2.40E-08 | ##### | Brain - Putamen (basal gangli | 70324575 | 69824575 | 70824575 | Yes |
| ENSG00000131626.16 | PPFIA1 | rs61887401 | 2.40E-08 | ##### | Brain - Putamen (basal gangli | 70324575 | 69824575 | 70824575 | Yes |
| ENSG00000131626.16 | PPFIA1 | rs113180277 | 2.40E-08 | ##### | Brain - Putamen (basal gangli | 70324575 | 69824575 | 70824575 | Yes |
| ENSG00000131626.16 | PPFIA1 | rs670310 | 2.40E-08 | ##### | Brain - Putamen (basal gangli | 70324575 | 69824575 | 70824575 | Yes |

| | | | | | | | | | |
|---|---|---|---|---|---|---|---|---|---|
| ENSG00000131626.16 | PPFIA1 | rs576588 | 2.50E-08 | ##### | Brain - Frontal Cortex (BA9) | 70324575 | 69824575 | 70824575 | Yes |
| ENSG00000131626.16 | PPFIA1 | rs11235935 | 2.70E-08 | ##### | Brain - Amygdala | 70324575 | 69824575 | 70824575 | Yes |
| ENSG00000131626.16 | PPFIA1 | rs4980597 | 2.70E-08 | ##### | Brain - Amygdala | 70324575 | 69824575 | 70824575 | Yes |
| ENSG00000131626.16 | PPFIA1 | rs531917 | 2.70E-08 | ##### | Brain - Amygdala | 70324575 | 69824575 | 70824575 | Yes |
| ENSG00000131626.16 | PPFIA1 | rs665890 | 2.80E-08 | ##### | Brain - Amygdala | 70324575 | 69824575 | 70824575 | Yes |
| ENSG00000131626.16 | PPFIA1 | rs1788974 | 3.10E-08 | ##### | Brain - Putamen (basal gangli | 70324575 | 69824575 | 70824575 | Yes |
| ENSG00000131626.16 | PPFIA1 | rs12295241 | 3.20E-08 | ##### | Brain - Amygdala | 70324575 | 69824575 | 70824575 | Yes |
| ENSG00000131626.16 | PPFIA1 | rs7109643 | 3.20E-08 | ##### | Brain - Amygdala | 70324575 | 69824575 | 70824575 | Yes |
| ENSG00000131626.16 | PPFIA1 | rs36037711 | 3.40E-08 | ##### | Brain - Putamen (basal gangli | 70324575 | 69824575 | 70824575 | Yes |
| ENSG00000131626.16 | PPFIA1 | rs665890 | 3.60E-08 | ##### | Brain - Frontal Cortex (BA9) | 70324575 | 69824575 | 70824575 | Yes |
| ENSG00000131626.16 | PPFIA1 | rs11356872 | 4.20E-08 | ##### | Brain - Amygdala | 70324575 | 69824575 | 70824575 | Yes |
| ENSG00000131626.16 | PPFIA1 | rs36037711 | 4.80E-08 | ##### | Brain - Amygdala | 70324575 | 69824575 | 70824575 | Yes |
| ENSG00000131626.16 | PPFIA1 | rs3133177 | 5.10E-08 | ##### | Brain - Putamen (basal gangli | 70324575 | 69824575 | 70824575 | Yes |
| ENSG00000131626.16 | PPFIA1 | rs621749 | 5.10E-08 | ##### | Brain - Putamen (basal gangli | 70324575 | 69824575 | 70824575 | Yes |
| ENSG00000131626.16 | PPFIA1 | rs3133177 | 5.10E-08 | ##### | Brain - Amygdala | 70324575 | 69824575 | 70824575 | Yes |
| ENSG00000131626.16 | PPFIA1 | rs621749 | 5.10E-08 | ##### | Brain - Amygdala | 70324575 | 69824575 | 70824575 | Yes |
| ENSG00000131626.16 | PPFIA1 | rs488746 | 5.20E-08 | ##### | Brain - Frontal Cortex (BA9) | 70324575 | 69824575 | 70824575 | Yes |
| ENSG00000131626.16 | PPFIA1 | rs546724 | 5.20E-08 | ##### | Brain - Frontal Cortex (BA9) | 70324575 | 69824575 | 70824575 | Yes |
| ENSG00000131626.16 | PPFIA1 | rs61887401 | 5.20E-08 | ##### | Brain - Frontal Cortex (BA9) | 70324575 | 69824575 | 70824575 | Yes |
| ENSG00000131626.16 | PPFIA1 | rs113180277 | 5.20E-08 | ##### | Brain - Frontal Cortex (BA9) | 70324575 | 69824575 | 70824575 | Yes |
| ENSG00000131626.16 | PPFIA1 | rs693315 | 5.20E-08 | ##### | Brain - Frontal Cortex (BA9) | 70324575 | 69824575 | 70824575 | Yes |
| ENSG00000131626.16 | PPFIA1 | rs531498 | 5.20E-08 | ##### | Brain - Frontal Cortex (BA9) | 70324575 | 69824575 | 70824575 | Yes |
| ENSG00000131626.16 | PPFIA1 | rs670310 | 5.20E-08 | ##### | Brain - Frontal Cortex (BA9) | 70324575 | 69824575 | 70824575 | Yes |
| ENSG00000131626.16 | PPFIA1 | rs524333 | 5.20E-08 | ##### | Brain - Frontal Cortex (BA9) | 70324575 | 69824575 | 70824575 | Yes |
| ENSG00000131626.16 | PPFIA1 | rs559176 | 5.20E-08 | ##### | Brain - Frontal Cortex (BA9) | 70324575 | 69824575 | 70824575 | Yes |
| ENSG00000131626.16 | PPFIA1 | rs670012 | 5.40E-08 | ##### | Brain - Putamen (basal gangli | 70324575 | 69824575 | 70824575 | Yes |
| ENSG00000131626.16 | PPFIA1 | rs4980779 | 5.40E-08 | ##### | Brain - Putamen (basal gangli | 70324575 | 69824575 | 70824575 | Yes |
| ENSG00000131626.16 | PPFIA1 | rs7130178 | 5.40E-08 | ##### | Brain - Putamen (basal gangli | 70324575 | 69824575 | 70824575 | Yes |
| ENSG00000131626.16 | PPFIA1 | rs7942766 | 5.40E-08 | ##### | Brain - Nucleus accumbens (b | 70324575 | 69824575 | 70824575 | Yes |
| ENSG00000131626.16 | PPFIA1 | rs1788974 | 5.70E-08 | ##### | Brain - Amygdala | 70324575 | 69824575 | 70824575 | Yes |
| ENSG00000131626.16 | PPFIA1 | rs576588 | 5.90E-08 | ##### | Brain - Nucleus accumbens (b | 70324575 | 69824575 | 70824575 | Yes |
| ENSG00000131626.16 | PPFIA1 | rs625294 | 6.00E-08 | ##### | Brain - Amygdala | 70324575 | 69824575 | 70824575 | Yes |

| | | | | | | | | | |
|---|---|---|---|---|---|---|---|---|---|
| ENSG00000131626.16 | PPFIA1 | rs670012 | 6.20E-08 | ##### | Brain - Nucleus accumbens (b | 70324575 | 69824575 | 70824575 | Yes |
| ENSG00000131626.16 | PPFIA1 | rs4980779 | 6.20E-08 | ##### | Brain - Nucleus accumbens (b | 70324575 | 69824575 | 70824575 | Yes |
| ENSG00000131626.16 | PPFIA1 | rs7130178 | 6.20E-08 | ##### | Brain - Nucleus accumbens (b | 70324575 | 69824575 | 70824575 | Yes |
| ENSG00000131626.16 | PPFIA1 | rs561050 | 6.70E-08 | ##### | Brain - Frontal Cortex (BA9) | 70324575 | 69824575 | 70824575 | Yes |
| ENSG00000131626.16 | PPFIA1 | rs665890 | 7.50E-08 | ##### | Brain - Putamen (basal gangli | 70324575 | 69824575 | 70824575 | Yes |
| ENSG00000131626.16 | PPFIA1 | rs571554 | 7.60E-08 | ##### | Brain - Nucleus accumbens (b | 70324575 | 69824575 | 70824575 | Yes |
| ENSG00000131626.16 | PPFIA1 | rs571554 | 7.70E-08 | ##### | Brain - Putamen (basal gangli | 70324575 | 69824575 | 70824575 | Yes |
| ENSG00000131626.16 | PPFIA1 | rs3133177 | 7.70E-08 | ##### | Brain - Frontal Cortex (BA9) | 70324575 | 69824575 | 70824575 | Yes |
| ENSG00000131626.16 | PPFIA1 | rs621749 | 7.70E-08 | ##### | Brain - Frontal Cortex (BA9) | 70324575 | 69824575 | 70824575 | Yes |
| ENSG00000131626.16 | PPFIA1 | rs11235935 | 7.70E-08 | ##### | Brain - Frontal Cortex (BA9) | 70324575 | 69824575 | 70824575 | Yes |
| ENSG00000131626.16 | PPFIA1 | rs4980597 | 7.70E-08 | ##### | Brain - Frontal Cortex (BA9) | 70324575 | 69824575 | 70824575 | Yes |
| ENSG00000131626.16 | PPFIA1 | rs531917 | 7.70E-08 | ##### | Brain - Frontal Cortex (BA9) | 70324575 | 69824575 | 70824575 | Yes |
| ENSG00000131626.16 | PPFIA1 | rs576588 | 7.90E-08 | ##### | Brain - Spinal cord (cervical c- | 70324575 | 69824575 | 70824575 | Yes |
| ENSG00000131626.16 | PPFIA1 | rs561050 | 8.80E-08 | ##### | Brain - Amygdala | 70324575 | 69824575 | 70824575 | Yes |
| ENSG00000131626.16 | PPFIA1 | rs7117443 | 9.00E-08 | ##### | Brain - Amygdala | 70324575 | 69824575 | 70824575 | Yes |
| ENSG00000131626.16 | PPFIA1 | rs3132424 | 9.40E-08 | ##### | Brain - Cortex | 70324575 | 69824575 | 70824575 | Yes |
| ENSG00000131626.16 | PPFIA1 | rs619650 | 9.60E-08 | ##### | Brain - Cortex | 70324575 | 69824575 | 70824575 | Yes |
| ENSG00000131626.16 | PPFIA1 | rs12284490 | 9.60E-08 | ##### | Brain - Cortex | 70324575 | 69824575 | 70824575 | Yes |
| ENSG00000131626.16 | PPFIA1 | rs670012 | 9.70E-08 | ##### | Brain - Caudate (basal ganglia | 70324575 | 69824575 | 70824575 | Yes |
| ENSG00000131626.16 | PPFIA1 | rs4980779 | 9.70E-08 | ##### | Brain - Caudate (basal ganglia | 70324575 | 69824575 | 70824575 | Yes |
| ENSG00000131626.16 | PPFIA1 | rs7130178 | 9.70E-08 | ##### | Brain - Caudate (basal ganglia | 70324575 | 69824575 | 70824575 | Yes |
| ENSG00000131626.16 | PPFIA1 | rs687660 | 1.00E-07 | ##### | Brain - Amygdala | 70324575 | 69824575 | 70824575 | Yes |
| ENSG00000131626.16 | PPFIA1 | rs658590 | 1.00E-07 | ##### | Brain - Amygdala | 70324575 | 69824575 | 70824575 | Yes |
| ENSG00000131626.16 | PPFIA1 | rs488746 | 1.00E-07 | ##### | Brain - Nucleus accumbens (b | 70324575 | 69824575 | 70824575 | Yes |
| ENSG00000131626.16 | PPFIA1 | rs546724 | 1.00E-07 | ##### | Brain - Nucleus accumbens (b | 70324575 | 69824575 | 70824575 | Yes |
| ENSG00000131626.16 | PPFIA1 | rs61887401 | 1.00E-07 | ##### | Brain - Nucleus accumbens (b | 70324575 | 69824575 | 70824575 | Yes |
| ENSG00000131626.16 | PPFIA1 | rs113180277 | 1.00E-07 | ##### | Brain - Nucleus accumbens (b | 70324575 | 69824575 | 70824575 | Yes |
| ENSG00000131626.16 | PPFIA1 | rs670310 | 1.00E-07 | ##### | Brain - Nucleus accumbens (b | 70324575 | 69824575 | 70824575 | Yes |
| ENSG00000131626.16 | PPFIA1 | rs583397 | 1.10E-07 | ##### | Brain - Spinal cord (cervical c- | 70324575 | 69824575 | 70824575 | Yes |
| ENSG00000131626.16 | PPFIA1 | rs693315 | 1.10E-07 | ##### | Brain - Putamen (basal gangli | 70324575 | 69824575 | 70824575 | Yes |
| ENSG00000131626.16 | PPFIA1 | rs531498 | 1.10E-07 | ##### | Brain - Putamen (basal gangli | 70324575 | 69824575 | 70824575 | Yes |
| ENSG00000131626.16 | PPFIA1 | rs524333 | 1.10E-07 | ##### | Brain - Putamen (basal gangli | 70324575 | 69824575 | 70824575 | Yes |

| | | | | | | | | | |
|---|---|---|---|---|---|---|---|---|---|
| ENSG00000131626.16 | PPFIA1 | rs559176 | 1.10E-07 | ##### | Brain - Putamen (basal gangli | 70324575 | 69824575 | 70824575 | Yes |
| ENSG00000131626.16 | PPFIA1 | rs625294 | 1.10E-07 | ##### | Brain - Putamen (basal gangli | 70324575 | 69824575 | 70824575 | Yes |
| ENSG00000131626.16 | PPFIA1 | rs625294 | 1.20E-07 | ##### | Brain - Caudate (basal ganglia | 70324575 | 69824575 | 70824575 | Yes |
| ENSG00000131626.16 | PPFIA1 | rs12283131 | 1.20E-07 | ##### | Brain - Caudate (basal ganglia | 70324575 | 69824575 | 70824575 | Yes |
| ENSG00000131626.16 | PPFIA1 | rs571554 | 1.30E-07 | ##### | Brain - Caudate (basal ganglia | 70324575 | 69824575 | 70824575 | Yes |
| ENSG00000131626.16 | PPFIA1 | rs7942766 | 1.30E-07 | ##### | Brain - Hippocampus | 70324575 | 69824575 | 70824575 | Yes |
| ENSG00000131626.16 | PPFIA1 | rs576588 | 1.30E-07 | ##### | Brain - Hippocampus | 70324575 | 69824575 | 70824575 | Yes |
| ENSG00000131626.16 | PPFIA1 | rs665890 | 1.40E-07 | ##### | Brain - Spinal cord (cervical c- | 70324575 | 69824575 | 70824575 | Yes |
| ENSG00000131626.16 | PPFIA1 | rs693315 | 1.60E-07 | ##### | Brain - Nucleus accumbens (b | 70324575 | 69824575 | 70824575 | Yes |
| ENSG00000131626.16 | PPFIA1 | rs531498 | 1.60E-07 | ##### | Brain - Nucleus accumbens (b | 70324575 | 69824575 | 70824575 | Yes |
| ENSG00000131626.16 | PPFIA1 | rs524333 | 1.60E-07 | ##### | Brain - Nucleus accumbens (b | 70324575 | 69824575 | 70824575 | Yes |
| ENSG00000131626.16 | PPFIA1 | rs559176 | 1.60E-07 | ##### | Brain - Nucleus accumbens (b | 70324575 | 69824575 | 70824575 | Yes |
| ENSG00000131626.16 | PPFIA1 | rs569732 | 1.60E-07 | ##### | Brain - Cortex | 70324575 | 69824575 | 70824575 | Yes |
| ENSG00000131626.16 | PPFIA1 | rs513487 | 1.60E-07 | ##### | Brain - Cortex | 70324575 | 69824575 | 70824575 | Yes |
| ENSG00000131626.16 | PPFIA1 | rs679455 | 1.60E-07 | ##### | Brain - Cortex | 70324575 | 69824575 | 70824575 | Yes |
| ENSG00000131626.16 | PPFIA1 | rs676889 | 1.60E-07 | ##### | Brain - Cortex | 70324575 | 69824575 | 70824575 | Yes |
| ENSG00000131626.16 | PPFIA1 | rs11236275 | 1.60E-07 | ##### | Brain - Cortex | 70324575 | 69824575 | 70824575 | Yes |
| ENSG00000131626.16 | PPFIA1 | rs3133177 | 1.60E-07 | ##### | Brain - Spinal cord (cervical c- | 70324575 | 69824575 | 70824575 | Yes |
| ENSG00000131626.16 | PPFIA1 | rs621749 | 1.60E-07 | ##### | Brain - Spinal cord (cervical c- | 70324575 | 69824575 | 70824575 | Yes |
| ENSG00000131626.16 | PPFIA1 | rs11235935 | 1.60E-07 | ##### | Brain - Spinal cord (cervical c- | 70324575 | 69824575 | 70824575 | Yes |
| ENSG00000131626.16 | PPFIA1 | rs7117443 | 1.60E-07 | ##### | Brain - Spinal cord (cervical c- | 70324575 | 69824575 | 70824575 | Yes |
| ENSG00000131626.16 | PPFIA1 | rs4980597 | 1.60E-07 | ##### | Brain - Spinal cord (cervical c- | 70324575 | 69824575 | 70824575 | Yes |
| ENSG00000131626.16 | PPFIA1 | rs531917 | 1.60E-07 | ##### | Brain - Spinal cord (cervical c- | 70324575 | 69824575 | 70824575 | Yes |
| ENSG00000131626.16 | PPFIA1 | rs2282481 | 1.60E-07 | ##### | Brain - Amygdala | 70324575 | 69824575 | 70824575 | Yes |
| ENSG00000131626.16 | PPFIA1 | rs665890 | 1.70E-07 | ##### | Brain - Hippocampus | 70324575 | 69824575 | 70824575 | Yes |
| ENSG00000131626.16 | PPFIA1 | rs12283131 | 1.70E-07 | ##### | Brain - Putamen (basal gangli | 70324575 | 69824575 | 70824575 | Yes |
| ENSG00000131626.16 | PPFIA1 | rs7117443 | 1.70E-07 | ##### | Brain - Putamen (basal gangli | 70324575 | 69824575 | 70824575 | Yes |
| ENSG00000131626.16 | PPFIA1 | rs670012 | 1.70E-07 | ##### | Brain - Frontal Cortex (BA9) | 70324575 | 69824575 | 70824575 | Yes |
| ENSG00000131626.16 | PPFIA1 | rs4980779 | 1.70E-07 | ##### | Brain - Frontal Cortex (BA9) | 70324575 | 69824575 | 70824575 | Yes |
| ENSG00000131626.16 | PPFIA1 | rs7130178 | 1.70E-07 | ##### | Brain - Frontal Cortex (BA9) | 70324575 | 69824575 | 70824575 | Yes |
| ENSG00000131626.16 | PPFIA1 | rs12283131 | 1.70E-07 | ##### | Brain - Nucleus accumbens (b | 70324575 | 69824575 | 70824575 | Yes |
| ENSG00000131626.16 | PPFIA1 | rs488746 | 1.80E-07 | ##### | Brain - Hippocampus | 70324575 | 69824575 | 70824575 | Yes |

| | | | | | | | | | |
|---|---|---|---|---|---|---|---|---|---|
| ENSG00000131626.16 | PPFIA1 | rs546724 | 1.80E-07 | ##### | Brain - Hippocampus | 70324575 | 69824575 | 70824575 | Yes |
| ENSG00000131626.16 | PPFIA1 | rs61887401 | 1.80E-07 | ##### | Brain - Hippocampus | 70324575 | 69824575 | 70824575 | Yes |
| ENSG00000131626.16 | PPFIA1 | rs113180277 | 1.80E-07 | ##### | Brain - Hippocampus | 70324575 | 69824575 | 70824575 | Yes |
| ENSG00000131626.16 | PPFIA1 | rs670310 | 1.80E-07 | ##### | Brain - Hippocampus | 70324575 | 69824575 | 70824575 | Yes |
| ENSG00000131626.16 | PPFIA1 | rs12283131 | 1.80E-07 | ##### | Brain - Amygdala | 70324575 | 69824575 | 70824575 | Yes |
| ENSG00000131626.16 | PPFIA1 | rs7942766 | 1.90E-07 | ##### | Brain - Caudate (basal ganglia | 70324575 | 69824575 | 70824575 | Yes |
| ENSG00000131626.16 | PPFIA1 | rs561050 | 2.00E-07 | ##### | Brain - Hippocampus | 70324575 | 69824575 | 70824575 | Yes |
| ENSG00000131626.16 | PPFIA1 | rs11235935 | 2.10E-07 | ##### | Brain - Putamen (basal gangli | 70324575 | 69824575 | 70824575 | Yes |
| ENSG00000131626.16 | PPFIA1 | rs4980597 | 2.10E-07 | ##### | Brain - Putamen (basal gangli | 70324575 | 69824575 | 70824575 | Yes |
| ENSG00000131626.16 | PPFIA1 | rs531917 | 2.10E-07 | ##### | Brain - Putamen (basal gangli | 70324575 | 69824575 | 70824575 | Yes |
| ENSG00000131626.16 | PPFIA1 | rs11356872 | 2.10E-07 | ##### | Brain - Putamen (basal gangli | 70324575 | 69824575 | 70824575 | Yes |
| ENSG00000131626.16 | PPFIA1 | rs3133177 | 2.20E-07 | ##### | Brain - Hippocampus | 70324575 | 69824575 | 70824575 | Yes |
| ENSG00000131626.16 | PPFIA1 | rs621749 | 2.20E-07 | ##### | Brain - Hippocampus | 70324575 | 69824575 | 70824575 | Yes |
| ENSG00000131626.16 | PPFIA1 | rs36037711 | 2.20E-07 | ##### | Brain - Frontal Cortex (BA9) | 70324575 | 69824575 | 70824575 | Yes |
| ENSG00000131626.16 | PPFIA1 | rs7117443 | 2.20E-07 | ##### | Brain - Frontal Cortex (BA9) | 70324575 | 69824575 | 70824575 | Yes |
| ENSG00000131626.16 | PPFIA1 | rs1788974 | 2.30E-07 | ##### | Brain - Frontal Cortex (BA9) | 70324575 | 69824575 | 70824575 | Yes |
| ENSG00000131626.16 | PPFIA1 | rs12295241 | 2.30E-07 | ##### | Brain - Caudate (basal ganglia | 70324575 | 69824575 | 70824575 | Yes |
| ENSG00000131626.16 | PPFIA1 | rs7109643 | 2.30E-07 | ##### | Brain - Caudate (basal ganglia | 70324575 | 69824575 | 70824575 | Yes |
| ENSG00000131626.16 | PPFIA1 | rs36037711 | 2.40E-07 | ##### | Brain - Nucleus accumbens (b | 70324575 | 69824575 | 70824575 | Yes |
| ENSG00000131626.16 | PPFIA1 | rs77395560 | 2.40E-07 | ##### | Brain - Cerebellum | 70324575 | 69824575 | 70824575 | Yes |
| ENSG00000131626.16 | PPFIA1 | rs571554 | 2.50E-07 | ##### | Brain - Frontal Cortex (BA9) | 70324575 | 69824575 | 70824575 | Yes |
| ENSG00000131626.16 | PPFIA1 | rs6592546 | 2.50E-07 | ##### | Brain - Amygdala | 70324575 | 69824575 | 70824575 | Yes |
| ENSG00000131626.16 | PPFIA1 | rs625294 | 2.60E-07 | ##### | Brain - Frontal Cortex (BA9) | 70324575 | 69824575 | 70824575 | Yes |
| ENSG00000131626.16 | PPFIA1 | rs12295241 | 2.60E-07 | ##### | Brain - Frontal Cortex (BA9) | 70324575 | 69824575 | 70824575 | Yes |
| ENSG00000131626.16 | PPFIA1 | rs7109643 | 2.60E-07 | ##### | Brain - Frontal Cortex (BA9) | 70324575 | 69824575 | 70824575 | Yes |
| ENSG00000131626.16 | PPFIA1 | rs665890 | 2.60E-07 | ##### | Brain - Caudate (basal ganglia | 70324575 | 69824575 | 70824575 | Yes |
| ENSG00000131626.16 | PPFIA1 | rs576588 | 2.60E-07 | ##### | Brain - Caudate (basal ganglia | 70324575 | 69824575 | 70824575 | Yes |
| ENSG00000131626.16 | PPFIA1 | rs4980782 | 2.70E-07 | ##### | Brain - Hippocampus | 70324575 | 69824575 | 70824575 | Yes |
| ENSG00000131626.16 | PPFIA1 | rs658590 | 2.80E-07 | ##### | Brain - Putamen (basal gangli | 70324575 | 69824575 | 70824575 | Yes |
| ENSG00000131626.16 | PPFIA1 | rs1788974 | 2.80E-07 | ##### | Brain - Nucleus accumbens (b | 70324575 | 69824575 | 70824575 | Yes |
| ENSG00000131626.16 | PPFIA1 | rs11356872 | 2.80E-07 | ##### | Brain - Spinal cord (cervical c- | 70324575 | 69824575 | 70824575 | Yes |
| ENSG00000131626.16 | PPFIA1 | rs488746 | 3.00E-07 | ##### | Brain - Caudate (basal ganglia | 70324575 | 69824575 | 70824575 | Yes |

| | | | | | | | | | |
|---|---|---|---|---|---|---|---|---|---|
| ENSG00000131626.16 | PPFIA1 | rs546724 | 3.00E-07 | ##### | Brain - Caudate (basal ganglia | 70324575 | 69824575 | 70824575 | Yes |
| ENSG00000131626.16 | PPFIA1 | rs61887401 | 3.00E-07 | ##### | Brain - Caudate (basal ganglia | 70324575 | 69824575 | 70824575 | Yes |
| ENSG00000131626.16 | PPFIA1 | rs113180277 | 3.00E-07 | ##### | Brain - Caudate (basal ganglia | 70324575 | 69824575 | 70824575 | Yes |
| ENSG00000131626.16 | PPFIA1 | rs670310 | 3.00E-07 | ##### | Brain - Caudate (basal ganglia | 70324575 | 69824575 | 70824575 | Yes |
| ENSG00000131626.16 | PPFIA1 | rs583347 | 3.00E-07 | ##### | Brain - Spinal cord (cervical c- | 70324575 | 69824575 | 70824575 | Yes |
| ENSG00000131626.16 | PPFIA1 | rs552282 | 3.00E-07 | ##### | Brain - Spinal cord (cervical c- | 70324575 | 69824575 | 70824575 | Yes |
| ENSG00000131626.16 | PPFIA1 | rs488746 | 3.10E-07 | ##### | Brain - Spinal cord (cervical c- | 70324575 | 69824575 | 70824575 | Yes |
| ENSG00000131626.16 | PPFIA1 | rs546724 | 3.10E-07 | ##### | Brain - Spinal cord (cervical c- | 70324575 | 69824575 | 70824575 | Yes |
| ENSG00000131626.16 | PPFIA1 | rs61887401 | 3.10E-07 | ##### | Brain - Spinal cord (cervical c- | 70324575 | 69824575 | 70824575 | Yes |
| ENSG00000131626.16 | PPFIA1 | rs113180277 | 3.10E-07 | ##### | Brain - Spinal cord (cervical c- | 70324575 | 69824575 | 70824575 | Yes |
| ENSG00000131626.16 | PPFIA1 | rs693315 | 3.10E-07 | ##### | Brain - Spinal cord (cervical c- | 70324575 | 69824575 | 70824575 | Yes |
| ENSG00000131626.16 | PPFIA1 | rs531498 | 3.10E-07 | ##### | Brain - Spinal cord (cervical c- | 70324575 | 69824575 | 70824575 | Yes |
| ENSG00000131626.16 | PPFIA1 | rs670310 | 3.10E-07 | ##### | Brain - Spinal cord (cervical c- | 70324575 | 69824575 | 70824575 | Yes |
| ENSG00000131626.16 | PPFIA1 | rs524333 | 3.10E-07 | ##### | Brain - Spinal cord (cervical c- | 70324575 | 69824575 | 70824575 | Yes |
| ENSG00000131626.16 | PPFIA1 | rs559176 | 3.10E-07 | ##### | Brain - Spinal cord (cervical c- | 70324575 | 69824575 | 70824575 | Yes |
| ENSG00000131626.16 | PPFIA1 | rs7942766 | 3.10E-07 | ##### | Brain - Spinal cord (cervical c- | 70324575 | 69824575 | 70824575 | Yes |
| ENSG00000131626.16 | PPFIA1 | rs1788973 | 3.20E-07 | ##### | Brain - Amygdala | 70324575 | 69824575 | 70824575 | Yes |
| ENSG00000131626.16 | PPFIA1 | rs10683183 | 3.20E-07 | ##### | Brain - Amygdala | 70324575 | 69824575 | 70824575 | Yes |
| ENSG00000131626.16 | PPFIA1 | rs6592542 | 3.20E-07 | ##### | Brain - Amygdala | 70324575 | 69824575 | 70824575 | Yes |
| ENSG00000131626.16 | PPFIA1 | rs693315 | 3.20E-07 | ##### | Brain - Hippocampus | 70324575 | 69824575 | 70824575 | Yes |
| ENSG00000131626.16 | PPFIA1 | rs531498 | 3.20E-07 | ##### | Brain - Hippocampus | 70324575 | 69824575 | 70824575 | Yes |
| ENSG00000131626.16 | PPFIA1 | rs524333 | 3.20E-07 | ##### | Brain - Hippocampus | 70324575 | 69824575 | 70824575 | Yes |
| ENSG00000131626.16 | PPFIA1 | rs559176 | 3.20E-07 | ##### | Brain - Hippocampus | 70324575 | 69824575 | 70824575 | Yes |
| ENSG00000131626.16 | PPFIA1 | rs687660 | 3.30E-07 | ##### | Brain - Putamen (basal gangli | 70324575 | 69824575 | 70824575 | Yes |
| ENSG00000131626.16 | PPFIA1 | rs11236048 | 3.30E-07 | ##### | Brain - Amygdala | 70324575 | 69824575 | 70824575 | Yes |
| ENSG00000131626.16 | PPFIA1 | rs658590 | 3.30E-07 | ##### | Brain - Frontal Cortex (BA9) | 70324575 | 69824575 | 70824575 | Yes |
| ENSG00000131626.16 | PPFIA1 | rs3133177 | 3.50E-07 | ##### | Brain - Caudate (basal ganglia | 70324575 | 69824575 | 70824575 | Yes |
| ENSG00000131626.16 | PPFIA1 | rs621749 | 3.50E-07 | ##### | Brain - Caudate (basal ganglia | 70324575 | 69824575 | 70824575 | Yes |
| ENSG00000131626.16 | PPFIA1 | rs625294 | 3.60E-07 | ##### | Brain - Hippocampus | 70324575 | 69824575 | 70824575 | Yes |
| ENSG00000131626.16 | PPFIA1 | rs687660 | 3.70E-07 | ##### | Brain - Frontal Cortex (BA9) | 70324575 | 69824575 | 70824575 | Yes |
| ENSG00000131626.16 | PPFIA1 | rs583397 | 3.90E-07 | ##### | Brain - Hippocampus | 70324575 | 69824575 | 70824575 | Yes |
| ENSG00000131626.16 | PPFIA1 | rs11235935 | 3.90E-07 | ##### | Brain - Hippocampus | 70324575 | 69824575 | 70824575 | Yes |

| | | | | | | | | | |
|---|---|---|---|---|---|---|---|---|---|
| ENSG00000131626.16 | PPFIA1 | rs4980597 | 3.90E-07 | ##### | Brain - Hippocampus | 70324575 | 69824575 | 70824575 | Yes |
| ENSG00000131626.16 | PPFIA1 | rs531917 | 3.90E-07 | ##### | Brain - Hippocampus | 70324575 | 69824575 | 70824575 | Yes |
| ENSG00000131626.16 | PPFIA1 | rs7948535 | 4.00E-07 | ##### | Brain - Cerebellar Hemisphere | 70324575 | 69824575 | 70824575 | Yes |
| ENSG00000131626.16 | PPFIA1 | rs687660 | 4.10E-07 | ##### | Brain - Spinal cord (cervical c- | 70324575 | 69824575 | 70824575 | Yes |
| ENSG00000131626.16 | PPFIA1 | rs658590 | 4.10E-07 | ##### | Brain - Spinal cord (cervical c- | 70324575 | 69824575 | 70824575 | Yes |
| ENSG00000131626.16 | PPFIA1 | rs3781649 | 4.40E-07 | ##### | Brain - Spinal cord (cervical c- | 70324575 | 69824575 | 70824575 | Yes |
| ENSG00000131626.16 | PPFIA1 | rs3781646 | 4.40E-07 | ##### | Brain - Spinal cord (cervical c- | 70324575 | 69824575 | 70824575 | Yes |
| ENSG00000131626.16 | PPFIA1 | rs151079983 | 4.40E-07 | ##### | Brain - Spinal cord (cervical c- | 70324575 | 69824575 | 70824575 | Yes |
| ENSG00000131626.16 | PPFIA1 | rs625294 | 4.40E-07 | ##### | Brain - Nucleus accumbens (b | 70324575 | 69824575 | 70824575 | Yes |
| ENSG00000131626.16 | PPFIA1 | rs644396 | 4.50E-07 | ##### | Brain - Cortex | 70324575 | 69824575 | 70824575 | Yes |
| ENSG00000131626.16 | PPFIA1 | rs12295241 | 4.50E-07 | ##### | Brain - Putamen (basal gangli | 70324575 | 69824575 | 70824575 | Yes |
| ENSG00000131626.16 | PPFIA1 | rs7109643 | 4.50E-07 | ##### | Brain - Putamen (basal gangli | 70324575 | 69824575 | 70824575 | Yes |
| ENSG00000131626.16 | PPFIA1 | rs4600242 | 4.80E-07 | ##### | Brain - Cerebellum | 70324575 | 69824575 | 70824575 | Yes |
| ENSG00000131626.16 | PPFIA1 | rs4604917 | 4.80E-07 | ##### | Brain - Cerebellum | 70324575 | 69824575 | 70824575 | Yes |
| ENSG00000131626.16 | PPFIA1 | rs4526792 | 4.80E-07 | ##### | Brain - Cerebellum | 70324575 | 69824575 | 70824575 | Yes |
| ENSG00000131626.16 | PPFIA1 | rs625294 | 5.10E-07 | ##### | Brain - Spinal cord (cervical c- | 70324575 | 69824575 | 70824575 | Yes |
| ENSG00000131626.16 | PPFIA1 | rs12295241 | 5.10E-07 | ##### | Brain - Spinal cord (cervical c- | 70324575 | 69824575 | 70824575 | Yes |
| ENSG00000131626.16 | PPFIA1 | rs7109643 | 5.10E-07 | ##### | Brain - Spinal cord (cervical c- | 70324575 | 69824575 | 70824575 | Yes |
| ENSG00000131626.16 | PPFIA1 | rs3132438 | 5.40E-07 | ##### | Brain - Amygdala | 70324575 | 69824575 | 70824575 | Yes |
| ENSG00000131626.16 | PPFIA1 | rs34141396 | 5.40E-07 | ##### | Brain - Amygdala | 70324575 | 69824575 | 70824575 | Yes |
| ENSG00000131626.16 | PPFIA1 | rs3132421 | 5.40E-07 | ##### | Brain - Amygdala | 70324575 | 69824575 | 70824575 | Yes |
| ENSG00000131626.16 | PPFIA1 | rs142385955 | 5.50E-07 | ##### | Brain - Spinal cord (cervical c- | 70324575 | 69824575 | 70824575 | Yes |
| ENSG00000131626.16 | PPFIA1 | rs1198234 | 5.50E-07 | ##### | Brain - Spinal cord (cervical c- | 70324575 | 69824575 | 70824575 | Yes |
| ENSG00000131626.16 | PPFIA1 | rs611216 | 5.50E-07 | ##### | Brain - Spinal cord (cervical c- | 70324575 | 69824575 | 70824575 | Yes |
| ENSG00000131626.16 | PPFIA1 | rs482438 | 5.50E-07 | ##### | Brain - Spinal cord (cervical c- | 70324575 | 69824575 | 70824575 | Yes |
| ENSG00000131626.16 | PPFIA1 | rs643301 | 5.50E-07 | ##### | Brain - Spinal cord (cervical c- | 70324575 | 69824575 | 70824575 | Yes |
| ENSG00000131626.16 | PPFIA1 | rs3802780 | 5.50E-07 | ##### | Brain - Spinal cord (cervical c- | 70324575 | 69824575 | 70824575 | Yes |
| ENSG00000131626.16 | PPFIA1 | rs1788973 | 5.70E-07 | ##### | Brain - Spinal cord (cervical c- | 70324575 | 69824575 | 70824575 | Yes |
| ENSG00000131626.16 | PPFIA1 | rs10683183 | 5.70E-07 | ##### | Brain - Spinal cord (cervical c- | 70324575 | 69824575 | 70824575 | Yes |
| ENSG00000131626.16 | PPFIA1 | rs6592542 | 5.70E-07 | ##### | Brain - Spinal cord (cervical c- | 70324575 | 69824575 | 70824575 | Yes |
| ENSG00000131626.16 | PPFIA1 | rs583347 | 5.70E-07 | ##### | Brain - Amygdala | 70324575 | 69824575 | 70824575 | Yes |
| ENSG00000131626.16 | PPFIA1 | rs552282 | 5.70E-07 | ##### | Brain - Amygdala | 70324575 | 69824575 | 70824575 | Yes |

| | | | | | | | | | |
|---|---|---|---|---|---|---|---|---|---|
| ENSG00000131626.16 | PPFIA1 | rs674374 | 5.80E-07 | ##### | Brain - Amygdala | 70324575 | 69824575 | 70824575 | Yes |
| ENSG00000131626.16 | PPFIA1 | rs693315 | 5.80E-07 | ##### | Brain - Caudate (basal ganglia | 70324575 | 69824575 | 70824575 | Yes |
| ENSG00000131626.16 | PPFIA1 | rs531498 | 5.80E-07 | ##### | Brain - Caudate (basal ganglia | 70324575 | 69824575 | 70824575 | Yes |
| ENSG00000131626.16 | PPFIA1 | rs524333 | 5.80E-07 | ##### | Brain - Caudate (basal ganglia | 70324575 | 69824575 | 70824575 | Yes |
| ENSG00000131626.16 | PPFIA1 | rs559176 | 5.80E-07 | ##### | Brain - Caudate (basal ganglia | 70324575 | 69824575 | 70824575 | Yes |
| ENSG00000131626.16 | PPFIA1 | rs670012 | 6.20E-07 | ##### | Brain - Hippocampus | 70324575 | 69824575 | 70824575 | Yes |
| ENSG00000131626.16 | PPFIA1 | rs4980779 | 6.20E-07 | ##### | Brain - Hippocampus | 70324575 | 69824575 | 70824575 | Yes |
| ENSG00000131626.16 | PPFIA1 | rs7130178 | 6.20E-07 | ##### | Brain - Hippocampus | 70324575 | 69824575 | 70824575 | Yes |
| ENSG00000131626.16 | PPFIA1 | rs638510 | 6.30E-07 | ##### | Brain - Putamen (basal gangli | 70324575 | 69824575 | 70824575 | Yes |
| ENSG00000131626.16 | PPFIA1 | rs3132438 | 6.30E-07 | ##### | Brain - Frontal Cortex (BA9) | 70324575 | 69824575 | 70824575 | Yes |
| ENSG00000131626.16 | PPFIA1 | rs35354974 | 6.40E-07 | ##### | Brain - Cerebellum | 70324575 | 69824575 | 70824575 | Yes |
| ENSG00000131626.16 | PPFIA1 | rs10899069 | 6.40E-07 | ##### | Brain - Cerebellum | 70324575 | 69824575 | 70824575 | Yes |
| ENSG00000131626.16 | PPFIA1 | rs592501 | 6.40E-07 | ##### | Brain - Spinal cord (cervical c- | 70324575 | 69824575 | 70824575 | Yes |
| ENSG00000131626.16 | PPFIA1 | rs548687 | 6.40E-07 | ##### | Brain - Spinal cord (cervical c- | 70324575 | 69824575 | 70824575 | Yes |
| ENSG00000131626.16 | PPFIA1 | rs1198236 | 6.40E-07 | ##### | Brain - Spinal cord (cervical c- | 70324575 | 69824575 | 70824575 | Yes |
| ENSG00000131626.16 | PPFIA1 | rs12295241 | 6.60E-07 | ##### | Brain - Nucleus accumbens (b | 70324575 | 69824575 | 70824575 | Yes |
| ENSG00000131626.16 | PPFIA1 | rs7109643 | 6.60E-07 | ##### | Brain - Nucleus accumbens (b | 70324575 | 69824575 | 70824575 | Yes |
| ENSG00000131626.16 | PPFIA1 | rs11356872 | 6.60E-07 | ##### | Brain - Frontal Cortex (BA9) | 70324575 | 69824575 | 70824575 | Yes |
| ENSG00000131626.16 | PPFIA1 | rs4980782 | 6.80E-07 | ##### | Brain - Frontal Cortex (BA9) | 70324575 | 69824575 | 70824575 | Yes |
| ENSG00000131626.16 | PPFIA1 | rs3133177 | 6.80E-07 | ##### | Brain - Nucleus accumbens (b | 70324575 | 69824575 | 70824575 | Yes |
| ENSG00000131626.16 | PPFIA1 | rs621749 | 6.80E-07 | ##### | Brain - Nucleus accumbens (b | 70324575 | 69824575 | 70824575 | Yes |
| ENSG00000131626.16 | PPFIA1 | rs7117443 | 6.80E-07 | ##### | Brain - Hippocampus | 70324575 | 69824575 | 70824575 | Yes |
| ENSG00000131626.16 | PPFIA1 | rs670141 | 6.90E-07 | ##### | Brain - Cortex | 70324575 | 69824575 | 70824575 | Yes |
| ENSG00000131626.16 | PPFIA1 | rs11235935 | 6.90E-07 | ##### | Brain - Caudate (basal ganglia | 70324575 | 69824575 | 70824575 | Yes |
| ENSG00000131626.16 | PPFIA1 | rs4980597 | 6.90E-07 | ##### | Brain - Caudate (basal ganglia | 70324575 | 69824575 | 70824575 | Yes |
| ENSG00000131626.16 | PPFIA1 | rs531917 | 6.90E-07 | ##### | Brain - Caudate (basal ganglia | 70324575 | 69824575 | 70824575 | Yes |
| ENSG00000131626.16 | PPFIA1 | rs561050 | 7.40E-07 | ##### | Brain - Nucleus accumbens (b | 70324575 | 69824575 | 70824575 | Yes |
| ENSG00000131626.16 | PPFIA1 | rs665890 | 7.40E-07 | ##### | Brain - Nucleus accumbens (b | 70324575 | 69824575 | 70824575 | Yes |
| ENSG00000131626.16 | PPFIA1 | rs36037711 | 7.80E-07 | ##### | Brain - Spinal cord (cervical c- | 70324575 | 69824575 | 70824575 | Yes |
| ENSG00000131626.16 | PPFIA1 | rs1788974 | 7.90E-07 | ##### | Brain - Spinal cord (cervical c- | 70324575 | 69824575 | 70824575 | Yes |
| ENSG00000131626.16 | PPFIA1 | rs12283131 | 7.90E-07 | ##### | Brain - Frontal Cortex (BA9) | 70324575 | 69824575 | 70824575 | Yes |
| ENSG00000131626.16 | PPFIA1 | rs4980782 | 7.90E-07 | ##### | Brain - Amygdala | 70324575 | 69824575 | 70824575 | Yes |

| | | | | | | | | | |
|---|---|---|---|---|---|---|---|---|---|
| ENSG00000131626.16 | PPFIA1 | rs36037711 | 8.20E-07 | ##### | Brain - Caudate (basal ganglia | 70324575 | 69824575 | 70824575 | Yes |
| ENSG00000131626.16 | PPFIA1 | rs2282481 | 8.30E-07 | ##### | Brain - Frontal Cortex (BA9) | 70324575 | 69824575 | 70824575 | Yes |
| ENSG00000131626.16 | PPFIA1 | rs11356872 | 8.50E-07 | ##### | Brain - Hippocampus | 70324575 | 69824575 | 70824575 | Yes |
| ENSG00000131626.16 | PPFIA1 | rs1198236 | 8.60E-07 | ##### | Brain - Hippocampus | 70324575 | 69824575 | 70824575 | Yes |
| ENSG00000131626.16 | PPFIA1 | rs482438 | 8.60E-07 | ##### | Brain - Hippocampus | 70324575 | 69824575 | 70824575 | Yes |
| ENSG00000131626.16 | PPFIA1 | rs643301 | 8.60E-07 | ##### | Brain - Hippocampus | 70324575 | 69824575 | 70824575 | Yes |
| ENSG00000131626.16 | PPFIA1 | rs3802780 | 8.60E-07 | ##### | Brain - Hippocampus | 70324575 | 69824575 | 70824575 | Yes |
| ENSG00000131626.16 | PPFIA1 | rs2353299 | 8.60E-07 | ##### | Brain - Cortex | 70324575 | 69824575 | 70824575 | Yes |
| ENSG00000131626.16 | PPFIA1 | rs499484 | 8.70E-07 | ##### | Brain - Amygdala | 70324575 | 69824575 | 70824575 | Yes |
| ENSG00000131626.16 | PPFIA1 | rs2282481 | 8.70E-07 | ##### | Brain - Spinal cord (cervical c- | 70324575 | 69824575 | 70824575 | Yes |
| ENSG00000131626.16 | PPFIA1 | rs662018 | 8.80E-07 | ##### | Brain - Spinal cord (cervical c- | 70324575 | 69824575 | 70824575 | Yes |
| ENSG00000131626.16 | PPFIA1 | rs1788974 | 8.90E-07 | ##### | Brain - Caudate (basal ganglia | 70324575 | 69824575 | 70824575 | Yes |
| ENSG00000131626.16 | PPFIA1 | rs670012 | 9.30E-07 | ##### | Brain - Spinal cord (cervical c- | 70324575 | 69824575 | 70824575 | Yes |
| ENSG00000131626.16 | PPFIA1 | rs4980779 | 9.30E-07 | ##### | Brain - Spinal cord (cervical c- | 70324575 | 69824575 | 70824575 | Yes |
| ENSG00000131626.16 | PPFIA1 | rs7130178 | 9.30E-07 | ##### | Brain - Spinal cord (cervical c- | 70324575 | 69824575 | 70824575 | Yes |
| ENSG00000131626.16 | PPFIA1 | rs569732 | 9.40E-07 | ##### | Brain - Amygdala | 70324575 | 69824575 | 70824575 | Yes |
| ENSG00000131626.16 | PPFIA1 | rs11235935 | 9.90E-07 | ##### | Brain - Nucleus accumbens (b | 70324575 | 69824575 | 70824575 | Yes |
| ENSG00000131626.16 | PPFIA1 | rs4980597 | 9.90E-07 | ##### | Brain - Nucleus accumbens (b | 70324575 | 69824575 | 70824575 | Yes |
| ENSG00000131626.16 | PPFIA1 | rs531917 | 9.90E-07 | ##### | Brain - Nucleus accumbens (b | 70324575 | 69824575 | 70824575 | Yes |
| ENSG00000131626.16 | PPFIA1 | rs638510 | 0.000001 | ##### | Brain - Amygdala | 70324575 | 69824575 | 70824575 | Yes |
| ENSG00000131626.16 | PPFIA1 | rs36037711 | 0.0000011 | ##### | Brain - Hippocampus | 70324575 | 69824575 | 70824575 | Yes |
| ENSG00000131626.16 | PPFIA1 | rs76294966 | 0.0000011 | ##### | Brain - Amygdala | 70324575 | 69824575 | 70824575 | Yes |
| ENSG00000131626.16 | PPFIA1 | rs4980780 | 0.0000011 | ##### | Brain - Amygdala | 70324575 | 69824575 | 70824575 | Yes |
| ENSG00000131626.16 | PPFIA1 | rs7129635 | 0.0000011 | ##### | Brain - Amygdala | 70324575 | 69824575 | 70824575 | Yes |
| ENSG00000131626.16 | PPFIA1 | rs594763 | 0.0000011 | ##### | Brain - Amygdala | 70324575 | 69824575 | 70824575 | Yes |
| ENSG00000131626.16 | PPFIA1 | rs571554 | 0.0000011 | ##### | Brain - Spinal cord (cervical c- | 70324575 | 69824575 | 70824575 | Yes |
| ENSG00000131626.16 | PPFIA1 | rs583397 | 0.0000011 | ##### | Brain - Frontal Cortex (BA9) | 70324575 | 69824575 | 70824575 | Yes |
| ENSG00000131626.16 | PPFIA1 | rs6592546 | 0.0000011 | ##### | Brain - Spinal cord (cervical c- | 70324575 | 69824575 | 70824575 | Yes |
| ENSG00000131626.16 | PPFIA1 | rs1788974 | 0.0000011 | ##### | Brain - Hippocampus | 70324575 | 69824575 | 70824575 | Yes |
| ENSG00000131626.16 | PPFIA1 | rs3781649 | 0.0000012 | ##### | Brain - Hippocampus | 70324575 | 69824575 | 70824575 | Yes |
| ENSG00000131626.16 | PPFIA1 | rs3781646 | 0.0000012 | ##### | Brain - Hippocampus | 70324575 | 69824575 | 70824575 | Yes |
| ENSG00000131626.16 | PPFIA1 | rs151079983 | 0.0000012 | ##### | Brain - Hippocampus | 70324575 | 69824575 | 70824575 | Yes |

| | | | | | | | | | |
|---|---|---|---|---|---|---|---|---|---|
| ENSG00000131626.16 | PPFIA1 | rs12295241 | 0.0000013 | ##### | Brain - Hippocampus | 70324575 | 69824575 | 70824575 | Yes |
| ENSG00000131626.16 | PPFIA1 | rs7109643 | 0.0000013 | ##### | Brain - Hippocampus | 70324575 | 69824575 | 70824575 | Yes |
| ENSG00000131626.16 | PPFIA1 | rs619650 | 0.0000013 | ##### | Brain - Amygdala | 70324575 | 69824575 | 70824575 | Yes |
| ENSG00000131626.16 | PPFIA1 | rs3132424 | 0.0000013 | ##### | Brain - Amygdala | 70324575 | 69824575 | 70824575 | Yes |
| ENSG00000131626.16 | PPFIA1 | rs687660 | 0.0000013 | ##### | Brain - Nucleus accumbens (b | 70324575 | 69824575 | 70824575 | Yes |
| ENSG00000131626.16 | PPFIA1 | rs7117443 | 0.0000014 | ##### | Brain - Caudate (basal ganglia | 70324575 | 69824575 | 70824575 | Yes |
| ENSG00000131626.16 | PPFIA1 | rs10591164 | 0.0000014 | ##### | Brain - Amygdala | 70324575 | 69824575 | 70824575 | Yes |
| ENSG00000131626.16 | PPFIA1 | rs35795372 | 0.0000014 | ##### | Brain - Amygdala | 70324575 | 69824575 | 70824575 | Yes |
| ENSG00000131626.16 | PPFIA1 | rs664130 | 0.0000014 | ##### | Brain - Amygdala | 70324575 | 69824575 | 70824575 | Yes |
| ENSG00000131626.16 | PPFIA1 | rs11236048 | 0.0000014 | ##### | Brain - Hippocampus | 70324575 | 69824575 | 70824575 | Yes |
| ENSG00000131626.16 | PPFIA1 | rs34141396 | 0.0000015 | ##### | Brain - Frontal Cortex (BA9) | 70324575 | 69824575 | 70824575 | Yes |
| ENSG00000131626.16 | PPFIA1 | rs76294966 | 0.0000015 | ##### | Brain - Frontal Cortex (BA9) | 70324575 | 69824575 | 70824575 | Yes |
| ENSG00000131626.16 | PPFIA1 | rs4980780 | 0.0000015 | ##### | Brain - Frontal Cortex (BA9) | 70324575 | 69824575 | 70824575 | Yes |
| ENSG00000131626.16 | PPFIA1 | rs7129635 | 0.0000015 | ##### | Brain - Frontal Cortex (BA9) | 70324575 | 69824575 | 70824575 | Yes |
| ENSG00000131626.16 | PPFIA1 | rs594763 | 0.0000015 | ##### | Brain - Frontal Cortex (BA9) | 70324575 | 69824575 | 70824575 | Yes |
| ENSG00000131626.16 | PPFIA1 | rs3132421 | 0.0000015 | ##### | Brain - Frontal Cortex (BA9) | 70324575 | 69824575 | 70824575 | Yes |
| ENSG00000131626.16 | PPFIA1 | rs531385401 | 0.0000015 | ##### | Brain - Nucleus accumbens (b | 70324575 | 69824575 | 70824575 | Yes |
| ENSG00000131626.16 | PPFIA1 | rs11236302 | 0.0000015 | ##### | Brain - Cerebellum | 70324575 | 69824575 | 70824575 | Yes |
| ENSG00000131626.16 | PPFIA1 | rs6592546 | 0.0000015 | ##### | Brain - Putamen (basal gangli | 70324575 | 69824575 | 70824575 | Yes |
| ENSG00000131626.16 | PPFIA1 | rs662018 | 0.0000016 | ##### | Brain - Hippocampus | 70324575 | 69824575 | 70824575 | Yes |
| ENSG00000131626.16 | PPFIA1 | rs670141 | 0.0000017 | ##### | Brain - Amygdala | 70324575 | 69824575 | 70824575 | Yes |
| ENSG00000131626.16 | PPFIA1 | rs674374 | 0.0000017 | ##### | Brain - Spinal cord (cervical c- | 70324575 | 69824575 | 70824575 | Yes |
| ENSG00000131626.16 | PPFIA1 | rs7117443 | 0.0000018 | ##### | Brain - Nucleus accumbens (b | 70324575 | 69824575 | 70824575 | Yes |
| ENSG00000131626.16 | PPFIA1 | rs561050 | 0.0000018 | ##### | Brain - Cortex | 70324575 | 69824575 | 70824575 | Yes |
| ENSG00000131626.16 | PPFIA1 | rs687660 | 0.0000018 | ##### | Brain - Caudate (basal ganglia | 70324575 | 69824575 | 70824575 | Yes |
| ENSG00000131626.16 | PPFIA1 | rs141709344 | 0.0000018 | ##### | Brain - Amygdala | 70324575 | 69824575 | 70824575 | Yes |
| ENSG00000131626.16 | PPFIA1 | rs662018 | 0.0000018 | ##### | Brain - Frontal Cortex (BA9) | 70324575 | 69824575 | 70824575 | Yes |
| ENSG00000131626.16 | PPFIA1 | rs76294966 | 0.0000019 | ##### | Brain - Hippocampus | 70324575 | 69824575 | 70824575 | Yes |
| ENSG00000131626.16 | PPFIA1 | rs4980780 | 0.0000019 | ##### | Brain - Hippocampus | 70324575 | 69824575 | 70824575 | Yes |
| ENSG00000131626.16 | PPFIA1 | rs7129635 | 0.0000019 | ##### | Brain - Hippocampus | 70324575 | 69824575 | 70824575 | Yes |
| ENSG00000131626.16 | PPFIA1 | rs594763 | 0.0000019 | ##### | Brain - Hippocampus | 70324575 | 69824575 | 70824575 | Yes |
| ENSG00000131626.16 | PPFIA1 | rs658590 | 0.0000019 | ##### | Brain - Caudate (basal ganglia | 70324575 | 69824575 | 70824575 | Yes |

| | | | | | | | | | |
|---|---|---|---|---|---|---|---|---|---|
| ENSG00000131626.16 | PPFIA1 | rs546502 | 0.0000019 | ##### | Brain - Frontal Cortex (BA9) | 70324575 | 69824575 | 70824575 | Yes |
| ENSG00000131626.16 | PPFIA1 | rs624765 | 0.0000019 | ##### | Brain - Frontal Cortex (BA9) | 70324575 | 69824575 | 70824575 | Yes |
| ENSG00000131626.16 | PPFIA1 | rs3892505 | 0.0000019 | ##### | Brain - Frontal Cortex (BA9) | 70324575 | 69824575 | 70824575 | Yes |
| ENSG00000131626.16 | PPFIA1 | rs35795372 | 0.0000019 | ##### | Brain - Frontal Cortex (BA9) | 70324575 | 69824575 | 70824575 | Yes |
| ENSG00000131626.16 | PPFIA1 | rs664130 | 0.0000019 | ##### | Brain - Frontal Cortex (BA9) | 70324575 | 69824575 | 70824575 | Yes |
| ENSG00000131626.16 | PPFIA1 | rs658590 | 0.000002 | ##### | Brain - Nucleus accumbens (b | 70324575 | 69824575 | 70824575 | Yes |
| ENSG00000131626.16 | PPFIA1 | rs12284490 | 0.000002 | ##### | Brain - Amygdala | 70324575 | 69824575 | 70824575 | Yes |
| ENSG00000131626.16 | PPFIA1 | rs499484 | 0.0000021 | ##### | Brain - Frontal Cortex (BA9) | 70324575 | 69824575 | 70824575 | Yes |
| ENSG00000131626.16 | PPFIA1 | rs1788973 | 0.0000021 | ##### | Brain - Frontal Cortex (BA9) | 70324575 | 69824575 | 70824575 | Yes |
| ENSG00000131626.16 | PPFIA1 | rs10683183 | 0.0000021 | ##### | Brain - Frontal Cortex (BA9) | 70324575 | 69824575 | 70824575 | Yes |
| ENSG00000131626.16 | PPFIA1 | rs6592542 | 0.0000021 | ##### | Brain - Frontal Cortex (BA9) | 70324575 | 69824575 | 70824575 | Yes |
| ENSG00000131626.16 | PPFIA1 | rs546502 | 0.0000022 | ##### | Brain - Hippocampus | 70324575 | 69824575 | 70824575 | Yes |
| ENSG00000131626.16 | PPFIA1 | rs624765 | 0.0000022 | ##### | Brain - Hippocampus | 70324575 | 69824575 | 70824575 | Yes |
| ENSG00000131626.16 | PPFIA1 | rs3892505 | 0.0000022 | ##### | Brain - Hippocampus | 70324575 | 69824575 | 70824575 | Yes |
| ENSG00000131626.16 | PPFIA1 | rs611216 | 0.0000022 | ##### | Brain - Hippocampus | 70324575 | 69824575 | 70824575 | Yes |
| ENSG00000131626.16 | PPFIA1 | rs548687 | 0.0000022 | ##### | Brain - Hippocampus | 70324575 | 69824575 | 70824575 | Yes |
| ENSG00000131626.16 | PPFIA1 | rs513487 | 0.0000022 | ##### | Brain - Amygdala | 70324575 | 69824575 | 70824575 | Yes |
| ENSG00000131626.16 | PPFIA1 | rs679455 | 0.0000022 | ##### | Brain - Amygdala | 70324575 | 69824575 | 70824575 | Yes |
| ENSG00000131626.16 | PPFIA1 | rs676889 | 0.0000022 | ##### | Brain - Amygdala | 70324575 | 69824575 | 70824575 | Yes |
| ENSG00000131626.16 | PPFIA1 | rs11236275 | 0.0000022 | ##### | Brain - Amygdala | 70324575 | 69824575 | 70824575 | Yes |
| ENSG00000131626.16 | PPFIA1 | rs658590 | 0.0000022 | ##### | Brain - Hippocampus | 70324575 | 69824575 | 70824575 | Yes |
| ENSG00000131626.16 | PPFIA1 | rs687660 | 0.0000023 | ##### | Brain - Hippocampus | 70324575 | 69824575 | 70824575 | Yes |
| ENSG00000131626.16 | PPFIA1 | rs11356872 | 0.0000024 | ##### | Brain - Caudate (basal ganglia | 70324575 | 69824575 | 70824575 | Yes |
| ENSG00000131626.16 | PPFIA1 | rs592501 | 0.0000024 | ##### | Brain - Hippocampus | 70324575 | 69824575 | 70824575 | Yes |
| ENSG00000131626.16 | PPFIA1 | rs571554 | 0.0000024 | ##### | Brain - Hippocampus | 70324575 | 69824575 | 70824575 | Yes |
| ENSG00000131626.16 | PPFIA1 | rs7948535 | 0.0000024 | ##### | Brain - Spinal cord (cervical c- | 70324575 | 69824575 | 70824575 | Yes |
| ENSG00000131626.16 | PPFIA1 | rs3132436 | 0.0000026 | ##### | Brain - Spinal cord (cervical c- | 70324575 | 69824575 | 70824575 | Yes |
| ENSG00000131626.16 | PPFIA1 | rs1788973 | 0.0000028 | ##### | Brain - Putamen (basal gangli | 70324575 | 69824575 | 70824575 | Yes |
| ENSG00000131626.16 | PPFIA1 | rs10683183 | 0.0000028 | ##### | Brain - Putamen (basal gangli | 70324575 | 69824575 | 70824575 | Yes |
| ENSG00000131626.16 | PPFIA1 | rs6592542 | 0.0000028 | ##### | Brain - Putamen (basal gangli | 70324575 | 69824575 | 70824575 | Yes |
| ENSG00000131626.16 | PPFIA1 | rs630280 | 0.0000028 | ##### | Brain - Cerebellum | 70324575 | 69824575 | 70824575 | Yes |
| ENSG00000131626.16 | PPFIA1 | rs3132438 | 0.0000028 | ##### | Brain - Hippocampus | 70324575 | 69824575 | 70824575 | Yes |

| | | | | | | | | | |
|---|---|---|---|---|---|---|---|---|---|
| ENSG00000131626.16 | PPFIA1 | rs6592546 | 0.0000029 | ##### | Brain - Frontal Cortex (BA9) | 70324575 | 69824575 | 70824575 | Yes |
| ENSG00000131626.16 | PPFIA1 | rs546502 | 0.0000029 | ##### | Brain - Amygdala | 70324575 | 69824575 | 70824575 | Yes |
| ENSG00000131626.16 | PPFIA1 | rs624765 | 0.0000029 | ##### | Brain - Amygdala | 70324575 | 69824575 | 70824575 | Yes |
| ENSG00000131626.16 | PPFIA1 | rs3892505 | 0.0000029 | ##### | Brain - Amygdala | 70324575 | 69824575 | 70824575 | Yes |
| ENSG00000131626.16 | PPFIA1 | rs4980782 | 0.0000029 | ##### | Brain - Cortex | 70324575 | 69824575 | 70824575 | Yes |
| ENSG00000131626.16 | PPFIA1 | rs3132436 | 0.0000029 | ##### | Brain - Hippocampus | 70324575 | 69824575 | 70824575 | Yes |
| ENSG00000131626.16 | PPFIA1 | rs570739 | 0.0000029 | ##### | Brain - Cerebellum | 70324575 | 69824575 | 70824575 | Yes |
| ENSG00000131626.16 | PPFIA1 | rs7943389 | 0.0000029 | ##### | Brain - Cerebellum | 70324575 | 69824575 | 70824575 | Yes |
| ENSG00000131626.16 | PPFIA1 | rs583347 | 0.0000029 | ##### | Brain - Frontal Cortex (BA9) | 70324575 | 69824575 | 70824575 | Yes |
| ENSG00000131626.16 | PPFIA1 | rs552282 | 0.0000029 | ##### | Brain - Frontal Cortex (BA9) | 70324575 | 69824575 | 70824575 | Yes |
| ENSG00000131626.16 | PPFIA1 | rs34141396 | 0.0000031 | ##### | Brain - Hippocampus | 70324575 | 69824575 | 70824575 | Yes |
| ENSG00000131626.16 | PPFIA1 | rs3132421 | 0.0000031 | ##### | Brain - Hippocampus | 70324575 | 69824575 | 70824575 | Yes |
| ENSG00000131626.16 | PPFIA1 | rs3781649 | 0.0000031 | ##### | Brain - Frontal Cortex (BA9) | 70324575 | 69824575 | 70824575 | Yes |
| ENSG00000131626.16 | PPFIA1 | rs3781646 | 0.0000031 | ##### | Brain - Frontal Cortex (BA9) | 70324575 | 69824575 | 70824575 | Yes |
| ENSG00000131626.16 | PPFIA1 | rs151079983 | 0.0000031 | ##### | Brain - Frontal Cortex (BA9) | 70324575 | 69824575 | 70824575 | Yes |
| ENSG00000131626.16 | PPFIA1 | rs2353299 | 0.0000035 | ##### | Brain - Amygdala | 70324575 | 69824575 | 70824575 | Yes |
| ENSG00000131626.16 | PPFIA1 | rs644396 | 0.0000035 | ##### | Brain - Amygdala | 70324575 | 69824575 | 70824575 | Yes |
| ENSG00000131626.16 | PPFIA1 | rs644396 | 0.0000035 | ##### | Brain - Putamen (basal gangli | 70324575 | 69824575 | 70824575 | Yes |
| ENSG00000131626.16 | PPFIA1 | rs2353299 | 0.0000035 | ##### | Brain - Putamen (basal gangli | 70324575 | 69824575 | 70824575 | Yes |
| ENSG00000131626.16 | PPFIA1 | rs10591164 | 0.0000036 | ##### | Brain - Hippocampus | 70324575 | 69824575 | 70824575 | Yes |
| ENSG00000131626.16 | PPFIA1 | rs35795372 | 0.0000036 | ##### | Brain - Hippocampus | 70324575 | 69824575 | 70824575 | Yes |
| ENSG00000131626.16 | PPFIA1 | rs664130 | 0.0000036 | ##### | Brain - Hippocampus | 70324575 | 69824575 | 70824575 | Yes |
| ENSG00000131626.16 | PPFIA1 | rs11236048 | 0.0000037 | ##### | Brain - Frontal Cortex (BA9) | 70324575 | 69824575 | 70824575 | Yes |
| ENSG00000131626.16 | PPFIA1 | rs670141 | 0.0000038 | ##### | Brain - Putamen (basal gangli | 70324575 | 69824575 | 70824575 | Yes |
| ENSG00000131626.16 | PPFIA1 | rs4603333 | 0.0000038 | ##### | Brain - Cerebellum | 70324575 | 69824575 | 70824575 | Yes |
| ENSG00000131626.16 | PPFIA1 | rs2282481 | 0.0000039 | ##### | Brain - Nucleus accumbens (b | 70324575 | 69824575 | 70824575 | Yes |
| ENSG00000131626.16 | PPFIA1 | rs627079 | 0.000004 | ##### | Brain - Cerebellum | 70324575 | 69824575 | 70824575 | Yes |
| ENSG00000131626.16 | PPFIA1 | rs141709344 | 0.0000043 | ##### | Brain - Frontal Cortex (BA9) | 70324575 | 69824575 | 70824575 | Yes |
| ENSG00000131626.16 | PPFIA1 | rs3132436 | 0.0000044 | ##### | Brain - Cortex | 70324575 | 69824575 | 70824575 | Yes |
| ENSG00000131626.16 | PPFIA1 | rs662018 | 0.0000045 | ##### | Brain - Amygdala | 70324575 | 69824575 | 70824575 | Yes |
| ENSG00000131626.16 | PPFIA1 | rs569732 | 0.0000045 | ##### | Brain - Putamen (basal gangli | 70324575 | 69824575 | 70824575 | Yes |
| ENSG00000131626.16 | PPFIA1 | rs513487 | 0.0000045 | ##### | Brain - Putamen (basal gangli | 70324575 | 69824575 | 70824575 | Yes |

| | | | | | | | | | |
|---|---|---|---|---|---|---|---|---|---|
| ENSG00000131626.16 | PPFIA1 | rs679455 | 0.0000045 | ##### | Brain - Putamen (basal gangli | 70324575 | 69824575 | 70824575 | Yes |
| ENSG00000131626.16 | PPFIA1 | rs676889 | 0.0000045 | ##### | Brain - Putamen (basal gangli | 70324575 | 69824575 | 70824575 | Yes |
| ENSG00000131626.16 | PPFIA1 | rs10591164 | 0.0000045 | ##### | Brain - Frontal Cortex (BA9) | 70324575 | 69824575 | 70824575 | Yes |
| ENSG00000131626.16 | PPFIA1 | rs3132436 | 0.0000047 | ##### | Brain - Nucleus accumbens (b | 70324575 | 69824575 | 70824575 | Yes |
| ENSG00000131626.16 | PPFIA1 | rs4523719 | 0.0000048 | ##### | Brain - Cerebellum | 70324575 | 69824575 | 70824575 | Yes |
| ENSG00000131626.16 | PPFIA1 | rs11236275 | 0.0000048 | ##### | Brain - Putamen (basal gangli | 70324575 | 69824575 | 70824575 | Yes |
| ENSG00000131626.16 | PPFIA1 | rs12283131 | 0.0000049 | ##### | Brain - Hippocampus | 70324575 | 69824575 | 70824575 | Yes |
| ENSG00000131626.16 | PPFIA1 | rs674374 | 0.000005 | ##### | Brain - Nucleus accumbens (b | 70324575 | 69824575 | 70824575 | Yes |
| ENSG00000131626.16 | PPFIA1 | rs141709344 | 0.0000051 | ##### | Brain - Putamen (basal gangli | 70324575 | 69824575 | 70824575 | Yes |
| ENSG00000131626.16 | PPFIA1 | rs6592546 | 0.0000052 | ##### | Brain - Hippocampus | 70324575 | 69824575 | 70824575 | Yes |
| ENSG00000131626.16 | PPFIA1 | rs4980782 | 0.0000053 | ##### | Brain - Spinal cord (cervical c- | 70324575 | 69824575 | 70824575 | Yes |
| ENSG00000131626.16 | PPFIA1 | rs142385955 | 0.0000058 | ##### | Brain - Hippocampus | 70324575 | 69824575 | 70824575 | Yes |
| ENSG00000131626.16 | PPFIA1 | rs1198234 | 0.0000058 | ##### | Brain - Hippocampus | 70324575 | 69824575 | 70824575 | Yes |
| ENSG00000131626.16 | PPFIA1 | rs12283131 | 0.0000061 | ##### | Brain - Spinal cord (cervical c- | 70324575 | 69824575 | 70824575 | Yes |
| ENSG00000131626.16 | PPFIA1 | rs1788973 | 0.0000061 | ##### | Brain - Nucleus accumbens (b | 70324575 | 69824575 | 70824575 | Yes |
| ENSG00000131626.16 | PPFIA1 | rs10683183 | 0.0000061 | ##### | Brain - Nucleus accumbens (b | 70324575 | 69824575 | 70824575 | Yes |
| ENSG00000131626.16 | PPFIA1 | rs6592542 | 0.0000061 | ##### | Brain - Nucleus accumbens (b | 70324575 | 69824575 | 70824575 | Yes |
| ENSG00000131626.16 | PPFIA1 | rs674374 | 0.0000062 | ##### | Brain - Putamen (basal gangli | 70324575 | 69824575 | 70824575 | Yes |
| ENSG00000131626.16 | PPFIA1 | rs1788973 | 0.0000062 | ##### | Brain - Hippocampus | 70324575 | 69824575 | 70824575 | Yes |
| ENSG00000131626.16 | PPFIA1 | rs10683183 | 0.0000062 | ##### | Brain - Hippocampus | 70324575 | 69824575 | 70824575 | Yes |
| ENSG00000131626.16 | PPFIA1 | rs6592542 | 0.0000062 | ##### | Brain - Hippocampus | 70324575 | 69824575 | 70824575 | Yes |
| ENSG00000131626.16 | PPFIA1 | rs4367974 | 0.0000062 | ##### | Brain - Hypothalamus | 70324575 | 69824575 | 70824575 | Yes |
| ENSG00000131626.16 | PPFIA1 | rs603927 | 0.0000064 | ##### | Brain - Cerebellum | 70324575 | 69824575 | 70824575 | Yes |
| ENSG00000131626.16 | PPFIA1 | rs6592546 | 0.0000069 | ##### | Brain - Caudate (basal ganglia | 70324575 | 69824575 | 70824575 | Yes |
| ENSG00000131626.16 | PPFIA1 | rs141709344 | 0.0000071 | ##### | Brain - Nucleus accumbens (b | 70324575 | 69824575 | 70824575 | Yes |
| ENSG00000131626.16 | PPFIA1 | rs674374 | 0.0000073 | ##### | Brain - Frontal Cortex (BA9) | 70324575 | 69824575 | 70824575 | Yes |
| ENSG00000131626.16 | PPFIA1 | rs548687 | 0.0000074 | ##### | Brain - Nucleus accumbens (b | 70324575 | 69824575 | 70824575 | Yes |
| ENSG00000131626.16 | PPFIA1 | rs571554 | 0.0000075 | ##### | Brain - Hypothalamus | 70324575 | 69824575 | 70824575 | Yes |
| ENSG00000131626.16 | PPFIA1 | rs611216 | 0.0000076 | ##### | Brain - Nucleus accumbens (b | 70324575 | 69824575 | 70824575 | Yes |
| ENSG00000131626.16 | PPFIA1 | rs638510 | 0.0000077 | ##### | Brain - Nucleus accumbens (b | 70324575 | 69824575 | 70824575 | Yes |
| ENSG00000131626.16 | PPFIA1 | rs11356872 | 0.0000079 | ##### | Brain - Nucleus accumbens (b | 70324575 | 69824575 | 70824575 | Yes |
| ENSG00000131626.16 | PPFIA1 | rs674374 | 0.0000079 | ##### | Brain - Caudate (basal ganglia | 70324575 | 69824575 | 70824575 | Yes |

| | | | | | | | | | |
|---|---|---|---|---|---|---|---|---|---|
| ENSG00000131626.16 | PPFIA1 | rs142385955 | 0.0000084 | ##### | Brain - Nucleus accumbens (b | 70324575 | 69824575 | 70824575 | Yes |
| ENSG00000131626.16 | PPFIA1 | rs1198234 | 0.0000084 | ##### | Brain - Nucleus accumbens (b | 70324575 | 69824575 | 70824575 | Yes |
| ENSG00000131626.16 | PPFIA1 | rs2282481 | 0.0000085 | ##### | Brain - Hippocampus | 70324575 | 69824575 | 70824575 | Yes |
| ENSG00000131626.16 | PPFIA1 | rs3132438 | 0.0000087 | ##### | Brain - Cortex | 70324575 | 69824575 | 70824575 | Yes |
| ENSG00000131626.16 | PPFIA1 | rs499484 | 0.0000091 | ##### | Brain - Hippocampus | 70324575 | 69824575 | 70824575 | Yes |
| ENSG00000131626.16 | PPFIA1 | rs4980782 | 0.0000091 | ##### | Brain - Nucleus accumbens (b | 70324575 | 69824575 | 70824575 | Yes |
| ENSG00000131626.16 | PPFIA1 | rs561050 | 0.0000095 | ##### | Brain - Anterior cingulate cort | 70324575 | 69824575 | 70824575 | Yes |
| ENSG00000131626.16 | PPFIA1 | rs546502 | 0.00001 | ##### | Brain - Spinal cord (cervical c- | 70324575 | 69824575 | 70824575 | Yes |
| ENSG00000131626.16 | PPFIA1 | rs624765 | 0.00001 | ##### | Brain - Spinal cord (cervical c- | 70324575 | 69824575 | 70824575 | Yes |
| ENSG00000131626.16 | PPFIA1 | rs3892505 | 0.00001 | ##### | Brain - Spinal cord (cervical c- | 70324575 | 69824575 | 70824575 | Yes |
| ENSG00000131626.16 | PPFIA1 | rs10591164 | 0.00001 | ##### | Brain - Spinal cord (cervical c- | 70324575 | 69824575 | 70824575 | Yes |
| ENSG00000131626.16 | PPFIA1 | rs35795372 | 0.00001 | ##### | Brain - Spinal cord (cervical c- | 70324575 | 69824575 | 70824575 | Yes |
| ENSG00000131626.16 | PPFIA1 | rs664130 | 0.00001 | ##### | Brain - Spinal cord (cervical c- | 70324575 | 69824575 | 70824575 | Yes |
| ENSG00000131626.16 | PPFIA1 | rs35077099 | 0.00001 | ##### | Brain - Hypothalamus | 70324575 | 69824575 | 70824575 | Yes |
| ENSG00000131626.16 | PPFIA1 | rs141709344 | 0.000011 | ##### | Brain - Spinal cord (cervical c- | 70324575 | 69824575 | 70824575 | Yes |
| ENSG00000131626.16 | PPFIA1 | rs583397 | 0.000011 | ##### | Brain - Nucleus accumbens (b | 70324575 | 69824575 | 70824575 | Yes |
| ENSG00000131626.16 | PPFIA1 | rs561050 | 0.000011 | ##### | Brain - Spinal cord (cervical c- | 70324575 | 69824575 | 70824575 | Yes |
| ENSG00000131626.16 | PPFIA1 | rs6592546 | 0.000011 | ##### | Brain - Nucleus accumbens (b | 70324575 | 69824575 | 70824575 | Yes |
| ENSG00000131626.16 | PPFIA1 | rs61341131 | 0.000012 | ##### | Brain - Hypothalamus | 70324575 | 69824575 | 70824575 | Yes |
| ENSG00000131626.16 | PPFIA1 | rs558391 | 0.000012 | ##### | Brain - Cerebellum | 70324575 | 69824575 | 70824575 | Yes |
| ENSG00000131626.16 | PPFIA1 | rs573795 | 0.000012 | ##### | Brain - Cerebellum | 70324575 | 69824575 | 70824575 | Yes |
| ENSG00000131626.16 | PPFIA1 | rs583347 | 0.000012 | ##### | Brain - Hippocampus | 70324575 | 69824575 | 70824575 | Yes |
| ENSG00000131626.16 | PPFIA1 | rs552282 | 0.000012 | ##### | Brain - Hippocampus | 70324575 | 69824575 | 70824575 | Yes |
| ENSG00000131626.16 | PPFIA1 | rs76294966 | 0.000012 | ##### | Brain - Nucleus accumbens (b | 70324575 | 69824575 | 70824575 | Yes |
| ENSG00000131626.16 | PPFIA1 | rs4980780 | 0.000012 | ##### | Brain - Nucleus accumbens (b | 70324575 | 69824575 | 70824575 | Yes |
| ENSG00000131626.16 | PPFIA1 | rs7129635 | 0.000012 | ##### | Brain - Nucleus accumbens (b | 70324575 | 69824575 | 70824575 | Yes |
| ENSG00000131626.16 | PPFIA1 | rs594763 | 0.000012 | ##### | Brain - Nucleus accumbens (b | 70324575 | 69824575 | 70824575 | Yes |
| ENSG00000131626.16 | PPFIA1 | rs35397909 | 0.000013 | ##### | Brain - Cerebellum | 70324575 | 69824575 | 70824575 | Yes |
| ENSG00000131626.16 | PPFIA1 | rs10591164 | 0.000013 | ##### | Brain - Cortex | 70324575 | 69824575 | 70824575 | Yes |
| ENSG00000131626.16 | PPFIA1 | rs76294966 | 0.000013 | ##### | Brain - Cortex | 70324575 | 69824575 | 70824575 | Yes |
| ENSG00000131626.16 | PPFIA1 | rs4980780 | 0.000013 | ##### | Brain - Cortex | 70324575 | 69824575 | 70824575 | Yes |
| ENSG00000131626.16 | PPFIA1 | rs7129635 | 0.000013 | ##### | Brain - Cortex | 70324575 | 69824575 | 70824575 | Yes |

| | | | | | | | | | |
|---|---|---|---|---|---|---|---|---|---|
| ENSG00000131626.16 | PPFIA1 | rs594763 | 0.000013 | ##### | Brain - Cortex | 70324575 | 69824575 | 70824575 | Yes |
| ENSG00000131626.16 | PPFIA1 | rs3132897 | 0.000013 | ##### | Brain - Cerebellum | 70324575 | 69824575 | 70824575 | Yes |
| ENSG00000131626.16 | PPFIA1 | rs75587843 | 0.000013 | ##### | Brain - Caudate (basal ganglia | 70324575 | 69824575 | 70824575 | No |
| ENSG00000131626.16 | PPFIA1 | rs71819112 | 0.000013 | ##### | Brain - Caudate (basal ganglia | 70324575 | 69824575 | 70824575 | No |
| ENSG00000131626.16 | PPFIA1 | rs114742415 | 0.000013 | ##### | Brain - Caudate (basal ganglia | 70324575 | 69824575 | 70824575 | No |
| ENSG00000131626.16 | PPFIA1 | rs3132438 | 0.000014 | ##### | Brain - Nucleus accumbens (b | 70324575 | 69824575 | 70824575 | Yes |
| ENSG00000131626.16 | PPFIA1 | rs34612166 | 0.000014 | ##### | Brain - Hypothalamus | 70324575 | 69824575 | 70824575 | Yes |
| ENSG00000131626.16 | PPFIA1 | rs34733714 | 0.000014 | ##### | Brain - Hypothalamus | 70324575 | 69824575 | 70824575 | Yes |
| ENSG00000131626.16 | PPFIA1 | rs2282481 | 0.000014 | ##### | Brain - Putamen (basal gangli | 70324575 | 69824575 | 70824575 | Yes |
| ENSG00000131626.16 | PPFIA1 | rs34612166 | 0.000014 | ##### | Brain - Cortex | 70324575 | 69824575 | 70824575 | Yes |
| ENSG00000131626.16 | PPFIA1 | rs199814664 | 0.000014 | ##### | Brain - Cerebellum | 70324575 | 69824575 | 70824575 | Yes |
| ENSG00000131626.16 | PPFIA1 | rs583347 | 0.000014 | ##### | Brain - Putamen (basal gangli | 70324575 | 69824575 | 70824575 | Yes |
| ENSG00000131626.16 | PPFIA1 | rs552282 | 0.000014 | ##### | Brain - Putamen (basal gangli | 70324575 | 69824575 | 70824575 | Yes |
| ENSG00000131626.16 | PPFIA1 | rs576588 | 0.000015 | ##### | Brain - Hypothalamus | 70324575 | 69824575 | 70824575 | Yes |
| ENSG00000131626.16 | PPFIA1 | rs34141396 | 0.000015 | ##### | Brain - Cortex | 70324575 | 69824575 | 70824575 | Yes |
| ENSG00000131626.16 | PPFIA1 | rs3132421 | 0.000015 | ##### | Brain - Cortex | 70324575 | 69824575 | 70824575 | Yes |
| ENSG00000131626.16 | PPFIA1 | rs546502 | 0.000016 | ##### | Brain - Cortex | 70324575 | 69824575 | 70824575 | Yes |
| ENSG00000131626.16 | PPFIA1 | rs624765 | 0.000016 | ##### | Brain - Cortex | 70324575 | 69824575 | 70824575 | Yes |
| ENSG00000131626.16 | PPFIA1 | rs3892505 | 0.000016 | ##### | Brain - Cortex | 70324575 | 69824575 | 70824575 | Yes |
| ENSG00000131626.16 | PPFIA1 | rs34141396 | 0.000016 | ##### | Brain - Nucleus accumbens (b | 70324575 | 69824575 | 70824575 | Yes |
| ENSG00000131626.16 | PPFIA1 | rs3132421 | 0.000016 | ##### | Brain - Nucleus accumbens (b | 70324575 | 69824575 | 70824575 | Yes |
| ENSG00000131626.16 | PPFIA1 | rs569732 | 0.000016 | ##### | Brain - Nucleus accumbens (b | 70324575 | 69824575 | 70824575 | Yes |
| ENSG00000131626.16 | PPFIA1 | rs513487 | 0.000016 | ##### | Brain - Nucleus accumbens (b | 70324575 | 69824575 | 70824575 | Yes |
| ENSG00000131626.16 | PPFIA1 | rs679455 | 0.000016 | ##### | Brain - Nucleus accumbens (b | 70324575 | 69824575 | 70824575 | Yes |
| ENSG00000131626.16 | PPFIA1 | rs11236048 | 0.000016 | ##### | Brain - Cortex | 70324575 | 69824575 | 70824575 | Yes |
| ENSG00000131626.16 | PPFIA1 | rs1788974 | 0.000017 | ##### | Brain - Anterior cingulate cort | 70324575 | 69824575 | 70824575 | Yes |
| ENSG00000131626.16 | PPFIA1 | rs36037711 | 0.000017 | ##### | Brain - Anterior cingulate cort | 70324575 | 69824575 | 70824575 | Yes |
| ENSG00000131626.16 | PPFIA1 | rs1198236 | 0.000017 | ##### | Brain - Nucleus accumbens (b | 70324575 | 69824575 | 70824575 | Yes |
| ENSG00000131626.16 | PPFIA1 | rs1788973 | 0.000017 | ##### | Brain - Caudate (basal ganglia | 70324575 | 69824575 | 70824575 | Yes |
| ENSG00000131626.16 | PPFIA1 | rs10683183 | 0.000017 | ##### | Brain - Caudate (basal ganglia | 70324575 | 69824575 | 70824575 | Yes |
| ENSG00000131626.16 | PPFIA1 | rs6592542 | 0.000017 | ##### | Brain - Caudate (basal ganglia | 70324575 | 69824575 | 70824575 | Yes |
| ENSG00000131626.16 | PPFIA1 | rs11236048 | 0.000018 | ##### | Brain - Spinal cord (cervical c- | 70324575 | 69824575 | 70824575 | Yes |

| | | | | | | | | | |
|---|---|---|---|---|---|---|---|---|---|
| ENSG00000131626.16 | PPFIA1 | rs499484 | 0.000018 | ##### | Brain - Spinal cord (cervical c- | 70324575 | 69824575 | 70824575 | Yes |
| ENSG00000131626.16 | PPFIA1 | rs482438 | 0.000018 | ##### | Brain - Nucleus accumbens (b | 70324575 | 69824575 | 70824575 | Yes |
| ENSG00000131626.16 | PPFIA1 | rs643301 | 0.000018 | ##### | Brain - Nucleus accumbens (b | 70324575 | 69824575 | 70824575 | Yes |
| ENSG00000131626.16 | PPFIA1 | rs3802780 | 0.000018 | ##### | Brain - Nucleus accumbens (b | 70324575 | 69824575 | 70824575 | Yes |
| ENSG00000131626.16 | PPFIA1 | rs670012 | 0.000018 | ##### | Brain - Hypothalamus | 70324575 | 69824575 | 70824575 | Yes |
| ENSG00000131626.16 | PPFIA1 | rs4980779 | 0.000018 | ##### | Brain - Hypothalamus | 70324575 | 69824575 | 70824575 | Yes |
| ENSG00000131626.16 | PPFIA1 | rs7130178 | 0.000018 | ##### | Brain - Hypothalamus | 70324575 | 69824575 | 70824575 | Yes |
| ENSG00000131626.16 | PPFIA1 | rs11236048 | 0.000018 | ##### | Brain - Nucleus accumbens (b | 70324575 | 69824575 | 70824575 | Yes |
| ENSG00000131626.16 | PPFIA1 | rs486852 | 0.000018 | ##### | Brain - Hypothalamus | 70324575 | 69824575 | 70824575 | Yes |
| ENSG00000131626.16 | PPFIA1 | rs141709344 | 0.000019 | ##### | Brain - Hippocampus | 70324575 | 69824575 | 70824575 | Yes |
| ENSG00000131626.16 | PPFIA1 | rs627079 | 0.000019 | ##### | Brain - Cerebellar Hemisphere | 70324575 | 69824575 | 70824575 | Yes |
| ENSG00000131626.16 | PPFIA1 | rs35795372 | 0.000019 | ##### | Brain - Cortex | 70324575 | 69824575 | 70824575 | Yes |
| ENSG00000131626.16 | PPFIA1 | rs664130 | 0.000019 | ##### | Brain - Cortex | 70324575 | 69824575 | 70824575 | Yes |
| ENSG00000131626.16 | PPFIA1 | rs571554 | 0.000019 | ##### | Brain - Anterior cingulate cort | 70324575 | 69824575 | 70824575 | Yes |
| ENSG00000131626.16 | PPFIA1 | rs592501 | 0.000019 | ##### | Brain - Nucleus accumbens (b | 70324575 | 69824575 | 70824575 | Yes |
| ENSG00000131626.16 | PPFIA1 | rs488746 | 0.000019 | ##### | Brain - Anterior cingulate cort | 70324575 | 69824575 | 70824575 | Yes |
| ENSG00000131626.16 | PPFIA1 | rs546724 | 0.000019 | ##### | Brain - Anterior cingulate cort | 70324575 | 69824575 | 70824575 | Yes |
| ENSG00000131626.16 | PPFIA1 | rs61887401 | 0.000019 | ##### | Brain - Anterior cingulate cort | 70324575 | 69824575 | 70824575 | Yes |
| ENSG00000131626.16 | PPFIA1 | rs113180277 | 0.000019 | ##### | Brain - Anterior cingulate cort | 70324575 | 69824575 | 70824575 | Yes |
| ENSG00000131626.16 | PPFIA1 | rs693315 | 0.000019 | ##### | Brain - Anterior cingulate cort | 70324575 | 69824575 | 70824575 | Yes |
| ENSG00000131626.16 | PPFIA1 | rs531498 | 0.000019 | ##### | Brain - Anterior cingulate cort | 70324575 | 69824575 | 70824575 | Yes |
| ENSG00000131626.16 | PPFIA1 | rs670310 | 0.000019 | ##### | Brain - Anterior cingulate cort | 70324575 | 69824575 | 70824575 | Yes |
| ENSG00000131626.16 | PPFIA1 | rs524333 | 0.000019 | ##### | Brain - Anterior cingulate cort | 70324575 | 69824575 | 70824575 | Yes |
| ENSG00000131626.16 | PPFIA1 | rs559176 | 0.000019 | ##### | Brain - Anterior cingulate cort | 70324575 | 69824575 | 70824575 | Yes |
| ENSG00000131626.16 | PPFIA1 | rs674374 | 0.00002 | ##### | Brain - Hippocampus | 70324575 | 69824575 | 70824575 | Yes |
| ENSG00000131626.16 | PPFIA1 | rs4417299 | 0.00002 | ##### | Brain - Cerebellum | 70324575 | 69824575 | 70824575 | Yes |
| ENSG00000131626.16 | PPFIA1 | rs678575 | 0.00002 | ##### | Brain - Cerebellum | 70324575 | 69824575 | 70824575 | Yes |
| ENSG00000131626.16 | PPFIA1 | rs77224759 | 0.00002 | ##### | Brain - Hypothalamus | 70324575 | 69824575 | 70824575 | Yes |
| ENSG00000131626.16 | PPFIA1 | rs61597492 | 0.00002 | ##### | Brain - Hypothalamus | 70324575 | 69824575 | 70824575 | Yes |
| ENSG00000131626.16 | PPFIA1 | rs11602439 | 0.00002 | ##### | Brain - Cerebellum | 70324575 | 69824575 | 70824575 | Yes |
| ENSG00000131626.16 | PPFIA1 | rs3132436 | 0.000021 | ##### | Brain - Putamen (basal gangli | 70324575 | 69824575 | 70824575 | Yes |
| ENSG00000131626.16 | PPFIA1 | rs12295241 | 0.000022 | ##### | Brain - Hypothalamus | 70324575 | 69824575 | 70824575 | Yes |

| | | | | | | | | | |
|---|---|---|---|---|---|---|---|---|---|
| ENSG00000131626.16 | PPFIA1 | rs7109643 | 0.000022 | ##### | Brain - Hypothalamus | 70324575 | 69824575 | 70824575 | Yes |
| ENSG00000131626.16 | PPFIA1 | rs12284490 | 0.000023 | ##### | Brain - Putamen (basal gangli | 70324575 | 69824575 | 70824575 | Yes |
| ENSG00000131626.16 | PPFIA1 | rs561050 | 0.000023 | ##### | Brain - Putamen (basal gangli | 70324575 | 69824575 | 70824575 | Yes |
| ENSG00000131626.16 | PPFIA1 | rs548687 | 0.000023 | ##### | Brain - Cortex | 70324575 | 69824575 | 70824575 | Yes |
| ENSG00000131626.16 | PPFIA1 | rs1198236 | 0.000023 | ##### | Brain - Cortex | 70324575 | 69824575 | 70824575 | Yes |
| ENSG00000131626.16 | PPFIA1 | rs643301 | 0.000023 | ##### | Brain - Cortex | 70324575 | 69824575 | 70824575 | Yes |
| ENSG00000131626.16 | PPFIA1 | rs2307683 | 0.000024 | ##### | Brain - Cerebellar Hemisphere | 70324575 | 69824575 | 70824575 | Yes |
| ENSG00000131626.16 | PPFIA1 | rs141709344 | 0.000024 | ##### | Brain - Caudate (basal ganglia | 70324575 | 69824575 | 70824575 | Yes |
| ENSG00000131626.16 | PPFIA1 | rs7942766 | 0.000025 | ##### | Brain - Anterior cingulate cort | 70324575 | 69824575 | 70824575 | Yes |
| ENSG00000131626.16 | PPFIA1 | rs111368954 | 0.000025 | ##### | Brain - Frontal Cortex (BA9) | 70324575 | 69824575 | 70824575 | Yes |
| ENSG00000131626.16 | PPFIA1 | rs619650 | 0.000025 | ##### | Brain - Nucleus accumbens (b | 70324575 | 69824575 | 70824575 | Yes |
| ENSG00000131626.16 | PPFIA1 | rs3132424 | 0.000025 | ##### | Brain - Nucleus accumbens (b | 70324575 | 69824575 | 70824575 | Yes |
| ENSG00000131626.16 | PPFIA1 | rs611216 | 0.000026 | ##### | Brain - Cortex | 70324575 | 69824575 | 70824575 | Yes |
| ENSG00000131626.16 | PPFIA1 | rs482438 | 0.000026 | ##### | Brain - Cortex | 70324575 | 69824575 | 70824575 | Yes |
| ENSG00000131626.16 | PPFIA1 | rs3802780 | 0.000026 | ##### | Brain - Cortex | 70324575 | 69824575 | 70824575 | Yes |
| ENSG00000131626.16 | PPFIA1 | rs2282481 | 0.000027 | ##### | Brain - Caudate (basal ganglia | 70324575 | 69824575 | 70824575 | Yes |
| ENSG00000131626.16 | PPFIA1 | rs77979967 | 0.000027 | ##### | Brain - Caudate (basal ganglia | 70324575 | 69824575 | 70824575 | No |
| ENSG00000131626.16 | PPFIA1 | rs12284490 | 0.000027 | ##### | Brain - Caudate (basal ganglia | 70324575 | 69824575 | 70824575 | Yes |
| ENSG00000131626.16 | PPFIA1 | rs583397 | 0.000027 | ##### | Brain - Cortex | 70324575 | 69824575 | 70824575 | Yes |
| ENSG00000131626.16 | PPFIA1 | rs676889 | 0.000027 | ##### | Brain - Nucleus accumbens (b | 70324575 | 69824575 | 70824575 | Yes |
| ENSG00000131626.16 | PPFIA1 | rs662018 | 0.000028 | ##### | Brain - Nucleus accumbens (b | 70324575 | 69824575 | 70824575 | Yes |
| ENSG00000131626.16 | PPFIA1 | rs12284490 | 0.000028 | ##### | Brain - Nucleus accumbens (b | 70324575 | 69824575 | 70824575 | Yes |
| ENSG00000131626.16 | PPFIA1 | rs34612166 | 0.000028 | ##### | Brain - Cerebellum | 70324575 | 69824575 | 70824575 | Yes |
| ENSG00000131626.16 | PPFIA1 | rs11236275 | 0.000029 | ##### | Brain - Nucleus accumbens (b | 70324575 | 69824575 | 70824575 | Yes |
| ENSG00000131626.16 | PPFIA1 | rs111368954 | 0.000029 | ##### | Brain - Hippocampus | 70324575 | 69824575 | 70824575 | Yes |
| ENSG00000131626.16 | PPFIA1 | rs142385955 | 0.000029 | ##### | Brain - Cortex | 70324575 | 69824575 | 70824575 | Yes |
| ENSG00000131626.16 | PPFIA1 | rs1198234 | 0.000029 | ##### | Brain - Cortex | 70324575 | 69824575 | 70824575 | Yes |
| ENSG00000131626.16 | PPFIA1 | rs592501 | 0.000029 | ##### | Brain - Cortex | 70324575 | 69824575 | 70824575 | Yes |
| ENSG00000131626.16 | PPFIA1 | rs61887436 | 0.00003 | ##### | Brain - Cerebellar Hemisphere | 70324575 | 69824575 | 70824575 | Yes |
| ENSG00000131626.16 | PPFIA1 | rs670012 | 0.000031 | ##### | Brain - Anterior cingulate cort | 70324575 | 69824575 | 70824575 | Yes |
| ENSG00000131626.16 | PPFIA1 | rs4980779 | 0.000031 | ##### | Brain - Anterior cingulate cort | 70324575 | 69824575 | 70824575 | Yes |
| ENSG00000131626.16 | PPFIA1 | rs7130178 | 0.000031 | ##### | Brain - Anterior cingulate cort | 70324575 | 69824575 | 70824575 | Yes |

| | | | | | | | | | |
|---|---|---|---|---|---|---|---|---|---|
| ENSG00000131626.16 | PPFIA1 | rs2307683 | 0.000032 | ##### | Brain - Cerebellum | 70324575 | 69824575 | 70824575 | Yes |
| ENSG00000131626.16 | PPFIA1 | rs10591164 | 0.000032 | ##### | Brain - Nucleus accumbens (b | 70324575 | 69824575 | 70824575 | Yes |
| ENSG00000131626.16 | PPFIA1 | rs576588 | 0.000032 | ##### | Brain - Anterior cingulate cort | 70324575 | 69824575 | 70824575 | Yes |
| ENSG00000131626.16 | PPFIA1 | rs499484 | 0.000033 | ##### | Brain - Cortex | 70324575 | 69824575 | 70824575 | Yes |
| ENSG00000131626.16 | PPFIA1 | rs111368954 | 0.000034 | ##### | Brain - Putamen (basal gangli | 70324575 | 69824575 | 70824575 | Yes |
| ENSG00000131626.16 | PPFIA1 | rs583347 | 0.000035 | ##### | Brain - Caudate (basal ganglia | 70324575 | 69824575 | 70824575 | Yes |
| ENSG00000131626.16 | PPFIA1 | rs552282 | 0.000035 | ##### | Brain - Caudate (basal ganglia | 70324575 | 69824575 | 70824575 | Yes |
| ENSG00000131626.16 | PPFIA1 | rs111368954 | 0.000036 | ##### | Brain - Nucleus accumbens (b | 70324575 | 69824575 | 70824575 | Yes |
| ENSG00000131626.16 | PPFIA1 | rs3781649 | 0.000036 | ##### | Brain - Cortex | 70324575 | 69824575 | 70824575 | Yes |
| ENSG00000131626.16 | PPFIA1 | rs3781646 | 0.000036 | ##### | Brain - Cortex | 70324575 | 69824575 | 70824575 | Yes |
| ENSG00000131626.16 | PPFIA1 | rs151079983 | 0.000036 | ##### | Brain - Cortex | 70324575 | 69824575 | 70824575 | Yes |
| ENSG00000131626.16 | PPFIA1 | rs567105 | 0.000038 | ##### | Brain - Cerebellum | 70324575 | 69824575 | 70824575 | Yes |
| ENSG00000131626.16 | PPFIA1 | rs682670 | 0.000038 | ##### | Brain - Cerebellum | 70324575 | 69824575 | 70824575 | Yes |
| ENSG00000131626.16 | PPFIA1 | rs61375303 | 0.000039 | ##### | Brain - Cerebellum | 70324575 | 69824575 | 70824575 | Yes |
| ENSG00000131626.16 | PPFIA1 | rs583347 | 0.00004 | ##### | Brain - Nucleus accumbens (b | 70324575 | 69824575 | 70824575 | Yes |
| ENSG00000131626.16 | PPFIA1 | rs552282 | 0.00004 | ##### | Brain - Nucleus accumbens (b | 70324575 | 69824575 | 70824575 | Yes |
| ENSG00000131626.16 | PPFIA1 | rs34612166 | 0.000041 | ##### | Brain - Cerebellar Hemisphere | 70324575 | 69824575 | 70824575 | Yes |
| ENSG00000131626.16 | PPFIA1 | rs576082 | 0.000045 | ##### | Brain - Cerebellum | 70324575 | 69824575 | 70824575 | Yes |
| ENSG00000131626.16 | PPFIA1 | rs546502 | 0.000048 | ##### | Brain - Nucleus accumbens (b | 70324575 | 69824575 | 70824575 | Yes |
| ENSG00000131626.16 | PPFIA1 | rs624765 | 0.000048 | ##### | Brain - Nucleus accumbens (b | 70324575 | 69824575 | 70824575 | Yes |
| ENSG00000131626.16 | PPFIA1 | rs3892505 | 0.000048 | ##### | Brain - Nucleus accumbens (b | 70324575 | 69824575 | 70824575 | Yes |
| ENSG00000131626.16 | PPFIA1 | rs34733714 | 0.000054 | ##### | Brain - Cerebellum | 70324575 | 69824575 | 70824575 | Yes |
| ENSG00000181754.6 | AMIGO1 | rs2570972 | 6.40E-17 | ##### | Brain - Spinal cord (cervical c- | 109508840 | 109008840 | 110008840 | Yes |
| ENSG00000181754.6 | AMIGO1 | rs6684193 | 1.00E-12 | ##### | Brain - Spinal cord (cervical c- | 109508840 | 109008840 | 110008840 | Yes |
| ENSG00000181754.6 | AMIGO1 | rs6684194 | 1.00E-12 | ##### | Brain - Spinal cord (cervical c- | 109508840 | 109008840 | 110008840 | Yes |
| ENSG00000181754.6 | AMIGO1 | rs11101966 | 1.00E-12 | ##### | Brain - Spinal cord (cervical c- | 109508840 | 109008840 | 110008840 | Yes |
| ENSG00000181754.6 | AMIGO1 | rs534135 | 1.00E-12 | ##### | Brain - Spinal cord (cervical c- | 109508840 | 109008840 | 110008840 | Yes |
| ENSG00000181754.6 | AMIGO1 | rs553689 | 1.40E-12 | ##### | Brain - Spinal cord (cervical c- | 109508840 | 109008840 | 110008840 | Yes |
| ENSG00000181754.6 | AMIGO1 | rs732715 | 2.10E-12 | ##### | Brain - Spinal cord (cervical c- | 109508840 | 109008840 | 110008840 | Yes |
| ENSG00000181754.6 | AMIGO1 | rs2781556 | 3.00E-12 | ##### | Brain - Spinal cord (cervical c- | 109508840 | 109008840 | 110008840 | Yes |
| ENSG00000181754.6 | AMIGO1 | rs528077 | 3.00E-12 | ##### | Brain - Spinal cord (cervical c- | 109508840 | 109008840 | 110008840 | Yes |
| ENSG00000181754.6 | AMIGO1 | rs529773 | 3.00E-12 | ##### | Brain - Spinal cord (cervical c- | 109508840 | 109008840 | 110008840 | Yes |

| | | | | | | | | | |
|---|---|---|---|---|---|---|---|---|---|
| ENSG00000181754.6 | AMIGO1 | rs2570973 | 3.00E-12 | ##### | Brain - Spinal cord (cervical c- | 109508840 | 109008840 | 110008840 | Yes |
| ENSG00000181754.6 | AMIGO1 | rs2781557 | 3.00E-12 | ##### | Brain - Spinal cord (cervical c- | 109508840 | 109008840 | 110008840 | Yes |
| ENSG00000181754.6 | AMIGO1 | rs538654 | 3.00E-12 | ##### | Brain - Spinal cord (cervical c- | 109508840 | 109008840 | 110008840 | Yes |
| ENSG00000181754.6 | AMIGO1 | rs543021 | 3.00E-12 | ##### | Brain - Spinal cord (cervical c- | 109508840 | 109008840 | 110008840 | Yes |
| ENSG00000181754.6 | AMIGO1 | rs543046 | 3.00E-12 | ##### | Brain - Spinal cord (cervical c- | 109508840 | 109008840 | 110008840 | Yes |
| ENSG00000181754.6 | AMIGO1 | rs2692703 | 3.00E-12 | ##### | Brain - Spinal cord (cervical c- | 109508840 | 109008840 | 110008840 | Yes |
| ENSG00000181754.6 | AMIGO1 | rs2692704 | 3.00E-12 | ##### | Brain - Spinal cord (cervical c- | 109508840 | 109008840 | 110008840 | Yes |
| ENSG00000181754.6 | AMIGO1 | rs474234 | 3.00E-12 | ##### | Brain - Spinal cord (cervical c- | 109508840 | 109008840 | 110008840 | Yes |
| ENSG00000181754.6 | AMIGO1 | rs558370 | 3.00E-12 | ##### | Brain - Spinal cord (cervical c- | 109508840 | 109008840 | 110008840 | Yes |
| ENSG00000181754.6 | AMIGO1 | rs528199 | 3.60E-12 | ##### | Brain - Spinal cord (cervical c- | 109508840 | 109008840 | 110008840 | Yes |
| ENSG00000181754.6 | AMIGO1 | rs1149170 | 8.20E-12 | 0.2 | Brain - Cortex | 109508840 | 109008840 | 110008840 | Yes |
| ENSG00000181754.6 | AMIGO1 | rs2570972 | 1.40E-11 | 0.17 | Brain - Hippocampus | 109508840 | 109008840 | 110008840 | Yes |
| ENSG00000181754.6 | AMIGO1 | rs2570972 | 2.30E-11 | ##### | Brain - Putamen (basal gangli | 109508840 | 109008840 | 110008840 | Yes |
| ENSG00000181754.6 | AMIGO1 | rs12741221 | 7.10E-11 | ##### | Brain - Cortex | 109508840 | 109008840 | 110008840 | Yes |
| ENSG00000181754.6 | AMIGO1 | rs3738774 | 7.10E-11 | ##### | Brain - Cortex | 109508840 | 109008840 | 110008840 | Yes |
| ENSG00000181754.6 | AMIGO1 | rs6684193 | 1.30E-10 | ##### | Brain - Hippocampus | 109508840 | 109008840 | 110008840 | Yes |
| ENSG00000181754.6 | AMIGO1 | rs6684194 | 1.30E-10 | ##### | Brain - Hippocampus | 109508840 | 109008840 | 110008840 | Yes |
| ENSG00000181754.6 | AMIGO1 | rs11101966 | 1.30E-10 | ##### | Brain - Hippocampus | 109508840 | 109008840 | 110008840 | Yes |
| ENSG00000181754.6 | AMIGO1 | rs553689 | 1.30E-10 | ##### | Brain - Hippocampus | 109508840 | 109008840 | 110008840 | Yes |
| ENSG00000181754.6 | AMIGO1 | rs534135 | 1.30E-10 | ##### | Brain - Hippocampus | 109508840 | 109008840 | 110008840 | Yes |
| ENSG00000181754.6 | AMIGO1 | rs2781556 | 1.30E-10 | ##### | Brain - Hippocampus | 109508840 | 109008840 | 110008840 | Yes |
| ENSG00000181754.6 | AMIGO1 | rs528077 | 1.30E-10 | ##### | Brain - Hippocampus | 109508840 | 109008840 | 110008840 | Yes |
| ENSG00000181754.6 | AMIGO1 | rs529773 | 1.30E-10 | ##### | Brain - Hippocampus | 109508840 | 109008840 | 110008840 | Yes |
| ENSG00000181754.6 | AMIGO1 | rs2570973 | 1.30E-10 | ##### | Brain - Hippocampus | 109508840 | 109008840 | 110008840 | Yes |
| ENSG00000181754.6 | AMIGO1 | rs2781557 | 1.30E-10 | ##### | Brain - Hippocampus | 109508840 | 109008840 | 110008840 | Yes |
| ENSG00000181754.6 | AMIGO1 | rs538654 | 1.30E-10 | ##### | Brain - Hippocampus | 109508840 | 109008840 | 110008840 | Yes |
| ENSG00000181754.6 | AMIGO1 | rs543021 | 1.30E-10 | ##### | Brain - Hippocampus | 109508840 | 109008840 | 110008840 | Yes |
| ENSG00000181754.6 | AMIGO1 | rs543046 | 1.30E-10 | ##### | Brain - Hippocampus | 109508840 | 109008840 | 110008840 | Yes |
| ENSG00000181754.6 | AMIGO1 | rs2692703 | 1.30E-10 | ##### | Brain - Hippocampus | 109508840 | 109008840 | 110008840 | Yes |
| ENSG00000181754.6 | AMIGO1 | rs2692704 | 1.30E-10 | ##### | Brain - Hippocampus | 109508840 | 109008840 | 110008840 | Yes |
| ENSG00000181754.6 | AMIGO1 | rs474234 | 1.30E-10 | ##### | Brain - Hippocampus | 109508840 | 109008840 | 110008840 | Yes |
| ENSG00000181754.6 | AMIGO1 | rs558370 | 1.30E-10 | ##### | Brain - Hippocampus | 109508840 | 109008840 | 110008840 | Yes |

| | | | | | | | | | |
|---|---|---|---|---|---|---|---|---|---|
| ENSG00000181754.6 | AMIGO1 | rs2570972 | 1.40E-10 | ##### | Brain - Caudate (basal ganglia | 109508840 | 109008840 | 110008840 | Yes |
| ENSG00000181754.6 | AMIGO1 | rs1149170 | 1.70E-10 | ##### | Brain - Spinal cord (cervical c- | 109508840 | 109008840 | 110008840 | Yes |
| ENSG00000181754.6 | AMIGO1 | rs35972691 | 2.10E-10 | ##### | Brain - Spinal cord (cervical c- | 109508840 | 109008840 | 110008840 | Yes |
| ENSG00000181754.6 | AMIGO1 | rs492557 | 2.30E-10 | ##### | Brain - Hippocampus | 109508840 | 109008840 | 110008840 | Yes |
| ENSG00000181754.6 | AMIGO1 | rs732715 | 3.90E-10 | ##### | Brain - Hippocampus | 109508840 | 109008840 | 110008840 | Yes |
| ENSG00000181754.6 | AMIGO1 | rs4970764 | 6.60E-10 | ##### | Brain - Spinal cord (cervical c- | 109508840 | 109008840 | 110008840 | Yes |
| ENSG00000181754.6 | AMIGO1 | rs7536291 | 2.30E-09 | ##### | Brain - Hippocampus | 109508840 | 109008840 | 110008840 | Yes |
| ENSG00000181754.6 | AMIGO1 | rs553689 | 2.30E-09 | ##### | Brain - Putamen (basal gangli | 109508840 | 109008840 | 110008840 | Yes |
| ENSG00000181754.6 | AMIGO1 | rs2781556 | 2.30E-09 | ##### | Brain - Putamen (basal gangli | 109508840 | 109008840 | 110008840 | Yes |
| ENSG00000181754.6 | AMIGO1 | rs528077 | 2.30E-09 | ##### | Brain - Putamen (basal gangli | 109508840 | 109008840 | 110008840 | Yes |
| ENSG00000181754.6 | AMIGO1 | rs529773 | 2.30E-09 | ##### | Brain - Putamen (basal gangli | 109508840 | 109008840 | 110008840 | Yes |
| ENSG00000181754.6 | AMIGO1 | rs2570973 | 2.30E-09 | ##### | Brain - Putamen (basal gangli | 109508840 | 109008840 | 110008840 | Yes |
| ENSG00000181754.6 | AMIGO1 | rs2781557 | 2.30E-09 | ##### | Brain - Putamen (basal gangli | 109508840 | 109008840 | 110008840 | Yes |
| ENSG00000181754.6 | AMIGO1 | rs538654 | 2.30E-09 | ##### | Brain - Putamen (basal gangli | 109508840 | 109008840 | 110008840 | Yes |
| ENSG00000181754.6 | AMIGO1 | rs543021 | 2.30E-09 | ##### | Brain - Putamen (basal gangli | 109508840 | 109008840 | 110008840 | Yes |
| ENSG00000181754.6 | AMIGO1 | rs543046 | 2.30E-09 | ##### | Brain - Putamen (basal gangli | 109508840 | 109008840 | 110008840 | Yes |
| ENSG00000181754.6 | AMIGO1 | rs2692703 | 2.30E-09 | ##### | Brain - Putamen (basal gangli | 109508840 | 109008840 | 110008840 | Yes |
| ENSG00000181754.6 | AMIGO1 | rs2692704 | 2.30E-09 | ##### | Brain - Putamen (basal gangli | 109508840 | 109008840 | 110008840 | Yes |
| ENSG00000181754.6 | AMIGO1 | rs474234 | 2.30E-09 | ##### | Brain - Putamen (basal gangli | 109508840 | 109008840 | 110008840 | Yes |
| ENSG00000181754.6 | AMIGO1 | rs558370 | 2.30E-09 | ##### | Brain - Putamen (basal gangli | 109508840 | 109008840 | 110008840 | Yes |
| ENSG00000181754.6 | AMIGO1 | rs12049330 | 2.30E-09 | ##### | Brain - Cortex | 109508840 | 109008840 | 110008840 | Yes |
| ENSG00000181754.6 | AMIGO1 | rs413582 | 2.80E-09 | ##### | Brain - Spinal cord (cervical c- | 109508840 | 109008840 | 110008840 | Yes |
| ENSG00000181754.6 | AMIGO1 | rs6684193 | 3.30E-09 | ##### | Brain - Putamen (basal gangli | 109508840 | 109008840 | 110008840 | Yes |
| ENSG00000181754.6 | AMIGO1 | rs6684194 | 3.30E-09 | ##### | Brain - Putamen (basal gangli | 109508840 | 109008840 | 110008840 | Yes |
| ENSG00000181754.6 | AMIGO1 | rs11101966 | 3.30E-09 | ##### | Brain - Putamen (basal gangli | 109508840 | 109008840 | 110008840 | Yes |
| ENSG00000181754.6 | AMIGO1 | rs534135 | 3.30E-09 | ##### | Brain - Putamen (basal gangli | 109508840 | 109008840 | 110008840 | Yes |
| ENSG00000181754.6 | AMIGO1 | rs12741221 | 3.70E-09 | ##### | Brain - Spinal cord (cervical c- | 109508840 | 109008840 | 110008840 | Yes |
| ENSG00000181754.6 | AMIGO1 | rs3738774 | 3.70E-09 | ##### | Brain - Spinal cord (cervical c- | 109508840 | 109008840 | 110008840 | Yes |
| ENSG00000181754.6 | AMIGO1 | rs4970843 | 5.50E-09 | ##### | Brain - Spinal cord (cervical c- | 109508840 | 109008840 | 110008840 | Yes |
| ENSG00000181754.6 | AMIGO1 | rs553689 | 7.10E-09 | ##### | Brain - Caudate (basal ganglia | 109508840 | 109008840 | 110008840 | Yes |
| ENSG00000181754.6 | AMIGO1 | rs528815 | 1.10E-08 | ##### | Brain - Spinal cord (cervical c- | 109508840 | 109008840 | 110008840 | Yes |
| ENSG00000181754.6 | AMIGO1 | rs532432 | 1.10E-08 | ##### | Brain - Spinal cord (cervical c- | 109508840 | 109008840 | 110008840 | Yes |

| | | | | | | | | | |
|---|---|---|---|---|---|---|---|---|---|
| ENSG00000181754.6 | AMIGO1 | rs475381 | 1.10E-08 | ##### | Brain - Spinal cord (cervical c- | 109508840 | 109008840 | 110008840 | Yes |
| ENSG00000181754.6 | AMIGO1 | rs523300 | 1.10E-08 | ##### | Brain - Spinal cord (cervical c- | 109508840 | 109008840 | 110008840 | Yes |
| ENSG00000181754.6 | AMIGO1 | rs2692705 | 1.10E-08 | ##### | Brain - Spinal cord (cervical c- | 109508840 | 109008840 | 110008840 | Yes |
| ENSG00000181754.6 | AMIGO1 | rs2781558 | 1.10E-08 | ##### | Brain - Spinal cord (cervical c- | 109508840 | 109008840 | 110008840 | Yes |
| ENSG00000181754.6 | AMIGO1 | rs2781559 | 1.10E-08 | ##### | Brain - Spinal cord (cervical c- | 109508840 | 109008840 | 110008840 | Yes |
| ENSG00000181754.6 | AMIGO1 | rs532623 | 1.10E-08 | ##### | Brain - Spinal cord (cervical c- | 109508840 | 109008840 | 110008840 | Yes |
| ENSG00000181754.6 | AMIGO1 | rs561741 | 1.10E-08 | ##### | Brain - Spinal cord (cervical c- | 109508840 | 109008840 | 110008840 | Yes |
| ENSG00000181754.6 | AMIGO1 | rs4571992 | 1.10E-08 | ##### | Brain - Spinal cord (cervical c- | 109508840 | 109008840 | 110008840 | Yes |
| ENSG00000181754.6 | AMIGO1 | rs472737 | 1.20E-08 | ##### | Brain - Spinal cord (cervical c- | 109508840 | 109008840 | 110008840 | Yes |
| ENSG00000181754.6 | AMIGO1 | rs2781555 | 1.20E-08 | ##### | Brain - Spinal cord (cervical c- | 109508840 | 109008840 | 110008840 | Yes |
| ENSG00000181754.6 | AMIGO1 | rs528199 | 1.20E-08 | ##### | Brain - Hippocampus | 109508840 | 109008840 | 110008840 | Yes |
| ENSG00000181754.6 | AMIGO1 | rs732715 | 1.30E-08 | ##### | Brain - Putamen (basal gangli | 109508840 | 109008840 | 110008840 | Yes |
| ENSG00000181754.6 | AMIGO1 | rs528077 | 1.40E-08 | ##### | Brain - Caudate (basal ganglia | 109508840 | 109008840 | 110008840 | Yes |
| ENSG00000181754.6 | AMIGO1 | rs529773 | 1.40E-08 | ##### | Brain - Caudate (basal ganglia | 109508840 | 109008840 | 110008840 | Yes |
| ENSG00000181754.6 | AMIGO1 | rs2570973 | 1.40E-08 | ##### | Brain - Caudate (basal ganglia | 109508840 | 109008840 | 110008840 | Yes |
| ENSG00000181754.6 | AMIGO1 | rs538654 | 1.40E-08 | ##### | Brain - Caudate (basal ganglia | 109508840 | 109008840 | 110008840 | Yes |
| ENSG00000181754.6 | AMIGO1 | rs543021 | 1.40E-08 | ##### | Brain - Caudate (basal ganglia | 109508840 | 109008840 | 110008840 | Yes |
| ENSG00000181754.6 | AMIGO1 | rs543046 | 1.40E-08 | ##### | Brain - Caudate (basal ganglia | 109508840 | 109008840 | 110008840 | Yes |
| ENSG00000181754.6 | AMIGO1 | rs2692704 | 1.40E-08 | ##### | Brain - Caudate (basal ganglia | 109508840 | 109008840 | 110008840 | Yes |
| ENSG00000181754.6 | AMIGO1 | rs474234 | 1.40E-08 | ##### | Brain - Caudate (basal ganglia | 109508840 | 109008840 | 110008840 | Yes |
| ENSG00000181754.6 | AMIGO1 | rs558370 | 1.40E-08 | ##### | Brain - Caudate (basal ganglia | 109508840 | 109008840 | 110008840 | Yes |
| ENSG00000181754.6 | AMIGO1 | rs6684193 | 1.60E-08 | ##### | Brain - Caudate (basal ganglia | 109508840 | 109008840 | 110008840 | Yes |
| ENSG00000181754.6 | AMIGO1 | rs6684194 | 1.60E-08 | ##### | Brain - Caudate (basal ganglia | 109508840 | 109008840 | 110008840 | Yes |
| ENSG00000181754.6 | AMIGO1 | rs11101966 | 1.60E-08 | ##### | Brain - Caudate (basal ganglia | 109508840 | 109008840 | 110008840 | Yes |
| ENSG00000181754.6 | AMIGO1 | rs534135 | 1.60E-08 | ##### | Brain - Caudate (basal ganglia | 109508840 | 109008840 | 110008840 | Yes |
| ENSG00000181754.6 | AMIGO1 | rs7536291 | 2.10E-08 | ##### | Brain - Spinal cord (cervical c- | 109508840 | 109008840 | 110008840 | Yes |
| ENSG00000181754.6 | AMIGO1 | rs528199 | 2.60E-08 | ##### | Brain - Caudate (basal ganglia | 109508840 | 109008840 | 110008840 | Yes |
| ENSG00000181754.6 | AMIGO1 | rs2781556 | 3.00E-08 | ##### | Brain - Caudate (basal ganglia | 109508840 | 109008840 | 110008840 | Yes |
| ENSG00000181754.6 | AMIGO1 | rs2781557 | 3.00E-08 | ##### | Brain - Caudate (basal ganglia | 109508840 | 109008840 | 110008840 | Yes |
| ENSG00000181754.6 | AMIGO1 | rs2692703 | 3.00E-08 | ##### | Brain - Caudate (basal ganglia | 109508840 | 109008840 | 110008840 | Yes |
| ENSG00000181754.6 | AMIGO1 | rs12066164 | 3.10E-08 | ##### | Brain - Spinal cord (cervical c- | 109508840 | 109008840 | 110008840 | Yes |
| ENSG00000181754.6 | AMIGO1 | rs2692706 | 3.10E-08 | ##### | Brain - Spinal cord (cervical c- | 109508840 | 109008840 | 110008840 | Yes |

| | | | | | | | | | |
|---|---|---|---|---|---|---|---|---|---|
| ENSG00000181754.6 | AMIGO1 | rs10494041 | 3.10E-08 | ##### | Brain - Cortex | 109508840 | 109008840 | 110008840 | Yes |
| ENSG00000181754.6 | AMIGO1 | rs732715 | 4.10E-08 | ##### | Brain - Caudate (basal ganglia | 109508840 | 109008840 | 110008840 | Yes |
| ENSG00000181754.6 | AMIGO1 | rs492557 | 4.30E-08 | ##### | Brain - Spinal cord (cervical c- | 109508840 | 109008840 | 110008840 | Yes |
| ENSG00000181754.6 | AMIGO1 | rs35608606 | 5.20E-08 | ##### | Brain - Putamen (basal gangli | 109508840 | 109008840 | 110008840 | Yes |
| ENSG00000181754.6 | AMIGO1 | rs4970764 | 5.90E-08 | ##### | Brain - Caudate (basal ganglia | 109508840 | 109008840 | 110008840 | Yes |
| ENSG00000181754.6 | AMIGO1 | rs4970762 | 6.80E-08 | ##### | Brain - Spinal cord (cervical c- | 109508840 | 109008840 | 110008840 | Yes |
| ENSG00000181754.6 | AMIGO1 | rs35972691 | 7.10E-08 | ##### | Brain - Caudate (basal ganglia | 109508840 | 109008840 | 110008840 | Yes |
| ENSG00000181754.6 | AMIGO1 | rs12037569 | 7.30E-08 | ##### | Brain - Cortex | 109508840 | 109008840 | 110008840 | Yes |
| ENSG00000181754.6 | AMIGO1 | rs35780288 | 7.30E-08 | ##### | Brain - Cortex | 109508840 | 109008840 | 110008840 | Yes |
| ENSG00000181754.6 | AMIGO1 | rs528199 | 8.50E-08 | ##### | Brain - Substantia nigra | 109508840 | 109008840 | 110008840 | Yes |
| ENSG00000181754.6 | AMIGO1 | rs3820667 | 9.10E-08 | ##### | Brain - Spinal cord (cervical c- | 109508840 | 109008840 | 110008840 | Yes |
| ENSG00000181754.6 | AMIGO1 | rs6669802 | 9.60E-08 | ##### | Brain - Putamen (basal gangli | 109508840 | 109008840 | 110008840 | Yes |
| ENSG00000181754.6 | AMIGO1 | rs12746776 | 1.20E-07 | ##### | Brain - Cortex | 109508840 | 109008840 | 110008840 | Yes |
| ENSG00000181754.6 | AMIGO1 | rs528199 | 1.30E-07 | ##### | Brain - Putamen (basal gangli | 109508840 | 109008840 | 110008840 | Yes |
| ENSG00000181754.6 | AMIGO1 | rs464218 | 1.70E-07 | ##### | Brain - Spinal cord (cervical c- | 109508840 | 109008840 | 110008840 | Yes |
| ENSG00000181754.6 | AMIGO1 | rs10858096 | 1.70E-07 | ##### | Brain - Spinal cord (cervical c- | 109508840 | 109008840 | 110008840 | Yes |
| ENSG00000181754.6 | AMIGO1 | rs2297797 | 1.80E-07 | ##### | Brain - Spinal cord (cervical c- | 109508840 | 109008840 | 110008840 | Yes |
| ENSG00000181754.6 | AMIGO1 | rs492557 | 2.20E-07 | ##### | Brain - Putamen (basal gangli | 109508840 | 109008840 | 110008840 | Yes |
| ENSG00000181754.6 | AMIGO1 | rs34571177 | 2.30E-07 | ##### | Brain - Cortex | 109508840 | 109008840 | 110008840 | Yes |
| ENSG00000181754.6 | AMIGO1 | rs657420 | 2.50E-07 | ##### | Brain - Caudate (basal ganglia | 109508840 | 109008840 | 110008840 | Yes |
| ENSG00000181754.6 | AMIGO1 | rs35375862 | 2.50E-07 | ##### | Brain - Cortex | 109508840 | 109008840 | 110008840 | Yes |
| ENSG00000181754.6 | AMIGO1 | rs4970843 | 3.70E-07 | ##### | Brain - Caudate (basal ganglia | 109508840 | 109008840 | 110008840 | Yes |
| ENSG00000181754.6 | AMIGO1 | rs2570972 | 4.90E-07 | ##### | Brain - Substantia nigra | 109508840 | 109008840 | 110008840 | Yes |
| ENSG00000181754.6 | AMIGO1 | rs4970764 | 4.90E-07 | ##### | Brain - Cerebellum | 109508840 | 109008840 | 110008840 | Yes |
| ENSG00000181754.6 | AMIGO1 | rs892510 | 5.10E-07 | ##### | Brain - Spinal cord (cervical c- | 109508840 | 109008840 | 110008840 | Yes |
| ENSG00000181754.6 | AMIGO1 | rs93063 | 5.10E-07 | ##### | Brain - Spinal cord (cervical c- | 109508840 | 109008840 | 110008840 | Yes |
| ENSG00000181754.6 | AMIGO1 | rs11351199 | 5.30E-07 | ##### | Brain - Spinal cord (cervical c- | 109508840 | 109008840 | 110008840 | Yes |
| ENSG00000181754.6 | AMIGO1 | rs2570972 | 5.40E-07 | ##### | Brain - Amygdala | 109508840 | 109008840 | 110008840 | Yes |
| ENSG00000181754.6 | AMIGO1 | rs80025011 | 5.80E-07 | ##### | Brain - Caudate (basal ganglia | 109508840 | 109008840 | 110008840 | Yes |
| ENSG00000181754.6 | AMIGO1 | rs4970764 | 6.00E-07 | ##### | Brain - Putamen (basal gangli | 109508840 | 109008840 | 110008840 | Yes |
| ENSG00000181754.6 | AMIGO1 | rs10857792 | 6.20E-07 | ##### | Brain - Caudate (basal ganglia | 109508840 | 109008840 | 110008840 | Yes |
| ENSG00000181754.6 | AMIGO1 | rs10857792 | 6.80E-07 | ##### | Brain - Spinal cord (cervical c- | 109508840 | 109008840 | 110008840 | Yes |

| | | | | | | | | | |
|---|---|---|---|---|---|---|---|---|---|
| ENSG00000181754.6 | AMIGO1 | rs4970762 | 6.90E-07 | ##### | Brain - Cerebellum | 109508840 | 109008840 | 110008840 | Yes |
| ENSG00000181754.6 | AMIGO1 | rs11101980 | 7.30E-07 | ##### | Brain - Substantia nigra | 109508840 | 109008840 | 110008840 | Yes |
| ENSG00000181754.6 | AMIGO1 | rs3754450 | 7.30E-07 | ##### | Brain - Substantia nigra | 109508840 | 109008840 | 110008840 | Yes |
| ENSG00000181754.6 | AMIGO1 | rs1010167 | 7.30E-07 | ##### | Brain - Substantia nigra | 109508840 | 109008840 | 110008840 | Yes |
| ENSG00000181754.6 | AMIGO1 | rs7536292 | 7.50E-07 | ##### | Brain - Cortex | 109508840 | 109008840 | 110008840 | Yes |
| ENSG00000181754.6 | AMIGO1 | rs413582 | 8.50E-07 | ##### | Brain - Caudate (basal ganglia | 109508840 | 109008840 | 110008840 | Yes |
| ENSG00000181754.6 | AMIGO1 | rs472737 | 8.80E-07 | ##### | Brain - Substantia nigra | 109508840 | 109008840 | 110008840 | Yes |
| ENSG00000181754.6 | AMIGO1 | rs2781555 | 8.80E-07 | ##### | Brain - Substantia nigra | 109508840 | 109008840 | 110008840 | Yes |
| ENSG00000181754.6 | AMIGO1 | rs7536291 | 8.80E-07 | ##### | Brain - Putamen (basal gangli | 109508840 | 109008840 | 110008840 | Yes |
| ENSG00000181754.6 | AMIGO1 | rs6681232 | 0.0000011 | ##### | Brain - Cerebellum | 109508840 | 109008840 | 110008840 | Yes |
| ENSG00000181754.6 | AMIGO1 | rs10857792 | 0.0000011 | ##### | Brain - Substantia nigra | 109508840 | 109008840 | 110008840 | Yes |
| ENSG00000181754.6 | AMIGO1 | rs12741221 | 0.0000012 | ##### | Brain - Substantia nigra | 109508840 | 109008840 | 110008840 | Yes |
| ENSG00000181754.6 | AMIGO1 | rs3738774 | 0.0000012 | ##### | Brain - Substantia nigra | 109508840 | 109008840 | 110008840 | Yes |
| ENSG00000181754.6 | AMIGO1 | rs1799875 | 0.0000012 | ##### | Brain - Putamen (basal gangli | 109508840 | 109008840 | 110008840 | Yes |
| ENSG00000181754.6 | AMIGO1 | rs35608606 | 0.0000012 | ##### | Brain - Caudate (basal ganglia | 109508840 | 109008840 | 110008840 | Yes |
| ENSG00000181754.6 | AMIGO1 | rs528815 | 0.0000012 | ##### | Brain - Substantia nigra | 109508840 | 109008840 | 110008840 | Yes |
| ENSG00000181754.6 | AMIGO1 | rs532432 | 0.0000012 | ##### | Brain - Substantia nigra | 109508840 | 109008840 | 110008840 | Yes |
| ENSG00000181754.6 | AMIGO1 | rs475381 | 0.0000012 | ##### | Brain - Substantia nigra | 109508840 | 109008840 | 110008840 | Yes |
| ENSG00000181754.6 | AMIGO1 | rs523300 | 0.0000012 | ##### | Brain - Substantia nigra | 109508840 | 109008840 | 110008840 | Yes |
| ENSG00000181754.6 | AMIGO1 | rs2692705 | 0.0000012 | ##### | Brain - Substantia nigra | 109508840 | 109008840 | 110008840 | Yes |
| ENSG00000181754.6 | AMIGO1 | rs2781558 | 0.0000012 | ##### | Brain - Substantia nigra | 109508840 | 109008840 | 110008840 | Yes |
| ENSG00000181754.6 | AMIGO1 | rs2781559 | 0.0000012 | ##### | Brain - Substantia nigra | 109508840 | 109008840 | 110008840 | Yes |
| ENSG00000181754.6 | AMIGO1 | rs532623 | 0.0000012 | ##### | Brain - Substantia nigra | 109508840 | 109008840 | 110008840 | Yes |
| ENSG00000181754.6 | AMIGO1 | rs561741 | 0.0000012 | ##### | Brain - Substantia nigra | 109508840 | 109008840 | 110008840 | Yes |
| ENSG00000181754.6 | AMIGO1 | rs4571992 | 0.0000012 | ##### | Brain - Substantia nigra | 109508840 | 109008840 | 110008840 | Yes |
| ENSG00000181754.6 | AMIGO1 | rs2301230 | 0.0000013 | ##### | Brain - Caudate (basal ganglia | 109508840 | 109008840 | 110008840 | Yes |
| ENSG00000181754.6 | AMIGO1 | rs2570972 | 0.0000014 | ##### | Brain - Hypothalamus | 109508840 | 109008840 | 110008840 | Yes |
| ENSG00000181754.6 | AMIGO1 | rs892510 | 0.0000015 | ##### | Brain - Hippocampus | 109508840 | 109008840 | 110008840 | Yes |
| ENSG00000181754.6 | AMIGO1 | rs93063 | 0.0000015 | ##### | Brain - Hippocampus | 109508840 | 109008840 | 110008840 | Yes |
| ENSG00000181754.6 | AMIGO1 | rs528815 | 0.0000016 | ##### | Brain - Hippocampus | 109508840 | 109008840 | 110008840 | Yes |
| ENSG00000181754.6 | AMIGO1 | rs532432 | 0.0000016 | ##### | Brain - Hippocampus | 109508840 | 109008840 | 110008840 | Yes |
| ENSG00000181754.6 | AMIGO1 | rs475381 | 0.0000016 | ##### | Brain - Hippocampus | 109508840 | 109008840 | 110008840 | Yes |

| | | | | | | | | | |
|---|---|---|---|---|---|---|---|---|---|
| ENSG00000181754.6 | AMIGO1 | rs523300 | 0.0000016 | ##### | Brain - Hippocampus | 109508840 | 109008840 | 110008840 | Yes |
| ENSG00000181754.6 | AMIGO1 | rs2692705 | 0.0000016 | ##### | Brain - Hippocampus | 109508840 | 109008840 | 110008840 | Yes |
| ENSG00000181754.6 | AMIGO1 | rs2781558 | 0.0000016 | ##### | Brain - Hippocampus | 109508840 | 109008840 | 110008840 | Yes |
| ENSG00000181754.6 | AMIGO1 | rs2781559 | 0.0000016 | ##### | Brain - Hippocampus | 109508840 | 109008840 | 110008840 | Yes |
| ENSG00000181754.6 | AMIGO1 | rs532623 | 0.0000016 | ##### | Brain - Hippocampus | 109508840 | 109008840 | 110008840 | Yes |
| ENSG00000181754.6 | AMIGO1 | rs561741 | 0.0000016 | ##### | Brain - Hippocampus | 109508840 | 109008840 | 110008840 | Yes |
| ENSG00000181754.6 | AMIGO1 | rs4571992 | 0.0000016 | ##### | Brain - Hippocampus | 109508840 | 109008840 | 110008840 | Yes |
| ENSG00000181754.6 | AMIGO1 | rs35608606 | 0.0000016 | ##### | Brain - Cortex | 109508840 | 109008840 | 110008840 | Yes |
| ENSG00000181754.6 | AMIGO1 | rs655246 | 0.0000017 | ##### | Brain - Caudate (basal ganglia | 109508840 | 109008840 | 110008840 | Yes |
| ENSG00000181754.6 | AMIGO1 | rs11101976 | 0.0000018 | ##### | Brain - Substantia nigra | 109508840 | 109008840 | 110008840 | Yes |
| ENSG00000181754.6 | AMIGO1 | rs12066164 | 0.0000019 | ##### | Brain - Hippocampus | 109508840 | 109008840 | 110008840 | Yes |
| ENSG00000181754.6 | AMIGO1 | rs2692706 | 0.0000019 | ##### | Brain - Hippocampus | 109508840 | 109008840 | 110008840 | Yes |
| ENSG00000181754.6 | AMIGO1 | rs892510 | 0.0000019 | ##### | Brain - Substantia nigra | 109508840 | 109008840 | 110008840 | Yes |
| ENSG00000181754.6 | AMIGO1 | rs93063 | 0.0000019 | ##### | Brain - Substantia nigra | 109508840 | 109008840 | 110008840 | Yes |
| ENSG00000181754.6 | AMIGO1 | rs10857792 | 0.0000019 | ##### | Brain - Putamen (basal gangli | 109508840 | 109008840 | 110008840 | Yes |
| ENSG00000181754.6 | AMIGO1 | rs6684193 | 0.0000019 | ##### | Brain - Substantia nigra | 109508840 | 109008840 | 110008840 | Yes |
| ENSG00000181754.6 | AMIGO1 | rs6684194 | 0.0000019 | ##### | Brain - Substantia nigra | 109508840 | 109008840 | 110008840 | Yes |
| ENSG00000181754.6 | AMIGO1 | rs11101966 | 0.0000019 | ##### | Brain - Substantia nigra | 109508840 | 109008840 | 110008840 | Yes |
| ENSG00000181754.6 | AMIGO1 | rs553689 | 0.0000019 | ##### | Brain - Substantia nigra | 109508840 | 109008840 | 110008840 | Yes |
| ENSG00000181754.6 | AMIGO1 | rs534135 | 0.0000019 | ##### | Brain - Substantia nigra | 109508840 | 109008840 | 110008840 | Yes |
| ENSG00000181754.6 | AMIGO1 | rs2781556 | 0.0000019 | ##### | Brain - Substantia nigra | 109508840 | 109008840 | 110008840 | Yes |
| ENSG00000181754.6 | AMIGO1 | rs528077 | 0.0000019 | ##### | Brain - Substantia nigra | 109508840 | 109008840 | 110008840 | Yes |
| ENSG00000181754.6 | AMIGO1 | rs529773 | 0.0000019 | ##### | Brain - Substantia nigra | 109508840 | 109008840 | 110008840 | Yes |
| ENSG00000181754.6 | AMIGO1 | rs2570973 | 0.0000019 | ##### | Brain - Substantia nigra | 109508840 | 109008840 | 110008840 | Yes |
| ENSG00000181754.6 | AMIGO1 | rs2781557 | 0.0000019 | ##### | Brain - Substantia nigra | 109508840 | 109008840 | 110008840 | Yes |
| ENSG00000181754.6 | AMIGO1 | rs538654 | 0.0000019 | ##### | Brain - Substantia nigra | 109508840 | 109008840 | 110008840 | Yes |
| ENSG00000181754.6 | AMIGO1 | rs543021 | 0.0000019 | ##### | Brain - Substantia nigra | 109508840 | 109008840 | 110008840 | Yes |
| ENSG00000181754.6 | AMIGO1 | rs543046 | 0.0000019 | ##### | Brain - Substantia nigra | 109508840 | 109008840 | 110008840 | Yes |
| ENSG00000181754.6 | AMIGO1 | rs2692703 | 0.0000019 | ##### | Brain - Substantia nigra | 109508840 | 109008840 | 110008840 | Yes |
| ENSG00000181754.6 | AMIGO1 | rs2692704 | 0.0000019 | ##### | Brain - Substantia nigra | 109508840 | 109008840 | 110008840 | Yes |
| ENSG00000181754.6 | AMIGO1 | rs474234 | 0.0000019 | ##### | Brain - Substantia nigra | 109508840 | 109008840 | 110008840 | Yes |
| ENSG00000181754.6 | AMIGO1 | rs558370 | 0.0000019 | ##### | Brain - Substantia nigra | 109508840 | 109008840 | 110008840 | Yes |

| | | | | | | | | | |
|---|---|---|---|---|---|---|---|---|---|
| ENSG00000181754.6 | AMIGO1 | rs732715 | 0.0000019 | ##### | Brain - Substantia nigra | 109508840 | 109008840 | 110008840 | Yes |
| ENSG00000181754.6 | AMIGO1 | rs4970762 | 0.000002 | ##### | Brain - Caudate (basal ganglia | 109508840 | 109008840 | 110008840 | Yes |
| ENSG00000181754.6 | AMIGO1 | rs11803546 | 0.0000021 | ##### | Brain - Spinal cord (cervical c- | 109508840 | 109008840 | 110008840 | Yes |
| ENSG00000181754.6 | AMIGO1 | rs57487258 | 0.0000021 | ##### | Brain - Cerebellum | 109508840 | 109008840 | 110008840 | Yes |
| ENSG00000181754.6 | AMIGO1 | rs34073826 | 0.0000021 | ##### | Brain - Cerebellum | 109508840 | 109008840 | 110008840 | Yes |
| ENSG00000181754.6 | AMIGO1 | rs6698913 | 0.0000021 | ##### | Brain - Cerebellum | 109508840 | 109008840 | 110008840 | Yes |
| ENSG00000181754.6 | AMIGO1 | rs7525563 | 0.0000021 | ##### | Brain - Cerebellum | 109508840 | 109008840 | 110008840 | Yes |
| ENSG00000181754.6 | AMIGO1 | rs6537803 | 0.0000021 | ##### | Brain - Cerebellum | 109508840 | 109008840 | 110008840 | Yes |
| ENSG00000181754.6 | AMIGO1 | rs11803546 | 0.0000023 | ##### | Brain - Caudate (basal ganglia | 109508840 | 109008840 | 110008840 | Yes |
| ENSG00000181754.6 | AMIGO1 | rs12066164 | 0.0000023 | ##### | Brain - Substantia nigra | 109508840 | 109008840 | 110008840 | Yes |
| ENSG00000181754.6 | AMIGO1 | rs2692706 | 0.0000023 | ##### | Brain - Substantia nigra | 109508840 | 109008840 | 110008840 | Yes |
| ENSG00000181754.6 | AMIGO1 | rs413582 | 0.0000023 | ##### | Brain - Substantia nigra | 109508840 | 109008840 | 110008840 | Yes |
| ENSG00000181754.6 | AMIGO1 | rs10127988 | 0.0000024 | ##### | Brain - Putamen (basal gangli | 109508840 | 109008840 | 110008840 | Yes |
| ENSG00000181754.6 | AMIGO1 | rs2301230 | 0.0000025 | ##### | Brain - Spinal cord (cervical c- | 109508840 | 109008840 | 110008840 | Yes |
| ENSG00000181754.6 | AMIGO1 | rs11803546 | 0.0000026 | ##### | Brain - Putamen (basal gangli | 109508840 | 109008840 | 110008840 | Yes |
| ENSG00000181754.6 | AMIGO1 | rs6669802 | 0.0000027 | ##### | Brain - Hippocampus | 109508840 | 109008840 | 110008840 | Yes |
| ENSG00000181754.6 | AMIGO1 | rs684044 | 0.0000028 | ##### | Brain - Caudate (basal ganglia | 109508840 | 109008840 | 110008840 | Yes |
| ENSG00000181754.6 | AMIGO1 | rs7553095 | 0.0000028 | ##### | Brain - Spinal cord (cervical c- | 109508840 | 109008840 | 110008840 | Yes |
| ENSG00000181754.6 | AMIGO1 | rs7536291 | 0.000003 | ##### | Brain - Caudate (basal ganglia | 109508840 | 109008840 | 110008840 | Yes |
| ENSG00000181754.6 | AMIGO1 | rs492557 | 0.000003 | ##### | Brain - Caudate (basal ganglia | 109508840 | 109008840 | 110008840 | Yes |
| ENSG00000181754.6 | AMIGO1 | rs146671905 | 0.000003 | ##### | Brain - Putamen (basal gangli | 109508840 | 109008840 | 110008840 | Yes |
| ENSG00000181754.6 | AMIGO1 | rs495562 | 0.0000034 | ##### | Brain - Spinal cord (cervical c- | 109508840 | 109008840 | 110008840 | Yes |
| ENSG00000181754.6 | AMIGO1 | rs7553095 | 0.0000034 | ##### | Brain - Substantia nigra | 109508840 | 109008840 | 110008840 | Yes |
| ENSG00000181754.6 | AMIGO1 | rs2301230 | 0.0000037 | ##### | Brain - Substantia nigra | 109508840 | 109008840 | 110008840 | Yes |
| ENSG00000181754.6 | AMIGO1 | rs35972691 | 0.0000039 | ##### | Brain - Putamen (basal gangli | 109508840 | 109008840 | 110008840 | Yes |
| ENSG00000181754.6 | AMIGO1 | rs651649 | 0.0000039 | ##### | Brain - Caudate (basal ganglia | 109508840 | 109008840 | 110008840 | Yes |
| ENSG00000181754.6 | AMIGO1 | rs10127988 | 0.000004 | ##### | Brain - Substantia nigra | 109508840 | 109008840 | 110008840 | Yes |
| ENSG00000181754.6 | AMIGO1 | rs12745189 | 0.000004 | ##### | Brain - Substantia nigra | 109508840 | 109008840 | 110008840 | Yes |
| ENSG00000181754.6 | AMIGO1 | rs10494041 | 0.0000042 | ##### | Brain - Spinal cord (cervical c- | 109508840 | 109008840 | 110008840 | Yes |
| ENSG00000181754.6 | AMIGO1 | rs11803546 | 0.0000044 | ##### | Brain - Substantia nigra | 109508840 | 109008840 | 110008840 | Yes |
| ENSG00000181754.6 | AMIGO1 | rs4970764 | 0.0000045 | ##### | Brain - Substantia nigra | 109508840 | 109008840 | 110008840 | Yes |
| ENSG00000181754.6 | AMIGO1 | rs2301230 | 0.0000046 | ##### | Brain - Putamen (basal gangli | 109508840 | 109008840 | 110008840 | Yes |

| | | | | | | | | | |
|---|---|---|---|---|---|---|---|---|---|
| ENSG00000181754.6 | AMIGO1 | rs4970843 | 0.0000046 | ##### | Brain - Cerebellum | 109508840 | 109008840 | 110008840 | Yes |
| ENSG00000181754.6 | AMIGO1 | rs4970843 | 0.0000046 | ##### | Brain - Substantia nigra | 109508840 | 109008840 | 110008840 | Yes |
| ENSG00000181754.6 | AMIGO1 | rs35608606 | 0.0000047 | ##### | Brain - Hippocampus | 109508840 | 109008840 | 110008840 | Yes |
| ENSG00000181754.6 | AMIGO1 | rs528199 | 0.0000051 | ##### | Brain - Amygdala | 109508840 | 109008840 | 110008840 | Yes |
| ENSG00000181754.6 | AMIGO1 | rs4970764 | 0.0000053 | ##### | Brain - Hippocampus | 109508840 | 109008840 | 110008840 | Yes |
| ENSG00000181754.6 | AMIGO1 | rs4970762 | 0.0000054 | ##### | Brain - Putamen (basal gangli | 109508840 | 109008840 | 110008840 | Yes |
| ENSG00000181754.6 | AMIGO1 | rs35608606 | 0.0000055 | ##### | Brain - Spinal cord (cervical c- | 109508840 | 109008840 | 110008840 | Yes |
| ENSG00000181754.6 | AMIGO1 | rs2297797 | 0.0000059 | ##### | Brain - Putamen (basal gangli | 109508840 | 109008840 | 110008840 | Yes |
| ENSG00000181754.6 | AMIGO1 | rs1799875 | 0.000006 | ##### | Brain - Substantia nigra | 109508840 | 109008840 | 110008840 | Yes |
| ENSG00000181754.6 | AMIGO1 | rs11351199 | 0.000006 | ##### | Brain - Cerebellum | 109508840 | 109008840 | 110008840 | Yes |
| ENSG00000181754.6 | AMIGO1 | rs11803546 | 0.0000062 | ##### | Brain - Hippocampus | 109508840 | 109008840 | 110008840 | Yes |
| ENSG00000181754.6 | AMIGO1 | rs12049330 | 0.0000065 | ##### | Brain - Spinal cord (cervical c- | 109508840 | 109008840 | 110008840 | Yes |
| ENSG00000181754.6 | AMIGO1 | rs7553095 | 0.0000069 | ##### | Brain - Caudate (basal ganglia | 109508840 | 109008840 | 110008840 | Yes |
| ENSG00000181754.6 | AMIGO1 | rs12044778 | 0.0000072 | ##### | Brain - Amygdala | 109508840 | 109008840 | 110008840 | Yes |
| ENSG00000181754.6 | AMIGO1 | rs528199 | 0.0000076 | ##### | Brain - Hypothalamus | 109508840 | 109008840 | 110008840 | Yes |
| ENSG00000181754.6 | AMIGO1 | rs2301230 | 0.0000077 | ##### | Brain - Hippocampus | 109508840 | 109008840 | 110008840 | Yes |
| ENSG00000181754.6 | AMIGO1 | rs10857792 | 0.0000081 | ##### | Brain - Hippocampus | 109508840 | 109008840 | 110008840 | Yes |
| ENSG00000181754.6 | AMIGO1 | rs10858096 | 0.0000087 | ##### | Brain - Cerebellum | 109508840 | 109008840 | 110008840 | Yes |
| ENSG00000181754.6 | AMIGO1 | rs10858096 | 0.00001 | ##### | Brain - Caudate (basal ganglia | 109508840 | 109008840 | 110008840 | Yes |
| ENSG00000181754.6 | AMIGO1 | rs11101965 | 0.00001 | ##### | Brain - Cortex | 109508840 | 109008840 | 110008840 | Yes |
| ENSG00000181754.6 | AMIGO1 | rs11101967 | 0.000011 | ##### | Brain - Cerebellum | 109508840 | 109008840 | 110008840 | Yes |
| ENSG00000181754.6 | AMIGO1 | rs3820667 | 0.000011 | ##### | Brain - Cerebellum | 109508840 | 109008840 | 110008840 | Yes |
| ENSG00000181754.6 | AMIGO1 | rs688498 | 0.000012 | ##### | Brain - Cerebellum | 109508840 | 109008840 | 110008840 | Yes |
| ENSG00000181754.6 | AMIGO1 | rs472737 | 0.000012 | ##### | Brain - Hippocampus | 109508840 | 109008840 | 110008840 | Yes |
| ENSG00000181754.6 | AMIGO1 | rs2781555 | 0.000012 | ##### | Brain - Hippocampus | 109508840 | 109008840 | 110008840 | Yes |
| ENSG00000181754.6 | AMIGO1 | rs35972691 | 0.000012 | ##### | Brain - Hippocampus | 109508840 | 109008840 | 110008840 | Yes |
| ENSG00000181754.6 | AMIGO1 | rs17575742 | 0.000012 | ##### | Brain - Hypothalamus | 109508840 | 109008840 | 110008840 | Yes |
| ENSG00000181754.6 | AMIGO1 | rs17575798 | 0.000012 | ##### | Brain - Hypothalamus | 109508840 | 109008840 | 110008840 | Yes |
| ENSG00000181754.6 | AMIGO1 | rs12045311 | 0.000012 | ##### | Brain - Hypothalamus | 109508840 | 109008840 | 110008840 | Yes |
| ENSG00000181754.6 | AMIGO1 | rs3820667 | 0.000012 | ##### | Brain - Caudate (basal ganglia | 109508840 | 109008840 | 110008840 | Yes |
| ENSG00000181754.6 | AMIGO1 | rs11351199 | 0.000012 | ##### | Brain - Caudate (basal ganglia | 109508840 | 109008840 | 110008840 | Yes |
| ENSG00000181754.6 | AMIGO1 | rs34843008 | 0.000013 | ##### | Brain - Spinal cord (cervical c- | 109508840 | 109008840 | 110008840 | Yes |

| | | | | | | | | | |
|---|---|---|---|---|---|---|---|---|---|
| ENSG00000181754.6 | AMIGO1 | rs919958 | 0.000013 | ##### | Brain - Hippocampus | 109508840 | 109008840 | 110008840 | Yes |
| ENSG00000181754.6 | AMIGO1 | rs4970766 | 0.000013 | ##### | Brain - Cerebellum | 109508840 | 109008840 | 110008840 | Yes |
| ENSG00000181754.6 | AMIGO1 | rs464218 | 0.000013 | ##### | Brain - Cerebellum | 109508840 | 109008840 | 110008840 | Yes |
| ENSG00000181754.6 | AMIGO1 | rs4970843 | 0.000014 | ##### | Brain - Putamen (basal gangli | 109508840 | 109008840 | 110008840 | Yes |
| ENSG00000181754.6 | AMIGO1 | rs146671905 | 0.000015 | ##### | Brain - Substantia nigra | 109508840 | 109008840 | 110008840 | Yes |
| ENSG00000181754.6 | AMIGO1 | rs590502 | 0.000015 | ##### | Brain - Caudate (basal ganglia | 109508840 | 109008840 | 110008840 | Yes |
| ENSG00000181754.6 | AMIGO1 | rs495562 | 0.000015 | ##### | Brain - Substantia nigra | 109508840 | 109008840 | 110008840 | Yes |
| ENSG00000181754.6 | AMIGO1 | rs646335 | 0.000015 | ##### | Brain - Caudate (basal ganglia | 109508840 | 109008840 | 110008840 | Yes |
| ENSG00000181754.6 | AMIGO1 | rs34843008 | 0.000015 | ##### | Brain - Cortex | 109508840 | 109008840 | 110008840 | Yes |
| ENSG00000181754.6 | AMIGO1 | rs595770 | 0.000015 | ##### | Brain - Cerebellum | 109508840 | 109008840 | 110008840 | Yes |
| ENSG00000181754.6 | AMIGO1 | rs72703294 | 0.000016 | ##### | Brain - Hippocampus | 109508840 | 109008840 | 110008840 | Yes |
| ENSG00000181754.6 | AMIGO1 | rs72703295 | 0.000016 | ##### | Brain - Hippocampus | 109508840 | 109008840 | 110008840 | Yes |
| ENSG00000181754.6 | AMIGO1 | rs518076 | 0.000016 | ##### | Brain - Hippocampus | 109508840 | 109008840 | 110008840 | Yes |
| ENSG00000181754.6 | AMIGO1 | rs55773147 | 0.000016 | ##### | Brain - Hippocampus | 109508840 | 109008840 | 110008840 | Yes |
| ENSG00000181754.6 | AMIGO1 | rs56367504 | 0.000016 | ##### | Brain - Hippocampus | 109508840 | 109008840 | 110008840 | Yes |
| ENSG00000181754.6 | AMIGO1 | rs144920545 | 0.000016 | ##### | Brain - Hippocampus | 109508840 | 109008840 | 110008840 | Yes |
| ENSG00000181754.6 | AMIGO1 | rs590502 | 0.000016 | ##### | Brain - Cerebellum | 109508840 | 109008840 | 110008840 | Yes |
| ENSG00000181754.6 | AMIGO1 | rs6684193 | 0.000019 | ##### | Brain - Amygdala | 109508840 | 109008840 | 110008840 | Yes |
| ENSG00000181754.6 | AMIGO1 | rs6684194 | 0.000019 | ##### | Brain - Amygdala | 109508840 | 109008840 | 110008840 | Yes |
| ENSG00000181754.6 | AMIGO1 | rs11101966 | 0.000019 | ##### | Brain - Amygdala | 109508840 | 109008840 | 110008840 | Yes |
| ENSG00000181754.6 | AMIGO1 | rs35972691 | 0.000019 | ##### | Brain - Cerebellum | 109508840 | 109008840 | 110008840 | Yes |
| ENSG00000181754.6 | AMIGO1 | rs116468196 | 0.00002 | ##### | Brain - Hippocampus | 109508840 | 109008840 | 110008840 | Yes |
| ENSG00000181754.6 | AMIGO1 | rs552101 | 0.00002 | ##### | Brain - Hippocampus | 109508840 | 109008840 | 110008840 | Yes |
| ENSG00000181754.6 | AMIGO1 | rs1279195 | 0.00002 | ##### | Brain - Hippocampus | 109508840 | 109008840 | 110008840 | Yes |
| ENSG00000181754.6 | AMIGO1 | rs1475693 | 0.00002 | ##### | Brain - Hippocampus | 109508840 | 109008840 | 110008840 | No |
| ENSG00000181754.6 | AMIGO1 | rs34571177 | 0.00002 | ##### | Brain - Spinal cord (cervical c- | 109508840 | 109008840 | 110008840 | Yes |
| ENSG00000181754.6 | AMIGO1 | rs11587687 | 0.00002 | ##### | Brain - Putamen (basal gangli | 109508840 | 109008840 | 110008840 | Yes |
| ENSG00000181754.6 | AMIGO1 | rs4970768 | 0.00002 | ##### | Brain - Putamen (basal gangli | 109508840 | 109008840 | 110008840 | Yes |
| ENSG00000181754.6 | AMIGO1 | rs77671818 | 0.000021 | ##### | Brain - Hippocampus | 109508840 | 109008840 | 110008840 | Yes |
| ENSG00000181754.6 | AMIGO1 | rs595770 | 0.000021 | ##### | Brain - Caudate (basal ganglia | 109508840 | 109008840 | 110008840 | Yes |
| ENSG00000181754.6 | AMIGO1 | rs2570972 | 0.000022 | ##### | Brain - Cortex | 109508840 | 109008840 | 110008840 | Yes |
| ENSG00000181754.6 | AMIGO1 | rs472737 | 0.000022 | ##### | Brain - Putamen (basal gangli | 109508840 | 109008840 | 110008840 | Yes |

| | | | | | | | | | |
|---|---|---|---|---|---|---|---|---|---|
| ENSG00000181754.6 | AMIGO1 | rs2781555 | 0.000022 | ##### | Brain - Putamen (basal gangli | 109508840 | 109008840 | 110008840 | Yes |
| ENSG00000181754.6 | AMIGO1 | rs4970843 | 0.000023 | ##### | Brain - Amygdala | 109508840 | 109008840 | 110008840 | Yes |
| ENSG00000181754.6 | AMIGO1 | rs11584134 | 0.000023 | ##### | Brain - Cerebellum | 109508840 | 109008840 | 110008840 | Yes |
| ENSG00000181754.6 | AMIGO1 | rs684044 | 0.000023 | ##### | Brain - Cerebellum | 109508840 | 109008840 | 110008840 | Yes |
| ENSG00000181754.6 | AMIGO1 | rs12066164 | 0.000025 | ##### | Brain - Cortex | 109508840 | 109008840 | 110008840 | Yes |
| ENSG00000181754.6 | AMIGO1 | rs2692706 | 0.000025 | ##### | Brain - Cortex | 109508840 | 109008840 | 110008840 | Yes |
| ENSG00000181754.6 | AMIGO1 | rs35375862 | 0.000025 | ##### | Brain - Spinal cord (cervical c- | 109508840 | 109008840 | 110008840 | Yes |
| ENSG00000181754.6 | AMIGO1 | rs413582 | 0.000026 | ##### | Brain - Amygdala | 109508840 | 109008840 | 110008840 | Yes |
| ENSG00000181754.6 | AMIGO1 | rs1799875 | 0.000026 | ##### | Brain - Spinal cord (cervical c- | 109508840 | 109008840 | 110008840 | Yes |
| ENSG00000181754.6 | AMIGO1 | rs4970837 | 0.000026 | ##### | Brain - Caudate (basal ganglia | 109508840 | 109008840 | 110008840 | Yes |
| ENSG00000181754.6 | AMIGO1 | rs413582 | 0.000026 | ##### | Brain - Cerebellum | 109508840 | 109008840 | 110008840 | Yes |
| ENSG00000181754.6 | AMIGO1 | rs2297797 | 0.000026 | ##### | Brain - Caudate (basal ganglia | 109508840 | 109008840 | 110008840 | Yes |
| ENSG00000181754.6 | AMIGO1 | rs1475694 | 0.000027 | ##### | Brain - Hippocampus | 109508840 | 109008840 | 110008840 | No |
| ENSG00000181754.6 | AMIGO1 | rs569670 | 0.000028 | ##### | Brain - Cerebellum | 109508840 | 109008840 | 110008840 | Yes |
| ENSG00000181754.6 | AMIGO1 | rs485660 | 0.000028 | ##### | Brain - Cerebellum | 109508840 | 109008840 | 110008840 | Yes |
| ENSG00000181754.6 | AMIGO1 | rs11101980 | 0.000029 | ##### | Brain - Caudate (basal ganglia | 109508840 | 109008840 | 110008840 | Yes |
| ENSG00000181754.6 | AMIGO1 | rs528199 | 0.000029 | ##### | Brain - Cortex | 109508840 | 109008840 | 110008840 | Yes |
| ENSG00000181754.6 | AMIGO1 | rs10127988 | 0.000031 | ##### | Brain - Caudate (basal ganglia | 109508840 | 109008840 | 110008840 | Yes |
| ENSG00000181754.6 | AMIGO1 | rs80025011 | 0.000031 | ##### | Brain - Cerebellum | 109508840 | 109008840 | 110008840 | Yes |
| ENSG00000181754.6 | AMIGO1 | rs61797132 | 0.000032 | ##### | Brain - Caudate (basal ganglia | 109508840 | 109008840 | 110008840 | Yes |
| ENSG00000181754.6 | AMIGO1 | rs611917 | 0.000032 | ##### | Brain - Caudate (basal ganglia | 109508840 | 109008840 | 110008840 | Yes |
| ENSG00000181754.6 | AMIGO1 | rs6684562 | 0.000033 | ##### | Brain - Cerebellum | 109508840 | 109008840 | 110008840 | Yes |
| ENSG00000181754.6 | AMIGO1 | rs657420 | 0.000035 | ##### | Brain - Cerebellum | 109508840 | 109008840 | 110008840 | Yes |
| ENSG00000181754.6 | AMIGO1 | rs9661614 | 0.000036 | ##### | Brain - Cerebellum | 109508840 | 109008840 | 110008840 | Yes |
| ENSG00000181754.6 | AMIGO1 | rs646335 | 0.000037 | ##### | Brain - Cerebellum | 109508840 | 109008840 | 110008840 | Yes |
| ENSG00000181754.6 | AMIGO1 | rs6680912 | 0.000037 | ##### | Brain - Cerebellum | 109508840 | 109008840 | 110008840 | Yes |
| ENSG00000181754.6 | AMIGO1 | rs6673727 | 0.000037 | ##### | Brain - Cerebellum | 109508840 | 109008840 | 110008840 | Yes |
| ENSG00000181754.6 | AMIGO1 | rs6677291 | 0.000037 | ##### | Brain - Cerebellum | 109508840 | 109008840 | 110008840 | Yes |
| ENSG00000181754.6 | AMIGO1 | rs111491315 | 0.000038 | ##### | Brain - Caudate (basal ganglia | 109508840 | 109008840 | 110008840 | Yes |
| ENSG00000181754.6 | AMIGO1 | rs4571992 | 0.000042 | ##### | Brain - Cortex | 109508840 | 109008840 | 110008840 | Yes |
| ENSG00000181754.6 | AMIGO1 | rs3820667 | 0.000043 | ##### | Brain - Putamen (basal gangli | 109508840 | 109008840 | 110008840 | Yes |
| ENSG00000181754.6 | AMIGO1 | rs413582 | 0.000043 | ##### | Brain - Putamen (basal gangli | 109508840 | 109008840 | 110008840 | Yes |

| | | | | | | | | | |
|---|---|---|---|---|---|---|---|---|---|
| ENSG00000181754.6 | AMIGO1 | rs464218 | 0.000045 | ##### | Brain - Caudate (basal ganglia | 109508840 | 109008840 | 110008840 | Yes |
| ENSG00000181754.6 | AMIGO1 | rs1765756 | 0.000046 | ##### | Brain - Cerebellum | 109508840 | 109008840 | 110008840 | Yes |
| ENSG00000181754.6 | AMIGO1 | rs1144595 | 0.000046 | ##### | Brain - Cerebellum | 109508840 | 109008840 | 110008840 | Yes |
| ENSG00000181754.6 | AMIGO1 | rs337299 | 0.000046 | ##### | Brain - Cortex | 109508840 | 109008840 | 110008840 | No |
| ENSG00000181754.6 | AMIGO1 | rs17035949 | 0.000047 | ##### | Brain - Cerebellum | 109508840 | 109008840 | 110008840 | Yes |
| ENSG00000181754.6 | AMIGO1 | rs1278286 | 0.000047 | ##### | Brain - Caudate (basal ganglia | 109508840 | 109008840 | 110008840 | Yes |
| ENSG00000181754.6 | AMIGO1 | rs2152924 | 0.000048 | ##### | Brain - Cerebellum | 109508840 | 109008840 | 110008840 | Yes |
| ENSG00000181754.6 | AMIGO1 | rs10857789 | 0.000048 | ##### | Brain - Cerebellum | 109508840 | 109008840 | 110008840 | Yes |
| ENSG00000181754.6 | AMIGO1 | rs146671905 | 0.000049 | ##### | Brain - Caudate (basal ganglia | 109508840 | 109008840 | 110008840 | Yes |
| ENSG00000181754.6 | AMIGO1 | rs3754450 | 0.000049 | ##### | Brain - Caudate (basal ganglia | 109508840 | 109008840 | 110008840 | Yes |
| ENSG00000181754.6 | AMIGO1 | rs11102967 | 0.00005 | ##### | Brain - Caudate (basal ganglia | 109508840 | 109008840 | 110008840 | Yes |
| ENSG00000181754.6 | AMIGO1 | rs3902354 | 0.00005 | ##### | Brain - Caudate (basal ganglia | 109508840 | 109008840 | 110008840 | Yes |
| ENSG00000181754.6 | AMIGO1 | rs1010167 | 0.000051 | ##### | Brain - Caudate (basal ganglia | 109508840 | 109008840 | 110008840 | Yes |
| ENSG00000181754.6 | AMIGO1 | rs12137189 | 0.000051 | ##### | Brain - Cerebellum | 109508840 | 109008840 | 110008840 | Yes |
| ENSG00000181754.6 | AMIGO1 | rs10776698 | 0.000051 | ##### | Brain - Cerebellum | 109508840 | 109008840 | 110008840 | Yes |
| ENSG00000181754.6 | AMIGO1 | rs12136063 | 0.000051 | ##### | Brain - Cerebellum | 109508840 | 109008840 | 110008840 | Yes |
| ENSG00000181754.6 | AMIGO1 | rs12753439 | 0.000051 | ##### | Brain - Cerebellum | 109508840 | 109008840 | 110008840 | Yes |
| ENSG00000181754.6 | AMIGO1 | rs9429519 | 0.000051 | ##### | Brain - Cerebellum | 109508840 | 109008840 | 110008840 | Yes |
| ENSG00000181754.6 | AMIGO1 | rs651649 | 0.000051 | ##### | Brain - Cerebellum | 109508840 | 109008840 | 110008840 | Yes |
| ENSG00000181754.6 | AMIGO1 | rs655246 | 0.000051 | ##### | Brain - Cerebellum | 109508840 | 109008840 | 110008840 | Yes |
| ENSG00000181754.6 | AMIGO1 | rs6688440 | 0.000053 | ##### | Brain - Cerebellum | 109508840 | 109008840 | 110008840 | Yes |
| ENSG00000181754.6 | AMIGO1 | rs532432 | 0.000056 | ##### | Brain - Cortex | 109508840 | 109008840 | 110008840 | Yes |
| ENSG00000181754.6 | AMIGO1 | rs2692705 | 0.000056 | ##### | Brain - Cortex | 109508840 | 109008840 | 110008840 | Yes |
| ENSG00000181754.6 | AMIGO1 | rs2781558 | 0.000056 | ##### | Brain - Cortex | 109508840 | 109008840 | 110008840 | Yes |
| ENSG00000181754.6 | AMIGO1 | rs2781559 | 0.000056 | ##### | Brain - Cortex | 109508840 | 109008840 | 110008840 | Yes |
| ENSG00000181754.6 | AMIGO1 | rs532623 | 0.000056 | ##### | Brain - Cortex | 109508840 | 109008840 | 110008840 | Yes |
| ENSG00000181754.6 | AMIGO1 | rs561741 | 0.000056 | ##### | Brain - Cortex | 109508840 | 109008840 | 110008840 | Yes |
| ENSG00000181754.6 | AMIGO1 | rs1608916 | 0.000061 | ##### | Brain - Cortex | 109508840 | 109008840 | 110008840 | Yes |
| ENSG00000181754.6 | AMIGO1 | rs72074532 | 0.000066 | ##### | Brain - Cerebellum | 109508840 | 109008840 | 110008840 | Yes |
| ENSG00000181754.6 | AMIGO1 | rs7550401 | 0.000066 | ##### | Brain - Cerebellum | 109508840 | 109008840 | 110008840 | Yes |
| ENSG00000181754.6 | AMIGO1 | rs11102972 | 0.000066 | ##### | Brain - Cerebellum | 109508840 | 109008840 | 110008840 | Yes |
| ENSG00000181754.6 | AMIGO1 | rs61798347 | 0.000067 | ##### | Brain - Cerebellum | 109508840 | 109008840 | 110008840 | Yes |

| | | | | | | | | | |
|---|---|---|---|---|---|---|---|---|---|
| ENSG00000181754.6 | AMIGO1 | rs2091329 | 0.000068 | ##### | Brain - Cerebellum | 109508840 | 109008840 | 110008840 | Yes |
| ENSG00000181754.6 | AMIGO1 | rs6677122 | 0.000069 | ##### | Brain - Cerebellum | 109508840 | 109008840 | 110008840 | Yes |
| ENSG00000181754.6 | AMIGO1 | rs1368988 | 0.000083 | ##### | Brain - Cerebellum | 109508840 | 109008840 | 110008840 | Yes |
| ENSG00000181754.6 | AMIGO1 | rs1764833 | 0.000084 | ##### | Brain - Cerebellum | 109508840 | 109008840 | 110008840 | Yes |
| ENSG00000181754.6 | AMIGO1 | rs4265455 | 0.000088 | ##### | Brain - Cerebellum | 109508840 | 109008840 | 110008840 | Yes |
| ENSG00000096433.10 | ITPR3 | rs9469554 | 0.0000023 | ##### | Brain - Cerebellar Hemisphere | 33688801 | 33188801 | 34188801 | Yes |
| ENSG00000096433.10 | ITPR3 | rs35095444 | 0.0000024 | ##### | Brain - Cerebellar Hemisphere | 33688801 | 33188801 | 34188801 | Yes |
| ENSG00000096433.10 | ITPR3 | rs34517899 | 0.000004 | ##### | Brain - Cerebellar Hemisphere | 33688801 | 33188801 | 34188801 | Yes |
| ENSG00000096433.10 | ITPR3 | rs12195707 | 0.000004 | ##### | Brain - Cerebellar Hemisphere | 33688801 | 33188801 | 34188801 | Yes |
| ENSG00000096433.10 | ITPR3 | rs10947426 | 0.000004 | ##### | Brain - Cerebellar Hemisphere | 33688801 | 33188801 | 34188801 | Yes |
| ENSG00000096433.10 | ITPR3 | rs7769838 | 0.0000054 | ##### | Brain - Caudate (basal ganglia | 33688801 | 33188801 | 34188801 | Yes |
| ENSG00000096433.10 | ITPR3 | rs10947417 | 0.000013 | ##### | Brain - Caudate (basal ganglia | 33688801 | 33188801 | 34188801 | Yes |
| ENSG00000096433.10 | ITPR3 | rs13192436 | 0.000013 | ##### | Brain - Caudate (basal ganglia | 33688801 | 33188801 | 34188801 | Yes |
| ENSG00000096433.10 | ITPR3 | rs2281831 | 0.000019 | ##### | Brain - Cerebellar Hemisphere | 33688801 | 33188801 | 34188801 | Yes |
| ENSG00000096433.10 | ITPR3 | rs71565393 | 0.000024 | ##### | Brain - Cerebellar Hemisphere | 33688801 | 33188801 | 34188801 | Yes |
| ENSG00000096433.10 | ITPR3 | rs12205667 | 0.000026 | ##### | Brain - Cerebellar Hemisphere | 33688801 | 33188801 | 34188801 | Yes |
| ENSG00000096433.10 | ITPR3 | rs12153870 | 0.000026 | ##### | Brain - Cerebellar Hemisphere | 33688801 | 33188801 | 34188801 | Yes |
| ENSG00000096433.10 | ITPR3 | rs13207407 | 0.000026 | ##### | Brain - Cerebellar Hemisphere | 33688801 | 33188801 | 34188801 | Yes |
| ENSG00000096433.10 | ITPR3 | rs36097803 | 0.000026 | ##### | Brain - Cerebellar Hemisphere | 33688801 | 33188801 | 34188801 | Yes |
| ENSG00000096433.10 | ITPR3 | rs13220421 | 0.000026 | ##### | Brain - Cerebellar Hemisphere | 33688801 | 33188801 | 34188801 | Yes |
| ENSG00000096433.10 | ITPR3 | rs35088242 | 0.000026 | ##### | Brain - Cerebellar Hemisphere | 33688801 | 33188801 | 34188801 | Yes |
| ENSG00000096433.10 | ITPR3 | rs10947415 | 0.000036 | ##### | Brain - Cerebellar Hemisphere | 33688801 | 33188801 | 34188801 | Yes |
| ENSG00000096433.10 | ITPR3 | rs13204162 | 0.000041 | ##### | Brain - Cerebellar Hemisphere | 33688801 | 33188801 | 34188801 | Yes |
| ENSG00000096433.10 | ITPR3 | rs12661400 | 0.000041 | ##### | Brain - Cerebellar Hemisphere | 33688801 | 33188801 | 34188801 | Yes |
| ENSG00000096433.10 | ITPR3 | rs35698850 | 0.000041 | ##### | Brain - Cerebellar Hemisphere | 33688801 | 33188801 | 34188801 | Yes |
| ENSG00000096433.10 | ITPR3 | rs2296341 | 0.000043 | ##### | Brain - Caudate (basal ganglia | 33688801 | 33188801 | 34188801 | Yes |
| ENSG00000096433.10 | ITPR3 | rs12205398 | 0.000048 | ##### | Brain - Cerebellar Hemisphere | 33688801 | 33188801 | 34188801 | Yes |
| ENSG00000096433.10 | ITPR3 | rs7769838 | 0.00005 | ##### | Brain - Cerebellar Hemisphere | 33688801 | 33188801 | 34188801 | Yes |
| ENSG00000096433.10 | ITPR3 | rs2281917 | 0.000052 | ##### | Brain - Cerebellar Hemisphere | 33688801 | 33188801 | 34188801 | Yes |
| ENSG00000096433.10 | ITPR3 | rs13219530 | 0.000052 | ##### | Brain - Cerebellar Hemisphere | 33688801 | 33188801 | 34188801 | Yes |
| ENSG00000096433.10 | ITPR3 | rs756137 | 0.000054 | ##### | Brain - Caudate (basal ganglia | 33688801 | 33188801 | 34188801 | Yes |
| ENSG00000096433.10 | ITPR3 | rs4713683 | 0.000054 | ##### | Brain - Caudate (basal ganglia | 33688801 | 33188801 | 34188801 | Yes |

| | | | | | | | | | |
|---|---|---|---|---|---|---|---|---|---|
| ENSG00000096433.10 | ITPR3 | rs2395449 | 0.000056 | ##### | Brain - Caudate (basal ganglia | 33688801 | 33188801 | 34188801 | Yes |
| ENSG00000096433.10 | ITPR3 | rs13195024 | 0.000058 | ##### | Brain - Cerebellar Hemisphere | 33688801 | 33188801 | 34188801 | Yes |
| ENSG00000096433.10 | ITPR3 | rs35329444 | 0.000058 | ##### | Brain - Cerebellar Hemisphere | 33688801 | 33188801 | 34188801 | Yes |
| ENSG00000096433.10 | ITPR3 | rs12200154 | 0.000058 | ##### | Brain - Cerebellar Hemisphere | 33688801 | 33188801 | 34188801 | Yes |
| ENSG00000096433.10 | ITPR3 | rs11755257 | 0.000058 | ##### | Brain - Cerebellar Hemisphere | 33688801 | 33188801 | 34188801 | Yes |
| ENSG00000096433.10 | ITPR3 | rs11758330 | 0.000058 | ##### | Brain - Cerebellar Hemisphere | 33688801 | 33188801 | 34188801 | Yes |
| ENSG00000096433.10 | ITPR3 | rs10947418 | 0.000058 | ##### | Brain - Cerebellar Hemisphere | 33688801 | 33188801 | 34188801 | Yes |
| ENSG00000096433.10 | ITPR3 | rs66731328 | 0.000058 | ##### | Brain - Cerebellar Hemisphere | 33688801 | 33188801 | 34188801 | Yes |
| ENSG00000096433.10 | ITPR3 | rs34942621 | 0.000062 | ##### | Brain - Cerebellar Hemisphere | 33688801 | 33188801 | 34188801 | Yes |
| ENSG00000096433.10 | ITPR3 | rs150517020 | 0.000066 | ##### | Brain - Cerebellar Hemisphere | 33688801 | 33188801 | 34188801 | Yes |
| ENSG00000072135.12 | PTPN18 | rs73959907 | 1.50E-10 | ##### | Brain - Cerebellar Hemisphere | 130356125 | 129856125 | 130856125 | No |
| ENSG00000072135.12 | PTPN18 | rs75923683 | 1.50E-10 | ##### | Brain - Cerebellar Hemisphere | 130356125 | 129856125 | 130856125 | No |
| ENSG00000072135.12 | PTPN18 | rs10205653 | 1.50E-10 | ##### | Brain - Cerebellar Hemisphere | 130356125 | 129856125 | 130856125 | No |
| ENSG00000072135.12 | PTPN18 | rs10197298 | 1.50E-10 | ##### | Brain - Cerebellar Hemisphere | 130356125 | 129856125 | 130856125 | No |
| ENSG00000072135.12 | PTPN18 | rs10178605 | 2.00E-09 | ##### | Brain - Cerebellar Hemisphere | 130356125 | 129856125 | 130856125 | Yes |
| ENSG00000072135.12 | PTPN18 | rs73959908 | 2.90E-09 | ##### | Brain - Cerebellar Hemisphere | 130356125 | 129856125 | 130856125 | No |
| ENSG00000072135.12 | PTPN18 | rs75274946 | 2.90E-09 | ##### | Brain - Cerebellar Hemisphere | 130356125 | 129856125 | 130856125 | No |
| ENSG00000072135.12 | PTPN18 | rs10198024 | 5.50E-09 | ##### | Brain - Cerebellar Hemisphere | 130356125 | 129856125 | 130856125 | No |
| ENSG00000072135.12 | PTPN18 | rs6743180 | 5.50E-09 | ##### | Brain - Cerebellar Hemisphere | 130356125 | 129856125 | 130856125 | No |
| ENSG00000072135.12 | PTPN18 | rs10179536 | 5.50E-09 | ##### | Brain - Cerebellar Hemisphere | 130356125 | 129856125 | 130856125 | No |
| ENSG00000072135.12 | PTPN18 | rs77333903 | 5.50E-09 | ##### | Brain - Cerebellar Hemisphere | 130356125 | 129856125 | 130856125 | No |
| ENSG00000072135.12 | PTPN18 | rs13382707 | 5.50E-09 | ##### | Brain - Cerebellar Hemisphere | 130356125 | 129856125 | 130856125 | No |
| ENSG00000072135.12 | PTPN18 | rs200623570 | 5.50E-09 | ##### | Brain - Cerebellar Hemisphere | 130356125 | 129856125 | 130856125 | No |
| ENSG00000072135.12 | PTPN18 | rs186106956 | 1.90E-08 | ##### | Brain - Cerebellar Hemisphere | 130356125 | 129856125 | 130856125 | Yes |
| ENSG00000072135.12 | PTPN18 | rs7570220 | 3.60E-08 | ##### | Brain - Cerebellum | 130356125 | 129856125 | 130856125 | Yes |
| ENSG00000072135.12 | PTPN18 | rs11899956 | 7.90E-08 | ##### | Brain - Cerebellar Hemisphere | 130356125 | 129856125 | 130856125 | No |
| ENSG00000072135.12 | PTPN18 | rs73959907 | 1.10E-07 | ##### | Brain - Frontal Cortex (BA9) | 130356125 | 129856125 | 130856125 | No |
| ENSG00000072135.12 | PTPN18 | rs75923683 | 1.10E-07 | ##### | Brain - Frontal Cortex (BA9) | 130356125 | 129856125 | 130856125 | No |
| ENSG00000072135.12 | PTPN18 | rs10205653 | 1.10E-07 | ##### | Brain - Frontal Cortex (BA9) | 130356125 | 129856125 | 130856125 | No |
| ENSG00000072135.12 | PTPN18 | rs10197298 | 1.10E-07 | ##### | Brain - Frontal Cortex (BA9) | 130356125 | 129856125 | 130856125 | No |
| ENSG00000072135.12 | PTPN18 | rs10198024 | 1.40E-07 | ##### | Brain - Frontal Cortex (BA9) | 130356125 | 129856125 | 130856125 | No |
| ENSG00000072135.12 | PTPN18 | rs6743180 | 1.40E-07 | ##### | Brain - Frontal Cortex (BA9) | 130356125 | 129856125 | 130856125 | No |

| | | | | | | | | | |
|---|---|---|---|---|---|---|---|---|---|
| ENSG00000072135.12 | PTPN18 | rs10179536 | 1.40E-07 | ##### | Brain - Frontal Cortex (BA9) | 130356125 | 129856125 | 130856125 | No |
| ENSG00000072135.12 | PTPN18 | rs77333903 | 1.40E-07 | ##### | Brain - Frontal Cortex (BA9) | 130356125 | 129856125 | 130856125 | No |
| ENSG00000072135.12 | PTPN18 | rs13382707 | 1.40E-07 | ##### | Brain - Frontal Cortex (BA9) | 130356125 | 129856125 | 130856125 | No |
| ENSG00000072135.12 | PTPN18 | rs200623570 | 1.40E-07 | ##### | Brain - Frontal Cortex (BA9) | 130356125 | 129856125 | 130856125 | No |
| ENSG00000072135.12 | PTPN18 | rs73959907 | 1.60E-07 | ##### | Brain - Caudate (basal ganglia | 130356125 | 129856125 | 130856125 | No |
| ENSG00000072135.12 | PTPN18 | rs75923683 | 1.60E-07 | ##### | Brain - Caudate (basal ganglia | 130356125 | 129856125 | 130856125 | No |
| ENSG00000072135.12 | PTPN18 | rs10205653 | 1.60E-07 | ##### | Brain - Caudate (basal ganglia | 130356125 | 129856125 | 130856125 | No |
| ENSG00000072135.12 | PTPN18 | rs10197298 | 1.60E-07 | ##### | Brain - Caudate (basal ganglia | 130356125 | 129856125 | 130856125 | No |
| ENSG00000072135.12 | PTPN18 | rs78951151 | 2.20E-07 | ##### | Brain - Cerebellar Hemisphere | 130356125 | 129856125 | 130856125 | Yes |
| ENSG00000072135.12 | PTPN18 | rs200623570 | 2.80E-07 | ##### | Brain - Nucleus accumbens (b | 130356125 | 129856125 | 130856125 | No |
| ENSG00000072135.12 | PTPN18 | rs7603950 | 3.50E-07 | ##### | Brain - Cerebellar Hemisphere | 130356125 | 129856125 | 130856125 | Yes |
| ENSG00000072135.12 | PTPN18 | rs7563097 | 3.50E-07 | ##### | Brain - Cerebellar Hemisphere | 130356125 | 129856125 | 130856125 | Yes |
| ENSG00000072135.12 | PTPN18 | rs114406893 | 4.40E-07 | ##### | Brain - Cortex | 130356125 | 129856125 | 130856125 | No |
| ENSG00000072135.12 | PTPN18 | rs80286115 | 4.40E-07 | ##### | Brain - Cortex | 130356125 | 129856125 | 130856125 | No |
| ENSG00000072135.12 | PTPN18 | rs7558521 | 8.90E-07 | ##### | Brain - Cortex | 130356125 | 129856125 | 130856125 | Yes |
| ENSG00000072135.12 | PTPN18 | rs4621198 | 8.90E-07 | ##### | Brain - Cortex | 130356125 | 129856125 | 130856125 | Yes |
| ENSG00000072135.12 | PTPN18 | rs142330150 | 8.90E-07 | ##### | Brain - Cortex | 130356125 | 129856125 | 130856125 | Yes |
| ENSG00000072135.12 | PTPN18 | rs17016623 | 9.40E-07 | ##### | Brain - Caudate (basal ganglia | 130356125 | 129856125 | 130856125 | Yes |
| ENSG00000072135.12 | PTPN18 | rs78951151 | 9.60E-07 | ##### | Brain - Cortex | 130356125 | 129856125 | 130856125 | Yes |
| ENSG00000072135.12 | PTPN18 | rs10198024 | 9.70E-07 | ##### | Brain - Caudate (basal ganglia | 130356125 | 129856125 | 130856125 | No |
| ENSG00000072135.12 | PTPN18 | rs6743180 | 9.70E-07 | ##### | Brain - Caudate (basal ganglia | 130356125 | 129856125 | 130856125 | No |
| ENSG00000072135.12 | PTPN18 | rs10179536 | 9.70E-07 | ##### | Brain - Caudate (basal ganglia | 130356125 | 129856125 | 130856125 | No |
| ENSG00000072135.12 | PTPN18 | rs77333903 | 9.70E-07 | ##### | Brain - Caudate (basal ganglia | 130356125 | 129856125 | 130856125 | No |
| ENSG00000072135.12 | PTPN18 | rs13382707 | 9.70E-07 | ##### | Brain - Caudate (basal ganglia | 130356125 | 129856125 | 130856125 | No |
| ENSG00000072135.12 | PTPN18 | rs200623570 | 9.70E-07 | ##### | Brain - Caudate (basal ganglia | 130356125 | 129856125 | 130856125 | No |
| ENSG00000072135.12 | PTPN18 | rs11901384 | 0.0000011 | ##### | Brain - Frontal Cortex (BA9) | 130356125 | 129856125 | 130856125 | Yes |
| ENSG00000072135.12 | PTPN18 | rs7603950 | 0.0000013 | ##### | Brain - Cortex | 130356125 | 129856125 | 130856125 | Yes |
| ENSG00000072135.12 | PTPN18 | rs7563097 | 0.0000013 | ##### | Brain - Cortex | 130356125 | 129856125 | 130856125 | Yes |
| ENSG00000072135.12 | PTPN18 | rs114154210 | 0.0000013 | ##### | Brain - Cerebellum | 130356125 | 129856125 | 130856125 | Yes |
| ENSG00000072135.12 | PTPN18 | rs78951151 | 0.0000018 | ##### | Brain - Caudate (basal ganglia | 130356125 | 129856125 | 130856125 | Yes |
| ENSG00000072135.12 | PTPN18 | rs76032333 | 0.0000018 | ##### | Brain - Cerebellum | 130356125 | 129856125 | 130856125 | Yes |
| ENSG00000072135.12 | PTPN18 | rs78951151 | 0.0000018 | ##### | Brain - Cerebellum | 130356125 | 129856125 | 130856125 | Yes |

| | | | | | | | | | |
|---|---|---|---|---|---|---|---|---|---|
| ENSG00000072135.12 | PTPN18 | rs114406893 | 0.0000024 | ##### | Brain - Cerebellar Hemisphere | 130356125 | 129856125 | 130856125 | No |
| ENSG00000072135.12 | PTPN18 | rs80286115 | 0.0000024 | ##### | Brain - Cerebellar Hemisphere | 130356125 | 129856125 | 130856125 | No |
| ENSG00000072135.12 | PTPN18 | rs17016623 | 0.0000025 | ##### | Brain - Cerebellum | 130356125 | 129856125 | 130856125 | Yes |
| ENSG00000072135.12 | PTPN18 | rs112657480 | 0.0000026 | ##### | Brain - Nucleus accumbens (b | 130356125 | 129856125 | 130856125 | Yes |
| ENSG00000072135.12 | PTPN18 | rs75196942 | 0.0000026 | ##### | Brain - Nucleus accumbens (b | 130356125 | 129856125 | 130856125 | Yes |
| ENSG00000072135.12 | PTPN18 | rs111387371 | 0.0000026 | ##### | Brain - Nucleus accumbens (b | 130356125 | 129856125 | 130856125 | Yes |
| ENSG00000072135.12 | PTPN18 | rs75069611 | 0.0000026 | ##### | Brain - Nucleus accumbens (b | 130356125 | 129856125 | 130856125 | Yes |
| ENSG00000072135.12 | PTPN18 | rs79648057 | 0.0000026 | ##### | Brain - Nucleus accumbens (b | 130356125 | 129856125 | 130856125 | Yes |
| ENSG00000072135.12 | PTPN18 | rs868110181 | 0.0000026 | ##### | Brain - Caudate (basal ganglia | 130356125 | 129856125 | 130856125 | Yes |
| ENSG00000072135.12 | PTPN18 | rs578029750 | 0.0000026 | ##### | Brain - Caudate (basal ganglia | 130356125 | 129856125 | 130856125 | Yes |
| ENSG00000072135.12 | PTPN18 | rs9967795 | 0.0000026 | ##### | Brain - Cerebellar Hemisphere | 130356125 | 129856125 | 130856125 | Yes |
| ENSG00000072135.12 | PTPN18 | rs9967796 | 0.0000026 | ##### | Brain - Cerebellar Hemisphere | 130356125 | 129856125 | 130856125 | Yes |
| ENSG00000072135.12 | PTPN18 | rs7570220 | 0.0000027 | ##### | Brain - Cerebellar Hemisphere | 130356125 | 129856125 | 130856125 | Yes |
| ENSG00000072135.12 | PTPN18 | rs530924882 | 0.0000028 | ##### | Brain - Cerebellar Hemisphere | 130356125 | 129856125 | 130856125 | Yes |
| ENSG00000072135.12 | PTPN18 | rs7558521 | 0.0000031 | ##### | Brain - Cerebellum | 130356125 | 129856125 | 130856125 | Yes |
| ENSG00000072135.12 | PTPN18 | rs4621198 | 0.0000031 | ##### | Brain - Cerebellum | 130356125 | 129856125 | 130856125 | Yes |
| ENSG00000072135.12 | PTPN18 | rs142330150 | 0.0000031 | ##### | Brain - Cerebellum | 130356125 | 129856125 | 130856125 | Yes |
| ENSG00000072135.12 | PTPN18 | rs74513274 | 0.0000031 | ##### | Brain - Cerebellum | 130356125 | 129856125 | 130856125 | Yes |
| ENSG00000072135.12 | PTPN18 | rs137992200 | 0.0000031 | ##### | Brain - Cerebellum | 130356125 | 129856125 | 130856125 | Yes |
| ENSG00000072135.12 | PTPN18 | rs147310902 | 0.0000031 | ##### | Brain - Cerebellum | 130356125 | 129856125 | 130856125 | Yes |
| ENSG00000072135.12 | PTPN18 | rs144193979 | 0.0000032 | ##### | Brain - Cerebellum | 130356125 | 129856125 | 130856125 | Yes |
| ENSG00000072135.12 | PTPN18 | rs146558998 | 0.0000032 | ##### | Brain - Cerebellum | 130356125 | 129856125 | 130856125 | Yes |
| ENSG00000072135.12 | PTPN18 | rs141587479 | 0.0000032 | ##### | Brain - Cerebellum | 130356125 | 129856125 | 130856125 | Yes |
| ENSG00000072135.12 | PTPN18 | rs17016591 | 0.0000032 | ##### | Brain - Cerebellum | 130356125 | 129856125 | 130856125 | Yes |
| ENSG00000072135.12 | PTPN18 | rs17016603 | 0.0000032 | ##### | Brain - Cerebellum | 130356125 | 129856125 | 130856125 | Yes |
| ENSG00000072135.12 | PTPN18 | rs6742510 | 0.0000032 | ##### | Brain - Cerebellum | 130356125 | 129856125 | 130856125 | Yes |
| ENSG00000072135.12 | PTPN18 | rs17016615 | 0.0000032 | ##### | Brain - Cerebellum | 130356125 | 129856125 | 130856125 | Yes |
| ENSG00000072135.12 | PTPN18 | rs17016616 | 0.0000032 | ##### | Brain - Cerebellum | 130356125 | 129856125 | 130856125 | Yes |
| ENSG00000072135.12 | PTPN18 | rs201214742 | 0.0000032 | ##### | Brain - Cerebellum | 130356125 | 129856125 | 130856125 | Yes |
| ENSG00000072135.12 | PTPN18 | rs17016621 | 0.0000032 | ##### | Brain - Cerebellum | 130356125 | 129856125 | 130856125 | Yes |
| ENSG00000072135.12 | PTPN18 | rs78951151 | 0.0000035 | ##### | Brain - Frontal Cortex (BA9) | 130356125 | 129856125 | 130856125 | Yes |
| ENSG00000072135.12 | PTPN18 | rs113347353 | 0.0000046 | ##### | Brain - Caudate (basal ganglia | 130356125 | 129856125 | 130856125 | No |

| | | | | | | | | | |
|---|---|---|---|---|---|---|---|---|---|
| ENSG00000072135.12 | PTPN18 | rs3739123 | 0.0000052 | ##### | Brain - Cerebellar Hemisphere | 130356125 | 129856125 | 130856125 | Yes |
| ENSG00000072135.12 | PTPN18 | rs3739122 | 0.0000052 | ##### | Brain - Cerebellar Hemisphere | 130356125 | 129856125 | 130856125 | Yes |
| ENSG00000072135.12 | PTPN18 | rs72854965 | 0.0000052 | ##### | Brain - Cerebellar Hemisphere | 130356125 | 129856125 | 130856125 | Yes |
| ENSG00000072135.12 | PTPN18 | rs144975008 | 0.0000053 | ##### | Brain - Caudate (basal ganglia | 130356125 | 129856125 | 130856125 | Yes |
| ENSG00000072135.12 | PTPN18 | rs116752374 | 0.0000053 | ##### | Brain - Caudate (basal ganglia | 130356125 | 129856125 | 130856125 | Yes |
| ENSG00000072135.12 | PTPN18 | rs112791004 | 0.0000053 | ##### | Brain - Caudate (basal ganglia | 130356125 | 129856125 | 130856125 | Yes |
| ENSG00000072135.12 | PTPN18 | rs111894685 | 0.0000053 | ##### | Brain - Caudate (basal ganglia | 130356125 | 129856125 | 130856125 | Yes |
| ENSG00000072135.12 | PTPN18 | rs112416166 | 0.0000053 | ##### | Brain - Caudate (basal ganglia | 130356125 | 129856125 | 130856125 | Yes |
| ENSG00000072135.12 | PTPN18 | rs113266675 | 0.0000053 | ##### | Brain - Caudate (basal ganglia | 130356125 | 129856125 | 130856125 | Yes |
| ENSG00000072135.12 | PTPN18 | rs113077512 | 0.0000053 | ##### | Brain - Caudate (basal ganglia | 130356125 | 129856125 | 130856125 | Yes |
| ENSG00000072135.12 | PTPN18 | rs143960243 | 0.0000053 | ##### | Brain - Caudate (basal ganglia | 130356125 | 129856125 | 130856125 | Yes |
| ENSG00000072135.12 | PTPN18 | rs112176365 | 0.0000053 | ##### | Brain - Caudate (basal ganglia | 130356125 | 129856125 | 130856125 | Yes |
| ENSG00000072135.12 | PTPN18 | rs6725079 | 0.0000053 | ##### | Brain - Caudate (basal ganglia | 130356125 | 129856125 | 130856125 | Yes |
| ENSG00000072135.12 | PTPN18 | rs544737002 | 0.0000053 | ##### | Brain - Caudate (basal ganglia | 130356125 | 129856125 | 130856125 | Yes |
| ENSG00000072135.12 | PTPN18 | rs553350244 | 0.0000053 | ##### | Brain - Caudate (basal ganglia | 130356125 | 129856125 | 130856125 | Yes |
| ENSG00000072135.12 | PTPN18 | rs114319834 | 0.0000053 | ##### | Brain - Caudate (basal ganglia | 130356125 | 129856125 | 130856125 | Yes |
| ENSG00000072135.12 | PTPN18 | rs111238220 | 0.0000053 | ##### | Brain - Caudate (basal ganglia | 130356125 | 129856125 | 130856125 | Yes |
| ENSG00000072135.12 | PTPN18 | rs79305276 | 0.0000053 | ##### | Brain - Caudate (basal ganglia | 130356125 | 129856125 | 130856125 | Yes |
| ENSG00000072135.12 | PTPN18 | rs144193979 | 0.0000053 | ##### | Brain - Caudate (basal ganglia | 130356125 | 129856125 | 130856125 | Yes |
| ENSG00000072135.12 | PTPN18 | rs146558998 | 0.0000053 | ##### | Brain - Caudate (basal ganglia | 130356125 | 129856125 | 130856125 | Yes |
| ENSG00000072135.12 | PTPN18 | rs76032333 | 0.0000053 | ##### | Brain - Caudate (basal ganglia | 130356125 | 129856125 | 130856125 | Yes |
| ENSG00000072135.12 | PTPN18 | rs141587479 | 0.0000053 | ##### | Brain - Caudate (basal ganglia | 130356125 | 129856125 | 130856125 | Yes |
| ENSG00000072135.12 | PTPN18 | rs17016591 | 0.0000053 | ##### | Brain - Caudate (basal ganglia | 130356125 | 129856125 | 130856125 | Yes |
| ENSG00000072135.12 | PTPN18 | rs17016603 | 0.0000053 | ##### | Brain - Caudate (basal ganglia | 130356125 | 129856125 | 130856125 | Yes |
| ENSG00000072135.12 | PTPN18 | rs6742510 | 0.0000053 | ##### | Brain - Caudate (basal ganglia | 130356125 | 129856125 | 130856125 | Yes |
| ENSG00000072135.12 | PTPN18 | rs17016615 | 0.0000053 | ##### | Brain - Caudate (basal ganglia | 130356125 | 129856125 | 130856125 | Yes |
| ENSG00000072135.12 | PTPN18 | rs17016616 | 0.0000053 | ##### | Brain - Caudate (basal ganglia | 130356125 | 129856125 | 130856125 | Yes |
| ENSG00000072135.12 | PTPN18 | rs201214742 | 0.0000053 | ##### | Brain - Caudate (basal ganglia | 130356125 | 129856125 | 130856125 | Yes |
| ENSG00000072135.12 | PTPN18 | rs17016621 | 0.0000053 | ##### | Brain - Caudate (basal ganglia | 130356125 | 129856125 | 130856125 | Yes |
| ENSG00000072135.12 | PTPN18 | rs11884245 | 0.0000054 | ##### | Brain - Cerebellar Hemisphere | 130356125 | 129856125 | 130856125 | Yes |
| ENSG00000072135.12 | PTPN18 | rs73960349 | 0.0000054 | ##### | Brain - Cerebellar Hemisphere | 130356125 | 129856125 | 130856125 | Yes |
| ENSG00000072135.12 | PTPN18 | rs11899834 | 0.0000054 | ##### | Brain - Cerebellar Hemisphere | 130356125 | 129856125 | 130856125 | No |

| | | | | | | | | | |
|---|---|---|---|---|---|---|---|---|---|
| ENSG00000072135.12 | PTPN18 | rs144975008 | 0.0000064 | ##### | Brain - Nucleus accumbens (b | 130356125 | 129856125 | 130856125 | Yes |
| ENSG00000072135.12 | PTPN18 | rs116752374 | 0.0000064 | ##### | Brain - Nucleus accumbens (b | 130356125 | 129856125 | 130856125 | Yes |
| ENSG00000072135.12 | PTPN18 | rs112791004 | 0.0000064 | ##### | Brain - Nucleus accumbens (b | 130356125 | 129856125 | 130856125 | Yes |
| ENSG00000072135.12 | PTPN18 | rs111894685 | 0.0000064 | ##### | Brain - Nucleus accumbens (b | 130356125 | 129856125 | 130856125 | Yes |
| ENSG00000072135.12 | PTPN18 | rs112416166 | 0.0000064 | ##### | Brain - Nucleus accumbens (b | 130356125 | 129856125 | 130856125 | Yes |
| ENSG00000072135.12 | PTPN18 | rs113266675 | 0.0000064 | ##### | Brain - Nucleus accumbens (b | 130356125 | 129856125 | 130856125 | Yes |
| ENSG00000072135.12 | PTPN18 | rs113077512 | 0.0000064 | ##### | Brain - Nucleus accumbens (b | 130356125 | 129856125 | 130856125 | Yes |
| ENSG00000072135.12 | PTPN18 | rs143960243 | 0.0000064 | ##### | Brain - Nucleus accumbens (b | 130356125 | 129856125 | 130856125 | Yes |
| ENSG00000072135.12 | PTPN18 | rs112176365 | 0.0000064 | ##### | Brain - Nucleus accumbens (b | 130356125 | 129856125 | 130856125 | Yes |
| ENSG00000072135.12 | PTPN18 | rs6725079 | 0.0000064 | ##### | Brain - Nucleus accumbens (b | 130356125 | 129856125 | 130856125 | Yes |
| ENSG00000072135.12 | PTPN18 | rs544737002 | 0.0000064 | ##### | Brain - Nucleus accumbens (b | 130356125 | 129856125 | 130856125 | Yes |
| ENSG00000072135.12 | PTPN18 | rs553350244 | 0.0000064 | ##### | Brain - Nucleus accumbens (b | 130356125 | 129856125 | 130856125 | Yes |
| ENSG00000072135.12 | PTPN18 | rs114319834 | 0.0000064 | ##### | Brain - Nucleus accumbens (b | 130356125 | 129856125 | 130856125 | Yes |
| ENSG00000072135.12 | PTPN18 | rs111238220 | 0.0000064 | ##### | Brain - Nucleus accumbens (b | 130356125 | 129856125 | 130856125 | Yes |
| ENSG00000072135.12 | PTPN18 | rs79305276 | 0.0000064 | ##### | Brain - Nucleus accumbens (b | 130356125 | 129856125 | 130856125 | Yes |
| ENSG00000072135.12 | PTPN18 | rs200512114 | 0.0000066 | ##### | Brain - Caudate (basal ganglia | 130356125 | 129856125 | 130856125 | Yes |
| ENSG00000072135.12 | PTPN18 | rs200512114 | 0.0000068 | ##### | Brain - Nucleus accumbens (b | 130356125 | 129856125 | 130856125 | Yes |
| ENSG00000072135.12 | PTPN18 | rs55702504 | 0.0000073 | ##### | Brain - Cortex | 130356125 | 129856125 | 130856125 | Yes |
| ENSG00000072135.12 | PTPN18 | rs73959908 | 0.0000077 | ##### | Brain - Frontal Cortex (BA9) | 130356125 | 129856125 | 130856125 | No |
| ENSG00000072135.12 | PTPN18 | rs75274946 | 0.0000077 | ##### | Brain - Frontal Cortex (BA9) | 130356125 | 129856125 | 130856125 | No |
| ENSG00000072135.12 | PTPN18 | rs200623570 | 0.0000087 | ##### | Brain - Cortex | 130356125 | 129856125 | 130856125 | No |
| ENSG00000072135.12 | PTPN18 | rs560871441 | 0.0000091 | ##### | Brain - Cerebellar Hemisphere | 130356125 | 129856125 | 130856125 | Yes |
| ENSG00000072135.12 | PTPN18 | rs17016623 | 0.0000097 | ##### | Brain - Nucleus accumbens (b | 130356125 | 129856125 | 130856125 | Yes |
| ENSG00000072135.12 | PTPN18 | rs150068082 | 0.00001 | ##### | Brain - Cerebellum | 130356125 | 129856125 | 130856125 | Yes |
| ENSG00000072135.12 | PTPN18 | rs116762420 | 0.00001 | ##### | Brain - Cerebellum | 130356125 | 129856125 | 130856125 | Yes |
| ENSG00000072135.12 | PTPN18 | rs114478505 | 0.00001 | ##### | Brain - Cerebellum | 130356125 | 129856125 | 130856125 | Yes |
| ENSG00000072135.12 | PTPN18 | rs7567015 | 0.00001 | ##### | Brain - Cerebellum | 130356125 | 129856125 | 130856125 | Yes |
| ENSG00000072135.12 | PTPN18 | rs76751313 | 0.00001 | ##### | Brain - Cerebellum | 130356125 | 129856125 | 130856125 | Yes |
| ENSG00000072135.12 | PTPN18 | rs79166986 | 0.00001 | ##### | Brain - Cortex | 130356125 | 129856125 | 130856125 | No |
| ENSG00000072135.12 | PTPN18 | rs72854962 | 0.000011 | ##### | Brain - Cerebellar Hemisphere | 130356125 | 129856125 | 130856125 | Yes |
| ENSG00000072135.12 | PTPN18 | rs62161856 | 0.000011 | ##### | Brain - Cortex | 130356125 | 129856125 | 130856125 | Yes |
| ENSG00000072135.12 | PTPN18 | rs73959907 | 0.000011 | ##### | Brain - Cerebellum | 130356125 | 129856125 | 130856125 | No |

| | | | | | | | | | |
|---|---|---|---|---|---|---|---|---|---|
| ENSG00000072135.12 | PTPN18 | rs75923683 | 0.000011 | ##### | Brain - Cerebellum | 130356125 | 129856125 | 130856125 | No |
| ENSG00000072135.12 | PTPN18 | rs10205653 | 0.000011 | ##### | Brain - Cerebellum | 130356125 | 129856125 | 130856125 | No |
| ENSG00000072135.12 | PTPN18 | rs10197298 | 0.000011 | ##### | Brain - Cerebellum | 130356125 | 129856125 | 130856125 | No |
| ENSG00000072135.12 | PTPN18 | rs13421181 | 0.000011 | ##### | Brain - Caudate (basal ganglia | 130356125 | 129856125 | 130856125 | No |
| ENSG00000072135.12 | PTPN18 | rs78837375 | 0.000011 | ##### | Brain - Caudate (basal ganglia | 130356125 | 129856125 | 130856125 | No |
| ENSG00000072135.12 | PTPN18 | rs114921595 | 0.000011 | ##### | Brain - Caudate (basal ganglia | 130356125 | 129856125 | 130856125 | No |
| ENSG00000072135.12 | PTPN18 | rs13413167 | 0.000011 | ##### | Brain - Caudate (basal ganglia | 130356125 | 129856125 | 130856125 | No |
| ENSG00000072135.12 | PTPN18 | rs57783690 | 0.000011 | ##### | Brain - Caudate (basal ganglia | 130356125 | 129856125 | 130856125 | No |
| ENSG00000072135.12 | PTPN18 | rs77712455 | 0.000011 | ##### | Brain - Caudate (basal ganglia | 130356125 | 129856125 | 130856125 | No |
| ENSG00000072135.12 | PTPN18 | rs10153766 | 0.000011 | ##### | Brain - Caudate (basal ganglia | 130356125 | 129856125 | 130856125 | No |
| ENSG00000072135.12 | PTPN18 | rs10183938 | 0.000011 | ##### | Brain - Caudate (basal ganglia | 130356125 | 129856125 | 130856125 | No |
| ENSG00000072135.12 | PTPN18 | rs13409036 | 0.000011 | ##### | Brain - Caudate (basal ganglia | 130356125 | 129856125 | 130856125 | No |
| ENSG00000072135.12 | PTPN18 | rs77274625 | 0.000011 | ##### | Brain - Caudate (basal ganglia | 130356125 | 129856125 | 130856125 | No |
| ENSG00000072135.12 | PTPN18 | rs13402442 | 0.000011 | ##### | Brain - Caudate (basal ganglia | 130356125 | 129856125 | 130856125 | Yes |
| ENSG00000072135.12 | PTPN18 | rs75986443 | 0.000011 | ##### | Brain - Caudate (basal ganglia | 130356125 | 129856125 | 130856125 | Yes |
| ENSG00000072135.12 | PTPN18 | rs6717894 | 0.000011 | ##### | Brain - Cerebellar Hemisphere | 130356125 | 129856125 | 130856125 | Yes |
| ENSG00000072135.12 | PTPN18 | rs72614442 | 0.000011 | ##### | Brain - Cerebellar Hemisphere | 130356125 | 129856125 | 130856125 | Yes |
| ENSG00000072135.12 | PTPN18 | rs7603950 | 0.000012 | ##### | Brain - Frontal Cortex (BA9) | 130356125 | 129856125 | 130856125 | Yes |
| ENSG00000072135.12 | PTPN18 | rs7563097 | 0.000012 | ##### | Brain - Frontal Cortex (BA9) | 130356125 | 129856125 | 130856125 | Yes |
| ENSG00000072135.12 | PTPN18 | rs868742405 | 0.000013 | ##### | Brain - Cortex | 130356125 | 129856125 | 130856125 | Yes |
| ENSG00000072135.12 | PTPN18 | rs17016623 | 0.000013 | ##### | Brain - Cerebellar Hemisphere | 130356125 | 129856125 | 130856125 | Yes |
| ENSG00000072135.12 | PTPN18 | rs10209164 | 0.000014 | ##### | Brain - Frontal Cortex (BA9) | 130356125 | 129856125 | 130856125 | Yes |
| ENSG00000072135.12 | PTPN18 | rs150745593 | 0.000015 | ##### | Brain - Cerebellar Hemisphere | 130356125 | 129856125 | 130856125 | Yes |
| ENSG00000072135.12 | PTPN18 | rs7603950 | 0.000015 | ##### | Brain - Cerebellum | 130356125 | 129856125 | 130856125 | Yes |
| ENSG00000072135.12 | PTPN18 | rs7563097 | 0.000015 | ##### | Brain - Cerebellum | 130356125 | 129856125 | 130856125 | Yes |
| ENSG00000072135.12 | PTPN18 | rs752028547 | 0.000015 | ##### | Brain - Cortex | 130356125 | 129856125 | 130856125 | Yes |
| ENSG00000072135.12 | PTPN18 | rs111798470 | 0.000016 | ##### | Brain - Nucleus accumbens (b | 130356125 | 129856125 | 130856125 | Yes |
| ENSG00000072135.12 | PTPN18 | rs75066561 | 0.000016 | ##### | Brain - Cerebellar Hemisphere | 130356125 | 129856125 | 130856125 | Yes |
| ENSG00000072135.12 | PTPN18 | rs10179902 | 0.000016 | ##### | Brain - Cerebellar Hemisphere | 130356125 | 129856125 | 130856125 | Yes |
| ENSG00000072135.12 | PTPN18 | rs76847861 | 0.000016 | ##### | Brain - Cerebellum | 130356125 | 129856125 | 130856125 | Yes |
| ENSG00000072135.12 | PTPN18 | rs77516789 | 0.000016 | ##### | Brain - Cerebellum | 130356125 | 129856125 | 130856125 | Yes |
| ENSG00000072135.12 | PTPN18 | rs75404769 | 0.000016 | ##### | Brain - Cerebellum | 130356125 | 129856125 | 130856125 | Yes |

| | | | | | | | | | |
|---|---|---|---|---|---|---|---|---|---|
| ENSG00000072135.12 | PTPN18 | rs10209164 | 0.000016 | ##### | Brain - Nucleus accumbens (b | 130356125 | 129856125 | 130856125 | Yes |
| ENSG00000072135.12 | PTPN18 | rs28764204 | 0.000017 | ##### | Brain - Cerebellum | 130356125 | 129856125 | 130856125 | Yes |
| ENSG00000072135.12 | PTPN18 | rs10178605 | 0.000017 | ##### | Brain - Frontal Cortex (BA9) | 130356125 | 129856125 | 130856125 | Yes |
| ENSG00000072135.12 | PTPN18 | rs112759995 | 0.000018 | ##### | Brain - Cerebellar Hemisphere | 130356125 | 129856125 | 130856125 | No |
| ENSG00000072135.12 | PTPN18 | rs72978683 | 0.000019 | ##### | Brain - Cerebellar Hemisphere | 130356125 | 129856125 | 130856125 | No |
| ENSG00000072135.12 | PTPN18 | rs72978686 | 0.000019 | ##### | Brain - Cerebellar Hemisphere | 130356125 | 129856125 | 130856125 | No |
| ENSG00000072135.12 | PTPN18 | rs6712719 | 0.000019 | ##### | Brain - Cerebellar Hemisphere | 130356125 | 129856125 | 130856125 | No |
| ENSG00000072135.12 | PTPN18 | rs72978700 | 0.000019 | ##### | Brain - Cerebellar Hemisphere | 130356125 | 129856125 | 130856125 | No |
| ENSG00000072135.12 | PTPN18 | rs7583274 | 0.000019 | ##### | Brain - Nucleus accumbens (b | 130356125 | 129856125 | 130856125 | Yes |
| ENSG00000072135.12 | PTPN18 | rs57333382 | 0.00002 | ##### | Brain - Cerebellar Hemisphere | 130356125 | 129856125 | 130856125 | Yes |
| ENSG00000072135.12 | PTPN18 | rs6708453 | 0.00002 | ##### | Brain - Cerebellar Hemisphere | 130356125 | 129856125 | 130856125 | Yes |
| ENSG00000072135.12 | PTPN18 | rs6734369 | 0.00002 | ##### | Brain - Nucleus accumbens (b | 130356125 | 129856125 | 130856125 | Yes |
| ENSG00000072135.12 | PTPN18 | rs74354117 | 0.00002 | ##### | Brain - Cerebellum | 130356125 | 129856125 | 130856125 | Yes |
| ENSG00000072135.12 | PTPN18 | rs556674096 | 0.000021 | ##### | Brain - Caudate (basal ganglia | 130356125 | 129856125 | 130856125 | Yes |
| ENSG00000072135.12 | PTPN18 | rs532456259 | 0.000021 | ##### | Brain - Caudate (basal ganglia | 130356125 | 129856125 | 130856125 | Yes |
| ENSG00000072135.12 | PTPN18 | rs1966719 | 0.000021 | ##### | Brain - Caudate (basal ganglia | 130356125 | 129856125 | 130856125 | Yes |
| ENSG00000072135.12 | PTPN18 | rs7569474 | 0.000021 | ##### | Brain - Cerebellum | 130356125 | 129856125 | 130856125 | Yes |
| ENSG00000072135.12 | PTPN18 | rs113164400 | 0.000021 | ##### | Brain - Cerebellum | 130356125 | 129856125 | 130856125 | Yes |
| ENSG00000072135.12 | PTPN18 | rs144975008 | 0.000021 | ##### | Brain - Cerebellum | 130356125 | 129856125 | 130856125 | Yes |
| ENSG00000072135.12 | PTPN18 | rs116752374 | 0.000021 | ##### | Brain - Cerebellum | 130356125 | 129856125 | 130856125 | Yes |
| ENSG00000072135.12 | PTPN18 | rs112791004 | 0.000021 | ##### | Brain - Cerebellum | 130356125 | 129856125 | 130856125 | Yes |
| ENSG00000072135.12 | PTPN18 | rs111894685 | 0.000021 | ##### | Brain - Cerebellum | 130356125 | 129856125 | 130856125 | Yes |
| ENSG00000072135.12 | PTPN18 | rs112416166 | 0.000021 | ##### | Brain - Cerebellum | 130356125 | 129856125 | 130856125 | Yes |
| ENSG00000072135.12 | PTPN18 | rs113266675 | 0.000021 | ##### | Brain - Cerebellum | 130356125 | 129856125 | 130856125 | Yes |
| ENSG00000072135.12 | PTPN18 | rs113077512 | 0.000021 | ##### | Brain - Cerebellum | 130356125 | 129856125 | 130856125 | Yes |
| ENSG00000072135.12 | PTPN18 | rs143960243 | 0.000021 | ##### | Brain - Cerebellum | 130356125 | 129856125 | 130856125 | Yes |
| ENSG00000072135.12 | PTPN18 | rs112176365 | 0.000021 | ##### | Brain - Cerebellum | 130356125 | 129856125 | 130856125 | Yes |
| ENSG00000072135.12 | PTPN18 | rs6725079 | 0.000021 | ##### | Brain - Cerebellum | 130356125 | 129856125 | 130856125 | Yes |
| ENSG00000072135.12 | PTPN18 | rs544737002 | 0.000021 | ##### | Brain - Cerebellum | 130356125 | 129856125 | 130856125 | Yes |
| ENSG00000072135.12 | PTPN18 | rs553350244 | 0.000021 | ##### | Brain - Cerebellum | 130356125 | 129856125 | 130856125 | Yes |
| ENSG00000072135.12 | PTPN18 | rs114319834 | 0.000021 | ##### | Brain - Cerebellum | 130356125 | 129856125 | 130856125 | Yes |
| ENSG00000072135.12 | PTPN18 | rs111238220 | 0.000021 | ##### | Brain - Cerebellum | 130356125 | 129856125 | 130856125 | Yes |

| | | | | | | | | | |
|---|---|---|---|---|---|---|---|---|---|
| ENSG00000072135.12 | PTPN18 | rs79305276 | 0.000021 | ##### | Brain - Cerebellum | 130356125 | 129856125 | 130856125 | Yes |
| ENSG00000072135.12 | PTPN18 | rs13402442 | 0.000022 | ##### | Brain - Cortex | 130356125 | 129856125 | 130856125 | Yes |
| ENSG00000072135.12 | PTPN18 | rs73959898 | 0.000022 | ##### | Brain - Cerebellum | 130356125 | 129856125 | 130856125 | No |
| ENSG00000072135.12 | PTPN18 | rs10206084 | 0.000023 | ##### | Brain - Cortex | 130356125 | 129856125 | 130856125 | Yes |
| ENSG00000072135.12 | PTPN18 | rs112314492 | 0.000024 | ##### | Brain - Cortex | 130356125 | 129856125 | 130856125 | No |
| ENSG00000072135.12 | PTPN18 | rs6431110 | 0.000024 | ##### | Brain - Cortex | 130356125 | 129856125 | 130856125 | No |
| ENSG00000072135.12 | PTPN18 | rs6745431 | 0.000024 | ##### | Brain - Cortex | 130356125 | 129856125 | 130856125 | No |
| ENSG00000072135.12 | PTPN18 | rs6735710 | 0.000024 | ##### | Brain - Cortex | 130356125 | 129856125 | 130856125 | No |
| ENSG00000072135.12 | PTPN18 | rs11891383 | 0.000024 | ##### | Brain - Cortex | 130356125 | 129856125 | 130856125 | No |
| ENSG00000072135.12 | PTPN18 | rs35287933 | 0.000024 | ##### | Brain - Cortex | 130356125 | 129856125 | 130856125 | No |
| ENSG00000072135.12 | PTPN18 | rs79727603 | 0.000024 | ##### | Brain - Cortex | 130356125 | 129856125 | 130856125 | Yes |
| ENSG00000072135.12 | PTPN18 | rs74548388 | 0.000024 | ##### | Brain - Cortex | 130356125 | 129856125 | 130856125 | No |
| ENSG00000072135.12 | PTPN18 | rs200512114 | 0.000025 | ##### | Brain - Cerebellum | 130356125 | 129856125 | 130856125 | Yes |
| ENSG00000072135.12 | PTPN18 | rs10179902 | 0.000027 | ##### | Brain - Nucleus accumbens (b | 130356125 | 129856125 | 130856125 | Yes |
| ENSG00000072135.12 | PTPN18 | rs7583274 | 0.000027 | ##### | Brain - Cerebellar Hemisphere | 130356125 | 129856125 | 130856125 | Yes |
| ENSG00000072135.12 | PTPN18 | rs112400519 | 0.000027 | ##### | Brain - Cerebellum | 130356125 | 129856125 | 130856125 | Yes |
| ENSG00000072135.12 | PTPN18 | rs6743749 | 0.000027 | ##### | Brain - Cerebellar Hemisphere | 130356125 | 129856125 | 130856125 | Yes |
| ENSG00000072135.12 | PTPN18 | rs6729549 | 0.000027 | ##### | Brain - Cerebellar Hemisphere | 130356125 | 129856125 | 130856125 | Yes |
| ENSG00000072135.12 | PTPN18 | rs6729562 | 0.000027 | ##### | Brain - Cerebellar Hemisphere | 130356125 | 129856125 | 130856125 | Yes |
| ENSG00000072135.12 | PTPN18 | rs6750168 | 0.000027 | ##### | Brain - Cerebellar Hemisphere | 130356125 | 129856125 | 130856125 | Yes |
| ENSG00000072135.12 | PTPN18 | rs57742879 | 0.000027 | ##### | Brain - Cerebellar Hemisphere | 130356125 | 129856125 | 130856125 | Yes |
| ENSG00000072135.12 | PTPN18 | rs58192070 | 0.000027 | ##### | Brain - Cerebellar Hemisphere | 130356125 | 129856125 | 130856125 | Yes |
| ENSG00000072135.12 | PTPN18 | rs141558337 | 0.000027 | ##### | Brain - Cerebellar Hemisphere | 130356125 | 129856125 | 130856125 | Yes |
| ENSG00000072135.12 | PTPN18 | rs11885772 | 0.000028 | ##### | Brain - Nucleus accumbens (b | 130356125 | 129856125 | 130856125 | Yes |
| ENSG00000072135.12 | PTPN18 | rs77687592 | 0.000028 | ##### | Brain - Cerebellar Hemisphere | 130356125 | 129856125 | 130856125 | Yes |
| ENSG00000072135.12 | PTPN18 | rs6760375 | 0.00003 | ##### | Brain - Cerebellar Hemisphere | 130356125 | 129856125 | 130856125 | No |
| ENSG00000072135.12 | PTPN18 | rs6431075 | 0.00003 | ##### | Brain - Cerebellar Hemisphere | 130356125 | 129856125 | 130856125 | No |
| ENSG00000072135.12 | PTPN18 | rs73960313 | 0.000031 | ##### | Brain - Cerebellum | 130356125 | 129856125 | 130856125 | Yes |
| ENSG00000072135.12 | PTPN18 | rs144975008 | 0.000031 | ##### | Brain - Cerebellar Hemisphere | 130356125 | 129856125 | 130856125 | Yes |
| ENSG00000072135.12 | PTPN18 | rs116752374 | 0.000031 | ##### | Brain - Cerebellar Hemisphere | 130356125 | 129856125 | 130856125 | Yes |
| ENSG00000072135.12 | PTPN18 | rs112791004 | 0.000031 | ##### | Brain - Cerebellar Hemisphere | 130356125 | 129856125 | 130856125 | Yes |
| ENSG00000072135.12 | PTPN18 | rs111894685 | 0.000031 | ##### | Brain - Cerebellar Hemisphere | 130356125 | 129856125 | 130856125 | Yes |

| | | | | | | | | | |
|---|---|---|---|---|---|---|---|---|---|
| ENSG00000072135.12 | PTPN18 | rs112416166 | 0.000031 | ##### | Brain - Cerebellar Hemisphere | 130356125 | 129856125 | 130856125 | Yes |
| ENSG00000072135.12 | PTPN18 | rs113266675 | 0.000031 | ##### | Brain - Cerebellar Hemisphere | 130356125 | 129856125 | 130856125 | Yes |
| ENSG00000072135.12 | PTPN18 | rs113077512 | 0.000031 | ##### | Brain - Cerebellar Hemisphere | 130356125 | 129856125 | 130856125 | Yes |
| ENSG00000072135.12 | PTPN18 | rs143960243 | 0.000031 | ##### | Brain - Cerebellar Hemisphere | 130356125 | 129856125 | 130856125 | Yes |
| ENSG00000072135.12 | PTPN18 | rs112176365 | 0.000031 | ##### | Brain - Cerebellar Hemisphere | 130356125 | 129856125 | 130856125 | Yes |
| ENSG00000072135.12 | PTPN18 | rs6725079 | 0.000031 | ##### | Brain - Cerebellar Hemisphere | 130356125 | 129856125 | 130856125 | Yes |
| ENSG00000072135.12 | PTPN18 | rs544737002 | 0.000031 | ##### | Brain - Cerebellar Hemisphere | 130356125 | 129856125 | 130856125 | Yes |
| ENSG00000072135.12 | PTPN18 | rs553350244 | 0.000031 | ##### | Brain - Cerebellar Hemisphere | 130356125 | 129856125 | 130856125 | Yes |
| ENSG00000072135.12 | PTPN18 | rs114319834 | 0.000031 | ##### | Brain - Cerebellar Hemisphere | 130356125 | 129856125 | 130856125 | Yes |
| ENSG00000072135.12 | PTPN18 | rs111238220 | 0.000031 | ##### | Brain - Cerebellar Hemisphere | 130356125 | 129856125 | 130856125 | Yes |
| ENSG00000072135.12 | PTPN18 | rs79305276 | 0.000031 | ##### | Brain - Cerebellar Hemisphere | 130356125 | 129856125 | 130856125 | Yes |
| ENSG00000072135.12 | PTPN18 | rs144193979 | 0.000031 | ##### | Brain - Cerebellar Hemisphere | 130356125 | 129856125 | 130856125 | Yes |
| ENSG00000072135.12 | PTPN18 | rs146558998 | 0.000031 | ##### | Brain - Cerebellar Hemisphere | 130356125 | 129856125 | 130856125 | Yes |
| ENSG00000072135.12 | PTPN18 | rs76032333 | 0.000031 | ##### | Brain - Cerebellar Hemisphere | 130356125 | 129856125 | 130856125 | Yes |
| ENSG00000072135.12 | PTPN18 | rs141587479 | 0.000031 | ##### | Brain - Cerebellar Hemisphere | 130356125 | 129856125 | 130856125 | Yes |
| ENSG00000072135.12 | PTPN18 | rs17016591 | 0.000031 | ##### | Brain - Cerebellar Hemisphere | 130356125 | 129856125 | 130856125 | Yes |
| ENSG00000072135.12 | PTPN18 | rs17016603 | 0.000031 | ##### | Brain - Cerebellar Hemisphere | 130356125 | 129856125 | 130856125 | Yes |
| ENSG00000072135.12 | PTPN18 | rs6742510 | 0.000031 | ##### | Brain - Cerebellar Hemisphere | 130356125 | 129856125 | 130856125 | Yes |
| ENSG00000072135.12 | PTPN18 | rs17016615 | 0.000031 | ##### | Brain - Cerebellar Hemisphere | 130356125 | 129856125 | 130856125 | Yes |
| ENSG00000072135.12 | PTPN18 | rs17016616 | 0.000031 | ##### | Brain - Cerebellar Hemisphere | 130356125 | 129856125 | 130856125 | Yes |
| ENSG00000072135.12 | PTPN18 | rs201214742 | 0.000031 | ##### | Brain - Cerebellar Hemisphere | 130356125 | 129856125 | 130856125 | Yes |
| ENSG00000072135.12 | PTPN18 | rs17016621 | 0.000031 | ##### | Brain - Cerebellar Hemisphere | 130356125 | 129856125 | 130856125 | Yes |
| ENSG00000072135.12 | PTPN18 | rs116779623 | 0.000033 | ##### | Brain - Nucleus accumbens (b | 130356125 | 129856125 | 130856125 | Yes |
| ENSG00000072135.12 | PTPN18 | rs530924882 | 0.000033 | ##### | Brain - Frontal Cortex (BA9) | 130356125 | 129856125 | 130856125 | Yes |
| ENSG00000072135.12 | PTPN18 | rs9967795 | 0.000034 | ##### | Brain - Cerebellum | 130356125 | 129856125 | 130856125 | Yes |
| ENSG00000072135.12 | PTPN18 | rs9967796 | 0.000034 | ##### | Brain - Cerebellum | 130356125 | 129856125 | 130856125 | Yes |
| ENSG00000072135.12 | PTPN18 | rs62161824 | 0.000034 | ##### | Brain - Cortex | 130356125 | 129856125 | 130856125 | Yes |
| ENSG00000072135.12 | PTPN18 | rs62161829 | 0.000034 | ##### | Brain - Cortex | 130356125 | 129856125 | 130856125 | Yes |
| ENSG00000072135.12 | PTPN18 | rs765613522 | 0.000034 | ##### | Brain - Cortex | 130356125 | 129856125 | 130856125 | Yes |
| ENSG00000072135.12 | PTPN18 | rs766332156 | 0.000034 | ##### | Brain - Cortex | 130356125 | 129856125 | 130856125 | Yes |
| ENSG00000072135.12 | PTPN18 | rs79015705 | 0.000034 | ##### | Brain - Cortex | 130356125 | 129856125 | 130856125 | Yes |
| ENSG00000072135.12 | PTPN18 | rs76557977 | 0.000034 | ##### | Brain - Cortex | 130356125 | 129856125 | 130856125 | Yes |

| | | | | | | | | | |
|---|---|---|---|---|---|---|---|---|---|
| ENSG00000072135.12 | PTPN18 | rs62161835 | 0.000034 | ##### | Brain - Cortex | 130356125 | 129856125 | 130856125 | Yes |
| ENSG00000072135.12 | PTPN18 | rs62161839 | 0.000034 | ##### | Brain - Cortex | 130356125 | 129856125 | 130856125 | Yes |
| ENSG00000072135.12 | PTPN18 | rs772763769 | 0.000034 | ##### | Brain - Cortex | 130356125 | 129856125 | 130856125 | Yes |
| ENSG00000072135.12 | PTPN18 | rs867828529 | 0.000034 | ##### | Brain - Cortex | 130356125 | 129856125 | 130856125 | Yes |
| ENSG00000072135.12 | PTPN18 | rs73960313 | 0.000035 | ##### | Brain - Cortex | 130356125 | 129856125 | 130856125 | Yes |
| ENSG00000072135.12 | PTPN18 | rs200512114 | 0.000035 | ##### | Brain - Cerebellar Hemisphere | 130356125 | 129856125 | 130856125 | Yes |
| ENSG00000072135.12 | PTPN18 | rs7560297 | 0.000041 | ##### | Brain - Cerebellum | 130356125 | 129856125 | 130856125 | Yes |
| ENSG00000072135.12 | PTPN18 | rs11904313 | 0.000041 | ##### | Brain - Cerebellum | 130356125 | 129856125 | 130856125 | Yes |
| ENSG00000072135.12 | PTPN18 | rs9784097 | 0.000041 | ##### | Brain - Cerebellum | 130356125 | 129856125 | 130856125 | Yes |
| ENSG00000072135.12 | PTPN18 | rs116617341 | 0.000041 | ##### | Brain - Cerebellar Hemisphere | 130356125 | 129856125 | 130856125 | No |
| ENSG00000072135.12 | PTPN18 | rs531901366 | 0.000041 | ##### | Brain - Cortex | 130356125 | 129856125 | 130856125 | Yes |
| ENSG00000072135.12 | PTPN18 | rs11901384 | 0.000043 | ##### | Brain - Caudate (basal ganglia | 130356125 | 129856125 | 130856125 | Yes |
| ENSG00000072135.12 | PTPN18 | rs767609696 | 0.000045 | ##### | Brain - Cortex | 130356125 | 129856125 | 130856125 | Yes |
| ENSG00000072135.12 | PTPN18 | rs62163589 | 0.000045 | ##### | Brain - Cortex | 130356125 | 129856125 | 130856125 | Yes |
| ENSG00000072135.12 | PTPN18 | rs202245476 | 0.000045 | ##### | Brain - Cortex | 130356125 | 129856125 | 130856125 | Yes |
| ENSG00000072135.12 | PTPN18 | rs9677253 | 0.000045 | ##### | Brain - Cortex | 130356125 | 129856125 | 130856125 | Yes |
| ENSG00000072135.12 | PTPN18 | rs17016623 | 0.000046 | ##### | Brain - Cortex | 130356125 | 129856125 | 130856125 | Yes |
| ENSG00000072135.12 | PTPN18 | rs186106956 | 0.000047 | ##### | Brain - Frontal Cortex (BA9) | 130356125 | 129856125 | 130856125 | Yes |
| ENSG00000072135.12 | PTPN18 | rs11904313 | 0.000047 | ##### | Brain - Cerebellar Hemisphere | 130356125 | 129856125 | 130856125 | Yes |
| ENSG00000072135.12 | PTPN18 | rs9784097 | 0.000047 | ##### | Brain - Cerebellar Hemisphere | 130356125 | 129856125 | 130856125 | Yes |
| ENSG00000072135.12 | PTPN18 | rs144193979 | 0.000048 | ##### | Brain - Cortex | 130356125 | 129856125 | 130856125 | Yes |
| ENSG00000072135.12 | PTPN18 | rs146558998 | 0.000048 | ##### | Brain - Cortex | 130356125 | 129856125 | 130856125 | Yes |
| ENSG00000072135.12 | PTPN18 | rs141587479 | 0.000048 | ##### | Brain - Cortex | 130356125 | 129856125 | 130856125 | Yes |
| ENSG00000072135.12 | PTPN18 | rs17016591 | 0.000048 | ##### | Brain - Cortex | 130356125 | 129856125 | 130856125 | Yes |
| ENSG00000072135.12 | PTPN18 | rs17016603 | 0.000048 | ##### | Brain - Cortex | 130356125 | 129856125 | 130856125 | Yes |
| ENSG00000072135.12 | PTPN18 | rs6742510 | 0.000048 | ##### | Brain - Cortex | 130356125 | 129856125 | 130856125 | Yes |
| ENSG00000072135.12 | PTPN18 | rs17016615 | 0.000048 | ##### | Brain - Cortex | 130356125 | 129856125 | 130856125 | Yes |
| ENSG00000072135.12 | PTPN18 | rs17016616 | 0.000048 | ##### | Brain - Cortex | 130356125 | 129856125 | 130856125 | Yes |
| ENSG00000072135.12 | PTPN18 | rs201214742 | 0.000048 | ##### | Brain - Cortex | 130356125 | 129856125 | 130856125 | Yes |
| ENSG00000072135.12 | PTPN18 | rs17016621 | 0.000048 | ##### | Brain - Cortex | 130356125 | 129856125 | 130856125 | Yes |
| ENSG00000072135.12 | PTPN18 | rs186106956 | 0.000049 | ##### | Brain - Cerebellum | 130356125 | 129856125 | 130856125 | Yes |
| ENSG00000072135.12 | PTPN18 | rs75640268 | 0.000049 | ##### | Brain - Cerebellum | 130356125 | 129856125 | 130856125 | Yes |

| | | | | | | | | | |
|---|---|---|---|---|---|---|---|---|---|
| ENSG00000072135.12 | PTPN18 | rs144193979 | 0.00005 | ##### | Brain - Nucleus accumbens (b | 130356125 | 129856125 | 130856125 | Yes |
| ENSG00000072135.12 | PTPN18 | rs146558998 | 0.00005 | ##### | Brain - Nucleus accumbens (b | 130356125 | 129856125 | 130856125 | Yes |
| ENSG00000072135.12 | PTPN18 | rs76032333 | 0.00005 | ##### | Brain - Nucleus accumbens (b | 130356125 | 129856125 | 130856125 | Yes |
| ENSG00000072135.12 | PTPN18 | rs141587479 | 0.00005 | ##### | Brain - Nucleus accumbens (b | 130356125 | 129856125 | 130856125 | Yes |
| ENSG00000072135.12 | PTPN18 | rs17016591 | 0.00005 | ##### | Brain - Nucleus accumbens (b | 130356125 | 129856125 | 130856125 | Yes |
| ENSG00000072135.12 | PTPN18 | rs17016603 | 0.00005 | ##### | Brain - Nucleus accumbens (b | 130356125 | 129856125 | 130856125 | Yes |
| ENSG00000072135.12 | PTPN18 | rs6742510 | 0.00005 | ##### | Brain - Nucleus accumbens (b | 130356125 | 129856125 | 130856125 | Yes |
| ENSG00000072135.12 | PTPN18 | rs17016615 | 0.00005 | ##### | Brain - Nucleus accumbens (b | 130356125 | 129856125 | 130856125 | Yes |
| ENSG00000072135.12 | PTPN18 | rs17016616 | 0.00005 | ##### | Brain - Nucleus accumbens (b | 130356125 | 129856125 | 130856125 | Yes |
| ENSG00000072135.12 | PTPN18 | rs201214742 | 0.00005 | ##### | Brain - Nucleus accumbens (b | 130356125 | 129856125 | 130856125 | Yes |
| ENSG00000072135.12 | PTPN18 | rs17016621 | 0.00005 | ##### | Brain - Nucleus accumbens (b | 130356125 | 129856125 | 130856125 | Yes |
| ENSG00000072135.12 | PTPN18 | rs75815833 | 0.000052 | ##### | Brain - Frontal Cortex (BA9) | 130356125 | 129856125 | 130856125 | Yes |
| ENSG00000072135.12 | PTPN18 | rs28764204 | 0.000053 | ##### | Brain - Cerebellar Hemisphere | 130356125 | 129856125 | 130856125 | Yes |
| ENSG00000072135.12 | PTPN18 | rs72614443 | 0.000055 | ##### | Brain - Cerebellar Hemisphere | 130356125 | 129856125 | 130856125 | Yes |
| ENSG00000072135.12 | PTPN18 | rs3739123 | 0.000056 | ##### | Brain - Cerebellum | 130356125 | 129856125 | 130856125 | Yes |
| ENSG00000072135.12 | PTPN18 | rs3739122 | 0.000056 | ##### | Brain - Cerebellum | 130356125 | 129856125 | 130856125 | Yes |
| ENSG00000072135.12 | PTPN18 | rs72854965 | 0.000056 | ##### | Brain - Cerebellum | 130356125 | 129856125 | 130856125 | Yes |
| ENSG00000072135.12 | PTPN18 | rs10198024 | 0.000056 | ##### | Brain - Nucleus accumbens (b | 130356125 | 129856125 | 130856125 | No |
| ENSG00000072135.12 | PTPN18 | rs6743180 | 0.000056 | ##### | Brain - Nucleus accumbens (b | 130356125 | 129856125 | 130856125 | No |
| ENSG00000072135.12 | PTPN18 | rs10179536 | 0.000056 | ##### | Brain - Nucleus accumbens (b | 130356125 | 129856125 | 130856125 | No |
| ENSG00000072135.12 | PTPN18 | rs77333903 | 0.000056 | ##### | Brain - Nucleus accumbens (b | 130356125 | 129856125 | 130856125 | No |
| ENSG00000072135.12 | PTPN18 | rs13382707 | 0.000056 | ##### | Brain - Nucleus accumbens (b | 130356125 | 129856125 | 130856125 | No |
| ENSG00000072135.12 | PTPN18 | rs73959907 | 0.000056 | ##### | Brain - Nucleus accumbens (b | 130356125 | 129856125 | 130856125 | No |
| ENSG00000072135.12 | PTPN18 | rs75923683 | 0.000056 | ##### | Brain - Nucleus accumbens (b | 130356125 | 129856125 | 130856125 | No |
| ENSG00000072135.12 | PTPN18 | rs10205653 | 0.000056 | ##### | Brain - Nucleus accumbens (b | 130356125 | 129856125 | 130856125 | No |
| ENSG00000072135.12 | PTPN18 | rs10197298 | 0.000056 | ##### | Brain - Nucleus accumbens (b | 130356125 | 129856125 | 130856125 | No |
| ENSG00000072135.12 | PTPN18 | rs13393960 | 0.000058 | ##### | Brain - Cortex | 130356125 | 129856125 | 130856125 | Yes |
| ENSG00000072135.12 | PTPN18 | rs76032333 | 0.000058 | ##### | Brain - Cortex | 130356125 | 129856125 | 130856125 | Yes |
| ENSG00000072135.12 | PTPN18 | rs10198024 | 0.000059 | ##### | Brain - Cortex | 130356125 | 129856125 | 130856125 | No |
| ENSG00000072135.12 | PTPN18 | rs6743180 | 0.000059 | ##### | Brain - Cortex | 130356125 | 129856125 | 130856125 | No |
| ENSG00000072135.12 | PTPN18 | rs10179536 | 0.000059 | ##### | Brain - Cortex | 130356125 | 129856125 | 130856125 | No |
| ENSG00000072135.12 | PTPN18 | rs77333903 | 0.000059 | ##### | Brain - Cortex | 130356125 | 129856125 | 130856125 | No |

| | | | | | | | | | |
|---|---|---|---|---|---|---|---|---|---|
| ENSG00000072135.12 | PTPN18 | rs13382707 | 0.000059 | ##### | Brain - Cortex | 130356125 | 129856125 | 130856125 | No |
| ENSG00000072135.12 | PTPN18 | rs73959907 | 0.000059 | ##### | Brain - Cortex | 130356125 | 129856125 | 130856125 | No |
| ENSG00000072135.12 | PTPN18 | rs75923683 | 0.000059 | ##### | Brain - Cortex | 130356125 | 129856125 | 130856125 | No |
| ENSG00000072135.12 | PTPN18 | rs10205653 | 0.000059 | ##### | Brain - Cortex | 130356125 | 129856125 | 130856125 | No |
| ENSG00000072135.12 | PTPN18 | rs10197298 | 0.000059 | ##### | Brain - Cortex | 130356125 | 129856125 | 130856125 | No |
| ENSG00000072135.12 | PTPN18 | rs79166986 | 0.000059 | ##### | Brain - Cerebellar Hemisphere | 130356125 | 129856125 | 130856125 | No |
| ENSG00000072135.12 | PTPN18 | rs144975008 | 0.000059 | ##### | Brain - Cortex | 130356125 | 129856125 | 130856125 | Yes |
| ENSG00000072135.12 | PTPN18 | rs116752374 | 0.000059 | ##### | Brain - Cortex | 130356125 | 129856125 | 130856125 | Yes |
| ENSG00000072135.12 | PTPN18 | rs112791004 | 0.000059 | ##### | Brain - Cortex | 130356125 | 129856125 | 130856125 | Yes |
| ENSG00000072135.12 | PTPN18 | rs111894685 | 0.000059 | ##### | Brain - Cortex | 130356125 | 129856125 | 130856125 | Yes |
| ENSG00000072135.12 | PTPN18 | rs112416166 | 0.000059 | ##### | Brain - Cortex | 130356125 | 129856125 | 130856125 | Yes |
| ENSG00000072135.12 | PTPN18 | rs113266675 | 0.000059 | ##### | Brain - Cortex | 130356125 | 129856125 | 130856125 | Yes |
| ENSG00000072135.12 | PTPN18 | rs113077512 | 0.000059 | ##### | Brain - Cortex | 130356125 | 129856125 | 130856125 | Yes |
| ENSG00000072135.12 | PTPN18 | rs143960243 | 0.000059 | ##### | Brain - Cortex | 130356125 | 129856125 | 130856125 | Yes |
| ENSG00000072135.12 | PTPN18 | rs112176365 | 0.000059 | ##### | Brain - Cortex | 130356125 | 129856125 | 130856125 | Yes |
| ENSG00000072135.12 | PTPN18 | rs6725079 | 0.000059 | ##### | Brain - Cortex | 130356125 | 129856125 | 130856125 | Yes |
| ENSG00000072135.12 | PTPN18 | rs544737002 | 0.000059 | ##### | Brain - Cortex | 130356125 | 129856125 | 130856125 | Yes |
| ENSG00000072135.12 | PTPN18 | rs553350244 | 0.000059 | ##### | Brain - Cortex | 130356125 | 129856125 | 130856125 | Yes |
| ENSG00000072135.12 | PTPN18 | rs114319834 | 0.000059 | ##### | Brain - Cortex | 130356125 | 129856125 | 130856125 | Yes |
| ENSG00000072135.12 | PTPN18 | rs111238220 | 0.000059 | ##### | Brain - Cortex | 130356125 | 129856125 | 130856125 | Yes |
| ENSG00000072135.12 | PTPN18 | rs79305276 | 0.000059 | ##### | Brain - Cortex | 130356125 | 129856125 | 130856125 | Yes |
| ENSG00000072135.12 | PTPN18 | rs200512114 | 0.000063 | ##### | Brain - Cortex | 130356125 | 129856125 | 130856125 | Yes |
| ENSG00000072135.12 | PTPN18 | rs7603950 | 0.000064 | ##### | Brain - Caudate (basal ganglia | 130356125 | 129856125 | 130856125 | Yes |
| ENSG00000072135.12 | PTPN18 | rs7563097 | 0.000064 | ##### | Brain - Caudate (basal ganglia | 130356125 | 129856125 | 130856125 | Yes |
| ENSG00000072135.12 | PTPN18 | rs11886406 | 0.000065 | ##### | Brain - Caudate (basal ganglia | 130356125 | 129856125 | 130856125 | Yes |
| ENSG00000072135.12 | PTPN18 | rs756687836 | 0.000065 | ##### | Brain - Cortex | 130356125 | 129856125 | 130856125 | Yes |
| ENSG00000072135.12 | PTPN18 | rs146985629 | 0.000066 | ##### | Brain - Cortex | 130356125 | 129856125 | 130856125 | Yes |
| ENSG00000072135.12 | PTPN18 | rs55872863 | 0.000067 | ##### | Brain - Cerebellar Hemisphere | 130356125 | 129856125 | 130856125 | Yes |
| ENSG00000072135.12 | PTPN18 | rs72854907 | 0.000068 | ##### | Brain - Cerebellar Hemisphere | 130356125 | 129856125 | 130856125 | Yes |
| ENSG00000072135.12 | PTPN18 | rs72854909 | 0.000068 | ##### | Brain - Cerebellar Hemisphere | 130356125 | 129856125 | 130856125 | Yes |
| ENSG00000072135.12 | PTPN18 | rs72854961 | 0.000072 | ##### | Brain - Cerebellum | 130356125 | 129856125 | 130856125 | Yes |
| ENSG00000072135.12 | PTPN18 | rs115220415 | 0.000075 | ##### | Brain - Cerebellar Hemisphere | 130356125 | 129856125 | 130856125 | No |

| | | | | | | | | | |
|---|---|---|---|---|---|---|---|---|---|
| ENSG00000072135.12 | PTPN18 | rs141297962 | 0.000078 | ##### | Brain - Cerebellar Hemisphere | 130356125 | 129856125 | 130856125 | Yes |
| ENSG00000072135.12 | PTPN18 | rs13427999 | 0.00008 | ##### | Brain - Cerebellar Hemisphere | 130356125 | 129856125 | 130856125 | Yes |
| ENSG00000072135.12 | PTPN18 | rs4377283 | 0.00008 | ##### | Brain - Cerebellar Hemisphere | 130356125 | 129856125 | 130856125 | Yes |
| ENSG00000072135.12 | PTPN18 | rs72614442 | 0.000082 | ##### | Brain - Cerebellum | 130356125 | 129856125 | 130856125 | Yes |
| ENSG00000072135.12 | PTPN18 | rs74548388 | 0.000082 | ##### | Brain - Caudate (basal ganglia | 130356125 | 129856125 | 130856125 | No |
| ENSG00000072135.12 | PTPN18 | rs72854962 | 0.000085 | ##### | Brain - Cerebellum | 130356125 | 129856125 | 130856125 | Yes |
| ENSG00000072135.12 | PTPN18 | rs150068082 | 0.000085 | ##### | Brain - Cerebellar Hemisphere | 130356125 | 129856125 | 130856125 | Yes |
| ENSG00000072135.12 | PTPN18 | rs116762420 | 0.000085 | ##### | Brain - Cerebellar Hemisphere | 130356125 | 129856125 | 130856125 | Yes |
| ENSG00000072135.12 | PTPN18 | rs75640268 | 0.000085 | ##### | Brain - Cerebellar Hemisphere | 130356125 | 129856125 | 130856125 | Yes |
| ENSG00000072135.12 | PTPN18 | rs114478505 | 0.000085 | ##### | Brain - Cerebellar Hemisphere | 130356125 | 129856125 | 130856125 | Yes |
| ENSG00000072135.12 | PTPN18 | rs7567015 | 0.000085 | ##### | Brain - Cerebellar Hemisphere | 130356125 | 129856125 | 130856125 | Yes |
| ENSG00000072135.12 | PTPN18 | rs76751313 | 0.000085 | ##### | Brain - Cerebellar Hemisphere | 130356125 | 129856125 | 130856125 | Yes |
| ENSG00000072135.12 | PTPN18 | rs72854906 | 0.000093 | ##### | Brain - Cerebellar Hemisphere | 130356125 | 129856125 | 130856125 | Yes |
| ENSG00000072135.12 | PTPN18 | rs6713511 | 0.000099 | ##### | Brain - Cerebellum | 130356125 | 129856125 | 130856125 | Yes |
| ENSG00000072135.12 | PTPN18 | rs4458251 | 0.0001 | ##### | Brain - Cerebellar Hemisphere | 130356125 | 129856125 | 130856125 | Yes |
| ENSG00000072135.12 | PTPN18 | rs7558812 | 0.0001 | ##### | Brain - Cerebellar Hemisphere | 130356125 | 129856125 | 130856125 | Yes |
| ENSG00000072135.12 | PTPN18 | rs12232961 | 0.0001 | ##### | Brain - Cerebellar Hemisphere | 130356125 | 129856125 | 130856125 | Yes |
| ENSG00000113356.11 | POLR3G | rs12374570 | 1.20E-08 | ##### | Brain - Nucleus accumbens (b | 90502412 | 90002412 | 91002412 | Yes |
| ENSG00000113356.11 | POLR3G | rs79216528 | 3.10E-07 | ##### | Brain - Nucleus accumbens (b | 90502412 | 90002412 | 91002412 | Yes |
| ENSG00000113356.11 | POLR3G | rs3763072 | 5.70E-07 | ##### | Brain - Nucleus accumbens (b | 90502412 | 90002412 | 91002412 | Yes |
| ENSG00000113356.11 | POLR3G | rs16868917 | 6.60E-07 | ##### | Brain - Nucleus accumbens (b | 90502412 | 90002412 | 91002412 | Yes |
| ENSG00000113356.11 | POLR3G | rs111608865 | 6.60E-07 | ##### | Brain - Nucleus accumbens (b | 90502412 | 90002412 | 91002412 | Yes |
| ENSG00000113356.11 | POLR3G | rs10041922 | 0.0000012 | ##### | Brain - Nucleus accumbens (b | 90502412 | 90002412 | 91002412 | Yes |
| ENSG00000113356.11 | POLR3G | rs6889253 | 0.0000012 | ##### | Brain - Nucleus accumbens (b | 90502412 | 90002412 | 91002412 | Yes |
| ENSG00000113356.11 | POLR3G | rs6890115 | 0.0000012 | ##### | Brain - Nucleus accumbens (b | 90502412 | 90002412 | 91002412 | Yes |
| ENSG00000113356.11 | POLR3G | rs3763070 | 0.0000014 | ##### | Brain - Nucleus accumbens (b | 90502412 | 90002412 | 91002412 | Yes |
| ENSG00000113356.11 | POLR3G | rs10043702 | 0.0000015 | ##### | Brain - Nucleus accumbens (b | 90502412 | 90002412 | 91002412 | Yes |
| ENSG00000113356.11 | POLR3G | rs6889947 | 0.0000015 | ##### | Brain - Nucleus accumbens (b | 90502412 | 90002412 | 91002412 | Yes |
| ENSG00000113356.11 | POLR3G | rs77269723 | 0.0000019 | ##### | Brain - Nucleus accumbens (b | 90502412 | 90002412 | 91002412 | Yes |
| ENSG00000113356.11 | POLR3G | rs78491050 | 0.0000019 | ##### | Brain - Nucleus accumbens (b | 90502412 | 90002412 | 91002412 | Yes |
| ENSG00000113356.11 | POLR3G | rs11956418 | 0.0000019 | ##### | Brain - Nucleus accumbens (b | 90502412 | 90002412 | 91002412 | Yes |
| ENSG00000113356.11 | POLR3G | rs150119518 | 0.000002 | ##### | Brain - Nucleus accumbens (b | 90502412 | 90002412 | 91002412 | Yes |

| | | | | | | | | | |
|---|---|---|---|---|---|---|---|---|---|
| ENSG00000113356.11 | POLR3G | rs75566605 | 0.000002 | ##### | Brain - Nucleus accumbens (b | 90502412 | 90002412 | 91002412 | Yes |
| ENSG00000113356.11 | POLR3G | rs10942594 | 0.000002 | ##### | Brain - Nucleus accumbens (b | 90502412 | 90002412 | 91002412 | Yes |
| ENSG00000113356.11 | POLR3G | rs11953207 | 0.000002 | ##### | Brain - Nucleus accumbens (b | 90502412 | 90002412 | 91002412 | Yes |
| ENSG00000113356.11 | POLR3G | rs10214167 | 0.000002 | ##### | Brain - Nucleus accumbens (b | 90502412 | 90002412 | 91002412 | Yes |
| ENSG00000113356.11 | POLR3G | rs28728101 | 0.0000025 | ##### | Brain - Nucleus accumbens (b | 90502412 | 90002412 | 91002412 | Yes |
| ENSG00000113356.11 | POLR3G | rs12516555 | 0.0000025 | ##### | Brain - Nucleus accumbens (b | 90502412 | 90002412 | 91002412 | Yes |
| ENSG00000113356.11 | POLR3G | rs10041220 | 0.0000025 | ##### | Brain - Nucleus accumbens (b | 90502412 | 90002412 | 91002412 | Yes |
| ENSG00000113356.11 | POLR3G | rs73772634 | 0.0000025 | ##### | Brain - Hypothalamus | 90502412 | 90002412 | 91002412 | Yes |
| ENSG00000113356.11 | POLR3G | rs6877906 | 0.0000025 | ##### | Brain - Hypothalamus | 90502412 | 90002412 | 91002412 | Yes |
| ENSG00000113356.11 | POLR3G | rs74604828 | 0.0000025 | ##### | Brain - Hypothalamus | 90502412 | 90002412 | 91002412 | Yes |
| ENSG00000113356.11 | POLR3G | rs73772637 | 0.0000025 | ##### | Brain - Hypothalamus | 90502412 | 90002412 | 91002412 | Yes |
| ENSG00000113356.11 | POLR3G | rs7712792 | 0.0000025 | ##### | Brain - Hypothalamus | 90502412 | 90002412 | 91002412 | Yes |
| ENSG00000113356.11 | POLR3G | rs10942593 | 0.0000028 | ##### | Brain - Nucleus accumbens (b | 90502412 | 90002412 | 91002412 | Yes |
| ENSG00000113356.11 | POLR3G | rs74651176 | 0.0000029 | ##### | Brain - Nucleus accumbens (b | 90502412 | 90002412 | 91002412 | Yes |
| ENSG00000113356.11 | POLR3G | rs2049010 | 0.0000035 | ##### | Brain - Nucleus accumbens (b | 90502412 | 90002412 | 91002412 | Yes |
| ENSG00000113356.11 | POLR3G | rs62373985 | 0.0000035 | ##### | Brain - Nucleus accumbens (b | 90502412 | 90002412 | 91002412 | Yes |
| ENSG00000113356.11 | POLR3G | rs879383127 | 0.0000035 | ##### | Brain - Nucleus accumbens (b | 90502412 | 90002412 | 91002412 | Yes |
| ENSG00000113356.11 | POLR3G | rs35475471 | 0.0000038 | ##### | Brain - Cortex | 90502412 | 90002412 | 91002412 | Yes |
| ENSG00000113356.11 | POLR3G | rs62373962 | 0.0000039 | ##### | Brain - Nucleus accumbens (b | 90502412 | 90002412 | 91002412 | Yes |
| ENSG00000113356.11 | POLR3G | rs62373963 | 0.0000039 | ##### | Brain - Nucleus accumbens (b | 90502412 | 90002412 | 91002412 | Yes |
| ENSG00000113356.11 | POLR3G | rs75418010 | 0.0000041 | ##### | Brain - Nucleus accumbens (b | 90502412 | 90002412 | 91002412 | Yes |
| ENSG00000113356.11 | POLR3G | rs57677204 | 0.0000041 | ##### | Brain - Nucleus accumbens (b | 90502412 | 90002412 | 91002412 | Yes |
| ENSG00000113356.11 | POLR3G | rs77198275 | 0.0000045 | ##### | Brain - Nucleus accumbens (b | 90502412 | 90002412 | 91002412 | Yes |
| ENSG00000113356.11 | POLR3G | rs113802395 | 0.0000059 | ##### | Brain - Anterior cingulate cort | 90502412 | 90002412 | 91002412 | Yes |
| ENSG00000113356.11 | POLR3G | rs59742368 | 0.0000078 | ##### | Brain - Nucleus accumbens (b | 90502412 | 90002412 | 91002412 | Yes |
| ENSG00000113356.11 | POLR3G | rs2303801 | 0.0000079 | ##### | Brain - Nucleus accumbens (b | 90502412 | 90002412 | 91002412 | Yes |
| ENSG00000113356.11 | POLR3G | rs3805478 | 0.0000085 | ##### | Brain - Nucleus accumbens (b | 90502412 | 90002412 | 91002412 | Yes |
| ENSG00000113356.11 | POLR3G | rs7729481 | 0.0000086 | ##### | Brain - Nucleus accumbens (b | 90502412 | 90002412 | 91002412 | Yes |
| ENSG00000113356.11 | POLR3G | rs144251015 | 0.0000095 | ##### | Brain - Nucleus accumbens (b | 90502412 | 90002412 | 91002412 | Yes |
| ENSG00000113356.11 | POLR3G | rs147851988 | 0.0000099 | ##### | Brain - Anterior cingulate cort | 90502412 | 90002412 | 91002412 | Yes |
| ENSG00000113356.11 | POLR3G | rs61610742 | 0.000011 | ##### | Brain - Nucleus accumbens (b | 90502412 | 90002412 | 91002412 | Yes |
| ENSG00000113356.11 | POLR3G | rs6864062 | 0.000012 | ##### | Brain - Nucleus accumbens (b | 90502412 | 90002412 | 91002412 | Yes |

| | | | | | | | | | |
|---|---|---|---|---|---|---|---|---|---|
| ENSG00000113356.11 | POLR3G | rs3805478 | 0.000013 | ##### | Brain - Cortex | 90502412 | 90002412 | 91002412 | Yes |
| ENSG00000113356.11 | POLR3G | rs10043702 | 0.000013 | ##### | Brain - Cortex | 90502412 | 90002412 | 91002412 | Yes |
| ENSG00000113356.11 | POLR3G | rs75647854 | 0.000014 | ##### | Brain - Nucleus accumbens (b | 90502412 | 90002412 | 91002412 | Yes |
| ENSG00000113356.11 | POLR3G | rs6872360 | 0.000014 | ##### | Brain - Nucleus accumbens (b | 90502412 | 90002412 | 91002412 | Yes |
| ENSG00000113356.11 | POLR3G | rs13436287 | 0.000014 | ##### | Brain - Nucleus accumbens (b | 90502412 | 90002412 | 91002412 | Yes |
| ENSG00000113356.11 | POLR3G | rs7732363 | 0.000014 | ##### | Brain - Nucleus accumbens (b | 90502412 | 90002412 | 91002412 | Yes |
| ENSG00000113356.11 | POLR3G | rs33981273 | 0.000014 | ##### | Brain - Nucleus accumbens (b | 90502412 | 90002412 | 91002412 | Yes |
| ENSG00000113356.11 | POLR3G | rs6867534 | 0.000015 | ##### | Brain - Nucleus accumbens (b | 90502412 | 90002412 | 91002412 | Yes |
| ENSG00000113356.11 | POLR3G | rs111644494 | 0.000015 | ##### | Brain - Putamen (basal gangli | 90502412 | 90002412 | 91002412 | Yes |
| ENSG00000113356.11 | POLR3G | rs188859916 | 0.000015 | ##### | Brain - Putamen (basal gangli | 90502412 | 90002412 | 91002412 | No |
| ENSG00000113356.11 | POLR3G | rs187270236 | 0.000015 | ##### | Brain - Putamen (basal gangli | 90502412 | 90002412 | 91002412 | No |
| ENSG00000113356.11 | POLR3G | rs188703823 | 0.000015 | ##### | Brain - Putamen (basal gangli | 90502412 | 90002412 | 91002412 | No |
| ENSG00000113356.11 | POLR3G | rs16868917 | 0.000016 | ##### | Brain - Cerebellum | 90502412 | 90002412 | 91002412 | Yes |
| ENSG00000113356.11 | POLR3G | rs77855755 | 0.000016 | ##### | Brain - Nucleus accumbens (b | 90502412 | 90002412 | 91002412 | Yes |
| ENSG00000113356.11 | POLR3G | rs12374570 | 0.000016 | ##### | Brain - Cerebellum | 90502412 | 90002412 | 91002412 | Yes |
| ENSG00000113356.11 | POLR3G | rs7730433 | 0.000016 | ##### | Brain - Nucleus accumbens (b | 90502412 | 90002412 | 91002412 | Yes |
| ENSG00000113356.11 | POLR3G | rs147227857 | 0.000017 | ##### | Brain - Hypothalamus | 90502412 | 90002412 | 91002412 | Yes |
| ENSG00000113356.11 | POLR3G | rs181885168 | 0.000017 | ##### | Brain - Hypothalamus | 90502412 | 90002412 | 91002412 | Yes |
| ENSG00000113356.11 | POLR3G | rs561355470 | 0.000017 | ##### | Brain - Hypothalamus | 90502412 | 90002412 | 91002412 | Yes |
| ENSG00000113356.11 | POLR3G | rs114006677 | 0.000017 | ##### | Brain - Hypothalamus | 90502412 | 90002412 | 91002412 | Yes |
| ENSG00000113356.11 | POLR3G | rs6894373 | 0.000017 | ##### | Brain - Hypothalamus | 90502412 | 90002412 | 91002412 | Yes |
| ENSG00000113356.11 | POLR3G | rs150826528 | 0.000017 | ##### | Brain - Hypothalamus | 90502412 | 90002412 | 91002412 | Yes |
| ENSG00000113356.11 | POLR3G | rs80277535 | 0.000017 | ##### | Brain - Hypothalamus | 90502412 | 90002412 | 91002412 | Yes |
| ENSG00000113356.11 | POLR3G | rs79825272 | 0.000017 | ##### | Brain - Hypothalamus | 90502412 | 90002412 | 91002412 | Yes |
| ENSG00000113356.11 | POLR3G | rs76537827 | 0.000017 | ##### | Brain - Hypothalamus | 90502412 | 90002412 | 91002412 | Yes |
| ENSG00000113356.11 | POLR3G | rs150270571 | 0.000017 | ##### | Brain - Hypothalamus | 90502412 | 90002412 | 91002412 | Yes |
| ENSG00000113356.11 | POLR3G | rs80051012 | 0.000017 | ##### | Brain - Hypothalamus | 90502412 | 90002412 | 91002412 | Yes |
| ENSG00000113356.11 | POLR3G | rs16869094 | 0.000019 | ##### | Brain - Nucleus accumbens (b | 90502412 | 90002412 | 91002412 | Yes |
| ENSG00000113356.11 | POLR3G | rs62373981 | 0.000019 | ##### | Brain - Nucleus accumbens (b | 90502412 | 90002412 | 91002412 | Yes |
| ENSG00000113356.11 | POLR3G | rs10514335 | 0.000019 | ##### | Brain - Nucleus accumbens (b | 90502412 | 90002412 | 91002412 | Yes |
| ENSG00000113356.11 | POLR3G | rs10473934 | 0.000021 | ##### | Brain - Nucleus accumbens (b | 90502412 | 90002412 | 91002412 | Yes |
| ENSG00000113356.11 | POLR3G | rs10041922 | 0.000036 | ##### | Brain - Cortex | 90502412 | 90002412 | 91002412 | Yes |

| | | | | | | | | | |
|---|---|---|---|---|---|---|---|---|---|
| ENSG00000113356.11 | POLR3G | rs6890115 | 0.000036 | ##### | Brain - Cortex | 90502412 | 90002412 | 91002412 | Yes |
| ENSG00000113356.11 | POLR3G | rs28728101 | 0.000036 | ##### | Brain - Cortex | 90502412 | 90002412 | 91002412 | Yes |
| ENSG00000113356.11 | POLR3G | rs12516555 | 0.000036 | ##### | Brain - Cortex | 90502412 | 90002412 | 91002412 | Yes |
| ENSG00000113356.11 | POLR3G | rs10041220 | 0.000036 | ##### | Brain - Cortex | 90502412 | 90002412 | 91002412 | Yes |
| ENSG00000113356.11 | POLR3G | rs10942593 | 0.000036 | ##### | Brain - Cortex | 90502412 | 90002412 | 91002412 | Yes |
| ENSG00000113356.11 | POLR3G | rs10942594 | 0.000036 | ##### | Brain - Cortex | 90502412 | 90002412 | 91002412 | Yes |
| ENSG00000113356.11 | POLR3G | rs11953207 | 0.000036 | ##### | Brain - Cortex | 90502412 | 90002412 | 91002412 | Yes |
| ENSG00000113356.11 | POLR3G | rs10214167 | 0.000036 | ##### | Brain - Cortex | 90502412 | 90002412 | 91002412 | Yes |
| ENSG00000113356.11 | POLR3G | rs77269723 | 0.000036 | ##### | Brain - Cortex | 90502412 | 90002412 | 91002412 | Yes |
| ENSG00000113356.11 | POLR3G | rs78491050 | 0.000036 | ##### | Brain - Cortex | 90502412 | 90002412 | 91002412 | Yes |
| ENSG00000113356.11 | POLR3G | rs11956418 | 0.000036 | ##### | Brain - Cortex | 90502412 | 90002412 | 91002412 | Yes |
| ENSG00000113356.11 | POLR3G | rs3763070 | 0.000036 | ##### | Brain - Cortex | 90502412 | 90002412 | 91002412 | Yes |
| ENSG00000113356.11 | POLR3G | rs6872360 | 0.000037 | ##### | Brain - Cortex | 90502412 | 90002412 | 91002412 | Yes |
| ENSG00000113356.11 | POLR3G | rs7730433 | 0.000037 | ##### | Brain - Cortex | 90502412 | 90002412 | 91002412 | Yes |
| ENSG00000113356.11 | POLR3G | rs13436287 | 0.000037 | ##### | Brain - Cortex | 90502412 | 90002412 | 91002412 | Yes |
| ENSG00000113356.11 | POLR3G | rs7732363 | 0.000037 | ##### | Brain - Cortex | 90502412 | 90002412 | 91002412 | Yes |
| ENSG00000113356.11 | POLR3G | rs33981273 | 0.000037 | ##### | Brain - Cortex | 90502412 | 90002412 | 91002412 | Yes |
| ENSG00000113356.11 | POLR3G | rs3114654 | 0.000037 | ##### | Brain - Cortex | 90502412 | 90002412 | 91002412 | Yes |
| ENSG00000113356.11 | POLR3G | rs3114652 | 0.000037 | ##### | Brain - Cortex | 90502412 | 90002412 | 91002412 | Yes |
| ENSG00000113356.11 | POLR3G | rs79241504 | 0.000037 | ##### | Brain - Cortex | 90502412 | 90002412 | 91002412 | Yes |
| ENSG00000113356.11 | POLR3G | rs28851470 | 0.000037 | ##### | Brain - Cortex | 90502412 | 90002412 | 91002412 | Yes |
| ENSG00000113356.11 | POLR3G | rs6864062 | 0.000037 | ##### | Brain - Cortex | 90502412 | 90002412 | 91002412 | Yes |
| ENSG00000113356.11 | POLR3G | rs59742368 | 0.000037 | ##### | Brain - Cortex | 90502412 | 90002412 | 91002412 | Yes |
| ENSG00000113356.11 | POLR3G | rs7729481 | 0.000037 | ##### | Brain - Cortex | 90502412 | 90002412 | 91002412 | Yes |
| ENSG00000113356.11 | POLR3G | rs3763072 | 0.000039 | ##### | Brain - Cortex | 90502412 | 90002412 | 91002412 | Yes |
| ENSG00000113356.11 | POLR3G | rs13190091 | 0.00004 | ##### | Brain - Cortex | 90502412 | 90002412 | 91002412 | Yes |
| ENSG00000113356.11 | POLR3G | rs137855265 | 0.00004 | ##### | Brain - Hypothalamus | 90502412 | 90002412 | 91002412 | No |
| ENSG00000113356.11 | POLR3G | rs148382118 | 0.00004 | ##### | Brain - Hypothalamus | 90502412 | 90002412 | 91002412 | No |
| ENSG00000113356.11 | POLR3G | rs16869016 | 0.000042 | ##### | Brain - Cerebellum | 90502412 | 90002412 | 91002412 | Yes |
| ENSG00000113356.11 | POLR3G | rs6889947 | 0.000042 | ##### | Brain - Cortex | 90502412 | 90002412 | 91002412 | Yes |
| ENSG00000113356.11 | POLR3G | rs2303801 | 0.000043 | ##### | Brain - Cortex | 90502412 | 90002412 | 91002412 | Yes |
| ENSG00000113356.11 | POLR3G | rs111608865 | 0.000044 | ##### | Brain - Cerebellum | 90502412 | 90002412 | 91002412 | Yes |

| | | | | | | | | | |
|---|---|---|---|---|---|---|---|---|---|
| ENSG00000113356.11 | POLR3G | rs36131676 | 0.000047 | ##### | Brain - Cortex | 90502412 | 90002412 | 91002412 | Yes |
| ENSG00000113356.11 | POLR3G | rs150119518 | 0.000049 | ##### | Brain - Cerebellum | 90502412 | 90002412 | 91002412 | Yes |
| ENSG00000113356.11 | POLR3G | rs75566605 | 0.000049 | ##### | Brain - Cerebellum | 90502412 | 90002412 | 91002412 | Yes |
| ENSG00000113356.11 | POLR3G | rs144251015 | 0.000051 | ##### | Brain - Cortex | 90502412 | 90002412 | 91002412 | Yes |
| ENSG00000113356.11 | POLR3G | rs76119383 | 0.000058 | ##### | Brain - Cerebellum | 90502412 | 90002412 | 91002412 | Yes |
| ENSG00000113356.11 | POLR3G | rs74484473 | 0.000059 | ##### | Brain - Nucleus accumbens (b | 90502412 | 90002412 | 91002412 | Yes |
| ENSG00000113356.11 | POLR3G | rs16869016 | 0.000064 | ##### | Brain - Nucleus accumbens (b | 90502412 | 90002412 | 91002412 | Yes |
| ENSG00000113356.11 | POLR3G | rs76119383 | 0.000064 | ##### | Brain - Nucleus accumbens (b | 90502412 | 90002412 | 91002412 | Yes |
| ENSG00000113356.11 | POLR3G | rs12374570 | 0.000068 | ##### | Brain - Cortex | 90502412 | 90002412 | 91002412 | Yes |
| ENSG00000113356.11 | POLR3G | rs6889253 | 0.000073 | ##### | Brain - Cortex | 90502412 | 90002412 | 91002412 | Yes |
| ENSG00000113356.11 | POLR3G | rs75418010 | 0.000077 | ##### | Brain - Cerebellum | 90502412 | 90002412 | 91002412 | Yes |
| ENSG00000113356.11 | POLR3G | rs57677204 | 0.000077 | ##### | Brain - Cerebellum | 90502412 | 90002412 | 91002412 | Yes |
| ENSG00000113356.11 | POLR3G | rs10473934 | 0.000079 | ##### | Brain - Cortex | 90502412 | 90002412 | 91002412 | Yes |
| ENSG00000132321.16 | IQCA1 | rs10179644 | 5.40E-09 | ##### | Brain - Cortex | 236419198 | 235919198 | 236919198 | Yes |
| ENSG00000132321.16 | IQCA1 | rs73127879 | 5.40E-09 | ##### | Brain - Cortex | 236419198 | 235919198 | 236919198 | Yes |
| ENSG00000132321.16 | IQCA1 | rs73127880 | 5.40E-09 | ##### | Brain - Cortex | 236419198 | 235919198 | 236919198 | Yes |
| ENSG00000132321.16 | IQCA1 | rs1922798 | 5.40E-09 | ##### | Brain - Cortex | 236419198 | 235919198 | 236919198 | Yes |
| ENSG00000132321.16 | IQCA1 | rs73127887 | 5.40E-09 | ##### | Brain - Cortex | 236419198 | 235919198 | 236919198 | Yes |
| ENSG00000132321.16 | IQCA1 | rs80148244 | 5.40E-09 | ##### | Brain - Cortex | 236419198 | 235919198 | 236919198 | Yes |
| ENSG00000132321.16 | IQCA1 | rs73127891 | 5.40E-09 | ##### | Brain - Cortex | 236419198 | 235919198 | 236919198 | Yes |
| ENSG00000132321.16 | IQCA1 | rs76804064 | 5.40E-09 | ##### | Brain - Cortex | 236419198 | 235919198 | 236919198 | Yes |
| ENSG00000132321.16 | IQCA1 | rs78752798 | 5.40E-09 | ##### | Brain - Cortex | 236419198 | 235919198 | 236919198 | Yes |
| ENSG00000132321.16 | IQCA1 | rs10193278 | 5.40E-09 | ##### | Brain - Cortex | 236419198 | 235919198 | 236919198 | Yes |
| ENSG00000132321.16 | IQCA1 | rs79733838 | 5.40E-09 | ##### | Brain - Cortex | 236419198 | 235919198 | 236919198 | Yes |
| ENSG00000132321.16 | IQCA1 | rs79357137 | 5.40E-09 | ##### | Brain - Cortex | 236419198 | 235919198 | 236919198 | Yes |
| ENSG00000132321.16 | IQCA1 | rs73127884 | 5.40E-08 | ##### | Brain - Nucleus accumbens (b | 236419198 | 235919198 | 236919198 | Yes |
| ENSG00000132321.16 | IQCA1 | rs6431459 | 7.70E-08 | ##### | Brain - Nucleus accumbens (b | 236419198 | 235919198 | 236919198 | Yes |
| ENSG00000132321.16 | IQCA1 | rs6745302 | 8.90E-08 | ##### | Brain - Cortex | 236419198 | 235919198 | 236919198 | Yes |
| ENSG00000132321.16 | IQCA1 | rs10179644 | 1.00E-07 | ##### | Brain - Nucleus accumbens (b | 236419198 | 235919198 | 236919198 | Yes |
| ENSG00000132321.16 | IQCA1 | rs73127879 | 1.00E-07 | ##### | Brain - Nucleus accumbens (b | 236419198 | 235919198 | 236919198 | Yes |
| ENSG00000132321.16 | IQCA1 | rs73127880 | 1.00E-07 | ##### | Brain - Nucleus accumbens (b | 236419198 | 235919198 | 236919198 | Yes |
| ENSG00000132321.16 | IQCA1 | rs1922798 | 1.00E-07 | ##### | Brain - Nucleus accumbens (b | 236419198 | 235919198 | 236919198 | Yes |

| | | | | | | | | | |
|---|---|---|---|---|---|---|---|---|---|
| ENSG00000132321.16 | IQCA1 | rs73127887 | 1.00E-07 | ##### | Brain - Nucleus accumbens (b | 236419198 | 235919198 | 236919198 | Yes |
| ENSG00000132321.16 | IQCA1 | rs80148244 | 1.00E-07 | ##### | Brain - Nucleus accumbens (b | 236419198 | 235919198 | 236919198 | Yes |
| ENSG00000132321.16 | IQCA1 | rs73127891 | 1.00E-07 | ##### | Brain - Nucleus accumbens (b | 236419198 | 235919198 | 236919198 | Yes |
| ENSG00000132321.16 | IQCA1 | rs76804064 | 1.00E-07 | ##### | Brain - Nucleus accumbens (b | 236419198 | 235919198 | 236919198 | Yes |
| ENSG00000132321.16 | IQCA1 | rs78752798 | 1.00E-07 | ##### | Brain - Nucleus accumbens (b | 236419198 | 235919198 | 236919198 | Yes |
| ENSG00000132321.16 | IQCA1 | rs10193278 | 1.00E-07 | ##### | Brain - Nucleus accumbens (b | 236419198 | 235919198 | 236919198 | Yes |
| ENSG00000132321.16 | IQCA1 | rs79733838 | 1.00E-07 | ##### | Brain - Nucleus accumbens (b | 236419198 | 235919198 | 236919198 | Yes |
| ENSG00000132321.16 | IQCA1 | rs79357137 | 1.00E-07 | ##### | Brain - Nucleus accumbens (b | 236419198 | 235919198 | 236919198 | Yes |
| ENSG00000132321.16 | IQCA1 | rs13428016 | 1.70E-07 | ##### | Brain - Nucleus accumbens (b | 236419198 | 235919198 | 236919198 | Yes |
| ENSG00000132321.16 | IQCA1 | rs10717671 | 1.70E-07 | ##### | Brain - Nucleus accumbens (b | 236419198 | 235919198 | 236919198 | Yes |
| ENSG00000132321.16 | IQCA1 | rs149347254 | 1.90E-07 | ##### | Brain - Nucleus accumbens (b | 236419198 | 235919198 | 236919198 | Yes |
| ENSG00000132321.16 | IQCA1 | rs13432714 | 2.00E-07 | ##### | Brain - Cortex | 236419198 | 235919198 | 236919198 | Yes |
| ENSG00000132321.16 | IQCA1 | rs10167317 | 2.00E-07 | ##### | Brain - Cortex | 236419198 | 235919198 | 236919198 | Yes |
| ENSG00000132321.16 | IQCA1 | rs73127878 | 2.00E-07 | ##### | Brain - Cortex | 236419198 | 235919198 | 236919198 | Yes |
| ENSG00000132321.16 | IQCA1 | rs73127881 | 2.00E-07 | ##### | Brain - Cortex | 236419198 | 235919198 | 236919198 | Yes |
| ENSG00000132321.16 | IQCA1 | rs10203752 | 2.00E-07 | ##### | Brain - Cortex | 236419198 | 235919198 | 236919198 | Yes |
| ENSG00000132321.16 | IQCA1 | rs7590455 | 2.00E-07 | ##### | Brain - Cortex | 236419198 | 235919198 | 236919198 | Yes |
| ENSG00000132321.16 | IQCA1 | rs7578464 | 2.00E-07 | ##### | Brain - Cortex | 236419198 | 235919198 | 236919198 | Yes |
| ENSG00000132321.16 | IQCA1 | rs7564232 | 2.00E-07 | ##### | Brain - Cortex | 236419198 | 235919198 | 236919198 | Yes |
| ENSG00000132321.16 | IQCA1 | rs13388314 | 2.00E-07 | ##### | Brain - Cortex | 236419198 | 235919198 | 236919198 | Yes |
| ENSG00000132321.16 | IQCA1 | rs7561761 | 2.00E-07 | ##### | Brain - Cortex | 236419198 | 235919198 | 236919198 | Yes |
| ENSG00000132321.16 | IQCA1 | rs13388636 | 2.00E-07 | ##### | Brain - Cortex | 236419198 | 235919198 | 236919198 | Yes |
| ENSG00000132321.16 | IQCA1 | rs13388752 | 2.00E-07 | ##### | Brain - Cortex | 236419198 | 235919198 | 236919198 | Yes |
| ENSG00000132321.16 | IQCA1 | rs10196356 | 2.00E-07 | ##### | Brain - Cortex | 236419198 | 235919198 | 236919198 | Yes |
| ENSG00000132321.16 | IQCA1 | rs10173626 | 2.00E-07 | ##### | Brain - Cortex | 236419198 | 235919198 | 236919198 | Yes |
| ENSG00000132321.16 | IQCA1 | rs10196774 | 2.00E-07 | ##### | Brain - Cortex | 236419198 | 235919198 | 236919198 | Yes |
| ENSG00000132321.16 | IQCA1 | rs10173842 | 2.00E-07 | ##### | Brain - Cortex | 236419198 | 235919198 | 236919198 | Yes |
| ENSG00000132321.16 | IQCA1 | rs116063576 | 2.00E-07 | ##### | Brain - Cortex | 236419198 | 235919198 | 236919198 | Yes |
| ENSG00000132321.16 | IQCA1 | rs73129814 | 2.00E-07 | ##### | Brain - Cortex | 236419198 | 235919198 | 236919198 | Yes |
| ENSG00000132321.16 | IQCA1 | rs73129815 | 2.00E-07 | ##### | Brain - Cortex | 236419198 | 235919198 | 236919198 | Yes |
| ENSG00000132321.16 | IQCA1 | rs73129817 | 2.00E-07 | ##### | Brain - Cortex | 236419198 | 235919198 | 236919198 | Yes |
| ENSG00000132321.16 | IQCA1 | rs10192294 | 2.00E-07 | ##### | Brain - Cortex | 236419198 | 235919198 | 236919198 | Yes |

| | | | | | | | | | |
|---|---|---|---|---|---|---|---|---|---|
| ENSG00000132321.16 | IQCA1 | rs10180564 | 2.00E-07 | ##### | Brain - Cortex | 236419198 | 235919198 | 236919198 | Yes |
| ENSG00000132321.16 | IQCA1 | rs10180565 | 2.00E-07 | ##### | Brain - Cortex | 236419198 | 235919198 | 236919198 | Yes |
| ENSG00000132321.16 | IQCA1 | rs7355537 | 2.00E-07 | ##### | Brain - Cortex | 236419198 | 235919198 | 236919198 | Yes |
| ENSG00000132321.16 | IQCA1 | rs10203594 | 2.00E-07 | ##### | Brain - Cortex | 236419198 | 235919198 | 236919198 | Yes |
| ENSG00000132321.16 | IQCA1 | rs10183186 | 2.00E-07 | ##### | Brain - Cortex | 236419198 | 235919198 | 236919198 | Yes |
| ENSG00000132321.16 | IQCA1 | rs113822213 | 2.00E-07 | ##### | Brain - Cortex | 236419198 | 235919198 | 236919198 | Yes |
| ENSG00000132321.16 | IQCA1 | rs73129829 | 2.00E-07 | ##### | Brain - Cortex | 236419198 | 235919198 | 236919198 | Yes |
| ENSG00000132321.16 | IQCA1 | rs57240758 | 2.00E-07 | ##### | Brain - Cortex | 236419198 | 235919198 | 236919198 | Yes |
| ENSG00000132321.16 | IQCA1 | rs6724406 | 2.00E-07 | ##### | Brain - Cortex | 236419198 | 235919198 | 236919198 | Yes |
| ENSG00000132321.16 | IQCA1 | rs6733348 | 2.30E-07 | ##### | Brain - Nucleus accumbens (b | 236419198 | 235919198 | 236919198 | Yes |
| ENSG00000132321.16 | IQCA1 | rs73130904 | 2.50E-07 | ##### | Brain - Spinal cord (cervical c- | 236419198 | 235919198 | 236919198 | Yes |
| ENSG00000132321.16 | IQCA1 | rs76557076 | 2.50E-07 | ##### | Brain - Cortex | 236419198 | 235919198 | 236919198 | Yes |
| ENSG00000132321.16 | IQCA1 | rs11893167 | 2.50E-07 | ##### | Brain - Cortex | 236419198 | 235919198 | 236919198 | Yes |
| ENSG00000132321.16 | IQCA1 | rs2077827 | 2.50E-07 | ##### | Brain - Cortex | 236419198 | 235919198 | 236919198 | Yes |
| ENSG00000132321.16 | IQCA1 | rs2077828 | 2.50E-07 | ##### | Brain - Cortex | 236419198 | 235919198 | 236919198 | Yes |
| ENSG00000132321.16 | IQCA1 | rs73129834 | 2.50E-07 | ##### | Brain - Cortex | 236419198 | 235919198 | 236919198 | Yes |
| ENSG00000132321.16 | IQCA1 | rs1108890 | 2.70E-07 | ##### | Brain - Nucleus accumbens (b | 236419198 | 235919198 | 236919198 | Yes |
| ENSG00000132321.16 | IQCA1 | rs6728422 | 2.90E-07 | ##### | Brain - Nucleus accumbens (b | 236419198 | 235919198 | 236919198 | Yes |
| ENSG00000132321.16 | IQCA1 | rs6742962 | 2.90E-07 | ##### | Brain - Nucleus accumbens (b | 236419198 | 235919198 | 236919198 | Yes |
| ENSG00000132321.16 | IQCA1 | rs6431460 | 3.10E-07 | ##### | Brain - Nucleus accumbens (b | 236419198 | 235919198 | 236919198 | Yes |
| ENSG00000132321.16 | IQCA1 | rs1482960 | 3.20E-07 | ##### | Brain - Nucleus accumbens (b | 236419198 | 235919198 | 236919198 | Yes |
| ENSG00000132321.16 | IQCA1 | rs7597596 | 3.90E-07 | ##### | Brain - Spinal cord (cervical c- | 236419198 | 235919198 | 236919198 | Yes |
| ENSG00000132321.16 | IQCA1 | rs7556918 | 3.90E-07 | ##### | Brain - Spinal cord (cervical c- | 236419198 | 235919198 | 236919198 | Yes |
| ENSG00000132321.16 | IQCA1 | rs6431461 | 4.90E-07 | ##### | Brain - Nucleus accumbens (b | 236419198 | 235919198 | 236919198 | Yes |
| ENSG00000132321.16 | IQCA1 | rs10202768 | 5.60E-07 | ##### | Brain - Spinal cord (cervical c- | 236419198 | 235919198 | 236919198 | Yes |
| ENSG00000132321.16 | IQCA1 | rs13392259 | 5.60E-07 | ##### | Brain - Spinal cord (cervical c- | 236419198 | 235919198 | 236919198 | Yes |
| ENSG00000132321.16 | IQCA1 | rs140081308 | 5.60E-07 | ##### | Brain - Spinal cord (cervical c- | 236419198 | 235919198 | 236919198 | Yes |
| ENSG00000132321.16 | IQCA1 | rs10206139 | 5.60E-07 | ##### | Brain - Spinal cord (cervical c- | 236419198 | 235919198 | 236919198 | Yes |
| ENSG00000132321.16 | IQCA1 | rs13426846 | 5.60E-07 | ##### | Brain - Spinal cord (cervical c- | 236419198 | 235919198 | 236919198 | Yes |
| ENSG00000132321.16 | IQCA1 | rs60242756 | 5.60E-07 | ##### | Brain - Spinal cord (cervical c- | 236419198 | 235919198 | 236919198 | Yes |
| ENSG00000132321.16 | IQCA1 | rs59625470 | 5.60E-07 | ##### | Brain - Spinal cord (cervical c- | 236419198 | 235919198 | 236919198 | Yes |
| ENSG00000132321.16 | IQCA1 | rs77704052 | 5.60E-07 | ##### | Brain - Spinal cord (cervical c- | 236419198 | 235919198 | 236919198 | Yes |

| | | | | | | | | | |
|---|---|---|---|---|---|---|---|---|---|
| ENSG00000132321.16 | IQCA1 | rs79825641 | 5.60E-07 | ##### | Brain - Spinal cord (cervical c- | 236419198 | 235919198 | 236919198 | Yes |
| ENSG00000132321.16 | IQCA1 | rs74941416 | 5.60E-07 | ##### | Brain - Spinal cord (cervical c- | 236419198 | 235919198 | 236919198 | Yes |
| ENSG00000132321.16 | IQCA1 | rs73127885 | 5.80E-07 | ##### | Brain - Cortex | 236419198 | 235919198 | 236919198 | Yes |
| ENSG00000132321.16 | IQCA1 | rs62189777 | 5.80E-07 | ##### | Brain - Nucleus accumbens (b | 236419198 | 235919198 | 236919198 | Yes |
| ENSG00000132321.16 | IQCA1 | rs60549967 | 7.70E-07 | ##### | Brain - Nucleus accumbens (b | 236419198 | 235919198 | 236919198 | Yes |
| ENSG00000132321.16 | IQCA1 | rs2317422 | 8.90E-07 | ##### | Brain - Nucleus accumbens (b | 236419198 | 235919198 | 236919198 | Yes |
| ENSG00000132321.16 | IQCA1 | rs2317638 | 9.10E-07 | ##### | Brain - Nucleus accumbens (b | 236419198 | 235919198 | 236919198 | Yes |
| ENSG00000132321.16 | IQCA1 | rs6431456 | 0.000001 | ##### | Brain - Nucleus accumbens (b | 236419198 | 235919198 | 236919198 | Yes |
| ENSG00000132321.16 | IQCA1 | rs7557194 | 0.000001 | ##### | Brain - Nucleus accumbens (b | 236419198 | 235919198 | 236919198 | Yes |
| ENSG00000132321.16 | IQCA1 | rs55656551 | 0.000001 | ##### | Brain - Nucleus accumbens (b | 236419198 | 235919198 | 236919198 | Yes |
| ENSG00000132321.16 | IQCA1 | rs4060823 | 0.000001 | ##### | Brain - Nucleus accumbens (b | 236419198 | 235919198 | 236919198 | Yes |
| ENSG00000132321.16 | IQCA1 | rs76317401 | 0.0000013 | ##### | Brain - Spinal cord (cervical c- | 236419198 | 235919198 | 236919198 | Yes |
| ENSG00000132321.16 | IQCA1 | rs58605253 | 0.0000013 | ##### | Brain - Spinal cord (cervical c- | 236419198 | 235919198 | 236919198 | Yes |
| ENSG00000132321.16 | IQCA1 | rs115983368 | 0.0000013 | ##### | Brain - Spinal cord (cervical c- | 236419198 | 235919198 | 236919198 | Yes |
| ENSG00000132321.16 | IQCA1 | rs116063415 | 0.0000013 | ##### | Brain - Spinal cord (cervical c- | 236419198 | 235919198 | 236919198 | Yes |
| ENSG00000132321.16 | IQCA1 | rs80263382 | 0.0000013 | ##### | Brain - Spinal cord (cervical c- | 236419198 | 235919198 | 236919198 | Yes |
| ENSG00000132321.16 | IQCA1 | rs143794909 | 0.0000013 | ##### | Brain - Spinal cord (cervical c- | 236419198 | 235919198 | 236919198 | Yes |
| ENSG00000132321.16 | IQCA1 | rs76818679 | 0.0000013 | ##### | Brain - Spinal cord (cervical c- | 236419198 | 235919198 | 236919198 | Yes |
| ENSG00000132321.16 | IQCA1 | rs77528566 | 0.0000013 | ##### | Brain - Spinal cord (cervical c- | 236419198 | 235919198 | 236919198 | Yes |
| ENSG00000132321.16 | IQCA1 | rs537121139 | 0.0000013 | ##### | Brain - Spinal cord (cervical c- | 236419198 | 235919198 | 236919198 | Yes |
| ENSG00000132321.16 | IQCA1 | rs73998612 | 0.0000013 | ##### | Brain - Spinal cord (cervical c- | 236419198 | 235919198 | 236919198 | Yes |
| ENSG00000132321.16 | IQCA1 | rs144917370 | 0.0000013 | ##### | Brain - Spinal cord (cervical c- | 236419198 | 235919198 | 236919198 | Yes |
| ENSG00000132321.16 | IQCA1 | rs76020172 | 0.0000013 | ##### | Brain - Spinal cord (cervical c- | 236419198 | 235919198 | 236919198 | Yes |
| ENSG00000132321.16 | IQCA1 | rs114487697 | 0.0000013 | ##### | Brain - Spinal cord (cervical c- | 236419198 | 235919198 | 236919198 | No |
| ENSG00000132321.16 | IQCA1 | rs6738413 | 0.0000013 | ##### | Brain - Nucleus accumbens (b | 236419198 | 235919198 | 236919198 | Yes |
| ENSG00000132321.16 | IQCA1 | rs6750125 | 0.0000013 | ##### | Brain - Nucleus accumbens (b | 236419198 | 235919198 | 236919198 | Yes |
| ENSG00000132321.16 | IQCA1 | rs11896685 | 0.0000013 | ##### | Brain - Nucleus accumbens (b | 236419198 | 235919198 | 236919198 | Yes |
| ENSG00000132321.16 | IQCA1 | rs112237027 | 0.0000013 | ##### | Brain - Nucleus accumbens (b | 236419198 | 235919198 | 236919198 | Yes |
| ENSG00000132321.16 | IQCA1 | rs141774975 | 0.0000014 | ##### | Brain - Cortex | 236419198 | 235919198 | 236919198 | Yes |
| ENSG00000132321.16 | IQCA1 | rs77047395 | 0.0000014 | ##### | Brain - Cortex | 236419198 | 235919198 | 236919198 | Yes |
| ENSG00000132321.16 | IQCA1 | rs77843043 | 0.0000014 | ##### | Brain - Cortex | 236419198 | 235919198 | 236919198 | Yes |
| ENSG00000132321.16 | IQCA1 | rs6742302 | 0.0000014 | ##### | Brain - Nucleus accumbens (b | 236419198 | 235919198 | 236919198 | Yes |

| | | | | | | | | | |
|---|---|---|---|---|---|---|---|---|---|
| ENSG00000132321.16 | IQCA1 | rs73130918 | 0.0000014 | ##### | Brain - Spinal cord (cervical c- | 236419198 | 235919198 | 236919198 | Yes |
| ENSG00000132321.16 | IQCA1 | rs62192361 | 0.0000014 | ##### | Brain - Nucleus accumbens (b | 236419198 | 235919198 | 236919198 | Yes |
| ENSG00000132321.16 | IQCA1 | rs2046282 | 0.0000015 | ##### | Brain - Nucleus accumbens (b | 236419198 | 235919198 | 236919198 | Yes |
| ENSG00000132321.16 | IQCA1 | rs2046283 | 0.0000015 | ##### | Brain - Nucleus accumbens (b | 236419198 | 235919198 | 236919198 | Yes |
| ENSG00000132321.16 | IQCA1 | rs57364322 | 0.0000018 | ##### | Brain - Anterior cingulate cort | 236419198 | 235919198 | 236919198 | Yes |
| ENSG00000132321.16 | IQCA1 | rs2046280 | 0.0000019 | ##### | Brain - Nucleus accumbens (b | 236419198 | 235919198 | 236919198 | Yes |
| ENSG00000132321.16 | IQCA1 | rs6746336 | 0.000002 | ##### | Brain - Nucleus accumbens (b | 236419198 | 235919198 | 236919198 | Yes |
| ENSG00000132321.16 | IQCA1 | rs11892411 | 0.0000022 | ##### | Brain - Nucleus accumbens (b | 236419198 | 235919198 | 236919198 | Yes |
| ENSG00000132321.16 | IQCA1 | rs6431457 | 0.0000023 | ##### | Brain - Nucleus accumbens (b | 236419198 | 235919198 | 236919198 | Yes |
| ENSG00000132321.16 | IQCA1 | rs10193029 | 0.0000025 | ##### | Brain - Nucleus accumbens (b | 236419198 | 235919198 | 236919198 | Yes |
| ENSG00000132321.16 | IQCA1 | rs13416077 | 0.0000025 | ##### | Brain - Nucleus accumbens (b | 236419198 | 235919198 | 236919198 | Yes |
| ENSG00000132321.16 | IQCA1 | rs7600965 | 0.0000025 | ##### | Brain - Nucleus accumbens (b | 236419198 | 235919198 | 236919198 | Yes |
| ENSG00000132321.16 | IQCA1 | rs59403004 | 0.0000026 | ##### | Brain - Anterior cingulate cort | 236419198 | 235919198 | 236919198 | Yes |
| ENSG00000132321.16 | IQCA1 | rs78453407 | 0.0000031 | ##### | Brain - Cortex | 236419198 | 235919198 | 236919198 | Yes |
| ENSG00000132321.16 | IQCA1 | rs76745547 | 0.0000031 | ##### | Brain - Cortex | 236419198 | 235919198 | 236919198 | Yes |
| ENSG00000132321.16 | IQCA1 | rs77999359 | 0.0000031 | ##### | Brain - Cortex | 236419198 | 235919198 | 236919198 | Yes |
| ENSG00000132321.16 | IQCA1 | rs78988858 | 0.0000031 | ##### | Brain - Cortex | 236419198 | 235919198 | 236919198 | Yes |
| ENSG00000132321.16 | IQCA1 | rs150691798 | 0.0000031 | ##### | Brain - Cortex | 236419198 | 235919198 | 236919198 | Yes |
| ENSG00000132321.16 | IQCA1 | rs76481577 | 0.0000031 | ##### | Brain - Cortex | 236419198 | 235919198 | 236919198 | Yes |
| ENSG00000132321.16 | IQCA1 | rs77858166 | 0.0000031 | ##### | Brain - Cortex | 236419198 | 235919198 | 236919198 | Yes |
| ENSG00000132321.16 | IQCA1 | rs6758710 | 0.0000032 | ##### | Brain - Nucleus accumbens (b | 236419198 | 235919198 | 236919198 | Yes |
| ENSG00000132321.16 | IQCA1 | rs62192376 | 0.0000032 | ##### | Brain - Nucleus accumbens (b | 236419198 | 235919198 | 236919198 | Yes |
| ENSG00000132321.16 | IQCA1 | rs11683125 | 0.0000036 | ##### | Brain - Spinal cord (cervical c- | 236419198 | 235919198 | 236919198 | Yes |
| ENSG00000132321.16 | IQCA1 | rs13407283 | 0.0000036 | ##### | Brain - Spinal cord (cervical c- | 236419198 | 235919198 | 236919198 | Yes |
| ENSG00000132321.16 | IQCA1 | rs13407373 | 0.0000036 | ##### | Brain - Spinal cord (cervical c- | 236419198 | 235919198 | 236919198 | Yes |
| ENSG00000132321.16 | IQCA1 | rs28373664 | 0.0000036 | ##### | Brain - Spinal cord (cervical c- | 236419198 | 235919198 | 236919198 | Yes |
| ENSG00000132321.16 | IQCA1 | rs6759530 | 0.000004 | ##### | Brain - Nucleus accumbens (b | 236419198 | 235919198 | 236919198 | Yes |
| ENSG00000132321.16 | IQCA1 | rs6745302 | 0.0000042 | ##### | Brain - Nucleus accumbens (b | 236419198 | 235919198 | 236919198 | Yes |
| ENSG00000132321.16 | IQCA1 | rs13432714 | 0.0000044 | ##### | Brain - Nucleus accumbens (b | 236419198 | 235919198 | 236919198 | Yes |
| ENSG00000132321.16 | IQCA1 | rs10167317 | 0.0000044 | ##### | Brain - Nucleus accumbens (b | 236419198 | 235919198 | 236919198 | Yes |
| ENSG00000132321.16 | IQCA1 | rs73127878 | 0.0000044 | ##### | Brain - Nucleus accumbens (b | 236419198 | 235919198 | 236919198 | Yes |
| ENSG00000132321.16 | IQCA1 | rs73127881 | 0.0000044 | ##### | Brain - Nucleus accumbens (b | 236419198 | 235919198 | 236919198 | Yes |

| | | | | | | | | | |
|---|---|---|---|---|---|---|---|---|---|
| ENSG00000132321.16 | IQCA1 | rs10203752 | 0.0000044 | ##### | Brain - Nucleus accumbens (b | 236419198 | 235919198 | 236919198 | Yes |
| ENSG00000132321.16 | IQCA1 | rs7590455 | 0.0000044 | ##### | Brain - Nucleus accumbens (b | 236419198 | 235919198 | 236919198 | Yes |
| ENSG00000132321.16 | IQCA1 | rs7578464 | 0.0000044 | ##### | Brain - Nucleus accumbens (b | 236419198 | 235919198 | 236919198 | Yes |
| ENSG00000132321.16 | IQCA1 | rs7564232 | 0.0000044 | ##### | Brain - Nucleus accumbens (b | 236419198 | 235919198 | 236919198 | Yes |
| ENSG00000132321.16 | IQCA1 | rs13388314 | 0.0000044 | ##### | Brain - Nucleus accumbens (b | 236419198 | 235919198 | 236919198 | Yes |
| ENSG00000132321.16 | IQCA1 | rs7561761 | 0.0000044 | ##### | Brain - Nucleus accumbens (b | 236419198 | 235919198 | 236919198 | Yes |
| ENSG00000132321.16 | IQCA1 | rs13388636 | 0.0000044 | ##### | Brain - Nucleus accumbens (b | 236419198 | 235919198 | 236919198 | Yes |
| ENSG00000132321.16 | IQCA1 | rs13388752 | 0.0000044 | ##### | Brain - Nucleus accumbens (b | 236419198 | 235919198 | 236919198 | Yes |
| ENSG00000132321.16 | IQCA1 | rs10196356 | 0.0000044 | ##### | Brain - Nucleus accumbens (b | 236419198 | 235919198 | 236919198 | Yes |
| ENSG00000132321.16 | IQCA1 | rs10173626 | 0.0000044 | ##### | Brain - Nucleus accumbens (b | 236419198 | 235919198 | 236919198 | Yes |
| ENSG00000132321.16 | IQCA1 | rs10196774 | 0.0000044 | ##### | Brain - Nucleus accumbens (b | 236419198 | 235919198 | 236919198 | Yes |
| ENSG00000132321.16 | IQCA1 | rs10173842 | 0.0000044 | ##### | Brain - Nucleus accumbens (b | 236419198 | 235919198 | 236919198 | Yes |
| ENSG00000132321.16 | IQCA1 | rs116063576 | 0.0000044 | ##### | Brain - Nucleus accumbens (b | 236419198 | 235919198 | 236919198 | Yes |
| ENSG00000132321.16 | IQCA1 | rs73129814 | 0.0000044 | ##### | Brain - Nucleus accumbens (b | 236419198 | 235919198 | 236919198 | Yes |
| ENSG00000132321.16 | IQCA1 | rs73129815 | 0.0000044 | ##### | Brain - Nucleus accumbens (b | 236419198 | 235919198 | 236919198 | Yes |
| ENSG00000132321.16 | IQCA1 | rs73129817 | 0.0000044 | ##### | Brain - Nucleus accumbens (b | 236419198 | 235919198 | 236919198 | Yes |
| ENSG00000132321.16 | IQCA1 | rs10192294 | 0.0000044 | ##### | Brain - Nucleus accumbens (b | 236419198 | 235919198 | 236919198 | Yes |
| ENSG00000132321.16 | IQCA1 | rs10180564 | 0.0000044 | ##### | Brain - Nucleus accumbens (b | 236419198 | 235919198 | 236919198 | Yes |
| ENSG00000132321.16 | IQCA1 | rs10180565 | 0.0000044 | ##### | Brain - Nucleus accumbens (b | 236419198 | 235919198 | 236919198 | Yes |
| ENSG00000132321.16 | IQCA1 | rs7355537 | 0.0000044 | ##### | Brain - Nucleus accumbens (b | 236419198 | 235919198 | 236919198 | Yes |
| ENSG00000132321.16 | IQCA1 | rs10203594 | 0.0000044 | ##### | Brain - Nucleus accumbens (b | 236419198 | 235919198 | 236919198 | Yes |
| ENSG00000132321.16 | IQCA1 | rs10183186 | 0.0000044 | ##### | Brain - Nucleus accumbens (b | 236419198 | 235919198 | 236919198 | Yes |
| ENSG00000132321.16 | IQCA1 | rs113822213 | 0.0000044 | ##### | Brain - Nucleus accumbens (b | 236419198 | 235919198 | 236919198 | Yes |
| ENSG00000132321.16 | IQCA1 | rs73129829 | 0.0000044 | ##### | Brain - Nucleus accumbens (b | 236419198 | 235919198 | 236919198 | Yes |
| ENSG00000132321.16 | IQCA1 | rs57240758 | 0.0000044 | ##### | Brain - Nucleus accumbens (b | 236419198 | 235919198 | 236919198 | Yes |
| ENSG00000132321.16 | IQCA1 | rs6724406 | 0.0000044 | ##### | Brain - Nucleus accumbens (b | 236419198 | 235919198 | 236919198 | Yes |
| ENSG00000132321.16 | IQCA1 | rs6719264 | 0.000005 | ##### | Brain - Nucleus accumbens (b | 236419198 | 235919198 | 236919198 | Yes |
| ENSG00000132321.16 | IQCA1 | rs76557076 | 0.0000051 | ##### | Brain - Nucleus accumbens (b | 236419198 | 235919198 | 236919198 | Yes |
| ENSG00000132321.16 | IQCA1 | rs11893167 | 0.0000051 | ##### | Brain - Nucleus accumbens (b | 236419198 | 235919198 | 236919198 | Yes |
| ENSG00000132321.16 | IQCA1 | rs7562715 | 0.0000054 | ##### | Brain - Cortex | 236419198 | 235919198 | 236919198 | Yes |
| ENSG00000132321.16 | IQCA1 | rs59024700 | 0.0000054 | ##### | Brain - Cortex | 236419198 | 235919198 | 236919198 | Yes |
| ENSG00000132321.16 | IQCA1 | rs73127867 | 0.0000054 | ##### | Brain - Cortex | 236419198 | 235919198 | 236919198 | Yes |

| | | | | | | | | | |
|---|---|---|---|---|---|---|---|---|---|
| ENSG00000132321.16 | IQCA1 | rs2046281 | 0.0000057 | ##### | Brain - Nucleus accumbens (b | 236419198 | 235919198 | 236919198 | Yes |
| ENSG00000132321.16 | IQCA1 | rs73998681 | 0.0000058 | ##### | Brain - Cortex | 236419198 | 235919198 | 236919198 | Yes |
| ENSG00000132321.16 | IQCA1 | rs10179644 | 0.0000059 | ##### | Brain - Frontal Cortex (BA9) | 236419198 | 235919198 | 236919198 | Yes |
| ENSG00000132321.16 | IQCA1 | rs73127879 | 0.0000059 | ##### | Brain - Frontal Cortex (BA9) | 236419198 | 235919198 | 236919198 | Yes |
| ENSG00000132321.16 | IQCA1 | rs73127880 | 0.0000059 | ##### | Brain - Frontal Cortex (BA9) | 236419198 | 235919198 | 236919198 | Yes |
| ENSG00000132321.16 | IQCA1 | rs1922798 | 0.0000059 | ##### | Brain - Frontal Cortex (BA9) | 236419198 | 235919198 | 236919198 | Yes |
| ENSG00000132321.16 | IQCA1 | rs73127887 | 0.0000059 | ##### | Brain - Frontal Cortex (BA9) | 236419198 | 235919198 | 236919198 | Yes |
| ENSG00000132321.16 | IQCA1 | rs80148244 | 0.0000059 | ##### | Brain - Frontal Cortex (BA9) | 236419198 | 235919198 | 236919198 | Yes |
| ENSG00000132321.16 | IQCA1 | rs73127891 | 0.0000059 | ##### | Brain - Frontal Cortex (BA9) | 236419198 | 235919198 | 236919198 | Yes |
| ENSG00000132321.16 | IQCA1 | rs76804064 | 0.0000059 | ##### | Brain - Frontal Cortex (BA9) | 236419198 | 235919198 | 236919198 | Yes |
| ENSG00000132321.16 | IQCA1 | rs78752798 | 0.0000059 | ##### | Brain - Frontal Cortex (BA9) | 236419198 | 235919198 | 236919198 | Yes |
| ENSG00000132321.16 | IQCA1 | rs10193278 | 0.0000059 | ##### | Brain - Frontal Cortex (BA9) | 236419198 | 235919198 | 236919198 | Yes |
| ENSG00000132321.16 | IQCA1 | rs79733838 | 0.0000059 | ##### | Brain - Frontal Cortex (BA9) | 236419198 | 235919198 | 236919198 | Yes |
| ENSG00000132321.16 | IQCA1 | rs79357137 | 0.0000059 | ##### | Brain - Frontal Cortex (BA9) | 236419198 | 235919198 | 236919198 | Yes |
| ENSG00000132321.16 | IQCA1 | rs13428016 | 0.000006 | ##### | Brain - Anterior cingulate cort | 236419198 | 235919198 | 236919198 | Yes |
| ENSG00000132321.16 | IQCA1 | rs10717671 | 0.000006 | ##### | Brain - Anterior cingulate cort | 236419198 | 235919198 | 236919198 | Yes |
| ENSG00000132321.16 | IQCA1 | rs2077827 | 0.0000062 | ##### | Brain - Nucleus accumbens (b | 236419198 | 235919198 | 236919198 | Yes |
| ENSG00000132321.16 | IQCA1 | rs2077828 | 0.0000062 | ##### | Brain - Nucleus accumbens (b | 236419198 | 235919198 | 236919198 | Yes |
| ENSG00000132321.16 | IQCA1 | rs73129834 | 0.0000062 | ##### | Brain - Nucleus accumbens (b | 236419198 | 235919198 | 236919198 | Yes |
| ENSG00000132321.16 | IQCA1 | rs7593458 | 0.0000063 | ##### | Brain - Nucleus accumbens (b | 236419198 | 235919198 | 236919198 | Yes |
| ENSG00000132321.16 | IQCA1 | rs66808092 | 0.0000063 | ##### | Brain - Nucleus accumbens (b | 236419198 | 235919198 | 236919198 | Yes |
| ENSG00000132321.16 | IQCA1 | rs73127885 | 0.0000067 | ##### | Brain - Nucleus accumbens (b | 236419198 | 235919198 | 236919198 | Yes |
| ENSG00000132321.16 | IQCA1 | rs62192377 | 0.0000069 | ##### | Brain - Nucleus accumbens (b | 236419198 | 235919198 | 236919198 | Yes |
| ENSG00000132321.16 | IQCA1 | rs6431468 | 0.000007 | ##### | Brain - Amygdala | 236419198 | 235919198 | 236919198 | Yes |
| ENSG00000132321.16 | IQCA1 | rs6431469 | 0.000007 | ##### | Brain - Amygdala | 236419198 | 235919198 | 236919198 | Yes |
| ENSG00000132321.16 | IQCA1 | rs77873497 | 0.0000071 | ##### | Brain - Amygdala | 236419198 | 235919198 | 236919198 | Yes |
| ENSG00000132321.16 | IQCA1 | rs75481888 | 0.0000071 | ##### | Brain - Amygdala | 236419198 | 235919198 | 236919198 | Yes |
| ENSG00000132321.16 | IQCA1 | rs6748498 | 0.0000072 | ##### | Brain - Anterior cingulate cort | 236419198 | 235919198 | 236919198 | Yes |
| ENSG00000132321.16 | IQCA1 | rs6734045 | 0.0000072 | ##### | Brain - Anterior cingulate cort | 236419198 | 235919198 | 236919198 | Yes |
| ENSG00000132321.16 | IQCA1 | rs77873497 | 0.0000076 | ##### | Brain - Anterior cingulate cort | 236419198 | 235919198 | 236919198 | Yes |
| ENSG00000132321.16 | IQCA1 | rs75481888 | 0.0000076 | ##### | Brain - Anterior cingulate cort | 236419198 | 235919198 | 236919198 | Yes |
| ENSG00000132321.16 | IQCA1 | rs77005664 | 0.0000076 | ##### | Brain - Anterior cingulate cort | 236419198 | 235919198 | 236919198 | Yes |

| | | | | | | | | | |
|---|---|---|---|---|---|---|---|---|---|
| ENSG00000132321.16 | IQCA1 | rs80050849 | 0.0000095 | ##### | Brain - Anterior cingulate cort | 236419198 | 235919198 | 236919198 | Yes |
| ENSG00000132321.16 | IQCA1 | rs116120818 | 0.0000098 | ##### | Brain - Spinal cord (cervical c- | 236419198 | 235919198 | 236919198 | Yes |
| ENSG00000132321.16 | IQCA1 | rs6713722 | 0.0000098 | ##### | Brain - Spinal cord (cervical c- | 236419198 | 235919198 | 236919198 | Yes |
| ENSG00000132321.16 | IQCA1 | rs13428016 | 0.00001 | ##### | Brain - Cerebellar Hemisphere | 236419198 | 235919198 | 236919198 | Yes |
| ENSG00000132321.16 | IQCA1 | rs10717671 | 0.00001 | ##### | Brain - Cerebellar Hemisphere | 236419198 | 235919198 | 236919198 | Yes |
| ENSG00000132321.16 | IQCA1 | rs58376523 | 0.000011 | ##### | Brain - Cortex | 236419198 | 235919198 | 236919198 | Yes |
| ENSG00000132321.16 | IQCA1 | rs74687797 | 0.000011 | ##### | Brain - Spinal cord (cervical c- | 236419198 | 235919198 | 236919198 | Yes |
| ENSG00000132321.16 | IQCA1 | rs115602727 | 0.000011 | ##### | Brain - Spinal cord (cervical c- | 236419198 | 235919198 | 236919198 | Yes |
| ENSG00000132321.16 | IQCA1 | rs114137147 | 0.000011 | ##### | Brain - Spinal cord (cervical c- | 236419198 | 235919198 | 236919198 | Yes |
| ENSG00000132321.16 | IQCA1 | rs139953264 | 0.000011 | ##### | Brain - Spinal cord (cervical c- | 236419198 | 235919198 | 236919198 | Yes |
| ENSG00000132321.16 | IQCA1 | rs78838583 | 0.000011 | ##### | Brain - Spinal cord (cervical c- | 236419198 | 235919198 | 236919198 | Yes |
| ENSG00000132321.16 | IQCA1 | rs78882965 | 0.000011 | ##### | Brain - Spinal cord (cervical c- | 236419198 | 235919198 | 236919198 | Yes |
| ENSG00000132321.16 | IQCA1 | rs77706882 | 0.000011 | ##### | Brain - Spinal cord (cervical c- | 236419198 | 235919198 | 236919198 | Yes |
| ENSG00000132321.16 | IQCA1 | rs79971923 | 0.000011 | ##### | Brain - Spinal cord (cervical c- | 236419198 | 235919198 | 236919198 | Yes |
| ENSG00000132321.16 | IQCA1 | rs114497817 | 0.000011 | ##### | Brain - Spinal cord (cervical c- | 236419198 | 235919198 | 236919198 | Yes |
| ENSG00000132321.16 | IQCA1 | rs140454346 | 0.000011 | ##### | Brain - Spinal cord (cervical c- | 236419198 | 235919198 | 236919198 | Yes |
| ENSG00000132321.16 | IQCA1 | rs116546052 | 0.000011 | ##### | Brain - Spinal cord (cervical c- | 236419198 | 235919198 | 236919198 | Yes |
| ENSG00000132321.16 | IQCA1 | rs75481811 | 0.000011 | ##### | Brain - Spinal cord (cervical c- | 236419198 | 235919198 | 236919198 | Yes |
| ENSG00000132321.16 | IQCA1 | rs6716884 | 0.000011 | ##### | Brain - Spinal cord (cervical c- | 236419198 | 235919198 | 236919198 | Yes |
| ENSG00000132321.16 | IQCA1 | rs6756394 | 0.000011 | ##### | Brain - Spinal cord (cervical c- | 236419198 | 235919198 | 236919198 | Yes |
| ENSG00000132321.16 | IQCA1 | rs6756884 | 0.000011 | ##### | Brain - Spinal cord (cervical c- | 236419198 | 235919198 | 236919198 | Yes |
| ENSG00000132321.16 | IQCA1 | rs143704622 | 0.000011 | ##### | Brain - Spinal cord (cervical c- | 236419198 | 235919198 | 236919198 | Yes |
| ENSG00000132321.16 | IQCA1 | rs76816500 | 0.000011 | ##### | Brain - Spinal cord (cervical c- | 236419198 | 235919198 | 236919198 | Yes |
| ENSG00000132321.16 | IQCA1 | rs115939355 | 0.000011 | ##### | Brain - Spinal cord (cervical c- | 236419198 | 235919198 | 236919198 | Yes |
| ENSG00000132321.16 | IQCA1 | rs115001042 | 0.000011 | ##### | Brain - Spinal cord (cervical c- | 236419198 | 235919198 | 236919198 | Yes |
| ENSG00000132321.16 | IQCA1 | rs73127884 | 0.000011 | ##### | Brain - Cortex | 236419198 | 235919198 | 236919198 | Yes |
| ENSG00000132321.16 | IQCA1 | rs77005664 | 0.000011 | ##### | Brain - Amygdala | 236419198 | 235919198 | 236919198 | Yes |
| ENSG00000132321.16 | IQCA1 | rs75404972 | 0.000011 | ##### | Brain - Amygdala | 236419198 | 235919198 | 236919198 | Yes |
| ENSG00000132321.16 | IQCA1 | rs78551120 | 0.000011 | ##### | Brain - Amygdala | 236419198 | 235919198 | 236919198 | Yes |
| ENSG00000132321.16 | IQCA1 | rs79056581 | 0.000011 | ##### | Brain - Amygdala | 236419198 | 235919198 | 236919198 | Yes |
| ENSG00000132321.16 | IQCA1 | rs145676572 | 0.000011 | ##### | Brain - Amygdala | 236419198 | 235919198 | 236919198 | Yes |
| ENSG00000132321.16 | IQCA1 | rs77342389 | 0.000011 | ##### | Brain - Amygdala | 236419198 | 235919198 | 236919198 | Yes |

| | | | | | | | | | |
|---|---|---|---|---|---|---|---|---|---|
| ENSG00000132321.16 | IQCA1 | rs117686211 | 0.000011 | ##### | Brain - Amygdala | 236419198 | 235919198 | 236919198 | Yes |
| ENSG00000132321.16 | IQCA1 | rs77611647 | 0.000012 | ##### | Brain - Cortex | 236419198 | 235919198 | 236919198 | Yes |
| ENSG00000132321.16 | IQCA1 | rs6431456 | 0.000013 | ##### | Brain - Cerebellar Hemisphere | 236419198 | 235919198 | 236919198 | Yes |
| ENSG00000132321.16 | IQCA1 | rs7557194 | 0.000013 | ##### | Brain - Cerebellar Hemisphere | 236419198 | 235919198 | 236919198 | Yes |
| ENSG00000132321.16 | IQCA1 | rs55656551 | 0.000013 | ##### | Brain - Cerebellar Hemisphere | 236419198 | 235919198 | 236919198 | Yes |
| ENSG00000132321.16 | IQCA1 | rs4060823 | 0.000013 | ##### | Brain - Cerebellar Hemisphere | 236419198 | 235919198 | 236919198 | Yes |
| ENSG00000132321.16 | IQCA1 | rs79086412 | 0.000013 | ##### | Brain - Cortex | 236419198 | 235919198 | 236919198 | No |
| ENSG00000132321.16 | IQCA1 | rs78244458 | 0.000013 | ##### | Brain - Cortex | 236419198 | 235919198 | 236919198 | No |
| ENSG00000132321.16 | IQCA1 | rs79175791 | 0.000013 | ##### | Brain - Cortex | 236419198 | 235919198 | 236919198 | No |
| ENSG00000132321.16 | IQCA1 | rs75404972 | 0.000014 | ##### | Brain - Anterior cingulate cort | 236419198 | 235919198 | 236919198 | Yes |
| ENSG00000132321.16 | IQCA1 | rs78551120 | 0.000014 | ##### | Brain - Anterior cingulate cort | 236419198 | 235919198 | 236919198 | Yes |
| ENSG00000132321.16 | IQCA1 | rs79056581 | 0.000014 | ##### | Brain - Anterior cingulate cort | 236419198 | 235919198 | 236919198 | Yes |
| ENSG00000132321.16 | IQCA1 | rs145676572 | 0.000014 | ##### | Brain - Anterior cingulate cort | 236419198 | 235919198 | 236919198 | Yes |
| ENSG00000132321.16 | IQCA1 | rs77342389 | 0.000014 | ##### | Brain - Anterior cingulate cort | 236419198 | 235919198 | 236919198 | Yes |
| ENSG00000132321.16 | IQCA1 | rs117686211 | 0.000014 | ##### | Brain - Anterior cingulate cort | 236419198 | 235919198 | 236919198 | Yes |
| ENSG00000132321.16 | IQCA1 | rs62192368 | 0.000014 | ##### | Brain - Nucleus accumbens (b | 236419198 | 235919198 | 236919198 | Yes |
| ENSG00000132321.16 | IQCA1 | rs80225031 | 0.000014 | ##### | Brain - Nucleus accumbens (b | 236419198 | 235919198 | 236919198 | Yes |
| ENSG00000132321.16 | IQCA1 | rs67796631 | 0.000014 | ##### | Brain - Anterior cingulate cort | 236419198 | 235919198 | 236919198 | Yes |
| ENSG00000132321.16 | IQCA1 | rs76557076 | 0.000014 | ##### | Brain - Frontal Cortex (BA9) | 236419198 | 235919198 | 236919198 | Yes |
| ENSG00000132321.16 | IQCA1 | rs11893167 | 0.000014 | ##### | Brain - Frontal Cortex (BA9) | 236419198 | 235919198 | 236919198 | Yes |
| ENSG00000132321.16 | IQCA1 | rs2077827 | 0.000014 | ##### | Brain - Frontal Cortex (BA9) | 236419198 | 235919198 | 236919198 | Yes |
| ENSG00000132321.16 | IQCA1 | rs2077828 | 0.000014 | ##### | Brain - Frontal Cortex (BA9) | 236419198 | 235919198 | 236919198 | Yes |
| ENSG00000132321.16 | IQCA1 | rs73129834 | 0.000014 | ##### | Brain - Frontal Cortex (BA9) | 236419198 | 235919198 | 236919198 | Yes |
| ENSG00000132321.16 | IQCA1 | rs13392175 | 0.000016 | ##### | Brain - Anterior cingulate cort | 236419198 | 235919198 | 236919198 | Yes |
| ENSG00000132321.16 | IQCA1 | rs150879391 | 0.000016 | ##### | Brain - Anterior cingulate cort | 236419198 | 235919198 | 236919198 | Yes |
| ENSG00000132321.16 | IQCA1 | rs62192377 | 0.000016 | ##### | Brain - Cerebellar Hemisphere | 236419198 | 235919198 | 236919198 | Yes |
| ENSG00000132321.16 | IQCA1 | rs6758710 | 0.000017 | ##### | Brain - Cerebellar Hemisphere | 236419198 | 235919198 | 236919198 | Yes |
| ENSG00000132321.16 | IQCA1 | rs62192376 | 0.000017 | ##### | Brain - Cerebellar Hemisphere | 236419198 | 235919198 | 236919198 | Yes |
| ENSG00000132321.16 | IQCA1 | rs114038160 | 0.000017 | ##### | Brain - Anterior cingulate cort | 236419198 | 235919198 | 236919198 | Yes |
| ENSG00000132321.16 | IQCA1 | rs75547606 | 0.000017 | ##### | Brain - Anterior cingulate cort | 236419198 | 235919198 | 236919198 | Yes |
| ENSG00000132321.16 | IQCA1 | rs79651798 | 0.000017 | ##### | Brain - Anterior cingulate cort | 236419198 | 235919198 | 236919198 | Yes |
| ENSG00000132321.16 | IQCA1 | rs114605323 | 0.000017 | ##### | Brain - Anterior cingulate cort | 236419198 | 235919198 | 236919198 | Yes |

| | | | | | | | | | |
|---|---|---|---|---|---|---|---|---|---|
| ENSG00000132321.16 | IQCA1 | rs76213802 | 0.000017 | ##### | Brain - Anterior cingulate cort | 236419198 | 235919198 | 236919198 | Yes |
| ENSG00000132321.16 | IQCA1 | rs78730330 | 0.000017 | ##### | Brain - Anterior cingulate cort | 236419198 | 235919198 | 236919198 | Yes |
| ENSG00000132321.16 | IQCA1 | rs75011852 | 0.000017 | ##### | Brain - Anterior cingulate cort | 236419198 | 235919198 | 236919198 | Yes |
| ENSG00000132321.16 | IQCA1 | rs118021539 | 0.000017 | ##### | Brain - Anterior cingulate cort | 236419198 | 235919198 | 236919198 | Yes |
| ENSG00000132321.16 | IQCA1 | rs117687759 | 0.000017 | ##### | Brain - Anterior cingulate cort | 236419198 | 235919198 | 236919198 | Yes |
| ENSG00000132321.16 | IQCA1 | rs78237430 | 0.000017 | ##### | Brain - Anterior cingulate cort | 236419198 | 235919198 | 236919198 | Yes |
| ENSG00000132321.16 | IQCA1 | rs80093468 | 0.000017 | ##### | Brain - Anterior cingulate cort | 236419198 | 235919198 | 236919198 | Yes |
| ENSG00000132321.16 | IQCA1 | rs74765120 | 0.000017 | ##### | Brain - Anterior cingulate cort | 236419198 | 235919198 | 236919198 | Yes |
| ENSG00000132321.16 | IQCA1 | rs77870186 | 0.000018 | ##### | Brain - Anterior cingulate cort | 236419198 | 235919198 | 236919198 | Yes |
| ENSG00000132321.16 | IQCA1 | rs139053340 | 0.000018 | ##### | Brain - Spinal cord (cervical c- | 236419198 | 235919198 | 236919198 | Yes |
| ENSG00000132321.16 | IQCA1 | rs13412178 | 0.000018 | ##### | Brain - Spinal cord (cervical c- | 236419198 | 235919198 | 236919198 | Yes |
| ENSG00000132321.16 | IQCA1 | rs10198334 | 0.000018 | ##### | Brain - Spinal cord (cervical c- | 236419198 | 235919198 | 236919198 | Yes |
| ENSG00000132321.16 | IQCA1 | rs28867102 | 0.000018 | ##### | Brain - Spinal cord (cervical c- | 236419198 | 235919198 | 236919198 | Yes |
| ENSG00000132321.16 | IQCA1 | rs62192371 | 0.000018 | ##### | Brain - Nucleus accumbens (b | 236419198 | 235919198 | 236919198 | Yes |
| ENSG00000132321.16 | IQCA1 | rs62192372 | 0.000018 | ##### | Brain - Nucleus accumbens (b | 236419198 | 235919198 | 236919198 | Yes |
| ENSG00000132321.16 | IQCA1 | rs780236452 | 0.000019 | ##### | Brain - Anterior cingulate cort | 236419198 | 235919198 | 236919198 | Yes |
| ENSG00000132321.16 | IQCA1 | rs116562332 | 0.000021 | ##### | Brain - Spinal cord (cervical c- | 236419198 | 235919198 | 236919198 | Yes |
| ENSG00000132321.16 | IQCA1 | rs76317401 | 0.000021 | ##### | Brain - Cortex | 236419198 | 235919198 | 236919198 | Yes |
| ENSG00000132321.16 | IQCA1 | rs58605253 | 0.000021 | ##### | Brain - Cortex | 236419198 | 235919198 | 236919198 | Yes |
| ENSG00000132321.16 | IQCA1 | rs59625470 | 0.000021 | ##### | Brain - Cortex | 236419198 | 235919198 | 236919198 | Yes |
| ENSG00000132321.16 | IQCA1 | rs74941416 | 0.000021 | ##### | Brain - Cortex | 236419198 | 235919198 | 236919198 | Yes |
| ENSG00000132321.16 | IQCA1 | rs115983368 | 0.000021 | ##### | Brain - Cortex | 236419198 | 235919198 | 236919198 | Yes |
| ENSG00000132321.16 | IQCA1 | rs73998612 | 0.000021 | ##### | Brain - Cortex | 236419198 | 235919198 | 236919198 | Yes |
| ENSG00000132321.16 | IQCA1 | rs76020172 | 0.000021 | ##### | Brain - Cortex | 236419198 | 235919198 | 236919198 | Yes |
| ENSG00000132321.16 | IQCA1 | rs73127885 | 0.000022 | ##### | Brain - Frontal Cortex (BA9) | 236419198 | 235919198 | 236919198 | Yes |
| ENSG00000132321.16 | IQCA1 | rs141245885 | 0.000023 | ##### | Brain - Anterior cingulate cort | 236419198 | 235919198 | 236919198 | Yes |
| ENSG00000132321.16 | IQCA1 | rs72978619 | 0.000023 | ##### | Brain - Anterior cingulate cort | 236419198 | 235919198 | 236919198 | Yes |
| ENSG00000132321.16 | IQCA1 | rs13427855 | 0.000024 | ##### | Brain - Nucleus accumbens (b | 236419198 | 235919198 | 236919198 | Yes |
| ENSG00000132321.16 | IQCA1 | rs59403004 | 0.000024 | ##### | Brain - Nucleus accumbens (b | 236419198 | 235919198 | 236919198 | Yes |
| ENSG00000132321.16 | IQCA1 | rs7593458 | 0.000026 | ##### | Brain - Anterior cingulate cort | 236419198 | 235919198 | 236919198 | Yes |
| ENSG00000132321.16 | IQCA1 | rs66808092 | 0.000026 | ##### | Brain - Anterior cingulate cort | 236419198 | 235919198 | 236919198 | Yes |
| ENSG00000132321.16 | IQCA1 | rs62192370 | 0.000028 | ##### | Brain - Nucleus accumbens (b | 236419198 | 235919198 | 236919198 | Yes |

| | | | | | | | | | |
|---|---|---|---|---|---|---|---|---|---|
| ENSG00000132321.16 | IQCA1 | rs6742302 | 0.000029 | ##### | Brain - Cerebellum | 236419198 | 235919198 | 236919198 | Yes |
| ENSG00000132321.16 | IQCA1 | rs6431457 | 0.00003 | ##### | Brain - Cerebellar Hemisphere | 236419198 | 235919198 | 236919198 | Yes |
| ENSG00000132321.16 | IQCA1 | rs62192361 | 0.00003 | ##### | Brain - Cerebellar Hemisphere | 236419198 | 235919198 | 236919198 | Yes |
| ENSG00000132321.16 | IQCA1 | rs6751722 | 0.000031 | ##### | Brain - Nucleus accumbens (b | 236419198 | 235919198 | 236919198 | Yes |
| ENSG00000132321.16 | IQCA1 | rs7593458 | 0.000033 | ##### | Brain - Cerebellar Hemisphere | 236419198 | 235919198 | 236919198 | Yes |
| ENSG00000132321.16 | IQCA1 | rs66808092 | 0.000033 | ##### | Brain - Cerebellar Hemisphere | 236419198 | 235919198 | 236919198 | Yes |
| ENSG00000132321.16 | IQCA1 | rs7585499 | 0.000035 | ##### | Brain - Cerebellar Hemisphere | 236419198 | 235919198 | 236919198 | Yes |
| ENSG00000132321.16 | IQCA1 | rs60242756 | 0.000043 | ##### | Brain - Cortex | 236419198 | 235919198 | 236919198 | Yes |
| ENSG00000132321.16 | IQCA1 | rs77704052 | 0.000043 | ##### | Brain - Cortex | 236419198 | 235919198 | 236919198 | Yes |
| ENSG00000132321.16 | IQCA1 | rs79825641 | 0.000043 | ##### | Brain - Cortex | 236419198 | 235919198 | 236919198 | Yes |
| ENSG00000132321.16 | IQCA1 | rs144917370 | 0.000043 | ##### | Brain - Cortex | 236419198 | 235919198 | 236919198 | Yes |
| ENSG00000132321.16 | IQCA1 | rs10193029 | 0.000045 | ##### | Brain - Cerebellum | 236419198 | 235919198 | 236919198 | Yes |
| ENSG00000132321.16 | IQCA1 | rs13416077 | 0.000045 | ##### | Brain - Cerebellum | 236419198 | 235919198 | 236919198 | Yes |
| ENSG00000132321.16 | IQCA1 | rs7600965 | 0.000045 | ##### | Brain - Cerebellum | 236419198 | 235919198 | 236919198 | Yes |
| ENSG00000132321.16 | IQCA1 | rs62192371 | 0.000053 | ##### | Brain - Cerebellar Hemisphere | 236419198 | 235919198 | 236919198 | Yes |
| ENSG00000132321.16 | IQCA1 | rs62192372 | 0.000053 | ##### | Brain - Cerebellar Hemisphere | 236419198 | 235919198 | 236919198 | Yes |
| ENSG00000132321.16 | IQCA1 | rs60507206 | 0.000061 | ##### | Brain - Cortex | 236419198 | 235919198 | 236919198 | No |
| ENSG00000132321.16 | IQCA1 | rs62192377 | 0.000064 | ##### | Brain - Cerebellum | 236419198 | 235919198 | 236919198 | Yes |
| ENSG00000132321.16 | IQCA1 | rs6758710 | 0.000067 | ##### | Brain - Cerebellum | 236419198 | 235919198 | 236919198 | Yes |
| ENSG00000132321.16 | IQCA1 | rs62192376 | 0.000067 | ##### | Brain - Cerebellum | 236419198 | 235919198 | 236919198 | Yes |
| ENSG00000196296.13 | ATP2A1 | rs180743 | 0.0000095 | ##### | Brain - Hypothalamus | 28904132 | 28404132 | 29404132 | Yes |
| ENSG00000196296.13 | ATP2A1 | rs180744 | 0.0000095 | ##### | Brain - Hypothalamus | 28904132 | 28404132 | 29404132 | Yes |
| ENSG00000196296.13 | ATP2A1 | rs149271 | 0.000011 | ##### | Brain - Hypothalamus | 28904132 | 28404132 | 29404132 | Yes |
| ENSG00000196296.13 | ATP2A1 | rs148114931 | 0.000019 | ##### | Brain - Hypothalamus | 28904132 | 28404132 | 29404132 | Yes |
| ENSG00000196296.13 | ATP2A1 | rs40831 | 0.00002 | ##### | Brain - Hypothalamus | 28904132 | 28404132 | 29404132 | Yes |
| ENSG00000196296.13 | ATP2A1 | rs118036722 | 0.000023 | ##### | Brain - Frontal Cortex (BA9) | 28904132 | 28404132 | 29404132 | No |
| ENSG00000196296.13 | ATP2A1 | rs151174 | 0.000038 | ##### | Brain - Hypothalamus | 28904132 | 28404132 | 29404132 | Yes |
| ENSG00000125734.15 | GPR108 | rs340135 | 1.30E-48 | ##### | Brain - Cortex | 6732114 | 6232114 | 7232114 | Yes |
| ENSG00000125734.15 | GPR108 | rs340141 | 6.50E-48 | ##### | Brain - Cortex | 6732114 | 6232114 | 7232114 | Yes |
| ENSG00000125734.15 | GPR108 | rs344633 | 6.50E-48 | ##### | Brain - Cortex | 6732114 | 6232114 | 7232114 | Yes |
| ENSG00000125734.15 | GPR108 | rs173174 | 6.50E-48 | ##### | Brain - Cortex | 6732114 | 6232114 | 7232114 | Yes |
| ENSG00000125734.15 | GPR108 | rs3810222 | 7.10E-43 | ##### | Brain - Cortex | 6732114 | 6232114 | 7232114 | Yes |

| | | | | | | | | | |
|---|---|---|---|---|---|---|---|---|---|
| ENSG00000125734.15 | GPR108 | rs3810223 | 7.10E-43 | ##### | Brain - Cortex | 6732114 | 6232114 | 7232114 | Yes |
| ENSG00000125734.15 | GPR108 | rs10410021 | 7.10E-43 | ##### | Brain - Cortex | 6732114 | 6232114 | 7232114 | Yes |
| ENSG00000125734.15 | GPR108 | rs10419080 | 7.10E-43 | ##### | Brain - Cortex | 6732114 | 6232114 | 7232114 | Yes |
| ENSG00000125734.15 | GPR108 | rs11085200 | 4.90E-42 | ##### | Brain - Cortex | 6732114 | 6232114 | 7232114 | Yes |
| ENSG00000125734.15 | GPR108 | rs340135 | 7.80E-42 | ##### | Brain - Cerebellum | 6732114 | 6232114 | 7232114 | Yes |
| ENSG00000125734.15 | GPR108 | rs340141 | 1.70E-41 | ##### | Brain - Cerebellum | 6732114 | 6232114 | 7232114 | Yes |
| ENSG00000125734.15 | GPR108 | rs344633 | 1.70E-41 | ##### | Brain - Cerebellum | 6732114 | 6232114 | 7232114 | Yes |
| ENSG00000125734.15 | GPR108 | rs173174 | 1.70E-41 | ##### | Brain - Cerebellum | 6732114 | 6232114 | 7232114 | Yes |
| ENSG00000125734.15 | GPR108 | rs3810222 | 3.00E-38 | ##### | Brain - Cerebellum | 6732114 | 6232114 | 7232114 | Yes |
| ENSG00000125734.15 | GPR108 | rs3810223 | 3.00E-38 | ##### | Brain - Cerebellum | 6732114 | 6232114 | 7232114 | Yes |
| ENSG00000125734.15 | GPR108 | rs10410021 | 3.00E-38 | ##### | Brain - Cerebellum | 6732114 | 6232114 | 7232114 | Yes |
| ENSG00000125734.15 | GPR108 | rs10419080 | 3.00E-38 | ##### | Brain - Cerebellum | 6732114 | 6232114 | 7232114 | Yes |
| ENSG00000125734.15 | GPR108 | rs11085200 | 1.80E-37 | ##### | Brain - Cerebellum | 6732114 | 6232114 | 7232114 | Yes |
| ENSG00000125734.15 | GPR108 | rs340135 | 4.10E-35 | ##### | Brain - Frontal Cortex (BA9) | 6732114 | 6232114 | 7232114 | Yes |
| ENSG00000125734.15 | GPR108 | rs340141 | 4.80E-34 | ##### | Brain - Frontal Cortex (BA9) | 6732114 | 6232114 | 7232114 | Yes |
| ENSG00000125734.15 | GPR108 | rs344633 | 4.80E-34 | ##### | Brain - Frontal Cortex (BA9) | 6732114 | 6232114 | 7232114 | Yes |
| ENSG00000125734.15 | GPR108 | rs173174 | 4.80E-34 | ##### | Brain - Frontal Cortex (BA9) | 6732114 | 6232114 | 7232114 | Yes |
| ENSG00000125734.15 | GPR108 | rs11085200 | 5.20E-32 | ##### | Brain - Frontal Cortex (BA9) | 6732114 | 6232114 | 7232114 | Yes |
| ENSG00000125734.15 | GPR108 | rs3810222 | 1.20E-30 | ##### | Brain - Frontal Cortex (BA9) | 6732114 | 6232114 | 7232114 | Yes |
| ENSG00000125734.15 | GPR108 | rs3810223 | 1.20E-30 | ##### | Brain - Frontal Cortex (BA9) | 6732114 | 6232114 | 7232114 | Yes |
| ENSG00000125734.15 | GPR108 | rs10410021 | 1.20E-30 | ##### | Brain - Frontal Cortex (BA9) | 6732114 | 6232114 | 7232114 | Yes |
| ENSG00000125734.15 | GPR108 | rs10419080 | 1.20E-30 | ##### | Brain - Frontal Cortex (BA9) | 6732114 | 6232114 | 7232114 | Yes |
| ENSG00000125734.15 | GPR108 | rs340135 | 8.90E-30 | ##### | Brain - Cerebellar Hemisphere | 6732114 | 6232114 | 7232114 | Yes |
| ENSG00000125734.15 | GPR108 | rs340141 | 2.40E-29 | ##### | Brain - Cerebellar Hemisphere | 6732114 | 6232114 | 7232114 | Yes |
| ENSG00000125734.15 | GPR108 | rs344633 | 2.40E-29 | ##### | Brain - Cerebellar Hemisphere | 6732114 | 6232114 | 7232114 | Yes |
| ENSG00000125734.15 | GPR108 | rs173174 | 2.40E-29 | ##### | Brain - Cerebellar Hemisphere | 6732114 | 6232114 | 7232114 | Yes |
| ENSG00000125734.15 | GPR108 | rs11085200 | 6.50E-28 | ##### | Brain - Cerebellar Hemisphere | 6732114 | 6232114 | 7232114 | Yes |
| ENSG00000125734.15 | GPR108 | rs340135 | 7.00E-28 | ##### | Brain - Hippocampus | 6732114 | 6232114 | 7232114 | Yes |
| ENSG00000125734.15 | GPR108 | rs3810222 | 1.50E-27 | ##### | Brain - Cerebellar Hemisphere | 6732114 | 6232114 | 7232114 | Yes |
| ENSG00000125734.15 | GPR108 | rs3810223 | 1.50E-27 | ##### | Brain - Cerebellar Hemisphere | 6732114 | 6232114 | 7232114 | Yes |
| ENSG00000125734.15 | GPR108 | rs10410021 | 1.50E-27 | ##### | Brain - Cerebellar Hemisphere | 6732114 | 6232114 | 7232114 | Yes |
| ENSG00000125734.15 | GPR108 | rs10419080 | 1.50E-27 | ##### | Brain - Cerebellar Hemisphere | 6732114 | 6232114 | 7232114 | Yes |

| | | | | | | | | | |
|---|---|---|---|---|---|---|---|---|---|
| ENSG00000125734.15 | GPR108 | rs10853997 | 1.60E-27 | ##### | Brain - Cortex | 6732114 | 6232114 | 7232114 | Yes |
| ENSG00000125734.15 | GPR108 | rs340135 | 1.70E-26 | ##### | Brain - Caudate (basal ganglia | 6732114 | 6232114 | 7232114 | Yes |
| ENSG00000125734.15 | GPR108 | rs340135 | 7.20E-26 | ##### | Brain - Nucleus accumbens (b | 6732114 | 6232114 | 7232114 | Yes |
| ENSG00000125734.15 | GPR108 | rs340141 | 1.80E-25 | ##### | Brain - Caudate (basal ganglia | 6732114 | 6232114 | 7232114 | Yes |
| ENSG00000125734.15 | GPR108 | rs344633 | 1.80E-25 | ##### | Brain - Caudate (basal ganglia | 6732114 | 6232114 | 7232114 | Yes |
| ENSG00000125734.15 | GPR108 | rs173174 | 1.80E-25 | ##### | Brain - Caudate (basal ganglia | 6732114 | 6232114 | 7232114 | Yes |
| ENSG00000125734.15 | GPR108 | rs340135 | 3.30E-25 | ##### | Brain - Anterior cingulate cort | 6732114 | 6232114 | 7232114 | Yes |
| ENSG00000125734.15 | GPR108 | rs340141 | 4.40E-25 | ##### | Brain - Anterior cingulate cort | 6732114 | 6232114 | 7232114 | Yes |
| ENSG00000125734.15 | GPR108 | rs344633 | 4.40E-25 | ##### | Brain - Anterior cingulate cort | 6732114 | 6232114 | 7232114 | Yes |
| ENSG00000125734.15 | GPR108 | rs173174 | 4.40E-25 | ##### | Brain - Anterior cingulate cort | 6732114 | 6232114 | 7232114 | Yes |
| ENSG00000125734.15 | GPR108 | rs340141 | 4.50E-25 | ##### | Brain - Nucleus accumbens (b | 6732114 | 6232114 | 7232114 | Yes |
| ENSG00000125734.15 | GPR108 | rs344633 | 4.50E-25 | ##### | Brain - Nucleus accumbens (b | 6732114 | 6232114 | 7232114 | Yes |
| ENSG00000125734.15 | GPR108 | rs173174 | 4.50E-25 | ##### | Brain - Nucleus accumbens (b | 6732114 | 6232114 | 7232114 | Yes |
| ENSG00000125734.15 | GPR108 | rs340141 | 5.50E-25 | ##### | Brain - Hippocampus | 6732114 | 6232114 | 7232114 | Yes |
| ENSG00000125734.15 | GPR108 | rs344633 | 5.50E-25 | ##### | Brain - Hippocampus | 6732114 | 6232114 | 7232114 | Yes |
| ENSG00000125734.15 | GPR108 | rs173174 | 5.50E-25 | ##### | Brain - Hippocampus | 6732114 | 6232114 | 7232114 | Yes |
| ENSG00000125734.15 | GPR108 | rs11085200 | 1.20E-24 | ##### | Brain - Anterior cingulate cort | 6732114 | 6232114 | 7232114 | Yes |
| ENSG00000125734.15 | GPR108 | rs3810222 | 2.50E-24 | ##### | Brain - Anterior cingulate cort | 6732114 | 6232114 | 7232114 | Yes |
| ENSG00000125734.15 | GPR108 | rs3810223 | 2.50E-24 | ##### | Brain - Anterior cingulate cort | 6732114 | 6232114 | 7232114 | Yes |
| ENSG00000125734.15 | GPR108 | rs10410021 | 2.50E-24 | ##### | Brain - Anterior cingulate cort | 6732114 | 6232114 | 7232114 | Yes |
| ENSG00000125734.15 | GPR108 | rs10419080 | 2.50E-24 | ##### | Brain - Anterior cingulate cort | 6732114 | 6232114 | 7232114 | Yes |
| ENSG00000125734.15 | GPR108 | rs340135 | 6.80E-24 | ##### | Brain - Putamen (basal gangli | 6732114 | 6232114 | 7232114 | Yes |
| ENSG00000125734.15 | GPR108 | rs3810222 | 9.20E-24 | ##### | Brain - Hippocampus | 6732114 | 6232114 | 7232114 | Yes |
| ENSG00000125734.15 | GPR108 | rs3810223 | 9.20E-24 | ##### | Brain - Hippocampus | 6732114 | 6232114 | 7232114 | Yes |
| ENSG00000125734.15 | GPR108 | rs10410021 | 9.20E-24 | ##### | Brain - Hippocampus | 6732114 | 6232114 | 7232114 | Yes |
| ENSG00000125734.15 | GPR108 | rs10419080 | 9.20E-24 | ##### | Brain - Hippocampus | 6732114 | 6232114 | 7232114 | Yes |
| ENSG00000125734.15 | GPR108 | rs10853997 | 2.30E-23 | ##### | Brain - Frontal Cortex (BA9) | 6732114 | 6232114 | 7232114 | Yes |
| ENSG00000125734.15 | GPR108 | rs3810222 | 2.90E-23 | ##### | Brain - Caudate (basal ganglia | 6732114 | 6232114 | 7232114 | Yes |
| ENSG00000125734.15 | GPR108 | rs3810223 | 2.90E-23 | ##### | Brain - Caudate (basal ganglia | 6732114 | 6232114 | 7232114 | Yes |
| ENSG00000125734.15 | GPR108 | rs10410021 | 2.90E-23 | ##### | Brain - Caudate (basal ganglia | 6732114 | 6232114 | 7232114 | Yes |
| ENSG00000125734.15 | GPR108 | rs10419080 | 2.90E-23 | ##### | Brain - Caudate (basal ganglia | 6732114 | 6232114 | 7232114 | Yes |
| ENSG00000125734.15 | GPR108 | rs11085200 | 3.90E-23 | ##### | Brain - Caudate (basal ganglia | 6732114 | 6232114 | 7232114 | Yes |

| | | | | | | | | | |
|---|---|---|---|---|---|---|---|---|---|
| ENSG00000125734.15 | GPR108 | rs3810222 | 4.00E-23 | ##### | Brain - Nucleus accumbens (b | 6732114 | 6232114 | 7232114 | Yes |
| ENSG00000125734.15 | GPR108 | rs3810223 | 4.00E-23 | ##### | Brain - Nucleus accumbens (b | 6732114 | 6232114 | 7232114 | Yes |
| ENSG00000125734.15 | GPR108 | rs10410021 | 4.00E-23 | ##### | Brain - Nucleus accumbens (b | 6732114 | 6232114 | 7232114 | Yes |
| ENSG00000125734.15 | GPR108 | rs10419080 | 4.00E-23 | ##### | Brain - Nucleus accumbens (b | 6732114 | 6232114 | 7232114 | Yes |
| ENSG00000125734.15 | GPR108 | rs11085200 | 5.50E-23 | ##### | Brain - Nucleus accumbens (b | 6732114 | 6232114 | 7232114 | Yes |
| ENSG00000125734.15 | GPR108 | rs10853997 | 1.10E-22 | ##### | Brain - Cerebellum | 6732114 | 6232114 | 7232114 | Yes |
| ENSG00000125734.15 | GPR108 | rs340141 | 1.30E-22 | ##### | Brain - Putamen (basal gangli | 6732114 | 6232114 | 7232114 | Yes |
| ENSG00000125734.15 | GPR108 | rs344633 | 1.30E-22 | ##### | Brain - Putamen (basal gangli | 6732114 | 6232114 | 7232114 | Yes |
| ENSG00000125734.15 | GPR108 | rs173174 | 1.30E-22 | ##### | Brain - Putamen (basal gangli | 6732114 | 6232114 | 7232114 | Yes |
| ENSG00000125734.15 | GPR108 | rs11085200 | 4.10E-22 | ##### | Brain - Hippocampus | 6732114 | 6232114 | 7232114 | Yes |
| ENSG00000125734.15 | GPR108 | rs2277986 | 4.40E-22 | ##### | Brain - Cortex | 6732114 | 6232114 | 7232114 | Yes |
| ENSG00000125734.15 | GPR108 | rs10853997 | 2.30E-20 | ##### | Brain - Cerebellar Hemisphere | 6732114 | 6232114 | 7232114 | Yes |
| ENSG00000125734.15 | GPR108 | rs3810222 | 4.20E-20 | ##### | Brain - Putamen (basal gangli | 6732114 | 6232114 | 7232114 | Yes |
| ENSG00000125734.15 | GPR108 | rs3810223 | 4.20E-20 | ##### | Brain - Putamen (basal gangli | 6732114 | 6232114 | 7232114 | Yes |
| ENSG00000125734.15 | GPR108 | rs10410021 | 4.20E-20 | ##### | Brain - Putamen (basal gangli | 6732114 | 6232114 | 7232114 | Yes |
| ENSG00000125734.15 | GPR108 | rs10419080 | 4.20E-20 | ##### | Brain - Putamen (basal gangli | 6732114 | 6232114 | 7232114 | Yes |
| ENSG00000125734.15 | GPR108 | rs11085200 | 9.20E-20 | ##### | Brain - Putamen (basal gangli | 6732114 | 6232114 | 7232114 | Yes |
| ENSG00000125734.15 | GPR108 | rs340135 | 1.70E-19 | ##### | Brain - Spinal cord (cervical c- | 6732114 | 6232114 | 7232114 | Yes |
| ENSG00000125734.15 | GPR108 | rs340141 | 1.70E-19 | ##### | Brain - Spinal cord (cervical c- | 6732114 | 6232114 | 7232114 | Yes |
| ENSG00000125734.15 | GPR108 | rs344633 | 1.70E-19 | ##### | Brain - Spinal cord (cervical c- | 6732114 | 6232114 | 7232114 | Yes |
| ENSG00000125734.15 | GPR108 | rs173174 | 1.70E-19 | ##### | Brain - Spinal cord (cervical c- | 6732114 | 6232114 | 7232114 | Yes |
| ENSG00000125734.15 | GPR108 | rs340135 | 7.90E-19 | ##### | Brain - Substantia nigra | 6732114 | 6232114 | 7232114 | Yes |
| ENSG00000125734.15 | GPR108 | rs340135 | 1.20E-18 | ##### | Brain - Hypothalamus | 6732114 | 6232114 | 7232114 | Yes |
| ENSG00000125734.15 | GPR108 | rs10853997 | 1.40E-18 | ##### | Brain - Nucleus accumbens (b | 6732114 | 6232114 | 7232114 | Yes |
| ENSG00000125734.15 | GPR108 | rs340135 | 1.60E-18 | ##### | Brain - Amygdala | 6732114 | 6232114 | 7232114 | Yes |
| ENSG00000125734.15 | GPR108 | rs340141 | 1.90E-18 | ##### | Brain - Hypothalamus | 6732114 | 6232114 | 7232114 | Yes |
| ENSG00000125734.15 | GPR108 | rs344633 | 1.90E-18 | ##### | Brain - Hypothalamus | 6732114 | 6232114 | 7232114 | Yes |
| ENSG00000125734.15 | GPR108 | rs173174 | 1.90E-18 | ##### | Brain - Hypothalamus | 6732114 | 6232114 | 7232114 | Yes |
| ENSG00000125734.15 | GPR108 | rs3810222 | 2.10E-18 | ##### | Brain - Spinal cord (cervical c- | 6732114 | 6232114 | 7232114 | Yes |
| ENSG00000125734.15 | GPR108 | rs3810223 | 2.10E-18 | ##### | Brain - Spinal cord (cervical c- | 6732114 | 6232114 | 7232114 | Yes |
| ENSG00000125734.15 | GPR108 | rs10410021 | 2.10E-18 | ##### | Brain - Spinal cord (cervical c- | 6732114 | 6232114 | 7232114 | Yes |
| ENSG00000125734.15 | GPR108 | rs10419080 | 2.10E-18 | ##### | Brain - Spinal cord (cervical c- | 6732114 | 6232114 | 7232114 | Yes |

| | | | | | | | | | |
|---|---|---|---|---|---|---|---|---|---|
| ENSG00000125734.15 | GPR108 | rs10853997 | 3.40E-18 | ##### | Brain - Putamen (basal gangli | 6732114 | 6232114 | 7232114 | Yes |
| ENSG00000125734.15 | GPR108 | rs340141 | 5.10E-18 | ##### | Brain - Amygdala | 6732114 | 6232114 | 7232114 | Yes |
| ENSG00000125734.15 | GPR108 | rs344633 | 5.10E-18 | ##### | Brain - Amygdala | 6732114 | 6232114 | 7232114 | Yes |
| ENSG00000125734.15 | GPR108 | rs173174 | 5.10E-18 | ##### | Brain - Amygdala | 6732114 | 6232114 | 7232114 | Yes |
| ENSG00000125734.15 | GPR108 | rs340141 | 5.60E-18 | ##### | Brain - Substantia nigra | 6732114 | 6232114 | 7232114 | Yes |
| ENSG00000125734.15 | GPR108 | rs344633 | 5.60E-18 | ##### | Brain - Substantia nigra | 6732114 | 6232114 | 7232114 | Yes |
| ENSG00000125734.15 | GPR108 | rs173174 | 5.60E-18 | ##### | Brain - Substantia nigra | 6732114 | 6232114 | 7232114 | Yes |
| ENSG00000125734.15 | GPR108 | rs10853997 | 2.30E-17 | ##### | Brain - Hippocampus | 6732114 | 6232114 | 7232114 | Yes |
| ENSG00000125734.15 | GPR108 | rs3810222 | 5.80E-17 | ##### | Brain - Hypothalamus | 6732114 | 6232114 | 7232114 | Yes |
| ENSG00000125734.15 | GPR108 | rs3810223 | 5.80E-17 | ##### | Brain - Hypothalamus | 6732114 | 6232114 | 7232114 | Yes |
| ENSG00000125734.15 | GPR108 | rs10410021 | 5.80E-17 | ##### | Brain - Hypothalamus | 6732114 | 6232114 | 7232114 | Yes |
| ENSG00000125734.15 | GPR108 | rs10419080 | 5.80E-17 | ##### | Brain - Hypothalamus | 6732114 | 6232114 | 7232114 | Yes |
| ENSG00000125734.15 | GPR108 | rs11085200 | 8.80E-17 | ##### | Brain - Spinal cord (cervical c- | 6732114 | 6232114 | 7232114 | Yes |
| ENSG00000125734.15 | GPR108 | rs3810222 | 1.40E-16 | ##### | Brain - Amygdala | 6732114 | 6232114 | 7232114 | Yes |
| ENSG00000125734.15 | GPR108 | rs3810223 | 1.40E-16 | ##### | Brain - Amygdala | 6732114 | 6232114 | 7232114 | Yes |
| ENSG00000125734.15 | GPR108 | rs10410021 | 1.40E-16 | ##### | Brain - Amygdala | 6732114 | 6232114 | 7232114 | Yes |
| ENSG00000125734.15 | GPR108 | rs10419080 | 1.40E-16 | ##### | Brain - Amygdala | 6732114 | 6232114 | 7232114 | Yes |
| ENSG00000125734.15 | GPR108 | rs11085200 | 1.40E-16 | ##### | Brain - Amygdala | 6732114 | 6232114 | 7232114 | Yes |
| ENSG00000125734.15 | GPR108 | rs11085200 | 2.00E-16 | ##### | Brain - Hypothalamus | 6732114 | 6232114 | 7232114 | Yes |
| ENSG00000125734.15 | GPR108 | rs3810222 | 2.50E-16 | ##### | Brain - Substantia nigra | 6732114 | 6232114 | 7232114 | Yes |
| ENSG00000125734.15 | GPR108 | rs3810223 | 2.50E-16 | ##### | Brain - Substantia nigra | 6732114 | 6232114 | 7232114 | Yes |
| ENSG00000125734.15 | GPR108 | rs10410021 | 2.50E-16 | ##### | Brain - Substantia nigra | 6732114 | 6232114 | 7232114 | Yes |
| ENSG00000125734.15 | GPR108 | rs10419080 | 2.50E-16 | ##### | Brain - Substantia nigra | 6732114 | 6232114 | 7232114 | Yes |
| ENSG00000125734.15 | GPR108 | rs10853997 | 4.20E-16 | ##### | Brain - Amygdala | 6732114 | 6232114 | 7232114 | Yes |
| ENSG00000125734.15 | GPR108 | rs12462822 | 4.60E-16 | ##### | Brain - Cortex | 6732114 | 6232114 | 7232114 | Yes |
| ENSG00000125734.15 | GPR108 | rs12462822 | 5.50E-16 | ##### | Brain - Frontal Cortex (BA9) | 6732114 | 6232114 | 7232114 | Yes |
| ENSG00000125734.15 | GPR108 | rs10853997 | 7.10E-16 | ##### | Brain - Hypothalamus | 6732114 | 6232114 | 7232114 | Yes |
| ENSG00000125734.15 | GPR108 | rs10853997 | 8.10E-16 | ##### | Brain - Caudate (basal ganglia | 6732114 | 6232114 | 7232114 | Yes |
| ENSG00000125734.15 | GPR108 | rs11085200 | 1.50E-15 | ##### | Brain - Substantia nigra | 6732114 | 6232114 | 7232114 | Yes |
| ENSG00000125734.15 | GPR108 | rs12462822 | 2.20E-15 | ##### | Brain - Cerebellar Hemisphere | 6732114 | 6232114 | 7232114 | Yes |
| ENSG00000125734.15 | GPR108 | rs339397 | 2.50E-15 | ##### | Brain - Frontal Cortex (BA9) | 6732114 | 6232114 | 7232114 | Yes |
| ENSG00000125734.15 | GPR108 | rs339397 | 5.40E-15 | ##### | Brain - Cortex | 6732114 | 6232114 | 7232114 | Yes |

| | | | | | | | | | |
|---|---|---|---|---|---|---|---|---|---|
| ENSG00000125734.15 | GPR108 | rs2277986 | 5.40E-15 | ##### | Brain - Cerebellum | 6732114 | 6232114 | 7232114 | Yes |
| ENSG00000125734.15 | GPR108 | rs163916 | 5.60E-15 | ##### | Brain - Frontal Cortex (BA9) | 6732114 | 6232114 | 7232114 | Yes |
| ENSG00000125734.15 | GPR108 | rs339399 | 5.60E-15 | ##### | Brain - Frontal Cortex (BA9) | 6732114 | 6232114 | 7232114 | Yes |
| ENSG00000125734.15 | GPR108 | rs163916 | 8.20E-15 | ##### | Brain - Cortex | 6732114 | 6232114 | 7232114 | Yes |
| ENSG00000125734.15 | GPR108 | rs339399 | 8.20E-15 | ##### | Brain - Cortex | 6732114 | 6232114 | 7232114 | Yes |
| ENSG00000125734.15 | GPR108 | rs10853997 | 1.60E-14 | ##### | Brain - Spinal cord (cervical c- | 6732114 | 6232114 | 7232114 | Yes |
| ENSG00000125734.15 | GPR108 | rs10853997 | 2.60E-14 | ##### | Brain - Anterior cingulate cort | 6732114 | 6232114 | 7232114 | Yes |
| ENSG00000125734.15 | GPR108 | rs12462822 | 4.70E-14 | ##### | Brain - Cerebellum | 6732114 | 6232114 | 7232114 | Yes |
| ENSG00000125734.15 | GPR108 | rs12462822 | 3.60E-13 | ##### | Brain - Putamen (basal gangli | 6732114 | 6232114 | 7232114 | Yes |
| ENSG00000125734.15 | GPR108 | rs12462822 | 5.00E-13 | ##### | Brain - Nucleus accumbens (b | 6732114 | 6232114 | 7232114 | Yes |
| ENSG00000125734.15 | GPR108 | rs2277986 | 6.10E-13 | ##### | Brain - Frontal Cortex (BA9) | 6732114 | 6232114 | 7232114 | Yes |
| ENSG00000125734.15 | GPR108 | rs2047140 | 9.60E-13 | ##### | Brain - Cortex | 6732114 | 6232114 | 7232114 | Yes |
| ENSG00000125734.15 | GPR108 | rs62125134 | 1.00E-12 | ##### | Brain - Frontal Cortex (BA9) | 6732114 | 6232114 | 7232114 | Yes |
| ENSG00000125734.15 | GPR108 | rs2047140 | 1.00E-12 | ##### | Brain - Frontal Cortex (BA9) | 6732114 | 6232114 | 7232114 | Yes |
| ENSG00000125734.15 | GPR108 | rs2277986 | 1.90E-12 | ##### | Brain - Putamen (basal gangli | 6732114 | 6232114 | 7232114 | Yes |
| ENSG00000125734.15 | GPR108 | rs2279623 | 1.90E-12 | ##### | Brain - Cortex | 6732114 | 6232114 | 7232114 | Yes |
| ENSG00000125734.15 | GPR108 | rs62125134 | 2.20E-12 | ##### | Brain - Cerebellar Hemisphere | 6732114 | 6232114 | 7232114 | Yes |
| ENSG00000125734.15 | GPR108 | rs10853997 | 2.70E-12 | ##### | Brain - Substantia nigra | 6732114 | 6232114 | 7232114 | Yes |
| ENSG00000125734.15 | GPR108 | rs340138 | 3.90E-12 | ##### | Brain - Cerebellum | 6732114 | 6232114 | 7232114 | Yes |
| ENSG00000125734.15 | GPR108 | rs2279623 | 6.10E-12 | ##### | Brain - Nucleus accumbens (b | 6732114 | 6232114 | 7232114 | Yes |
| ENSG00000125734.15 | GPR108 | rs12462822 | 6.10E-12 | ##### | Brain - Hippocampus | 6732114 | 6232114 | 7232114 | Yes |
| ENSG00000125734.15 | GPR108 | rs62125134 | 8.40E-12 | ##### | Brain - Cortex | 6732114 | 6232114 | 7232114 | Yes |
| ENSG00000125734.15 | GPR108 | rs2277986 | 8.70E-12 | ##### | Brain - Nucleus accumbens (b | 6732114 | 6232114 | 7232114 | Yes |
| ENSG00000125734.15 | GPR108 | rs340139 | 1.10E-11 | ##### | Brain - Cerebellum | 6732114 | 6232114 | 7232114 | Yes |
| ENSG00000125734.15 | GPR108 | rs2279623 | 1.10E-11 | ##### | Brain - Cerebellum | 6732114 | 6232114 | 7232114 | Yes |
| ENSG00000125734.15 | GPR108 | rs2279623 | 1.30E-11 | ##### | Brain - Putamen (basal gangli | 6732114 | 6232114 | 7232114 | Yes |
| ENSG00000125734.15 | GPR108 | rs2279623 | 1.60E-11 | ##### | Brain - Cerebellar Hemisphere | 6732114 | 6232114 | 7232114 | Yes |
| ENSG00000125734.15 | GPR108 | rs62125134 | 3.00E-11 | ##### | Brain - Cerebellum | 6732114 | 6232114 | 7232114 | Yes |
| ENSG00000125734.15 | GPR108 | rs339408 | 3.20E-11 | ##### | Brain - Cortex | 6732114 | 6232114 | 7232114 | Yes |
| ENSG00000125734.15 | GPR108 | rs339401 | 3.70E-11 | ##### | Brain - Cerebellum | 6732114 | 6232114 | 7232114 | Yes |
| ENSG00000125734.15 | GPR108 | rs2277986 | 5.90E-11 | ##### | Brain - Caudate (basal ganglia | 6732114 | 6232114 | 7232114 | Yes |
| ENSG00000125734.15 | GPR108 | rs62125134 | 6.40E-11 | ##### | Brain - Nucleus accumbens (b | 6732114 | 6232114 | 7232114 | Yes |

| | | | | | | | | | |
|---|---|---|---|---|---|---|---|---|---|
| ENSG00000125734.15 | GPR108 | rs2279623 | 7.00E-11 | ##### | Brain - Frontal Cortex (BA9) | 6732114 | 6232114 | 7232114 | Yes |
| ENSG00000125734.15 | GPR108 | rs2279623 | 1.00E-10 | ##### | Brain - Hippocampus | 6732114 | 6232114 | 7232114 | Yes |
| ENSG00000125734.15 | GPR108 | rs2277986 | 1.90E-10 | ##### | Brain - Cerebellar Hemisphere | 6732114 | 6232114 | 7232114 | Yes |
| ENSG00000125734.15 | GPR108 | rs2547438 | 2.20E-10 | ##### | Brain - Nucleus accumbens (b | 6732114 | 6232114 | 7232114 | Yes |
| ENSG00000125734.15 | GPR108 | rs62125134 | 2.70E-10 | ##### | Brain - Putamen (basal gangli | 6732114 | 6232114 | 7232114 | Yes |
| ENSG00000125734.15 | GPR108 | rs2277986 | 4.30E-10 | ##### | Brain - Hippocampus | 6732114 | 6232114 | 7232114 | Yes |
| ENSG00000125734.15 | GPR108 | rs12462822 | 4.40E-10 | ##### | Brain - Spinal cord (cervical c- | 6732114 | 6232114 | 7232114 | Yes |
| ENSG00000125734.15 | GPR108 | rs12462822 | 7.80E-10 | ##### | Brain - Amygdala | 6732114 | 6232114 | 7232114 | Yes |
| ENSG00000125734.15 | GPR108 | rs12462822 | 8.30E-10 | ##### | Brain - Hypothalamus | 6732114 | 6232114 | 7232114 | Yes |
| ENSG00000125734.15 | GPR108 | rs339395 | 1.20E-09 | ##### | Brain - Frontal Cortex (BA9) | 6732114 | 6232114 | 7232114 | Yes |
| ENSG00000125734.15 | GPR108 | rs62125134 | 2.00E-09 | ##### | Brain - Hippocampus | 6732114 | 6232114 | 7232114 | Yes |
| ENSG00000125734.15 | GPR108 | rs71309647 | 2.20E-09 | ##### | Brain - Cortex | 6732114 | 6232114 | 7232114 | Yes |
| ENSG00000125734.15 | GPR108 | rs12462822 | 2.30E-09 | ##### | Brain - Caudate (basal ganglia | 6732114 | 6232114 | 7232114 | Yes |
| ENSG00000125734.15 | GPR108 | rs10420527 | 2.30E-09 | ##### | Brain - Cerebellar Hemisphere | 6732114 | 6232114 | 7232114 | Yes |
| ENSG00000125734.15 | GPR108 | rs344614 | 2.30E-09 | ##### | Brain - Cerebellar Hemisphere | 6732114 | 6232114 | 7232114 | Yes |
| ENSG00000125734.15 | GPR108 | rs339397 | 3.70E-09 | ##### | Brain - Anterior cingulate cort | 6732114 | 6232114 | 7232114 | Yes |
| ENSG00000125734.15 | GPR108 | rs163916 | 3.70E-09 | ##### | Brain - Anterior cingulate cort | 6732114 | 6232114 | 7232114 | Yes |
| ENSG00000125734.15 | GPR108 | rs339399 | 3.70E-09 | ##### | Brain - Anterior cingulate cort | 6732114 | 6232114 | 7232114 | Yes |
| ENSG00000125734.15 | GPR108 | rs381897 | 5.90E-09 | ##### | Brain - Cerebellum | 6732114 | 6232114 | 7232114 | Yes |
| ENSG00000125734.15 | GPR108 | rs339395 | 6.60E-09 | ##### | Brain - Cortex | 6732114 | 6232114 | 7232114 | Yes |
| ENSG00000125734.15 | GPR108 | rs344616 | 7.00E-09 | ##### | Brain - Cerebellar Hemisphere | 6732114 | 6232114 | 7232114 | Yes |
| ENSG00000125734.15 | GPR108 | rs62125134 | 7.60E-09 | ##### | Brain - Hypothalamus | 6732114 | 6232114 | 7232114 | Yes |
| ENSG00000125734.15 | GPR108 | rs2547438 | 9.40E-09 | ##### | Brain - Frontal Cortex (BA9) | 6732114 | 6232114 | 7232114 | Yes |
| ENSG00000125734.15 | GPR108 | rs171094 | 9.60E-09 | ##### | Brain - Cerebellar Hemisphere | 6732114 | 6232114 | 7232114 | Yes |
| ENSG00000125734.15 | GPR108 | rs12462822 | 9.60E-09 | ##### | Brain - Anterior cingulate cort | 6732114 | 6232114 | 7232114 | Yes |
| ENSG00000125734.15 | GPR108 | rs12981348 | 9.80E-09 | ##### | Brain - Cerebellum | 6732114 | 6232114 | 7232114 | Yes |
| ENSG00000125734.15 | GPR108 | rs344616 | 1.00E-08 | ##### | Brain - Cerebellum | 6732114 | 6232114 | 7232114 | Yes |
| ENSG00000125734.15 | GPR108 | rs62125134 | 1.10E-08 | ##### | Brain - Anterior cingulate cort | 6732114 | 6232114 | 7232114 | Yes |
| ENSG00000125734.15 | GPR108 | rs2277986 | 1.10E-08 | ##### | Brain - Amygdala | 6732114 | 6232114 | 7232114 | Yes |
| ENSG00000125734.15 | GPR108 | rs10420527 | 1.20E-08 | ##### | Brain - Cerebellum | 6732114 | 6232114 | 7232114 | Yes |
| ENSG00000125734.15 | GPR108 | rs344614 | 1.20E-08 | ##### | Brain - Cerebellum | 6732114 | 6232114 | 7232114 | Yes |
| ENSG00000125734.15 | GPR108 | rs2547438 | 1.20E-08 | ##### | Brain - Cerebellum | 6732114 | 6232114 | 7232114 | Yes |

| | | | | | | | | | |
|---|---|---|---|---|---|---|---|---|---|
| ENSG00000125734.15 | GPR108 | rs2047140 | 1.40E-08 | ##### | Brain - Amygdala | 6732114 | 6232114 | 7232114 | Yes |
| ENSG00000125734.15 | GPR108 | rs2277986 | 1.60E-08 | ##### | Brain - Anterior cingulate cort | 6732114 | 6232114 | 7232114 | Yes |
| ENSG00000125734.15 | GPR108 | rs2047140 | 1.80E-08 | ##### | Brain - Anterior cingulate cort | 6732114 | 6232114 | 7232114 | Yes |
| ENSG00000125734.15 | GPR108 | rs2547438 | 1.90E-08 | ##### | Brain - Putamen (basal gangli | 6732114 | 6232114 | 7232114 | Yes |
| ENSG00000125734.15 | GPR108 | rs2291666 | 1.90E-08 | ##### | Brain - Cerebellum | 6732114 | 6232114 | 7232114 | Yes |
| ENSG00000125734.15 | GPR108 | rs2047140 | 2.00E-08 | ##### | Brain - Cerebellar Hemisphere | 6732114 | 6232114 | 7232114 | Yes |
| ENSG00000125734.15 | GPR108 | rs339400 | 2.10E-08 | ##### | Brain - Cerebellum | 6732114 | 6232114 | 7232114 | Yes |
| ENSG00000125734.15 | GPR108 | rs1690635 | 2.60E-08 | ##### | Brain - Cerebellum | 6732114 | 6232114 | 7232114 | Yes |
| ENSG00000125734.15 | GPR108 | rs2250656 | 2.70E-08 | ##### | Brain - Amygdala | 6732114 | 6232114 | 7232114 | Yes |
| ENSG00000125734.15 | GPR108 | rs171094 | 2.70E-08 | ##### | Brain - Cerebellum | 6732114 | 6232114 | 7232114 | Yes |
| ENSG00000125734.15 | GPR108 | rs2047140 | 2.90E-08 | ##### | Brain - Hippocampus | 6732114 | 6232114 | 7232114 | Yes |
| ENSG00000125734.15 | GPR108 | rs62125134 | 3.30E-08 | ##### | Brain - Amygdala | 6732114 | 6232114 | 7232114 | Yes |
| ENSG00000125734.15 | GPR108 | rs2547438 | 3.60E-08 | ##### | Brain - Cerebellar Hemisphere | 6732114 | 6232114 | 7232114 | Yes |
| ENSG00000125734.15 | GPR108 | rs62125134 | 5.30E-08 | ##### | Brain - Spinal cord (cervical c- | 6732114 | 6232114 | 7232114 | Yes |
| ENSG00000125734.15 | GPR108 | rs62125134 | 5.80E-08 | ##### | Brain - Caudate (basal ganglia | 6732114 | 6232114 | 7232114 | Yes |
| ENSG00000125734.15 | GPR108 | rs12981348 | 6.10E-08 | ##### | Brain - Cerebellar Hemisphere | 6732114 | 6232114 | 7232114 | Yes |
| ENSG00000125734.15 | GPR108 | rs2277986 | 7.00E-08 | ##### | Brain - Hypothalamus | 6732114 | 6232114 | 7232114 | Yes |
| ENSG00000125734.15 | GPR108 | rs235774 | 7.30E-08 | ##### | Brain - Cerebellum | 6732114 | 6232114 | 7232114 | Yes |
| ENSG00000125734.15 | GPR108 | rs2547438 | 7.60E-08 | ##### | Brain - Cortex | 6732114 | 6232114 | 7232114 | Yes |
| ENSG00000125734.15 | GPR108 | rs339397 | 7.70E-08 | ##### | Brain - Hypothalamus | 6732114 | 6232114 | 7232114 | Yes |
| ENSG00000125734.15 | GPR108 | rs163916 | 7.70E-08 | ##### | Brain - Hypothalamus | 6732114 | 6232114 | 7232114 | Yes |
| ENSG00000125734.15 | GPR108 | rs339399 | 7.70E-08 | ##### | Brain - Hypothalamus | 6732114 | 6232114 | 7232114 | Yes |
| ENSG00000125734.15 | GPR108 | rs2277986 | 8.80E-08 | ##### | Brain - Spinal cord (cervical c- | 6732114 | 6232114 | 7232114 | Yes |
| ENSG00000125734.15 | GPR108 | rs339397 | 1.10E-07 | ##### | Brain - Amygdala | 6732114 | 6232114 | 7232114 | Yes |
| ENSG00000125734.15 | GPR108 | rs381897 | 1.40E-07 | ##### | Brain - Nucleus accumbens (b | 6732114 | 6232114 | 7232114 | Yes |
| ENSG00000125734.15 | GPR108 | rs2279623 | 1.60E-07 | ##### | Brain - Caudate (basal ganglia | 6732114 | 6232114 | 7232114 | Yes |
| ENSG00000125734.15 | GPR108 | rs163916 | 1.60E-07 | ##### | Brain - Amygdala | 6732114 | 6232114 | 7232114 | Yes |
| ENSG00000125734.15 | GPR108 | rs339399 | 1.60E-07 | ##### | Brain - Amygdala | 6732114 | 6232114 | 7232114 | Yes |
| ENSG00000125734.15 | GPR108 | rs2279623 | 1.70E-07 | ##### | Brain - Hypothalamus | 6732114 | 6232114 | 7232114 | Yes |
| ENSG00000125734.15 | GPR108 | rs2547438 | 1.70E-07 | ##### | Brain - Anterior cingulate cort | 6732114 | 6232114 | 7232114 | Yes |
| ENSG00000125734.15 | GPR108 | rs339397 | 1.80E-07 | ##### | Brain - Cerebellar Hemisphere | 6732114 | 6232114 | 7232114 | Yes |
| ENSG00000125734.15 | GPR108 | rs2547438 | 1.90E-07 | ##### | Brain - Amygdala | 6732114 | 6232114 | 7232114 | Yes |

| | | | | | | | | | |
|---|---|---|---|---|---|---|---|---|---|
| ENSG00000125734.15 | GPR108 | rs381897 | 2.30E-07 | ##### | Brain - Cortex | 6732114 | 6232114 | 7232114 | Yes |
| ENSG00000125734.15 | GPR108 | rs339408 | 2.50E-07 | ##### | Brain - Frontal Cortex (BA9) | 6732114 | 6232114 | 7232114 | Yes |
| ENSG00000125734.15 | GPR108 | rs2250656 | 2.50E-07 | ##### | Brain - Nucleus accumbens (b | 6732114 | 6232114 | 7232114 | Yes |
| ENSG00000125734.15 | GPR108 | rs163916 | 2.60E-07 | ##### | Brain - Cerebellar Hemisphere | 6732114 | 6232114 | 7232114 | Yes |
| ENSG00000125734.15 | GPR108 | rs339399 | 2.60E-07 | ##### | Brain - Cerebellar Hemisphere | 6732114 | 6232114 | 7232114 | Yes |
| ENSG00000125734.15 | GPR108 | rs163915 | 2.60E-07 | ##### | Brain - Cerebellum | 6732114 | 6232114 | 7232114 | Yes |
| ENSG00000125734.15 | GPR108 | rs2279623 | 2.70E-07 | ##### | Brain - Spinal cord (cervical c- | 6732114 | 6232114 | 7232114 | Yes |
| ENSG00000125734.15 | GPR108 | rs10420527 | 2.90E-07 | ##### | Brain - Nucleus accumbens (b | 6732114 | 6232114 | 7232114 | Yes |
| ENSG00000125734.15 | GPR108 | rs344614 | 2.90E-07 | ##### | Brain - Nucleus accumbens (b | 6732114 | 6232114 | 7232114 | Yes |
| ENSG00000125734.15 | GPR108 | rs2250656 | 3.00E-07 | ##### | Brain - Cerebellar Hemisphere | 6732114 | 6232114 | 7232114 | Yes |
| ENSG00000125734.15 | GPR108 | rs340138 | 3.10E-07 | ##### | Brain - Cerebellar Hemisphere | 6732114 | 6232114 | 7232114 | Yes |
| ENSG00000125734.15 | GPR108 | rs339397 | 3.20E-07 | ##### | Brain - Hippocampus | 6732114 | 6232114 | 7232114 | Yes |
| ENSG00000125734.15 | GPR108 | rs163916 | 3.20E-07 | ##### | Brain - Hippocampus | 6732114 | 6232114 | 7232114 | Yes |
| ENSG00000125734.15 | GPR108 | rs339399 | 3.20E-07 | ##### | Brain - Hippocampus | 6732114 | 6232114 | 7232114 | Yes |
| ENSG00000125734.15 | GPR108 | rs340139 | 3.20E-07 | ##### | Brain - Cerebellar Hemisphere | 6732114 | 6232114 | 7232114 | Yes |
| ENSG00000125734.15 | GPR108 | rs2047140 | 3.30E-07 | ##### | Brain - Hypothalamus | 6732114 | 6232114 | 7232114 | Yes |
| ENSG00000125734.15 | GPR108 | rs2279623 | 3.70E-07 | ##### | Brain - Amygdala | 6732114 | 6232114 | 7232114 | Yes |
| ENSG00000125734.15 | GPR108 | rs344616 | 3.80E-07 | ##### | Brain - Nucleus accumbens (b | 6732114 | 6232114 | 7232114 | Yes |
| ENSG00000125734.15 | GPR108 | rs12462822 | 4.20E-07 | ##### | Brain - Substantia nigra | 6732114 | 6232114 | 7232114 | Yes |
| ENSG00000125734.15 | GPR108 | rs2291666 | 4.30E-07 | ##### | Brain - Cerebellar Hemisphere | 6732114 | 6232114 | 7232114 | Yes |
| ENSG00000125734.15 | GPR108 | rs75247649 | 4.70E-07 | ##### | Brain - Frontal Cortex (BA9) | 6732114 | 6232114 | 7232114 | Yes |
| ENSG00000125734.15 | GPR108 | rs2547438 | 4.90E-07 | ##### | Brain - Spinal cord (cervical c- | 6732114 | 6232114 | 7232114 | Yes |
| ENSG00000125734.15 | GPR108 | rs2250656 | 5.40E-07 | ##### | Brain - Putamen (basal gangli | 6732114 | 6232114 | 7232114 | Yes |
| ENSG00000125734.15 | GPR108 | rs171094 | 5.50E-07 | ##### | Brain - Nucleus accumbens (b | 6732114 | 6232114 | 7232114 | Yes |
| ENSG00000125734.15 | GPR108 | rs71309647 | 6.30E-07 | ##### | Brain - Frontal Cortex (BA9) | 6732114 | 6232114 | 7232114 | Yes |
| ENSG00000125734.15 | GPR108 | rs1702376 | 6.50E-07 | ##### | Brain - Cortex | 6732114 | 6232114 | 7232114 | Yes |
| ENSG00000125734.15 | GPR108 | rs2250656 | 7.40E-07 | ##### | Brain - Cerebellum | 6732114 | 6232114 | 7232114 | Yes |
| ENSG00000125734.15 | GPR108 | rs2277986 | 8.60E-07 | ##### | Brain - Substantia nigra | 6732114 | 6232114 | 7232114 | Yes |
| ENSG00000125734.15 | GPR108 | rs11569575 | 8.90E-07 | ##### | Brain - Amygdala | 6732114 | 6232114 | 7232114 | Yes |
| ENSG00000125734.15 | GPR108 | rs339408 | 9.80E-07 | ##### | Brain - Amygdala | 6732114 | 6232114 | 7232114 | Yes |
| ENSG00000125734.15 | GPR108 | rs2291666 | 0.0000011 | ##### | Brain - Nucleus accumbens (b | 6732114 | 6232114 | 7232114 | Yes |
| ENSG00000125734.15 | GPR108 | rs2250656 | 0.0000011 | ##### | Brain - Frontal Cortex (BA9) | 6732114 | 6232114 | 7232114 | Yes |

| | | | | | | | | | |
|---|---|---|---|---|---|---|---|---|---|
| ENSG00000125734.15 | GPR108 | rs2250656 | 0.0000012 | ##### | Brain - Anterior cingulate cort | 6732114 | 6232114 | 7232114 | Yes |
| ENSG00000125734.15 | GPR108 | rs339401 | 0.0000012 | ##### | Brain - Cerebellar Hemisphere | 6732114 | 6232114 | 7232114 | Yes |
| ENSG00000125734.15 | GPR108 | rs71309647 | 0.0000013 | ##### | Brain - Cerebellum | 6732114 | 6232114 | 7232114 | Yes |
| ENSG00000125734.15 | GPR108 | rs3745556 | 0.0000013 | ##### | Brain - Cortex | 6732114 | 6232114 | 7232114 | Yes |
| ENSG00000125734.15 | GPR108 | rs235774 | 0.0000014 | ##### | Brain - Cerebellar Hemisphere | 6732114 | 6232114 | 7232114 | Yes |
| ENSG00000125734.15 | GPR108 | rs339408 | 0.0000015 | ##### | Brain - Hippocampus | 6732114 | 6232114 | 7232114 | Yes |
| ENSG00000125734.15 | GPR108 | rs163916 | 0.0000018 | ##### | Brain - Substantia nigra | 6732114 | 6232114 | 7232114 | Yes |
| ENSG00000125734.15 | GPR108 | rs339399 | 0.0000018 | ##### | Brain - Substantia nigra | 6732114 | 6232114 | 7232114 | Yes |
| ENSG00000125734.15 | GPR108 | rs2047140 | 0.0000019 | ##### | Brain - Putamen (basal gangli | 6732114 | 6232114 | 7232114 | Yes |
| ENSG00000125734.15 | GPR108 | rs381897 | 0.0000019 | ##### | Brain - Cerebellar Hemisphere | 6732114 | 6232114 | 7232114 | Yes |
| ENSG00000125734.15 | GPR108 | rs339397 | 0.000002 | ##### | Brain - Cerebellum | 6732114 | 6232114 | 7232114 | Yes |
| ENSG00000125734.15 | GPR108 | rs339397 | 0.0000021 | ##### | Brain - Substantia nigra | 6732114 | 6232114 | 7232114 | Yes |
| ENSG00000125734.15 | GPR108 | rs381897 | 0.0000021 | ##### | Brain - Putamen (basal gangli | 6732114 | 6232114 | 7232114 | Yes |
| ENSG00000125734.15 | GPR108 | rs163916 | 0.0000023 | ##### | Brain - Cerebellum | 6732114 | 6232114 | 7232114 | Yes |
| ENSG00000125734.15 | GPR108 | rs339399 | 0.0000023 | ##### | Brain - Cerebellum | 6732114 | 6232114 | 7232114 | Yes |
| ENSG00000125734.15 | GPR108 | rs2047140 | 0.0000027 | ##### | Brain - Substantia nigra | 6732114 | 6232114 | 7232114 | Yes |
| ENSG00000125734.15 | GPR108 | rs2250656 | 0.0000027 | ##### | Brain - Cortex | 6732114 | 6232114 | 7232114 | Yes |
| ENSG00000125734.15 | GPR108 | rs339405 | 0.0000034 | ##### | Brain - Cerebellum | 6732114 | 6232114 | 7232114 | Yes |
| ENSG00000125734.15 | GPR108 | rs235774 | 0.0000034 | ##### | Brain - Amygdala | 6732114 | 6232114 | 7232114 | Yes |
| ENSG00000125734.15 | GPR108 | rs12981348 | 0.000004 | ##### | Brain - Frontal Cortex (BA9) | 6732114 | 6232114 | 7232114 | Yes |
| ENSG00000125734.15 | GPR108 | rs1702376 | 0.0000041 | ##### | Brain - Cerebellum | 6732114 | 6232114 | 7232114 | Yes |
| ENSG00000125734.15 | GPR108 | rs12981348 | 0.0000041 | ##### | Brain - Nucleus accumbens (b | 6732114 | 6232114 | 7232114 | Yes |
| ENSG00000125734.15 | GPR108 | rs344616 | 0.0000044 | ##### | Brain - Caudate (basal ganglia | 6732114 | 6232114 | 7232114 | Yes |
| ENSG00000125734.15 | GPR108 | rs344616 | 0.0000049 | ##### | Brain - Putamen (basal gangli | 6732114 | 6232114 | 7232114 | Yes |
| ENSG00000125734.15 | GPR108 | rs62125134 | 0.0000051 | ##### | Brain - Substantia nigra | 6732114 | 6232114 | 7232114 | Yes |
| ENSG00000125734.15 | GPR108 | rs2047140 | 0.0000051 | ##### | Brain - Caudate (basal ganglia | 6732114 | 6232114 | 7232114 | Yes |
| ENSG00000125734.15 | GPR108 | rs141729894 | 0.0000051 | ##### | Brain - Anterior cingulate cort | 6732114 | 6232114 | 7232114 | No |
| ENSG00000125734.15 | GPR108 | rs2047140 | 0.0000052 | ##### | Brain - Nucleus accumbens (b | 6732114 | 6232114 | 7232114 | Yes |
| ENSG00000125734.15 | GPR108 | rs1702376 | 0.0000053 | ##### | Brain - Frontal Cortex (BA9) | 6732114 | 6232114 | 7232114 | Yes |
| ENSG00000125734.15 | GPR108 | rs3745556 | 0.0000053 | ##### | Brain - Nucleus accumbens (b | 6732114 | 6232114 | 7232114 | Yes |
| ENSG00000125734.15 | GPR108 | rs339408 | 0.0000053 | ##### | Brain - Anterior cingulate cort | 6732114 | 6232114 | 7232114 | Yes |
| ENSG00000125734.15 | GPR108 | rs339397 | 0.0000056 | ##### | Brain - Caudate (basal ganglia | 6732114 | 6232114 | 7232114 | Yes |

| | | | | | | | | | |
|---|---|---|---|---|---|---|---|---|---|
| ENSG00000125734.15 | GPR108 | rs2279623 | 0.0000057 | ##### | Brain - Anterior cingulate cort | 6732114 | 6232114 | 7232114 | Yes |
| ENSG00000125734.15 | GPR108 | rs339395 | 0.0000066 | ##### | Brain - Cerebellar Hemisphere | 6732114 | 6232114 | 7232114 | Yes |
| ENSG00000125734.15 | GPR108 | rs75247649 | 0.0000067 | ##### | Brain - Cerebellum | 6732114 | 6232114 | 7232114 | Yes |
| ENSG00000125734.15 | GPR108 | rs235774 | 0.0000068 | ##### | Brain - Nucleus accumbens (b | 6732114 | 6232114 | 7232114 | Yes |
| ENSG00000125734.15 | GPR108 | rs339404 | 0.0000075 | ##### | Brain - Cerebellum | 6732114 | 6232114 | 7232114 | Yes |
| ENSG00000125734.15 | GPR108 | rs10420527 | 0.0000079 | ##### | Brain - Putamen (basal gangli | 6732114 | 6232114 | 7232114 | Yes |
| ENSG00000125734.15 | GPR108 | rs344614 | 0.0000079 | ##### | Brain - Putamen (basal gangli | 6732114 | 6232114 | 7232114 | Yes |
| ENSG00000125734.15 | GPR108 | rs163915 | 0.0000086 | ##### | Brain - Cerebellar Hemisphere | 6732114 | 6232114 | 7232114 | Yes |
| ENSG00000125734.15 | GPR108 | rs171094 | 0.0000097 | ##### | Brain - Hypothalamus | 6732114 | 6232114 | 7232114 | Yes |
| ENSG00000125734.15 | GPR108 | rs10420527 | 0.0000097 | ##### | Brain - Hypothalamus | 6732114 | 6232114 | 7232114 | Yes |
| ENSG00000125734.15 | GPR108 | rs344614 | 0.0000097 | ##### | Brain - Hypothalamus | 6732114 | 6232114 | 7232114 | Yes |
| ENSG00000125734.15 | GPR108 | rs339395 | 0.00001 | ##### | Brain - Anterior cingulate cort | 6732114 | 6232114 | 7232114 | Yes |
| ENSG00000125734.15 | GPR108 | rs171094 | 0.00001 | ##### | Brain - Putamen (basal gangli | 6732114 | 6232114 | 7232114 | Yes |
| ENSG00000125734.15 | GPR108 | rs235774 | 0.000011 | ##### | Brain - Hypothalamus | 6732114 | 6232114 | 7232114 | Yes |
| ENSG00000125734.15 | GPR108 | rs339396 | 0.000012 | ##### | Brain - Cerebellum | 6732114 | 6232114 | 7232114 | Yes |
| ENSG00000125734.15 | GPR108 | rs10420527 | 0.000013 | ##### | Brain - Caudate (basal ganglia | 6732114 | 6232114 | 7232114 | Yes |
| ENSG00000125734.15 | GPR108 | rs344614 | 0.000013 | ##### | Brain - Caudate (basal ganglia | 6732114 | 6232114 | 7232114 | Yes |
| ENSG00000125734.15 | GPR108 | rs2250656 | 0.000013 | ##### | Brain - Spinal cord (cervical c- | 6732114 | 6232114 | 7232114 | Yes |
| ENSG00000125734.15 | GPR108 | rs339408 | 0.000014 | ##### | Brain - Hypothalamus | 6732114 | 6232114 | 7232114 | Yes |
| ENSG00000125734.15 | GPR108 | rs163916 | 0.000015 | ##### | Brain - Caudate (basal ganglia | 6732114 | 6232114 | 7232114 | Yes |
| ENSG00000125734.15 | GPR108 | rs339399 | 0.000015 | ##### | Brain - Caudate (basal ganglia | 6732114 | 6232114 | 7232114 | Yes |
| ENSG00000125734.15 | GPR108 | rs2547438 | 0.000016 | ##### | Brain - Hypothalamus | 6732114 | 6232114 | 7232114 | Yes |
| ENSG00000125734.15 | GPR108 | rs339396 | 0.000017 | ##### | Brain - Cortex | 6732114 | 6232114 | 7232114 | Yes |
| ENSG00000125734.15 | GPR108 | rs163915 | 0.000017 | ##### | Brain - Nucleus accumbens (b | 6732114 | 6232114 | 7232114 | Yes |
| ENSG00000125734.15 | GPR108 | rs2547438 | 0.000018 | ##### | Brain - Hippocampus | 6732114 | 6232114 | 7232114 | Yes |
| ENSG00000125734.15 | GPR108 | rs57086167 | 0.000018 | ##### | Brain - Cortex | 6732114 | 6232114 | 7232114 | Yes |
| ENSG00000125734.15 | GPR108 | rs1690635 | 0.000019 | ##### | Brain - Cerebellar Hemisphere | 6732114 | 6232114 | 7232114 | Yes |
| ENSG00000125734.15 | GPR108 | rs339408 | 0.000021 | ##### | Brain - Putamen (basal gangli | 6732114 | 6232114 | 7232114 | Yes |
| ENSG00000125734.15 | GPR108 | rs340138 | 0.000022 | ##### | Brain - Caudate (basal ganglia | 6732114 | 6232114 | 7232114 | Yes |
| ENSG00000125734.15 | GPR108 | rs340139 | 0.000022 | ##### | Brain - Caudate (basal ganglia | 6732114 | 6232114 | 7232114 | Yes |
| ENSG00000125734.15 | GPR108 | rs339397 | 0.000024 | ##### | Brain - Putamen (basal gangli | 6732114 | 6232114 | 7232114 | Yes |
| ENSG00000125734.15 | GPR108 | rs235774 | 0.000025 | ##### | Brain - Putamen (basal gangli | 6732114 | 6232114 | 7232114 | Yes |

| | | | | | | | | | |
|---|---|---|---|---|---|---|---|---|---|
| ENSG00000125734.15 | GPR108 | rs163494 | 0.000027 | ##### | Brain - Frontal Cortex (BA9) | 6732114 | 6232114 | 7232114 | Yes |
| ENSG00000125734.15 | GPR108 | rs3745556 | 0.000028 | ##### | Brain - Caudate (basal ganglia | 6732114 | 6232114 | 7232114 | Yes |
| ENSG00000125734.15 | GPR108 | rs339408 | 0.000033 | ##### | Brain - Nucleus accumbens (b | 6732114 | 6232114 | 7232114 | Yes |
| ENSG00000125734.15 | GPR108 | rs2047140 | 0.000042 | ##### | Brain - Cerebellum | 6732114 | 6232114 | 7232114 | Yes |
| ENSG00000120647.9 | CCDC77 | rs11062813 | 1.10E-19 | ##### | Brain - Hypothalamus | 440788 | -59212 | 940788 | Yes |
| ENSG00000120647.9 | CCDC77 | rs2369280 | 1.80E-18 | ##### | Brain - Cerebellar Hemisphere | 440788 | -59212 | 940788 | Yes |
| ENSG00000120647.9 | CCDC77 | rs372437080 | 2.80E-17 | ##### | Brain - Hypothalamus | 440788 | -59212 | 940788 | Yes |
| ENSG00000120647.9 | CCDC77 | rs4980892 | 8.70E-17 | ##### | Brain - Cerebellar Hemisphere | 440788 | -59212 | 940788 | Yes |
| ENSG00000120647.9 | CCDC77 | rs111250603 | 8.80E-17 | ##### | Brain - Cerebellar Hemisphere | 440788 | -59212 | 940788 | Yes |
| ENSG00000120647.9 | CCDC77 | rs2369277 | 8.80E-17 | ##### | Brain - Cerebellar Hemisphere | 440788 | -59212 | 940788 | Yes |
| ENSG00000120647.9 | CCDC77 | rs9805067 | 8.80E-17 | ##### | Brain - Cerebellar Hemisphere | 440788 | -59212 | 940788 | Yes |
| ENSG00000120647.9 | CCDC77 | rs1541464 | 8.80E-17 | ##### | Brain - Cerebellar Hemisphere | 440788 | -59212 | 940788 | Yes |
| ENSG00000120647.9 | CCDC77 | rs1541465 | 8.80E-17 | ##### | Brain - Cerebellar Hemisphere | 440788 | -59212 | 940788 | Yes |
| ENSG00000120647.9 | CCDC77 | rs4980817 | 8.80E-17 | ##### | Brain - Cerebellar Hemisphere | 440788 | -59212 | 940788 | Yes |
| ENSG00000120647.9 | CCDC77 | rs4482086 | 8.80E-17 | ##### | Brain - Cerebellar Hemisphere | 440788 | -59212 | 940788 | Yes |
| ENSG00000120647.9 | CCDC77 | rs2369279 | 8.80E-17 | ##### | Brain - Cerebellar Hemisphere | 440788 | -59212 | 940788 | Yes |
| ENSG00000120647.9 | CCDC77 | rs11062803 | 8.80E-17 | ##### | Brain - Cerebellar Hemisphere | 440788 | -59212 | 940788 | Yes |
| ENSG00000120647.9 | CCDC77 | rs372437080 | 1.10E-16 | ##### | Brain - Cerebellar Hemisphere | 440788 | -59212 | 940788 | Yes |
| ENSG00000120647.9 | CCDC77 | rs1541463 | 1.20E-16 | ##### | Brain - Cerebellar Hemisphere | 440788 | -59212 | 940788 | Yes |
| ENSG00000120647.9 | CCDC77 | rs2369278 | 1.20E-16 | ##### | Brain - Cerebellar Hemisphere | 440788 | -59212 | 940788 | Yes |
| ENSG00000120647.9 | CCDC77 | rs10774163 | 1.20E-16 | ##### | Brain - Cerebellar Hemisphere | 440788 | -59212 | 940788 | Yes |
| ENSG00000120647.9 | CCDC77 | rs10848905 | 1.70E-16 | ##### | Brain - Cerebellar Hemisphere | 440788 | -59212 | 940788 | Yes |
| ENSG00000120647.9 | CCDC77 | rs10744627 | 2.20E-16 | ##### | Brain - Cerebellar Hemisphere | 440788 | -59212 | 940788 | Yes |
| ENSG00000120647.9 | CCDC77 | rs7133796 | 3.60E-16 | ##### | Brain - Cerebellar Hemisphere | 440788 | -59212 | 940788 | Yes |
| ENSG00000120647.9 | CCDC77 | rs7138958 | 6.20E-16 | ##### | Brain - Cerebellar Hemisphere | 440788 | -59212 | 940788 | Yes |
| ENSG00000120647.9 | CCDC77 | rs2007537 | 6.20E-16 | ##### | Brain - Cerebellar Hemisphere | 440788 | -59212 | 940788 | Yes |
| ENSG00000120647.9 | CCDC77 | rs372437080 | 8.50E-16 | ##### | Brain - Caudate (basal ganglia | 440788 | -59212 | 940788 | Yes |
| ENSG00000120647.9 | CCDC77 | rs75894333 | 9.80E-16 | ##### | Brain - Cerebellar Hemisphere | 440788 | -59212 | 940788 | Yes |
| ENSG00000120647.9 | CCDC77 | rs507597 | 9.80E-16 | ##### | Brain - Cerebellar Hemisphere | 440788 | -59212 | 940788 | Yes |
| ENSG00000120647.9 | CCDC77 | rs17800744 | 9.80E-16 | ##### | Brain - Cerebellar Hemisphere | 440788 | -59212 | 940788 | Yes |
| ENSG00000120647.9 | CCDC77 | rs77854644 | 9.80E-16 | ##### | Brain - Cerebellar Hemisphere | 440788 | -59212 | 940788 | Yes |
| ENSG00000120647.9 | CCDC77 | rs11608938 | 9.80E-16 | ##### | Brain - Cerebellar Hemisphere | 440788 | -59212 | 940788 | Yes |

| | | | | | | | | | |
|---|---|---|---|---|---|---|---|---|---|
| ENSG00000120647.9 | CCDC77 | rs7965038 | 9.80E-16 | ##### | Brain - Cerebellar Hemisphere | 440788 | -59212 | 940788 | Yes |
| ENSG00000120647.9 | CCDC77 | rs7307606 | 9.80E-16 | ##### | Brain - Cerebellar Hemisphere | 440788 | -59212 | 940788 | Yes |
| ENSG00000120647.9 | CCDC77 | rs10661251 | 9.80E-16 | ##### | Brain - Cerebellar Hemisphere | 440788 | -59212 | 940788 | Yes |
| ENSG00000120647.9 | CCDC77 | rs1011699 | 9.80E-16 | ##### | Brain - Cerebellar Hemisphere | 440788 | -59212 | 940788 | Yes |
| ENSG00000120647.9 | CCDC77 | rs6489475 | 9.80E-16 | ##### | Brain - Cerebellar Hemisphere | 440788 | -59212 | 940788 | Yes |
| ENSG00000120647.9 | CCDC77 | rs10774154 | 9.80E-16 | ##### | Brain - Cerebellar Hemisphere | 440788 | -59212 | 940788 | Yes |
| ENSG00000120647.9 | CCDC77 | rs11062813 | 1.50E-15 | ##### | Brain - Cerebellar Hemisphere | 440788 | -59212 | 940788 | Yes |
| ENSG00000120647.9 | CCDC77 | rs372437080 | 2.70E-15 | ##### | Brain - Frontal Cortex (BA9) | 440788 | -59212 | 940788 | Yes |
| ENSG00000120647.9 | CCDC77 | rs1155942 | 3.90E-15 | ##### | Brain - Hypothalamus | 440788 | -59212 | 940788 | Yes |
| ENSG00000120647.9 | CCDC77 | rs7972566 | 4.40E-15 | ##### | Brain - Hypothalamus | 440788 | -59212 | 940788 | Yes |
| ENSG00000120647.9 | CCDC77 | rs10774170 | 4.40E-15 | ##### | Brain - Hypothalamus | 440788 | -59212 | 940788 | Yes |
| ENSG00000120647.9 | CCDC77 | rs2041299 | 4.70E-15 | ##### | Brain - Cerebellar Hemisphere | 440788 | -59212 | 940788 | Yes |
| ENSG00000120647.9 | CCDC77 | rs7972566 | 6.70E-15 | ##### | Brain - Cerebellar Hemisphere | 440788 | -59212 | 940788 | Yes |
| ENSG00000120647.9 | CCDC77 | rs10774170 | 6.70E-15 | ##### | Brain - Cerebellar Hemisphere | 440788 | -59212 | 940788 | Yes |
| ENSG00000120647.9 | CCDC77 | rs79839498 | 9.20E-15 | ##### | Brain - Cerebellar Hemisphere | 440788 | -59212 | 940788 | Yes |
| ENSG00000120647.9 | CCDC77 | rs525381 | 1.00E-14 | ##### | Brain - Cerebellar Hemisphere | 440788 | -59212 | 940788 | Yes |
| ENSG00000120647.9 | CCDC77 | rs7133796 | 1.10E-14 | ##### | Brain - Hypothalamus | 440788 | -59212 | 940788 | Yes |
| ENSG00000120647.9 | CCDC77 | rs495973 | 1.10E-14 | ##### | Brain - Caudate (basal ganglia | 440788 | -59212 | 940788 | Yes |
| ENSG00000120647.9 | CCDC77 | rs75894333 | 1.10E-14 | ##### | Brain - Hypothalamus | 440788 | -59212 | 940788 | Yes |
| ENSG00000120647.9 | CCDC77 | rs11608938 | 1.10E-14 | ##### | Brain - Hypothalamus | 440788 | -59212 | 940788 | Yes |
| ENSG00000120647.9 | CCDC77 | rs7965038 | 1.10E-14 | ##### | Brain - Hypothalamus | 440788 | -59212 | 940788 | Yes |
| ENSG00000120647.9 | CCDC77 | rs7307606 | 1.10E-14 | ##### | Brain - Hypothalamus | 440788 | -59212 | 940788 | Yes |
| ENSG00000120647.9 | CCDC77 | rs10661251 | 1.10E-14 | ##### | Brain - Hypothalamus | 440788 | -59212 | 940788 | Yes |
| ENSG00000120647.9 | CCDC77 | rs1011699 | 1.10E-14 | ##### | Brain - Hypothalamus | 440788 | -59212 | 940788 | Yes |
| ENSG00000120647.9 | CCDC77 | rs6489475 | 1.10E-14 | ##### | Brain - Hypothalamus | 440788 | -59212 | 940788 | Yes |
| ENSG00000120647.9 | CCDC77 | rs10774154 | 1.10E-14 | ##### | Brain - Hypothalamus | 440788 | -59212 | 940788 | Yes |
| ENSG00000120647.9 | CCDC77 | rs1207322 | 1.10E-14 | ##### | Brain - Cerebellar Hemisphere | 440788 | -59212 | 940788 | Yes |
| ENSG00000120647.9 | CCDC77 | rs4980813 | 1.10E-14 | ##### | Brain - Cerebellar Hemisphere | 440788 | -59212 | 940788 | Yes |
| ENSG00000120647.9 | CCDC77 | rs61906996 | 1.10E-14 | ##### | Brain - Cerebellar Hemisphere | 440788 | -59212 | 940788 | Yes |
| ENSG00000120647.9 | CCDC77 | rs4980814 | 1.10E-14 | ##### | Brain - Cerebellar Hemisphere | 440788 | -59212 | 940788 | Yes |
| ENSG00000120647.9 | CCDC77 | rs7977462 | 1.10E-14 | ##### | Brain - Cerebellar Hemisphere | 440788 | -59212 | 940788 | Yes |
| ENSG00000120647.9 | CCDC77 | rs10744616 | 1.10E-14 | ##### | Brain - Cerebellar Hemisphere | 440788 | -59212 | 940788 | Yes |

| | | | | | | | | | |
|---|---|---|---|---|---|---|---|---|---|
| ENSG00000120647.9 | CCDC77 | rs10774150 | 1.10E-14 | ##### | Brain - Cerebellar Hemisphere | 440788 | -59212 | 940788 | Yes |
| ENSG00000120647.9 | CCDC77 | rs3831759 | 1.10E-14 | ##### | Brain - Cerebellar Hemisphere | 440788 | -59212 | 940788 | Yes |
| ENSG00000120647.9 | CCDC77 | rs10744627 | 1.20E-14 | ##### | Brain - Hypothalamus | 440788 | -59212 | 940788 | Yes |
| ENSG00000120647.9 | CCDC77 | rs520932 | 2.10E-14 | ##### | Brain - Caudate (basal ganglia | 440788 | -59212 | 940788 | Yes |
| ENSG00000120647.9 | CCDC77 | rs495973 | 2.30E-14 | ##### | Brain - Cerebellar Hemisphere | 440788 | -59212 | 940788 | Yes |
| ENSG00000120647.9 | CCDC77 | rs2041299 | 2.30E-14 | ##### | Brain - Hypothalamus | 440788 | -59212 | 940788 | Yes |
| ENSG00000120647.9 | CCDC77 | rs75894333 | 2.40E-14 | ##### | Brain - Caudate (basal ganglia | 440788 | -59212 | 940788 | Yes |
| ENSG00000120647.9 | CCDC77 | rs11608938 | 2.40E-14 | ##### | Brain - Caudate (basal ganglia | 440788 | -59212 | 940788 | Yes |
| ENSG00000120647.9 | CCDC77 | rs7965038 | 2.40E-14 | ##### | Brain - Caudate (basal ganglia | 440788 | -59212 | 940788 | Yes |
| ENSG00000120647.9 | CCDC77 | rs7307606 | 2.40E-14 | ##### | Brain - Caudate (basal ganglia | 440788 | -59212 | 940788 | Yes |
| ENSG00000120647.9 | CCDC77 | rs10661251 | 2.40E-14 | ##### | Brain - Caudate (basal ganglia | 440788 | -59212 | 940788 | Yes |
| ENSG00000120647.9 | CCDC77 | rs1011699 | 2.40E-14 | ##### | Brain - Caudate (basal ganglia | 440788 | -59212 | 940788 | Yes |
| ENSG00000120647.9 | CCDC77 | rs6489475 | 2.40E-14 | ##### | Brain - Caudate (basal ganglia | 440788 | -59212 | 940788 | Yes |
| ENSG00000120647.9 | CCDC77 | rs10774154 | 2.40E-14 | ##### | Brain - Caudate (basal ganglia | 440788 | -59212 | 940788 | Yes |
| ENSG00000120647.9 | CCDC77 | rs10774169 | 2.60E-14 | ##### | Brain - Hypothalamus | 440788 | -59212 | 940788 | Yes |
| ENSG00000120647.9 | CCDC77 | rs507597 | 2.70E-14 | ##### | Brain - Caudate (basal ganglia | 440788 | -59212 | 940788 | Yes |
| ENSG00000120647.9 | CCDC77 | rs17800744 | 2.70E-14 | ##### | Brain - Caudate (basal ganglia | 440788 | -59212 | 940788 | Yes |
| ENSG00000120647.9 | CCDC77 | rs77854644 | 2.70E-14 | ##### | Brain - Caudate (basal ganglia | 440788 | -59212 | 940788 | Yes |
| ENSG00000120647.9 | CCDC77 | rs1207324 | 3.10E-14 | ##### | Brain - Cerebellar Hemisphere | 440788 | -59212 | 940788 | Yes |
| ENSG00000120647.9 | CCDC77 | rs61907026 | 3.30E-14 | ##### | Brain - Cerebellar Hemisphere | 440788 | -59212 | 940788 | Yes |
| ENSG00000120647.9 | CCDC77 | rs61907030 | 3.30E-14 | ##### | Brain - Cerebellar Hemisphere | 440788 | -59212 | 940788 | Yes |
| ENSG00000120647.9 | CCDC77 | rs61907031 | 3.30E-14 | ##### | Brain - Cerebellar Hemisphere | 440788 | -59212 | 940788 | Yes |
| ENSG00000120647.9 | CCDC77 | rs11615564 | 3.30E-14 | ##### | Brain - Cerebellar Hemisphere | 440788 | -59212 | 940788 | Yes |
| ENSG00000120647.9 | CCDC77 | rs7138958 | 3.80E-14 | ##### | Brain - Hypothalamus | 440788 | -59212 | 940788 | Yes |
| ENSG00000120647.9 | CCDC77 | rs2007537 | 3.80E-14 | ##### | Brain - Hypothalamus | 440788 | -59212 | 940788 | Yes |
| ENSG00000120647.9 | CCDC77 | rs4980876 | 4.00E-14 | ##### | Brain - Cerebellar Hemisphere | 440788 | -59212 | 940788 | Yes |
| ENSG00000120647.9 | CCDC77 | rs111250603 | 4.20E-14 | ##### | Brain - Hypothalamus | 440788 | -59212 | 940788 | Yes |
| ENSG00000120647.9 | CCDC77 | rs2369277 | 4.20E-14 | ##### | Brain - Hypothalamus | 440788 | -59212 | 940788 | Yes |
| ENSG00000120647.9 | CCDC77 | rs9805067 | 4.20E-14 | ##### | Brain - Hypothalamus | 440788 | -59212 | 940788 | Yes |
| ENSG00000120647.9 | CCDC77 | rs1541464 | 4.20E-14 | ##### | Brain - Hypothalamus | 440788 | -59212 | 940788 | Yes |
| ENSG00000120647.9 | CCDC77 | rs1541465 | 4.20E-14 | ##### | Brain - Hypothalamus | 440788 | -59212 | 940788 | Yes |
| ENSG00000120647.9 | CCDC77 | rs4980817 | 4.20E-14 | ##### | Brain - Hypothalamus | 440788 | -59212 | 940788 | Yes |

| | | | | | | | | | |
|---|---|---|---|---|---|---|---|---|---|
| ENSG00000120647.9 | CCDC77 | rs4482086 | 4.20E-14 | ##### | Brain - Hypothalamus | 440788 | -59212 | 940788 | Yes |
| ENSG00000120647.9 | CCDC77 | rs2369279 | 4.20E-14 | ##### | Brain - Hypothalamus | 440788 | -59212 | 940788 | Yes |
| ENSG00000120647.9 | CCDC77 | rs11062803 | 4.20E-14 | ##### | Brain - Hypothalamus | 440788 | -59212 | 940788 | Yes |
| ENSG00000120647.9 | CCDC77 | rs4980896 | 4.20E-14 | ##### | Brain - Caudate (basal ganglia | 440788 | -59212 | 940788 | Yes |
| ENSG00000120647.9 | CCDC77 | rs7133796 | 4.80E-14 | ##### | Brain - Caudate (basal ganglia | 440788 | -59212 | 940788 | Yes |
| ENSG00000120647.9 | CCDC77 | rs10848905 | 5.40E-14 | ##### | Brain - Hypothalamus | 440788 | -59212 | 940788 | Yes |
| ENSG00000120647.9 | CCDC77 | rs3213751 | 6.70E-14 | ##### | Brain - Cerebellar Hemisphere | 440788 | -59212 | 940788 | Yes |
| ENSG00000120647.9 | CCDC77 | rs4980878 | 7.00E-14 | ##### | Brain - Cerebellar Hemisphere | 440788 | -59212 | 940788 | Yes |
| ENSG00000120647.9 | CCDC77 | rs75894333 | 7.10E-14 | ##### | Brain - Frontal Cortex (BA9) | 440788 | -59212 | 940788 | Yes |
| ENSG00000120647.9 | CCDC77 | rs11608938 | 7.10E-14 | ##### | Brain - Frontal Cortex (BA9) | 440788 | -59212 | 940788 | Yes |
| ENSG00000120647.9 | CCDC77 | rs7965038 | 7.10E-14 | ##### | Brain - Frontal Cortex (BA9) | 440788 | -59212 | 940788 | Yes |
| ENSG00000120647.9 | CCDC77 | rs7307606 | 7.10E-14 | ##### | Brain - Frontal Cortex (BA9) | 440788 | -59212 | 940788 | Yes |
| ENSG00000120647.9 | CCDC77 | rs10661251 | 7.10E-14 | ##### | Brain - Frontal Cortex (BA9) | 440788 | -59212 | 940788 | Yes |
| ENSG00000120647.9 | CCDC77 | rs1011699 | 7.10E-14 | ##### | Brain - Frontal Cortex (BA9) | 440788 | -59212 | 940788 | Yes |
| ENSG00000120647.9 | CCDC77 | rs6489475 | 7.10E-14 | ##### | Brain - Frontal Cortex (BA9) | 440788 | -59212 | 940788 | Yes |
| ENSG00000120647.9 | CCDC77 | rs10774154 | 7.10E-14 | ##### | Brain - Frontal Cortex (BA9) | 440788 | -59212 | 940788 | Yes |
| ENSG00000120647.9 | CCDC77 | rs507597 | 7.70E-14 | ##### | Brain - Frontal Cortex (BA9) | 440788 | -59212 | 940788 | Yes |
| ENSG00000120647.9 | CCDC77 | rs17800744 | 7.70E-14 | ##### | Brain - Frontal Cortex (BA9) | 440788 | -59212 | 940788 | Yes |
| ENSG00000120647.9 | CCDC77 | rs77854644 | 7.70E-14 | ##### | Brain - Frontal Cortex (BA9) | 440788 | -59212 | 940788 | Yes |
| ENSG00000120647.9 | CCDC77 | rs57022939 | 7.80E-14 | ##### | Brain - Cerebellar Hemisphere | 440788 | -59212 | 940788 | Yes |
| ENSG00000120647.9 | CCDC77 | rs4980875 | 7.80E-14 | ##### | Brain - Cerebellar Hemisphere | 440788 | -59212 | 940788 | Yes |
| ENSG00000120647.9 | CCDC77 | rs61906995 | 7.80E-14 | ##### | Brain - Cerebellar Hemisphere | 440788 | -59212 | 940788 | Yes |
| ENSG00000120647.9 | CCDC77 | rs12582850 | 7.80E-14 | ##### | Brain - Cerebellar Hemisphere | 440788 | -59212 | 940788 | Yes |
| ENSG00000120647.9 | CCDC77 | rs7298019 | 7.80E-14 | ##### | Brain - Cerebellar Hemisphere | 440788 | -59212 | 940788 | Yes |
| ENSG00000120647.9 | CCDC77 | rs2098094 | 7.80E-14 | ##### | Brain - Cerebellar Hemisphere | 440788 | -59212 | 940788 | Yes |
| ENSG00000120647.9 | CCDC77 | rs2159364 | 7.80E-14 | ##### | Brain - Cerebellar Hemisphere | 440788 | -59212 | 940788 | Yes |
| ENSG00000120647.9 | CCDC77 | rs10848868 | 7.80E-14 | ##### | Brain - Cerebellar Hemisphere | 440788 | -59212 | 940788 | Yes |
| ENSG00000120647.9 | CCDC77 | rs1860360 | 7.80E-14 | ##### | Brain - Cerebellar Hemisphere | 440788 | -59212 | 940788 | Yes |
| ENSG00000120647.9 | CCDC77 | rs2041299 | 8.00E-14 | ##### | Brain - Caudate (basal ganglia | 440788 | -59212 | 940788 | Yes |
| ENSG00000120647.9 | CCDC77 | rs524468 | 8.10E-14 | ##### | Brain - Caudate (basal ganglia | 440788 | -59212 | 940788 | Yes |
| ENSG00000120647.9 | CCDC77 | rs7972566 | 8.60E-14 | ##### | Brain - Caudate (basal ganglia | 440788 | -59212 | 940788 | Yes |
| ENSG00000120647.9 | CCDC77 | rs10774170 | 8.60E-14 | ##### | Brain - Caudate (basal ganglia | 440788 | -59212 | 940788 | Yes |

| | | | | | | | | | |
|---|---|---|---|---|---|---|---|---|---|
| ENSG00000120647.9 | CCDC77 | rs1155942 | 9.00E-14 | ##### | Brain - Cerebellar Hemisphere | 440788 | -59212 | 940788 | Yes |
| ENSG00000120647.9 | CCDC77 | rs797765 | 9.10E-14 | ##### | Brain - Frontal Cortex (BA9) | 440788 | -59212 | 940788 | Yes |
| ENSG00000120647.9 | CCDC77 | rs2369280 | 9.10E-14 | ##### | Brain - Hypothalamus | 440788 | -59212 | 940788 | Yes |
| ENSG00000120647.9 | CCDC77 | rs4980884 | 9.20E-14 | ##### | Brain - Cerebellar Hemisphere | 440788 | -59212 | 940788 | Yes |
| ENSG00000120647.9 | CCDC77 | rs11062813 | 1.10E-13 | ##### | Brain - Caudate (basal ganglia | 440788 | -59212 | 940788 | Yes |
| ENSG00000120647.9 | CCDC77 | rs1541463 | 1.10E-13 | ##### | Brain - Hypothalamus | 440788 | -59212 | 940788 | Yes |
| ENSG00000120647.9 | CCDC77 | rs2369278 | 1.10E-13 | ##### | Brain - Hypothalamus | 440788 | -59212 | 940788 | Yes |
| ENSG00000120647.9 | CCDC77 | rs10774163 | 1.10E-13 | ##### | Brain - Hypothalamus | 440788 | -59212 | 940788 | Yes |
| ENSG00000120647.9 | CCDC77 | rs507597 | 1.20E-13 | ##### | Brain - Hypothalamus | 440788 | -59212 | 940788 | Yes |
| ENSG00000120647.9 | CCDC77 | rs17800744 | 1.20E-13 | ##### | Brain - Hypothalamus | 440788 | -59212 | 940788 | Yes |
| ENSG00000120647.9 | CCDC77 | rs77854644 | 1.20E-13 | ##### | Brain - Hypothalamus | 440788 | -59212 | 940788 | Yes |
| ENSG00000120647.9 | CCDC77 | rs486098 | 1.20E-13 | ##### | Brain - Cerebellar Hemisphere | 440788 | -59212 | 940788 | Yes |
| ENSG00000120647.9 | CCDC77 | rs524468 | 1.40E-13 | ##### | Brain - Frontal Cortex (BA9) | 440788 | -59212 | 940788 | Yes |
| ENSG00000120647.9 | CCDC77 | rs61907026 | 1.70E-13 | ##### | Brain - Caudate (basal ganglia | 440788 | -59212 | 940788 | Yes |
| ENSG00000120647.9 | CCDC77 | rs61907030 | 1.70E-13 | ##### | Brain - Caudate (basal ganglia | 440788 | -59212 | 940788 | Yes |
| ENSG00000120647.9 | CCDC77 | rs61907031 | 1.70E-13 | ##### | Brain - Caudate (basal ganglia | 440788 | -59212 | 940788 | Yes |
| ENSG00000120647.9 | CCDC77 | rs11615564 | 1.70E-13 | ##### | Brain - Caudate (basal ganglia | 440788 | -59212 | 940788 | Yes |
| ENSG00000120647.9 | CCDC77 | rs7972566 | 2.00E-13 | ##### | Brain - Spinal cord (cervical c- | 440788 | -59212 | 940788 | Yes |
| ENSG00000120647.9 | CCDC77 | rs797765 | 2.00E-13 | ##### | Brain - Cerebellar Hemisphere | 440788 | -59212 | 940788 | Yes |
| ENSG00000120647.9 | CCDC77 | rs79839498 | 2.40E-13 | ##### | Brain - Caudate (basal ganglia | 440788 | -59212 | 940788 | Yes |
| ENSG00000120647.9 | CCDC77 | rs4980892 | 2.50E-13 | ##### | Brain - Hypothalamus | 440788 | -59212 | 940788 | Yes |
| ENSG00000120647.9 | CCDC77 | rs495316 | 2.50E-13 | ##### | Brain - Cerebellar Hemisphere | 440788 | -59212 | 940788 | Yes |
| ENSG00000120647.9 | CCDC77 | rs2369280 | 2.50E-13 | ##### | Brain - Caudate (basal ganglia | 440788 | -59212 | 940788 | Yes |
| ENSG00000120647.9 | CCDC77 | rs2300131 | 2.90E-13 | ##### | Brain - Cerebellar Hemisphere | 440788 | -59212 | 940788 | Yes |
| ENSG00000120647.9 | CCDC77 | rs2041299 | 3.10E-13 | ##### | Brain - Frontal Cortex (BA9) | 440788 | -59212 | 940788 | Yes |
| ENSG00000120647.9 | CCDC77 | rs111250603 | 3.30E-13 | ##### | Brain - Caudate (basal ganglia | 440788 | -59212 | 940788 | Yes |
| ENSG00000120647.9 | CCDC77 | rs2369277 | 3.30E-13 | ##### | Brain - Caudate (basal ganglia | 440788 | -59212 | 940788 | Yes |
| ENSG00000120647.9 | CCDC77 | rs9805067 | 3.30E-13 | ##### | Brain - Caudate (basal ganglia | 440788 | -59212 | 940788 | Yes |
| ENSG00000120647.9 | CCDC77 | rs1541464 | 3.30E-13 | ##### | Brain - Caudate (basal ganglia | 440788 | -59212 | 940788 | Yes |
| ENSG00000120647.9 | CCDC77 | rs1541465 | 3.30E-13 | ##### | Brain - Caudate (basal ganglia | 440788 | -59212 | 940788 | Yes |
| ENSG00000120647.9 | CCDC77 | rs4980817 | 3.30E-13 | ##### | Brain - Caudate (basal ganglia | 440788 | -59212 | 940788 | Yes |
| ENSG00000120647.9 | CCDC77 | rs4482086 | 3.30E-13 | ##### | Brain - Caudate (basal ganglia | 440788 | -59212 | 940788 | Yes |

| | | | | | | | | | |
|---|---|---|---|---|---|---|---|---|---|
| ENSG00000120647.9 | CCDC77 | rs2369279 | 3.30E-13 | ##### | Brain - Caudate (basal ganglia | 440788 | -59212 | 940788 | Yes |
| ENSG00000120647.9 | CCDC77 | rs11062803 | 3.30E-13 | ##### | Brain - Caudate (basal ganglia | 440788 | -59212 | 940788 | Yes |
| ENSG00000120647.9 | CCDC77 | rs520932 | 3.30E-13 | ##### | Brain - Hypothalamus | 440788 | -59212 | 940788 | Yes |
| ENSG00000120647.9 | CCDC77 | rs520932 | 3.80E-13 | ##### | Brain - Cerebellar Hemisphere | 440788 | -59212 | 940788 | Yes |
| ENSG00000120647.9 | CCDC77 | rs79839498 | 3.80E-13 | ##### | Brain - Frontal Cortex (BA9) | 440788 | -59212 | 940788 | Yes |
| ENSG00000120647.9 | CCDC77 | rs7133796 | 3.90E-13 | ##### | Brain - Putamen (basal gangli | 440788 | -59212 | 940788 | Yes |
| ENSG00000120647.9 | CCDC77 | rs11062813 | 4.00E-13 | ##### | Brain - Spinal cord (cervical c- | 440788 | -59212 | 940788 | Yes |
| ENSG00000120647.9 | CCDC77 | rs3815558 | 4.40E-13 | ##### | Brain - Cerebellar Hemisphere | 440788 | -59212 | 940788 | Yes |
| ENSG00000120647.9 | CCDC77 | rs78998485 | 4.40E-13 | ##### | Brain - Cerebellar Hemisphere | 440788 | -59212 | 940788 | Yes |
| ENSG00000120647.9 | CCDC77 | rs7135768 | 4.40E-13 | ##### | Brain - Cerebellar Hemisphere | 440788 | -59212 | 940788 | Yes |
| ENSG00000120647.9 | CCDC77 | rs6489450 | 4.40E-13 | ##### | Brain - Cerebellar Hemisphere | 440788 | -59212 | 940788 | Yes |
| ENSG00000120647.9 | CCDC77 | rs7138958 | 4.40E-13 | ##### | Brain - Caudate (basal ganglia | 440788 | -59212 | 940788 | Yes |
| ENSG00000120647.9 | CCDC77 | rs2007537 | 4.40E-13 | ##### | Brain - Caudate (basal ganglia | 440788 | -59212 | 940788 | Yes |
| ENSG00000120647.9 | CCDC77 | rs12301816 | 4.40E-13 | ##### | Brain - Caudate (basal ganglia | 440788 | -59212 | 940788 | Yes |
| ENSG00000120647.9 | CCDC77 | rs482514 | 4.50E-13 | ##### | Brain - Cerebellar Hemisphere | 440788 | -59212 | 940788 | Yes |
| ENSG00000120647.9 | CCDC77 | rs7315852 | 4.50E-13 | ##### | Brain - Cerebellar Hemisphere | 440788 | -59212 | 940788 | Yes |
| ENSG00000120647.9 | CCDC77 | rs2284334 | 4.50E-13 | ##### | Brain - Cerebellar Hemisphere | 440788 | -59212 | 940788 | Yes |
| ENSG00000120647.9 | CCDC77 | rs720128 | 4.50E-13 | ##### | Brain - Cerebellar Hemisphere | 440788 | -59212 | 940788 | Yes |
| ENSG00000120647.9 | CCDC77 | rs9739950 | 4.50E-13 | ##### | Brain - Cerebellar Hemisphere | 440788 | -59212 | 940788 | Yes |
| ENSG00000120647.9 | CCDC77 | rs12311569 | 4.50E-13 | ##### | Brain - Cerebellar Hemisphere | 440788 | -59212 | 940788 | Yes |
| ENSG00000120647.9 | CCDC77 | rs111250603 | 5.00E-13 | ##### | Brain - Frontal Cortex (BA9) | 440788 | -59212 | 940788 | Yes |
| ENSG00000120647.9 | CCDC77 | rs2369277 | 5.00E-13 | ##### | Brain - Frontal Cortex (BA9) | 440788 | -59212 | 940788 | Yes |
| ENSG00000120647.9 | CCDC77 | rs9805067 | 5.00E-13 | ##### | Brain - Frontal Cortex (BA9) | 440788 | -59212 | 940788 | Yes |
| ENSG00000120647.9 | CCDC77 | rs1541464 | 5.00E-13 | ##### | Brain - Frontal Cortex (BA9) | 440788 | -59212 | 940788 | Yes |
| ENSG00000120647.9 | CCDC77 | rs1541465 | 5.00E-13 | ##### | Brain - Frontal Cortex (BA9) | 440788 | -59212 | 940788 | Yes |
| ENSG00000120647.9 | CCDC77 | rs4980817 | 5.00E-13 | ##### | Brain - Frontal Cortex (BA9) | 440788 | -59212 | 940788 | Yes |
| ENSG00000120647.9 | CCDC77 | rs4482086 | 5.00E-13 | ##### | Brain - Frontal Cortex (BA9) | 440788 | -59212 | 940788 | Yes |
| ENSG00000120647.9 | CCDC77 | rs2369279 | 5.00E-13 | ##### | Brain - Frontal Cortex (BA9) | 440788 | -59212 | 940788 | Yes |
| ENSG00000120647.9 | CCDC77 | rs11062803 | 5.00E-13 | ##### | Brain - Frontal Cortex (BA9) | 440788 | -59212 | 940788 | Yes |
| ENSG00000120647.9 | CCDC77 | rs7133796 | 5.40E-13 | ##### | Brain - Cortex | 440788 | -59212 | 940788 | Yes |
| ENSG00000120647.9 | CCDC77 | rs10848877 | 5.40E-13 | ##### | Brain - Cerebellar Hemisphere | 440788 | -59212 | 940788 | Yes |
| ENSG00000120647.9 | CCDC77 | rs4980892 | 5.60E-13 | ##### | Brain - Caudate (basal ganglia | 440788 | -59212 | 940788 | Yes |

| | | | | | | | | | |
|---|---|---|---|---|---|---|---|---|---|
| ENSG00000120647.9 | CCDC77 | rs7133796 | 5.60E-13 | ##### | Brain - Spinal cord (cervical c- | 440788 | -59212 | 940788 | Yes |
| ENSG00000120647.9 | CCDC77 | rs10774169 | 5.90E-13 | ##### | Brain - Caudate (basal ganglia | 440788 | -59212 | 940788 | Yes |
| ENSG00000120647.9 | CCDC77 | rs2253565 | 6.00E-13 | ##### | Brain - Cerebellar Hemisphere | 440788 | -59212 | 940788 | Yes |
| ENSG00000120647.9 | CCDC77 | rs7972566 | 6.00E-13 | ##### | Brain - Putamen (basal gangli | 440788 | -59212 | 940788 | Yes |
| ENSG00000120647.9 | CCDC77 | rs10774170 | 6.00E-13 | ##### | Brain - Putamen (basal gangli | 440788 | -59212 | 940788 | Yes |
| ENSG00000120647.9 | CCDC77 | rs111250603 | 6.30E-13 | ##### | Brain - Putamen (basal gangli | 440788 | -59212 | 940788 | Yes |
| ENSG00000120647.9 | CCDC77 | rs2369277 | 6.30E-13 | ##### | Brain - Putamen (basal gangli | 440788 | -59212 | 940788 | Yes |
| ENSG00000120647.9 | CCDC77 | rs9805067 | 6.30E-13 | ##### | Brain - Putamen (basal gangli | 440788 | -59212 | 940788 | Yes |
| ENSG00000120647.9 | CCDC77 | rs1541464 | 6.30E-13 | ##### | Brain - Putamen (basal gangli | 440788 | -59212 | 940788 | Yes |
| ENSG00000120647.9 | CCDC77 | rs1541465 | 6.30E-13 | ##### | Brain - Putamen (basal gangli | 440788 | -59212 | 940788 | Yes |
| ENSG00000120647.9 | CCDC77 | rs4980817 | 6.30E-13 | ##### | Brain - Putamen (basal gangli | 440788 | -59212 | 940788 | Yes |
| ENSG00000120647.9 | CCDC77 | rs4482086 | 6.30E-13 | ##### | Brain - Putamen (basal gangli | 440788 | -59212 | 940788 | Yes |
| ENSG00000120647.9 | CCDC77 | rs2369279 | 6.30E-13 | ##### | Brain - Putamen (basal gangli | 440788 | -59212 | 940788 | Yes |
| ENSG00000120647.9 | CCDC77 | rs11062803 | 6.30E-13 | ##### | Brain - Putamen (basal gangli | 440788 | -59212 | 940788 | Yes |
| ENSG00000120647.9 | CCDC77 | rs10774169 | 6.50E-13 | ##### | Brain - Cerebellar Hemisphere | 440788 | -59212 | 940788 | Yes |
| ENSG00000120647.9 | CCDC77 | rs4980878 | 6.70E-13 | ##### | Brain - Caudate (basal ganglia | 440788 | -59212 | 940788 | Yes |
| ENSG00000120647.9 | CCDC77 | rs10774170 | 8.30E-13 | ##### | Brain - Spinal cord (cervical c- | 440788 | -59212 | 940788 | Yes |
| ENSG00000120647.9 | CCDC77 | rs7133796 | 8.80E-13 | ##### | Brain - Frontal Cortex (BA9) | 440788 | -59212 | 940788 | Yes |
| ENSG00000120647.9 | CCDC77 | rs1155942 | 1.00E-12 | ##### | Brain - Putamen (basal gangli | 440788 | -59212 | 940788 | Yes |
| ENSG00000120647.9 | CCDC77 | rs525381 | 1.10E-12 | ##### | Brain - Frontal Cortex (BA9) | 440788 | -59212 | 940788 | Yes |
| ENSG00000120647.9 | CCDC77 | rs11062813 | 1.10E-12 | ##### | Brain - Frontal Cortex (BA9) | 440788 | -59212 | 940788 | Yes |
| ENSG00000120647.9 | CCDC77 | rs111250603 | 1.10E-12 | ##### | Brain - Spinal cord (cervical c- | 440788 | -59212 | 940788 | Yes |
| ENSG00000120647.9 | CCDC77 | rs2369277 | 1.10E-12 | ##### | Brain - Spinal cord (cervical c- | 440788 | -59212 | 940788 | Yes |
| ENSG00000120647.9 | CCDC77 | rs9805067 | 1.10E-12 | ##### | Brain - Spinal cord (cervical c- | 440788 | -59212 | 940788 | Yes |
| ENSG00000120647.9 | CCDC77 | rs1541464 | 1.10E-12 | ##### | Brain - Spinal cord (cervical c- | 440788 | -59212 | 940788 | Yes |
| ENSG00000120647.9 | CCDC77 | rs1541465 | 1.10E-12 | ##### | Brain - Spinal cord (cervical c- | 440788 | -59212 | 940788 | Yes |
| ENSG00000120647.9 | CCDC77 | rs4980817 | 1.10E-12 | ##### | Brain - Spinal cord (cervical c- | 440788 | -59212 | 940788 | Yes |
| ENSG00000120647.9 | CCDC77 | rs2041299 | 1.40E-12 | ##### | Brain - Spinal cord (cervical c- | 440788 | -59212 | 940788 | Yes |
| ENSG00000120647.9 | CCDC77 | rs4482086 | 1.40E-12 | ##### | Brain - Spinal cord (cervical c- | 440788 | -59212 | 940788 | Yes |
| ENSG00000120647.9 | CCDC77 | rs2369279 | 1.40E-12 | ##### | Brain - Spinal cord (cervical c- | 440788 | -59212 | 940788 | Yes |
| ENSG00000120647.9 | CCDC77 | rs12301816 | 1.50E-12 | ##### | Brain - Hypothalamus | 440788 | -59212 | 940788 | Yes |
| ENSG00000120647.9 | CCDC77 | rs1155942 | 1.60E-12 | ##### | Brain - Caudate (basal ganglia | 440788 | -59212 | 940788 | Yes |

| | | | | | | | | | |
|---|---|---|---|---|---|---|---|---|---|
| ENSG00000120647.9 | CCDC77 | rs61907026 | 1.70E-12 | ##### | Brain - Frontal Cortex (BA9) | 440788 | -59212 | 940788 | Yes |
| ENSG00000120647.9 | CCDC77 | rs61907030 | 1.70E-12 | ##### | Brain - Frontal Cortex (BA9) | 440788 | -59212 | 940788 | Yes |
| ENSG00000120647.9 | CCDC77 | rs61907031 | 1.70E-12 | ##### | Brain - Frontal Cortex (BA9) | 440788 | -59212 | 940788 | Yes |
| ENSG00000120647.9 | CCDC77 | rs11615564 | 1.70E-12 | ##### | Brain - Frontal Cortex (BA9) | 440788 | -59212 | 940788 | Yes |
| ENSG00000120647.9 | CCDC77 | rs1207322 | 2.00E-12 | ##### | Brain - Frontal Cortex (BA9) | 440788 | -59212 | 940788 | Yes |
| ENSG00000120647.9 | CCDC77 | rs4980813 | 2.00E-12 | ##### | Brain - Frontal Cortex (BA9) | 440788 | -59212 | 940788 | Yes |
| ENSG00000120647.9 | CCDC77 | rs4980876 | 2.00E-12 | ##### | Brain - Frontal Cortex (BA9) | 440788 | -59212 | 940788 | Yes |
| ENSG00000120647.9 | CCDC77 | rs61906996 | 2.00E-12 | ##### | Brain - Frontal Cortex (BA9) | 440788 | -59212 | 940788 | Yes |
| ENSG00000120647.9 | CCDC77 | rs4980814 | 2.00E-12 | ##### | Brain - Frontal Cortex (BA9) | 440788 | -59212 | 940788 | Yes |
| ENSG00000120647.9 | CCDC77 | rs7977462 | 2.00E-12 | ##### | Brain - Frontal Cortex (BA9) | 440788 | -59212 | 940788 | Yes |
| ENSG00000120647.9 | CCDC77 | rs10744616 | 2.00E-12 | ##### | Brain - Frontal Cortex (BA9) | 440788 | -59212 | 940788 | Yes |
| ENSG00000120647.9 | CCDC77 | rs10774150 | 2.00E-12 | ##### | Brain - Frontal Cortex (BA9) | 440788 | -59212 | 940788 | Yes |
| ENSG00000120647.9 | CCDC77 | rs3831759 | 2.00E-12 | ##### | Brain - Frontal Cortex (BA9) | 440788 | -59212 | 940788 | Yes |
| ENSG00000120647.9 | CCDC77 | rs10744627 | 2.10E-12 | ##### | Brain - Frontal Cortex (BA9) | 440788 | -59212 | 940788 | Yes |
| ENSG00000120647.9 | CCDC77 | rs372437080 | 2.10E-12 | ##### | Brain - Cortex | 440788 | -59212 | 940788 | Yes |
| ENSG00000120647.9 | CCDC77 | rs7132845 | 2.10E-12 | ##### | Brain - Cerebellar Hemisphere | 440788 | -59212 | 940788 | Yes |
| ENSG00000120647.9 | CCDC77 | rs579526 | 2.40E-12 | ##### | Brain - Frontal Cortex (BA9) | 440788 | -59212 | 940788 | Yes |
| ENSG00000120647.9 | CCDC77 | rs1155942 | 2.50E-12 | ##### | Brain - Spinal cord (cervical c- | 440788 | -59212 | 940788 | Yes |
| ENSG00000120647.9 | CCDC77 | rs372437080 | 2.50E-12 | ##### | Brain - Spinal cord (cervical c- | 440788 | -59212 | 940788 | Yes |
| ENSG00000120647.9 | CCDC77 | rs4980875 | 3.10E-12 | ##### | Brain - Caudate (basal ganglia | 440788 | -59212 | 940788 | Yes |
| ENSG00000120647.9 | CCDC77 | rs61906995 | 3.10E-12 | ##### | Brain - Caudate (basal ganglia | 440788 | -59212 | 940788 | Yes |
| ENSG00000120647.9 | CCDC77 | rs12582850 | 3.10E-12 | ##### | Brain - Caudate (basal ganglia | 440788 | -59212 | 940788 | Yes |
| ENSG00000120647.9 | CCDC77 | rs7298019 | 3.10E-12 | ##### | Brain - Caudate (basal ganglia | 440788 | -59212 | 940788 | Yes |
| ENSG00000120647.9 | CCDC77 | rs2098094 | 3.10E-12 | ##### | Brain - Caudate (basal ganglia | 440788 | -59212 | 940788 | Yes |
| ENSG00000120647.9 | CCDC77 | rs2159364 | 3.10E-12 | ##### | Brain - Caudate (basal ganglia | 440788 | -59212 | 940788 | Yes |
| ENSG00000120647.9 | CCDC77 | rs10848868 | 3.10E-12 | ##### | Brain - Caudate (basal ganglia | 440788 | -59212 | 940788 | Yes |
| ENSG00000120647.9 | CCDC77 | rs1860360 | 3.10E-12 | ##### | Brain - Caudate (basal ganglia | 440788 | -59212 | 940788 | Yes |
| ENSG00000120647.9 | CCDC77 | rs1541463 | 3.20E-12 | ##### | Brain - Caudate (basal ganglia | 440788 | -59212 | 940788 | Yes |
| ENSG00000120647.9 | CCDC77 | rs10848905 | 3.20E-12 | ##### | Brain - Caudate (basal ganglia | 440788 | -59212 | 940788 | Yes |
| ENSG00000120647.9 | CCDC77 | rs2369278 | 3.20E-12 | ##### | Brain - Caudate (basal ganglia | 440788 | -59212 | 940788 | Yes |
| ENSG00000120647.9 | CCDC77 | rs10774163 | 3.20E-12 | ##### | Brain - Caudate (basal ganglia | 440788 | -59212 | 940788 | Yes |
| ENSG00000120647.9 | CCDC77 | rs2253565 | 3.50E-12 | ##### | Brain - Putamen (basal gangli | 440788 | -59212 | 940788 | Yes |

| | | | | | | | | | |
|---|---|---|---|---|---|---|---|---|---|
| ENSG00000120647.9 | CCDC77 | rs1207322 | 3.60E-12 | ##### | Brain - Caudate (basal ganglia | 440788 | -59212 | 940788 | Yes |
| ENSG00000120647.9 | CCDC77 | rs4980813 | 3.60E-12 | ##### | Brain - Caudate (basal ganglia | 440788 | -59212 | 940788 | Yes |
| ENSG00000120647.9 | CCDC77 | rs61906996 | 3.60E-12 | ##### | Brain - Caudate (basal ganglia | 440788 | -59212 | 940788 | Yes |
| ENSG00000120647.9 | CCDC77 | rs4980814 | 3.60E-12 | ##### | Brain - Caudate (basal ganglia | 440788 | -59212 | 940788 | Yes |
| ENSG00000120647.9 | CCDC77 | rs7977462 | 3.60E-12 | ##### | Brain - Caudate (basal ganglia | 440788 | -59212 | 940788 | Yes |
| ENSG00000120647.9 | CCDC77 | rs10744616 | 3.60E-12 | ##### | Brain - Caudate (basal ganglia | 440788 | -59212 | 940788 | Yes |
| ENSG00000120647.9 | CCDC77 | rs10774150 | 3.60E-12 | ##### | Brain - Caudate (basal ganglia | 440788 | -59212 | 940788 | Yes |
| ENSG00000120647.9 | CCDC77 | rs3831759 | 3.60E-12 | ##### | Brain - Caudate (basal ganglia | 440788 | -59212 | 940788 | Yes |
| ENSG00000120647.9 | CCDC77 | rs495973 | 3.60E-12 | ##### | Brain - Frontal Cortex (BA9) | 440788 | -59212 | 940788 | Yes |
| ENSG00000120647.9 | CCDC77 | rs579526 | 3.60E-12 | ##### | Brain - Hypothalamus | 440788 | -59212 | 940788 | Yes |
| ENSG00000120647.9 | CCDC77 | rs1155942 | 3.80E-12 | ##### | Brain - Frontal Cortex (BA9) | 440788 | -59212 | 940788 | Yes |
| ENSG00000120647.9 | CCDC77 | rs7972566 | 3.80E-12 | ##### | Brain - Frontal Cortex (BA9) | 440788 | -59212 | 940788 | Yes |
| ENSG00000120647.9 | CCDC77 | rs10774170 | 3.80E-12 | ##### | Brain - Frontal Cortex (BA9) | 440788 | -59212 | 940788 | Yes |
| ENSG00000120647.9 | CCDC77 | rs579526 | 3.90E-12 | ##### | Brain - Caudate (basal ganglia | 440788 | -59212 | 940788 | Yes |
| ENSG00000120647.9 | CCDC77 | rs486098 | 4.00E-12 | ##### | Brain - Caudate (basal ganglia | 440788 | -59212 | 940788 | Yes |
| ENSG00000120647.9 | CCDC77 | rs10848905 | 4.30E-12 | ##### | Brain - Spinal cord (cervical c- | 440788 | -59212 | 940788 | Yes |
| ENSG00000120647.9 | CCDC77 | rs2369278 | 4.30E-12 | ##### | Brain - Spinal cord (cervical c- | 440788 | -59212 | 940788 | Yes |
| ENSG00000120647.9 | CCDC77 | rs10774163 | 4.30E-12 | ##### | Brain - Spinal cord (cervical c- | 440788 | -59212 | 940788 | Yes |
| ENSG00000120647.9 | CCDC77 | rs495973 | 4.60E-12 | ##### | Brain - Hypothalamus | 440788 | -59212 | 940788 | Yes |
| ENSG00000120647.9 | CCDC77 | rs7136116 | 4.70E-12 | ##### | Brain - Caudate (basal ganglia | 440788 | -59212 | 940788 | Yes |
| ENSG00000120647.9 | CCDC77 | rs11610457 | 4.70E-12 | ##### | Brain - Caudate (basal ganglia | 440788 | -59212 | 940788 | Yes |
| ENSG00000120647.9 | CCDC77 | rs57022939 | 4.70E-12 | ##### | Brain - Caudate (basal ganglia | 440788 | -59212 | 940788 | Yes |
| ENSG00000120647.9 | CCDC77 | rs75894333 | 4.80E-12 | ##### | Brain - Spinal cord (cervical c- | 440788 | -59212 | 940788 | Yes |
| ENSG00000120647.9 | CCDC77 | rs507597 | 4.80E-12 | ##### | Brain - Spinal cord (cervical c- | 440788 | -59212 | 940788 | Yes |
| ENSG00000120647.9 | CCDC77 | rs17800744 | 4.80E-12 | ##### | Brain - Spinal cord (cervical c- | 440788 | -59212 | 940788 | Yes |
| ENSG00000120647.9 | CCDC77 | rs77854644 | 4.80E-12 | ##### | Brain - Spinal cord (cervical c- | 440788 | -59212 | 940788 | Yes |
| ENSG00000120647.9 | CCDC77 | rs11608938 | 4.80E-12 | ##### | Brain - Spinal cord (cervical c- | 440788 | -59212 | 940788 | Yes |
| ENSG00000120647.9 | CCDC77 | rs7965038 | 4.80E-12 | ##### | Brain - Spinal cord (cervical c- | 440788 | -59212 | 940788 | Yes |
| ENSG00000120647.9 | CCDC77 | rs7307606 | 4.80E-12 | ##### | Brain - Spinal cord (cervical c- | 440788 | -59212 | 940788 | Yes |
| ENSG00000120647.9 | CCDC77 | rs10661251 | 4.80E-12 | ##### | Brain - Spinal cord (cervical c- | 440788 | -59212 | 940788 | Yes |
| ENSG00000120647.9 | CCDC77 | rs1011699 | 4.80E-12 | ##### | Brain - Spinal cord (cervical c- | 440788 | -59212 | 940788 | Yes |
| ENSG00000120647.9 | CCDC77 | rs6489475 | 4.80E-12 | ##### | Brain - Spinal cord (cervical c- | 440788 | -59212 | 940788 | Yes |

| | | | | | | | | | |
|---|---|---|---|---|---|---|---|---|---|
| ENSG00000120647.9 | CCDC77 | rs10774154 | 4.80E-12 | ##### | Brain - Spinal cord (cervical c- | 440788 | -59212 | 940788 | Yes |
| ENSG00000120647.9 | CCDC77 | rs1541463 | 4.90E-12 | ##### | Brain - Spinal cord (cervical c- | 440788 | -59212 | 940788 | Yes |
| ENSG00000120647.9 | CCDC77 | rs4980876 | 5.10E-12 | ##### | Brain - Caudate (basal ganglia | 440788 | -59212 | 940788 | Yes |
| ENSG00000120647.9 | CCDC77 | rs10744627 | 5.20E-12 | ##### | Brain - Spinal cord (cervical c- | 440788 | -59212 | 940788 | Yes |
| ENSG00000120647.9 | CCDC77 | rs10744627 | 5.80E-12 | ##### | Brain - Caudate (basal ganglia | 440788 | -59212 | 940788 | Yes |
| ENSG00000120647.9 | CCDC77 | rs11062813 | 6.00E-12 | ##### | Brain - Putamen (basal gangli | 440788 | -59212 | 940788 | Yes |
| ENSG00000120647.9 | CCDC77 | rs4980878 | 6.80E-12 | ##### | Brain - Frontal Cortex (BA9) | 440788 | -59212 | 940788 | Yes |
| ENSG00000120647.9 | CCDC77 | rs2253565 | 7.00E-12 | ##### | Brain - Hypothalamus | 440788 | -59212 | 940788 | Yes |
| ENSG00000120647.9 | CCDC77 | rs111250603 | 7.50E-12 | ##### | Brain - Cortex | 440788 | -59212 | 940788 | Yes |
| ENSG00000120647.9 | CCDC77 | rs2369277 | 7.50E-12 | ##### | Brain - Cortex | 440788 | -59212 | 940788 | Yes |
| ENSG00000120647.9 | CCDC77 | rs9805067 | 7.50E-12 | ##### | Brain - Cortex | 440788 | -59212 | 940788 | Yes |
| ENSG00000120647.9 | CCDC77 | rs1541464 | 7.50E-12 | ##### | Brain - Cortex | 440788 | -59212 | 940788 | Yes |
| ENSG00000120647.9 | CCDC77 | rs1541465 | 7.50E-12 | ##### | Brain - Cortex | 440788 | -59212 | 940788 | Yes |
| ENSG00000120647.9 | CCDC77 | rs4980817 | 7.50E-12 | ##### | Brain - Cortex | 440788 | -59212 | 940788 | Yes |
| ENSG00000120647.9 | CCDC77 | rs524468 | 7.60E-12 | ##### | Brain - Cerebellar Hemisphere | 440788 | -59212 | 940788 | Yes |
| ENSG00000120647.9 | CCDC77 | rs1207324 | 8.90E-12 | ##### | Brain - Caudate (basal ganglia | 440788 | -59212 | 940788 | Yes |
| ENSG00000120647.9 | CCDC77 | rs4980875 | 8.90E-12 | ##### | Brain - Frontal Cortex (BA9) | 440788 | -59212 | 940788 | Yes |
| ENSG00000120647.9 | CCDC77 | rs61906995 | 8.90E-12 | ##### | Brain - Frontal Cortex (BA9) | 440788 | -59212 | 940788 | Yes |
| ENSG00000120647.9 | CCDC77 | rs12582850 | 8.90E-12 | ##### | Brain - Frontal Cortex (BA9) | 440788 | -59212 | 940788 | Yes |
| ENSG00000120647.9 | CCDC77 | rs7298019 | 8.90E-12 | ##### | Brain - Frontal Cortex (BA9) | 440788 | -59212 | 940788 | Yes |
| ENSG00000120647.9 | CCDC77 | rs2098094 | 8.90E-12 | ##### | Brain - Frontal Cortex (BA9) | 440788 | -59212 | 940788 | Yes |
| ENSG00000120647.9 | CCDC77 | rs2159364 | 8.90E-12 | ##### | Brain - Frontal Cortex (BA9) | 440788 | -59212 | 940788 | Yes |
| ENSG00000120647.9 | CCDC77 | rs10848868 | 8.90E-12 | ##### | Brain - Frontal Cortex (BA9) | 440788 | -59212 | 940788 | Yes |
| ENSG00000120647.9 | CCDC77 | rs1860360 | 8.90E-12 | ##### | Brain - Frontal Cortex (BA9) | 440788 | -59212 | 940788 | Yes |
| ENSG00000120647.9 | CCDC77 | rs12301816 | 9.10E-12 | ##### | Brain - Cerebellar Hemisphere | 440788 | -59212 | 940788 | Yes |
| ENSG00000120647.9 | CCDC77 | rs579526 | 9.30E-12 | ##### | Brain - Cerebellar Hemisphere | 440788 | -59212 | 940788 | Yes |
| ENSG00000120647.9 | CCDC77 | rs10744627 | 9.30E-12 | ##### | Brain - Putamen (basal gangli | 440788 | -59212 | 940788 | Yes |
| ENSG00000120647.9 | CCDC77 | rs3815558 | 9.80E-12 | ##### | Brain - Caudate (basal ganglia | 440788 | -59212 | 940788 | Yes |
| ENSG00000120647.9 | CCDC77 | rs78998485 | 9.80E-12 | ##### | Brain - Caudate (basal ganglia | 440788 | -59212 | 940788 | Yes |
| ENSG00000120647.9 | CCDC77 | rs7135768 | 9.80E-12 | ##### | Brain - Caudate (basal ganglia | 440788 | -59212 | 940788 | Yes |
| ENSG00000120647.9 | CCDC77 | rs6489450 | 9.80E-12 | ##### | Brain - Caudate (basal ganglia | 440788 | -59212 | 940788 | Yes |
| ENSG00000120647.9 | CCDC77 | rs3213751 | 1.10E-11 | ##### | Brain - Caudate (basal ganglia | 440788 | -59212 | 940788 | Yes |

| | | | | | | | | | |
|---|---|---|---|---|---|---|---|---|---|
| ENSG00000120647.9 | CCDC77 | rs57022939 | 1.10E-11 | ##### | Brain - Frontal Cortex (BA9) | 440788 | -59212 | 940788 | Yes |
| ENSG00000120647.9 | CCDC77 | rs11062803 | 1.20E-11 | ##### | Brain - Spinal cord (cervical c- | 440788 | -59212 | 940788 | Yes |
| ENSG00000120647.9 | CCDC77 | rs4980884 | 1.20E-11 | ##### | Brain - Caudate (basal ganglia | 440788 | -59212 | 940788 | Yes |
| ENSG00000120647.9 | CCDC77 | rs7965303 | 1.30E-11 | ##### | Brain - Cerebellar Hemisphere | 440788 | -59212 | 940788 | Yes |
| ENSG00000120647.9 | CCDC77 | rs10848879 | 1.30E-11 | ##### | Brain - Cerebellar Hemisphere | 440788 | -59212 | 940788 | Yes |
| ENSG00000120647.9 | CCDC77 | rs75894333 | 1.30E-11 | ##### | Brain - Putamen (basal gangli | 440788 | -59212 | 940788 | Yes |
| ENSG00000120647.9 | CCDC77 | rs507597 | 1.30E-11 | ##### | Brain - Putamen (basal gangli | 440788 | -59212 | 940788 | Yes |
| ENSG00000120647.9 | CCDC77 | rs17800744 | 1.30E-11 | ##### | Brain - Putamen (basal gangli | 440788 | -59212 | 940788 | Yes |
| ENSG00000120647.9 | CCDC77 | rs77854644 | 1.30E-11 | ##### | Brain - Putamen (basal gangli | 440788 | -59212 | 940788 | Yes |
| ENSG00000120647.9 | CCDC77 | rs11608938 | 1.30E-11 | ##### | Brain - Putamen (basal gangli | 440788 | -59212 | 940788 | Yes |
| ENSG00000120647.9 | CCDC77 | rs7965038 | 1.30E-11 | ##### | Brain - Putamen (basal gangli | 440788 | -59212 | 940788 | Yes |
| ENSG00000120647.9 | CCDC77 | rs7307606 | 1.30E-11 | ##### | Brain - Putamen (basal gangli | 440788 | -59212 | 940788 | Yes |
| ENSG00000120647.9 | CCDC77 | rs10661251 | 1.30E-11 | ##### | Brain - Putamen (basal gangli | 440788 | -59212 | 940788 | Yes |
| ENSG00000120647.9 | CCDC77 | rs1011699 | 1.30E-11 | ##### | Brain - Putamen (basal gangli | 440788 | -59212 | 940788 | Yes |
| ENSG00000120647.9 | CCDC77 | rs6489475 | 1.30E-11 | ##### | Brain - Putamen (basal gangli | 440788 | -59212 | 940788 | Yes |
| ENSG00000120647.9 | CCDC77 | rs10774154 | 1.30E-11 | ##### | Brain - Putamen (basal gangli | 440788 | -59212 | 940788 | Yes |
| ENSG00000120647.9 | CCDC77 | rs4980886 | 1.30E-11 | ##### | Brain - Caudate (basal ganglia | 440788 | -59212 | 940788 | Yes |
| ENSG00000120647.9 | CCDC77 | rs61907026 | 1.40E-11 | ##### | Brain - Hypothalamus | 440788 | -59212 | 940788 | Yes |
| ENSG00000120647.9 | CCDC77 | rs79839498 | 1.40E-11 | ##### | Brain - Hypothalamus | 440788 | -59212 | 940788 | Yes |
| ENSG00000120647.9 | CCDC77 | rs61907030 | 1.40E-11 | ##### | Brain - Hypothalamus | 440788 | -59212 | 940788 | Yes |
| ENSG00000120647.9 | CCDC77 | rs61907031 | 1.40E-11 | ##### | Brain - Hypothalamus | 440788 | -59212 | 940788 | Yes |
| ENSG00000120647.9 | CCDC77 | rs11615564 | 1.40E-11 | ##### | Brain - Hypothalamus | 440788 | -59212 | 940788 | Yes |
| ENSG00000120647.9 | CCDC77 | rs7136957 | 1.40E-11 | ##### | Brain - Frontal Cortex (BA9) | 440788 | -59212 | 940788 | Yes |
| ENSG00000120647.9 | CCDC77 | rs2041299 | 1.50E-11 | ##### | Brain - Putamen (basal gangli | 440788 | -59212 | 940788 | Yes |
| ENSG00000120647.9 | CCDC77 | rs7303180 | 1.50E-11 | ##### | Brain - Caudate (basal ganglia | 440788 | -59212 | 940788 | Yes |
| ENSG00000120647.9 | CCDC77 | rs579526 | 1.50E-11 | ##### | Brain - Spinal cord (cervical c- | 440788 | -59212 | 940788 | Yes |
| ENSG00000120647.9 | CCDC77 | rs7136957 | 1.50E-11 | ##### | Brain - Cerebellar Hemisphere | 440788 | -59212 | 940788 | Yes |
| ENSG00000120647.9 | CCDC77 | rs576994 | 1.60E-11 | ##### | Brain - Cerebellar Hemisphere | 440788 | -59212 | 940788 | Yes |
| ENSG00000120647.9 | CCDC77 | rs797759 | 1.60E-11 | ##### | Brain - Cerebellar Hemisphere | 440788 | -59212 | 940788 | Yes |
| ENSG00000120647.9 | CCDC77 | rs479836 | 1.60E-11 | ##### | Brain - Cerebellar Hemisphere | 440788 | -59212 | 940788 | Yes |
| ENSG00000120647.9 | CCDC77 | rs2284332 | 1.60E-11 | ##### | Brain - Cerebellar Hemisphere | 440788 | -59212 | 940788 | Yes |
| ENSG00000120647.9 | CCDC77 | rs2229351 | 1.60E-11 | ##### | Brain - Cerebellar Hemisphere | 440788 | -59212 | 940788 | Yes |

| | | | | | | | | | |
|---|---|---|---|---|---|---|---|---|---|
| ENSG00000120647.9 | CCDC77 | rs4980810 | 1.60E-11 | ##### | Brain - Cerebellar Hemisphere | 440788 | -59212 | 940788 | Yes |
| ENSG00000120647.9 | CCDC77 | rs11062302 | 1.60E-11 | ##### | Brain - Cerebellar Hemisphere | 440788 | -59212 | 940788 | Yes |
| ENSG00000120647.9 | CCDC77 | rs12314497 | 1.60E-11 | ##### | Brain - Cerebellar Hemisphere | 440788 | -59212 | 940788 | Yes |
| ENSG00000120647.9 | CCDC77 | rs534925791 | 1.60E-11 | ##### | Brain - Cerebellar Hemisphere | 440788 | -59212 | 940788 | Yes |
| ENSG00000120647.9 | CCDC77 | rs144729439 | 1.60E-11 | ##### | Brain - Cerebellar Hemisphere | 440788 | -59212 | 940788 | Yes |
| ENSG00000120647.9 | CCDC77 | rs11062385 | 1.60E-11 | ##### | Brain - Cerebellar Hemisphere | 440788 | -59212 | 940788 | Yes |
| ENSG00000120647.9 | CCDC77 | rs11611810 | 1.60E-11 | ##### | Brain - Cerebellar Hemisphere | 440788 | -59212 | 940788 | Yes |
| ENSG00000120647.9 | CCDC77 | rs10848715 | 1.60E-11 | ##### | Brain - Cerebellar Hemisphere | 440788 | -59212 | 940788 | Yes |
| ENSG00000120647.9 | CCDC77 | rs10848716 | 1.60E-11 | ##### | Brain - Cerebellar Hemisphere | 440788 | -59212 | 940788 | Yes |
| ENSG00000120647.9 | CCDC77 | rs61907027 | 1.60E-11 | ##### | Brain - Cerebellar Hemisphere | 440788 | -59212 | 940788 | Yes |
| ENSG00000120647.9 | CCDC77 | rs7301422 | 1.60E-11 | ##### | Brain - Cerebellar Hemisphere | 440788 | -59212 | 940788 | Yes |
| ENSG00000120647.9 | CCDC77 | rs11062512 | 1.60E-11 | ##### | Brain - Cerebellar Hemisphere | 440788 | -59212 | 940788 | Yes |
| ENSG00000120647.9 | CCDC77 | rs3759371 | 1.60E-11 | ##### | Brain - Cerebellar Hemisphere | 440788 | -59212 | 940788 | Yes |
| ENSG00000120647.9 | CCDC77 | rs6489448 | 1.60E-11 | ##### | Brain - Cerebellar Hemisphere | 440788 | -59212 | 940788 | Yes |
| ENSG00000120647.9 | CCDC77 | rs7316660 | 1.60E-11 | ##### | Brain - Cerebellar Hemisphere | 440788 | -59212 | 940788 | Yes |
| ENSG00000120647.9 | CCDC77 | rs6489449 | 1.60E-11 | ##### | Brain - Cerebellar Hemisphere | 440788 | -59212 | 940788 | Yes |
| ENSG00000120647.9 | CCDC77 | rs10735021 | 1.60E-11 | ##### | Brain - Cerebellar Hemisphere | 440788 | -59212 | 940788 | Yes |
| ENSG00000120647.9 | CCDC77 | rs10848845 | 1.60E-11 | ##### | Brain - Cerebellar Hemisphere | 440788 | -59212 | 940788 | Yes |
| ENSG00000120647.9 | CCDC77 | rs10774137 | 1.60E-11 | ##### | Brain - Cerebellar Hemisphere | 440788 | -59212 | 940788 | Yes |
| ENSG00000120647.9 | CCDC77 | rs11835325 | 1.60E-11 | ##### | Brain - Cerebellar Hemisphere | 440788 | -59212 | 940788 | Yes |
| ENSG00000120647.9 | CCDC77 | rs10848849 | 1.60E-11 | ##### | Brain - Cerebellar Hemisphere | 440788 | -59212 | 940788 | Yes |
| ENSG00000120647.9 | CCDC77 | rs6489454 | 1.60E-11 | ##### | Brain - Cerebellar Hemisphere | 440788 | -59212 | 940788 | Yes |
| ENSG00000120647.9 | CCDC77 | rs7134608 | 1.60E-11 | ##### | Brain - Cerebellar Hemisphere | 440788 | -59212 | 940788 | Yes |
| ENSG00000120647.9 | CCDC77 | rs7138890 | 1.60E-11 | ##### | Brain - Cerebellar Hemisphere | 440788 | -59212 | 940788 | Yes |
| ENSG00000120647.9 | CCDC77 | rs10431355 | 1.60E-11 | ##### | Brain - Cerebellar Hemisphere | 440788 | -59212 | 940788 | Yes |
| ENSG00000120647.9 | CCDC77 | rs6489458 | 1.60E-11 | ##### | Brain - Cerebellar Hemisphere | 440788 | -59212 | 940788 | Yes |
| ENSG00000120647.9 | CCDC77 | rs2300134 | 1.60E-11 | ##### | Brain - Cerebellar Hemisphere | 440788 | -59212 | 940788 | Yes |
| ENSG00000120647.9 | CCDC77 | rs3782852 | 1.60E-11 | ##### | Brain - Cerebellar Hemisphere | 440788 | -59212 | 940788 | Yes |
| ENSG00000120647.9 | CCDC77 | rs201124968 | 1.60E-11 | ##### | Brain - Cerebellar Hemisphere | 440788 | -59212 | 940788 | Yes |
| ENSG00000120647.9 | CCDC77 | rs201946814 | 1.60E-11 | ##### | Brain - Cerebellar Hemisphere | 440788 | -59212 | 940788 | Yes |
| ENSG00000120647.9 | CCDC77 | rs6489463 | 1.60E-11 | ##### | Brain - Cerebellar Hemisphere | 440788 | -59212 | 940788 | Yes |
| ENSG00000120647.9 | CCDC77 | rs2079785 | 1.60E-11 | ##### | Brain - Cerebellar Hemisphere | 440788 | -59212 | 940788 | Yes |

| | | | | | | | | | |
|---|---|---|---|---|---|---|---|---|---|
| ENSG00000120647.9 | CCDC77 | rs7963937 | 1.60E-11 | ##### | Brain - Cerebellar Hemisphere | 440788 | -59212 | 940788 | Yes |
| ENSG00000120647.9 | CCDC77 | rs2369276 | 1.60E-11 | ##### | Brain - Cerebellar Hemisphere | 440788 | -59212 | 940788 | Yes |
| ENSG00000120647.9 | CCDC77 | rs7953137 | 1.60E-11 | ##### | Brain - Cerebellar Hemisphere | 440788 | -59212 | 940788 | Yes |
| ENSG00000120647.9 | CCDC77 | rs6489471 | 1.60E-11 | ##### | Brain - Cerebellar Hemisphere | 440788 | -59212 | 940788 | Yes |
| ENSG00000120647.9 | CCDC77 | rs7312065 | 1.60E-11 | ##### | Brain - Cerebellar Hemisphere | 440788 | -59212 | 940788 | Yes |
| ENSG00000120647.9 | CCDC77 | rs34380840 | 1.60E-11 | ##### | Brain - Cerebellar Hemisphere | 440788 | -59212 | 940788 | Yes |
| ENSG00000120647.9 | CCDC77 | rs1541463 | 1.80E-11 | ##### | Brain - Frontal Cortex (BA9) | 440788 | -59212 | 940788 | Yes |
| ENSG00000120647.9 | CCDC77 | rs2369278 | 1.80E-11 | ##### | Brain - Frontal Cortex (BA9) | 440788 | -59212 | 940788 | Yes |
| ENSG00000120647.9 | CCDC77 | rs10774163 | 1.80E-11 | ##### | Brain - Frontal Cortex (BA9) | 440788 | -59212 | 940788 | Yes |
| ENSG00000120647.9 | CCDC77 | rs113574411 | 1.80E-11 | ##### | Brain - Caudate (basal ganglia | 440788 | -59212 | 940788 | Yes |
| ENSG00000120647.9 | CCDC77 | rs479840 | 1.80E-11 | ##### | Brain - Caudate (basal ganglia | 440788 | -59212 | 940788 | Yes |
| ENSG00000120647.9 | CCDC77 | rs479803 | 1.80E-11 | ##### | Brain - Caudate (basal ganglia | 440788 | -59212 | 940788 | Yes |
| ENSG00000120647.9 | CCDC77 | rs72472276 | 1.80E-11 | ##### | Brain - Caudate (basal ganglia | 440788 | -59212 | 940788 | Yes |
| ENSG00000120647.9 | CCDC77 | rs4980877 | 1.80E-11 | ##### | Brain - Caudate (basal ganglia | 440788 | -59212 | 940788 | Yes |
| ENSG00000120647.9 | CCDC77 | rs7963309 | 1.80E-11 | ##### | Brain - Caudate (basal ganglia | 440788 | -59212 | 940788 | Yes |
| ENSG00000120647.9 | CCDC77 | rs7134353 | 1.80E-11 | ##### | Brain - Caudate (basal ganglia | 440788 | -59212 | 940788 | Yes |
| ENSG00000120647.9 | CCDC77 | rs11374245 | 1.80E-11 | ##### | Brain - Caudate (basal ganglia | 440788 | -59212 | 940788 | Yes |
| ENSG00000120647.9 | CCDC77 | rs16929744 | 1.80E-11 | ##### | Brain - Caudate (basal ganglia | 440788 | -59212 | 940788 | Yes |
| ENSG00000120647.9 | CCDC77 | rs525381 | 1.90E-11 | ##### | Brain - Caudate (basal ganglia | 440788 | -59212 | 940788 | Yes |
| ENSG00000120647.9 | CCDC77 | rs495316 | 2.00E-11 | ##### | Brain - Frontal Cortex (BA9) | 440788 | -59212 | 940788 | Yes |
| ENSG00000120647.9 | CCDC77 | rs4482086 | 2.10E-11 | ##### | Brain - Cortex | 440788 | -59212 | 940788 | Yes |
| ENSG00000120647.9 | CCDC77 | rs2369279 | 2.10E-11 | ##### | Brain - Cortex | 440788 | -59212 | 940788 | Yes |
| ENSG00000120647.9 | CCDC77 | rs11062803 | 2.10E-11 | ##### | Brain - Cortex | 440788 | -59212 | 940788 | Yes |
| ENSG00000120647.9 | CCDC77 | rs61907026 | 2.30E-11 | ##### | Brain - Putamen (basal gangli | 440788 | -59212 | 940788 | Yes |
| ENSG00000120647.9 | CCDC77 | rs482514 | 2.30E-11 | ##### | Brain - Caudate (basal ganglia | 440788 | -59212 | 940788 | Yes |
| ENSG00000120647.9 | CCDC77 | rs7315852 | 2.30E-11 | ##### | Brain - Caudate (basal ganglia | 440788 | -59212 | 940788 | Yes |
| ENSG00000120647.9 | CCDC77 | rs2284334 | 2.30E-11 | ##### | Brain - Caudate (basal ganglia | 440788 | -59212 | 940788 | Yes |
| ENSG00000120647.9 | CCDC77 | rs720128 | 2.30E-11 | ##### | Brain - Caudate (basal ganglia | 440788 | -59212 | 940788 | Yes |
| ENSG00000120647.9 | CCDC77 | rs9739950 | 2.30E-11 | ##### | Brain - Caudate (basal ganglia | 440788 | -59212 | 940788 | Yes |
| ENSG00000120647.9 | CCDC77 | rs12311569 | 2.30E-11 | ##### | Brain - Caudate (basal ganglia | 440788 | -59212 | 940788 | Yes |
| ENSG00000120647.9 | CCDC77 | rs520932 | 2.30E-11 | ##### | Brain - Frontal Cortex (BA9) | 440788 | -59212 | 940788 | Yes |
| ENSG00000120647.9 | CCDC77 | rs3213751 | 2.50E-11 | ##### | Brain - Frontal Cortex (BA9) | 440788 | -59212 | 940788 | Yes |

| | | | | | | | | | |
|---|---|---|---|---|---|---|---|---|---|
| ENSG00000120647.9 | CCDC77 | rs4980884 | 2.50E-11 | ##### | Brain - Frontal Cortex (BA9) | 440788 | -59212 | 940788 | Yes |
| ENSG00000120647.9 | CCDC77 | rs576994 | 2.70E-11 | ##### | Brain - Frontal Cortex (BA9) | 440788 | -59212 | 940788 | Yes |
| ENSG00000120647.9 | CCDC77 | rs797759 | 2.70E-11 | ##### | Brain - Frontal Cortex (BA9) | 440788 | -59212 | 940788 | Yes |
| ENSG00000120647.9 | CCDC77 | rs479836 | 2.70E-11 | ##### | Brain - Frontal Cortex (BA9) | 440788 | -59212 | 940788 | Yes |
| ENSG00000120647.9 | CCDC77 | rs4980810 | 2.70E-11 | ##### | Brain - Frontal Cortex (BA9) | 440788 | -59212 | 940788 | Yes |
| ENSG00000120647.9 | CCDC77 | rs11062302 | 2.70E-11 | ##### | Brain - Frontal Cortex (BA9) | 440788 | -59212 | 940788 | Yes |
| ENSG00000120647.9 | CCDC77 | rs12314497 | 2.70E-11 | ##### | Brain - Frontal Cortex (BA9) | 440788 | -59212 | 940788 | Yes |
| ENSG00000120647.9 | CCDC77 | rs534925791 | 2.70E-11 | ##### | Brain - Frontal Cortex (BA9) | 440788 | -59212 | 940788 | Yes |
| ENSG00000120647.9 | CCDC77 | rs144729439 | 2.70E-11 | ##### | Brain - Frontal Cortex (BA9) | 440788 | -59212 | 940788 | Yes |
| ENSG00000120647.9 | CCDC77 | rs11062385 | 2.70E-11 | ##### | Brain - Frontal Cortex (BA9) | 440788 | -59212 | 940788 | Yes |
| ENSG00000120647.9 | CCDC77 | rs11611810 | 2.70E-11 | ##### | Brain - Frontal Cortex (BA9) | 440788 | -59212 | 940788 | Yes |
| ENSG00000120647.9 | CCDC77 | rs10848715 | 2.70E-11 | ##### | Brain - Frontal Cortex (BA9) | 440788 | -59212 | 940788 | Yes |
| ENSG00000120647.9 | CCDC77 | rs10848716 | 2.70E-11 | ##### | Brain - Frontal Cortex (BA9) | 440788 | -59212 | 940788 | Yes |
| ENSG00000120647.9 | CCDC77 | rs61907027 | 2.70E-11 | ##### | Brain - Frontal Cortex (BA9) | 440788 | -59212 | 940788 | Yes |
| ENSG00000120647.9 | CCDC77 | rs7301422 | 2.70E-11 | ##### | Brain - Frontal Cortex (BA9) | 440788 | -59212 | 940788 | Yes |
| ENSG00000120647.9 | CCDC77 | rs11062512 | 2.70E-11 | ##### | Brain - Frontal Cortex (BA9) | 440788 | -59212 | 940788 | Yes |
| ENSG00000120647.9 | CCDC77 | rs3759371 | 2.70E-11 | ##### | Brain - Frontal Cortex (BA9) | 440788 | -59212 | 940788 | Yes |
| ENSG00000120647.9 | CCDC77 | rs6489448 | 2.70E-11 | ##### | Brain - Frontal Cortex (BA9) | 440788 | -59212 | 940788 | Yes |
| ENSG00000120647.9 | CCDC77 | rs7316660 | 2.70E-11 | ##### | Brain - Frontal Cortex (BA9) | 440788 | -59212 | 940788 | Yes |
| ENSG00000120647.9 | CCDC77 | rs6489449 | 2.70E-11 | ##### | Brain - Frontal Cortex (BA9) | 440788 | -59212 | 940788 | Yes |
| ENSG00000120647.9 | CCDC77 | rs10735021 | 2.70E-11 | ##### | Brain - Frontal Cortex (BA9) | 440788 | -59212 | 940788 | Yes |
| ENSG00000120647.9 | CCDC77 | rs10848845 | 2.70E-11 | ##### | Brain - Frontal Cortex (BA9) | 440788 | -59212 | 940788 | Yes |
| ENSG00000120647.9 | CCDC77 | rs10774137 | 2.70E-11 | ##### | Brain - Frontal Cortex (BA9) | 440788 | -59212 | 940788 | Yes |
| ENSG00000120647.9 | CCDC77 | rs11835325 | 2.70E-11 | ##### | Brain - Frontal Cortex (BA9) | 440788 | -59212 | 940788 | Yes |
| ENSG00000120647.9 | CCDC77 | rs10848849 | 2.70E-11 | ##### | Brain - Frontal Cortex (BA9) | 440788 | -59212 | 940788 | Yes |
| ENSG00000120647.9 | CCDC77 | rs6489454 | 2.70E-11 | ##### | Brain - Frontal Cortex (BA9) | 440788 | -59212 | 940788 | Yes |
| ENSG00000120647.9 | CCDC77 | rs7134608 | 2.70E-11 | ##### | Brain - Frontal Cortex (BA9) | 440788 | -59212 | 940788 | Yes |
| ENSG00000120647.9 | CCDC77 | rs7138890 | 2.70E-11 | ##### | Brain - Frontal Cortex (BA9) | 440788 | -59212 | 940788 | Yes |
| ENSG00000120647.9 | CCDC77 | rs10431355 | 2.70E-11 | ##### | Brain - Frontal Cortex (BA9) | 440788 | -59212 | 940788 | Yes |
| ENSG00000120647.9 | CCDC77 | rs6489458 | 2.70E-11 | ##### | Brain - Frontal Cortex (BA9) | 440788 | -59212 | 940788 | Yes |
| ENSG00000120647.9 | CCDC77 | rs2300134 | 2.70E-11 | ##### | Brain - Frontal Cortex (BA9) | 440788 | -59212 | 940788 | Yes |
| ENSG00000120647.9 | CCDC77 | rs3782852 | 2.70E-11 | ##### | Brain - Frontal Cortex (BA9) | 440788 | -59212 | 940788 | Yes |

| | | | | | | | | | |
|---|---|---|---|---|---|---|---|---|---|
| ENSG00000120647.9 | CCDC77 | rs201946814 | 2.70E-11 | ##### | Brain - Frontal Cortex (BA9) | 440788 | -59212 | 940788 | Yes |
| ENSG00000120647.9 | CCDC77 | rs6489463 | 2.70E-11 | ##### | Brain - Frontal Cortex (BA9) | 440788 | -59212 | 940788 | Yes |
| ENSG00000120647.9 | CCDC77 | rs2079785 | 2.70E-11 | ##### | Brain - Frontal Cortex (BA9) | 440788 | -59212 | 940788 | Yes |
| ENSG00000120647.9 | CCDC77 | rs7963937 | 2.70E-11 | ##### | Brain - Frontal Cortex (BA9) | 440788 | -59212 | 940788 | Yes |
| ENSG00000120647.9 | CCDC77 | rs2369276 | 2.70E-11 | ##### | Brain - Frontal Cortex (BA9) | 440788 | -59212 | 940788 | Yes |
| ENSG00000120647.9 | CCDC77 | rs7953137 | 2.70E-11 | ##### | Brain - Frontal Cortex (BA9) | 440788 | -59212 | 940788 | Yes |
| ENSG00000120647.9 | CCDC77 | rs6489471 | 2.70E-11 | ##### | Brain - Frontal Cortex (BA9) | 440788 | -59212 | 940788 | Yes |
| ENSG00000120647.9 | CCDC77 | rs7312065 | 2.70E-11 | ##### | Brain - Frontal Cortex (BA9) | 440788 | -59212 | 940788 | Yes |
| ENSG00000120647.9 | CCDC77 | rs34380840 | 2.70E-11 | ##### | Brain - Frontal Cortex (BA9) | 440788 | -59212 | 940788 | Yes |
| ENSG00000120647.9 | CCDC77 | rs7138981 | 2.70E-11 | ##### | Brain - Cerebellar Hemisphere | 440788 | -59212 | 940788 | Yes |
| ENSG00000120647.9 | CCDC77 | rs9669094 | 2.70E-11 | ##### | Brain - Cerebellar Hemisphere | 440788 | -59212 | 940788 | Yes |
| ENSG00000120647.9 | CCDC77 | rs7132763 | 2.70E-11 | ##### | Brain - Cerebellar Hemisphere | 440788 | -59212 | 940788 | Yes |
| ENSG00000120647.9 | CCDC77 | rs2015554 | 2.70E-11 | ##### | Brain - Cerebellar Hemisphere | 440788 | -59212 | 940788 | Yes |
| ENSG00000120647.9 | CCDC77 | rs10774152 | 2.90E-11 | ##### | Brain - Cerebellar Hemisphere | 440788 | -59212 | 940788 | Yes |
| ENSG00000120647.9 | CCDC77 | rs10774153 | 2.90E-11 | ##### | Brain - Cerebellar Hemisphere | 440788 | -59212 | 940788 | Yes |
| ENSG00000120647.9 | CCDC77 | rs495316 | 2.90E-11 | ##### | Brain - Caudate (basal ganglia | 440788 | -59212 | 940788 | Yes |
| ENSG00000120647.9 | CCDC77 | rs764614 | 3.00E-11 | ##### | Brain - Cerebellar Hemisphere | 440788 | -59212 | 940788 | Yes |
| ENSG00000120647.9 | CCDC77 | rs10848905 | 3.00E-11 | ##### | Brain - Frontal Cortex (BA9) | 440788 | -59212 | 940788 | Yes |
| ENSG00000120647.9 | CCDC77 | rs797765 | 3.10E-11 | ##### | Brain - Caudate (basal ganglia | 440788 | -59212 | 940788 | Yes |
| ENSG00000120647.9 | CCDC77 | rs764614 | 3.20E-11 | ##### | Brain - Caudate (basal ganglia | 440788 | -59212 | 940788 | Yes |
| ENSG00000120647.9 | CCDC77 | rs7138981 | 3.20E-11 | ##### | Brain - Caudate (basal ganglia | 440788 | -59212 | 940788 | Yes |
| ENSG00000120647.9 | CCDC77 | rs9669094 | 3.20E-11 | ##### | Brain - Caudate (basal ganglia | 440788 | -59212 | 940788 | Yes |
| ENSG00000120647.9 | CCDC77 | rs7132763 | 3.20E-11 | ##### | Brain - Caudate (basal ganglia | 440788 | -59212 | 940788 | Yes |
| ENSG00000120647.9 | CCDC77 | rs2015554 | 3.20E-11 | ##### | Brain - Caudate (basal ganglia | 440788 | -59212 | 940788 | Yes |
| ENSG00000120647.9 | CCDC77 | rs2253565 | 3.20E-11 | ##### | Brain - Spinal cord (cervical c- | 440788 | -59212 | 940788 | Yes |
| ENSG00000120647.9 | CCDC77 | rs77074951 | 3.40E-11 | ##### | Brain - Caudate (basal ganglia | 440788 | -59212 | 940788 | Yes |
| ENSG00000120647.9 | CCDC77 | rs10431354 | 3.50E-11 | ##### | Brain - Cerebellar Hemisphere | 440788 | -59212 | 940788 | Yes |
| ENSG00000120647.9 | CCDC77 | rs7303180 | 3.50E-11 | ##### | Brain - Cerebellar Hemisphere | 440788 | -59212 | 940788 | Yes |
| ENSG00000120647.9 | CCDC77 | rs11613357 | 3.70E-11 | ##### | Brain - Hypothalamus | 440788 | -59212 | 940788 | Yes |
| ENSG00000120647.9 | CCDC77 | rs797765 | 3.70E-11 | ##### | Brain - Hypothalamus | 440788 | -59212 | 940788 | Yes |
| ENSG00000120647.9 | CCDC77 | rs4980876 | 3.80E-11 | ##### | Brain - Putamen (basal gangli | 440788 | -59212 | 940788 | Yes |
| ENSG00000120647.9 | CCDC77 | rs4980878 | 3.90E-11 | ##### | Brain - Putamen (basal gangli | 440788 | -59212 | 940788 | Yes |

| | | | | | | | | | |
|---|---|---|---|---|---|---|---|---|---|
| ENSG00000120647.9 | CCDC77 | rs482514 | 3.90E-11 | ##### | Brain - Frontal Cortex (BA9) | 440788 | -59212 | 940788 | Yes |
| ENSG00000120647.9 | CCDC77 | rs7315852 | 3.90E-11 | ##### | Brain - Frontal Cortex (BA9) | 440788 | -59212 | 940788 | Yes |
| ENSG00000120647.9 | CCDC77 | rs2284334 | 3.90E-11 | ##### | Brain - Frontal Cortex (BA9) | 440788 | -59212 | 940788 | Yes |
| ENSG00000120647.9 | CCDC77 | rs720128 | 3.90E-11 | ##### | Brain - Frontal Cortex (BA9) | 440788 | -59212 | 940788 | Yes |
| ENSG00000120647.9 | CCDC77 | rs9739950 | 3.90E-11 | ##### | Brain - Frontal Cortex (BA9) | 440788 | -59212 | 940788 | Yes |
| ENSG00000120647.9 | CCDC77 | rs12311569 | 3.90E-11 | ##### | Brain - Frontal Cortex (BA9) | 440788 | -59212 | 940788 | Yes |
| ENSG00000120647.9 | CCDC77 | rs201124968 | 4.00E-11 | ##### | Brain - Frontal Cortex (BA9) | 440788 | -59212 | 940788 | Yes |
| ENSG00000120647.9 | CCDC77 | rs61907030 | 4.10E-11 | ##### | Brain - Putamen (basal gangli | 440788 | -59212 | 940788 | Yes |
| ENSG00000120647.9 | CCDC77 | rs61907031 | 4.10E-11 | ##### | Brain - Putamen (basal gangli | 440788 | -59212 | 940788 | Yes |
| ENSG00000120647.9 | CCDC77 | rs11615564 | 4.10E-11 | ##### | Brain - Putamen (basal gangli | 440788 | -59212 | 940788 | Yes |
| ENSG00000120647.9 | CCDC77 | rs12298843 | 4.10E-11 | ##### | Brain - Cerebellar Hemisphere | 440788 | -59212 | 940788 | Yes |
| ENSG00000120647.9 | CCDC77 | rs7965303 | 4.20E-11 | ##### | Brain - Frontal Cortex (BA9) | 440788 | -59212 | 940788 | Yes |
| ENSG00000120647.9 | CCDC77 | rs10848879 | 4.20E-11 | ##### | Brain - Frontal Cortex (BA9) | 440788 | -59212 | 940788 | Yes |
| ENSG00000120647.9 | CCDC77 | rs79839498 | 4.30E-11 | ##### | Brain - Putamen (basal gangli | 440788 | -59212 | 940788 | Yes |
| ENSG00000120647.9 | CCDC77 | rs2369280 | 4.30E-11 | ##### | Brain - Frontal Cortex (BA9) | 440788 | -59212 | 940788 | Yes |
| ENSG00000120647.9 | CCDC77 | rs2253565 | 4.30E-11 | ##### | Brain - Caudate (basal ganglia | 440788 | -59212 | 940788 | Yes |
| ENSG00000120647.9 | CCDC77 | rs4980876 | 4.40E-11 | ##### | Brain - Hypothalamus | 440788 | -59212 | 940788 | Yes |
| ENSG00000120647.9 | CCDC77 | rs4980874 | 4.40E-11 | ##### | Brain - Caudate (basal ganglia | 440788 | -59212 | 940788 | Yes |
| ENSG00000120647.9 | CCDC77 | rs524468 | 4.50E-11 | ##### | Brain - Hypothalamus | 440788 | -59212 | 940788 | Yes |
| ENSG00000120647.9 | CCDC77 | rs2284332 | 4.60E-11 | ##### | Brain - Frontal Cortex (BA9) | 440788 | -59212 | 940788 | Yes |
| ENSG00000120647.9 | CCDC77 | rs2229351 | 4.60E-11 | ##### | Brain - Frontal Cortex (BA9) | 440788 | -59212 | 940788 | Yes |
| ENSG00000120647.9 | CCDC77 | rs1207324 | 4.60E-11 | ##### | Brain - Frontal Cortex (BA9) | 440788 | -59212 | 940788 | Yes |
| ENSG00000120647.9 | CCDC77 | rs797765 | 4.60E-11 | ##### | Brain - Spinal cord (cervical c- | 440788 | -59212 | 940788 | Yes |
| ENSG00000120647.9 | CCDC77 | rs4980811 | 4.80E-11 | ##### | Brain - Caudate (basal ganglia | 440788 | -59212 | 940788 | Yes |
| ENSG00000120647.9 | CCDC77 | rs10848877 | 5.20E-11 | ##### | Brain - Caudate (basal ganglia | 440788 | -59212 | 940788 | Yes |
| ENSG00000120647.9 | CCDC77 | rs1207322 | 5.20E-11 | ##### | Brain - Putamen (basal gangli | 440788 | -59212 | 940788 | Yes |
| ENSG00000120647.9 | CCDC77 | rs4980813 | 5.20E-11 | ##### | Brain - Putamen (basal gangli | 440788 | -59212 | 940788 | Yes |
| ENSG00000120647.9 | CCDC77 | rs61906996 | 5.20E-11 | ##### | Brain - Putamen (basal gangli | 440788 | -59212 | 940788 | Yes |
| ENSG00000120647.9 | CCDC77 | rs4980814 | 5.20E-11 | ##### | Brain - Putamen (basal gangli | 440788 | -59212 | 940788 | Yes |
| ENSG00000120647.9 | CCDC77 | rs7977462 | 5.20E-11 | ##### | Brain - Putamen (basal gangli | 440788 | -59212 | 940788 | Yes |
| ENSG00000120647.9 | CCDC77 | rs10744616 | 5.20E-11 | ##### | Brain - Putamen (basal gangli | 440788 | -59212 | 940788 | Yes |
| ENSG00000120647.9 | CCDC77 | rs10774150 | 5.20E-11 | ##### | Brain - Putamen (basal gangli | 440788 | -59212 | 940788 | Yes |

| | | | | | | | | | |
|---|---|---|---|---|---|---|---|---|---|
| ENSG00000120647.9 | CCDC77 | rs3831759 | 5.20E-11 | ##### | Brain - Putamen (basal gangli | 440788 | -59212 | 940788 | Yes |
| ENSG00000120647.9 | CCDC77 | rs525381 | 5.20E-11 | ##### | Brain - Cerebellum | 440788 | -59212 | 940788 | Yes |
| ENSG00000120647.9 | CCDC77 | rs4980886 | 5.30E-11 | ##### | Brain - Cerebellar Hemisphere | 440788 | -59212 | 940788 | Yes |
| ENSG00000120647.9 | CCDC77 | rs3753158 | 5.50E-11 | ##### | Brain - Cerebellar Hemisphere | 440788 | -59212 | 940788 | Yes |
| ENSG00000120647.9 | CCDC77 | rs3753158 | 5.80E-11 | ##### | Brain - Caudate (basal ganglia | 440788 | -59212 | 940788 | Yes |
| ENSG00000120647.9 | CCDC77 | rs2052279 | 6.00E-11 | ##### | Brain - Cerebellar Hemisphere | 440788 | -59212 | 940788 | Yes |
| ENSG00000120647.9 | CCDC77 | rs78813329 | 6.00E-11 | ##### | Brain - Cerebellar Hemisphere | 440788 | -59212 | 940788 | Yes |
| ENSG00000120647.9 | CCDC77 | rs3753157 | 6.00E-11 | ##### | Brain - Cerebellar Hemisphere | 440788 | -59212 | 940788 | Yes |
| ENSG00000120647.9 | CCDC77 | rs4980883 | 6.00E-11 | ##### | Brain - Cerebellar Hemisphere | 440788 | -59212 | 940788 | Yes |
| ENSG00000120647.9 | CCDC77 | rs137910445 | 6.00E-11 | ##### | Brain - Cerebellar Hemisphere | 440788 | -59212 | 940788 | Yes |
| ENSG00000120647.9 | CCDC77 | rs11062537 | 6.00E-11 | ##### | Brain - Cerebellar Hemisphere | 440788 | -59212 | 940788 | Yes |
| ENSG00000120647.9 | CCDC77 | rs715230 | 6.00E-11 | ##### | Brain - Cerebellar Hemisphere | 440788 | -59212 | 940788 | Yes |
| ENSG00000120647.9 | CCDC77 | rs715229 | 6.00E-11 | ##### | Brain - Cerebellar Hemisphere | 440788 | -59212 | 940788 | Yes |
| ENSG00000120647.9 | CCDC77 | rs715228 | 6.00E-11 | ##### | Brain - Cerebellar Hemisphere | 440788 | -59212 | 940788 | Yes |
| ENSG00000120647.9 | CCDC77 | rs10848838 | 6.00E-11 | ##### | Brain - Cerebellar Hemisphere | 440788 | -59212 | 940788 | Yes |
| ENSG00000120647.9 | CCDC77 | rs372437080 | 6.10E-11 | ##### | Brain - Putamen (basal gangli | 440788 | -59212 | 940788 | Yes |
| ENSG00000120647.9 | CCDC77 | rs201124968 | 6.20E-11 | ##### | Brain - Caudate (basal ganglia | 440788 | -59212 | 940788 | Yes |
| ENSG00000120647.9 | CCDC77 | rs525381 | 6.30E-11 | ##### | Brain - Cortex | 440788 | -59212 | 940788 | Yes |
| ENSG00000120647.9 | CCDC77 | rs520932 | 6.40E-11 | ##### | Brain - Spinal cord (cervical c- | 440788 | -59212 | 940788 | Yes |
| ENSG00000120647.9 | CCDC77 | rs3213750 | 6.70E-11 | ##### | Brain - Caudate (basal ganglia | 440788 | -59212 | 940788 | Yes |
| ENSG00000120647.9 | CCDC77 | rs3753156 | 6.70E-11 | ##### | Brain - Caudate (basal ganglia | 440788 | -59212 | 940788 | Yes |
| ENSG00000120647.9 | CCDC77 | rs12581260 | 6.70E-11 | ##### | Brain - Caudate (basal ganglia | 440788 | -59212 | 940788 | Yes |
| ENSG00000120647.9 | CCDC77 | rs4980881 | 6.70E-11 | ##### | Brain - Caudate (basal ganglia | 440788 | -59212 | 940788 | Yes |
| ENSG00000120647.9 | CCDC77 | rs16929783 | 6.70E-11 | ##### | Brain - Caudate (basal ganglia | 440788 | -59212 | 940788 | Yes |
| ENSG00000120647.9 | CCDC77 | rs57022939 | 6.80E-11 | ##### | Brain - Putamen (basal gangli | 440788 | -59212 | 940788 | Yes |
| ENSG00000120647.9 | CCDC77 | rs4980875 | 6.80E-11 | ##### | Brain - Putamen (basal gangli | 440788 | -59212 | 940788 | Yes |
| ENSG00000120647.9 | CCDC77 | rs61906995 | 6.80E-11 | ##### | Brain - Putamen (basal gangli | 440788 | -59212 | 940788 | Yes |
| ENSG00000120647.9 | CCDC77 | rs12582850 | 6.80E-11 | ##### | Brain - Putamen (basal gangli | 440788 | -59212 | 940788 | Yes |
| ENSG00000120647.9 | CCDC77 | rs7298019 | 6.80E-11 | ##### | Brain - Putamen (basal gangli | 440788 | -59212 | 940788 | Yes |
| ENSG00000120647.9 | CCDC77 | rs2098094 | 6.80E-11 | ##### | Brain - Putamen (basal gangli | 440788 | -59212 | 940788 | Yes |
| ENSG00000120647.9 | CCDC77 | rs2159364 | 6.80E-11 | ##### | Brain - Putamen (basal gangli | 440788 | -59212 | 940788 | Yes |
| ENSG00000120647.9 | CCDC77 | rs10848868 | 6.80E-11 | ##### | Brain - Putamen (basal gangli | 440788 | -59212 | 940788 | Yes |

| | | | | | | | | | |
|---|---|---|---|---|---|---|---|---|---|
| ENSG00000120647.9 | CCDC77 | rs1860360 | 6.80E-11 | ##### | Brain - Putamen (basal gangli | 440788 | -59212 | 940788 | Yes |
| ENSG00000120647.9 | CCDC77 | rs10774152 | 6.80E-11 | ##### | Brain - Caudate (basal ganglia | 440788 | -59212 | 940788 | Yes |
| ENSG00000120647.9 | CCDC77 | rs10774153 | 6.80E-11 | ##### | Brain - Caudate (basal ganglia | 440788 | -59212 | 940788 | Yes |
| ENSG00000120647.9 | CCDC77 | rs10431354 | 6.80E-11 | ##### | Brain - Frontal Cortex (BA9) | 440788 | -59212 | 940788 | Yes |
| ENSG00000120647.9 | CCDC77 | rs576994 | 7.00E-11 | ##### | Brain - Caudate (basal ganglia | 440788 | -59212 | 940788 | Yes |
| ENSG00000120647.9 | CCDC77 | rs797759 | 7.00E-11 | ##### | Brain - Caudate (basal ganglia | 440788 | -59212 | 940788 | Yes |
| ENSG00000120647.9 | CCDC77 | rs479836 | 7.00E-11 | ##### | Brain - Caudate (basal ganglia | 440788 | -59212 | 940788 | Yes |
| ENSG00000120647.9 | CCDC77 | rs4980810 | 7.00E-11 | ##### | Brain - Caudate (basal ganglia | 440788 | -59212 | 940788 | Yes |
| ENSG00000120647.9 | CCDC77 | rs11062302 | 7.00E-11 | ##### | Brain - Caudate (basal ganglia | 440788 | -59212 | 940788 | Yes |
| ENSG00000120647.9 | CCDC77 | rs12314497 | 7.00E-11 | ##### | Brain - Caudate (basal ganglia | 440788 | -59212 | 940788 | Yes |
| ENSG00000120647.9 | CCDC77 | rs534925791 | 7.00E-11 | ##### | Brain - Caudate (basal ganglia | 440788 | -59212 | 940788 | Yes |
| ENSG00000120647.9 | CCDC77 | rs144729439 | 7.00E-11 | ##### | Brain - Caudate (basal ganglia | 440788 | -59212 | 940788 | Yes |
| ENSG00000120647.9 | CCDC77 | rs11062385 | 7.00E-11 | ##### | Brain - Caudate (basal ganglia | 440788 | -59212 | 940788 | Yes |
| ENSG00000120647.9 | CCDC77 | rs11611810 | 7.00E-11 | ##### | Brain - Caudate (basal ganglia | 440788 | -59212 | 940788 | Yes |
| ENSG00000120647.9 | CCDC77 | rs10848715 | 7.00E-11 | ##### | Brain - Caudate (basal ganglia | 440788 | -59212 | 940788 | Yes |
| ENSG00000120647.9 | CCDC77 | rs10848716 | 7.00E-11 | ##### | Brain - Caudate (basal ganglia | 440788 | -59212 | 940788 | Yes |
| ENSG00000120647.9 | CCDC77 | rs61907027 | 7.00E-11 | ##### | Brain - Caudate (basal ganglia | 440788 | -59212 | 940788 | Yes |
| ENSG00000120647.9 | CCDC77 | rs7301422 | 7.00E-11 | ##### | Brain - Caudate (basal ganglia | 440788 | -59212 | 940788 | Yes |
| ENSG00000120647.9 | CCDC77 | rs11062512 | 7.00E-11 | ##### | Brain - Caudate (basal ganglia | 440788 | -59212 | 940788 | Yes |
| ENSG00000120647.9 | CCDC77 | rs3759371 | 7.00E-11 | ##### | Brain - Caudate (basal ganglia | 440788 | -59212 | 940788 | Yes |
| ENSG00000120647.9 | CCDC77 | rs6489448 | 7.00E-11 | ##### | Brain - Caudate (basal ganglia | 440788 | -59212 | 940788 | Yes |
| ENSG00000120647.9 | CCDC77 | rs7316660 | 7.00E-11 | ##### | Brain - Caudate (basal ganglia | 440788 | -59212 | 940788 | Yes |
| ENSG00000120647.9 | CCDC77 | rs6489449 | 7.00E-11 | ##### | Brain - Caudate (basal ganglia | 440788 | -59212 | 940788 | Yes |
| ENSG00000120647.9 | CCDC77 | rs10735021 | 7.00E-11 | ##### | Brain - Caudate (basal ganglia | 440788 | -59212 | 940788 | Yes |
| ENSG00000120647.9 | CCDC77 | rs10848845 | 7.00E-11 | ##### | Brain - Caudate (basal ganglia | 440788 | -59212 | 940788 | Yes |
| ENSG00000120647.9 | CCDC77 | rs10774137 | 7.00E-11 | ##### | Brain - Caudate (basal ganglia | 440788 | -59212 | 940788 | Yes |
| ENSG00000120647.9 | CCDC77 | rs11835325 | 7.00E-11 | ##### | Brain - Caudate (basal ganglia | 440788 | -59212 | 940788 | Yes |
| ENSG00000120647.9 | CCDC77 | rs10848849 | 7.00E-11 | ##### | Brain - Caudate (basal ganglia | 440788 | -59212 | 940788 | Yes |
| ENSG00000120647.9 | CCDC77 | rs6489454 | 7.00E-11 | ##### | Brain - Caudate (basal ganglia | 440788 | -59212 | 940788 | Yes |
| ENSG00000120647.9 | CCDC77 | rs7134608 | 7.00E-11 | ##### | Brain - Caudate (basal ganglia | 440788 | -59212 | 940788 | Yes |
| ENSG00000120647.9 | CCDC77 | rs7138890 | 7.00E-11 | ##### | Brain - Caudate (basal ganglia | 440788 | -59212 | 940788 | Yes |
| ENSG00000120647.9 | CCDC77 | rs10431355 | 7.00E-11 | ##### | Brain - Caudate (basal ganglia | 440788 | -59212 | 940788 | Yes |

| | | | | | | | | | |
|---|---|---|---|---|---|---|---|---|---|
| ENSG00000120647.9 | CCDC77 | rs6489458 | 7.00E-11 | ##### | Brain - Caudate (basal ganglia | 440788 | -59212 | 940788 | Yes |
| ENSG00000120647.9 | CCDC77 | rs2300134 | 7.00E-11 | ##### | Brain - Caudate (basal ganglia | 440788 | -59212 | 940788 | Yes |
| ENSG00000120647.9 | CCDC77 | rs3782852 | 7.00E-11 | ##### | Brain - Caudate (basal ganglia | 440788 | -59212 | 940788 | Yes |
| ENSG00000120647.9 | CCDC77 | rs201946814 | 7.00E-11 | ##### | Brain - Caudate (basal ganglia | 440788 | -59212 | 940788 | Yes |
| ENSG00000120647.9 | CCDC77 | rs6489463 | 7.00E-11 | ##### | Brain - Caudate (basal ganglia | 440788 | -59212 | 940788 | Yes |
| ENSG00000120647.9 | CCDC77 | rs2079785 | 7.00E-11 | ##### | Brain - Caudate (basal ganglia | 440788 | -59212 | 940788 | Yes |
| ENSG00000120647.9 | CCDC77 | rs7963937 | 7.00E-11 | ##### | Brain - Caudate (basal ganglia | 440788 | -59212 | 940788 | Yes |
| ENSG00000120647.9 | CCDC77 | rs2369276 | 7.00E-11 | ##### | Brain - Caudate (basal ganglia | 440788 | -59212 | 940788 | Yes |
| ENSG00000120647.9 | CCDC77 | rs7953137 | 7.00E-11 | ##### | Brain - Caudate (basal ganglia | 440788 | -59212 | 940788 | Yes |
| ENSG00000120647.9 | CCDC77 | rs6489471 | 7.00E-11 | ##### | Brain - Caudate (basal ganglia | 440788 | -59212 | 940788 | Yes |
| ENSG00000120647.9 | CCDC77 | rs7312065 | 7.00E-11 | ##### | Brain - Caudate (basal ganglia | 440788 | -59212 | 940788 | Yes |
| ENSG00000120647.9 | CCDC77 | rs7136957 | 7.00E-11 | ##### | Brain - Caudate (basal ganglia | 440788 | -59212 | 940788 | Yes |
| ENSG00000120647.9 | CCDC77 | rs34380840 | 7.00E-11 | ##### | Brain - Caudate (basal ganglia | 440788 | -59212 | 940788 | Yes |
| ENSG00000120647.9 | CCDC77 | rs495316 | 7.10E-11 | ##### | Brain - Hypothalamus | 440788 | -59212 | 940788 | Yes |
| ENSG00000120647.9 | CCDC77 | rs4980878 | 7.10E-11 | ##### | Brain - Spinal cord (cervical c- | 440788 | -59212 | 940788 | Yes |
| ENSG00000120647.9 | CCDC77 | rs4980878 | 7.10E-11 | ##### | Brain - Hypothalamus | 440788 | -59212 | 940788 | Yes |
| ENSG00000120647.9 | CCDC77 | rs10848877 | 7.50E-11 | ##### | Brain - Frontal Cortex (BA9) | 440788 | -59212 | 940788 | Yes |
| ENSG00000120647.9 | CCDC77 | rs495316 | 8.50E-11 | ##### | Brain - Putamen (basal gangli | 440788 | -59212 | 940788 | Yes |
| ENSG00000120647.9 | CCDC77 | rs4980896 | 8.50E-11 | ##### | Brain - Hypothalamus | 440788 | -59212 | 940788 | Yes |
| ENSG00000120647.9 | CCDC77 | rs524468 | 8.60E-11 | ##### | Brain - Spinal cord (cervical c- | 440788 | -59212 | 940788 | Yes |
| ENSG00000120647.9 | CCDC77 | rs113574411 | 8.70E-11 | ##### | Brain - Cerebellar Hemisphere | 440788 | -59212 | 940788 | Yes |
| ENSG00000120647.9 | CCDC77 | rs479840 | 8.70E-11 | ##### | Brain - Cerebellar Hemisphere | 440788 | -59212 | 940788 | Yes |
| ENSG00000120647.9 | CCDC77 | rs479803 | 8.70E-11 | ##### | Brain - Cerebellar Hemisphere | 440788 | -59212 | 940788 | Yes |
| ENSG00000120647.9 | CCDC77 | rs72472276 | 8.70E-11 | ##### | Brain - Cerebellar Hemisphere | 440788 | -59212 | 940788 | Yes |
| ENSG00000120647.9 | CCDC77 | rs4980877 | 8.70E-11 | ##### | Brain - Cerebellar Hemisphere | 440788 | -59212 | 940788 | Yes |
| ENSG00000120647.9 | CCDC77 | rs7963309 | 8.70E-11 | ##### | Brain - Cerebellar Hemisphere | 440788 | -59212 | 940788 | Yes |
| ENSG00000120647.9 | CCDC77 | rs7134353 | 8.70E-11 | ##### | Brain - Cerebellar Hemisphere | 440788 | -59212 | 940788 | Yes |
| ENSG00000120647.9 | CCDC77 | rs11374245 | 8.70E-11 | ##### | Brain - Cerebellar Hemisphere | 440788 | -59212 | 940788 | Yes |
| ENSG00000120647.9 | CCDC77 | rs16929744 | 8.70E-11 | ##### | Brain - Cerebellar Hemisphere | 440788 | -59212 | 940788 | Yes |
| ENSG00000120647.9 | CCDC77 | rs1207322 | 8.80E-11 | ##### | Brain - Hypothalamus | 440788 | -59212 | 940788 | Yes |
| ENSG00000120647.9 | CCDC77 | rs4980875 | 8.80E-11 | ##### | Brain - Hypothalamus | 440788 | -59212 | 940788 | Yes |
| ENSG00000120647.9 | CCDC77 | rs61906995 | 8.80E-11 | ##### | Brain - Hypothalamus | 440788 | -59212 | 940788 | Yes |

| | | | | | | | | | |
|---|---|---|---|---|---|---|---|---|---|
| ENSG00000120647.9 | CCDC77 | rs4980813 | 8.80E-11 | ##### | Brain - Hypothalamus | 440788 | -59212 | 940788 | Yes |
| ENSG00000120647.9 | CCDC77 | rs12582850 | 8.80E-11 | ##### | Brain - Hypothalamus | 440788 | -59212 | 940788 | Yes |
| ENSG00000120647.9 | CCDC77 | rs61906996 | 8.80E-11 | ##### | Brain - Hypothalamus | 440788 | -59212 | 940788 | Yes |
| ENSG00000120647.9 | CCDC77 | rs4980814 | 8.80E-11 | ##### | Brain - Hypothalamus | 440788 | -59212 | 940788 | Yes |
| ENSG00000120647.9 | CCDC77 | rs7298019 | 8.80E-11 | ##### | Brain - Hypothalamus | 440788 | -59212 | 940788 | Yes |
| ENSG00000120647.9 | CCDC77 | rs2098094 | 8.80E-11 | ##### | Brain - Hypothalamus | 440788 | -59212 | 940788 | Yes |
| ENSG00000120647.9 | CCDC77 | rs2159364 | 8.80E-11 | ##### | Brain - Hypothalamus | 440788 | -59212 | 940788 | Yes |
| ENSG00000120647.9 | CCDC77 | rs7977462 | 8.80E-11 | ##### | Brain - Hypothalamus | 440788 | -59212 | 940788 | Yes |
| ENSG00000120647.9 | CCDC77 | rs10744616 | 8.80E-11 | ##### | Brain - Hypothalamus | 440788 | -59212 | 940788 | Yes |
| ENSG00000120647.9 | CCDC77 | rs10848868 | 8.80E-11 | ##### | Brain - Hypothalamus | 440788 | -59212 | 940788 | Yes |
| ENSG00000120647.9 | CCDC77 | rs10774150 | 8.80E-11 | ##### | Brain - Hypothalamus | 440788 | -59212 | 940788 | Yes |
| ENSG00000120647.9 | CCDC77 | rs1860360 | 8.80E-11 | ##### | Brain - Hypothalamus | 440788 | -59212 | 940788 | Yes |
| ENSG00000120647.9 | CCDC77 | rs3831759 | 8.80E-11 | ##### | Brain - Hypothalamus | 440788 | -59212 | 940788 | Yes |
| ENSG00000120647.9 | CCDC77 | rs2284333 | 9.10E-11 | ##### | Brain - Caudate (basal ganglia | 440788 | -59212 | 940788 | Yes |
| ENSG00000120647.9 | CCDC77 | rs7963793 | 9.20E-11 | ##### | Brain - Cerebellar Hemisphere | 440788 | -59212 | 940788 | Yes |
| ENSG00000120647.9 | CCDC77 | rs10848814 | 9.20E-11 | ##### | Brain - Cerebellar Hemisphere | 440788 | -59212 | 940788 | Yes |
| ENSG00000120647.9 | CCDC77 | rs10848815 | 9.20E-11 | ##### | Brain - Cerebellar Hemisphere | 440788 | -59212 | 940788 | Yes |
| ENSG00000120647.9 | CCDC77 | rs887311 | 9.20E-11 | ##### | Brain - Cerebellar Hemisphere | 440788 | -59212 | 940788 | Yes |
| ENSG00000120647.9 | CCDC77 | rs7136116 | 9.90E-11 | ##### | Brain - Cerebellar Hemisphere | 440788 | -59212 | 940788 | Yes |
| ENSG00000120647.9 | CCDC77 | rs11610457 | 9.90E-11 | ##### | Brain - Cerebellar Hemisphere | 440788 | -59212 | 940788 | Yes |
| ENSG00000120647.9 | CCDC77 | rs3815558 | 1.00E-10 | ##### | Brain - Frontal Cortex (BA9) | 440788 | -59212 | 940788 | Yes |
| ENSG00000120647.9 | CCDC77 | rs78998485 | 1.00E-10 | ##### | Brain - Frontal Cortex (BA9) | 440788 | -59212 | 940788 | Yes |
| ENSG00000120647.9 | CCDC77 | rs7135768 | 1.00E-10 | ##### | Brain - Frontal Cortex (BA9) | 440788 | -59212 | 940788 | Yes |
| ENSG00000120647.9 | CCDC77 | rs6489450 | 1.00E-10 | ##### | Brain - Frontal Cortex (BA9) | 440788 | -59212 | 940788 | Yes |
| ENSG00000120647.9 | CCDC77 | rs2284333 | 1.10E-10 | ##### | Brain - Cerebellar Hemisphere | 440788 | -59212 | 940788 | Yes |
| ENSG00000120647.9 | CCDC77 | rs4980811 | 1.10E-10 | ##### | Brain - Cerebellar Hemisphere | 440788 | -59212 | 940788 | Yes |
| ENSG00000120647.9 | CCDC77 | rs3213751 | 1.10E-10 | ##### | Brain - Putamen (basal gangli | 440788 | -59212 | 940788 | Yes |
| ENSG00000120647.9 | CCDC77 | rs7963793 | 1.10E-10 | ##### | Brain - Caudate (basal ganglia | 440788 | -59212 | 940788 | Yes |
| ENSG00000120647.9 | CCDC77 | rs10848814 | 1.10E-10 | ##### | Brain - Caudate (basal ganglia | 440788 | -59212 | 940788 | Yes |
| ENSG00000120647.9 | CCDC77 | rs10848815 | 1.10E-10 | ##### | Brain - Caudate (basal ganglia | 440788 | -59212 | 940788 | Yes |
| ENSG00000120647.9 | CCDC77 | rs887311 | 1.10E-10 | ##### | Brain - Caudate (basal ganglia | 440788 | -59212 | 940788 | Yes |
| ENSG00000120647.9 | CCDC77 | rs2253565 | 1.10E-10 | ##### | Brain - Frontal Cortex (BA9) | 440788 | -59212 | 940788 | Yes |

| | | | | | | | | | |
|---|---|---|---|---|---|---|---|---|---|
| ENSG00000120647.9 | CCDC77 | rs495316 | 1.10E-10 | ##### | Brain - Spinal cord (cervical c- | 440788 | -59212 | 940788 | Yes |
| ENSG00000120647.9 | CCDC77 | rs576159 | 1.10E-10 | ##### | Brain - Caudate (basal ganglia | 440788 | -59212 | 940788 | Yes |
| ENSG00000120647.9 | CCDC77 | rs7965303 | 1.20E-10 | ##### | Brain - Caudate (basal ganglia | 440788 | -59212 | 940788 | Yes |
| ENSG00000120647.9 | CCDC77 | rs10848879 | 1.20E-10 | ##### | Brain - Caudate (basal ganglia | 440788 | -59212 | 940788 | Yes |
| ENSG00000120647.9 | CCDC77 | rs4980876 | 1.20E-10 | ##### | Brain - Cortex | 440788 | -59212 | 940788 | Yes |
| ENSG00000120647.9 | CCDC77 | rs1207324 | 1.20E-10 | ##### | Brain - Hypothalamus | 440788 | -59212 | 940788 | Yes |
| ENSG00000120647.9 | CCDC77 | rs2284332 | 1.30E-10 | ##### | Brain - Caudate (basal ganglia | 440788 | -59212 | 940788 | Yes |
| ENSG00000120647.9 | CCDC77 | rs2229351 | 1.30E-10 | ##### | Brain - Caudate (basal ganglia | 440788 | -59212 | 940788 | Yes |
| ENSG00000120647.9 | CCDC77 | rs4980884 | 1.40E-10 | ##### | Brain - Putamen (basal gangli | 440788 | -59212 | 940788 | Yes |
| ENSG00000120647.9 | CCDC77 | rs3753158 | 1.40E-10 | ##### | Brain - Frontal Cortex (BA9) | 440788 | -59212 | 940788 | Yes |
| ENSG00000120647.9 | CCDC77 | rs3815558 | 1.40E-10 | ##### | Brain - Putamen (basal gangli | 440788 | -59212 | 940788 | Yes |
| ENSG00000120647.9 | CCDC77 | rs78998485 | 1.40E-10 | ##### | Brain - Putamen (basal gangli | 440788 | -59212 | 940788 | Yes |
| ENSG00000120647.9 | CCDC77 | rs7135768 | 1.40E-10 | ##### | Brain - Putamen (basal gangli | 440788 | -59212 | 940788 | Yes |
| ENSG00000120647.9 | CCDC77 | rs6489450 | 1.40E-10 | ##### | Brain - Putamen (basal gangli | 440788 | -59212 | 940788 | Yes |
| ENSG00000120647.9 | CCDC77 | rs10082788 | 1.40E-10 | ##### | Brain - Frontal Cortex (BA9) | 440788 | -59212 | 940788 | Yes |
| ENSG00000120647.9 | CCDC77 | rs4980874 | 1.50E-10 | ##### | Brain - Cerebellar Hemisphere | 440788 | -59212 | 940788 | Yes |
| ENSG00000120647.9 | CCDC77 | rs4980892 | 1.50E-10 | ##### | Brain - Spinal cord (cervical c- | 440788 | -59212 | 940788 | Yes |
| ENSG00000120647.9 | CCDC77 | rs6489477 | 1.60E-10 | ##### | Brain - Cerebellar Hemisphere | 440788 | -59212 | 940788 | Yes |
| ENSG00000120647.9 | CCDC77 | rs6489478 | 1.60E-10 | ##### | Brain - Cerebellar Hemisphere | 440788 | -59212 | 940788 | Yes |
| ENSG00000120647.9 | CCDC77 | rs7132845 | 1.70E-10 | ##### | Brain - Caudate (basal ganglia | 440788 | -59212 | 940788 | Yes |
| ENSG00000120647.9 | CCDC77 | rs10744627 | 1.70E-10 | ##### | Brain - Cortex | 440788 | -59212 | 940788 | Yes |
| ENSG00000120647.9 | CCDC77 | rs576159 | 1.70E-10 | ##### | Brain - Hypothalamus | 440788 | -59212 | 940788 | Yes |
| ENSG00000120647.9 | CCDC77 | rs576159 | 1.70E-10 | ##### | Brain - Cerebellar Hemisphere | 440788 | -59212 | 940788 | Yes |
| ENSG00000120647.9 | CCDC77 | rs4980886 | 1.90E-10 | ##### | Brain - Frontal Cortex (BA9) | 440788 | -59212 | 940788 | Yes |
| ENSG00000120647.9 | CCDC77 | rs525381 | 1.90E-10 | ##### | Brain - Putamen (basal gangli | 440788 | -59212 | 940788 | Yes |
| ENSG00000120647.9 | CCDC77 | rs10848644 | 1.90E-10 | ##### | Brain - Caudate (basal ganglia | 440788 | -59212 | 940788 | Yes |
| ENSG00000120647.9 | CCDC77 | rs77074951 | 1.90E-10 | ##### | Brain - Cerebellar Hemisphere | 440788 | -59212 | 940788 | Yes |
| ENSG00000120647.9 | CCDC77 | rs495973 | 2.00E-10 | ##### | Brain - Spinal cord (cervical c- | 440788 | -59212 | 940788 | Yes |
| ENSG00000120647.9 | CCDC77 | rs6489477 | 2.10E-10 | ##### | Brain - Caudate (basal ganglia | 440788 | -59212 | 940788 | Yes |
| ENSG00000120647.9 | CCDC77 | rs6489478 | 2.10E-10 | ##### | Brain - Caudate (basal ganglia | 440788 | -59212 | 940788 | Yes |
| ENSG00000120647.9 | CCDC77 | rs2052279 | 2.20E-10 | ##### | Brain - Caudate (basal ganglia | 440788 | -59212 | 940788 | Yes |
| ENSG00000120647.9 | CCDC77 | rs78813329 | 2.20E-10 | ##### | Brain - Caudate (basal ganglia | 440788 | -59212 | 940788 | Yes |

| | | | | | | | | | |
|---|---|---|---|---|---|---|---|---|---|
| ENSG00000120647.9 | CCDC77 | rs3753157 | 2.20E-10 | ##### | Brain - Caudate (basal ganglia | 440788 | -59212 | 940788 | Yes |
| ENSG00000120647.9 | CCDC77 | rs4980883 | 2.20E-10 | ##### | Brain - Caudate (basal ganglia | 440788 | -59212 | 940788 | Yes |
| ENSG00000120647.9 | CCDC77 | rs137910445 | 2.20E-10 | ##### | Brain - Caudate (basal ganglia | 440788 | -59212 | 940788 | Yes |
| ENSG00000120647.9 | CCDC77 | rs11062537 | 2.20E-10 | ##### | Brain - Caudate (basal ganglia | 440788 | -59212 | 940788 | Yes |
| ENSG00000120647.9 | CCDC77 | rs715230 | 2.20E-10 | ##### | Brain - Caudate (basal ganglia | 440788 | -59212 | 940788 | Yes |
| ENSG00000120647.9 | CCDC77 | rs715229 | 2.20E-10 | ##### | Brain - Caudate (basal ganglia | 440788 | -59212 | 940788 | Yes |
| ENSG00000120647.9 | CCDC77 | rs715228 | 2.20E-10 | ##### | Brain - Caudate (basal ganglia | 440788 | -59212 | 940788 | Yes |
| ENSG00000120647.9 | CCDC77 | rs10848838 | 2.20E-10 | ##### | Brain - Caudate (basal ganglia | 440788 | -59212 | 940788 | Yes |
| ENSG00000120647.9 | CCDC77 | rs7132845 | 2.30E-10 | ##### | Brain - Frontal Cortex (BA9) | 440788 | -59212 | 940788 | Yes |
| ENSG00000120647.9 | CCDC77 | rs2052279 | 2.40E-10 | ##### | Brain - Frontal Cortex (BA9) | 440788 | -59212 | 940788 | Yes |
| ENSG00000120647.9 | CCDC77 | rs78813329 | 2.40E-10 | ##### | Brain - Frontal Cortex (BA9) | 440788 | -59212 | 940788 | Yes |
| ENSG00000120647.9 | CCDC77 | rs3753157 | 2.40E-10 | ##### | Brain - Frontal Cortex (BA9) | 440788 | -59212 | 940788 | Yes |
| ENSG00000120647.9 | CCDC77 | rs4980883 | 2.40E-10 | ##### | Brain - Frontal Cortex (BA9) | 440788 | -59212 | 940788 | Yes |
| ENSG00000120647.9 | CCDC77 | rs137910445 | 2.40E-10 | ##### | Brain - Frontal Cortex (BA9) | 440788 | -59212 | 940788 | Yes |
| ENSG00000120647.9 | CCDC77 | rs11062537 | 2.40E-10 | ##### | Brain - Frontal Cortex (BA9) | 440788 | -59212 | 940788 | Yes |
| ENSG00000120647.9 | CCDC77 | rs715230 | 2.40E-10 | ##### | Brain - Frontal Cortex (BA9) | 440788 | -59212 | 940788 | Yes |
| ENSG00000120647.9 | CCDC77 | rs715229 | 2.40E-10 | ##### | Brain - Frontal Cortex (BA9) | 440788 | -59212 | 940788 | Yes |
| ENSG00000120647.9 | CCDC77 | rs715228 | 2.40E-10 | ##### | Brain - Frontal Cortex (BA9) | 440788 | -59212 | 940788 | Yes |
| ENSG00000120647.9 | CCDC77 | rs2300133 | 2.40E-10 | ##### | Brain - Frontal Cortex (BA9) | 440788 | -59212 | 940788 | Yes |
| ENSG00000120647.9 | CCDC77 | rs10848838 | 2.40E-10 | ##### | Brain - Frontal Cortex (BA9) | 440788 | -59212 | 940788 | Yes |
| ENSG00000120647.9 | CCDC77 | rs1207322 | 2.50E-10 | ##### | Brain - Cortex | 440788 | -59212 | 940788 | Yes |
| ENSG00000120647.9 | CCDC77 | rs4980813 | 2.50E-10 | ##### | Brain - Cortex | 440788 | -59212 | 940788 | Yes |
| ENSG00000120647.9 | CCDC77 | rs61906996 | 2.50E-10 | ##### | Brain - Cortex | 440788 | -59212 | 940788 | Yes |
| ENSG00000120647.9 | CCDC77 | rs4980814 | 2.50E-10 | ##### | Brain - Cortex | 440788 | -59212 | 940788 | Yes |
| ENSG00000120647.9 | CCDC77 | rs7977462 | 2.50E-10 | ##### | Brain - Cortex | 440788 | -59212 | 940788 | Yes |
| ENSG00000120647.9 | CCDC77 | rs10744616 | 2.50E-10 | ##### | Brain - Cortex | 440788 | -59212 | 940788 | Yes |
| ENSG00000120647.9 | CCDC77 | rs10774150 | 2.50E-10 | ##### | Brain - Cortex | 440788 | -59212 | 940788 | Yes |
| ENSG00000120647.9 | CCDC77 | rs3831759 | 2.50E-10 | ##### | Brain - Cortex | 440788 | -59212 | 940788 | Yes |
| ENSG00000120647.9 | CCDC77 | rs7138958 | 2.50E-10 | ##### | Brain - Spinal cord (cervical c- | 440788 | -59212 | 940788 | Yes |
| ENSG00000120647.9 | CCDC77 | rs2007537 | 2.50E-10 | ##### | Brain - Spinal cord (cervical c- | 440788 | -59212 | 940788 | Yes |
| ENSG00000120647.9 | CCDC77 | rs1207322 | 2.50E-10 | ##### | Brain - Cerebellum | 440788 | -59212 | 940788 | Yes |
| ENSG00000120647.9 | CCDC77 | rs4980813 | 2.50E-10 | ##### | Brain - Cerebellum | 440788 | -59212 | 940788 | Yes |

| | | | | | | | | | |
|---|---|---|---|---|---|---|---|---|---|
| ENSG00000120647.9 | CCDC77 | rs61906996 | 2.50E-10 | ##### | Brain - Cerebellum | 440788 | -59212 | 940788 | Yes |
| ENSG00000120647.9 | CCDC77 | rs4980814 | 2.50E-10 | ##### | Brain - Cerebellum | 440788 | -59212 | 940788 | Yes |
| ENSG00000120647.9 | CCDC77 | rs7977462 | 2.50E-10 | ##### | Brain - Cerebellum | 440788 | -59212 | 940788 | Yes |
| ENSG00000120647.9 | CCDC77 | rs10744616 | 2.50E-10 | ##### | Brain - Cerebellum | 440788 | -59212 | 940788 | Yes |
| ENSG00000120647.9 | CCDC77 | rs10774150 | 2.50E-10 | ##### | Brain - Cerebellum | 440788 | -59212 | 940788 | Yes |
| ENSG00000120647.9 | CCDC77 | rs3831759 | 2.50E-10 | ##### | Brain - Cerebellum | 440788 | -59212 | 940788 | Yes |
| ENSG00000120647.9 | CCDC77 | rs6489477 | 2.60E-10 | ##### | Brain - Frontal Cortex (BA9) | 440788 | -59212 | 940788 | Yes |
| ENSG00000120647.9 | CCDC77 | rs6489478 | 2.60E-10 | ##### | Brain - Frontal Cortex (BA9) | 440788 | -59212 | 940788 | Yes |
| ENSG00000120647.9 | CCDC77 | rs7972566 | 2.80E-10 | ##### | Brain - Cortex | 440788 | -59212 | 940788 | Yes |
| ENSG00000120647.9 | CCDC77 | rs10774170 | 2.80E-10 | ##### | Brain - Cortex | 440788 | -59212 | 940788 | Yes |
| ENSG00000120647.9 | CCDC77 | rs12298843 | 2.80E-10 | ##### | Brain - Frontal Cortex (BA9) | 440788 | -59212 | 940788 | Yes |
| ENSG00000120647.9 | CCDC77 | rs4980822 | 2.80E-10 | ##### | Brain - Hypothalamus | 440788 | -59212 | 940788 | Yes |
| ENSG00000120647.9 | CCDC77 | rs3753156 | 2.90E-10 | ##### | Brain - Cerebellar Hemisphere | 440788 | -59212 | 940788 | Yes |
| ENSG00000120647.9 | CCDC77 | rs12581260 | 2.90E-10 | ##### | Brain - Cerebellar Hemisphere | 440788 | -59212 | 940788 | Yes |
| ENSG00000120647.9 | CCDC77 | rs16929783 | 2.90E-10 | ##### | Brain - Cerebellar Hemisphere | 440788 | -59212 | 940788 | Yes |
| ENSG00000120647.9 | CCDC77 | rs2300131 | 2.90E-10 | ##### | Brain - Caudate (basal ganglia | 440788 | -59212 | 940788 | Yes |
| ENSG00000120647.9 | CCDC77 | rs764614 | 3.00E-10 | ##### | Brain - Frontal Cortex (BA9) | 440788 | -59212 | 940788 | Yes |
| ENSG00000120647.9 | CCDC77 | rs7138981 | 3.00E-10 | ##### | Brain - Frontal Cortex (BA9) | 440788 | -59212 | 940788 | Yes |
| ENSG00000120647.9 | CCDC77 | rs9669094 | 3.00E-10 | ##### | Brain - Frontal Cortex (BA9) | 440788 | -59212 | 940788 | Yes |
| ENSG00000120647.9 | CCDC77 | rs7132763 | 3.00E-10 | ##### | Brain - Frontal Cortex (BA9) | 440788 | -59212 | 940788 | Yes |
| ENSG00000120647.9 | CCDC77 | rs2015554 | 3.00E-10 | ##### | Brain - Frontal Cortex (BA9) | 440788 | -59212 | 940788 | Yes |
| ENSG00000120647.9 | CCDC77 | rs113574411 | 3.00E-10 | ##### | Brain - Hypothalamus | 440788 | -59212 | 940788 | Yes |
| ENSG00000120647.9 | CCDC77 | rs479840 | 3.00E-10 | ##### | Brain - Hypothalamus | 440788 | -59212 | 940788 | Yes |
| ENSG00000120647.9 | CCDC77 | rs479803 | 3.00E-10 | ##### | Brain - Hypothalamus | 440788 | -59212 | 940788 | Yes |
| ENSG00000120647.9 | CCDC77 | rs4980811 | 3.00E-10 | ##### | Brain - Hypothalamus | 440788 | -59212 | 940788 | Yes |
| ENSG00000120647.9 | CCDC77 | rs72472276 | 3.00E-10 | ##### | Brain - Hypothalamus | 440788 | -59212 | 940788 | Yes |
| ENSG00000120647.9 | CCDC77 | rs4980877 | 3.00E-10 | ##### | Brain - Hypothalamus | 440788 | -59212 | 940788 | Yes |
| ENSG00000120647.9 | CCDC77 | rs7963309 | 3.00E-10 | ##### | Brain - Hypothalamus | 440788 | -59212 | 940788 | Yes |
| ENSG00000120647.9 | CCDC77 | rs7134353 | 3.00E-10 | ##### | Brain - Hypothalamus | 440788 | -59212 | 940788 | Yes |
| ENSG00000120647.9 | CCDC77 | rs11374245 | 3.00E-10 | ##### | Brain - Hypothalamus | 440788 | -59212 | 940788 | Yes |
| ENSG00000120647.9 | CCDC77 | rs16929744 | 3.00E-10 | ##### | Brain - Hypothalamus | 440788 | -59212 | 940788 | Yes |
| ENSG00000120647.9 | CCDC77 | rs1207324 | 3.10E-10 | ##### | Brain - Putamen (basal gangli | 440788 | -59212 | 940788 | Yes |

| | | | | | | | | | |
|---|---|---|---|---|---|---|---|---|---|
| ENSG00000120647.9 | CCDC77 | rs1207322 | 3.10E-10 | ##### | Brain - Spinal cord (cervical c- | 440788 | -59212 | 940788 | Yes |
| ENSG00000120647.9 | CCDC77 | rs57022939 | 3.10E-10 | ##### | Brain - Spinal cord (cervical c- | 440788 | -59212 | 940788 | Yes |
| ENSG00000120647.9 | CCDC77 | rs4980875 | 3.10E-10 | ##### | Brain - Spinal cord (cervical c- | 440788 | -59212 | 940788 | Yes |
| ENSG00000120647.9 | CCDC77 | rs61906995 | 3.10E-10 | ##### | Brain - Spinal cord (cervical c- | 440788 | -59212 | 940788 | Yes |
| ENSG00000120647.9 | CCDC77 | rs4980813 | 3.10E-10 | ##### | Brain - Spinal cord (cervical c- | 440788 | -59212 | 940788 | Yes |
| ENSG00000120647.9 | CCDC77 | rs4980876 | 3.10E-10 | ##### | Brain - Spinal cord (cervical c- | 440788 | -59212 | 940788 | Yes |
| ENSG00000120647.9 | CCDC77 | rs12582850 | 3.10E-10 | ##### | Brain - Spinal cord (cervical c- | 440788 | -59212 | 940788 | Yes |
| ENSG00000120647.9 | CCDC77 | rs61906996 | 3.10E-10 | ##### | Brain - Spinal cord (cervical c- | 440788 | -59212 | 940788 | Yes |
| ENSG00000120647.9 | CCDC77 | rs4980814 | 3.10E-10 | ##### | Brain - Spinal cord (cervical c- | 440788 | -59212 | 940788 | Yes |
| ENSG00000120647.9 | CCDC77 | rs7298019 | 3.10E-10 | ##### | Brain - Spinal cord (cervical c- | 440788 | -59212 | 940788 | Yes |
| ENSG00000120647.9 | CCDC77 | rs2098094 | 3.10E-10 | ##### | Brain - Spinal cord (cervical c- | 440788 | -59212 | 940788 | Yes |
| ENSG00000120647.9 | CCDC77 | rs2159364 | 3.10E-10 | ##### | Brain - Spinal cord (cervical c- | 440788 | -59212 | 940788 | Yes |
| ENSG00000120647.9 | CCDC77 | rs7977462 | 3.10E-10 | ##### | Brain - Spinal cord (cervical c- | 440788 | -59212 | 940788 | Yes |
| ENSG00000120647.9 | CCDC77 | rs10744616 | 3.10E-10 | ##### | Brain - Spinal cord (cervical c- | 440788 | -59212 | 940788 | Yes |
| ENSG00000120647.9 | CCDC77 | rs10848868 | 3.10E-10 | ##### | Brain - Spinal cord (cervical c- | 440788 | -59212 | 940788 | Yes |
| ENSG00000120647.9 | CCDC77 | rs10774150 | 3.10E-10 | ##### | Brain - Spinal cord (cervical c- | 440788 | -59212 | 940788 | Yes |
| ENSG00000120647.9 | CCDC77 | rs1860360 | 3.10E-10 | ##### | Brain - Spinal cord (cervical c- | 440788 | -59212 | 940788 | Yes |
| ENSG00000120647.9 | CCDC77 | rs3831759 | 3.10E-10 | ##### | Brain - Spinal cord (cervical c- | 440788 | -59212 | 940788 | Yes |
| ENSG00000120647.9 | CCDC77 | rs2300131 | 3.40E-10 | ##### | Brain - Frontal Cortex (BA9) | 440788 | -59212 | 940788 | Yes |
| ENSG00000120647.9 | CCDC77 | rs12298843 | 3.40E-10 | ##### | Brain - Caudate (basal ganglia | 440788 | -59212 | 940788 | Yes |
| ENSG00000120647.9 | CCDC77 | rs7138958 | 3.60E-10 | ##### | Brain - Frontal Cortex (BA9) | 440788 | -59212 | 940788 | Yes |
| ENSG00000120647.9 | CCDC77 | rs2007537 | 3.60E-10 | ##### | Brain - Frontal Cortex (BA9) | 440788 | -59212 | 940788 | Yes |
| ENSG00000120647.9 | CCDC77 | rs486098 | 3.60E-10 | ##### | Brain - Hypothalamus | 440788 | -59212 | 940788 | Yes |
| ENSG00000120647.9 | CCDC77 | rs2369280 | 3.70E-10 | ##### | Brain - Spinal cord (cervical c- | 440788 | -59212 | 940788 | Yes |
| ENSG00000120647.9 | CCDC77 | rs3213750 | 3.80E-10 | ##### | Brain - Cerebellar Hemisphere | 440788 | -59212 | 940788 | Yes |
| ENSG00000120647.9 | CCDC77 | rs4980881 | 3.80E-10 | ##### | Brain - Cerebellar Hemisphere | 440788 | -59212 | 940788 | Yes |
| ENSG00000120647.9 | CCDC77 | rs4980896 | 4.40E-10 | ##### | Brain - Cerebellar Hemisphere | 440788 | -59212 | 940788 | Yes |
| ENSG00000120647.9 | CCDC77 | rs10082788 | 4.50E-10 | ##### | Brain - Caudate (basal ganglia | 440788 | -59212 | 940788 | Yes |
| ENSG00000120647.9 | CCDC77 | rs11613357 | 4.50E-10 | ##### | Brain - Caudate (basal ganglia | 440788 | -59212 | 940788 | Yes |
| ENSG00000120647.9 | CCDC77 | rs2300133 | 4.80E-10 | ##### | Brain - Cerebellar Hemisphere | 440788 | -59212 | 940788 | Yes |
| ENSG00000120647.9 | CCDC77 | rs372437080 | 5.00E-10 | ##### | Brain - Cerebellum | 440788 | -59212 | 940788 | Yes |
| ENSG00000120647.9 | CCDC77 | rs12372234 | 5.40E-10 | ##### | Brain - Hypothalamus | 440788 | -59212 | 940788 | Yes |

| | | | | | | | | | |
|---|---|---|---|---|---|---|---|---|---|
| ENSG00000120647.9 | CCDC77 | rs2300133 | 5.40E-10 | ##### | Brain - Caudate (basal ganglia | 440788 | -59212 | 940788 | Yes |
| ENSG00000120647.9 | CCDC77 | rs10774152 | 5.50E-10 | ##### | Brain - Frontal Cortex (BA9) | 440788 | -59212 | 940788 | Yes |
| ENSG00000120647.9 | CCDC77 | rs10774153 | 5.50E-10 | ##### | Brain - Frontal Cortex (BA9) | 440788 | -59212 | 940788 | Yes |
| ENSG00000120647.9 | CCDC77 | rs7133796 | 5.80E-10 | ##### | Brain - Cerebellum | 440788 | -59212 | 940788 | Yes |
| ENSG00000120647.9 | CCDC77 | rs4980876 | 5.80E-10 | ##### | Brain - Cerebellum | 440788 | -59212 | 940788 | Yes |
| ENSG00000120647.9 | CCDC77 | rs7138958 | 5.90E-10 | ##### | Brain - Nucleus accumbens (b | 440788 | -59212 | 940788 | Yes |
| ENSG00000120647.9 | CCDC77 | rs2007537 | 5.90E-10 | ##### | Brain - Nucleus accumbens (b | 440788 | -59212 | 940788 | Yes |
| ENSG00000120647.9 | CCDC77 | rs7303180 | 6.00E-10 | ##### | Brain - Hypothalamus | 440788 | -59212 | 940788 | Yes |
| ENSG00000120647.9 | CCDC77 | rs57022939 | 6.00E-10 | ##### | Brain - Hypothalamus | 440788 | -59212 | 940788 | Yes |
| ENSG00000120647.9 | CCDC77 | rs482514 | 6.10E-10 | ##### | Brain - Spinal cord (cervical c- | 440788 | -59212 | 940788 | Yes |
| ENSG00000120647.9 | CCDC77 | rs7315852 | 6.10E-10 | ##### | Brain - Spinal cord (cervical c- | 440788 | -59212 | 940788 | Yes |
| ENSG00000120647.9 | CCDC77 | rs2284334 | 6.10E-10 | ##### | Brain - Spinal cord (cervical c- | 440788 | -59212 | 940788 | Yes |
| ENSG00000120647.9 | CCDC77 | rs720128 | 6.10E-10 | ##### | Brain - Spinal cord (cervical c- | 440788 | -59212 | 940788 | Yes |
| ENSG00000120647.9 | CCDC77 | rs9739950 | 6.10E-10 | ##### | Brain - Spinal cord (cervical c- | 440788 | -59212 | 940788 | Yes |
| ENSG00000120647.9 | CCDC77 | rs12311569 | 6.10E-10 | ##### | Brain - Spinal cord (cervical c- | 440788 | -59212 | 940788 | Yes |
| ENSG00000120647.9 | CCDC77 | rs10848877 | 6.10E-10 | ##### | Brain - Spinal cord (cervical c- | 440788 | -59212 | 940788 | Yes |
| ENSG00000120647.9 | CCDC77 | rs7136116 | 6.20E-10 | ##### | Brain - Hypothalamus | 440788 | -59212 | 940788 | Yes |
| ENSG00000120647.9 | CCDC77 | rs11610457 | 6.20E-10 | ##### | Brain - Hypothalamus | 440788 | -59212 | 940788 | Yes |
| ENSG00000120647.9 | CCDC77 | rs1541463 | 6.40E-10 | ##### | Brain - Putamen (basal gangli | 440788 | -59212 | 940788 | Yes |
| ENSG00000120647.9 | CCDC77 | rs10848905 | 6.40E-10 | ##### | Brain - Putamen (basal gangli | 440788 | -59212 | 940788 | Yes |
| ENSG00000120647.9 | CCDC77 | rs2369278 | 6.40E-10 | ##### | Brain - Putamen (basal gangli | 440788 | -59212 | 940788 | Yes |
| ENSG00000120647.9 | CCDC77 | rs10774163 | 6.40E-10 | ##### | Brain - Putamen (basal gangli | 440788 | -59212 | 940788 | Yes |
| ENSG00000120647.9 | CCDC77 | rs576159 | 6.50E-10 | ##### | Brain - Frontal Cortex (BA9) | 440788 | -59212 | 940788 | Yes |
| ENSG00000120647.9 | CCDC77 | rs4980892 | 6.50E-10 | ##### | Brain - Frontal Cortex (BA9) | 440788 | -59212 | 940788 | Yes |
| ENSG00000120647.9 | CCDC77 | rs10431354 | 6.90E-10 | ##### | Brain - Caudate (basal ganglia | 440788 | -59212 | 940788 | Yes |
| ENSG00000120647.9 | CCDC77 | rs3213751 | 6.90E-10 | ##### | Brain - Cerebellum | 440788 | -59212 | 940788 | Yes |
| ENSG00000120647.9 | CCDC77 | rs764614 | 7.30E-10 | ##### | Brain - Hypothalamus | 440788 | -59212 | 940788 | Yes |
| ENSG00000120647.9 | CCDC77 | rs7138981 | 7.30E-10 | ##### | Brain - Hypothalamus | 440788 | -59212 | 940788 | Yes |
| ENSG00000120647.9 | CCDC77 | rs9669094 | 7.30E-10 | ##### | Brain - Hypothalamus | 440788 | -59212 | 940788 | Yes |
| ENSG00000120647.9 | CCDC77 | rs7132763 | 7.30E-10 | ##### | Brain - Hypothalamus | 440788 | -59212 | 940788 | Yes |
| ENSG00000120647.9 | CCDC77 | rs2015554 | 7.30E-10 | ##### | Brain - Hypothalamus | 440788 | -59212 | 940788 | Yes |
| ENSG00000120647.9 | CCDC77 | rs576994 | 7.70E-10 | ##### | Brain - Spinal cord (cervical c- | 440788 | -59212 | 940788 | Yes |

| | | | | | | | | | |
|---|---|---|---|---|---|---|---|---|---|
| ENSG00000120647.9 | CCDC77 | rs797759 | 7.70E-10 | ##### | Brain - Spinal cord (cervical c- | 440788 | -59212 | 940788 | Yes |
| ENSG00000120647.9 | CCDC77 | rs479836 | 7.70E-10 | ##### | Brain - Spinal cord (cervical c- | 440788 | -59212 | 940788 | Yes |
| ENSG00000120647.9 | CCDC77 | rs2284332 | 7.70E-10 | ##### | Brain - Spinal cord (cervical c- | 440788 | -59212 | 940788 | Yes |
| ENSG00000120647.9 | CCDC77 | rs2229351 | 7.70E-10 | ##### | Brain - Spinal cord (cervical c- | 440788 | -59212 | 940788 | Yes |
| ENSG00000120647.9 | CCDC77 | rs4980810 | 7.70E-10 | ##### | Brain - Spinal cord (cervical c- | 440788 | -59212 | 940788 | Yes |
| ENSG00000120647.9 | CCDC77 | rs11062302 | 7.70E-10 | ##### | Brain - Spinal cord (cervical c- | 440788 | -59212 | 940788 | Yes |
| ENSG00000120647.9 | CCDC77 | rs12314497 | 7.70E-10 | ##### | Brain - Spinal cord (cervical c- | 440788 | -59212 | 940788 | Yes |
| ENSG00000120647.9 | CCDC77 | rs534925791 | 7.70E-10 | ##### | Brain - Spinal cord (cervical c- | 440788 | -59212 | 940788 | Yes |
| ENSG00000120647.9 | CCDC77 | rs144729439 | 7.70E-10 | ##### | Brain - Spinal cord (cervical c- | 440788 | -59212 | 940788 | Yes |
| ENSG00000120647.9 | CCDC77 | rs11062385 | 7.70E-10 | ##### | Brain - Spinal cord (cervical c- | 440788 | -59212 | 940788 | Yes |
| ENSG00000120647.9 | CCDC77 | rs11611810 | 7.70E-10 | ##### | Brain - Spinal cord (cervical c- | 440788 | -59212 | 940788 | Yes |
| ENSG00000120647.9 | CCDC77 | rs10848715 | 7.70E-10 | ##### | Brain - Spinal cord (cervical c- | 440788 | -59212 | 940788 | Yes |
| ENSG00000120647.9 | CCDC77 | rs10848716 | 7.70E-10 | ##### | Brain - Spinal cord (cervical c- | 440788 | -59212 | 940788 | Yes |
| ENSG00000120647.9 | CCDC77 | rs61907027 | 7.70E-10 | ##### | Brain - Spinal cord (cervical c- | 440788 | -59212 | 940788 | Yes |
| ENSG00000120647.9 | CCDC77 | rs7301422 | 7.70E-10 | ##### | Brain - Spinal cord (cervical c- | 440788 | -59212 | 940788 | Yes |
| ENSG00000120647.9 | CCDC77 | rs11062512 | 7.70E-10 | ##### | Brain - Spinal cord (cervical c- | 440788 | -59212 | 940788 | Yes |
| ENSG00000120647.9 | CCDC77 | rs3759371 | 7.70E-10 | ##### | Brain - Spinal cord (cervical c- | 440788 | -59212 | 940788 | Yes |
| ENSG00000120647.9 | CCDC77 | rs6489448 | 7.70E-10 | ##### | Brain - Spinal cord (cervical c- | 440788 | -59212 | 940788 | Yes |
| ENSG00000120647.9 | CCDC77 | rs7316660 | 7.70E-10 | ##### | Brain - Spinal cord (cervical c- | 440788 | -59212 | 940788 | Yes |
| ENSG00000120647.9 | CCDC77 | rs6489449 | 7.70E-10 | ##### | Brain - Spinal cord (cervical c- | 440788 | -59212 | 940788 | Yes |
| ENSG00000120647.9 | CCDC77 | rs10735021 | 7.70E-10 | ##### | Brain - Spinal cord (cervical c- | 440788 | -59212 | 940788 | Yes |
| ENSG00000120647.9 | CCDC77 | rs10848845 | 7.70E-10 | ##### | Brain - Spinal cord (cervical c- | 440788 | -59212 | 940788 | Yes |
| ENSG00000120647.9 | CCDC77 | rs10774137 | 7.70E-10 | ##### | Brain - Spinal cord (cervical c- | 440788 | -59212 | 940788 | Yes |
| ENSG00000120647.9 | CCDC77 | rs11835325 | 7.70E-10 | ##### | Brain - Spinal cord (cervical c- | 440788 | -59212 | 940788 | Yes |
| ENSG00000120647.9 | CCDC77 | rs10848849 | 7.70E-10 | ##### | Brain - Spinal cord (cervical c- | 440788 | -59212 | 940788 | Yes |
| ENSG00000120647.9 | CCDC77 | rs6489454 | 7.70E-10 | ##### | Brain - Spinal cord (cervical c- | 440788 | -59212 | 940788 | Yes |
| ENSG00000120647.9 | CCDC77 | rs7134608 | 7.70E-10 | ##### | Brain - Spinal cord (cervical c- | 440788 | -59212 | 940788 | Yes |
| ENSG00000120647.9 | CCDC77 | rs7138890 | 7.70E-10 | ##### | Brain - Spinal cord (cervical c- | 440788 | -59212 | 940788 | Yes |
| ENSG00000120647.9 | CCDC77 | rs10431355 | 7.70E-10 | ##### | Brain - Spinal cord (cervical c- | 440788 | -59212 | 940788 | Yes |
| ENSG00000120647.9 | CCDC77 | rs6489458 | 7.70E-10 | ##### | Brain - Spinal cord (cervical c- | 440788 | -59212 | 940788 | Yes |
| ENSG00000120647.9 | CCDC77 | rs2300134 | 7.70E-10 | ##### | Brain - Spinal cord (cervical c- | 440788 | -59212 | 940788 | Yes |
| ENSG00000120647.9 | CCDC77 | rs3782852 | 7.70E-10 | ##### | Brain - Spinal cord (cervical c- | 440788 | -59212 | 940788 | Yes |

| | | | | | | | | | |
|---|---|---|---|---|---|---|---|---|---|
| ENSG00000120647.9 | CCDC77 | rs201124968 | 7.70E-10 | ##### | Brain - Spinal cord (cervical c- | 440788 | -59212 | 940788 | Yes |
| ENSG00000120647.9 | CCDC77 | rs201946814 | 7.70E-10 | ##### | Brain - Spinal cord (cervical c- | 440788 | -59212 | 940788 | Yes |
| ENSG00000120647.9 | CCDC77 | rs6489463 | 7.70E-10 | ##### | Brain - Spinal cord (cervical c- | 440788 | -59212 | 940788 | Yes |
| ENSG00000120647.9 | CCDC77 | rs2079785 | 7.70E-10 | ##### | Brain - Spinal cord (cervical c- | 440788 | -59212 | 940788 | Yes |
| ENSG00000120647.9 | CCDC77 | rs7963937 | 7.70E-10 | ##### | Brain - Spinal cord (cervical c- | 440788 | -59212 | 940788 | Yes |
| ENSG00000120647.9 | CCDC77 | rs2369276 | 7.70E-10 | ##### | Brain - Spinal cord (cervical c- | 440788 | -59212 | 940788 | Yes |
| ENSG00000120647.9 | CCDC77 | rs7953137 | 7.70E-10 | ##### | Brain - Spinal cord (cervical c- | 440788 | -59212 | 940788 | Yes |
| ENSG00000120647.9 | CCDC77 | rs6489471 | 7.70E-10 | ##### | Brain - Spinal cord (cervical c- | 440788 | -59212 | 940788 | Yes |
| ENSG00000120647.9 | CCDC77 | rs7312065 | 7.70E-10 | ##### | Brain - Spinal cord (cervical c- | 440788 | -59212 | 940788 | Yes |
| ENSG00000120647.9 | CCDC77 | rs7136957 | 7.70E-10 | ##### | Brain - Spinal cord (cervical c- | 440788 | -59212 | 940788 | Yes |
| ENSG00000120647.9 | CCDC77 | rs34380840 | 7.70E-10 | ##### | Brain - Spinal cord (cervical c- | 440788 | -59212 | 940788 | Yes |
| ENSG00000120647.9 | CCDC77 | rs6489477 | 7.70E-10 | ##### | Brain - Spinal cord (cervical c- | 440788 | -59212 | 940788 | Yes |
| ENSG00000120647.9 | CCDC77 | rs6489478 | 7.70E-10 | ##### | Brain - Spinal cord (cervical c- | 440788 | -59212 | 940788 | Yes |
| ENSG00000120647.9 | CCDC77 | rs7965303 | 7.70E-10 | ##### | Brain - Spinal cord (cervical c- | 440788 | -59212 | 940788 | Yes |
| ENSG00000120647.9 | CCDC77 | rs10848879 | 7.70E-10 | ##### | Brain - Spinal cord (cervical c- | 440788 | -59212 | 940788 | Yes |
| ENSG00000120647.9 | CCDC77 | rs12301816 | 7.80E-10 | ##### | Brain - Nucleus accumbens (b | 440788 | -59212 | 940788 | Yes |
| ENSG00000120647.9 | CCDC77 | rs55657100 | 8.00E-10 | ##### | Brain - Hypothalamus | 440788 | -59212 | 940788 | Yes |
| ENSG00000120647.9 | CCDC77 | rs10774152 | 8.20E-10 | ##### | Brain - Hypothalamus | 440788 | -59212 | 940788 | Yes |
| ENSG00000120647.9 | CCDC77 | rs10774153 | 8.20E-10 | ##### | Brain - Hypothalamus | 440788 | -59212 | 940788 | Yes |
| ENSG00000120647.9 | CCDC77 | rs495973 | 8.30E-10 | ##### | Brain - Nucleus accumbens (b | 440788 | -59212 | 940788 | Yes |
| ENSG00000120647.9 | CCDC77 | rs10774169 | 8.60E-10 | ##### | Brain - Spinal cord (cervical c- | 440788 | -59212 | 940788 | Yes |
| ENSG00000120647.9 | CCDC77 | rs1207324 | 9.10E-10 | ##### | Brain - Cerebellum | 440788 | -59212 | 940788 | Yes |
| ENSG00000120647.9 | CCDC77 | rs486098 | 9.90E-10 | ##### | Brain - Putamen (basal gangli | 440788 | -59212 | 940788 | Yes |
| ENSG00000120647.9 | CCDC77 | rs482514 | 1.00E-09 | ##### | Brain - Hypothalamus | 440788 | -59212 | 940788 | Yes |
| ENSG00000120647.9 | CCDC77 | rs7315852 | 1.00E-09 | ##### | Brain - Hypothalamus | 440788 | -59212 | 940788 | Yes |
| ENSG00000120647.9 | CCDC77 | rs2284334 | 1.00E-09 | ##### | Brain - Hypothalamus | 440788 | -59212 | 940788 | Yes |
| ENSG00000120647.9 | CCDC77 | rs720128 | 1.00E-09 | ##### | Brain - Hypothalamus | 440788 | -59212 | 940788 | Yes |
| ENSG00000120647.9 | CCDC77 | rs9739950 | 1.00E-09 | ##### | Brain - Hypothalamus | 440788 | -59212 | 940788 | Yes |
| ENSG00000120647.9 | CCDC77 | rs12311569 | 1.00E-09 | ##### | Brain - Hypothalamus | 440788 | -59212 | 940788 | Yes |
| ENSG00000120647.9 | CCDC77 | rs525381 | 1.00E-09 | ##### | Brain - Spinal cord (cervical c- | 440788 | -59212 | 940788 | Yes |
| ENSG00000120647.9 | CCDC77 | rs486098 | 1.10E-09 | ##### | Brain - Frontal Cortex (BA9) | 440788 | -59212 | 940788 | Yes |
| ENSG00000120647.9 | CCDC77 | rs1155942 | 1.10E-09 | ##### | Brain - Cortex | 440788 | -59212 | 940788 | Yes |

| | | | | | | | | | |
|---|---|---|---|---|---|---|---|---|---|
| ENSG00000120647.9 | CCDC77 | rs10082788 | 1.10E-09 | ##### | Brain - Hypothalamus | 440788 | -59212 | 940788 | Yes |
| ENSG00000120647.9 | CCDC77 | rs57022939 | 1.20E-09 | ##### | Brain - Cerebellum | 440788 | -59212 | 940788 | Yes |
| ENSG00000120647.9 | CCDC77 | rs2253565 | 1.20E-09 | ##### | Brain - Cortex | 440788 | -59212 | 940788 | Yes |
| ENSG00000120647.9 | CCDC77 | rs10848644 | 1.20E-09 | ##### | Brain - Frontal Cortex (BA9) | 440788 | -59212 | 940788 | Yes |
| ENSG00000120647.9 | CCDC77 | rs764614 | 1.30E-09 | ##### | Brain - Spinal cord (cervical c- | 440788 | -59212 | 940788 | Yes |
| ENSG00000120647.9 | CCDC77 | rs10848877 | 1.30E-09 | ##### | Brain - Hypothalamus | 440788 | -59212 | 940788 | Yes |
| ENSG00000120647.9 | CCDC77 | rs6489523 | 1.30E-09 | ##### | Brain - Hypothalamus | 440788 | -59212 | 940788 | Yes |
| ENSG00000120647.9 | CCDC77 | rs524468 | 1.30E-09 | ##### | Brain - Nucleus accumbens (b | 440788 | -59212 | 940788 | Yes |
| ENSG00000120647.9 | CCDC77 | rs10774192 | 1.40E-09 | ##### | Brain - Frontal Cortex (BA9) | 440788 | -59212 | 940788 | Yes |
| ENSG00000120647.9 | CCDC77 | rs113574411 | 1.40E-09 | ##### | Brain - Frontal Cortex (BA9) | 440788 | -59212 | 940788 | Yes |
| ENSG00000120647.9 | CCDC77 | rs479840 | 1.40E-09 | ##### | Brain - Frontal Cortex (BA9) | 440788 | -59212 | 940788 | Yes |
| ENSG00000120647.9 | CCDC77 | rs479803 | 1.40E-09 | ##### | Brain - Frontal Cortex (BA9) | 440788 | -59212 | 940788 | Yes |
| ENSG00000120647.9 | CCDC77 | rs72472276 | 1.40E-09 | ##### | Brain - Frontal Cortex (BA9) | 440788 | -59212 | 940788 | Yes |
| ENSG00000120647.9 | CCDC77 | rs4980877 | 1.40E-09 | ##### | Brain - Frontal Cortex (BA9) | 440788 | -59212 | 940788 | Yes |
| ENSG00000120647.9 | CCDC77 | rs7963309 | 1.40E-09 | ##### | Brain - Frontal Cortex (BA9) | 440788 | -59212 | 940788 | Yes |
| ENSG00000120647.9 | CCDC77 | rs7134353 | 1.40E-09 | ##### | Brain - Frontal Cortex (BA9) | 440788 | -59212 | 940788 | Yes |
| ENSG00000120647.9 | CCDC77 | rs11374245 | 1.40E-09 | ##### | Brain - Frontal Cortex (BA9) | 440788 | -59212 | 940788 | Yes |
| ENSG00000120647.9 | CCDC77 | rs7303180 | 1.40E-09 | ##### | Brain - Frontal Cortex (BA9) | 440788 | -59212 | 940788 | Yes |
| ENSG00000120647.9 | CCDC77 | rs16929744 | 1.40E-09 | ##### | Brain - Frontal Cortex (BA9) | 440788 | -59212 | 940788 | Yes |
| ENSG00000120647.9 | CCDC77 | rs10161265 | 1.50E-09 | ##### | Brain - Caudate (basal ganglia | 440788 | -59212 | 940788 | Yes |
| ENSG00000120647.9 | CCDC77 | rs4980897 | 1.50E-09 | ##### | Brain - Caudate (basal ganglia | 440788 | -59212 | 940788 | Yes |
| ENSG00000120647.9 | CCDC77 | rs486098 | 1.50E-09 | ##### | Brain - Spinal cord (cervical c- | 440788 | -59212 | 940788 | Yes |
| ENSG00000120647.9 | CCDC77 | rs525381 | 1.80E-09 | ##### | Brain - Hypothalamus | 440788 | -59212 | 940788 | Yes |
| ENSG00000120647.9 | CCDC77 | rs2369280 | 1.80E-09 | ##### | Brain - Putamen (basal gangli | 440788 | -59212 | 940788 | Yes |
| ENSG00000120647.9 | CCDC77 | rs77074951 | 1.80E-09 | ##### | Brain - Hypothalamus | 440788 | -59212 | 940788 | Yes |
| ENSG00000120647.9 | CCDC77 | rs7311740 | 1.90E-09 | ##### | Brain - Hypothalamus | 440788 | -59212 | 940788 | Yes |
| ENSG00000120647.9 | CCDC77 | rs4980898 | 1.90E-09 | ##### | Brain - Caudate (basal ganglia | 440788 | -59212 | 940788 | Yes |
| ENSG00000120647.9 | CCDC77 | rs7963793 | 1.90E-09 | ##### | Brain - Frontal Cortex (BA9) | 440788 | -59212 | 940788 | Yes |
| ENSG00000120647.9 | CCDC77 | rs10848814 | 1.90E-09 | ##### | Brain - Frontal Cortex (BA9) | 440788 | -59212 | 940788 | Yes |
| ENSG00000120647.9 | CCDC77 | rs10848815 | 1.90E-09 | ##### | Brain - Frontal Cortex (BA9) | 440788 | -59212 | 940788 | Yes |
| ENSG00000120647.9 | CCDC77 | rs887311 | 1.90E-09 | ##### | Brain - Frontal Cortex (BA9) | 440788 | -59212 | 940788 | Yes |
| ENSG00000120647.9 | CCDC77 | rs6489477 | 1.90E-09 | ##### | Brain - Hypothalamus | 440788 | -59212 | 940788 | Yes |

| | | | | | | | | | |
|---|---|---|---|---|---|---|---|---|---|
| ENSG00000120647.9 | CCDC77 | rs6489478 | 1.90E-09 | ##### | Brain - Hypothalamus | 440788 | -59212 | 940788 | Yes |
| ENSG00000120647.9 | CCDC77 | rs7965303 | 1.90E-09 | ##### | Brain - Hypothalamus | 440788 | -59212 | 940788 | Yes |
| ENSG00000120647.9 | CCDC77 | rs10848879 | 1.90E-09 | ##### | Brain - Hypothalamus | 440788 | -59212 | 940788 | Yes |
| ENSG00000120647.9 | CCDC77 | rs4482086 | 2.00E-09 | ##### | Brain - Cerebellum | 440788 | -59212 | 940788 | Yes |
| ENSG00000120647.9 | CCDC77 | rs2369279 | 2.00E-09 | ##### | Brain - Cerebellum | 440788 | -59212 | 940788 | Yes |
| ENSG00000120647.9 | CCDC77 | rs576994 | 2.00E-09 | ##### | Brain - Hypothalamus | 440788 | -59212 | 940788 | Yes |
| ENSG00000120647.9 | CCDC77 | rs797759 | 2.00E-09 | ##### | Brain - Hypothalamus | 440788 | -59212 | 940788 | Yes |
| ENSG00000120647.9 | CCDC77 | rs479836 | 2.00E-09 | ##### | Brain - Hypothalamus | 440788 | -59212 | 940788 | Yes |
| ENSG00000120647.9 | CCDC77 | rs4980810 | 2.00E-09 | ##### | Brain - Hypothalamus | 440788 | -59212 | 940788 | Yes |
| ENSG00000120647.9 | CCDC77 | rs11062302 | 2.00E-09 | ##### | Brain - Hypothalamus | 440788 | -59212 | 940788 | Yes |
| ENSG00000120647.9 | CCDC77 | rs12314497 | 2.00E-09 | ##### | Brain - Hypothalamus | 440788 | -59212 | 940788 | Yes |
| ENSG00000120647.9 | CCDC77 | rs534925791 | 2.00E-09 | ##### | Brain - Hypothalamus | 440788 | -59212 | 940788 | Yes |
| ENSG00000120647.9 | CCDC77 | rs144729439 | 2.00E-09 | ##### | Brain - Hypothalamus | 440788 | -59212 | 940788 | Yes |
| ENSG00000120647.9 | CCDC77 | rs11062385 | 2.00E-09 | ##### | Brain - Hypothalamus | 440788 | -59212 | 940788 | Yes |
| ENSG00000120647.9 | CCDC77 | rs11611810 | 2.00E-09 | ##### | Brain - Hypothalamus | 440788 | -59212 | 940788 | Yes |
| ENSG00000120647.9 | CCDC77 | rs10848715 | 2.00E-09 | ##### | Brain - Hypothalamus | 440788 | -59212 | 940788 | Yes |
| ENSG00000120647.9 | CCDC77 | rs10848716 | 2.00E-09 | ##### | Brain - Hypothalamus | 440788 | -59212 | 940788 | Yes |
| ENSG00000120647.9 | CCDC77 | rs61907027 | 2.00E-09 | ##### | Brain - Hypothalamus | 440788 | -59212 | 940788 | Yes |
| ENSG00000120647.9 | CCDC77 | rs7301422 | 2.00E-09 | ##### | Brain - Hypothalamus | 440788 | -59212 | 940788 | Yes |
| ENSG00000120647.9 | CCDC77 | rs11062512 | 2.00E-09 | ##### | Brain - Hypothalamus | 440788 | -59212 | 940788 | Yes |
| ENSG00000120647.9 | CCDC77 | rs3759371 | 2.00E-09 | ##### | Brain - Hypothalamus | 440788 | -59212 | 940788 | Yes |
| ENSG00000120647.9 | CCDC77 | rs6489448 | 2.00E-09 | ##### | Brain - Hypothalamus | 440788 | -59212 | 940788 | Yes |
| ENSG00000120647.9 | CCDC77 | rs7316660 | 2.00E-09 | ##### | Brain - Hypothalamus | 440788 | -59212 | 940788 | Yes |
| ENSG00000120647.9 | CCDC77 | rs6489449 | 2.00E-09 | ##### | Brain - Hypothalamus | 440788 | -59212 | 940788 | Yes |
| ENSG00000120647.9 | CCDC77 | rs10735021 | 2.00E-09 | ##### | Brain - Hypothalamus | 440788 | -59212 | 940788 | Yes |
| ENSG00000120647.9 | CCDC77 | rs10848845 | 2.00E-09 | ##### | Brain - Hypothalamus | 440788 | -59212 | 940788 | Yes |
| ENSG00000120647.9 | CCDC77 | rs10774137 | 2.00E-09 | ##### | Brain - Hypothalamus | 440788 | -59212 | 940788 | Yes |
| ENSG00000120647.9 | CCDC77 | rs11835325 | 2.00E-09 | ##### | Brain - Hypothalamus | 440788 | -59212 | 940788 | Yes |
| ENSG00000120647.9 | CCDC77 | rs10848849 | 2.00E-09 | ##### | Brain - Hypothalamus | 440788 | -59212 | 940788 | Yes |
| ENSG00000120647.9 | CCDC77 | rs6489454 | 2.00E-09 | ##### | Brain - Hypothalamus | 440788 | -59212 | 940788 | Yes |
| ENSG00000120647.9 | CCDC77 | rs7134608 | 2.00E-09 | ##### | Brain - Hypothalamus | 440788 | -59212 | 940788 | Yes |
| ENSG00000120647.9 | CCDC77 | rs7138890 | 2.00E-09 | ##### | Brain - Hypothalamus | 440788 | -59212 | 940788 | Yes |

| | | | | | | | | | |
|---|---|---|---|---|---|---|---|---|---|
| ENSG00000120647.9 | CCDC77 | rs10431355 | 2.00E-09 | ##### | Brain - Hypothalamus | 440788 | -59212 | 940788 | Yes |
| ENSG00000120647.9 | CCDC77 | rs6489458 | 2.00E-09 | ##### | Brain - Hypothalamus | 440788 | -59212 | 940788 | Yes |
| ENSG00000120647.9 | CCDC77 | rs2300134 | 2.00E-09 | ##### | Brain - Hypothalamus | 440788 | -59212 | 940788 | Yes |
| ENSG00000120647.9 | CCDC77 | rs3782852 | 2.00E-09 | ##### | Brain - Hypothalamus | 440788 | -59212 | 940788 | Yes |
| ENSG00000120647.9 | CCDC77 | rs201124968 | 2.00E-09 | ##### | Brain - Hypothalamus | 440788 | -59212 | 940788 | Yes |
| ENSG00000120647.9 | CCDC77 | rs201946814 | 2.00E-09 | ##### | Brain - Hypothalamus | 440788 | -59212 | 940788 | Yes |
| ENSG00000120647.9 | CCDC77 | rs6489463 | 2.00E-09 | ##### | Brain - Hypothalamus | 440788 | -59212 | 940788 | Yes |
| ENSG00000120647.9 | CCDC77 | rs2079785 | 2.00E-09 | ##### | Brain - Hypothalamus | 440788 | -59212 | 940788 | Yes |
| ENSG00000120647.9 | CCDC77 | rs7963937 | 2.00E-09 | ##### | Brain - Hypothalamus | 440788 | -59212 | 940788 | Yes |
| ENSG00000120647.9 | CCDC77 | rs2369276 | 2.00E-09 | ##### | Brain - Hypothalamus | 440788 | -59212 | 940788 | Yes |
| ENSG00000120647.9 | CCDC77 | rs7953137 | 2.00E-09 | ##### | Brain - Hypothalamus | 440788 | -59212 | 940788 | Yes |
| ENSG00000120647.9 | CCDC77 | rs6489471 | 2.00E-09 | ##### | Brain - Hypothalamus | 440788 | -59212 | 940788 | Yes |
| ENSG00000120647.9 | CCDC77 | rs7312065 | 2.00E-09 | ##### | Brain - Hypothalamus | 440788 | -59212 | 940788 | Yes |
| ENSG00000120647.9 | CCDC77 | rs34380840 | 2.00E-09 | ##### | Brain - Hypothalamus | 440788 | -59212 | 940788 | Yes |
| ENSG00000120647.9 | CCDC77 | rs495973 | 2.00E-09 | ##### | Brain - Putamen (basal gangli | 440788 | -59212 | 940788 | Yes |
| ENSG00000120647.9 | CCDC77 | rs7138958 | 2.00E-09 | ##### | Brain - Putamen (basal gangli | 440788 | -59212 | 940788 | Yes |
| ENSG00000120647.9 | CCDC77 | rs2007537 | 2.00E-09 | ##### | Brain - Putamen (basal gangli | 440788 | -59212 | 940788 | Yes |
| ENSG00000120647.9 | CCDC77 | rs61907026 | 2.10E-09 | ##### | Brain - Spinal cord (cervical c- | 440788 | -59212 | 940788 | Yes |
| ENSG00000120647.9 | CCDC77 | rs79839498 | 2.10E-09 | ##### | Brain - Spinal cord (cervical c- | 440788 | -59212 | 940788 | Yes |
| ENSG00000120647.9 | CCDC77 | rs61907030 | 2.10E-09 | ##### | Brain - Spinal cord (cervical c- | 440788 | -59212 | 940788 | Yes |
| ENSG00000120647.9 | CCDC77 | rs61907031 | 2.10E-09 | ##### | Brain - Spinal cord (cervical c- | 440788 | -59212 | 940788 | Yes |
| ENSG00000120647.9 | CCDC77 | rs11615564 | 2.10E-09 | ##### | Brain - Spinal cord (cervical c- | 440788 | -59212 | 940788 | Yes |
| ENSG00000120647.9 | CCDC77 | rs3213751 | 2.20E-09 | ##### | Brain - Cortex | 440788 | -59212 | 940788 | Yes |
| ENSG00000120647.9 | CCDC77 | rs7295011 | 2.20E-09 | ##### | Brain - Hypothalamus | 440788 | -59212 | 940788 | Yes |
| ENSG00000120647.9 | CCDC77 | rs1207324 | 2.20E-09 | ##### | Brain - Spinal cord (cervical c- | 440788 | -59212 | 940788 | Yes |
| ENSG00000120647.9 | CCDC77 | rs17800720 | 2.30E-09 | ##### | Brain - Caudate (basal ganglia | 440788 | -59212 | 940788 | Yes |
| ENSG00000120647.9 | CCDC77 | rs3837515 | 2.30E-09 | ##### | Brain - Caudate (basal ganglia | 440788 | -59212 | 940788 | Yes |
| ENSG00000120647.9 | CCDC77 | rs4980892 | 2.30E-09 | ##### | Brain - Putamen (basal gangli | 440788 | -59212 | 940788 | Yes |
| ENSG00000120647.9 | CCDC77 | rs77074951 | 2.30E-09 | ##### | Brain - Frontal Cortex (BA9) | 440788 | -59212 | 940788 | Yes |
| ENSG00000120647.9 | CCDC77 | rs17800720 | 2.40E-09 | ##### | Brain - Frontal Cortex (BA9) | 440788 | -59212 | 940788 | Yes |
| ENSG00000120647.9 | CCDC77 | rs3837515 | 2.40E-09 | ##### | Brain - Frontal Cortex (BA9) | 440788 | -59212 | 940788 | Yes |
| ENSG00000120647.9 | CCDC77 | rs2284333 | 2.40E-09 | ##### | Brain - Hypothalamus | 440788 | -59212 | 940788 | Yes |

| | | | | | | | | | |
|---|---|---|---|---|---|---|---|---|---|
| ENSG00000120647.9 | CCDC77 | rs10160995 | 2.50E-09 | ##### | Brain - Caudate (basal ganglia | 440788 | -59212 | 940788 | Yes |
| ENSG00000120647.9 | CCDC77 | rs10082788 | 2.50E-09 | ##### | Brain - Cerebellar Hemisphere | 440788 | -59212 | 940788 | Yes |
| ENSG00000120647.9 | CCDC77 | rs2369278 | 2.60E-09 | ##### | Brain - Nucleus accumbens (b | 440788 | -59212 | 940788 | Yes |
| ENSG00000120647.9 | CCDC77 | rs10774163 | 2.60E-09 | ##### | Brain - Nucleus accumbens (b | 440788 | -59212 | 940788 | Yes |
| ENSG00000120647.9 | CCDC77 | rs520932 | 2.80E-09 | ##### | Brain - Nucleus accumbens (b | 440788 | -59212 | 940788 | Yes |
| ENSG00000120647.9 | CCDC77 | rs11062813 | 2.90E-09 | ##### | Brain - Cortex | 440788 | -59212 | 940788 | Yes |
| ENSG00000120647.9 | CCDC77 | rs7132763 | 3.00E-09 | ##### | Brain - Spinal cord (cervical c- | 440788 | -59212 | 940788 | Yes |
| ENSG00000120647.9 | CCDC77 | rs7136957 | 3.10E-09 | ##### | Brain - Hypothalamus | 440788 | -59212 | 940788 | Yes |
| ENSG00000120647.9 | CCDC77 | rs28850111 | 3.10E-09 | ##### | Brain - Caudate (basal ganglia | 440788 | -59212 | 940788 | Yes |
| ENSG00000120647.9 | CCDC77 | rs7138958 | 3.20E-09 | ##### | Brain - Cortex | 440788 | -59212 | 940788 | Yes |
| ENSG00000120647.9 | CCDC77 | rs2007537 | 3.20E-09 | ##### | Brain - Cortex | 440788 | -59212 | 940788 | Yes |
| ENSG00000120647.9 | CCDC77 | rs2080403 | 3.20E-09 | ##### | Brain - Frontal Cortex (BA9) | 440788 | -59212 | 940788 | Yes |
| ENSG00000120647.9 | CCDC77 | rs2289957 | 3.20E-09 | ##### | Brain - Frontal Cortex (BA9) | 440788 | -59212 | 940788 | Yes |
| ENSG00000120647.9 | CCDC77 | rs201124968 | 3.20E-09 | ##### | Brain - Putamen (basal gangli | 440788 | -59212 | 940788 | Yes |
| ENSG00000120647.9 | CCDC77 | rs10848987 | 3.40E-09 | ##### | Brain - Frontal Cortex (BA9) | 440788 | -59212 | 940788 | Yes |
| ENSG00000120647.9 | CCDC77 | rs111250603 | 3.40E-09 | ##### | Brain - Cerebellum | 440788 | -59212 | 940788 | Yes |
| ENSG00000120647.9 | CCDC77 | rs2369277 | 3.40E-09 | ##### | Brain - Cerebellum | 440788 | -59212 | 940788 | Yes |
| ENSG00000120647.9 | CCDC77 | rs9805067 | 3.40E-09 | ##### | Brain - Cerebellum | 440788 | -59212 | 940788 | Yes |
| ENSG00000120647.9 | CCDC77 | rs1541464 | 3.40E-09 | ##### | Brain - Cerebellum | 440788 | -59212 | 940788 | Yes |
| ENSG00000120647.9 | CCDC77 | rs1541465 | 3.40E-09 | ##### | Brain - Cerebellum | 440788 | -59212 | 940788 | Yes |
| ENSG00000120647.9 | CCDC77 | rs4980817 | 3.40E-09 | ##### | Brain - Cerebellum | 440788 | -59212 | 940788 | Yes |
| ENSG00000120647.9 | CCDC77 | rs11062167 | 3.40E-09 | ##### | Brain - Caudate (basal ganglia | 440788 | -59212 | 940788 | Yes |
| ENSG00000120647.9 | CCDC77 | rs55990424 | 3.40E-09 | ##### | Brain - Caudate (basal ganglia | 440788 | -59212 | 940788 | Yes |
| ENSG00000120647.9 | CCDC77 | rs11062167 | 3.80E-09 | ##### | Brain - Frontal Cortex (BA9) | 440788 | -59212 | 940788 | Yes |
| ENSG00000120647.9 | CCDC77 | rs55990424 | 3.80E-09 | ##### | Brain - Frontal Cortex (BA9) | 440788 | -59212 | 940788 | Yes |
| ENSG00000120647.9 | CCDC77 | rs10774169 | 3.80E-09 | ##### | Brain - Nucleus accumbens (b | 440788 | -59212 | 940788 | Yes |
| ENSG00000120647.9 | CCDC77 | rs520932 | 3.90E-09 | ##### | Brain - Putamen (basal gangli | 440788 | -59212 | 940788 | Yes |
| ENSG00000120647.9 | CCDC77 | rs55657100 | 3.90E-09 | ##### | Brain - Frontal Cortex (BA9) | 440788 | -59212 | 940788 | Yes |
| ENSG00000120647.9 | CCDC77 | rs4980884 | 4.00E-09 | ##### | Brain - Cerebellum | 440788 | -59212 | 940788 | Yes |
| ENSG00000120647.9 | CCDC77 | rs4980875 | 4.20E-09 | ##### | Brain - Cerebellum | 440788 | -59212 | 940788 | Yes |
| ENSG00000120647.9 | CCDC77 | rs61906995 | 4.20E-09 | ##### | Brain - Cerebellum | 440788 | -59212 | 940788 | Yes |
| ENSG00000120647.9 | CCDC77 | rs12582850 | 4.20E-09 | ##### | Brain - Cerebellum | 440788 | -59212 | 940788 | Yes |

| | | | | | | | | | |
|---|---|---|---|---|---|---|---|---|---|
| ENSG00000120647.9 | CCDC77 | rs7298019 | 4.20E-09 | ##### | Brain - Cerebellum | 440788 | -59212 | 940788 | Yes |
| ENSG00000120647.9 | CCDC77 | rs2098094 | 4.20E-09 | ##### | Brain - Cerebellum | 440788 | -59212 | 940788 | Yes |
| ENSG00000120647.9 | CCDC77 | rs2159364 | 4.20E-09 | ##### | Brain - Cerebellum | 440788 | -59212 | 940788 | Yes |
| ENSG00000120647.9 | CCDC77 | rs10848868 | 4.20E-09 | ##### | Brain - Cerebellum | 440788 | -59212 | 940788 | Yes |
| ENSG00000120647.9 | CCDC77 | rs1860360 | 4.20E-09 | ##### | Brain - Cerebellum | 440788 | -59212 | 940788 | Yes |
| ENSG00000120647.9 | CCDC77 | rs4980874 | 4.20E-09 | ##### | Brain - Frontal Cortex (BA9) | 440788 | -59212 | 940788 | Yes |
| ENSG00000120647.9 | CCDC77 | rs4980878 | 4.20E-09 | ##### | Brain - Cerebellum | 440788 | -59212 | 940788 | Yes |
| ENSG00000120647.9 | CCDC77 | rs2041299 | 4.50E-09 | ##### | Brain - Cortex | 440788 | -59212 | 940788 | Yes |
| ENSG00000120647.9 | CCDC77 | rs4980886 | 4.50E-09 | ##### | Brain - Hypothalamus | 440788 | -59212 | 940788 | Yes |
| ENSG00000120647.9 | CCDC77 | rs4980811 | 4.60E-09 | ##### | Brain - Frontal Cortex (BA9) | 440788 | -59212 | 940788 | Yes |
| ENSG00000120647.9 | CCDC77 | rs10848905 | 4.70E-09 | ##### | Brain - Nucleus accumbens (b | 440788 | -59212 | 940788 | Yes |
| ENSG00000120647.9 | CCDC77 | rs4980874 | 5.00E-09 | ##### | Brain - Hypothalamus | 440788 | -59212 | 940788 | Yes |
| ENSG00000120647.9 | CCDC77 | rs2191159 | 5.20E-09 | ##### | Brain - Hypothalamus | 440788 | -59212 | 940788 | Yes |
| ENSG00000120647.9 | CCDC77 | rs10848647 | 5.40E-09 | ##### | Brain - Cerebellar Hemisphere | 440788 | -59212 | 940788 | Yes |
| ENSG00000120647.9 | CCDC77 | rs10848650 | 5.40E-09 | ##### | Brain - Cerebellar Hemisphere | 440788 | -59212 | 940788 | Yes |
| ENSG00000120647.9 | CCDC77 | rs4980896 | 5.40E-09 | ##### | Brain - Nucleus accumbens (b | 440788 | -59212 | 940788 | Yes |
| ENSG00000120647.9 | CCDC77 | rs10848987 | 5.50E-09 | ##### | Brain - Hypothalamus | 440788 | -59212 | 940788 | Yes |
| ENSG00000120647.9 | CCDC77 | rs11063005 | 5.60E-09 | ##### | Brain - Frontal Cortex (BA9) | 440788 | -59212 | 940788 | Yes |
| ENSG00000120647.9 | CCDC77 | rs11063010 | 5.60E-09 | ##### | Brain - Frontal Cortex (BA9) | 440788 | -59212 | 940788 | Yes |
| ENSG00000120647.9 | CCDC77 | rs797765 | 5.70E-09 | ##### | Brain - Putamen (basal gangli | 440788 | -59212 | 940788 | Yes |
| ENSG00000120647.9 | CCDC77 | rs1541463 | 6.00E-09 | ##### | Brain - Cortex | 440788 | -59212 | 940788 | Yes |
| ENSG00000120647.9 | CCDC77 | rs1541463 | 6.10E-09 | ##### | Brain - Nucleus accumbens (b | 440788 | -59212 | 940788 | Yes |
| ENSG00000120647.9 | CCDC77 | rs6489523 | 6.20E-09 | ##### | Brain - Frontal Cortex (BA9) | 440788 | -59212 | 940788 | Yes |
| ENSG00000120647.9 | CCDC77 | rs10774169 | 6.30E-09 | ##### | Brain - Putamen (basal gangli | 440788 | -59212 | 940788 | Yes |
| ENSG00000120647.9 | CCDC77 | rs67523949 | 6.40E-09 | ##### | Brain - Frontal Cortex (BA9) | 440788 | -59212 | 940788 | Yes |
| ENSG00000120647.9 | CCDC77 | rs1548904 | 7.10E-09 | ##### | Brain - Caudate (basal ganglia | 440788 | -59212 | 940788 | Yes |
| ENSG00000120647.9 | CCDC77 | rs1972582 | 7.10E-09 | ##### | Brain - Caudate (basal ganglia | 440788 | -59212 | 940788 | Yes |
| ENSG00000120647.9 | CCDC77 | rs10848647 | 7.10E-09 | ##### | Brain - Caudate (basal ganglia | 440788 | -59212 | 940788 | Yes |
| ENSG00000120647.9 | CCDC77 | rs10774038 | 7.10E-09 | ##### | Brain - Caudate (basal ganglia | 440788 | -59212 | 940788 | Yes |
| ENSG00000120647.9 | CCDC77 | rs10848766 | 7.20E-09 | ##### | Brain - Cerebellar Hemisphere | 440788 | -59212 | 940788 | Yes |
| ENSG00000120647.9 | CCDC77 | rs10744561 | 7.20E-09 | ##### | Brain - Caudate (basal ganglia | 440788 | -59212 | 940788 | Yes |
| ENSG00000120647.9 | CCDC77 | rs10848650 | 7.20E-09 | ##### | Brain - Caudate (basal ganglia | 440788 | -59212 | 940788 | Yes |

| | | | | | | | | | |
|---|---|---|---|---|---|---|---|---|---|
| ENSG00000120647.9 | CCDC77 | rs7972566 | 7.40E-09 | ##### | Brain - Cerebellum | 440788 | -59212 | 940788 | Yes |
| ENSG00000120647.9 | CCDC77 | rs2284333 | 7.60E-09 | ##### | Brain - Frontal Cortex (BA9) | 440788 | -59212 | 940788 | Yes |
| ENSG00000120647.9 | CCDC77 | rs2253565 | 8.10E-09 | ##### | Brain - Cerebellum | 440788 | -59212 | 940788 | Yes |
| ENSG00000120647.9 | CCDC77 | rs10848877 | 8.20E-09 | ##### | Brain - Nucleus accumbens (b | 440788 | -59212 | 940788 | Yes |
| ENSG00000120647.9 | CCDC77 | rs3753156 | 8.30E-09 | ##### | Brain - Frontal Cortex (BA9) | 440788 | -59212 | 940788 | Yes |
| ENSG00000120647.9 | CCDC77 | rs12581260 | 8.30E-09 | ##### | Brain - Frontal Cortex (BA9) | 440788 | -59212 | 940788 | Yes |
| ENSG00000120647.9 | CCDC77 | rs16929783 | 8.30E-09 | ##### | Brain - Frontal Cortex (BA9) | 440788 | -59212 | 940788 | Yes |
| ENSG00000120647.9 | CCDC77 | rs113574411 | 8.50E-09 | ##### | Brain - Spinal cord (cervical c- | 440788 | -59212 | 940788 | Yes |
| ENSG00000120647.9 | CCDC77 | rs479840 | 8.50E-09 | ##### | Brain - Spinal cord (cervical c- | 440788 | -59212 | 940788 | Yes |
| ENSG00000120647.9 | CCDC77 | rs479803 | 8.50E-09 | ##### | Brain - Spinal cord (cervical c- | 440788 | -59212 | 940788 | Yes |
| ENSG00000120647.9 | CCDC77 | rs2284333 | 8.50E-09 | ##### | Brain - Spinal cord (cervical c- | 440788 | -59212 | 940788 | Yes |
| ENSG00000120647.9 | CCDC77 | rs4980811 | 8.50E-09 | ##### | Brain - Spinal cord (cervical c- | 440788 | -59212 | 940788 | Yes |
| ENSG00000120647.9 | CCDC77 | rs72472276 | 8.50E-09 | ##### | Brain - Spinal cord (cervical c- | 440788 | -59212 | 940788 | Yes |
| ENSG00000120647.9 | CCDC77 | rs4980877 | 8.50E-09 | ##### | Brain - Spinal cord (cervical c- | 440788 | -59212 | 940788 | Yes |
| ENSG00000120647.9 | CCDC77 | rs7963309 | 8.50E-09 | ##### | Brain - Spinal cord (cervical c- | 440788 | -59212 | 940788 | Yes |
| ENSG00000120647.9 | CCDC77 | rs7134353 | 8.50E-09 | ##### | Brain - Spinal cord (cervical c- | 440788 | -59212 | 940788 | Yes |
| ENSG00000120647.9 | CCDC77 | rs11374245 | 8.50E-09 | ##### | Brain - Spinal cord (cervical c- | 440788 | -59212 | 940788 | Yes |
| ENSG00000120647.9 | CCDC77 | rs7303180 | 8.50E-09 | ##### | Brain - Spinal cord (cervical c- | 440788 | -59212 | 940788 | Yes |
| ENSG00000120647.9 | CCDC77 | rs16929744 | 8.50E-09 | ##### | Brain - Spinal cord (cervical c- | 440788 | -59212 | 940788 | Yes |
| ENSG00000120647.9 | CCDC77 | rs7138981 | 8.50E-09 | ##### | Brain - Spinal cord (cervical c- | 440788 | -59212 | 940788 | Yes |
| ENSG00000120647.9 | CCDC77 | rs9669094 | 8.50E-09 | ##### | Brain - Spinal cord (cervical c- | 440788 | -59212 | 940788 | Yes |
| ENSG00000120647.9 | CCDC77 | rs2015554 | 8.50E-09 | ##### | Brain - Spinal cord (cervical c- | 440788 | -59212 | 940788 | Yes |
| ENSG00000120647.9 | CCDC77 | rs10774152 | 8.50E-09 | ##### | Brain - Spinal cord (cervical c- | 440788 | -59212 | 940788 | Yes |
| ENSG00000120647.9 | CCDC77 | rs10774153 | 8.50E-09 | ##### | Brain - Spinal cord (cervical c- | 440788 | -59212 | 940788 | Yes |
| ENSG00000120647.9 | CCDC77 | rs10774169 | 8.90E-09 | ##### | Brain - Frontal Cortex (BA9) | 440788 | -59212 | 940788 | Yes |
| ENSG00000120647.9 | CCDC77 | rs797765 | 9.10E-09 | ##### | Brain - Cortex | 440788 | -59212 | 940788 | Yes |
| ENSG00000120647.9 | CCDC77 | rs11062934 | 9.30E-09 | ##### | Brain - Caudate (basal ganglia | 440788 | -59212 | 940788 | Yes |
| ENSG00000120647.9 | CCDC77 | rs12372234 | 9.30E-09 | ##### | Brain - Frontal Cortex (BA9) | 440788 | -59212 | 940788 | Yes |
| ENSG00000120647.9 | CCDC77 | rs3213751 | 9.60E-09 | ##### | Brain - Hypothalamus | 440788 | -59212 | 940788 | Yes |
| ENSG00000120647.9 | CCDC77 | rs3815558 | 9.60E-09 | ##### | Brain - Hypothalamus | 440788 | -59212 | 940788 | Yes |
| ENSG00000120647.9 | CCDC77 | rs78998485 | 9.60E-09 | ##### | Brain - Hypothalamus | 440788 | -59212 | 940788 | Yes |
| ENSG00000120647.9 | CCDC77 | rs4980884 | 9.60E-09 | ##### | Brain - Hypothalamus | 440788 | -59212 | 940788 | Yes |

| | | | | | | | | | |
|---|---|---|---|---|---|---|---|---|---|
| ENSG00000120647.9 | CCDC77 | rs7135768 | 9.60E-09 | ##### | Brain - Hypothalamus | 440788 | -59212 | 940788 | Yes |
| ENSG00000120647.9 | CCDC77 | rs6489450 | 9.60E-09 | ##### | Brain - Hypothalamus | 440788 | -59212 | 940788 | Yes |
| ENSG00000120647.9 | CCDC77 | rs482514 | 9.60E-09 | ##### | Brain - Nucleus accumbens (b | 440788 | -59212 | 940788 | Yes |
| ENSG00000120647.9 | CCDC77 | rs7315852 | 9.60E-09 | ##### | Brain - Nucleus accumbens (b | 440788 | -59212 | 940788 | Yes |
| ENSG00000120647.9 | CCDC77 | rs2284334 | 9.60E-09 | ##### | Brain - Nucleus accumbens (b | 440788 | -59212 | 940788 | Yes |
| ENSG00000120647.9 | CCDC77 | rs720128 | 9.60E-09 | ##### | Brain - Nucleus accumbens (b | 440788 | -59212 | 940788 | Yes |
| ENSG00000120647.9 | CCDC77 | rs9739950 | 9.60E-09 | ##### | Brain - Nucleus accumbens (b | 440788 | -59212 | 940788 | Yes |
| ENSG00000120647.9 | CCDC77 | rs12311569 | 9.60E-09 | ##### | Brain - Nucleus accumbens (b | 440788 | -59212 | 940788 | Yes |
| ENSG00000120647.9 | CCDC77 | rs2253565 | 9.80E-09 | ##### | Brain - Nucleus accumbens (b | 440788 | -59212 | 940788 | Yes |
| ENSG00000120647.9 | CCDC77 | rs77074951 | 9.90E-09 | ##### | Brain - Spinal cord (cervical c- | 440788 | -59212 | 940788 | Yes |
| ENSG00000120647.9 | CCDC77 | rs3815558 | 1.00E-08 | ##### | Brain - Cerebellum | 440788 | -59212 | 940788 | Yes |
| ENSG00000120647.9 | CCDC77 | rs78998485 | 1.00E-08 | ##### | Brain - Cerebellum | 440788 | -59212 | 940788 | Yes |
| ENSG00000120647.9 | CCDC77 | rs7135768 | 1.00E-08 | ##### | Brain - Cerebellum | 440788 | -59212 | 940788 | Yes |
| ENSG00000120647.9 | CCDC77 | rs6489450 | 1.00E-08 | ##### | Brain - Cerebellum | 440788 | -59212 | 940788 | Yes |
| ENSG00000120647.9 | CCDC77 | rs4482086 | 1.00E-08 | ##### | Brain - Nucleus accumbens (b | 440788 | -59212 | 940788 | Yes |
| ENSG00000120647.9 | CCDC77 | rs2369279 | 1.00E-08 | ##### | Brain - Nucleus accumbens (b | 440788 | -59212 | 940788 | Yes |
| ENSG00000120647.9 | CCDC77 | rs11062803 | 1.00E-08 | ##### | Brain - Nucleus accumbens (b | 440788 | -59212 | 940788 | Yes |
| ENSG00000120647.9 | CCDC77 | rs2300131 | 1.00E-08 | ##### | Brain - Putamen (basal gangli | 440788 | -59212 | 940788 | Yes |
| ENSG00000120647.9 | CCDC77 | rs10848905 | 1.00E-08 | ##### | Brain - Cortex | 440788 | -59212 | 940788 | Yes |
| ENSG00000120647.9 | CCDC77 | rs2369278 | 1.00E-08 | ##### | Brain - Cortex | 440788 | -59212 | 940788 | Yes |
| ENSG00000120647.9 | CCDC77 | rs10774163 | 1.00E-08 | ##### | Brain - Cortex | 440788 | -59212 | 940788 | Yes |
| ENSG00000120647.9 | CCDC77 | rs10431354 | 1.10E-08 | ##### | Brain - Spinal cord (cervical c- | 440788 | -59212 | 940788 | Yes |
| ENSG00000120647.9 | CCDC77 | rs7295011 | 1.10E-08 | ##### | Brain - Frontal Cortex (BA9) | 440788 | -59212 | 940788 | Yes |
| ENSG00000120647.9 | CCDC77 | rs12301816 | 1.10E-08 | ##### | Brain - Frontal Cortex (BA9) | 440788 | -59212 | 940788 | Yes |
| ENSG00000120647.9 | CCDC77 | rs487252 | 1.20E-08 | ##### | Brain - Cerebellar Hemisphere | 440788 | -59212 | 940788 | Yes |
| ENSG00000120647.9 | CCDC77 | rs4980822 | 1.20E-08 | ##### | Brain - Frontal Cortex (BA9) | 440788 | -59212 | 940788 | Yes |
| ENSG00000120647.9 | CCDC77 | rs201124968 | 1.20E-08 | ##### | Brain - Cortex | 440788 | -59212 | 940788 | Yes |
| ENSG00000120647.9 | CCDC77 | rs3782860 | 1.20E-08 | ##### | Brain - Caudate (basal ganglia | 440788 | -59212 | 940788 | Yes |
| ENSG00000120647.9 | CCDC77 | rs524468 | 1.20E-08 | ##### | Brain - Putamen (basal gangli | 440788 | -59212 | 940788 | Yes |
| ENSG00000120647.9 | CCDC77 | rs11062803 | 1.20E-08 | ##### | Brain - Cerebellum | 440788 | -59212 | 940788 | Yes |
| ENSG00000120647.9 | CCDC77 | rs12229673 | 1.20E-08 | ##### | Brain - Caudate (basal ganglia | 440788 | -59212 | 940788 | Yes |
| ENSG00000120647.9 | CCDC77 | rs12231564 | 1.20E-08 | ##### | Brain - Caudate (basal ganglia | 440788 | -59212 | 940788 | Yes |

| | | | | | | | | | |
|---|---|---|---|---|---|---|---|---|---|
| ENSG00000120647.9 | CCDC77 | rs576994 | 1.30E-08 | ##### | Brain - Putamen (basal gangli | 440788 | -59212 | 940788 | Yes |
| ENSG00000120647.9 | CCDC77 | rs797759 | 1.30E-08 | ##### | Brain - Putamen (basal gangli | 440788 | -59212 | 940788 | Yes |
| ENSG00000120647.9 | CCDC77 | rs479836 | 1.30E-08 | ##### | Brain - Putamen (basal gangli | 440788 | -59212 | 940788 | Yes |
| ENSG00000120647.9 | CCDC77 | rs2284332 | 1.30E-08 | ##### | Brain - Putamen (basal gangli | 440788 | -59212 | 940788 | Yes |
| ENSG00000120647.9 | CCDC77 | rs2229351 | 1.30E-08 | ##### | Brain - Putamen (basal gangli | 440788 | -59212 | 940788 | Yes |
| ENSG00000120647.9 | CCDC77 | rs4980810 | 1.30E-08 | ##### | Brain - Putamen (basal gangli | 440788 | -59212 | 940788 | Yes |
| ENSG00000120647.9 | CCDC77 | rs11062302 | 1.30E-08 | ##### | Brain - Putamen (basal gangli | 440788 | -59212 | 940788 | Yes |
| ENSG00000120647.9 | CCDC77 | rs12314497 | 1.30E-08 | ##### | Brain - Putamen (basal gangli | 440788 | -59212 | 940788 | Yes |
| ENSG00000120647.9 | CCDC77 | rs534925791 | 1.30E-08 | ##### | Brain - Putamen (basal gangli | 440788 | -59212 | 940788 | Yes |
| ENSG00000120647.9 | CCDC77 | rs144729439 | 1.30E-08 | ##### | Brain - Putamen (basal gangli | 440788 | -59212 | 940788 | Yes |
| ENSG00000120647.9 | CCDC77 | rs11062385 | 1.30E-08 | ##### | Brain - Putamen (basal gangli | 440788 | -59212 | 940788 | Yes |
| ENSG00000120647.9 | CCDC77 | rs11611810 | 1.30E-08 | ##### | Brain - Putamen (basal gangli | 440788 | -59212 | 940788 | Yes |
| ENSG00000120647.9 | CCDC77 | rs10848715 | 1.30E-08 | ##### | Brain - Putamen (basal gangli | 440788 | -59212 | 940788 | Yes |
| ENSG00000120647.9 | CCDC77 | rs10848716 | 1.30E-08 | ##### | Brain - Putamen (basal gangli | 440788 | -59212 | 940788 | Yes |
| ENSG00000120647.9 | CCDC77 | rs61907027 | 1.30E-08 | ##### | Brain - Putamen (basal gangli | 440788 | -59212 | 940788 | Yes |
| ENSG00000120647.9 | CCDC77 | rs7301422 | 1.30E-08 | ##### | Brain - Putamen (basal gangli | 440788 | -59212 | 940788 | Yes |
| ENSG00000120647.9 | CCDC77 | rs11062512 | 1.30E-08 | ##### | Brain - Putamen (basal gangli | 440788 | -59212 | 940788 | Yes |
| ENSG00000120647.9 | CCDC77 | rs3759371 | 1.30E-08 | ##### | Brain - Putamen (basal gangli | 440788 | -59212 | 940788 | Yes |
| ENSG00000120647.9 | CCDC77 | rs6489448 | 1.30E-08 | ##### | Brain - Putamen (basal gangli | 440788 | -59212 | 940788 | Yes |
| ENSG00000120647.9 | CCDC77 | rs7316660 | 1.30E-08 | ##### | Brain - Putamen (basal gangli | 440788 | -59212 | 940788 | Yes |
| ENSG00000120647.9 | CCDC77 | rs6489449 | 1.30E-08 | ##### | Brain - Putamen (basal gangli | 440788 | -59212 | 940788 | Yes |
| ENSG00000120647.9 | CCDC77 | rs10735021 | 1.30E-08 | ##### | Brain - Putamen (basal gangli | 440788 | -59212 | 940788 | Yes |
| ENSG00000120647.9 | CCDC77 | rs10848845 | 1.30E-08 | ##### | Brain - Putamen (basal gangli | 440788 | -59212 | 940788 | Yes |
| ENSG00000120647.9 | CCDC77 | rs10774137 | 1.30E-08 | ##### | Brain - Putamen (basal gangli | 440788 | -59212 | 940788 | Yes |
| ENSG00000120647.9 | CCDC77 | rs11835325 | 1.30E-08 | ##### | Brain - Putamen (basal gangli | 440788 | -59212 | 940788 | Yes |
| ENSG00000120647.9 | CCDC77 | rs10848849 | 1.30E-08 | ##### | Brain - Putamen (basal gangli | 440788 | -59212 | 940788 | Yes |
| ENSG00000120647.9 | CCDC77 | rs6489454 | 1.30E-08 | ##### | Brain - Putamen (basal gangli | 440788 | -59212 | 940788 | Yes |
| ENSG00000120647.9 | CCDC77 | rs7134608 | 1.30E-08 | ##### | Brain - Putamen (basal gangli | 440788 | -59212 | 940788 | Yes |
| ENSG00000120647.9 | CCDC77 | rs7138890 | 1.30E-08 | ##### | Brain - Putamen (basal gangli | 440788 | -59212 | 940788 | Yes |
| ENSG00000120647.9 | CCDC77 | rs10431355 | 1.30E-08 | ##### | Brain - Putamen (basal gangli | 440788 | -59212 | 940788 | Yes |
| ENSG00000120647.9 | CCDC77 | rs6489458 | 1.30E-08 | ##### | Brain - Putamen (basal gangli | 440788 | -59212 | 940788 | Yes |
| ENSG00000120647.9 | CCDC77 | rs2300134 | 1.30E-08 | ##### | Brain - Putamen (basal gangli | 440788 | -59212 | 940788 | Yes |

| | | | | | | | | | |
|---|---|---|---|---|---|---|---|---|---|
| ENSG00000120647.9 | CCDC77 | rs3782852 | 1.30E-08 | ##### | Brain - Putamen (basal gangli | 440788 | -59212 | 940788 | Yes |
| ENSG00000120647.9 | CCDC77 | rs201946814 | 1.30E-08 | ##### | Brain - Putamen (basal gangli | 440788 | -59212 | 940788 | Yes |
| ENSG00000120647.9 | CCDC77 | rs6489463 | 1.30E-08 | ##### | Brain - Putamen (basal gangli | 440788 | -59212 | 940788 | Yes |
| ENSG00000120647.9 | CCDC77 | rs2079785 | 1.30E-08 | ##### | Brain - Putamen (basal gangli | 440788 | -59212 | 940788 | Yes |
| ENSG00000120647.9 | CCDC77 | rs7963937 | 1.30E-08 | ##### | Brain - Putamen (basal gangli | 440788 | -59212 | 940788 | Yes |
| ENSG00000120647.9 | CCDC77 | rs2369276 | 1.30E-08 | ##### | Brain - Putamen (basal gangli | 440788 | -59212 | 940788 | Yes |
| ENSG00000120647.9 | CCDC77 | rs7953137 | 1.30E-08 | ##### | Brain - Putamen (basal gangli | 440788 | -59212 | 940788 | Yes |
| ENSG00000120647.9 | CCDC77 | rs6489471 | 1.30E-08 | ##### | Brain - Putamen (basal gangli | 440788 | -59212 | 940788 | Yes |
| ENSG00000120647.9 | CCDC77 | rs7312065 | 1.30E-08 | ##### | Brain - Putamen (basal gangli | 440788 | -59212 | 940788 | Yes |
| ENSG00000120647.9 | CCDC77 | rs7136957 | 1.30E-08 | ##### | Brain - Putamen (basal gangli | 440788 | -59212 | 940788 | Yes |
| ENSG00000120647.9 | CCDC77 | rs34380840 | 1.30E-08 | ##### | Brain - Putamen (basal gangli | 440788 | -59212 | 940788 | Yes |
| ENSG00000120647.9 | CCDC77 | rs2284332 | 1.30E-08 | ##### | Brain - Hypothalamus | 440788 | -59212 | 940788 | Yes |
| ENSG00000120647.9 | CCDC77 | rs2229351 | 1.30E-08 | ##### | Brain - Hypothalamus | 440788 | -59212 | 940788 | Yes |
| ENSG00000120647.9 | CCDC77 | rs1548904 | 1.30E-08 | ##### | Brain - Cerebellar Hemisphere | 440788 | -59212 | 940788 | Yes |
| ENSG00000120647.9 | CCDC77 | rs10744561 | 1.30E-08 | ##### | Brain - Cerebellar Hemisphere | 440788 | -59212 | 940788 | Yes |
| ENSG00000120647.9 | CCDC77 | rs1972582 | 1.30E-08 | ##### | Brain - Cerebellar Hemisphere | 440788 | -59212 | 940788 | Yes |
| ENSG00000120647.9 | CCDC77 | rs10774038 | 1.30E-08 | ##### | Brain - Cerebellar Hemisphere | 440788 | -59212 | 940788 | Yes |
| ENSG00000120647.9 | CCDC77 | rs576159 | 1.30E-08 | ##### | Brain - Spinal cord (cervical c- | 440788 | -59212 | 940788 | Yes |
| ENSG00000120647.9 | CCDC77 | rs2080402 | 1.30E-08 | ##### | Brain - Frontal Cortex (BA9) | 440788 | -59212 | 940788 | Yes |
| ENSG00000120647.9 | CCDC77 | rs12370433 | 1.40E-08 | ##### | Brain - Frontal Cortex (BA9) | 440788 | -59212 | 940788 | Yes |
| ENSG00000120647.9 | CCDC77 | rs56042953 | 1.40E-08 | ##### | Brain - Hypothalamus | 440788 | -59212 | 940788 | Yes |
| ENSG00000120647.9 | CCDC77 | rs10848766 | 1.50E-08 | ##### | Brain - Caudate (basal ganglia | 440788 | -59212 | 940788 | Yes |
| ENSG00000120647.9 | CCDC77 | rs579526 | 1.50E-08 | ##### | Brain - Putamen (basal gangli | 440788 | -59212 | 940788 | Yes |
| ENSG00000120647.9 | CCDC77 | rs507597 | 1.50E-08 | ##### | Brain - Cerebellum | 440788 | -59212 | 940788 | Yes |
| ENSG00000120647.9 | CCDC77 | rs17800744 | 1.50E-08 | ##### | Brain - Cerebellum | 440788 | -59212 | 940788 | Yes |
| ENSG00000120647.9 | CCDC77 | rs77854644 | 1.50E-08 | ##### | Brain - Cerebellum | 440788 | -59212 | 940788 | Yes |
| ENSG00000120647.9 | CCDC77 | rs576994 | 1.50E-08 | ##### | Brain - Cortex | 440788 | -59212 | 940788 | Yes |
| ENSG00000120647.9 | CCDC77 | rs797759 | 1.50E-08 | ##### | Brain - Cortex | 440788 | -59212 | 940788 | Yes |
| ENSG00000120647.9 | CCDC77 | rs479836 | 1.50E-08 | ##### | Brain - Cortex | 440788 | -59212 | 940788 | Yes |
| ENSG00000120647.9 | CCDC77 | rs2229351 | 1.50E-08 | ##### | Brain - Cortex | 440788 | -59212 | 940788 | Yes |
| ENSG00000120647.9 | CCDC77 | rs4980810 | 1.50E-08 | ##### | Brain - Cortex | 440788 | -59212 | 940788 | Yes |
| ENSG00000120647.9 | CCDC77 | rs11062302 | 1.50E-08 | ##### | Brain - Cortex | 440788 | -59212 | 940788 | Yes |

| | | | | | | | | | |
|---|---|---|---|---|---|---|---|---|---|
| ENSG00000120647.9 | CCDC77 | rs12314497 | 1.50E-08 | ##### | Brain - Cortex | 440788 | -59212 | 940788 | Yes |
| ENSG00000120647.9 | CCDC77 | rs534925791 | 1.50E-08 | ##### | Brain - Cortex | 440788 | -59212 | 940788 | Yes |
| ENSG00000120647.9 | CCDC77 | rs144729439 | 1.50E-08 | ##### | Brain - Cortex | 440788 | -59212 | 940788 | Yes |
| ENSG00000120647.9 | CCDC77 | rs11062385 | 1.50E-08 | ##### | Brain - Cortex | 440788 | -59212 | 940788 | Yes |
| ENSG00000120647.9 | CCDC77 | rs11611810 | 1.50E-08 | ##### | Brain - Cortex | 440788 | -59212 | 940788 | Yes |
| ENSG00000120647.9 | CCDC77 | rs10848715 | 1.50E-08 | ##### | Brain - Cortex | 440788 | -59212 | 940788 | Yes |
| ENSG00000120647.9 | CCDC77 | rs10848716 | 1.50E-08 | ##### | Brain - Cortex | 440788 | -59212 | 940788 | Yes |
| ENSG00000120647.9 | CCDC77 | rs61907027 | 1.50E-08 | ##### | Brain - Cortex | 440788 | -59212 | 940788 | Yes |
| ENSG00000120647.9 | CCDC77 | rs7301422 | 1.50E-08 | ##### | Brain - Cortex | 440788 | -59212 | 940788 | Yes |
| ENSG00000120647.9 | CCDC77 | rs11062512 | 1.50E-08 | ##### | Brain - Cortex | 440788 | -59212 | 940788 | Yes |
| ENSG00000120647.9 | CCDC77 | rs3759371 | 1.50E-08 | ##### | Brain - Cortex | 440788 | -59212 | 940788 | Yes |
| ENSG00000120647.9 | CCDC77 | rs6489448 | 1.50E-08 | ##### | Brain - Cortex | 440788 | -59212 | 940788 | Yes |
| ENSG00000120647.9 | CCDC77 | rs7316660 | 1.50E-08 | ##### | Brain - Cortex | 440788 | -59212 | 940788 | Yes |
| ENSG00000120647.9 | CCDC77 | rs6489449 | 1.50E-08 | ##### | Brain - Cortex | 440788 | -59212 | 940788 | Yes |
| ENSG00000120647.9 | CCDC77 | rs10735021 | 1.50E-08 | ##### | Brain - Cortex | 440788 | -59212 | 940788 | Yes |
| ENSG00000120647.9 | CCDC77 | rs10848845 | 1.50E-08 | ##### | Brain - Cortex | 440788 | -59212 | 940788 | Yes |
| ENSG00000120647.9 | CCDC77 | rs10774137 | 1.50E-08 | ##### | Brain - Cortex | 440788 | -59212 | 940788 | Yes |
| ENSG00000120647.9 | CCDC77 | rs11835325 | 1.50E-08 | ##### | Brain - Cortex | 440788 | -59212 | 940788 | Yes |
| ENSG00000120647.9 | CCDC77 | rs10848849 | 1.50E-08 | ##### | Brain - Cortex | 440788 | -59212 | 940788 | Yes |
| ENSG00000120647.9 | CCDC77 | rs6489454 | 1.50E-08 | ##### | Brain - Cortex | 440788 | -59212 | 940788 | Yes |
| ENSG00000120647.9 | CCDC77 | rs7134608 | 1.50E-08 | ##### | Brain - Cortex | 440788 | -59212 | 940788 | Yes |
| ENSG00000120647.9 | CCDC77 | rs7138890 | 1.50E-08 | ##### | Brain - Cortex | 440788 | -59212 | 940788 | Yes |
| ENSG00000120647.9 | CCDC77 | rs10431355 | 1.50E-08 | ##### | Brain - Cortex | 440788 | -59212 | 940788 | Yes |
| ENSG00000120647.9 | CCDC77 | rs6489458 | 1.50E-08 | ##### | Brain - Cortex | 440788 | -59212 | 940788 | Yes |
| ENSG00000120647.9 | CCDC77 | rs2300134 | 1.50E-08 | ##### | Brain - Cortex | 440788 | -59212 | 940788 | Yes |
| ENSG00000120647.9 | CCDC77 | rs3782852 | 1.50E-08 | ##### | Brain - Cortex | 440788 | -59212 | 940788 | Yes |
| ENSG00000120647.9 | CCDC77 | rs201946814 | 1.50E-08 | ##### | Brain - Cortex | 440788 | -59212 | 940788 | Yes |
| ENSG00000120647.9 | CCDC77 | rs6489463 | 1.50E-08 | ##### | Brain - Cortex | 440788 | -59212 | 940788 | Yes |
| ENSG00000120647.9 | CCDC77 | rs2079785 | 1.50E-08 | ##### | Brain - Cortex | 440788 | -59212 | 940788 | Yes |
| ENSG00000120647.9 | CCDC77 | rs7963937 | 1.50E-08 | ##### | Brain - Cortex | 440788 | -59212 | 940788 | Yes |
| ENSG00000120647.9 | CCDC77 | rs2369276 | 1.50E-08 | ##### | Brain - Cortex | 440788 | -59212 | 940788 | Yes |
| ENSG00000120647.9 | CCDC77 | rs7953137 | 1.50E-08 | ##### | Brain - Cortex | 440788 | -59212 | 940788 | Yes |

| | | | | | | | | | |
|---|---|---|---|---|---|---|---|---|---|
| ENSG00000120647.9 | CCDC77 | rs6489471 | 1.50E-08 | ##### | Brain - Cortex | 440788 | -59212 | 940788 | Yes |
| ENSG00000120647.9 | CCDC77 | rs7312065 | 1.50E-08 | ##### | Brain - Cortex | 440788 | -59212 | 940788 | Yes |
| ENSG00000120647.9 | CCDC77 | rs7136957 | 1.50E-08 | ##### | Brain - Cortex | 440788 | -59212 | 940788 | Yes |
| ENSG00000120647.9 | CCDC77 | rs34380840 | 1.50E-08 | ##### | Brain - Cortex | 440788 | -59212 | 940788 | Yes |
| ENSG00000120647.9 | CCDC77 | rs7965303 | 1.50E-08 | ##### | Brain - Cortex | 440788 | -59212 | 940788 | Yes |
| ENSG00000120647.9 | CCDC77 | rs10848879 | 1.50E-08 | ##### | Brain - Cortex | 440788 | -59212 | 940788 | Yes |
| ENSG00000120647.9 | CCDC77 | rs57022939 | 1.50E-08 | ##### | Brain - Cortex | 440788 | -59212 | 940788 | Yes |
| ENSG00000120647.9 | CCDC77 | rs4980875 | 1.50E-08 | ##### | Brain - Cortex | 440788 | -59212 | 940788 | Yes |
| ENSG00000120647.9 | CCDC77 | rs61906995 | 1.50E-08 | ##### | Brain - Cortex | 440788 | -59212 | 940788 | Yes |
| ENSG00000120647.9 | CCDC77 | rs12582850 | 1.50E-08 | ##### | Brain - Cortex | 440788 | -59212 | 940788 | Yes |
| ENSG00000120647.9 | CCDC77 | rs7298019 | 1.50E-08 | ##### | Brain - Cortex | 440788 | -59212 | 940788 | Yes |
| ENSG00000120647.9 | CCDC77 | rs2098094 | 1.50E-08 | ##### | Brain - Cortex | 440788 | -59212 | 940788 | Yes |
| ENSG00000120647.9 | CCDC77 | rs2159364 | 1.50E-08 | ##### | Brain - Cortex | 440788 | -59212 | 940788 | Yes |
| ENSG00000120647.9 | CCDC77 | rs10848868 | 1.50E-08 | ##### | Brain - Cortex | 440788 | -59212 | 940788 | Yes |
| ENSG00000120647.9 | CCDC77 | rs1860360 | 1.50E-08 | ##### | Brain - Cortex | 440788 | -59212 | 940788 | Yes |
| ENSG00000120647.9 | CCDC77 | rs2369280 | 1.50E-08 | ##### | Brain - Nucleus accumbens (b | 440788 | -59212 | 940788 | Yes |
| ENSG00000120647.9 | CCDC77 | rs3782860 | 1.50E-08 | ##### | Brain - Cerebellar Hemisphere | 440788 | -59212 | 940788 | Yes |
| ENSG00000120647.9 | CCDC77 | rs2253565 | 1.50E-08 | ##### | Brain - Anterior cingulate cort | 440788 | -59212 | 940788 | Yes |
| ENSG00000120647.9 | CCDC77 | rs3213750 | 1.50E-08 | ##### | Brain - Frontal Cortex (BA9) | 440788 | -59212 | 940788 | Yes |
| ENSG00000120647.9 | CCDC77 | rs4980881 | 1.50E-08 | ##### | Brain - Frontal Cortex (BA9) | 440788 | -59212 | 940788 | Yes |
| ENSG00000120647.9 | CCDC77 | rs7969761 | 1.50E-08 | ##### | Brain - Frontal Cortex (BA9) | 440788 | -59212 | 940788 | Yes |
| ENSG00000120647.9 | CCDC77 | rs7965303 | 1.60E-08 | ##### | Brain - Putamen (basal gangli | 440788 | -59212 | 940788 | Yes |
| ENSG00000120647.9 | CCDC77 | rs10848879 | 1.60E-08 | ##### | Brain - Putamen (basal gangli | 440788 | -59212 | 940788 | Yes |
| ENSG00000120647.9 | CCDC77 | rs7972566 | 1.60E-08 | ##### | Brain - Nucleus accumbens (b | 440788 | -59212 | 940788 | Yes |
| ENSG00000120647.9 | CCDC77 | rs10774170 | 1.60E-08 | ##### | Brain - Nucleus accumbens (b | 440788 | -59212 | 940788 | Yes |
| ENSG00000120647.9 | CCDC77 | rs3213750 | 1.60E-08 | ##### | Brain - Hypothalamus | 440788 | -59212 | 940788 | Yes |
| ENSG00000120647.9 | CCDC77 | rs4980881 | 1.60E-08 | ##### | Brain - Hypothalamus | 440788 | -59212 | 940788 | Yes |
| ENSG00000120647.9 | CCDC77 | rs2300133 | 1.60E-08 | ##### | Brain - Putamen (basal gangli | 440788 | -59212 | 940788 | Yes |
| ENSG00000120647.9 | CCDC77 | rs2191159 | 1.60E-08 | ##### | Brain - Frontal Cortex (BA9) | 440788 | -59212 | 940788 | Yes |
| ENSG00000120647.9 | CCDC77 | rs10744627 | 1.70E-08 | ##### | Brain - Nucleus accumbens (b | 440788 | -59212 | 940788 | Yes |
| ENSG00000120647.9 | CCDC77 | rs75894333 | 1.80E-08 | ##### | Brain - Cortex | 440788 | -59212 | 940788 | Yes |
| ENSG00000120647.9 | CCDC77 | rs507597 | 1.80E-08 | ##### | Brain - Cortex | 440788 | -59212 | 940788 | Yes |

| | | | | | | | | | |
|---|---|---|---|---|---|---|---|---|---|
| ENSG00000120647.9 | CCDC77 | rs17800744 | 1.80E-08 | ##### | Brain - Cortex | 440788 | -59212 | 940788 | Yes |
| ENSG00000120647.9 | CCDC77 | rs77854644 | 1.80E-08 | ##### | Brain - Cortex | 440788 | -59212 | 940788 | Yes |
| ENSG00000120647.9 | CCDC77 | rs11608938 | 1.80E-08 | ##### | Brain - Cortex | 440788 | -59212 | 940788 | Yes |
| ENSG00000120647.9 | CCDC77 | rs7965038 | 1.80E-08 | ##### | Brain - Cortex | 440788 | -59212 | 940788 | Yes |
| ENSG00000120647.9 | CCDC77 | rs7307606 | 1.80E-08 | ##### | Brain - Cortex | 440788 | -59212 | 940788 | Yes |
| ENSG00000120647.9 | CCDC77 | rs10661251 | 1.80E-08 | ##### | Brain - Cortex | 440788 | -59212 | 940788 | Yes |
| ENSG00000120647.9 | CCDC77 | rs1011699 | 1.80E-08 | ##### | Brain - Cortex | 440788 | -59212 | 940788 | Yes |
| ENSG00000120647.9 | CCDC77 | rs6489475 | 1.80E-08 | ##### | Brain - Cortex | 440788 | -59212 | 940788 | Yes |
| ENSG00000120647.9 | CCDC77 | rs10774154 | 1.80E-08 | ##### | Brain - Cortex | 440788 | -59212 | 940788 | Yes |
| ENSG00000120647.9 | CCDC77 | rs10431354 | 1.80E-08 | ##### | Brain - Putamen (basal gangli | 440788 | -59212 | 940788 | Yes |
| ENSG00000120647.9 | CCDC77 | rs61908460 | 1.80E-08 | ##### | Brain - Caudate (basal ganglia | 440788 | -59212 | 940788 | Yes |
| ENSG00000120647.9 | CCDC77 | rs4980878 | 1.80E-08 | ##### | Brain - Cortex | 440788 | -59212 | 940788 | Yes |
| ENSG00000120647.9 | CCDC77 | rs10774192 | 1.90E-08 | ##### | Brain - Hypothalamus | 440788 | -59212 | 940788 | Yes |
| ENSG00000120647.9 | CCDC77 | rs11063005 | 1.90E-08 | ##### | Brain - Hypothalamus | 440788 | -59212 | 940788 | Yes |
| ENSG00000120647.9 | CCDC77 | rs11063010 | 1.90E-08 | ##### | Brain - Hypothalamus | 440788 | -59212 | 940788 | Yes |
| ENSG00000120647.9 | CCDC77 | rs2229351 | 1.90E-08 | ##### | Brain - Cerebellum | 440788 | -59212 | 940788 | Yes |
| ENSG00000120647.9 | CCDC77 | rs7133796 | 1.90E-08 | ##### | Brain - Nucleus accumbens (b | 440788 | -59212 | 940788 | Yes |
| ENSG00000120647.9 | CCDC77 | rs12298843 | 2.00E-08 | ##### | Brain - Putamen (basal gangli | 440788 | -59212 | 940788 | Yes |
| ENSG00000120647.9 | CCDC77 | rs111250603 | 2.00E-08 | ##### | Brain - Nucleus accumbens (b | 440788 | -59212 | 940788 | Yes |
| ENSG00000120647.9 | CCDC77 | rs2369277 | 2.00E-08 | ##### | Brain - Nucleus accumbens (b | 440788 | -59212 | 940788 | Yes |
| ENSG00000120647.9 | CCDC77 | rs9805067 | 2.00E-08 | ##### | Brain - Nucleus accumbens (b | 440788 | -59212 | 940788 | Yes |
| ENSG00000120647.9 | CCDC77 | rs1541464 | 2.00E-08 | ##### | Brain - Nucleus accumbens (b | 440788 | -59212 | 940788 | Yes |
| ENSG00000120647.9 | CCDC77 | rs1541465 | 2.00E-08 | ##### | Brain - Nucleus accumbens (b | 440788 | -59212 | 940788 | Yes |
| ENSG00000120647.9 | CCDC77 | rs4980817 | 2.00E-08 | ##### | Brain - Nucleus accumbens (b | 440788 | -59212 | 940788 | Yes |
| ENSG00000120647.9 | CCDC77 | rs2284332 | 2.10E-08 | ##### | Brain - Cortex | 440788 | -59212 | 940788 | Yes |
| ENSG00000120647.9 | CCDC77 | rs3753156 | 2.10E-08 | ##### | Brain - Hypothalamus | 440788 | -59212 | 940788 | Yes |
| ENSG00000120647.9 | CCDC77 | rs12581260 | 2.10E-08 | ##### | Brain - Hypothalamus | 440788 | -59212 | 940788 | Yes |
| ENSG00000120647.9 | CCDC77 | rs16929783 | 2.10E-08 | ##### | Brain - Hypothalamus | 440788 | -59212 | 940788 | Yes |
| ENSG00000120647.9 | CCDC77 | rs372437080 | 2.10E-08 | ##### | Brain - Anterior cingulate cort | 440788 | -59212 | 940788 | Yes |
| ENSG00000120647.9 | CCDC77 | rs2052279 | 2.20E-08 | ##### | Brain - Putamen (basal gangli | 440788 | -59212 | 940788 | Yes |
| ENSG00000120647.9 | CCDC77 | rs78813329 | 2.20E-08 | ##### | Brain - Putamen (basal gangli | 440788 | -59212 | 940788 | Yes |
| ENSG00000120647.9 | CCDC77 | rs3753157 | 2.20E-08 | ##### | Brain - Putamen (basal gangli | 440788 | -59212 | 940788 | Yes |

| | | | | | | | | | |
|---|---|---|---|---|---|---|---|---|---|
| ENSG00000120647.9 | CCDC77 | rs4980883 | 2.20E-08 | ##### | Brain - Putamen (basal gangli | 440788 | -59212 | 940788 | Yes |
| ENSG00000120647.9 | CCDC77 | rs137910445 | 2.20E-08 | ##### | Brain - Putamen (basal gangli | 440788 | -59212 | 940788 | Yes |
| ENSG00000120647.9 | CCDC77 | rs11062537 | 2.20E-08 | ##### | Brain - Putamen (basal gangli | 440788 | -59212 | 940788 | Yes |
| ENSG00000120647.9 | CCDC77 | rs715230 | 2.20E-08 | ##### | Brain - Putamen (basal gangli | 440788 | -59212 | 940788 | Yes |
| ENSG00000120647.9 | CCDC77 | rs715229 | 2.20E-08 | ##### | Brain - Putamen (basal gangli | 440788 | -59212 | 940788 | Yes |
| ENSG00000120647.9 | CCDC77 | rs715228 | 2.20E-08 | ##### | Brain - Putamen (basal gangli | 440788 | -59212 | 940788 | Yes |
| ENSG00000120647.9 | CCDC77 | rs10848838 | 2.20E-08 | ##### | Brain - Putamen (basal gangli | 440788 | -59212 | 940788 | Yes |
| ENSG00000120647.9 | CCDC77 | rs10774152 | 2.20E-08 | ##### | Brain - Nucleus accumbens (b | 440788 | -59212 | 940788 | Yes |
| ENSG00000120647.9 | CCDC77 | rs10774153 | 2.20E-08 | ##### | Brain - Nucleus accumbens (b | 440788 | -59212 | 940788 | Yes |
| ENSG00000120647.9 | CCDC77 | rs1048466 | 2.20E-08 | ##### | Brain - Cerebellar Hemisphere | 440788 | -59212 | 940788 | Yes |
| ENSG00000120647.9 | CCDC77 | rs67523949 | 2.20E-08 | ##### | Brain - Caudate (basal ganglia | 440788 | -59212 | 940788 | Yes |
| ENSG00000120647.9 | CCDC77 | rs4980892 | 2.30E-08 | ##### | Brain - Nucleus accumbens (b | 440788 | -59212 | 940788 | Yes |
| ENSG00000120647.9 | CCDC77 | rs1048466 | 2.30E-08 | ##### | Brain - Frontal Cortex (BA9) | 440788 | -59212 | 940788 | Yes |
| ENSG00000120647.9 | CCDC77 | rs67523949 | 2.30E-08 | ##### | Brain - Cerebellar Hemisphere | 440788 | -59212 | 940788 | Yes |
| ENSG00000120647.9 | CCDC77 | rs12370433 | 2.50E-08 | ##### | Brain - Hypothalamus | 440788 | -59212 | 940788 | Yes |
| ENSG00000120647.9 | CCDC77 | rs2080403 | 2.50E-08 | ##### | Brain - Cerebellar Hemisphere | 440788 | -59212 | 940788 | Yes |
| ENSG00000120647.9 | CCDC77 | rs2289957 | 2.50E-08 | ##### | Brain - Cerebellar Hemisphere | 440788 | -59212 | 940788 | Yes |
| ENSG00000120647.9 | CCDC77 | rs55657100 | 2.60E-08 | ##### | Brain - Caudate (basal ganglia | 440788 | -59212 | 940788 | Yes |
| ENSG00000120647.9 | CCDC77 | rs10774192 | 2.70E-08 | ##### | Brain - Caudate (basal ganglia | 440788 | -59212 | 940788 | Yes |
| ENSG00000120647.9 | CCDC77 | rs482514 | 2.70E-08 | ##### | Brain - Putamen (basal gangli | 440788 | -59212 | 940788 | Yes |
| ENSG00000120647.9 | CCDC77 | rs7315852 | 2.70E-08 | ##### | Brain - Putamen (basal gangli | 440788 | -59212 | 940788 | Yes |
| ENSG00000120647.9 | CCDC77 | rs2284334 | 2.70E-08 | ##### | Brain - Putamen (basal gangli | 440788 | -59212 | 940788 | Yes |
| ENSG00000120647.9 | CCDC77 | rs720128 | 2.70E-08 | ##### | Brain - Putamen (basal gangli | 440788 | -59212 | 940788 | Yes |
| ENSG00000120647.9 | CCDC77 | rs9739950 | 2.70E-08 | ##### | Brain - Putamen (basal gangli | 440788 | -59212 | 940788 | Yes |
| ENSG00000120647.9 | CCDC77 | rs12311569 | 2.70E-08 | ##### | Brain - Putamen (basal gangli | 440788 | -59212 | 940788 | Yes |
| ENSG00000120647.9 | CCDC77 | rs7138981 | 2.70E-08 | ##### | Brain - Nucleus accumbens (b | 440788 | -59212 | 940788 | Yes |
| ENSG00000120647.9 | CCDC77 | rs9669094 | 2.70E-08 | ##### | Brain - Nucleus accumbens (b | 440788 | -59212 | 940788 | Yes |
| ENSG00000120647.9 | CCDC77 | rs7132763 | 2.70E-08 | ##### | Brain - Nucleus accumbens (b | 440788 | -59212 | 940788 | Yes |
| ENSG00000120647.9 | CCDC77 | rs2015554 | 2.70E-08 | ##### | Brain - Nucleus accumbens (b | 440788 | -59212 | 940788 | Yes |
| ENSG00000120647.9 | CCDC77 | rs55657100 | 2.80E-08 | ##### | Brain - Cerebellar Hemisphere | 440788 | -59212 | 940788 | Yes |
| ENSG00000120647.9 | CCDC77 | rs4980898 | 2.80E-08 | ##### | Brain - Cerebellar Hemisphere | 440788 | -59212 | 940788 | Yes |
| ENSG00000120647.9 | CCDC77 | rs11062167 | 2.80E-08 | ##### | Brain - Cerebellar Hemisphere | 440788 | -59212 | 940788 | Yes |

| | | | | | | | | | |
|---|---|---|---|---|---|---|---|---|---|
| ENSG00000120647.9 | CCDC77 | rs55990424 | 2.80E-08 | ##### | Brain - Cerebellar Hemisphere | 440788 | -59212 | 940788 | Yes |
| ENSG00000120647.9 | CCDC77 | rs486098 | 2.80E-08 | ##### | Brain - Anterior cingulate cort | 440788 | -59212 | 940788 | Yes |
| ENSG00000120647.9 | CCDC77 | rs11613331 | 2.80E-08 | ##### | Brain - Frontal Cortex (BA9) | 440788 | -59212 | 940788 | Yes |
| ENSG00000120647.9 | CCDC77 | rs11062930 | 2.80E-08 | ##### | Brain - Caudate (basal ganglia | 440788 | -59212 | 940788 | Yes |
| ENSG00000120647.9 | CCDC77 | rs525381 | 2.80E-08 | ##### | Brain - Nucleus accumbens (b | 440788 | -59212 | 940788 | Yes |
| ENSG00000120647.9 | CCDC77 | rs3213751 | 2.80E-08 | ##### | Brain - Spinal cord (cervical c- | 440788 | -59212 | 940788 | Yes |
| ENSG00000120647.9 | CCDC77 | rs3815558 | 2.80E-08 | ##### | Brain - Spinal cord (cervical c- | 440788 | -59212 | 940788 | Yes |
| ENSG00000120647.9 | CCDC77 | rs78998485 | 2.80E-08 | ##### | Brain - Spinal cord (cervical c- | 440788 | -59212 | 940788 | Yes |
| ENSG00000120647.9 | CCDC77 | rs4980884 | 2.80E-08 | ##### | Brain - Spinal cord (cervical c- | 440788 | -59212 | 940788 | Yes |
| ENSG00000120647.9 | CCDC77 | rs7135768 | 2.80E-08 | ##### | Brain - Spinal cord (cervical c- | 440788 | -59212 | 940788 | Yes |
| ENSG00000120647.9 | CCDC77 | rs6489450 | 2.80E-08 | ##### | Brain - Spinal cord (cervical c- | 440788 | -59212 | 940788 | Yes |
| ENSG00000120647.9 | CCDC77 | rs1048466 | 3.00E-08 | ##### | Brain - Hypothalamus | 440788 | -59212 | 940788 | Yes |
| ENSG00000120647.9 | CCDC77 | rs4980908 | 3.00E-08 | ##### | Brain - Frontal Cortex (BA9) | 440788 | -59212 | 940788 | Yes |
| ENSG00000120647.9 | CCDC77 | rs7132845 | 3.10E-08 | ##### | Brain - Hypothalamus | 440788 | -59212 | 940788 | Yes |
| ENSG00000120647.9 | CCDC77 | rs11613357 | 3.10E-08 | ##### | Brain - Cerebellar Hemisphere | 440788 | -59212 | 940788 | Yes |
| ENSG00000120647.9 | CCDC77 | rs1207324 | 3.10E-08 | ##### | Brain - Cortex | 440788 | -59212 | 940788 | Yes |
| ENSG00000120647.9 | CCDC77 | rs4980886 | 3.10E-08 | ##### | Brain - Spinal cord (cervical c- | 440788 | -59212 | 940788 | Yes |
| ENSG00000120647.9 | CCDC77 | rs12372234 | 3.10E-08 | ##### | Brain - Caudate (basal ganglia | 440788 | -59212 | 940788 | Yes |
| ENSG00000120647.9 | CCDC77 | rs10848961 | 3.20E-08 | ##### | Brain - Caudate (basal ganglia | 440788 | -59212 | 940788 | Yes |
| ENSG00000120647.9 | CCDC77 | rs10744627 | 3.20E-08 | ##### | Brain - Cerebellum | 440788 | -59212 | 940788 | Yes |
| ENSG00000120647.9 | CCDC77 | rs4140962 | 3.30E-08 | ##### | Brain - Caudate (basal ganglia | 440788 | -59212 | 940788 | Yes |
| ENSG00000120647.9 | CCDC77 | rs7136957 | 3.30E-08 | ##### | Brain - Nucleus accumbens (b | 440788 | -59212 | 940788 | Yes |
| ENSG00000120647.9 | CCDC77 | rs2080403 | 3.40E-08 | ##### | Brain - Caudate (basal ganglia | 440788 | -59212 | 940788 | Yes |
| ENSG00000120647.9 | CCDC77 | rs2289957 | 3.40E-08 | ##### | Brain - Caudate (basal ganglia | 440788 | -59212 | 940788 | Yes |
| ENSG00000120647.9 | CCDC77 | rs10082788 | 3.40E-08 | ##### | Brain - Spinal cord (cervical c- | 440788 | -59212 | 940788 | Yes |
| ENSG00000120647.9 | CCDC77 | rs201124968 | 3.40E-08 | ##### | Brain - Cerebellum | 440788 | -59212 | 940788 | Yes |
| ENSG00000120647.9 | CCDC77 | rs7965303 | 3.40E-08 | ##### | Brain - Nucleus accumbens (b | 440788 | -59212 | 940788 | Yes |
| ENSG00000120647.9 | CCDC77 | rs10848879 | 3.40E-08 | ##### | Brain - Nucleus accumbens (b | 440788 | -59212 | 940788 | Yes |
| ENSG00000120647.9 | CCDC77 | rs4980896 | 3.50E-08 | ##### | Brain - Frontal Cortex (BA9) | 440788 | -59212 | 940788 | Yes |
| ENSG00000120647.9 | CCDC77 | rs2369278 | 3.50E-08 | ##### | Brain - Cerebellum | 440788 | -59212 | 940788 | Yes |
| ENSG00000120647.9 | CCDC77 | rs10774163 | 3.50E-08 | ##### | Brain - Cerebellum | 440788 | -59212 | 940788 | Yes |
| ENSG00000120647.9 | CCDC77 | rs6489477 | 3.50E-08 | ##### | Brain - Putamen (basal gangli | 440788 | -59212 | 940788 | Yes |

| | | | | | | | | | |
|---|---|---|---|---|---|---|---|---|---|
| ENSG00000120647.9 | CCDC77 | rs6489478 | 3.50E-08 | ##### | Brain - Putamen (basal gangli | 440788 | -59212 | 940788 | Yes |
| ENSG00000120647.9 | CCDC77 | rs3753158 | 3.50E-08 | ##### | Brain - Hypothalamus | 440788 | -59212 | 940788 | Yes |
| ENSG00000120647.9 | CCDC77 | rs10160995 | 3.60E-08 | ##### | Brain - Cerebellar Hemisphere | 440788 | -59212 | 940788 | Yes |
| ENSG00000120647.9 | CCDC77 | rs3753158 | 3.60E-08 | ##### | Brain - Putamen (basal gangli | 440788 | -59212 | 940788 | Yes |
| ENSG00000120647.9 | CCDC77 | rs10848877 | 3.60E-08 | ##### | Brain - Putamen (basal gangli | 440788 | -59212 | 940788 | Yes |
| ENSG00000120647.9 | CCDC77 | rs10160995 | 3.80E-08 | ##### | Brain - Frontal Cortex (BA9) | 440788 | -59212 | 940788 | Yes |
| ENSG00000120647.9 | CCDC77 | rs4980898 | 3.80E-08 | ##### | Brain - Frontal Cortex (BA9) | 440788 | -59212 | 940788 | Yes |
| ENSG00000120647.9 | CCDC77 | rs10774170 | 3.90E-08 | ##### | Brain - Cerebellum | 440788 | -59212 | 940788 | Yes |
| ENSG00000120647.9 | CCDC77 | rs10431354 | 3.90E-08 | ##### | Brain - Hypothalamus | 440788 | -59212 | 940788 | Yes |
| ENSG00000120647.9 | CCDC77 | rs3753158 | 3.90E-08 | ##### | Brain - Cerebellum | 440788 | -59212 | 940788 | Yes |
| ENSG00000120647.9 | CCDC77 | rs10848644 | 4.10E-08 | ##### | Brain - Cerebellar Hemisphere | 440788 | -59212 | 940788 | Yes |
| ENSG00000120647.9 | CCDC77 | rs7136116 | 4.10E-08 | ##### | Brain - Frontal Cortex (BA9) | 440788 | -59212 | 940788 | Yes |
| ENSG00000120647.9 | CCDC77 | rs11610457 | 4.10E-08 | ##### | Brain - Frontal Cortex (BA9) | 440788 | -59212 | 940788 | Yes |
| ENSG00000120647.9 | CCDC77 | rs4980884 | 4.20E-08 | ##### | Brain - Cortex | 440788 | -59212 | 940788 | Yes |
| ENSG00000120647.9 | CCDC77 | rs579526 | 4.30E-08 | ##### | Brain - Cortex | 440788 | -59212 | 940788 | Yes |
| ENSG00000120647.9 | CCDC77 | rs3782860 | 4.30E-08 | ##### | Brain - Frontal Cortex (BA9) | 440788 | -59212 | 940788 | Yes |
| ENSG00000120647.9 | CCDC77 | rs7132845 | 4.30E-08 | ##### | Brain - Spinal cord (cervical c- | 440788 | -59212 | 940788 | Yes |
| ENSG00000120647.9 | CCDC77 | rs10744561 | 4.30E-08 | ##### | Brain - Frontal Cortex (BA9) | 440788 | -59212 | 940788 | Yes |
| ENSG00000120647.9 | CCDC77 | rs56042953 | 4.30E-08 | ##### | Brain - Hippocampus | 440788 | -59212 | 940788 | Yes |
| ENSG00000120647.9 | CCDC77 | rs7963793 | 4.40E-08 | ##### | Brain - Hypothalamus | 440788 | -59212 | 940788 | Yes |
| ENSG00000120647.9 | CCDC77 | rs10848814 | 4.40E-08 | ##### | Brain - Hypothalamus | 440788 | -59212 | 940788 | Yes |
| ENSG00000120647.9 | CCDC77 | rs10848815 | 4.40E-08 | ##### | Brain - Hypothalamus | 440788 | -59212 | 940788 | Yes |
| ENSG00000120647.9 | CCDC77 | rs887311 | 4.40E-08 | ##### | Brain - Hypothalamus | 440788 | -59212 | 940788 | Yes |
| ENSG00000120647.9 | CCDC77 | rs10848905 | 4.40E-08 | ##### | Brain - Cerebellum | 440788 | -59212 | 940788 | Yes |
| ENSG00000120647.9 | CCDC77 | rs7136957 | 4.50E-08 | ##### | Brain - Cerebellum | 440788 | -59212 | 940788 | Yes |
| ENSG00000120647.9 | CCDC77 | rs3764876 | 4.60E-08 | ##### | Brain - Caudate (basal ganglia | 440788 | -59212 | 940788 | Yes |
| ENSG00000120647.9 | CCDC77 | rs10774191 | 4.60E-08 | ##### | Brain - Caudate (basal ganglia | 440788 | -59212 | 940788 | Yes |
| ENSG00000120647.9 | CCDC77 | rs61916593 | 4.60E-08 | ##### | Brain - Caudate (basal ganglia | 440788 | -59212 | 940788 | Yes |
| ENSG00000120647.9 | CCDC77 | rs4980876 | 4.70E-08 | ##### | Brain - Nucleus accumbens (b | 440788 | -59212 | 940788 | Yes |
| ENSG00000120647.9 | CCDC77 | rs10848650 | 4.80E-08 | ##### | Brain - Frontal Cortex (BA9) | 440788 | -59212 | 940788 | Yes |
| ENSG00000120647.9 | CCDC77 | rs7132845 | 4.80E-08 | ##### | Brain - Putamen (basal gangli | 440788 | -59212 | 940788 | Yes |
| ENSG00000120647.9 | CCDC77 | rs576994 | 4.90E-08 | ##### | Brain - Nucleus accumbens (b | 440788 | -59212 | 940788 | Yes |

| | | | | | | | | | |
|---|---|---|---|---|---|---|---|---|---|
| ENSG00000120647.9 | CCDC77 | rs797759 | 4.90E-08 | ##### | Brain - Nucleus accumbens (b | 440788 | -59212 | 940788 | Yes |
| ENSG00000120647.9 | CCDC77 | rs479836 | 4.90E-08 | ##### | Brain - Nucleus accumbens (b | 440788 | -59212 | 940788 | Yes |
| ENSG00000120647.9 | CCDC77 | rs4980810 | 4.90E-08 | ##### | Brain - Nucleus accumbens (b | 440788 | -59212 | 940788 | Yes |
| ENSG00000120647.9 | CCDC77 | rs11062302 | 4.90E-08 | ##### | Brain - Nucleus accumbens (b | 440788 | -59212 | 940788 | Yes |
| ENSG00000120647.9 | CCDC77 | rs12314497 | 4.90E-08 | ##### | Brain - Nucleus accumbens (b | 440788 | -59212 | 940788 | Yes |
| ENSG00000120647.9 | CCDC77 | rs534925791 | 4.90E-08 | ##### | Brain - Nucleus accumbens (b | 440788 | -59212 | 940788 | Yes |
| ENSG00000120647.9 | CCDC77 | rs144729439 | 4.90E-08 | ##### | Brain - Nucleus accumbens (b | 440788 | -59212 | 940788 | Yes |
| ENSG00000120647.9 | CCDC77 | rs11062385 | 4.90E-08 | ##### | Brain - Nucleus accumbens (b | 440788 | -59212 | 940788 | Yes |
| ENSG00000120647.9 | CCDC77 | rs11611810 | 4.90E-08 | ##### | Brain - Nucleus accumbens (b | 440788 | -59212 | 940788 | Yes |
| ENSG00000120647.9 | CCDC77 | rs10848715 | 4.90E-08 | ##### | Brain - Nucleus accumbens (b | 440788 | -59212 | 940788 | Yes |
| ENSG00000120647.9 | CCDC77 | rs10848716 | 4.90E-08 | ##### | Brain - Nucleus accumbens (b | 440788 | -59212 | 940788 | Yes |
| ENSG00000120647.9 | CCDC77 | rs61907027 | 4.90E-08 | ##### | Brain - Nucleus accumbens (b | 440788 | -59212 | 940788 | Yes |
| ENSG00000120647.9 | CCDC77 | rs7301422 | 4.90E-08 | ##### | Brain - Nucleus accumbens (b | 440788 | -59212 | 940788 | Yes |
| ENSG00000120647.9 | CCDC77 | rs11062512 | 4.90E-08 | ##### | Brain - Nucleus accumbens (b | 440788 | -59212 | 940788 | Yes |
| ENSG00000120647.9 | CCDC77 | rs3759371 | 4.90E-08 | ##### | Brain - Nucleus accumbens (b | 440788 | -59212 | 940788 | Yes |
| ENSG00000120647.9 | CCDC77 | rs6489448 | 4.90E-08 | ##### | Brain - Nucleus accumbens (b | 440788 | -59212 | 940788 | Yes |
| ENSG00000120647.9 | CCDC77 | rs7316660 | 4.90E-08 | ##### | Brain - Nucleus accumbens (b | 440788 | -59212 | 940788 | Yes |
| ENSG00000120647.9 | CCDC77 | rs6489449 | 4.90E-08 | ##### | Brain - Nucleus accumbens (b | 440788 | -59212 | 940788 | Yes |
| ENSG00000120647.9 | CCDC77 | rs10735021 | 4.90E-08 | ##### | Brain - Nucleus accumbens (b | 440788 | -59212 | 940788 | Yes |
| ENSG00000120647.9 | CCDC77 | rs10848845 | 4.90E-08 | ##### | Brain - Nucleus accumbens (b | 440788 | -59212 | 940788 | Yes |
| ENSG00000120647.9 | CCDC77 | rs10774137 | 4.90E-08 | ##### | Brain - Nucleus accumbens (b | 440788 | -59212 | 940788 | Yes |
| ENSG00000120647.9 | CCDC77 | rs11835325 | 4.90E-08 | ##### | Brain - Nucleus accumbens (b | 440788 | -59212 | 940788 | Yes |
| ENSG00000120647.9 | CCDC77 | rs10848849 | 4.90E-08 | ##### | Brain - Nucleus accumbens (b | 440788 | -59212 | 940788 | Yes |
| ENSG00000120647.9 | CCDC77 | rs6489454 | 4.90E-08 | ##### | Brain - Nucleus accumbens (b | 440788 | -59212 | 940788 | Yes |
| ENSG00000120647.9 | CCDC77 | rs7134608 | 4.90E-08 | ##### | Brain - Nucleus accumbens (b | 440788 | -59212 | 940788 | Yes |
| ENSG00000120647.9 | CCDC77 | rs7138890 | 4.90E-08 | ##### | Brain - Nucleus accumbens (b | 440788 | -59212 | 940788 | Yes |
| ENSG00000120647.9 | CCDC77 | rs10431355 | 4.90E-08 | ##### | Brain - Nucleus accumbens (b | 440788 | -59212 | 940788 | Yes |
| ENSG00000120647.9 | CCDC77 | rs6489458 | 4.90E-08 | ##### | Brain - Nucleus accumbens (b | 440788 | -59212 | 940788 | Yes |
| ENSG00000120647.9 | CCDC77 | rs2300134 | 4.90E-08 | ##### | Brain - Nucleus accumbens (b | 440788 | -59212 | 940788 | Yes |
| ENSG00000120647.9 | CCDC77 | rs3782852 | 4.90E-08 | ##### | Brain - Nucleus accumbens (b | 440788 | -59212 | 940788 | Yes |
| ENSG00000120647.9 | CCDC77 | rs201124968 | 4.90E-08 | ##### | Brain - Nucleus accumbens (b | 440788 | -59212 | 940788 | Yes |
| ENSG00000120647.9 | CCDC77 | rs201946814 | 4.90E-08 | ##### | Brain - Nucleus accumbens (b | 440788 | -59212 | 940788 | Yes |

| | | | | | | | | | |
|---|---|---|---|---|---|---|---|---|---|
| ENSG00000120647.9 | CCDC77 | rs6489463 | 4.90E-08 | ##### | Brain - Nucleus accumbens (b | 440788 | -59212 | 940788 | Yes |
| ENSG00000120647.9 | CCDC77 | rs2079785 | 4.90E-08 | ##### | Brain - Nucleus accumbens (b | 440788 | -59212 | 940788 | Yes |
| ENSG00000120647.9 | CCDC77 | rs7963937 | 4.90E-08 | ##### | Brain - Nucleus accumbens (b | 440788 | -59212 | 940788 | Yes |
| ENSG00000120647.9 | CCDC77 | rs2369276 | 4.90E-08 | ##### | Brain - Nucleus accumbens (b | 440788 | -59212 | 940788 | Yes |
| ENSG00000120647.9 | CCDC77 | rs7953137 | 4.90E-08 | ##### | Brain - Nucleus accumbens (b | 440788 | -59212 | 940788 | Yes |
| ENSG00000120647.9 | CCDC77 | rs6489471 | 4.90E-08 | ##### | Brain - Nucleus accumbens (b | 440788 | -59212 | 940788 | Yes |
| ENSG00000120647.9 | CCDC77 | rs7312065 | 4.90E-08 | ##### | Brain - Nucleus accumbens (b | 440788 | -59212 | 940788 | Yes |
| ENSG00000120647.9 | CCDC77 | rs34380840 | 4.90E-08 | ##### | Brain - Nucleus accumbens (b | 440788 | -59212 | 940788 | Yes |
| ENSG00000120647.9 | CCDC77 | rs2300131 | 4.90E-08 | ##### | Brain - Hypothalamus | 440788 | -59212 | 940788 | Yes |
| ENSG00000120647.9 | CCDC77 | rs7133796 | 5.10E-08 | ##### | Brain - Anterior cingulate cort | 440788 | -59212 | 940788 | Yes |
| ENSG00000120647.9 | CCDC77 | rs487252 | 5.20E-08 | ##### | Brain - Frontal Cortex (BA9) | 440788 | -59212 | 940788 | Yes |
| ENSG00000120647.9 | CCDC77 | rs576994 | 5.20E-08 | ##### | Brain - Cerebellum | 440788 | -59212 | 940788 | Yes |
| ENSG00000120647.9 | CCDC77 | rs797759 | 5.20E-08 | ##### | Brain - Cerebellum | 440788 | -59212 | 940788 | Yes |
| ENSG00000120647.9 | CCDC77 | rs479836 | 5.20E-08 | ##### | Brain - Cerebellum | 440788 | -59212 | 940788 | Yes |
| ENSG00000120647.9 | CCDC77 | rs4980810 | 5.20E-08 | ##### | Brain - Cerebellum | 440788 | -59212 | 940788 | Yes |
| ENSG00000120647.9 | CCDC77 | rs11062302 | 5.20E-08 | ##### | Brain - Cerebellum | 440788 | -59212 | 940788 | Yes |
| ENSG00000120647.9 | CCDC77 | rs12314497 | 5.20E-08 | ##### | Brain - Cerebellum | 440788 | -59212 | 940788 | Yes |
| ENSG00000120647.9 | CCDC77 | rs534925791 | 5.20E-08 | ##### | Brain - Cerebellum | 440788 | -59212 | 940788 | Yes |
| ENSG00000120647.9 | CCDC77 | rs144729439 | 5.20E-08 | ##### | Brain - Cerebellum | 440788 | -59212 | 940788 | Yes |
| ENSG00000120647.9 | CCDC77 | rs11062385 | 5.20E-08 | ##### | Brain - Cerebellum | 440788 | -59212 | 940788 | Yes |
| ENSG00000120647.9 | CCDC77 | rs11611810 | 5.20E-08 | ##### | Brain - Cerebellum | 440788 | -59212 | 940788 | Yes |
| ENSG00000120647.9 | CCDC77 | rs10848715 | 5.20E-08 | ##### | Brain - Cerebellum | 440788 | -59212 | 940788 | Yes |
| ENSG00000120647.9 | CCDC77 | rs10848716 | 5.20E-08 | ##### | Brain - Cerebellum | 440788 | -59212 | 940788 | Yes |
| ENSG00000120647.9 | CCDC77 | rs61907027 | 5.20E-08 | ##### | Brain - Cerebellum | 440788 | -59212 | 940788 | Yes |
| ENSG00000120647.9 | CCDC77 | rs7301422 | 5.20E-08 | ##### | Brain - Cerebellum | 440788 | -59212 | 940788 | Yes |
| ENSG00000120647.9 | CCDC77 | rs11062512 | 5.20E-08 | ##### | Brain - Cerebellum | 440788 | -59212 | 940788 | Yes |
| ENSG00000120647.9 | CCDC77 | rs3759371 | 5.20E-08 | ##### | Brain - Cerebellum | 440788 | -59212 | 940788 | Yes |
| ENSG00000120647.9 | CCDC77 | rs6489448 | 5.20E-08 | ##### | Brain - Cerebellum | 440788 | -59212 | 940788 | Yes |
| ENSG00000120647.9 | CCDC77 | rs7316660 | 5.20E-08 | ##### | Brain - Cerebellum | 440788 | -59212 | 940788 | Yes |
| ENSG00000120647.9 | CCDC77 | rs6489449 | 5.20E-08 | ##### | Brain - Cerebellum | 440788 | -59212 | 940788 | Yes |
| ENSG00000120647.9 | CCDC77 | rs10735021 | 5.20E-08 | ##### | Brain - Cerebellum | 440788 | -59212 | 940788 | Yes |
| ENSG00000120647.9 | CCDC77 | rs10848845 | 5.20E-08 | ##### | Brain - Cerebellum | 440788 | -59212 | 940788 | Yes |

| | | | | | | | | | |
|---|---|---|---|---|---|---|---|---|---|
| ENSG00000120647.9 | CCDC77 | rs10774137 | 5.20E-08 | ##### | Brain - Cerebellum | 440788 | -59212 | 940788 | Yes |
| ENSG00000120647.9 | CCDC77 | rs11835325 | 5.20E-08 | ##### | Brain - Cerebellum | 440788 | -59212 | 940788 | Yes |
| ENSG00000120647.9 | CCDC77 | rs10848849 | 5.20E-08 | ##### | Brain - Cerebellum | 440788 | -59212 | 940788 | Yes |
| ENSG00000120647.9 | CCDC77 | rs6489454 | 5.20E-08 | ##### | Brain - Cerebellum | 440788 | -59212 | 940788 | Yes |
| ENSG00000120647.9 | CCDC77 | rs7134608 | 5.20E-08 | ##### | Brain - Cerebellum | 440788 | -59212 | 940788 | Yes |
| ENSG00000120647.9 | CCDC77 | rs7138890 | 5.20E-08 | ##### | Brain - Cerebellum | 440788 | -59212 | 940788 | Yes |
| ENSG00000120647.9 | CCDC77 | rs10431355 | 5.20E-08 | ##### | Brain - Cerebellum | 440788 | -59212 | 940788 | Yes |
| ENSG00000120647.9 | CCDC77 | rs6489458 | 5.20E-08 | ##### | Brain - Cerebellum | 440788 | -59212 | 940788 | Yes |
| ENSG00000120647.9 | CCDC77 | rs2300134 | 5.20E-08 | ##### | Brain - Cerebellum | 440788 | -59212 | 940788 | Yes |
| ENSG00000120647.9 | CCDC77 | rs3782852 | 5.20E-08 | ##### | Brain - Cerebellum | 440788 | -59212 | 940788 | Yes |
| ENSG00000120647.9 | CCDC77 | rs201946814 | 5.20E-08 | ##### | Brain - Cerebellum | 440788 | -59212 | 940788 | Yes |
| ENSG00000120647.9 | CCDC77 | rs6489463 | 5.20E-08 | ##### | Brain - Cerebellum | 440788 | -59212 | 940788 | Yes |
| ENSG00000120647.9 | CCDC77 | rs2079785 | 5.20E-08 | ##### | Brain - Cerebellum | 440788 | -59212 | 940788 | Yes |
| ENSG00000120647.9 | CCDC77 | rs7963937 | 5.20E-08 | ##### | Brain - Cerebellum | 440788 | -59212 | 940788 | Yes |
| ENSG00000120647.9 | CCDC77 | rs2369276 | 5.20E-08 | ##### | Brain - Cerebellum | 440788 | -59212 | 940788 | Yes |
| ENSG00000120647.9 | CCDC77 | rs7953137 | 5.20E-08 | ##### | Brain - Cerebellum | 440788 | -59212 | 940788 | Yes |
| ENSG00000120647.9 | CCDC77 | rs6489471 | 5.20E-08 | ##### | Brain - Cerebellum | 440788 | -59212 | 940788 | Yes |
| ENSG00000120647.9 | CCDC77 | rs7312065 | 5.20E-08 | ##### | Brain - Cerebellum | 440788 | -59212 | 940788 | Yes |
| ENSG00000120647.9 | CCDC77 | rs34380840 | 5.20E-08 | ##### | Brain - Cerebellum | 440788 | -59212 | 940788 | Yes |
| ENSG00000120647.9 | CCDC77 | rs7965303 | 5.20E-08 | ##### | Brain - Cerebellum | 440788 | -59212 | 940788 | Yes |
| ENSG00000120647.9 | CCDC77 | rs10848879 | 5.20E-08 | ##### | Brain - Cerebellum | 440788 | -59212 | 940788 | Yes |
| ENSG00000120647.9 | CCDC77 | rs2284332 | 5.20E-08 | ##### | Brain - Cerebellum | 440788 | -59212 | 940788 | Yes |
| ENSG00000120647.9 | CCDC77 | rs764614 | 5.20E-08 | ##### | Brain - Cortex | 440788 | -59212 | 940788 | Yes |
| ENSG00000120647.9 | CCDC77 | rs7132763 | 5.20E-08 | ##### | Brain - Cortex | 440788 | -59212 | 940788 | Yes |
| ENSG00000120647.9 | CCDC77 | rs1207322 | 5.20E-08 | ##### | Brain - Nucleus accumbens (b | 440788 | -59212 | 940788 | Yes |
| ENSG00000120647.9 | CCDC77 | rs4980813 | 5.20E-08 | ##### | Brain - Nucleus accumbens (b | 440788 | -59212 | 940788 | Yes |
| ENSG00000120647.9 | CCDC77 | rs61906996 | 5.20E-08 | ##### | Brain - Nucleus accumbens (b | 440788 | -59212 | 940788 | Yes |
| ENSG00000120647.9 | CCDC77 | rs4980814 | 5.20E-08 | ##### | Brain - Nucleus accumbens (b | 440788 | -59212 | 940788 | Yes |
| ENSG00000120647.9 | CCDC77 | rs7977462 | 5.20E-08 | ##### | Brain - Nucleus accumbens (b | 440788 | -59212 | 940788 | Yes |
| ENSG00000120647.9 | CCDC77 | rs10744616 | 5.20E-08 | ##### | Brain - Nucleus accumbens (b | 440788 | -59212 | 940788 | Yes |
| ENSG00000120647.9 | CCDC77 | rs10774150 | 5.20E-08 | ##### | Brain - Nucleus accumbens (b | 440788 | -59212 | 940788 | Yes |
| ENSG00000120647.9 | CCDC77 | rs3831759 | 5.20E-08 | ##### | Brain - Nucleus accumbens (b | 440788 | -59212 | 940788 | Yes |

| | | | | | | | | | |
|---|---|---|---|---|---|---|---|---|---|
| ENSG00000120647.9 | CCDC77 | rs7138958 | 5.20E-08 | ##### | Brain - Cerebellum | 440788 | -59212 | 940788 | Yes |
| ENSG00000120647.9 | CCDC77 | rs2007537 | 5.20E-08 | ##### | Brain - Cerebellum | 440788 | -59212 | 940788 | Yes |
| ENSG00000120647.9 | CCDC77 | rs2052279 | 5.40E-08 | ##### | Brain - Spinal cord (cervical c- | 440788 | -59212 | 940788 | Yes |
| ENSG00000120647.9 | CCDC77 | rs78813329 | 5.40E-08 | ##### | Brain - Spinal cord (cervical c- | 440788 | -59212 | 940788 | Yes |
| ENSG00000120647.9 | CCDC77 | rs3753157 | 5.40E-08 | ##### | Brain - Spinal cord (cervical c- | 440788 | -59212 | 940788 | Yes |
| ENSG00000120647.9 | CCDC77 | rs4980883 | 5.40E-08 | ##### | Brain - Spinal cord (cervical c- | 440788 | -59212 | 940788 | Yes |
| ENSG00000120647.9 | CCDC77 | rs137910445 | 5.40E-08 | ##### | Brain - Spinal cord (cervical c- | 440788 | -59212 | 940788 | Yes |
| ENSG00000120647.9 | CCDC77 | rs11062537 | 5.40E-08 | ##### | Brain - Spinal cord (cervical c- | 440788 | -59212 | 940788 | Yes |
| ENSG00000120647.9 | CCDC77 | rs715230 | 5.40E-08 | ##### | Brain - Spinal cord (cervical c- | 440788 | -59212 | 940788 | Yes |
| ENSG00000120647.9 | CCDC77 | rs715229 | 5.40E-08 | ##### | Brain - Spinal cord (cervical c- | 440788 | -59212 | 940788 | Yes |
| ENSG00000120647.9 | CCDC77 | rs715228 | 5.40E-08 | ##### | Brain - Spinal cord (cervical c- | 440788 | -59212 | 940788 | Yes |
| ENSG00000120647.9 | CCDC77 | rs2300133 | 5.40E-08 | ##### | Brain - Spinal cord (cervical c- | 440788 | -59212 | 940788 | Yes |
| ENSG00000120647.9 | CCDC77 | rs10848838 | 5.40E-08 | ##### | Brain - Spinal cord (cervical c- | 440788 | -59212 | 940788 | Yes |
| ENSG00000120647.9 | CCDC77 | rs1548904 | 5.40E-08 | ##### | Brain - Frontal Cortex (BA9) | 440788 | -59212 | 940788 | Yes |
| ENSG00000120647.9 | CCDC77 | rs1972582 | 5.40E-08 | ##### | Brain - Frontal Cortex (BA9) | 440788 | -59212 | 940788 | Yes |
| ENSG00000120647.9 | CCDC77 | rs10774038 | 5.40E-08 | ##### | Brain - Frontal Cortex (BA9) | 440788 | -59212 | 940788 | Yes |
| ENSG00000120647.9 | CCDC77 | rs11062943 | 5.50E-08 | ##### | Brain - Caudate (basal ganglia | 440788 | -59212 | 940788 | Yes |
| ENSG00000120647.9 | CCDC77 | rs4980819 | 5.50E-08 | ##### | Brain - Caudate (basal ganglia | 440788 | -59212 | 940788 | Yes |
| ENSG00000120647.9 | CCDC77 | rs495973 | 5.50E-08 | ##### | Brain - Substantia nigra | 440788 | -59212 | 940788 | Yes |
| ENSG00000120647.9 | CCDC77 | rs7972566 | 5.50E-08 | ##### | Brain - Anterior cingulate cort | 440788 | -59212 | 940788 | Yes |
| ENSG00000120647.9 | CCDC77 | rs10774170 | 5.50E-08 | ##### | Brain - Anterior cingulate cort | 440788 | -59212 | 940788 | Yes |
| ENSG00000120647.9 | CCDC77 | rs10160995 | 5.50E-08 | ##### | Brain - Hypothalamus | 440788 | -59212 | 940788 | Yes |
| ENSG00000120647.9 | CCDC77 | rs4980898 | 5.50E-08 | ##### | Brain - Hypothalamus | 440788 | -59212 | 940788 | Yes |
| ENSG00000120647.9 | CCDC77 | rs11062930 | 5.50E-08 | ##### | Brain - Hypothalamus | 440788 | -59212 | 940788 | Yes |
| ENSG00000120647.9 | CCDC77 | rs55657100 | 5.50E-08 | ##### | Brain - Hippocampus | 440788 | -59212 | 940788 | Yes |
| ENSG00000120647.9 | CCDC77 | rs487252 | 5.80E-08 | ##### | Brain - Caudate (basal ganglia | 440788 | -59212 | 940788 | Yes |
| ENSG00000120647.9 | CCDC77 | rs3764876 | 5.80E-08 | ##### | Brain - Hypothalamus | 440788 | -59212 | 940788 | Yes |
| ENSG00000120647.9 | CCDC77 | rs10774191 | 5.80E-08 | ##### | Brain - Hypothalamus | 440788 | -59212 | 940788 | Yes |
| ENSG00000120647.9 | CCDC77 | rs61916593 | 5.80E-08 | ##### | Brain - Hypothalamus | 440788 | -59212 | 940788 | Yes |
| ENSG00000120647.9 | CCDC77 | rs10848961 | 5.80E-08 | ##### | Brain - Hypothalamus | 440788 | -59212 | 940788 | Yes |
| ENSG00000120647.9 | CCDC77 | rs4980886 | 5.90E-08 | ##### | Brain - Nucleus accumbens (b | 440788 | -59212 | 940788 | Yes |
| ENSG00000120647.9 | CCDC77 | rs2369280 | 5.90E-08 | ##### | Brain - Cortex | 440788 | -59212 | 940788 | Yes |

| | | | | | | | | | |
|---|---|---|---|---|---|---|---|---|---|
| ENSG00000120647.9 | CCDC77 | rs17800720 | 6.00E-08 | ##### | Brain - Cerebellar Hemisphere | 440788 | -59212 | 940788 | Yes |
| ENSG00000120647.9 | CCDC77 | rs3837515 | 6.00E-08 | ##### | Brain - Cerebellar Hemisphere | 440788 | -59212 | 940788 | Yes |
| ENSG00000120647.9 | CCDC77 | rs10848647 | 6.10E-08 | ##### | Brain - Frontal Cortex (BA9) | 440788 | -59212 | 940788 | Yes |
| ENSG00000120647.9 | CCDC77 | rs75894333 | 6.10E-08 | ##### | Brain - Cerebellum | 440788 | -59212 | 940788 | Yes |
| ENSG00000120647.9 | CCDC77 | rs11608938 | 6.10E-08 | ##### | Brain - Cerebellum | 440788 | -59212 | 940788 | Yes |
| ENSG00000120647.9 | CCDC77 | rs7965038 | 6.10E-08 | ##### | Brain - Cerebellum | 440788 | -59212 | 940788 | Yes |
| ENSG00000120647.9 | CCDC77 | rs7307606 | 6.10E-08 | ##### | Brain - Cerebellum | 440788 | -59212 | 940788 | Yes |
| ENSG00000120647.9 | CCDC77 | rs2041299 | 6.10E-08 | ##### | Brain - Cerebellum | 440788 | -59212 | 940788 | Yes |
| ENSG00000120647.9 | CCDC77 | rs10661251 | 6.10E-08 | ##### | Brain - Cerebellum | 440788 | -59212 | 940788 | Yes |
| ENSG00000120647.9 | CCDC77 | rs1011699 | 6.10E-08 | ##### | Brain - Cerebellum | 440788 | -59212 | 940788 | Yes |
| ENSG00000120647.9 | CCDC77 | rs6489475 | 6.10E-08 | ##### | Brain - Cerebellum | 440788 | -59212 | 940788 | Yes |
| ENSG00000120647.9 | CCDC77 | rs10774154 | 6.10E-08 | ##### | Brain - Cerebellum | 440788 | -59212 | 940788 | Yes |
| ENSG00000120647.9 | CCDC77 | rs11062930 | 6.10E-08 | ##### | Brain - Frontal Cortex (BA9) | 440788 | -59212 | 940788 | Yes |
| ENSG00000120647.9 | CCDC77 | rs10082788 | 6.20E-08 | ##### | Brain - Nucleus accumbens (b | 440788 | -59212 | 940788 | Yes |
| ENSG00000120647.9 | CCDC77 | rs4980874 | 6.20E-08 | ##### | Brain - Spinal cord (cervical c- | 440788 | -59212 | 940788 | Yes |
| ENSG00000120647.9 | CCDC77 | rs495316 | 6.30E-08 | ##### | Brain - Cortex | 440788 | -59212 | 940788 | Yes |
| ENSG00000120647.9 | CCDC77 | rs2041298 | 6.30E-08 | ##### | Brain - Caudate (basal ganglia | 440788 | -59212 | 940788 | Yes |
| ENSG00000120647.9 | CCDC77 | rs482514 | 6.40E-08 | ##### | Brain - Cortex | 440788 | -59212 | 940788 | Yes |
| ENSG00000120647.9 | CCDC77 | rs7315852 | 6.40E-08 | ##### | Brain - Cortex | 440788 | -59212 | 940788 | Yes |
| ENSG00000120647.9 | CCDC77 | rs2284334 | 6.40E-08 | ##### | Brain - Cortex | 440788 | -59212 | 940788 | Yes |
| ENSG00000120647.9 | CCDC77 | rs720128 | 6.40E-08 | ##### | Brain - Cortex | 440788 | -59212 | 940788 | Yes |
| ENSG00000120647.9 | CCDC77 | rs9739950 | 6.40E-08 | ##### | Brain - Cortex | 440788 | -59212 | 940788 | Yes |
| ENSG00000120647.9 | CCDC77 | rs12311569 | 6.40E-08 | ##### | Brain - Cortex | 440788 | -59212 | 940788 | Yes |
| ENSG00000120647.9 | CCDC77 | rs10848877 | 6.40E-08 | ##### | Brain - Cortex | 440788 | -59212 | 940788 | Yes |
| ENSG00000120647.9 | CCDC77 | rs11062929 | 6.70E-08 | ##### | Brain - Caudate (basal ganglia | 440788 | -59212 | 940788 | Yes |
| ENSG00000120647.9 | CCDC77 | rs486098 | 6.70E-08 | ##### | Brain - Cerebellum | 440788 | -59212 | 940788 | Yes |
| ENSG00000120647.9 | CCDC77 | rs1541463 | 6.90E-08 | ##### | Brain - Cerebellum | 440788 | -59212 | 940788 | Yes |
| ENSG00000120647.9 | CCDC77 | rs4980878 | 6.90E-08 | ##### | Brain - Anterior cingulate cort | 440788 | -59212 | 940788 | Yes |
| ENSG00000120647.9 | CCDC77 | rs3764876 | 6.90E-08 | ##### | Brain - Frontal Cortex (BA9) | 440788 | -59212 | 940788 | Yes |
| ENSG00000120647.9 | CCDC77 | rs10774191 | 6.90E-08 | ##### | Brain - Frontal Cortex (BA9) | 440788 | -59212 | 940788 | Yes |
| ENSG00000120647.9 | CCDC77 | rs61916593 | 6.90E-08 | ##### | Brain - Frontal Cortex (BA9) | 440788 | -59212 | 940788 | Yes |
| ENSG00000120647.9 | CCDC77 | rs4140962 | 7.20E-08 | ##### | Brain - Cerebellar Hemisphere | 440788 | -59212 | 940788 | Yes |

| | | | | | | | | | |
|---|---|---|---|---|---|---|---|---|---|
| ENSG00000120647.9 | CCDC77 | rs1155942 | 7.30E-08 | ##### | Brain - Nucleus accumbens (b | 440788 | -59212 | 940788 | Yes |
| ENSG00000120647.9 | CCDC77 | rs7295011 | 7.60E-08 | ##### | Brain - Cerebellar Hemisphere | 440788 | -59212 | 940788 | Yes |
| ENSG00000120647.9 | CCDC77 | rs4980878 | 7.60E-08 | ##### | Brain - Nucleus accumbens (b | 440788 | -59212 | 940788 | Yes |
| ENSG00000120647.9 | CCDC77 | rs764614 | 7.60E-08 | ##### | Brain - Nucleus accumbens (b | 440788 | -59212 | 940788 | Yes |
| ENSG00000120647.9 | CCDC77 | rs10848650 | 7.80E-08 | ##### | Brain - Hypothalamus | 440788 | -59212 | 940788 | Yes |
| ENSG00000120647.9 | CCDC77 | rs11062930 | 8.00E-08 | ##### | Brain - Cerebellar Hemisphere | 440788 | -59212 | 940788 | Yes |
| ENSG00000120647.9 | CCDC77 | rs7969761 | 8.40E-08 | ##### | Brain - Caudate (basal ganglia | 440788 | -59212 | 940788 | Yes |
| ENSG00000120647.9 | CCDC77 | rs6489523 | 8.50E-08 | ##### | Brain - Spinal cord (cervical c- | 440788 | -59212 | 940788 | Yes |
| ENSG00000120647.9 | CCDC77 | rs79839498 | 8.60E-08 | ##### | Brain - Cortex | 440788 | -59212 | 940788 | Yes |
| ENSG00000120647.9 | CCDC77 | rs372437080 | 8.60E-08 | ##### | Brain - Nucleus accumbens (b | 440788 | -59212 | 940788 | Yes |
| ENSG00000120647.9 | CCDC77 | rs7963793 | 8.70E-08 | ##### | Brain - Spinal cord (cervical c- | 440788 | -59212 | 940788 | Yes |
| ENSG00000120647.9 | CCDC77 | rs79839498 | 8.70E-08 | ##### | Brain - Cerebellum | 440788 | -59212 | 940788 | Yes |
| ENSG00000120647.9 | CCDC77 | rs1155942 | 8.80E-08 | ##### | Brain - Cerebellum | 440788 | -59212 | 940788 | Yes |
| ENSG00000120647.9 | CCDC77 | rs113574411 | 8.80E-08 | ##### | Brain - Nucleus accumbens (b | 440788 | -59212 | 940788 | Yes |
| ENSG00000120647.9 | CCDC77 | rs479840 | 8.80E-08 | ##### | Brain - Nucleus accumbens (b | 440788 | -59212 | 940788 | Yes |
| ENSG00000120647.9 | CCDC77 | rs479803 | 8.80E-08 | ##### | Brain - Nucleus accumbens (b | 440788 | -59212 | 940788 | Yes |
| ENSG00000120647.9 | CCDC77 | rs72472276 | 8.80E-08 | ##### | Brain - Nucleus accumbens (b | 440788 | -59212 | 940788 | Yes |
| ENSG00000120647.9 | CCDC77 | rs4980877 | 8.80E-08 | ##### | Brain - Nucleus accumbens (b | 440788 | -59212 | 940788 | Yes |
| ENSG00000120647.9 | CCDC77 | rs7963309 | 8.80E-08 | ##### | Brain - Nucleus accumbens (b | 440788 | -59212 | 940788 | Yes |
| ENSG00000120647.9 | CCDC77 | rs7134353 | 8.80E-08 | ##### | Brain - Nucleus accumbens (b | 440788 | -59212 | 940788 | Yes |
| ENSG00000120647.9 | CCDC77 | rs11374245 | 8.80E-08 | ##### | Brain - Nucleus accumbens (b | 440788 | -59212 | 940788 | Yes |
| ENSG00000120647.9 | CCDC77 | rs16929744 | 8.80E-08 | ##### | Brain - Nucleus accumbens (b | 440788 | -59212 | 940788 | Yes |
| ENSG00000120647.9 | CCDC77 | rs524468 | 8.90E-08 | ##### | Brain - Cortex | 440788 | -59212 | 940788 | Yes |
| ENSG00000120647.9 | CCDC77 | rs2052279 | 8.90E-08 | ##### | Brain - Cortex | 440788 | -59212 | 940788 | Yes |
| ENSG00000120647.9 | CCDC77 | rs78813329 | 8.90E-08 | ##### | Brain - Cortex | 440788 | -59212 | 940788 | Yes |
| ENSG00000120647.9 | CCDC77 | rs3753157 | 8.90E-08 | ##### | Brain - Cortex | 440788 | -59212 | 940788 | Yes |
| ENSG00000120647.9 | CCDC77 | rs4980883 | 8.90E-08 | ##### | Brain - Cortex | 440788 | -59212 | 940788 | Yes |
| ENSG00000120647.9 | CCDC77 | rs137910445 | 8.90E-08 | ##### | Brain - Cortex | 440788 | -59212 | 940788 | Yes |
| ENSG00000120647.9 | CCDC77 | rs11062537 | 8.90E-08 | ##### | Brain - Cortex | 440788 | -59212 | 940788 | Yes |
| ENSG00000120647.9 | CCDC77 | rs715230 | 8.90E-08 | ##### | Brain - Cortex | 440788 | -59212 | 940788 | Yes |
| ENSG00000120647.9 | CCDC77 | rs715229 | 8.90E-08 | ##### | Brain - Cortex | 440788 | -59212 | 940788 | Yes |
| ENSG00000120647.9 | CCDC77 | rs715228 | 8.90E-08 | ##### | Brain - Cortex | 440788 | -59212 | 940788 | Yes |

| | | | | | | | | | |
|---|---|---|---|---|---|---|---|---|---|
| ENSG00000120647.9 | CCDC77 | rs10848838 | 8.90E-08 | ##### | Brain - Cortex | 440788 | -59212 | 940788 | Yes |
| ENSG00000120647.9 | CCDC77 | rs12298843 | 8.90E-08 | ##### | Brain - Cortex | 440788 | -59212 | 940788 | Yes |
| ENSG00000120647.9 | CCDC77 | rs1207324 | 9.20E-08 | ##### | Brain - Anterior cingulate cort | 440788 | -59212 | 940788 | Yes |
| ENSG00000120647.9 | CCDC77 | rs576159 | 9.40E-08 | ##### | Brain - Putamen (basal gangli | 440788 | -59212 | 940788 | Yes |
| ENSG00000120647.9 | CCDC77 | rs2284332 | 9.50E-08 | ##### | Brain - Nucleus accumbens (b | 440788 | -59212 | 940788 | Yes |
| ENSG00000120647.9 | CCDC77 | rs2229351 | 9.50E-08 | ##### | Brain - Nucleus accumbens (b | 440788 | -59212 | 940788 | Yes |
| ENSG00000120647.9 | CCDC77 | rs495316 | 9.50E-08 | ##### | Brain - Cerebellum | 440788 | -59212 | 940788 | Yes |
| ENSG00000120647.9 | CCDC77 | rs11613357 | 9.60E-08 | ##### | Brain - Spinal cord (cervical c- | 440788 | -59212 | 940788 | Yes |
| ENSG00000120647.9 | CCDC77 | rs10848647 | 9.60E-08 | ##### | Brain - Hypothalamus | 440788 | -59212 | 940788 | Yes |
| ENSG00000120647.9 | CCDC77 | rs2300131 | 9.70E-08 | ##### | Brain - Nucleus accumbens (b | 440788 | -59212 | 940788 | Yes |
| ENSG00000120647.9 | CCDC77 | rs7303180 | 9.80E-08 | ##### | Brain - Nucleus accumbens (b | 440788 | -59212 | 940788 | Yes |
| ENSG00000120647.9 | CCDC77 | rs3815558 | 1.00E-07 | ##### | Brain - Cortex | 440788 | -59212 | 940788 | Yes |
| ENSG00000120647.9 | CCDC77 | rs78998485 | 1.00E-07 | ##### | Brain - Cortex | 440788 | -59212 | 940788 | Yes |
| ENSG00000120647.9 | CCDC77 | rs7135768 | 1.00E-07 | ##### | Brain - Cortex | 440788 | -59212 | 940788 | Yes |
| ENSG00000120647.9 | CCDC77 | rs6489450 | 1.00E-07 | ##### | Brain - Cortex | 440788 | -59212 | 940788 | Yes |
| ENSG00000120647.9 | CCDC77 | rs4980822 | 1.00E-07 | ##### | Brain - Caudate (basal ganglia | 440788 | -59212 | 940788 | Yes |
| ENSG00000120647.9 | CCDC77 | rs797765 | 1.00E-07 | ##### | Brain - Nucleus accumbens (b | 440788 | -59212 | 940788 | Yes |
| ENSG00000120647.9 | CCDC77 | rs2300131 | 1.10E-07 | ##### | Brain - Spinal cord (cervical c- | 440788 | -59212 | 940788 | Yes |
| ENSG00000120647.9 | CCDC77 | rs11062813 | 1.10E-07 | ##### | Brain - Substantia nigra | 440788 | -59212 | 940788 | Yes |
| ENSG00000120647.9 | CCDC77 | rs2052279 | 1.10E-07 | ##### | Brain - Hypothalamus | 440788 | -59212 | 940788 | Yes |
| ENSG00000120647.9 | CCDC77 | rs78813329 | 1.10E-07 | ##### | Brain - Hypothalamus | 440788 | -59212 | 940788 | Yes |
| ENSG00000120647.9 | CCDC77 | rs3753157 | 1.10E-07 | ##### | Brain - Hypothalamus | 440788 | -59212 | 940788 | Yes |
| ENSG00000120647.9 | CCDC77 | rs4980883 | 1.10E-07 | ##### | Brain - Hypothalamus | 440788 | -59212 | 940788 | Yes |
| ENSG00000120647.9 | CCDC77 | rs137910445 | 1.10E-07 | ##### | Brain - Hypothalamus | 440788 | -59212 | 940788 | Yes |
| ENSG00000120647.9 | CCDC77 | rs11062537 | 1.10E-07 | ##### | Brain - Hypothalamus | 440788 | -59212 | 940788 | Yes |
| ENSG00000120647.9 | CCDC77 | rs715230 | 1.10E-07 | ##### | Brain - Hypothalamus | 440788 | -59212 | 940788 | Yes |
| ENSG00000120647.9 | CCDC77 | rs715229 | 1.10E-07 | ##### | Brain - Hypothalamus | 440788 | -59212 | 940788 | Yes |
| ENSG00000120647.9 | CCDC77 | rs715228 | 1.10E-07 | ##### | Brain - Hypothalamus | 440788 | -59212 | 940788 | Yes |
| ENSG00000120647.9 | CCDC77 | rs10848838 | 1.10E-07 | ##### | Brain - Hypothalamus | 440788 | -59212 | 940788 | Yes |
| ENSG00000120647.9 | CCDC77 | rs7132845 | 1.10E-07 | ##### | Brain - Cerebellum | 440788 | -59212 | 940788 | Yes |
| ENSG00000120647.9 | CCDC77 | rs11063005 | 1.10E-07 | ##### | Brain - Caudate (basal ganglia | 440788 | -59212 | 940788 | Yes |
| ENSG00000120647.9 | CCDC77 | rs11063010 | 1.10E-07 | ##### | Brain - Caudate (basal ganglia | 440788 | -59212 | 940788 | Yes |

| | | | | | | | | | |
|---|---|---|---|---|---|---|---|---|---|
| ENSG00000120647.9 | CCDC77 | rs10774169 | 1.10E-07 | ##### | Brain - Cortex | 440788 | -59212 | 940788 | Yes |
| ENSG00000120647.9 | CCDC77 | rs12301816 | 1.10E-07 | ##### | Brain - Cortex | 440788 | -59212 | 940788 | Yes |
| ENSG00000120647.9 | CCDC77 | rs4980875 | 1.10E-07 | ##### | Brain - Nucleus accumbens (b | 440788 | -59212 | 940788 | Yes |
| ENSG00000120647.9 | CCDC77 | rs61906995 | 1.10E-07 | ##### | Brain - Nucleus accumbens (b | 440788 | -59212 | 940788 | Yes |
| ENSG00000120647.9 | CCDC77 | rs12582850 | 1.10E-07 | ##### | Brain - Nucleus accumbens (b | 440788 | -59212 | 940788 | Yes |
| ENSG00000120647.9 | CCDC77 | rs7298019 | 1.10E-07 | ##### | Brain - Nucleus accumbens (b | 440788 | -59212 | 940788 | Yes |
| ENSG00000120647.9 | CCDC77 | rs2098094 | 1.10E-07 | ##### | Brain - Nucleus accumbens (b | 440788 | -59212 | 940788 | Yes |
| ENSG00000120647.9 | CCDC77 | rs2159364 | 1.10E-07 | ##### | Brain - Nucleus accumbens (b | 440788 | -59212 | 940788 | Yes |
| ENSG00000120647.9 | CCDC77 | rs10848868 | 1.10E-07 | ##### | Brain - Nucleus accumbens (b | 440788 | -59212 | 940788 | Yes |
| ENSG00000120647.9 | CCDC77 | rs1860360 | 1.10E-07 | ##### | Brain - Nucleus accumbens (b | 440788 | -59212 | 940788 | Yes |
| ENSG00000120647.9 | CCDC77 | rs2080402 | 1.10E-07 | ##### | Brain - Cerebellar Hemisphere | 440788 | -59212 | 940788 | Yes |
| ENSG00000120647.9 | CCDC77 | rs6489477 | 1.10E-07 | ##### | Brain - Nucleus accumbens (b | 440788 | -59212 | 940788 | Yes |
| ENSG00000120647.9 | CCDC77 | rs6489478 | 1.10E-07 | ##### | Brain - Nucleus accumbens (b | 440788 | -59212 | 940788 | Yes |
| ENSG00000120647.9 | CCDC77 | rs2300131 | 1.10E-07 | ##### | Brain - Cortex | 440788 | -59212 | 940788 | Yes |
| ENSG00000120647.9 | CCDC77 | rs11062813 | 1.20E-07 | ##### | Brain - Hippocampus | 440788 | -59212 | 940788 | Yes |
| ENSG00000120647.9 | CCDC77 | rs2052279 | 1.20E-07 | ##### | Brain - Cerebellum | 440788 | -59212 | 940788 | Yes |
| ENSG00000120647.9 | CCDC77 | rs78813329 | 1.20E-07 | ##### | Brain - Cerebellum | 440788 | -59212 | 940788 | Yes |
| ENSG00000120647.9 | CCDC77 | rs3753157 | 1.20E-07 | ##### | Brain - Cerebellum | 440788 | -59212 | 940788 | Yes |
| ENSG00000120647.9 | CCDC77 | rs4980883 | 1.20E-07 | ##### | Brain - Cerebellum | 440788 | -59212 | 940788 | Yes |
| ENSG00000120647.9 | CCDC77 | rs137910445 | 1.20E-07 | ##### | Brain - Cerebellum | 440788 | -59212 | 940788 | Yes |
| ENSG00000120647.9 | CCDC77 | rs11062537 | 1.20E-07 | ##### | Brain - Cerebellum | 440788 | -59212 | 940788 | Yes |
| ENSG00000120647.9 | CCDC77 | rs715230 | 1.20E-07 | ##### | Brain - Cerebellum | 440788 | -59212 | 940788 | Yes |
| ENSG00000120647.9 | CCDC77 | rs715229 | 1.20E-07 | ##### | Brain - Cerebellum | 440788 | -59212 | 940788 | Yes |
| ENSG00000120647.9 | CCDC77 | rs715228 | 1.20E-07 | ##### | Brain - Cerebellum | 440788 | -59212 | 940788 | Yes |
| ENSG00000120647.9 | CCDC77 | rs10848838 | 1.20E-07 | ##### | Brain - Cerebellum | 440788 | -59212 | 940788 | Yes |
| ENSG00000120647.9 | CCDC77 | rs4980892 | 1.20E-07 | ##### | Brain - Cortex | 440788 | -59212 | 940788 | Yes |
| ENSG00000120647.9 | CCDC77 | rs12298843 | 1.20E-07 | ##### | Brain - Hypothalamus | 440788 | -59212 | 940788 | Yes |
| ENSG00000120647.9 | CCDC77 | rs1048466 | 1.20E-07 | ##### | Brain - Caudate (basal ganglia | 440788 | -59212 | 940788 | Yes |
| ENSG00000120647.9 | CCDC77 | rs482514 | 1.20E-07 | ##### | Brain - Cerebellum | 440788 | -59212 | 940788 | Yes |
| ENSG00000120647.9 | CCDC77 | rs7315852 | 1.20E-07 | ##### | Brain - Cerebellum | 440788 | -59212 | 940788 | Yes |
| ENSG00000120647.9 | CCDC77 | rs2284334 | 1.20E-07 | ##### | Brain - Cerebellum | 440788 | -59212 | 940788 | Yes |
| ENSG00000120647.9 | CCDC77 | rs720128 | 1.20E-07 | ##### | Brain - Cerebellum | 440788 | -59212 | 940788 | Yes |

| | | | | | | | | | |
|---|---|---|---|---|---|---|---|---|---|
| ENSG00000120647.9 | CCDC77 | rs9739950 | 1.20E-07 | ##### | Brain - Cerebellum | 440788 | -59212 | 940788 | Yes |
| ENSG00000120647.9 | CCDC77 | rs12311569 | 1.20E-07 | ##### | Brain - Cerebellum | 440788 | -59212 | 940788 | Yes |
| ENSG00000120647.9 | CCDC77 | rs10848877 | 1.20E-07 | ##### | Brain - Cerebellum | 440788 | -59212 | 940788 | Yes |
| ENSG00000120647.9 | CCDC77 | rs28850111 | 1.20E-07 | ##### | Brain - Hypothalamus | 440788 | -59212 | 940788 | Yes |
| ENSG00000120647.9 | CCDC77 | rs4980886 | 1.20E-07 | ##### | Brain - Putamen (basal gangli | 440788 | -59212 | 940788 | Yes |
| ENSG00000120647.9 | CCDC77 | rs7138981 | 1.20E-07 | ##### | Brain - Putamen (basal gangli | 440788 | -59212 | 940788 | Yes |
| ENSG00000120647.9 | CCDC77 | rs9669094 | 1.20E-07 | ##### | Brain - Putamen (basal gangli | 440788 | -59212 | 940788 | Yes |
| ENSG00000120647.9 | CCDC77 | rs7132763 | 1.20E-07 | ##### | Brain - Putamen (basal gangli | 440788 | -59212 | 940788 | Yes |
| ENSG00000120647.9 | CCDC77 | rs2015554 | 1.20E-07 | ##### | Brain - Putamen (basal gangli | 440788 | -59212 | 940788 | Yes |
| ENSG00000120647.9 | CCDC77 | rs764614 | 1.30E-07 | ##### | Brain - Putamen (basal gangli | 440788 | -59212 | 940788 | Yes |
| ENSG00000120647.9 | CCDC77 | rs12298843 | 1.30E-07 | ##### | Brain - Cerebellum | 440788 | -59212 | 940788 | Yes |
| ENSG00000120647.9 | CCDC77 | rs2041298 | 1.30E-07 | ##### | Brain - Hypothalamus | 440788 | -59212 | 940788 | Yes |
| ENSG00000120647.9 | CCDC77 | rs507597 | 1.30E-07 | ##### | Brain - Substantia nigra | 440788 | -59212 | 940788 | Yes |
| ENSG00000120647.9 | CCDC77 | rs17800744 | 1.30E-07 | ##### | Brain - Substantia nigra | 440788 | -59212 | 940788 | Yes |
| ENSG00000120647.9 | CCDC77 | rs77854644 | 1.30E-07 | ##### | Brain - Substantia nigra | 440788 | -59212 | 940788 | Yes |
| ENSG00000120647.9 | CCDC77 | rs764614 | 1.30E-07 | ##### | Brain - Cerebellum | 440788 | -59212 | 940788 | Yes |
| ENSG00000120647.9 | CCDC77 | rs7132763 | 1.30E-07 | ##### | Brain - Cerebellum | 440788 | -59212 | 940788 | Yes |
| ENSG00000120647.9 | CCDC77 | rs7311740 | 1.30E-07 | ##### | Brain - Frontal Cortex (BA9) | 440788 | -59212 | 940788 | Yes |
| ENSG00000120647.9 | CCDC77 | rs2300133 | 1.40E-07 | ##### | Brain - Hypothalamus | 440788 | -59212 | 940788 | Yes |
| ENSG00000120647.9 | CCDC77 | rs576159 | 1.40E-07 | ##### | Brain - Nucleus accumbens (b | 440788 | -59212 | 940788 | Yes |
| ENSG00000120647.9 | CCDC77 | rs113574411 | 1.40E-07 | ##### | Brain - Putamen (basal gangli | 440788 | -59212 | 940788 | Yes |
| ENSG00000120647.9 | CCDC77 | rs479840 | 1.40E-07 | ##### | Brain - Putamen (basal gangli | 440788 | -59212 | 940788 | Yes |
| ENSG00000120647.9 | CCDC77 | rs479803 | 1.40E-07 | ##### | Brain - Putamen (basal gangli | 440788 | -59212 | 940788 | Yes |
| ENSG00000120647.9 | CCDC77 | rs77074951 | 1.40E-07 | ##### | Brain - Putamen (basal gangli | 440788 | -59212 | 940788 | Yes |
| ENSG00000120647.9 | CCDC77 | rs4980874 | 1.40E-07 | ##### | Brain - Putamen (basal gangli | 440788 | -59212 | 940788 | Yes |
| ENSG00000120647.9 | CCDC77 | rs2284333 | 1.40E-07 | ##### | Brain - Putamen (basal gangli | 440788 | -59212 | 940788 | Yes |
| ENSG00000120647.9 | CCDC77 | rs4980811 | 1.40E-07 | ##### | Brain - Putamen (basal gangli | 440788 | -59212 | 940788 | Yes |
| ENSG00000120647.9 | CCDC77 | rs72472276 | 1.40E-07 | ##### | Brain - Putamen (basal gangli | 440788 | -59212 | 940788 | Yes |
| ENSG00000120647.9 | CCDC77 | rs4980877 | 1.40E-07 | ##### | Brain - Putamen (basal gangli | 440788 | -59212 | 940788 | Yes |
| ENSG00000120647.9 | CCDC77 | rs7963309 | 1.40E-07 | ##### | Brain - Putamen (basal gangli | 440788 | -59212 | 940788 | Yes |
| ENSG00000120647.9 | CCDC77 | rs7134353 | 1.40E-07 | ##### | Brain - Putamen (basal gangli | 440788 | -59212 | 940788 | Yes |
| ENSG00000120647.9 | CCDC77 | rs11374245 | 1.40E-07 | ##### | Brain - Putamen (basal gangli | 440788 | -59212 | 940788 | Yes |

| | | | | | | | | | |
|---|---|---|---|---|---|---|---|---|---|
| ENSG00000120647.9 | CCDC77 | rs16929744 | 1.40E-07 | ##### | Brain - Putamen (basal gangli | 440788 | -59212 | 940788 | Yes |
| ENSG00000120647.9 | CCDC77 | rs1207322 | 1.40E-07 | ##### | Brain - Anterior cingulate cort | 440788 | -59212 | 940788 | Yes |
| ENSG00000120647.9 | CCDC77 | rs57022939 | 1.40E-07 | ##### | Brain - Anterior cingulate cort | 440788 | -59212 | 940788 | Yes |
| ENSG00000120647.9 | CCDC77 | rs4980875 | 1.40E-07 | ##### | Brain - Anterior cingulate cort | 440788 | -59212 | 940788 | Yes |
| ENSG00000120647.9 | CCDC77 | rs61906995 | 1.40E-07 | ##### | Brain - Anterior cingulate cort | 440788 | -59212 | 940788 | Yes |
| ENSG00000120647.9 | CCDC77 | rs4980813 | 1.40E-07 | ##### | Brain - Anterior cingulate cort | 440788 | -59212 | 940788 | Yes |
| ENSG00000120647.9 | CCDC77 | rs4980876 | 1.40E-07 | ##### | Brain - Anterior cingulate cort | 440788 | -59212 | 940788 | Yes |
| ENSG00000120647.9 | CCDC77 | rs12582850 | 1.40E-07 | ##### | Brain - Anterior cingulate cort | 440788 | -59212 | 940788 | Yes |
| ENSG00000120647.9 | CCDC77 | rs61906996 | 1.40E-07 | ##### | Brain - Anterior cingulate cort | 440788 | -59212 | 940788 | Yes |
| ENSG00000120647.9 | CCDC77 | rs4980814 | 1.40E-07 | ##### | Brain - Anterior cingulate cort | 440788 | -59212 | 940788 | Yes |
| ENSG00000120647.9 | CCDC77 | rs7298019 | 1.40E-07 | ##### | Brain - Anterior cingulate cort | 440788 | -59212 | 940788 | Yes |
| ENSG00000120647.9 | CCDC77 | rs2098094 | 1.40E-07 | ##### | Brain - Anterior cingulate cort | 440788 | -59212 | 940788 | Yes |
| ENSG00000120647.9 | CCDC77 | rs2159364 | 1.40E-07 | ##### | Brain - Anterior cingulate cort | 440788 | -59212 | 940788 | Yes |
| ENSG00000120647.9 | CCDC77 | rs7977462 | 1.40E-07 | ##### | Brain - Anterior cingulate cort | 440788 | -59212 | 940788 | Yes |
| ENSG00000120647.9 | CCDC77 | rs10744616 | 1.40E-07 | ##### | Brain - Anterior cingulate cort | 440788 | -59212 | 940788 | Yes |
| ENSG00000120647.9 | CCDC77 | rs10848868 | 1.40E-07 | ##### | Brain - Anterior cingulate cort | 440788 | -59212 | 940788 | Yes |
| ENSG00000120647.9 | CCDC77 | rs10774150 | 1.40E-07 | ##### | Brain - Anterior cingulate cort | 440788 | -59212 | 940788 | Yes |
| ENSG00000120647.9 | CCDC77 | rs1860360 | 1.40E-07 | ##### | Brain - Anterior cingulate cort | 440788 | -59212 | 940788 | Yes |
| ENSG00000120647.9 | CCDC77 | rs3831759 | 1.40E-07 | ##### | Brain - Anterior cingulate cort | 440788 | -59212 | 940788 | Yes |
| ENSG00000120647.9 | CCDC77 | rs580420 | 1.40E-07 | ##### | Brain - Hippocampus | 440788 | -59212 | 940788 | Yes |
| ENSG00000120647.9 | CCDC77 | rs495973 | 1.40E-07 | ##### | Brain - Cortex | 440788 | -59212 | 940788 | Yes |
| ENSG00000120647.9 | CCDC77 | rs2300131 | 1.40E-07 | ##### | Brain - Cerebellum | 440788 | -59212 | 940788 | Yes |
| ENSG00000120647.9 | CCDC77 | rs56042953 | 1.40E-07 | ##### | Brain - Caudate (basal ganglia | 440788 | -59212 | 940788 | Yes |
| ENSG00000120647.9 | CCDC77 | rs10161265 | 1.40E-07 | ##### | Brain - Hypothalamus | 440788 | -59212 | 940788 | Yes |
| ENSG00000120647.9 | CCDC77 | rs4980897 | 1.40E-07 | ##### | Brain - Hypothalamus | 440788 | -59212 | 940788 | Yes |
| ENSG00000120647.9 | CCDC77 | rs11062930 | 1.50E-07 | ##### | Brain - Spinal cord (cervical c- | 440788 | -59212 | 940788 | Yes |
| ENSG00000120647.9 | CCDC77 | rs10161265 | 1.50E-07 | ##### | Brain - Spinal cord (cervical c- | 440788 | -59212 | 940788 | Yes |
| ENSG00000120647.9 | CCDC77 | rs4980897 | 1.50E-07 | ##### | Brain - Spinal cord (cervical c- | 440788 | -59212 | 940788 | Yes |
| ENSG00000120647.9 | CCDC77 | rs16930719 | 1.50E-07 | ##### | Brain - Frontal Cortex (BA9) | 440788 | -59212 | 940788 | Yes |
| ENSG00000120647.9 | CCDC77 | rs2369281 | 1.50E-07 | ##### | Brain - Hypothalamus | 440788 | -59212 | 940788 | Yes |
| ENSG00000120647.9 | CCDC77 | rs56042953 | 1.60E-07 | ##### | Brain - Cerebellar Hemisphere | 440788 | -59212 | 940788 | Yes |
| ENSG00000120647.9 | CCDC77 | rs579526 | 1.60E-07 | ##### | Brain - Nucleus accumbens (b | 440788 | -59212 | 940788 | Yes |

| | | | | | | | | | |
|---|---|---|---|---|---|---|---|---|---|
| ENSG00000120647.9 | CCDC77 | rs2369280 | 1.60E-07 | ##### | Brain - Cerebellum | 440788 | -59212 | 940788 | Yes |
| ENSG00000120647.9 | CCDC77 | rs12372234 | 1.60E-07 | ##### | Brain - Spinal cord (cervical c- | 440788 | -59212 | 940788 | Yes |
| ENSG00000120647.9 | CCDC77 | rs75894333 | 1.70E-07 | ##### | Brain - Nucleus accumbens (b | 440788 | -59212 | 940788 | Yes |
| ENSG00000120647.9 | CCDC77 | rs11608938 | 1.70E-07 | ##### | Brain - Nucleus accumbens (b | 440788 | -59212 | 940788 | Yes |
| ENSG00000120647.9 | CCDC77 | rs7965038 | 1.70E-07 | ##### | Brain - Nucleus accumbens (b | 440788 | -59212 | 940788 | Yes |
| ENSG00000120647.9 | CCDC77 | rs7307606 | 1.70E-07 | ##### | Brain - Nucleus accumbens (b | 440788 | -59212 | 940788 | Yes |
| ENSG00000120647.9 | CCDC77 | rs10661251 | 1.70E-07 | ##### | Brain - Nucleus accumbens (b | 440788 | -59212 | 940788 | Yes |
| ENSG00000120647.9 | CCDC77 | rs1011699 | 1.70E-07 | ##### | Brain - Nucleus accumbens (b | 440788 | -59212 | 940788 | Yes |
| ENSG00000120647.9 | CCDC77 | rs6489475 | 1.70E-07 | ##### | Brain - Nucleus accumbens (b | 440788 | -59212 | 940788 | Yes |
| ENSG00000120647.9 | CCDC77 | rs10774154 | 1.70E-07 | ##### | Brain - Nucleus accumbens (b | 440788 | -59212 | 940788 | Yes |
| ENSG00000120647.9 | CCDC77 | rs10774152 | 1.70E-07 | ##### | Brain - Putamen (basal gangli | 440788 | -59212 | 940788 | Yes |
| ENSG00000120647.9 | CCDC77 | rs10774153 | 1.70E-07 | ##### | Brain - Putamen (basal gangli | 440788 | -59212 | 940788 | Yes |
| ENSG00000120647.9 | CCDC77 | rs4980811 | 1.70E-07 | ##### | Brain - Nucleus accumbens (b | 440788 | -59212 | 940788 | Yes |
| ENSG00000120647.9 | CCDC77 | rs507597 | 1.70E-07 | ##### | Brain - Hippocampus | 440788 | -59212 | 940788 | Yes |
| ENSG00000120647.9 | CCDC77 | rs17800744 | 1.70E-07 | ##### | Brain - Hippocampus | 440788 | -59212 | 940788 | Yes |
| ENSG00000120647.9 | CCDC77 | rs77854644 | 1.70E-07 | ##### | Brain - Hippocampus | 440788 | -59212 | 940788 | Yes |
| ENSG00000120647.9 | CCDC77 | rs77074951 | 1.80E-07 | ##### | Brain - Nucleus accumbens (b | 440788 | -59212 | 940788 | Yes |
| ENSG00000120647.9 | CCDC77 | rs2191159 | 1.80E-07 | ##### | Brain - Caudate (basal ganglia | 440788 | -59212 | 940788 | Yes |
| ENSG00000120647.9 | CCDC77 | rs7963793 | 1.80E-07 | ##### | Brain - Putamen (basal gangli | 440788 | -59212 | 940788 | Yes |
| ENSG00000120647.9 | CCDC77 | rs10848814 | 1.80E-07 | ##### | Brain - Putamen (basal gangli | 440788 | -59212 | 940788 | Yes |
| ENSG00000120647.9 | CCDC77 | rs10848815 | 1.80E-07 | ##### | Brain - Putamen (basal gangli | 440788 | -59212 | 940788 | Yes |
| ENSG00000120647.9 | CCDC77 | rs887311 | 1.80E-07 | ##### | Brain - Putamen (basal gangli | 440788 | -59212 | 940788 | Yes |
| ENSG00000120647.9 | CCDC77 | rs75894333 | 1.80E-07 | ##### | Brain - Anterior cingulate cort | 440788 | -59212 | 940788 | Yes |
| ENSG00000120647.9 | CCDC77 | rs507597 | 1.80E-07 | ##### | Brain - Anterior cingulate cort | 440788 | -59212 | 940788 | Yes |
| ENSG00000120647.9 | CCDC77 | rs17800744 | 1.80E-07 | ##### | Brain - Anterior cingulate cort | 440788 | -59212 | 940788 | Yes |
| ENSG00000120647.9 | CCDC77 | rs77854644 | 1.80E-07 | ##### | Brain - Anterior cingulate cort | 440788 | -59212 | 940788 | Yes |
| ENSG00000120647.9 | CCDC77 | rs11608938 | 1.80E-07 | ##### | Brain - Anterior cingulate cort | 440788 | -59212 | 940788 | Yes |
| ENSG00000120647.9 | CCDC77 | rs7965038 | 1.80E-07 | ##### | Brain - Anterior cingulate cort | 440788 | -59212 | 940788 | Yes |
| ENSG00000120647.9 | CCDC77 | rs7307606 | 1.80E-07 | ##### | Brain - Anterior cingulate cort | 440788 | -59212 | 940788 | Yes |
| ENSG00000120647.9 | CCDC77 | rs10661251 | 1.80E-07 | ##### | Brain - Anterior cingulate cort | 440788 | -59212 | 940788 | Yes |
| ENSG00000120647.9 | CCDC77 | rs1011699 | 1.80E-07 | ##### | Brain - Anterior cingulate cort | 440788 | -59212 | 940788 | Yes |
| ENSG00000120647.9 | CCDC77 | rs6489475 | 1.80E-07 | ##### | Brain - Anterior cingulate cort | 440788 | -59212 | 940788 | Yes |

| | | | | | | | | | |
|---|---|---|---|---|---|---|---|---|---|
| ENSG00000120647.9 | CCDC77 | rs10774154 | 1.80E-07 | ##### | Brain - Anterior cingulate cort | 440788 | -59212 | 940788 | Yes |
| ENSG00000120647.9 | CCDC77 | rs11062357 | 1.80E-07 | ##### | Brain - Caudate (basal ganglia | 440788 | -59212 | 940788 | Yes |
| ENSG00000120647.9 | CCDC77 | rs57022939 | 1.90E-07 | ##### | Brain - Nucleus accumbens (b | 440788 | -59212 | 940788 | Yes |
| ENSG00000120647.9 | CCDC77 | rs3753158 | 1.90E-07 | ##### | Brain - Nucleus accumbens (b | 440788 | -59212 | 940788 | Yes |
| ENSG00000120647.9 | CCDC77 | rs1207324 | 1.90E-07 | ##### | Brain - Nucleus accumbens (b | 440788 | -59212 | 940788 | Yes |
| ENSG00000120647.9 | CCDC77 | rs61907026 | 1.90E-07 | ##### | Brain - Cortex | 440788 | -59212 | 940788 | Yes |
| ENSG00000120647.9 | CCDC77 | rs61907030 | 1.90E-07 | ##### | Brain - Cortex | 440788 | -59212 | 940788 | Yes |
| ENSG00000120647.9 | CCDC77 | rs61907031 | 1.90E-07 | ##### | Brain - Cortex | 440788 | -59212 | 940788 | Yes |
| ENSG00000120647.9 | CCDC77 | rs11615564 | 1.90E-07 | ##### | Brain - Cortex | 440788 | -59212 | 940788 | Yes |
| ENSG00000120647.9 | CCDC77 | rs2369281 | 1.90E-07 | ##### | Brain - Frontal Cortex (BA9) | 440788 | -59212 | 940788 | Yes |
| ENSG00000120647.9 | CCDC77 | rs7303180 | 2.00E-07 | ##### | Brain - Putamen (basal gangli | 440788 | -59212 | 940788 | Yes |
| ENSG00000120647.9 | CCDC77 | rs7136116 | 2.00E-07 | ##### | Brain - Nucleus accumbens (b | 440788 | -59212 | 940788 | Yes |
| ENSG00000120647.9 | CCDC77 | rs11610457 | 2.00E-07 | ##### | Brain - Nucleus accumbens (b | 440788 | -59212 | 940788 | Yes |
| ENSG00000120647.9 | CCDC77 | rs11062813 | 2.00E-07 | ##### | Brain - Anterior cingulate cort | 440788 | -59212 | 940788 | Yes |
| ENSG00000120647.9 | CCDC77 | rs6489523 | 2.00E-07 | ##### | Brain - Caudate (basal ganglia | 440788 | -59212 | 940788 | Yes |
| ENSG00000120647.9 | CCDC77 | rs11063005 | 2.00E-07 | ##### | Brain - Cerebellar Hemisphere | 440788 | -59212 | 940788 | Yes |
| ENSG00000120647.9 | CCDC77 | rs11063010 | 2.00E-07 | ##### | Brain - Cerebellar Hemisphere | 440788 | -59212 | 940788 | Yes |
| ENSG00000120647.9 | CCDC77 | rs7132845 | 2.00E-07 | ##### | Brain - Nucleus accumbens (b | 440788 | -59212 | 940788 | Yes |
| ENSG00000120647.9 | CCDC77 | rs3213750 | 2.00E-07 | ##### | Brain - Spinal cord (cervical c- | 440788 | -59212 | 940788 | Yes |
| ENSG00000120647.9 | CCDC77 | rs3753156 | 2.00E-07 | ##### | Brain - Spinal cord (cervical c- | 440788 | -59212 | 940788 | Yes |
| ENSG00000120647.9 | CCDC77 | rs12581260 | 2.00E-07 | ##### | Brain - Spinal cord (cervical c- | 440788 | -59212 | 940788 | Yes |
| ENSG00000120647.9 | CCDC77 | rs4980881 | 2.00E-07 | ##### | Brain - Spinal cord (cervical c- | 440788 | -59212 | 940788 | Yes |
| ENSG00000120647.9 | CCDC77 | rs16929783 | 2.00E-07 | ##### | Brain - Spinal cord (cervical c- | 440788 | -59212 | 940788 | Yes |
| ENSG00000120647.9 | CCDC77 | rs10848814 | 2.00E-07 | ##### | Brain - Spinal cord (cervical c- | 440788 | -59212 | 940788 | Yes |
| ENSG00000120647.9 | CCDC77 | rs10848815 | 2.00E-07 | ##### | Brain - Spinal cord (cervical c- | 440788 | -59212 | 940788 | Yes |
| ENSG00000120647.9 | CCDC77 | rs887311 | 2.00E-07 | ##### | Brain - Spinal cord (cervical c- | 440788 | -59212 | 940788 | Yes |
| ENSG00000120647.9 | CCDC77 | rs2300133 | 2.10E-07 | ##### | Brain - Cortex | 440788 | -59212 | 940788 | Yes |
| ENSG00000120647.9 | CCDC77 | rs3213750 | 2.10E-07 | ##### | Brain - Putamen (basal gangli | 440788 | -59212 | 940788 | Yes |
| ENSG00000120647.9 | CCDC77 | rs3753156 | 2.10E-07 | ##### | Brain - Putamen (basal gangli | 440788 | -59212 | 940788 | Yes |
| ENSG00000120647.9 | CCDC77 | rs12581260 | 2.10E-07 | ##### | Brain - Putamen (basal gangli | 440788 | -59212 | 940788 | Yes |
| ENSG00000120647.9 | CCDC77 | rs4980881 | 2.10E-07 | ##### | Brain - Putamen (basal gangli | 440788 | -59212 | 940788 | Yes |
| ENSG00000120647.9 | CCDC77 | rs16929783 | 2.10E-07 | ##### | Brain - Putamen (basal gangli | 440788 | -59212 | 940788 | Yes |

| | | | | | | | | | |
|---|---|---|---|---|---|---|---|---|---|
| ENSG00000120647.9 | CCDC77 | rs10774192 | 2.10E-07 | ##### | Brain - Cerebellar Hemisphere | 440788 | -59212 | 940788 | Yes |
| ENSG00000120647.9 | CCDC77 | rs61907026 | 2.10E-07 | ##### | Brain - Cerebellum | 440788 | -59212 | 940788 | Yes |
| ENSG00000120647.9 | CCDC77 | rs61907030 | 2.10E-07 | ##### | Brain - Cerebellum | 440788 | -59212 | 940788 | Yes |
| ENSG00000120647.9 | CCDC77 | rs61907031 | 2.10E-07 | ##### | Brain - Cerebellum | 440788 | -59212 | 940788 | Yes |
| ENSG00000120647.9 | CCDC77 | rs11615564 | 2.10E-07 | ##### | Brain - Cerebellum | 440788 | -59212 | 940788 | Yes |
| ENSG00000120647.9 | CCDC77 | rs12301816 | 2.20E-07 | ##### | Brain - Putamen (basal gangli | 440788 | -59212 | 940788 | Yes |
| ENSG00000120647.9 | CCDC77 | rs7295011 | 2.30E-07 | ##### | Brain - Caudate (basal ganglia | 440788 | -59212 | 940788 | Yes |
| ENSG00000120647.9 | CCDC77 | rs10744561 | 2.30E-07 | ##### | Brain - Hypothalamus | 440788 | -59212 | 940788 | Yes |
| ENSG00000120647.9 | CCDC77 | rs4140962 | 2.30E-07 | ##### | Brain - Hypothalamus | 440788 | -59212 | 940788 | Yes |
| ENSG00000120647.9 | CCDC77 | rs7963793 | 2.30E-07 | ##### | Brain - Cortex | 440788 | -59212 | 940788 | Yes |
| ENSG00000120647.9 | CCDC77 | rs16930719 | 2.30E-07 | ##### | Brain - Spinal cord (cervical c- | 440788 | -59212 | 940788 | Yes |
| ENSG00000120647.9 | CCDC77 | rs7963793 | 2.30E-07 | ##### | Brain - Nucleus accumbens (b | 440788 | -59212 | 940788 | Yes |
| ENSG00000120647.9 | CCDC77 | rs10848814 | 2.30E-07 | ##### | Brain - Nucleus accumbens (b | 440788 | -59212 | 940788 | Yes |
| ENSG00000120647.9 | CCDC77 | rs10848815 | 2.30E-07 | ##### | Brain - Nucleus accumbens (b | 440788 | -59212 | 940788 | Yes |
| ENSG00000120647.9 | CCDC77 | rs887311 | 2.30E-07 | ##### | Brain - Nucleus accumbens (b | 440788 | -59212 | 940788 | Yes |
| ENSG00000120647.9 | CCDC77 | rs12298843 | 2.50E-07 | ##### | Brain - Spinal cord (cervical c- | 440788 | -59212 | 940788 | Yes |
| ENSG00000120647.9 | CCDC77 | rs495316 | 2.50E-07 | ##### | Brain - Nucleus accumbens (b | 440788 | -59212 | 940788 | Yes |
| ENSG00000120647.9 | CCDC77 | rs56042953 | 2.50E-07 | ##### | Brain - Frontal Cortex (BA9) | 440788 | -59212 | 940788 | Yes |
| ENSG00000120647.9 | CCDC77 | rs7963793 | 2.60E-07 | ##### | Brain - Cerebellum | 440788 | -59212 | 940788 | Yes |
| ENSG00000120647.9 | CCDC77 | rs1155942 | 2.60E-07 | ##### | Brain - Anterior cingulate cort | 440788 | -59212 | 940788 | Yes |
| ENSG00000120647.9 | CCDC77 | rs75894333 | 2.60E-07 | ##### | Brain - Substantia nigra | 440788 | -59212 | 940788 | Yes |
| ENSG00000120647.9 | CCDC77 | rs11608938 | 2.60E-07 | ##### | Brain - Substantia nigra | 440788 | -59212 | 940788 | Yes |
| ENSG00000120647.9 | CCDC77 | rs7965038 | 2.60E-07 | ##### | Brain - Substantia nigra | 440788 | -59212 | 940788 | Yes |
| ENSG00000120647.9 | CCDC77 | rs7307606 | 2.60E-07 | ##### | Brain - Substantia nigra | 440788 | -59212 | 940788 | Yes |
| ENSG00000120647.9 | CCDC77 | rs10661251 | 2.60E-07 | ##### | Brain - Substantia nigra | 440788 | -59212 | 940788 | Yes |
| ENSG00000120647.9 | CCDC77 | rs1011699 | 2.60E-07 | ##### | Brain - Substantia nigra | 440788 | -59212 | 940788 | Yes |
| ENSG00000120647.9 | CCDC77 | rs6489475 | 2.60E-07 | ##### | Brain - Substantia nigra | 440788 | -59212 | 940788 | Yes |
| ENSG00000120647.9 | CCDC77 | rs10774154 | 2.60E-07 | ##### | Brain - Substantia nigra | 440788 | -59212 | 940788 | Yes |
| ENSG00000120647.9 | CCDC77 | rs10848987 | 2.60E-07 | ##### | Brain - Cerebellar Hemisphere | 440788 | -59212 | 940788 | Yes |
| ENSG00000120647.9 | CCDC77 | rs11062945 | 2.60E-07 | ##### | Brain - Caudate (basal ganglia | 440788 | -59212 | 940788 | Yes |
| ENSG00000120647.9 | CCDC77 | rs7311740 | 2.70E-07 | ##### | Brain - Caudate (basal ganglia | 440788 | -59212 | 940788 | Yes |
| ENSG00000120647.9 | CCDC77 | rs1548904 | 2.70E-07 | ##### | Brain - Hypothalamus | 440788 | -59212 | 940788 | Yes |

| | | | | | | | | | |
|---|---|---|---|---|---|---|---|---|---|
| ENSG00000120647.9 | CCDC77 | rs1972582 | 2.70E-07 | ##### | Brain - Hypothalamus | 440788 | -59212 | 940788 | Yes |
| ENSG00000120647.9 | CCDC77 | rs10774038 | 2.70E-07 | ##### | Brain - Hypothalamus | 440788 | -59212 | 940788 | Yes |
| ENSG00000120647.9 | CCDC77 | rs576159 | 2.80E-07 | ##### | Brain - Cortex | 440788 | -59212 | 940788 | Yes |
| ENSG00000120647.9 | CCDC77 | rs507597 | 2.90E-07 | ##### | Brain - Nucleus accumbens (b | 440788 | -59212 | 940788 | Yes |
| ENSG00000120647.9 | CCDC77 | rs17800744 | 2.90E-07 | ##### | Brain - Nucleus accumbens (b | 440788 | -59212 | 940788 | Yes |
| ENSG00000120647.9 | CCDC77 | rs77854644 | 2.90E-07 | ##### | Brain - Nucleus accumbens (b | 440788 | -59212 | 940788 | Yes |
| ENSG00000120647.9 | CCDC77 | rs4980896 | 3.00E-07 | ##### | Brain - Putamen (basal gangli | 440788 | -59212 | 940788 | Yes |
| ENSG00000120647.9 | CCDC77 | rs520932 | 3.00E-07 | ##### | Brain - Anterior cingulate cort | 440788 | -59212 | 940788 | Yes |
| ENSG00000120647.9 | CCDC77 | rs56042953 | 3.00E-07 | ##### | Brain - Substantia nigra | 440788 | -59212 | 940788 | Yes |
| ENSG00000120647.9 | CCDC77 | rs10774169 | 3.10E-07 | ##### | Brain - Anterior cingulate cort | 440788 | -59212 | 940788 | Yes |
| ENSG00000120647.9 | CCDC77 | rs579526 | 3.10E-07 | ##### | Brain - Cerebellum | 440788 | -59212 | 940788 | Yes |
| ENSG00000120647.9 | CCDC77 | rs10848647 | 3.10E-07 | ##### | Brain - Substantia nigra | 440788 | -59212 | 940788 | Yes |
| ENSG00000120647.9 | CCDC77 | rs10848650 | 3.10E-07 | ##### | Brain - Substantia nigra | 440788 | -59212 | 940788 | Yes |
| ENSG00000120647.9 | CCDC77 | rs11613331 | 3.20E-07 | ##### | Brain - Caudate (basal ganglia | 440788 | -59212 | 940788 | Yes |
| ENSG00000120647.9 | CCDC77 | rs2041299 | 3.30E-07 | ##### | Brain - Anterior cingulate cort | 440788 | -59212 | 940788 | Yes |
| ENSG00000120647.9 | CCDC77 | rs75894333 | 3.30E-07 | ##### | Brain - Hippocampus | 440788 | -59212 | 940788 | Yes |
| ENSG00000120647.9 | CCDC77 | rs11608938 | 3.30E-07 | ##### | Brain - Hippocampus | 440788 | -59212 | 940788 | Yes |
| ENSG00000120647.9 | CCDC77 | rs7965038 | 3.30E-07 | ##### | Brain - Hippocampus | 440788 | -59212 | 940788 | Yes |
| ENSG00000120647.9 | CCDC77 | rs7307606 | 3.30E-07 | ##### | Brain - Hippocampus | 440788 | -59212 | 940788 | Yes |
| ENSG00000120647.9 | CCDC77 | rs10661251 | 3.30E-07 | ##### | Brain - Hippocampus | 440788 | -59212 | 940788 | Yes |
| ENSG00000120647.9 | CCDC77 | rs1011699 | 3.30E-07 | ##### | Brain - Hippocampus | 440788 | -59212 | 940788 | Yes |
| ENSG00000120647.9 | CCDC77 | rs6489475 | 3.30E-07 | ##### | Brain - Hippocampus | 440788 | -59212 | 940788 | Yes |
| ENSG00000120647.9 | CCDC77 | rs10774154 | 3.30E-07 | ##### | Brain - Hippocampus | 440788 | -59212 | 940788 | Yes |
| ENSG00000120647.9 | CCDC77 | rs2284333 | 3.30E-07 | ##### | Brain - Nucleus accumbens (b | 440788 | -59212 | 940788 | Yes |
| ENSG00000120647.9 | CCDC77 | rs10848766 | 3.30E-07 | ##### | Brain - Frontal Cortex (BA9) | 440788 | -59212 | 940788 | Yes |
| ENSG00000120647.9 | CCDC77 | rs797765 | 3.40E-07 | ##### | Brain - Cerebellum | 440788 | -59212 | 940788 | Yes |
| ENSG00000120647.9 | CCDC77 | rs493624 | 3.40E-07 | ##### | Brain - Hippocampus | 440788 | -59212 | 940788 | Yes |
| ENSG00000120647.9 | CCDC77 | rs4980822 | 3.50E-07 | ##### | Brain - Spinal cord (cervical c- | 440788 | -59212 | 940788 | Yes |
| ENSG00000120647.9 | CCDC77 | rs11062945 | 3.60E-07 | ##### | Brain - Hypothalamus | 440788 | -59212 | 940788 | Yes |
| ENSG00000120647.9 | CCDC77 | rs7132845 | 3.60E-07 | ##### | Brain - Cortex | 440788 | -59212 | 940788 | Yes |
| ENSG00000120647.9 | CCDC77 | rs764614 | 3.70E-07 | ##### | Brain - Anterior cingulate cort | 440788 | -59212 | 940788 | Yes |
| ENSG00000120647.9 | CCDC77 | rs11062813 | 3.70E-07 | ##### | Brain - Nucleus accumbens (b | 440788 | -59212 | 940788 | Yes |

| | | | | | | | | | |
|---|---|---|---|---|---|---|---|---|---|
| ENSG00000120647.9 | CCDC77 | rs7310504 | 3.80E-07 | ##### | Brain - Hypothalamus | 440788 | -59212 | 940788 | Yes |
| ENSG00000120647.9 | CCDC77 | rs11062928 | 3.80E-07 | ##### | Brain - Caudate (basal ganglia | 440788 | -59212 | 940788 | Yes |
| ENSG00000120647.9 | CCDC77 | rs4980886 | 3.80E-07 | ##### | Brain - Cortex | 440788 | -59212 | 940788 | Yes |
| ENSG00000120647.9 | CCDC77 | rs16930719 | 3.90E-07 | ##### | Brain - Caudate (basal ganglia | 440788 | -59212 | 940788 | Yes |
| ENSG00000120647.9 | CCDC77 | rs16930719 | 3.90E-07 | ##### | Brain - Hypothalamus | 440788 | -59212 | 940788 | Yes |
| ENSG00000120647.9 | CCDC77 | rs520932 | 3.90E-07 | ##### | Brain - Substantia nigra | 440788 | -59212 | 940788 | Yes |
| ENSG00000120647.9 | CCDC77 | rs11062167 | 3.90E-07 | ##### | Brain - Putamen (basal gangli | 440788 | -59212 | 940788 | Yes |
| ENSG00000120647.9 | CCDC77 | rs55990424 | 3.90E-07 | ##### | Brain - Putamen (basal gangli | 440788 | -59212 | 940788 | Yes |
| ENSG00000120647.9 | CCDC77 | rs2300133 | 3.90E-07 | ##### | Brain - Cerebellum | 440788 | -59212 | 940788 | Yes |
| ENSG00000120647.9 | CCDC77 | rs2052279 | 4.10E-07 | ##### | Brain - Nucleus accumbens (b | 440788 | -59212 | 940788 | Yes |
| ENSG00000120647.9 | CCDC77 | rs78813329 | 4.10E-07 | ##### | Brain - Nucleus accumbens (b | 440788 | -59212 | 940788 | Yes |
| ENSG00000120647.9 | CCDC77 | rs3753157 | 4.10E-07 | ##### | Brain - Nucleus accumbens (b | 440788 | -59212 | 940788 | Yes |
| ENSG00000120647.9 | CCDC77 | rs4980883 | 4.10E-07 | ##### | Brain - Nucleus accumbens (b | 440788 | -59212 | 940788 | Yes |
| ENSG00000120647.9 | CCDC77 | rs137910445 | 4.10E-07 | ##### | Brain - Nucleus accumbens (b | 440788 | -59212 | 940788 | Yes |
| ENSG00000120647.9 | CCDC77 | rs11062537 | 4.10E-07 | ##### | Brain - Nucleus accumbens (b | 440788 | -59212 | 940788 | Yes |
| ENSG00000120647.9 | CCDC77 | rs715230 | 4.10E-07 | ##### | Brain - Nucleus accumbens (b | 440788 | -59212 | 940788 | Yes |
| ENSG00000120647.9 | CCDC77 | rs715229 | 4.10E-07 | ##### | Brain - Nucleus accumbens (b | 440788 | -59212 | 940788 | Yes |
| ENSG00000120647.9 | CCDC77 | rs715228 | 4.10E-07 | ##### | Brain - Nucleus accumbens (b | 440788 | -59212 | 940788 | Yes |
| ENSG00000120647.9 | CCDC77 | rs10848838 | 4.10E-07 | ##### | Brain - Nucleus accumbens (b | 440788 | -59212 | 940788 | Yes |
| ENSG00000120647.9 | CCDC77 | rs11062945 | 4.40E-07 | ##### | Brain - Cerebellar Hemisphere | 440788 | -59212 | 940788 | Yes |
| ENSG00000120647.9 | CCDC77 | rs487252 | 4.50E-07 | ##### | Brain - Hypothalamus | 440788 | -59212 | 940788 | Yes |
| ENSG00000120647.9 | CCDC77 | rs6489523 | 4.50E-07 | ##### | Brain - Cerebellar Hemisphere | 440788 | -59212 | 940788 | Yes |
| ENSG00000120647.9 | CCDC77 | rs4980884 | 4.50E-07 | ##### | Brain - Nucleus accumbens (b | 440788 | -59212 | 940788 | Yes |
| ENSG00000120647.9 | CCDC77 | rs797765 | 4.60E-07 | ##### | Brain - Hippocampus | 440788 | -59212 | 940788 | Yes |
| ENSG00000120647.9 | CCDC77 | rs4980874 | 4.60E-07 | ##### | Brain - Nucleus accumbens (b | 440788 | -59212 | 940788 | Yes |
| ENSG00000120647.9 | CCDC77 | rs3213751 | 4.70E-07 | ##### | Brain - Nucleus accumbens (b | 440788 | -59212 | 940788 | Yes |
| ENSG00000120647.9 | CCDC77 | rs7311740 | 4.70E-07 | ##### | Brain - Spinal cord (cervical c- | 440788 | -59212 | 940788 | Yes |
| ENSG00000120647.9 | CCDC77 | rs10160995 | 4.70E-07 | ##### | Brain - Spinal cord (cervical c- | 440788 | -59212 | 940788 | Yes |
| ENSG00000120647.9 | CCDC77 | rs4980898 | 4.70E-07 | ##### | Brain - Spinal cord (cervical c- | 440788 | -59212 | 940788 | Yes |
| ENSG00000120647.9 | CCDC77 | rs16930719 | 4.80E-07 | ##### | Brain - Cerebellar Hemisphere | 440788 | -59212 | 940788 | Yes |
| ENSG00000120647.9 | CCDC77 | rs4980896 | 4.90E-07 | ##### | Brain - Cortex | 440788 | -59212 | 940788 | Yes |
| ENSG00000120647.9 | CCDC77 | rs3764876 | 4.90E-07 | ##### | Brain - Cerebellar Hemisphere | 440788 | -59212 | 940788 | Yes |

| | | | | | | | | | |
|---|---|---|---|---|---|---|---|---|---|
| ENSG00000120647.9 | CCDC77 | rs10774191 | 4.90E-07 | ##### | Brain - Cerebellar Hemisphere | 440788 | -59212 | 940788 | Yes |
| ENSG00000120647.9 | CCDC77 | rs61916593 | 4.90E-07 | ##### | Brain - Cerebellar Hemisphere | 440788 | -59212 | 940788 | Yes |
| ENSG00000120647.9 | CCDC77 | rs2041298 | 5.00E-07 | ##### | Brain - Cerebellar Hemisphere | 440788 | -59212 | 940788 | Yes |
| ENSG00000120647.9 | CCDC77 | rs2080402 | 5.00E-07 | ##### | Brain - Hypothalamus | 440788 | -59212 | 940788 | Yes |
| ENSG00000120647.9 | CCDC77 | rs55657100 | 5.00E-07 | ##### | Brain - Putamen (basal gangli | 440788 | -59212 | 940788 | Yes |
| ENSG00000120647.9 | CCDC77 | rs7138981 | 5.20E-07 | ##### | Brain - Anterior cingulate cort | 440788 | -59212 | 940788 | Yes |
| ENSG00000120647.9 | CCDC77 | rs9669094 | 5.20E-07 | ##### | Brain - Anterior cingulate cort | 440788 | -59212 | 940788 | Yes |
| ENSG00000120647.9 | CCDC77 | rs7132763 | 5.20E-07 | ##### | Brain - Anterior cingulate cort | 440788 | -59212 | 940788 | Yes |
| ENSG00000120647.9 | CCDC77 | rs2015554 | 5.20E-07 | ##### | Brain - Anterior cingulate cort | 440788 | -59212 | 940788 | Yes |
| ENSG00000120647.9 | CCDC77 | rs28850111 | 5.30E-07 | ##### | Brain - Spinal cord (cervical c- | 440788 | -59212 | 940788 | Yes |
| ENSG00000120647.9 | CCDC77 | rs55657100 | 5.30E-07 | ##### | Brain - Spinal cord (cervical c- | 440788 | -59212 | 940788 | Yes |
| ENSG00000120647.9 | CCDC77 | rs12301816 | 5.70E-07 | ##### | Brain - Spinal cord (cervical c- | 440788 | -59212 | 940788 | Yes |
| ENSG00000120647.9 | CCDC77 | rs576159 | 5.80E-07 | ##### | Brain - Cerebellum | 440788 | -59212 | 940788 | Yes |
| ENSG00000120647.9 | CCDC77 | rs486098 | 5.90E-07 | ##### | Brain - Nucleus accumbens (b | 440788 | -59212 | 940788 | Yes |
| ENSG00000120647.9 | CCDC77 | rs10774192 | 6.00E-07 | ##### | Brain - Spinal cord (cervical c- | 440788 | -59212 | 940788 | Yes |
| ENSG00000120647.9 | CCDC77 | rs10774152 | 6.00E-07 | ##### | Brain - Anterior cingulate cort | 440788 | -59212 | 940788 | Yes |
| ENSG00000120647.9 | CCDC77 | rs10774153 | 6.00E-07 | ##### | Brain - Anterior cingulate cort | 440788 | -59212 | 940788 | Yes |
| ENSG00000120647.9 | CCDC77 | rs2080402 | 6.20E-07 | ##### | Brain - Caudate (basal ganglia | 440788 | -59212 | 940788 | Yes |
| ENSG00000120647.9 | CCDC77 | rs11062813 | 6.20E-07 | ##### | Brain - Cerebellum | 440788 | -59212 | 940788 | Yes |
| ENSG00000120647.9 | CCDC77 | rs7136116 | 6.30E-07 | ##### | Brain - Putamen (basal gangli | 440788 | -59212 | 940788 | Yes |
| ENSG00000120647.9 | CCDC77 | rs11610457 | 6.30E-07 | ##### | Brain - Putamen (basal gangli | 440788 | -59212 | 940788 | Yes |
| ENSG00000120647.9 | CCDC77 | rs7138981 | 6.30E-07 | ##### | Brain - Cortex | 440788 | -59212 | 940788 | Yes |
| ENSG00000120647.9 | CCDC77 | rs9669094 | 6.30E-07 | ##### | Brain - Cortex | 440788 | -59212 | 940788 | Yes |
| ENSG00000120647.9 | CCDC77 | rs2015554 | 6.30E-07 | ##### | Brain - Cortex | 440788 | -59212 | 940788 | Yes |
| ENSG00000120647.9 | CCDC77 | rs10774152 | 6.30E-07 | ##### | Brain - Cortex | 440788 | -59212 | 940788 | Yes |
| ENSG00000120647.9 | CCDC77 | rs10774153 | 6.30E-07 | ##### | Brain - Cortex | 440788 | -59212 | 940788 | Yes |
| ENSG00000120647.9 | CCDC77 | rs495316 | 6.40E-07 | ##### | Brain - Anterior cingulate cort | 440788 | -59212 | 940788 | Yes |
| ENSG00000120647.9 | CCDC77 | rs11613331 | 6.50E-07 | ##### | Brain - Cerebellar Hemisphere | 440788 | -59212 | 940788 | Yes |
| ENSG00000120647.9 | CCDC77 | rs520932 | 6.50E-07 | ##### | Brain - Cerebellum | 440788 | -59212 | 940788 | Yes |
| ENSG00000120647.9 | CCDC77 | rs61907026 | 6.50E-07 | ##### | Brain - Substantia nigra | 440788 | -59212 | 940788 | Yes |
| ENSG00000120647.9 | CCDC77 | rs10082788 | 6.60E-07 | ##### | Brain - Putamen (basal gangli | 440788 | -59212 | 940788 | Yes |
| ENSG00000120647.9 | CCDC77 | rs10848644 | 6.60E-07 | ##### | Brain - Putamen (basal gangli | 440788 | -59212 | 940788 | Yes |

| | | | | | | | | | |
|---|---|---|---|---|---|---|---|---|---|
| ENSG00000120647.9 | CCDC77 | rs495973 | 6.60E-07 | ##### | Brain - Anterior cingulate cort | 440788 | -59212 | 940788 | Yes |
| ENSG00000120647.9 | CCDC77 | rs487252 | 6.70E-07 | ##### | Brain - Hippocampus | 440788 | -59212 | 940788 | Yes |
| ENSG00000120647.9 | CCDC77 | rs10774169 | 6.90E-07 | ##### | Brain - Cerebellum | 440788 | -59212 | 940788 | Yes |
| ENSG00000120647.9 | CCDC77 | rs2300133 | 6.90E-07 | ##### | Brain - Nucleus accumbens (b | 440788 | -59212 | 940788 | Yes |
| ENSG00000120647.9 | CCDC77 | rs3753156 | 6.90E-07 | ##### | Brain - Nucleus accumbens (b | 440788 | -59212 | 940788 | Yes |
| ENSG00000120647.9 | CCDC77 | rs12581260 | 6.90E-07 | ##### | Brain - Nucleus accumbens (b | 440788 | -59212 | 940788 | Yes |
| ENSG00000120647.9 | CCDC77 | rs16929783 | 6.90E-07 | ##### | Brain - Nucleus accumbens (b | 440788 | -59212 | 940788 | Yes |
| ENSG00000120647.9 | CCDC77 | rs10161265 | 6.90E-07 | ##### | Brain - Frontal Cortex (BA9) | 440788 | -59212 | 940788 | Yes |
| ENSG00000120647.9 | CCDC77 | rs4980897 | 6.90E-07 | ##### | Brain - Frontal Cortex (BA9) | 440788 | -59212 | 940788 | Yes |
| ENSG00000120647.9 | CCDC77 | rs113574411 | 6.90E-07 | ##### | Brain - Cortex | 440788 | -59212 | 940788 | Yes |
| ENSG00000120647.9 | CCDC77 | rs479840 | 6.90E-07 | ##### | Brain - Cortex | 440788 | -59212 | 940788 | Yes |
| ENSG00000120647.9 | CCDC77 | rs479803 | 6.90E-07 | ##### | Brain - Cortex | 440788 | -59212 | 940788 | Yes |
| ENSG00000120647.9 | CCDC77 | rs72472276 | 6.90E-07 | ##### | Brain - Cortex | 440788 | -59212 | 940788 | Yes |
| ENSG00000120647.9 | CCDC77 | rs4980877 | 6.90E-07 | ##### | Brain - Cortex | 440788 | -59212 | 940788 | Yes |
| ENSG00000120647.9 | CCDC77 | rs7963309 | 6.90E-07 | ##### | Brain - Cortex | 440788 | -59212 | 940788 | Yes |
| ENSG00000120647.9 | CCDC77 | rs7134353 | 6.90E-07 | ##### | Brain - Cortex | 440788 | -59212 | 940788 | Yes |
| ENSG00000120647.9 | CCDC77 | rs11374245 | 6.90E-07 | ##### | Brain - Cortex | 440788 | -59212 | 940788 | Yes |
| ENSG00000120647.9 | CCDC77 | rs16929744 | 6.90E-07 | ##### | Brain - Cortex | 440788 | -59212 | 940788 | Yes |
| ENSG00000120647.9 | CCDC77 | rs3753158 | 6.90E-07 | ##### | Brain - Cortex | 440788 | -59212 | 940788 | Yes |
| ENSG00000120647.9 | CCDC77 | rs7310504 | 7.10E-07 | ##### | Brain - Frontal Cortex (BA9) | 440788 | -59212 | 940788 | Yes |
| ENSG00000120647.9 | CCDC77 | rs10082788 | 7.10E-07 | ##### | Brain - Cortex | 440788 | -59212 | 940788 | Yes |
| ENSG00000120647.9 | CCDC77 | rs113574411 | 7.10E-07 | ##### | Brain - Anterior cingulate cort | 440788 | -59212 | 940788 | Yes |
| ENSG00000120647.9 | CCDC77 | rs479840 | 7.10E-07 | ##### | Brain - Anterior cingulate cort | 440788 | -59212 | 940788 | Yes |
| ENSG00000120647.9 | CCDC77 | rs479803 | 7.10E-07 | ##### | Brain - Anterior cingulate cort | 440788 | -59212 | 940788 | Yes |
| ENSG00000120647.9 | CCDC77 | rs77074951 | 7.10E-07 | ##### | Brain - Anterior cingulate cort | 440788 | -59212 | 940788 | Yes |
| ENSG00000120647.9 | CCDC77 | rs72472276 | 7.10E-07 | ##### | Brain - Anterior cingulate cort | 440788 | -59212 | 940788 | Yes |
| ENSG00000120647.9 | CCDC77 | rs4980877 | 7.10E-07 | ##### | Brain - Anterior cingulate cort | 440788 | -59212 | 940788 | Yes |
| ENSG00000120647.9 | CCDC77 | rs7963309 | 7.10E-07 | ##### | Brain - Anterior cingulate cort | 440788 | -59212 | 940788 | Yes |
| ENSG00000120647.9 | CCDC77 | rs7134353 | 7.10E-07 | ##### | Brain - Anterior cingulate cort | 440788 | -59212 | 940788 | Yes |
| ENSG00000120647.9 | CCDC77 | rs11374245 | 7.10E-07 | ##### | Brain - Anterior cingulate cort | 440788 | -59212 | 940788 | Yes |
| ENSG00000120647.9 | CCDC77 | rs7303180 | 7.10E-07 | ##### | Brain - Anterior cingulate cort | 440788 | -59212 | 940788 | Yes |
| ENSG00000120647.9 | CCDC77 | rs16929744 | 7.10E-07 | ##### | Brain - Anterior cingulate cort | 440788 | -59212 | 940788 | Yes |

| | | | | | | | | | |
|---|---|---|---|---|---|---|---|---|---|
| ENSG00000120647.9 | CCDC77 | rs7969761 | 7.50E-07 | ##### | Brain - Cerebellar Hemisphere | 440788 | -59212 | 940788 | Yes |
| ENSG00000120647.9 | CCDC77 | rs492540 | 7.50E-07 | ##### | Brain - Hippocampus | 440788 | -59212 | 940788 | Yes |
| ENSG00000120647.9 | CCDC77 | rs486098 | 7.60E-07 | ##### | Brain - Cortex | 440788 | -59212 | 940788 | Yes |
| ENSG00000120647.9 | CCDC77 | rs7136116 | 7.70E-07 | ##### | Brain - Substantia nigra | 440788 | -59212 | 940788 | Yes |
| ENSG00000120647.9 | CCDC77 | rs11610457 | 7.70E-07 | ##### | Brain - Substantia nigra | 440788 | -59212 | 940788 | Yes |
| ENSG00000120647.9 | CCDC77 | rs372437080 | 7.80E-07 | ##### | Brain - Hippocampus | 440788 | -59212 | 940788 | Yes |
| ENSG00000120647.9 | CCDC77 | rs55657100 | 7.80E-07 | ##### | Brain - Substantia nigra | 440788 | -59212 | 940788 | Yes |
| ENSG00000120647.9 | CCDC77 | rs12301816 | 7.90E-07 | ##### | Brain - Substantia nigra | 440788 | -59212 | 940788 | Yes |
| ENSG00000120647.9 | CCDC77 | rs2369281 | 8.00E-07 | ##### | Brain - Caudate (basal ganglia | 440788 | -59212 | 940788 | Yes |
| ENSG00000120647.9 | CCDC77 | rs10848987 | 8.00E-07 | ##### | Brain - Spinal cord (cervical c- | 440788 | -59212 | 940788 | Yes |
| ENSG00000120647.9 | CCDC77 | rs505211 | 8.00E-07 | ##### | Brain - Hippocampus | 440788 | -59212 | 940788 | Yes |
| ENSG00000120647.9 | CCDC77 | rs532463 | 8.00E-07 | ##### | Brain - Hippocampus | 440788 | -59212 | 940788 | Yes |
| ENSG00000120647.9 | CCDC77 | rs505574 | 8.00E-07 | ##### | Brain - Hippocampus | 440788 | -59212 | 940788 | Yes |
| ENSG00000120647.9 | CCDC77 | rs1088324 | 8.00E-07 | ##### | Brain - Hippocampus | 440788 | -59212 | 940788 | Yes |
| ENSG00000120647.9 | CCDC77 | rs2041299 | 8.00E-07 | ##### | Brain - Hippocampus | 440788 | -59212 | 940788 | Yes |
| ENSG00000120647.9 | CCDC77 | rs6489477 | 8.10E-07 | ##### | Brain - Cortex | 440788 | -59212 | 940788 | Yes |
| ENSG00000120647.9 | CCDC77 | rs6489478 | 8.10E-07 | ##### | Brain - Cortex | 440788 | -59212 | 940788 | Yes |
| ENSG00000120647.9 | CCDC77 | rs11063005 | 8.10E-07 | ##### | Brain - Spinal cord (cervical c- | 440788 | -59212 | 940788 | Yes |
| ENSG00000120647.9 | CCDC77 | rs11063010 | 8.10E-07 | ##### | Brain - Spinal cord (cervical c- | 440788 | -59212 | 940788 | Yes |
| ENSG00000120647.9 | CCDC77 | rs10431354 | 8.20E-07 | ##### | Brain - Cortex | 440788 | -59212 | 940788 | Yes |
| ENSG00000120647.9 | CCDC77 | rs12298843 | 8.20E-07 | ##### | Brain - Nucleus accumbens (b | 440788 | -59212 | 940788 | Yes |
| ENSG00000120647.9 | CCDC77 | rs4980892 | 8.30E-07 | ##### | Brain - Anterior cingulate cort | 440788 | -59212 | 940788 | Yes |
| ENSG00000120647.9 | CCDC77 | rs4980874 | 8.30E-07 | ##### | Brain - Anterior cingulate cort | 440788 | -59212 | 940788 | Yes |
| ENSG00000120647.9 | CCDC77 | rs12370433 | 8.40E-07 | ##### | Brain - Caudate (basal ganglia | 440788 | -59212 | 940788 | Yes |
| ENSG00000120647.9 | CCDC77 | rs2284333 | 8.50E-07 | ##### | Brain - Anterior cingulate cort | 440788 | -59212 | 940788 | Yes |
| ENSG00000120647.9 | CCDC77 | rs4980811 | 8.50E-07 | ##### | Brain - Anterior cingulate cort | 440788 | -59212 | 940788 | Yes |
| ENSG00000120647.9 | CCDC77 | rs4980892 | 8.50E-07 | ##### | Brain - Cerebellum | 440788 | -59212 | 940788 | Yes |
| ENSG00000120647.9 | CCDC77 | rs524468 | 8.70E-07 | ##### | Brain - Substantia nigra | 440788 | -59212 | 940788 | Yes |
| ENSG00000120647.9 | CCDC77 | rs4980908 | 8.90E-07 | ##### | Brain - Hypothalamus | 440788 | -59212 | 940788 | Yes |
| ENSG00000120647.9 | CCDC77 | rs61916594 | 9.30E-07 | ##### | Brain - Caudate (basal ganglia | 440788 | -59212 | 940788 | Yes |
| ENSG00000120647.9 | CCDC77 | rs10774190 | 9.30E-07 | ##### | Brain - Caudate (basal ganglia | 440788 | -59212 | 940788 | Yes |
| ENSG00000120647.9 | CCDC77 | rs79839498 | 9.40E-07 | ##### | Brain - Nucleus accumbens (b | 440788 | -59212 | 940788 | Yes |

| | | | | | | | | | |
|---|---|---|---|---|---|---|---|---|---|
| ENSG00000120647.9 | CCDC77 | rs61906962 | 9.50E-07 | ##### | Brain - Hippocampus | 440788 | -59212 | 940788 | Yes |
| ENSG00000120647.9 | CCDC77 | rs55876072 | 9.50E-07 | ##### | Brain - Hippocampus | 440788 | -59212 | 940788 | Yes |
| ENSG00000120647.9 | CCDC77 | rs3815558 | 9.50E-07 | ##### | Brain - Nucleus accumbens (b | 440788 | -59212 | 940788 | Yes |
| ENSG00000120647.9 | CCDC77 | rs78998485 | 9.50E-07 | ##### | Brain - Nucleus accumbens (b | 440788 | -59212 | 940788 | Yes |
| ENSG00000120647.9 | CCDC77 | rs7135768 | 9.50E-07 | ##### | Brain - Nucleus accumbens (b | 440788 | -59212 | 940788 | Yes |
| ENSG00000120647.9 | CCDC77 | rs6489450 | 9.50E-07 | ##### | Brain - Nucleus accumbens (b | 440788 | -59212 | 940788 | Yes |
| ENSG00000120647.9 | CCDC77 | rs2284333 | 9.60E-07 | ##### | Brain - Cortex | 440788 | -59212 | 940788 | Yes |
| ENSG00000120647.9 | CCDC77 | rs4980811 | 9.60E-07 | ##### | Brain - Cortex | 440788 | -59212 | 940788 | Yes |
| ENSG00000120647.9 | CCDC77 | rs4980908 | 9.60E-07 | ##### | Brain - Spinal cord (cervical c- | 440788 | -59212 | 940788 | Yes |
| ENSG00000120647.9 | CCDC77 | rs10431354 | 9.60E-07 | ##### | Brain - Cerebellum | 440788 | -59212 | 940788 | Yes |
| ENSG00000120647.9 | CCDC77 | rs7310504 | 9.70E-07 | ##### | Brain - Caudate (basal ganglia | 440788 | -59212 | 940788 | Yes |
| ENSG00000120647.9 | CCDC77 | rs10848766 | 9.90E-07 | ##### | Brain - Putamen (basal gangli | 440788 | -59212 | 940788 | Yes |
| ENSG00000120647.9 | CCDC77 | rs4980886 | 0.000001 | ##### | Brain - Anterior cingulate cort | 440788 | -59212 | 940788 | Yes |
| ENSG00000120647.9 | CCDC77 | rs11062357 | 0.000001 | ##### | Brain - Frontal Cortex (BA9) | 440788 | -59212 | 940788 | Yes |
| ENSG00000120647.9 | CCDC77 | rs10744627 | 0.000001 | ##### | Brain - Anterior cingulate cort | 440788 | -59212 | 940788 | Yes |
| ENSG00000120647.9 | CCDC77 | rs6489477 | 0.000001 | ##### | Brain - Cerebellum | 440788 | -59212 | 940788 | Yes |
| ENSG00000120647.9 | CCDC77 | rs6489478 | 0.000001 | ##### | Brain - Cerebellum | 440788 | -59212 | 940788 | Yes |
| ENSG00000120647.9 | CCDC77 | rs520932 | 0.0000011 | ##### | Brain - Cortex | 440788 | -59212 | 940788 | Yes |
| ENSG00000120647.9 | CCDC77 | rs4980908 | 0.0000011 | ##### | Brain - Caudate (basal ganglia | 440788 | -59212 | 940788 | Yes |
| ENSG00000120647.9 | CCDC77 | rs3782860 | 0.0000011 | ##### | Brain - Putamen (basal gangli | 440788 | -59212 | 940788 | Yes |
| ENSG00000120647.9 | CCDC77 | rs7303180 | 0.0000011 | ##### | Brain - Cortex | 440788 | -59212 | 940788 | Yes |
| ENSG00000120647.9 | CCDC77 | rs1548904 | 0.0000011 | ##### | Brain - Substantia nigra | 440788 | -59212 | 940788 | Yes |
| ENSG00000120647.9 | CCDC77 | rs10744561 | 0.0000011 | ##### | Brain - Substantia nigra | 440788 | -59212 | 940788 | Yes |
| ENSG00000120647.9 | CCDC77 | rs1972582 | 0.0000011 | ##### | Brain - Substantia nigra | 440788 | -59212 | 940788 | Yes |
| ENSG00000120647.9 | CCDC77 | rs10774038 | 0.0000011 | ##### | Brain - Substantia nigra | 440788 | -59212 | 940788 | Yes |
| ENSG00000120647.9 | CCDC77 | rs11613357 | 0.0000011 | ##### | Brain - Frontal Cortex (BA9) | 440788 | -59212 | 940788 | Yes |
| ENSG00000120647.9 | CCDC77 | rs3213750 | 0.0000011 | ##### | Brain - Nucleus accumbens (b | 440788 | -59212 | 940788 | Yes |
| ENSG00000120647.9 | CCDC77 | rs4980881 | 0.0000011 | ##### | Brain - Nucleus accumbens (b | 440788 | -59212 | 940788 | Yes |
| ENSG00000120647.9 | CCDC77 | rs79839498 | 0.0000012 | ##### | Brain - Substantia nigra | 440788 | -59212 | 940788 | Yes |
| ENSG00000120647.9 | CCDC77 | rs3753158 | 0.0000012 | ##### | Brain - Spinal cord (cervical c- | 440788 | -59212 | 940788 | Yes |
| ENSG00000120647.9 | CCDC77 | rs4980906 | 0.0000012 | ##### | Brain - Caudate (basal ganglia | 440788 | -59212 | 940788 | Yes |
| ENSG00000120647.9 | CCDC77 | rs2286606 | 0.0000012 | ##### | Brain - Caudate (basal ganglia | 440788 | -59212 | 940788 | Yes |

| | | | | | | | | | |
|---|---|---|---|---|---|---|---|---|---|
| ENSG00000120647.9 | CCDC77 | rs77074951 | 0.0000012 | ##### | Brain - Cortex | 440788 | -59212 | 940788 | Yes |
| ENSG00000120647.9 | CCDC77 | rs10848974 | 0.0000012 | ##### | Brain - Caudate (basal ganglia | 440788 | -59212 | 940788 | Yes |
| ENSG00000120647.9 | CCDC77 | rs61907026 | 0.0000012 | ##### | Brain - Nucleus accumbens (b | 440788 | -59212 | 940788 | Yes |
| ENSG00000120647.9 | CCDC77 | rs2041299 | 0.0000012 | ##### | Brain - Nucleus accumbens (b | 440788 | -59212 | 940788 | Yes |
| ENSG00000120647.9 | CCDC77 | rs4980874 | 0.0000012 | ##### | Brain - Cortex | 440788 | -59212 | 940788 | Yes |
| ENSG00000120647.9 | CCDC77 | rs61907026 | 0.0000013 | ##### | Brain - Hippocampus | 440788 | -59212 | 940788 | Yes |
| ENSG00000120647.9 | CCDC77 | rs79839498 | 0.0000013 | ##### | Brain - Hippocampus | 440788 | -59212 | 940788 | Yes |
| ENSG00000120647.9 | CCDC77 | rs61907030 | 0.0000013 | ##### | Brain - Hippocampus | 440788 | -59212 | 940788 | Yes |
| ENSG00000120647.9 | CCDC77 | rs61907031 | 0.0000013 | ##### | Brain - Hippocampus | 440788 | -59212 | 940788 | Yes |
| ENSG00000120647.9 | CCDC77 | rs11615564 | 0.0000013 | ##### | Brain - Hippocampus | 440788 | -59212 | 940788 | Yes |
| ENSG00000120647.9 | CCDC77 | rs67523949 | 0.0000013 | ##### | Brain - Hypothalamus | 440788 | -59212 | 940788 | Yes |
| ENSG00000120647.9 | CCDC77 | rs17800720 | 0.0000013 | ##### | Brain - Putamen (basal gangli | 440788 | -59212 | 940788 | Yes |
| ENSG00000120647.9 | CCDC77 | rs3837515 | 0.0000013 | ##### | Brain - Putamen (basal gangli | 440788 | -59212 | 940788 | Yes |
| ENSG00000120647.9 | CCDC77 | rs495973 | 0.0000013 | ##### | Brain - Cerebellum | 440788 | -59212 | 940788 | Yes |
| ENSG00000120647.9 | CCDC77 | rs2080403 | 0.0000014 | ##### | Brain - Cortex | 440788 | -59212 | 940788 | Yes |
| ENSG00000120647.9 | CCDC77 | rs2289957 | 0.0000014 | ##### | Brain - Cortex | 440788 | -59212 | 940788 | Yes |
| ENSG00000120647.9 | CCDC77 | rs201124968 | 0.0000014 | ##### | Brain - Anterior cingulate cort | 440788 | -59212 | 940788 | Yes |
| ENSG00000120647.9 | CCDC77 | rs7311740 | 0.0000014 | ##### | Brain - Cerebellar Hemisphere | 440788 | -59212 | 940788 | Yes |
| ENSG00000120647.9 | CCDC77 | rs579366 | 0.0000014 | ##### | Brain - Hippocampus | 440788 | -59212 | 940788 | Yes |
| ENSG00000120647.9 | CCDC77 | rs492750 | 0.0000014 | ##### | Brain - Hippocampus | 440788 | -59212 | 940788 | Yes |
| ENSG00000120647.9 | CCDC77 | rs61907030 | 0.0000015 | ##### | Brain - Substantia nigra | 440788 | -59212 | 940788 | Yes |
| ENSG00000120647.9 | CCDC77 | rs61907031 | 0.0000015 | ##### | Brain - Substantia nigra | 440788 | -59212 | 940788 | Yes |
| ENSG00000120647.9 | CCDC77 | rs11615564 | 0.0000015 | ##### | Brain - Substantia nigra | 440788 | -59212 | 940788 | Yes |
| ENSG00000120647.9 | CCDC77 | rs111250603 | 0.0000016 | ##### | Brain - Anterior cingulate cort | 440788 | -59212 | 940788 | Yes |
| ENSG00000120647.9 | CCDC77 | rs2369277 | 0.0000016 | ##### | Brain - Anterior cingulate cort | 440788 | -59212 | 940788 | Yes |
| ENSG00000120647.9 | CCDC77 | rs9805067 | 0.0000016 | ##### | Brain - Anterior cingulate cort | 440788 | -59212 | 940788 | Yes |
| ENSG00000120647.9 | CCDC77 | rs1541464 | 0.0000016 | ##### | Brain - Anterior cingulate cort | 440788 | -59212 | 940788 | Yes |
| ENSG00000120647.9 | CCDC77 | rs1541465 | 0.0000016 | ##### | Brain - Anterior cingulate cort | 440788 | -59212 | 940788 | Yes |
| ENSG00000120647.9 | CCDC77 | rs4980817 | 0.0000016 | ##### | Brain - Anterior cingulate cort | 440788 | -59212 | 940788 | Yes |
| ENSG00000120647.9 | CCDC77 | rs4482086 | 0.0000016 | ##### | Brain - Anterior cingulate cort | 440788 | -59212 | 940788 | Yes |
| ENSG00000120647.9 | CCDC77 | rs2369279 | 0.0000016 | ##### | Brain - Anterior cingulate cort | 440788 | -59212 | 940788 | Yes |
| ENSG00000120647.9 | CCDC77 | rs11062803 | 0.0000016 | ##### | Brain - Anterior cingulate cort | 440788 | -59212 | 940788 | Yes |

| | | | | | | | | | |
|---|---|---|---|---|---|---|---|---|---|
| ENSG00000120647.9 | CCDC77 | rs7138981 | 0.0000016 | ##### | Brain - Cerebellum | 440788 | -59212 | 940788 | Yes |
| ENSG00000120647.9 | CCDC77 | rs9669094 | 0.0000016 | ##### | Brain - Cerebellum | 440788 | -59212 | 940788 | Yes |
| ENSG00000120647.9 | CCDC77 | rs2015554 | 0.0000016 | ##### | Brain - Cerebellum | 440788 | -59212 | 940788 | Yes |
| ENSG00000120647.9 | CCDC77 | rs10774152 | 0.0000016 | ##### | Brain - Cerebellum | 440788 | -59212 | 940788 | Yes |
| ENSG00000120647.9 | CCDC77 | rs10774153 | 0.0000016 | ##### | Brain - Cerebellum | 440788 | -59212 | 940788 | Yes |
| ENSG00000120647.9 | CCDC77 | rs2284333 | 0.0000016 | ##### | Brain - Cerebellum | 440788 | -59212 | 940788 | Yes |
| ENSG00000120647.9 | CCDC77 | rs11062986 | 0.0000016 | ##### | Brain - Caudate (basal ganglia | 440788 | -59212 | 940788 | Yes |
| ENSG00000120647.9 | CCDC77 | rs7138958 | 0.0000016 | ##### | Brain - Anterior cingulate cort | 440788 | -59212 | 940788 | Yes |
| ENSG00000120647.9 | CCDC77 | rs2007537 | 0.0000016 | ##### | Brain - Anterior cingulate cort | 440788 | -59212 | 940788 | Yes |
| ENSG00000120647.9 | CCDC77 | rs67523949 | 0.0000016 | ##### | Brain - Cortex | 440788 | -59212 | 940788 | Yes |
| ENSG00000120647.9 | CCDC77 | rs12833509 | 0.0000016 | ##### | Brain - Caudate (basal ganglia | 440788 | -59212 | 940788 | Yes |
| ENSG00000120647.9 | CCDC77 | rs10744561 | 0.0000017 | ##### | Brain - Putamen (basal gangli | 440788 | -59212 | 940788 | Yes |
| ENSG00000120647.9 | CCDC77 | rs10848650 | 0.0000017 | ##### | Brain - Putamen (basal gangli | 440788 | -59212 | 940788 | Yes |
| ENSG00000120647.9 | CCDC77 | rs530500 | 0.0000017 | ##### | Brain - Hippocampus | 440788 | -59212 | 940788 | Yes |
| ENSG00000120647.9 | CCDC77 | rs2041298 | 0.0000017 | ##### | Brain - Frontal Cortex (BA9) | 440788 | -59212 | 940788 | Yes |
| ENSG00000120647.9 | CCDC77 | rs2369280 | 0.0000017 | ##### | Brain - Anterior cingulate cort | 440788 | -59212 | 940788 | Yes |
| ENSG00000120647.9 | CCDC77 | rs579526 | 0.0000017 | ##### | Brain - Hippocampus | 440788 | -59212 | 940788 | Yes |
| ENSG00000120647.9 | CCDC77 | rs1541463 | 0.0000018 | ##### | Brain - Anterior cingulate cort | 440788 | -59212 | 940788 | Yes |
| ENSG00000120647.9 | CCDC77 | rs2369278 | 0.0000018 | ##### | Brain - Anterior cingulate cort | 440788 | -59212 | 940788 | Yes |
| ENSG00000120647.9 | CCDC77 | rs10774163 | 0.0000018 | ##### | Brain - Anterior cingulate cort | 440788 | -59212 | 940788 | Yes |
| ENSG00000120647.9 | CCDC77 | rs7136116 | 0.0000018 | ##### | Brain - Spinal cord (cervical c- | 440788 | -59212 | 940788 | Yes |
| ENSG00000120647.9 | CCDC77 | rs11610457 | 0.0000018 | ##### | Brain - Spinal cord (cervical c- | 440788 | -59212 | 940788 | Yes |
| ENSG00000120647.9 | CCDC77 | rs28850111 | 0.0000018 | ##### | Brain - Frontal Cortex (BA9) | 440788 | -59212 | 940788 | Yes |
| ENSG00000120647.9 | CCDC77 | rs61907030 | 0.0000018 | ##### | Brain - Nucleus accumbens (b | 440788 | -59212 | 940788 | Yes |
| ENSG00000120647.9 | CCDC77 | rs61907031 | 0.0000018 | ##### | Brain - Nucleus accumbens (b | 440788 | -59212 | 940788 | Yes |
| ENSG00000120647.9 | CCDC77 | rs11615564 | 0.0000018 | ##### | Brain - Nucleus accumbens (b | 440788 | -59212 | 940788 | Yes |
| ENSG00000120647.9 | CCDC77 | rs7303180 | 0.0000019 | ##### | Brain - Cerebellum | 440788 | -59212 | 940788 | Yes |
| ENSG00000120647.9 | CCDC77 | rs7136957 | 0.0000019 | ##### | Brain - Anterior cingulate cort | 440788 | -59212 | 940788 | Yes |
| ENSG00000120647.9 | CCDC77 | rs17800720 | 0.000002 | ##### | Brain - Substantia nigra | 440788 | -59212 | 940788 | Yes |
| ENSG00000120647.9 | CCDC77 | rs3837515 | 0.000002 | ##### | Brain - Substantia nigra | 440788 | -59212 | 940788 | Yes |
| ENSG00000120647.9 | CCDC77 | rs524468 | 0.000002 | ##### | Brain - Anterior cingulate cort | 440788 | -59212 | 940788 | Yes |
| ENSG00000120647.9 | CCDC77 | rs4980886 | 0.000002 | ##### | Brain - Cerebellum | 440788 | -59212 | 940788 | Yes |

| | | | | | | | | | |
|---|---|---|---|---|---|---|---|---|---|
| ENSG00000120647.9 | CCDC77 | rs525381 | 0.000002 | ##### | Brain - Anterior cingulate cort | 440788 | -59212 | 940788 | Yes |
| ENSG00000120647.9 | CCDC77 | rs10848766 | 0.0000021 | ##### | Brain - Cortex | 440788 | -59212 | 940788 | Yes |
| ENSG00000120647.9 | CCDC77 | rs11062357 | 0.0000021 | ##### | Brain - Cerebellar Hemisphere | 440788 | -59212 | 940788 | Yes |
| ENSG00000120647.9 | CCDC77 | rs1860361 | 0.0000021 | ##### | Brain - Caudate (basal ganglia | 440788 | -59212 | 940788 | Yes |
| ENSG00000120647.9 | CCDC77 | rs1548904 | 0.0000022 | ##### | Brain - Putamen (basal gangli | 440788 | -59212 | 940788 | Yes |
| ENSG00000120647.9 | CCDC77 | rs1972582 | 0.0000022 | ##### | Brain - Putamen (basal gangli | 440788 | -59212 | 940788 | Yes |
| ENSG00000120647.9 | CCDC77 | rs10848647 | 0.0000022 | ##### | Brain - Putamen (basal gangli | 440788 | -59212 | 940788 | Yes |
| ENSG00000120647.9 | CCDC77 | rs10774038 | 0.0000022 | ##### | Brain - Putamen (basal gangli | 440788 | -59212 | 940788 | Yes |
| ENSG00000120647.9 | CCDC77 | rs2041298 | 0.0000023 | ##### | Brain - Substantia nigra | 440788 | -59212 | 940788 | Yes |
| ENSG00000120647.9 | CCDC77 | rs12301816 | 0.0000024 | ##### | Brain - Cerebellum | 440788 | -59212 | 940788 | Yes |
| ENSG00000120647.9 | CCDC77 | rs12372234 | 0.0000024 | ##### | Brain - Cerebellar Hemisphere | 440788 | -59212 | 940788 | Yes |
| ENSG00000120647.9 | CCDC77 | rs10848987 | 0.0000024 | ##### | Brain - Caudate (basal ganglia | 440788 | -59212 | 940788 | Yes |
| ENSG00000120647.9 | CCDC77 | rs10848814 | 0.0000024 | ##### | Brain - Cortex | 440788 | -59212 | 940788 | Yes |
| ENSG00000120647.9 | CCDC77 | rs10848815 | 0.0000024 | ##### | Brain - Cortex | 440788 | -59212 | 940788 | Yes |
| ENSG00000120647.9 | CCDC77 | rs887311 | 0.0000024 | ##### | Brain - Cortex | 440788 | -59212 | 940788 | Yes |
| ENSG00000120647.9 | CCDC77 | rs77074951 | 0.0000024 | ##### | Brain - Cerebellum | 440788 | -59212 | 940788 | Yes |
| ENSG00000120647.9 | CCDC77 | rs4980906 | 0.0000025 | ##### | Brain - Cerebellar Hemisphere | 440788 | -59212 | 940788 | Yes |
| ENSG00000120647.9 | CCDC77 | rs10848644 | 0.0000025 | ##### | Brain - Hypothalamus | 440788 | -59212 | 940788 | Yes |
| ENSG00000120647.9 | CCDC77 | rs56042953 | 0.0000025 | ##### | Brain - Spinal cord (cervical c- | 440788 | -59212 | 940788 | Yes |
| ENSG00000120647.9 | CCDC77 | rs11062929 | 0.0000025 | ##### | Brain - Hypothalamus | 440788 | -59212 | 940788 | Yes |
| ENSG00000120647.9 | CCDC77 | rs10431354 | 0.0000025 | ##### | Brain - Nucleus accumbens (b | 440788 | -59212 | 940788 | Yes |
| ENSG00000120647.9 | CCDC77 | rs1860361 | 0.0000025 | ##### | Brain - Cerebellar Hemisphere | 440788 | -59212 | 940788 | Yes |
| ENSG00000120647.9 | CCDC77 | rs6489477 | 0.0000026 | ##### | Brain - Anterior cingulate cort | 440788 | -59212 | 940788 | Yes |
| ENSG00000120647.9 | CCDC77 | rs6489478 | 0.0000026 | ##### | Brain - Anterior cingulate cort | 440788 | -59212 | 940788 | Yes |
| ENSG00000120647.9 | CCDC77 | rs7965303 | 0.0000026 | ##### | Brain - Anterior cingulate cort | 440788 | -59212 | 940788 | Yes |
| ENSG00000120647.9 | CCDC77 | rs10848879 | 0.0000026 | ##### | Brain - Anterior cingulate cort | 440788 | -59212 | 940788 | Yes |
| ENSG00000120647.9 | CCDC77 | rs4140962 | 0.0000026 | ##### | Brain - Frontal Cortex (BA9) | 440788 | -59212 | 940788 | Yes |
| ENSG00000120647.9 | CCDC77 | rs17800738 | 0.0000026 | ##### | Brain - Cerebellar Hemisphere | 440788 | -59212 | 940788 | Yes |
| ENSG00000120647.9 | CCDC77 | rs112382198 | 0.0000026 | ##### | Brain - Cerebellar Hemisphere | 440788 | -59212 | 940788 | Yes |
| ENSG00000120647.9 | CCDC77 | rs151213767 | 0.0000026 | ##### | Brain - Cerebellar Hemisphere | 440788 | -59212 | 940788 | Yes |
| ENSG00000120647.9 | CCDC77 | rs10848814 | 0.0000027 | ##### | Brain - Cerebellum | 440788 | -59212 | 940788 | Yes |
| ENSG00000120647.9 | CCDC77 | rs10848815 | 0.0000027 | ##### | Brain - Cerebellum | 440788 | -59212 | 940788 | Yes |

| | | | | | | | | | |
|---|---|---|---|---|---|---|---|---|---|
| ENSG00000120647.9 | CCDC77 | rs887311 | 0.0000027 | ##### | Brain - Cerebellum | 440788 | -59212 | 940788 | Yes |
| ENSG00000120647.9 | CCDC77 | rs2080403 | 0.0000027 | ##### | Brain - Hypothalamus | 440788 | -59212 | 940788 | Yes |
| ENSG00000120647.9 | CCDC77 | rs2289957 | 0.0000027 | ##### | Brain - Hypothalamus | 440788 | -59212 | 940788 | Yes |
| ENSG00000120647.9 | CCDC77 | rs55657100 | 0.0000027 | ##### | Brain - Cortex | 440788 | -59212 | 940788 | Yes |
| ENSG00000120647.9 | CCDC77 | rs576994 | 0.0000027 | ##### | Brain - Anterior cingulate cort | 440788 | -59212 | 940788 | Yes |
| ENSG00000120647.9 | CCDC77 | rs797759 | 0.0000027 | ##### | Brain - Anterior cingulate cort | 440788 | -59212 | 940788 | Yes |
| ENSG00000120647.9 | CCDC77 | rs479836 | 0.0000027 | ##### | Brain - Anterior cingulate cort | 440788 | -59212 | 940788 | Yes |
| ENSG00000120647.9 | CCDC77 | rs2284332 | 0.0000027 | ##### | Brain - Anterior cingulate cort | 440788 | -59212 | 940788 | Yes |
| ENSG00000120647.9 | CCDC77 | rs2229351 | 0.0000027 | ##### | Brain - Anterior cingulate cort | 440788 | -59212 | 940788 | Yes |
| ENSG00000120647.9 | CCDC77 | rs4980810 | 0.0000027 | ##### | Brain - Anterior cingulate cort | 440788 | -59212 | 940788 | Yes |
| ENSG00000120647.9 | CCDC77 | rs11062302 | 0.0000027 | ##### | Brain - Anterior cingulate cort | 440788 | -59212 | 940788 | Yes |
| ENSG00000120647.9 | CCDC77 | rs12314497 | 0.0000027 | ##### | Brain - Anterior cingulate cort | 440788 | -59212 | 940788 | Yes |
| ENSG00000120647.9 | CCDC77 | rs534925791 | 0.0000027 | ##### | Brain - Anterior cingulate cort | 440788 | -59212 | 940788 | Yes |
| ENSG00000120647.9 | CCDC77 | rs144729439 | 0.0000027 | ##### | Brain - Anterior cingulate cort | 440788 | -59212 | 940788 | Yes |
| ENSG00000120647.9 | CCDC77 | rs11062385 | 0.0000027 | ##### | Brain - Anterior cingulate cort | 440788 | -59212 | 940788 | Yes |
| ENSG00000120647.9 | CCDC77 | rs11611810 | 0.0000027 | ##### | Brain - Anterior cingulate cort | 440788 | -59212 | 940788 | Yes |
| ENSG00000120647.9 | CCDC77 | rs10848715 | 0.0000027 | ##### | Brain - Anterior cingulate cort | 440788 | -59212 | 940788 | Yes |
| ENSG00000120647.9 | CCDC77 | rs10848716 | 0.0000027 | ##### | Brain - Anterior cingulate cort | 440788 | -59212 | 940788 | Yes |
| ENSG00000120647.9 | CCDC77 | rs61907027 | 0.0000027 | ##### | Brain - Anterior cingulate cort | 440788 | -59212 | 940788 | Yes |
| ENSG00000120647.9 | CCDC77 | rs7301422 | 0.0000027 | ##### | Brain - Anterior cingulate cort | 440788 | -59212 | 940788 | Yes |
| ENSG00000120647.9 | CCDC77 | rs11062512 | 0.0000027 | ##### | Brain - Anterior cingulate cort | 440788 | -59212 | 940788 | Yes |
| ENSG00000120647.9 | CCDC77 | rs3759371 | 0.0000027 | ##### | Brain - Anterior cingulate cort | 440788 | -59212 | 940788 | Yes |
| ENSG00000120647.9 | CCDC77 | rs6489448 | 0.0000027 | ##### | Brain - Anterior cingulate cort | 440788 | -59212 | 940788 | Yes |
| ENSG00000120647.9 | CCDC77 | rs7316660 | 0.0000027 | ##### | Brain - Anterior cingulate cort | 440788 | -59212 | 940788 | Yes |
| ENSG00000120647.9 | CCDC77 | rs6489449 | 0.0000027 | ##### | Brain - Anterior cingulate cort | 440788 | -59212 | 940788 | Yes |
| ENSG00000120647.9 | CCDC77 | rs10735021 | 0.0000027 | ##### | Brain - Anterior cingulate cort | 440788 | -59212 | 940788 | Yes |
| ENSG00000120647.9 | CCDC77 | rs10848845 | 0.0000027 | ##### | Brain - Anterior cingulate cort | 440788 | -59212 | 940788 | Yes |
| ENSG00000120647.9 | CCDC77 | rs10774137 | 0.0000027 | ##### | Brain - Anterior cingulate cort | 440788 | -59212 | 940788 | Yes |
| ENSG00000120647.9 | CCDC77 | rs11835325 | 0.0000027 | ##### | Brain - Anterior cingulate cort | 440788 | -59212 | 940788 | Yes |
| ENSG00000120647.9 | CCDC77 | rs10848849 | 0.0000027 | ##### | Brain - Anterior cingulate cort | 440788 | -59212 | 940788 | Yes |
| ENSG00000120647.9 | CCDC77 | rs6489454 | 0.0000027 | ##### | Brain - Anterior cingulate cort | 440788 | -59212 | 940788 | Yes |
| ENSG00000120647.9 | CCDC77 | rs7134608 | 0.0000027 | ##### | Brain - Anterior cingulate cort | 440788 | -59212 | 940788 | Yes |

| | | | | | | | | | |
|---|---|---|---|---|---|---|---|---|---|
| ENSG00000120647.9 | CCDC77 | rs7138890 | 0.0000027 | ##### | Brain - Anterior cingulate cort | 440788 | -59212 | 940788 | Yes |
| ENSG00000120647.9 | CCDC77 | rs10431355 | 0.0000027 | ##### | Brain - Anterior cingulate cort | 440788 | -59212 | 940788 | Yes |
| ENSG00000120647.9 | CCDC77 | rs6489458 | 0.0000027 | ##### | Brain - Anterior cingulate cort | 440788 | -59212 | 940788 | Yes |
| ENSG00000120647.9 | CCDC77 | rs2300134 | 0.0000027 | ##### | Brain - Anterior cingulate cort | 440788 | -59212 | 940788 | Yes |
| ENSG00000120647.9 | CCDC77 | rs3782852 | 0.0000027 | ##### | Brain - Anterior cingulate cort | 440788 | -59212 | 940788 | Yes |
| ENSG00000120647.9 | CCDC77 | rs201946814 | 0.0000027 | ##### | Brain - Anterior cingulate cort | 440788 | -59212 | 940788 | Yes |
| ENSG00000120647.9 | CCDC77 | rs6489463 | 0.0000027 | ##### | Brain - Anterior cingulate cort | 440788 | -59212 | 940788 | Yes |
| ENSG00000120647.9 | CCDC77 | rs2079785 | 0.0000027 | ##### | Brain - Anterior cingulate cort | 440788 | -59212 | 940788 | Yes |
| ENSG00000120647.9 | CCDC77 | rs7963937 | 0.0000027 | ##### | Brain - Anterior cingulate cort | 440788 | -59212 | 940788 | Yes |
| ENSG00000120647.9 | CCDC77 | rs2369276 | 0.0000027 | ##### | Brain - Anterior cingulate cort | 440788 | -59212 | 940788 | Yes |
| ENSG00000120647.9 | CCDC77 | rs7953137 | 0.0000027 | ##### | Brain - Anterior cingulate cort | 440788 | -59212 | 940788 | Yes |
| ENSG00000120647.9 | CCDC77 | rs6489471 | 0.0000027 | ##### | Brain - Anterior cingulate cort | 440788 | -59212 | 940788 | Yes |
| ENSG00000120647.9 | CCDC77 | rs7312065 | 0.0000027 | ##### | Brain - Anterior cingulate cort | 440788 | -59212 | 940788 | Yes |
| ENSG00000120647.9 | CCDC77 | rs34380840 | 0.0000027 | ##### | Brain - Anterior cingulate cort | 440788 | -59212 | 940788 | Yes |
| ENSG00000120647.9 | CCDC77 | rs482514 | 0.0000027 | ##### | Brain - Anterior cingulate cort | 440788 | -59212 | 940788 | Yes |
| ENSG00000120647.9 | CCDC77 | rs7315852 | 0.0000027 | ##### | Brain - Anterior cingulate cort | 440788 | -59212 | 940788 | Yes |
| ENSG00000120647.9 | CCDC77 | rs2284334 | 0.0000027 | ##### | Brain - Anterior cingulate cort | 440788 | -59212 | 940788 | Yes |
| ENSG00000120647.9 | CCDC77 | rs720128 | 0.0000027 | ##### | Brain - Anterior cingulate cort | 440788 | -59212 | 940788 | Yes |
| ENSG00000120647.9 | CCDC77 | rs9739950 | 0.0000027 | ##### | Brain - Anterior cingulate cort | 440788 | -59212 | 940788 | Yes |
| ENSG00000120647.9 | CCDC77 | rs12311569 | 0.0000027 | ##### | Brain - Anterior cingulate cort | 440788 | -59212 | 940788 | Yes |
| ENSG00000120647.9 | CCDC77 | rs10161265 | 0.0000028 | ##### | Brain - Cerebellar Hemisphere | 440788 | -59212 | 940788 | Yes |
| ENSG00000120647.9 | CCDC77 | rs4980897 | 0.0000028 | ##### | Brain - Cerebellar Hemisphere | 440788 | -59212 | 940788 | Yes |
| ENSG00000120647.9 | CCDC77 | rs10848766 | 0.0000028 | ##### | Brain - Hypothalamus | 440788 | -59212 | 940788 | Yes |
| ENSG00000120647.9 | CCDC77 | rs3213750 | 0.0000028 | ##### | Brain - Cortex | 440788 | -59212 | 940788 | Yes |
| ENSG00000120647.9 | CCDC77 | rs3753156 | 0.0000028 | ##### | Brain - Cortex | 440788 | -59212 | 940788 | Yes |
| ENSG00000120647.9 | CCDC77 | rs12581260 | 0.0000028 | ##### | Brain - Cortex | 440788 | -59212 | 940788 | Yes |
| ENSG00000120647.9 | CCDC77 | rs4980881 | 0.0000028 | ##### | Brain - Cortex | 440788 | -59212 | 940788 | Yes |
| ENSG00000120647.9 | CCDC77 | rs16929783 | 0.0000028 | ##### | Brain - Cortex | 440788 | -59212 | 940788 | Yes |
| ENSG00000120647.9 | CCDC77 | rs10848905 | 0.0000028 | ##### | Brain - Anterior cingulate cort | 440788 | -59212 | 940788 | Yes |
| ENSG00000120647.9 | CCDC77 | rs7295011 | 0.0000029 | ##### | Brain - Spinal cord (cervical c- | 440788 | -59212 | 940788 | Yes |
| ENSG00000120647.9 | CCDC77 | rs17800720 | 0.0000029 | ##### | Brain - Hypothalamus | 440788 | -59212 | 940788 | Yes |
| ENSG00000120647.9 | CCDC77 | rs3837515 | 0.0000029 | ##### | Brain - Hypothalamus | 440788 | -59212 | 940788 | Yes |

| | | | | | | | | | |
|---|---|---|---|---|---|---|---|---|---|
| ENSG00000120647.9 | CCDC77 | rs2041299 | 0.000003 | ##### | Brain - Substantia nigra | 440788 | -59212 | 940788 | Yes |
| ENSG00000120647.9 | CCDC77 | rs487252 | 0.000003 | ##### | Brain - Putamen (basal gangli | 440788 | -59212 | 940788 | Yes |
| ENSG00000120647.9 | CCDC77 | rs3764876 | 0.000003 | ##### | Brain - Spinal cord (cervical c- | 440788 | -59212 | 940788 | Yes |
| ENSG00000120647.9 | CCDC77 | rs10774191 | 0.000003 | ##### | Brain - Spinal cord (cervical c- | 440788 | -59212 | 940788 | Yes |
| ENSG00000120647.9 | CCDC77 | rs61916593 | 0.000003 | ##### | Brain - Spinal cord (cervical c- | 440788 | -59212 | 940788 | Yes |
| ENSG00000120647.9 | CCDC77 | rs12229673 | 0.0000031 | ##### | Brain - Cerebellar Hemisphere | 440788 | -59212 | 940788 | Yes |
| ENSG00000120647.9 | CCDC77 | rs12231564 | 0.0000031 | ##### | Brain - Cerebellar Hemisphere | 440788 | -59212 | 940788 | Yes |
| ENSG00000120647.9 | CCDC77 | rs2286606 | 0.0000031 | ##### | Brain - Hypothalamus | 440788 | -59212 | 940788 | Yes |
| ENSG00000120647.9 | CCDC77 | rs524468 | 0.0000032 | ##### | Brain - Cerebellum | 440788 | -59212 | 940788 | Yes |
| ENSG00000120647.9 | CCDC77 | rs12301816 | 0.0000032 | ##### | Brain - Amygdala | 440788 | -59212 | 940788 | Yes |
| ENSG00000120647.9 | CCDC77 | rs113574411 | 0.0000032 | ##### | Brain - Cerebellum | 440788 | -59212 | 940788 | Yes |
| ENSG00000120647.9 | CCDC77 | rs479840 | 0.0000032 | ##### | Brain - Cerebellum | 440788 | -59212 | 940788 | Yes |
| ENSG00000120647.9 | CCDC77 | rs479803 | 0.0000032 | ##### | Brain - Cerebellum | 440788 | -59212 | 940788 | Yes |
| ENSG00000120647.9 | CCDC77 | rs4980811 | 0.0000032 | ##### | Brain - Cerebellum | 440788 | -59212 | 940788 | Yes |
| ENSG00000120647.9 | CCDC77 | rs72472276 | 0.0000032 | ##### | Brain - Cerebellum | 440788 | -59212 | 940788 | Yes |
| ENSG00000120647.9 | CCDC77 | rs4980877 | 0.0000032 | ##### | Brain - Cerebellum | 440788 | -59212 | 940788 | Yes |
| ENSG00000120647.9 | CCDC77 | rs7963309 | 0.0000032 | ##### | Brain - Cerebellum | 440788 | -59212 | 940788 | Yes |
| ENSG00000120647.9 | CCDC77 | rs7134353 | 0.0000032 | ##### | Brain - Cerebellum | 440788 | -59212 | 940788 | Yes |
| ENSG00000120647.9 | CCDC77 | rs11374245 | 0.0000032 | ##### | Brain - Cerebellum | 440788 | -59212 | 940788 | Yes |
| ENSG00000120647.9 | CCDC77 | rs16929744 | 0.0000032 | ##### | Brain - Cerebellum | 440788 | -59212 | 940788 | Yes |
| ENSG00000120647.9 | CCDC77 | rs10848877 | 0.0000033 | ##### | Brain - Anterior cingulate cort | 440788 | -59212 | 940788 | Yes |
| ENSG00000120647.9 | CCDC77 | rs576159 | 0.0000033 | ##### | Brain - Anterior cingulate cort | 440788 | -59212 | 940788 | Yes |
| ENSG00000120647.9 | CCDC77 | rs11062943 | 0.0000033 | ##### | Brain - Cerebellar Hemisphere | 440788 | -59212 | 940788 | Yes |
| ENSG00000120647.9 | CCDC77 | rs4980819 | 0.0000033 | ##### | Brain - Cerebellar Hemisphere | 440788 | -59212 | 940788 | Yes |
| ENSG00000120647.9 | CCDC77 | rs10848905 | 0.0000034 | ##### | Brain - Hippocampus | 440788 | -59212 | 940788 | Yes |
| ENSG00000120647.9 | CCDC77 | rs11062934 | 0.0000035 | ##### | Brain - Amygdala | 440788 | -59212 | 940788 | Yes |
| ENSG00000120647.9 | CCDC77 | rs10082788 | 0.0000035 | ##### | Brain - Amygdala | 440788 | -59212 | 940788 | Yes |
| ENSG00000120647.9 | CCDC77 | rs61908460 | 0.0000035 | ##### | Brain - Cerebellar Hemisphere | 440788 | -59212 | 940788 | Yes |
| ENSG00000120647.9 | CCDC77 | rs520932 | 0.0000037 | ##### | Brain - Hippocampus | 440788 | -59212 | 940788 | Yes |
| ENSG00000120647.9 | CCDC77 | rs11062167 | 0.0000038 | ##### | Brain - Cortex | 440788 | -59212 | 940788 | Yes |
| ENSG00000120647.9 | CCDC77 | rs55990424 | 0.0000038 | ##### | Brain - Cortex | 440788 | -59212 | 940788 | Yes |
| ENSG00000120647.9 | CCDC77 | rs10082788 | 0.0000038 | ##### | Brain - Anterior cingulate cort | 440788 | -59212 | 940788 | Yes |

| | | | | | | | | | |
|---|---|---|---|---|---|---|---|---|---|
| ENSG00000120647.9 | CCDC77 | rs4140962 | 0.0000039 | ##### | Brain - Putamen (basal gangli | 440788 | -59212 | 940788 | Yes |
| ENSG00000120647.9 | CCDC77 | rs10744627 | 0.0000039 | ##### | Brain - Hippocampus | 440788 | -59212 | 940788 | Yes |
| ENSG00000120647.9 | CCDC77 | rs3782860 | 0.000004 | ##### | Brain - Hypothalamus | 440788 | -59212 | 940788 | Yes |
| ENSG00000120647.9 | CCDC77 | rs579526 | 0.0000041 | ##### | Brain - Substantia nigra | 440788 | -59212 | 940788 | Yes |
| ENSG00000120647.9 | CCDC77 | rs1048466 | 0.0000042 | ##### | Brain - Spinal cord (cervical c- | 440788 | -59212 | 940788 | Yes |
| ENSG00000120647.9 | CCDC77 | rs7969761 | 0.0000043 | ##### | Brain - Putamen (basal gangli | 440788 | -59212 | 940788 | Yes |
| ENSG00000120647.9 | CCDC77 | rs2080403 | 0.0000046 | ##### | Brain - Putamen (basal gangli | 440788 | -59212 | 940788 | Yes |
| ENSG00000120647.9 | CCDC77 | rs2289957 | 0.0000046 | ##### | Brain - Putamen (basal gangli | 440788 | -59212 | 940788 | Yes |
| ENSG00000120647.9 | CCDC77 | rs10848644 | 0.0000047 | ##### | Brain - Substantia nigra | 440788 | -59212 | 940788 | Yes |
| ENSG00000120647.9 | CCDC77 | rs4980908 | 0.0000048 | ##### | Brain - Nucleus accumbens (b | 440788 | -59212 | 940788 | Yes |
| ENSG00000120647.9 | CCDC77 | rs507597 | 0.0000049 | ##### | Brain - Amygdala | 440788 | -59212 | 940788 | Yes |
| ENSG00000120647.9 | CCDC77 | rs17800744 | 0.0000049 | ##### | Brain - Amygdala | 440788 | -59212 | 940788 | Yes |
| ENSG00000120647.9 | CCDC77 | rs77854644 | 0.0000049 | ##### | Brain - Amygdala | 440788 | -59212 | 940788 | Yes |
| ENSG00000120647.9 | CCDC77 | rs12229673 | 0.000005 | ##### | Brain - Hypothalamus | 440788 | -59212 | 940788 | Yes |
| ENSG00000120647.9 | CCDC77 | rs12231564 | 0.000005 | ##### | Brain - Hypothalamus | 440788 | -59212 | 940788 | Yes |
| ENSG00000120647.9 | CCDC77 | rs525381 | 0.000005 | ##### | Brain - Hippocampus | 440788 | -59212 | 940788 | Yes |
| ENSG00000120647.9 | CCDC77 | rs2191159 | 0.000005 | ##### | Brain - Spinal cord (cervical c- | 440788 | -59212 | 940788 | Yes |
| ENSG00000120647.9 | CCDC77 | rs111250603 | 0.0000051 | ##### | Brain - Hippocampus | 440788 | -59212 | 940788 | Yes |
| ENSG00000120647.9 | CCDC77 | rs2369277 | 0.0000051 | ##### | Brain - Hippocampus | 440788 | -59212 | 940788 | Yes |
| ENSG00000120647.9 | CCDC77 | rs9805067 | 0.0000051 | ##### | Brain - Hippocampus | 440788 | -59212 | 940788 | Yes |
| ENSG00000120647.9 | CCDC77 | rs1541464 | 0.0000051 | ##### | Brain - Hippocampus | 440788 | -59212 | 940788 | Yes |
| ENSG00000120647.9 | CCDC77 | rs1541465 | 0.0000051 | ##### | Brain - Hippocampus | 440788 | -59212 | 940788 | Yes |
| ENSG00000120647.9 | CCDC77 | rs4980817 | 0.0000051 | ##### | Brain - Hippocampus | 440788 | -59212 | 940788 | Yes |
| ENSG00000120647.9 | CCDC77 | rs4482086 | 0.0000051 | ##### | Brain - Hippocampus | 440788 | -59212 | 940788 | Yes |
| ENSG00000120647.9 | CCDC77 | rs2369279 | 0.0000051 | ##### | Brain - Hippocampus | 440788 | -59212 | 940788 | Yes |
| ENSG00000120647.9 | CCDC77 | rs11062803 | 0.0000051 | ##### | Brain - Hippocampus | 440788 | -59212 | 940788 | Yes |
| ENSG00000120647.9 | CCDC77 | rs11062943 | 0.0000051 | ##### | Brain - Amygdala | 440788 | -59212 | 940788 | Yes |
| ENSG00000120647.9 | CCDC77 | rs4980819 | 0.0000051 | ##### | Brain - Amygdala | 440788 | -59212 | 940788 | Yes |
| ENSG00000120647.9 | CCDC77 | | 0.0000052 | ##### | Brain - Hippocampus | 440788 | -59212 | 940788 | Yes |
| ENSG00000120647.9 | CCDC77 | rs4980906 | 0.0000052 | ##### | Brain - Cortex | 440788 | -59212 | 940788 | Yes |
| ENSG00000120647.9 | CCDC77 | rs495973 | 0.0000052 | ##### | Brain - Hippocampus | 440788 | -59212 | 940788 | Yes |
| ENSG00000120647.9 | CCDC77 | rs11613826 | 0.0000053 | ##### | Brain - Caudate (basal ganglia | 440788 | -59212 | 940788 | Yes |

| | | | | | | | | | |
|---|---|---|---|---|---|---|---|---|---|
| ENSG00000120647.9 | CCDC77 | rs11062125 | 0.0000053 | ##### | Brain - Caudate (basal ganglia | 440788 | -59212 | 940788 | Yes |
| ENSG00000120647.9 | CCDC77 | rs11062126 | 0.0000053 | ##### | Brain - Caudate (basal ganglia | 440788 | -59212 | 940788 | Yes |
| ENSG00000120647.9 | CCDC77 | rs35311301 | 0.0000053 | ##### | Brain - Caudate (basal ganglia | 440788 | -59212 | 940788 | Yes |
| ENSG00000120647.9 | CCDC77 | rs12825287 | 0.0000053 | ##### | Brain - Caudate (basal ganglia | 440788 | -59212 | 940788 | Yes |
| ENSG00000120647.9 | CCDC77 | rs12366824 | 0.0000053 | ##### | Brain - Caudate (basal ganglia | 440788 | -59212 | 940788 | Yes |
| ENSG00000120647.9 | CCDC77 | rs10848638 | 0.0000053 | ##### | Brain - Caudate (basal ganglia | 440788 | -59212 | 940788 | Yes |
| ENSG00000120647.9 | CCDC77 | rs11062934 | 0.0000054 | ##### | Brain - Hypothalamus | 440788 | -59212 | 940788 | Yes |
| ENSG00000120647.9 | CCDC77 | rs1541463 | 0.0000054 | ##### | Brain - Hippocampus | 440788 | -59212 | 940788 | Yes |
| ENSG00000120647.9 | CCDC77 | rs2369278 | 0.0000054 | ##### | Brain - Hippocampus | 440788 | -59212 | 940788 | Yes |
| ENSG00000120647.9 | CCDC77 | rs10774163 | 0.0000054 | ##### | Brain - Hippocampus | 440788 | -59212 | 940788 | Yes |
| ENSG00000120647.9 | CCDC77 | rs10774190 | 0.0000054 | ##### | Brain - Cerebellar Hemisphere | 440788 | -59212 | 940788 | Yes |
| ENSG00000120647.9 | CCDC77 | rs3753156 | 0.0000056 | ##### | Brain - Cerebellum | 440788 | -59212 | 940788 | Yes |
| ENSG00000120647.9 | CCDC77 | rs12581260 | 0.0000056 | ##### | Brain - Cerebellum | 440788 | -59212 | 940788 | Yes |
| ENSG00000120647.9 | CCDC77 | rs16929783 | 0.0000056 | ##### | Brain - Cerebellum | 440788 | -59212 | 940788 | Yes |
| ENSG00000120647.9 | CCDC77 | rs7310504 | 0.0000056 | ##### | Brain - Cerebellar Hemisphere | 440788 | -59212 | 940788 | Yes |
| ENSG00000120647.9 | CCDC77 | rs10848961 | 0.0000059 | ##### | Brain - Frontal Cortex (BA9) | 440788 | -59212 | 940788 | Yes |
| ENSG00000120647.9 | CCDC77 | rs12826998 | 0.0000059 | ##### | Brain - Caudate (basal ganglia | 440788 | -59212 | 940788 | Yes |
| ENSG00000120647.9 | CCDC77 | rs11062929 | 0.000006 | ##### | Brain - Cerebellar Hemisphere | 440788 | -59212 | 940788 | Yes |
| ENSG00000120647.9 | CCDC77 | rs10774192 | 0.0000062 | ##### | Brain - Nucleus accumbens (b | 440788 | -59212 | 940788 | Yes |
| ENSG00000120647.9 | CCDC77 | rs11613331 | 0.0000062 | ##### | Brain - Putamen (basal gangli | 440788 | -59212 | 940788 | Yes |
| ENSG00000120647.9 | CCDC77 | rs10848639 | 0.0000063 | ##### | Brain - Caudate (basal ganglia | 440788 | -59212 | 940788 | Yes |
| ENSG00000120647.9 | CCDC77 | rs10848766 | 0.0000065 | ##### | Brain - Cerebellum | 440788 | -59212 | 940788 | Yes |
| ENSG00000120647.9 | CCDC77 | rs797765 | 0.0000066 | ##### | Brain - Substantia nigra | 440788 | -59212 | 940788 | Yes |
| ENSG00000120647.9 | CCDC77 | rs3213750 | 0.0000068 | ##### | Brain - Cerebellum | 440788 | -59212 | 940788 | Yes |
| ENSG00000120647.9 | CCDC77 | rs4980881 | 0.0000068 | ##### | Brain - Cerebellum | 440788 | -59212 | 940788 | Yes |
| ENSG00000120647.9 | CCDC77 | rs579526 | 0.0000068 | ##### | Brain - Anterior cingulate cort | 440788 | -59212 | 940788 | Yes |
| ENSG00000120647.9 | CCDC77 | rs10848644 | 0.0000069 | ##### | Brain - Cortex | 440788 | -59212 | 940788 | Yes |
| ENSG00000120647.9 | CCDC77 | rs11062943 | 0.000007 | ##### | Brain - Frontal Cortex (BA9) | 440788 | -59212 | 940788 | Yes |
| ENSG00000120647.9 | CCDC77 | rs4980819 | 0.000007 | ##### | Brain - Frontal Cortex (BA9) | 440788 | -59212 | 940788 | Yes |
| ENSG00000120647.9 | CCDC77 | rs4980878 | 0.0000071 | ##### | Brain - Hippocampus | 440788 | -59212 | 940788 | Yes |
| ENSG00000120647.9 | CCDC77 | rs10431354 | 0.0000071 | ##### | Brain - Anterior cingulate cort | 440788 | -59212 | 940788 | Yes |
| ENSG00000120647.9 | CCDC77 | rs4980908 | 0.0000073 | ##### | Brain - Cortex | 440788 | -59212 | 940788 | Yes |

| | | | | | | | | | |
|---|---|---|---|---|---|---|---|---|---|
| ENSG00000120647.9 | CCDC77 | rs372437080 | 0.0000073 | ##### | Brain - Substantia nigra | 440788 | -59212 | 940788 | Yes |
| ENSG00000120647.9 | CCDC77 | rs4980896 | 0.0000074 | ##### | Brain - Spinal cord (cervical c- | 440788 | -59212 | 940788 | Yes |
| ENSG00000120647.9 | CCDC77 | rs10744627 | 0.0000074 | ##### | Brain - Amygdala | 440788 | -59212 | 940788 | Yes |
| ENSG00000120647.9 | CCDC77 | rs11613331 | 0.0000076 | ##### | Brain - Hypothalamus | 440788 | -59212 | 940788 | Yes |
| ENSG00000120647.9 | CCDC77 | rs11062357 | 0.0000077 | ##### | Brain - Putamen (basal gangli | 440788 | -59212 | 940788 | Yes |
| ENSG00000120647.9 | CCDC77 | rs487252 | 0.0000077 | ##### | Brain - Substantia nigra | 440788 | -59212 | 940788 | Yes |
| ENSG00000120647.9 | CCDC77 | rs75894333 | 0.0000078 | ##### | Brain - Amygdala | 440788 | -59212 | 940788 | Yes |
| ENSG00000120647.9 | CCDC77 | rs11608938 | 0.0000078 | ##### | Brain - Amygdala | 440788 | -59212 | 940788 | Yes |
| ENSG00000120647.9 | CCDC77 | rs7965038 | 0.0000078 | ##### | Brain - Amygdala | 440788 | -59212 | 940788 | Yes |
| ENSG00000120647.9 | CCDC77 | rs7307606 | 0.0000078 | ##### | Brain - Amygdala | 440788 | -59212 | 940788 | Yes |
| ENSG00000120647.9 | CCDC77 | rs10661251 | 0.0000078 | ##### | Brain - Amygdala | 440788 | -59212 | 940788 | Yes |
| ENSG00000120647.9 | CCDC77 | rs1011699 | 0.0000078 | ##### | Brain - Amygdala | 440788 | -59212 | 940788 | Yes |
| ENSG00000120647.9 | CCDC77 | rs6489475 | 0.0000078 | ##### | Brain - Amygdala | 440788 | -59212 | 940788 | Yes |
| ENSG00000120647.9 | CCDC77 | rs10774154 | 0.0000078 | ##### | Brain - Amygdala | 440788 | -59212 | 940788 | Yes |
| ENSG00000120647.9 | CCDC77 | rs2286606 | 0.0000079 | ##### | Brain - Frontal Cortex (BA9) | 440788 | -59212 | 940788 | Yes |
| ENSG00000120647.9 | CCDC77 | rs7133796 | 0.0000079 | ##### | Brain - Hippocampus | 440788 | -59212 | 940788 | Yes |
| ENSG00000120647.9 | CCDC77 | rs56042953 | 0.0000079 | ##### | Brain - Putamen (basal gangli | 440788 | -59212 | 940788 | Yes |
| ENSG00000120647.9 | CCDC77 | rs7310504 | 0.0000081 | ##### | Brain - Spinal cord (cervical c- | 440788 | -59212 | 940788 | Yes |
| ENSG00000120647.9 | CCDC77 | rs524468 | 0.0000081 | ##### | Brain - Hippocampus | 440788 | -59212 | 940788 | Yes |
| ENSG00000120647.9 | CCDC77 | rs57022939 | 0.0000083 | ##### | Brain - Hippocampus | 440788 | -59212 | 940788 | Yes |
| ENSG00000120647.9 | CCDC77 | rs67523949 | 0.0000085 | ##### | Brain - Putamen (basal gangli | 440788 | -59212 | 940788 | Yes |
| ENSG00000120647.9 | CCDC77 | rs10431354 | 0.0000089 | ##### | Brain - Amygdala | 440788 | -59212 | 940788 | Yes |
| ENSG00000120647.9 | CCDC77 | rs497740 | 0.000009 | ##### | Brain - Caudate (basal ganglia | 440788 | -59212 | 940788 | Yes |
| ENSG00000120647.9 | CCDC77 | rs4980822 | 0.000009 | ##### | Brain - Cerebellar Hemisphere | 440788 | -59212 | 940788 | Yes |
| ENSG00000120647.9 | CCDC77 | rs12370433 | 0.0000092 | ##### | Brain - Putamen (basal gangli | 440788 | -59212 | 940788 | Yes |
| ENSG00000120647.9 | CCDC77 | rs4980898 | 0.0000092 | ##### | Brain - Nucleus accumbens (b | 440788 | -59212 | 940788 | Yes |
| ENSG00000120647.9 | CCDC77 | rs3213751 | 0.0000093 | ##### | Brain - Anterior cingulate cort | 440788 | -59212 | 940788 | Yes |
| ENSG00000120647.9 | CCDC77 | rs3815558 | 0.0000093 | ##### | Brain - Anterior cingulate cort | 440788 | -59212 | 940788 | Yes |
| ENSG00000120647.9 | CCDC77 | rs78998485 | 0.0000093 | ##### | Brain - Anterior cingulate cort | 440788 | -59212 | 940788 | Yes |
| ENSG00000120647.9 | CCDC77 | rs4980884 | 0.0000093 | ##### | Brain - Anterior cingulate cort | 440788 | -59212 | 940788 | Yes |
| ENSG00000120647.9 | CCDC77 | rs7135768 | 0.0000093 | ##### | Brain - Anterior cingulate cort | 440788 | -59212 | 940788 | Yes |
| ENSG00000120647.9 | CCDC77 | rs6489450 | 0.0000093 | ##### | Brain - Anterior cingulate cort | 440788 | -59212 | 940788 | Yes |

| | | | | | | | | | |
|---|---|---|---|---|---|---|---|---|---|
| ENSG00000120647.9 | CCDC77 | rs7310504 | 0.0000093 | ##### | Brain - Substantia nigra | 440788 | -59212 | 940788 | Yes |
| ENSG00000120647.9 | CCDC77 | rs4980896 | 0.0000095 | ##### | Brain - Substantia nigra | 440788 | -59212 | 940788 | Yes |
| ENSG00000120647.9 | CCDC77 | rs111250603 | 0.0000096 | ##### | Brain - Amygdala | 440788 | -59212 | 940788 | Yes |
| ENSG00000120647.9 | CCDC77 | rs2369277 | 0.0000096 | ##### | Brain - Amygdala | 440788 | -59212 | 940788 | Yes |
| ENSG00000120647.9 | CCDC77 | rs9805067 | 0.0000096 | ##### | Brain - Amygdala | 440788 | -59212 | 940788 | Yes |
| ENSG00000120647.9 | CCDC77 | rs1541464 | 0.0000096 | ##### | Brain - Amygdala | 440788 | -59212 | 940788 | Yes |
| ENSG00000120647.9 | CCDC77 | rs1541465 | 0.0000096 | ##### | Brain - Amygdala | 440788 | -59212 | 940788 | Yes |
| ENSG00000120647.9 | CCDC77 | rs4980817 | 0.0000096 | ##### | Brain - Amygdala | 440788 | -59212 | 940788 | Yes |
| ENSG00000120647.9 | CCDC77 | rs4482086 | 0.0000096 | ##### | Brain - Amygdala | 440788 | -59212 | 940788 | Yes |
| ENSG00000120647.9 | CCDC77 | rs2369279 | 0.0000096 | ##### | Brain - Amygdala | 440788 | -59212 | 940788 | Yes |
| ENSG00000120647.9 | CCDC77 | rs11062803 | 0.0000096 | ##### | Brain - Amygdala | 440788 | -59212 | 940788 | Yes |
| ENSG00000120647.9 | CCDC77 | rs10160995 | 0.0000097 | ##### | Brain - Nucleus accumbens (b | 440788 | -59212 | 940788 | Yes |
| ENSG00000120647.9 | CCDC77 | rs11062167 | 0.0000097 | ##### | Brain - Hypothalamus | 440788 | -59212 | 940788 | Yes |
| ENSG00000120647.9 | CCDC77 | rs55990424 | 0.0000097 | ##### | Brain - Hypothalamus | 440788 | -59212 | 940788 | Yes |
| ENSG00000120647.9 | CCDC77 | rs10848961 | 0.0000097 | ##### | Brain - Cerebellar Hemisphere | 440788 | -59212 | 940788 | Yes |
| ENSG00000120647.9 | CCDC77 | rs12229673 | 0.0000098 | ##### | Brain - Amygdala | 440788 | -59212 | 940788 | Yes |
| ENSG00000120647.9 | CCDC77 | rs12231564 | 0.0000098 | ##### | Brain - Amygdala | 440788 | -59212 | 940788 | Yes |
| ENSG00000120647.9 | CCDC77 | rs4980874 | 0.00001 | ##### | Brain - Cerebellum | 440788 | -59212 | 940788 | Yes |
| ENSG00000120647.9 | CCDC77 | rs7307754 | 0.00001 | ##### | Brain - Caudate (basal ganglia | 440788 | -59212 | 940788 | Yes |
| ENSG00000120647.9 | CCDC77 | rs520932 | 0.00001 | ##### | Brain - Amygdala | 440788 | -59212 | 940788 | Yes |
| ENSG00000120647.9 | CCDC77 | rs4140962 | 0.00001 | ##### | Brain - Cortex | 440788 | -59212 | 940788 | Yes |
| ENSG00000120647.9 | CCDC77 | rs1541463 | 0.00001 | ##### | Brain - Amygdala | 440788 | -59212 | 940788 | Yes |
| ENSG00000120647.9 | CCDC77 | rs2369278 | 0.00001 | ##### | Brain - Amygdala | 440788 | -59212 | 940788 | Yes |
| ENSG00000120647.9 | CCDC77 | rs10774163 | 0.00001 | ##### | Brain - Amygdala | 440788 | -59212 | 940788 | Yes |
| ENSG00000120647.9 | CCDC77 | rs10848594 | 0.000011 | ##### | Brain - Caudate (basal ganglia | 440788 | -59212 | 940788 | Yes |
| ENSG00000120647.9 | CCDC77 | rs11062943 | 0.000011 | ##### | Brain - Hypothalamus | 440788 | -59212 | 940788 | Yes |
| ENSG00000120647.9 | CCDC77 | rs4980819 | 0.000011 | ##### | Brain - Hypothalamus | 440788 | -59212 | 940788 | Yes |
| ENSG00000120647.9 | CCDC77 | rs7969761 | 0.000011 | ##### | Brain - Hypothalamus | 440788 | -59212 | 940788 | Yes |
| ENSG00000120647.9 | CCDC77 | rs7138958 | 0.000011 | ##### | Brain - Substantia nigra | 440788 | -59212 | 940788 | Yes |
| ENSG00000120647.9 | CCDC77 | rs2007537 | 0.000011 | ##### | Brain - Substantia nigra | 440788 | -59212 | 940788 | Yes |
| ENSG00000120647.9 | CCDC77 | rs1207322 | 0.000012 | ##### | Brain - Hippocampus | 440788 | -59212 | 940788 | Yes |
| ENSG00000120647.9 | CCDC77 | rs4980813 | 0.000012 | ##### | Brain - Hippocampus | 440788 | -59212 | 940788 | Yes |

| | | | | | | | | | |
|---|---|---|---|---|---|---|---|---|---|
| ENSG00000120647.9 | CCDC77 | rs61906996 | 0.000012 | ##### | Brain - Hippocampus | 440788 | -59212 | 940788 | Yes |
| ENSG00000120647.9 | CCDC77 | rs4980814 | 0.000012 | ##### | Brain - Hippocampus | 440788 | -59212 | 940788 | Yes |
| ENSG00000120647.9 | CCDC77 | rs7977462 | 0.000012 | ##### | Brain - Hippocampus | 440788 | -59212 | 940788 | Yes |
| ENSG00000120647.9 | CCDC77 | rs10744616 | 0.000012 | ##### | Brain - Hippocampus | 440788 | -59212 | 940788 | Yes |
| ENSG00000120647.9 | CCDC77 | rs10774150 | 0.000012 | ##### | Brain - Hippocampus | 440788 | -59212 | 940788 | Yes |
| ENSG00000120647.9 | CCDC77 | rs3831759 | 0.000012 | ##### | Brain - Hippocampus | 440788 | -59212 | 940788 | Yes |
| ENSG00000120647.9 | CCDC77 | rs11062934 | 0.000012 | ##### | Brain - Frontal Cortex (BA9) | 440788 | -59212 | 940788 | Yes |
| ENSG00000120647.9 | CCDC77 | rs2286606 | 0.000012 | ##### | Brain - Spinal cord (cervical c- | 440788 | -59212 | 940788 | Yes |
| ENSG00000120647.9 | CCDC77 | rs1048466 | 0.000012 | ##### | Brain - Putamen (basal gangli | 440788 | -59212 | 940788 | Yes |
| ENSG00000120647.9 | CCDC77 | rs4980875 | 0.000012 | ##### | Brain - Hippocampus | 440788 | -59212 | 940788 | Yes |
| ENSG00000120647.9 | CCDC77 | rs61906995 | 0.000012 | ##### | Brain - Hippocampus | 440788 | -59212 | 940788 | Yes |
| ENSG00000120647.9 | CCDC77 | rs12582850 | 0.000012 | ##### | Brain - Hippocampus | 440788 | -59212 | 940788 | Yes |
| ENSG00000120647.9 | CCDC77 | rs7298019 | 0.000012 | ##### | Brain - Hippocampus | 440788 | -59212 | 940788 | Yes |
| ENSG00000120647.9 | CCDC77 | rs2098094 | 0.000012 | ##### | Brain - Hippocampus | 440788 | -59212 | 940788 | Yes |
| ENSG00000120647.9 | CCDC77 | rs2159364 | 0.000012 | ##### | Brain - Hippocampus | 440788 | -59212 | 940788 | Yes |
| ENSG00000120647.9 | CCDC77 | rs10848868 | 0.000012 | ##### | Brain - Hippocampus | 440788 | -59212 | 940788 | Yes |
| ENSG00000120647.9 | CCDC77 | rs1860360 | 0.000012 | ##### | Brain - Hippocampus | 440788 | -59212 | 940788 | Yes |
| ENSG00000120647.9 | CCDC77 | rs10848905 | 0.000013 | ##### | Brain - Amygdala | 440788 | -59212 | 940788 | Yes |
| ENSG00000120647.9 | CCDC77 | rs2080402 | 0.000013 | ##### | Brain - Putamen (basal gangli | 440788 | -59212 | 940788 | Yes |
| ENSG00000120647.9 | CCDC77 | rs10774190 | 0.000013 | ##### | Brain - Hypothalamus | 440788 | -59212 | 940788 | Yes |
| ENSG00000120647.9 | CCDC77 | rs2253565 | 0.000013 | ##### | Brain - Amygdala | 440788 | -59212 | 940788 | Yes |
| ENSG00000120647.9 | CCDC77 | rs4980876 | 0.000014 | ##### | Brain - Hippocampus | 440788 | -59212 | 940788 | Yes |
| ENSG00000120647.9 | CCDC77 | rs372437080 | 0.000014 | ##### | Brain - Amygdala | 440788 | -59212 | 940788 | Yes |
| ENSG00000120647.9 | CCDC77 | rs797765 | 0.000014 | ##### | Brain - Amygdala | 440788 | -59212 | 940788 | Yes |
| ENSG00000120647.9 | CCDC77 | rs2041299 | 0.000014 | ##### | Brain - Amygdala | 440788 | -59212 | 940788 | Yes |
| ENSG00000120647.9 | CCDC77 | rs10774190 | 0.000014 | ##### | Brain - Frontal Cortex (BA9) | 440788 | -59212 | 940788 | Yes |
| ENSG00000120647.9 | CCDC77 | rs1155942 | 0.000014 | ##### | Brain - Hippocampus | 440788 | -59212 | 940788 | Yes |
| ENSG00000120647.9 | CCDC77 | rs10161265 | 0.000014 | ##### | Brain - Amygdala | 440788 | -59212 | 940788 | Yes |
| ENSG00000120647.9 | CCDC77 | rs4980897 | 0.000014 | ##### | Brain - Amygdala | 440788 | -59212 | 940788 | Yes |
| ENSG00000120647.9 | CCDC77 | rs4980822 | 0.000014 | ##### | Brain - Putamen (basal gangli | 440788 | -59212 | 940788 | Yes |
| ENSG00000120647.9 | CCDC77 | rs7963793 | 0.000014 | ##### | Brain - Anterior cingulate cort | 440788 | -59212 | 940788 | Yes |
| ENSG00000120647.9 | CCDC77 | rs10848814 | 0.000014 | ##### | Brain - Anterior cingulate cort | 440788 | -59212 | 940788 | Yes |

| | | | | | | | | | |
|---|---|---|---|---|---|---|---|---|---|
| ENSG00000120647.9 | CCDC77 | rs10848815 | 0.000014 | ##### | Brain - Anterior cingulate cort | 440788 | -59212 | 940788 | Yes |
| ENSG00000120647.9 | CCDC77 | rs887311 | 0.000014 | ##### | Brain - Anterior cingulate cort | 440788 | -59212 | 940788 | Yes |
| ENSG00000120647.9 | CCDC77 | rs11062945 | 0.000014 | ##### | Brain - Frontal Cortex (BA9) | 440788 | -59212 | 940788 | Yes |
| ENSG00000120647.9 | CCDC77 | rs7965303 | 0.000014 | ##### | Brain - Hippocampus | 440788 | -59212 | 940788 | Yes |
| ENSG00000120647.9 | CCDC77 | rs10848879 | 0.000014 | ##### | Brain - Hippocampus | 440788 | -59212 | 940788 | Yes |
| ENSG00000120647.9 | CCDC77 | rs2080403 | 0.000015 | ##### | Brain - Substantia nigra | 440788 | -59212 | 940788 | Yes |
| ENSG00000120647.9 | CCDC77 | rs2289957 | 0.000015 | ##### | Brain - Substantia nigra | 440788 | -59212 | 940788 | Yes |
| ENSG00000120647.9 | CCDC77 | rs10161265 | 0.000015 | ##### | Brain - Anterior cingulate cort | 440788 | -59212 | 940788 | Yes |
| ENSG00000120647.9 | CCDC77 | rs4980897 | 0.000015 | ##### | Brain - Anterior cingulate cort | 440788 | -59212 | 940788 | Yes |
| ENSG00000120647.9 | CCDC77 | rs67523949 | 0.000015 | ##### | Brain - Substantia nigra | 440788 | -59212 | 940788 | Yes |
| ENSG00000120647.9 | CCDC77 | rs4980906 | 0.000015 | ##### | Brain - Hypothalamus | 440788 | -59212 | 940788 | Yes |
| ENSG00000120647.9 | CCDC77 | rs61916594 | 0.000015 | ##### | Brain - Frontal Cortex (BA9) | 440788 | -59212 | 940788 | Yes |
| ENSG00000120647.9 | CCDC77 | rs2191159 | 0.000015 | ##### | Brain - Putamen (basal gangli | 440788 | -59212 | 940788 | Yes |
| ENSG00000120647.9 | CCDC77 | rs7972566 | 0.000015 | ##### | Brain - Hippocampus | 440788 | -59212 | 940788 | Yes |
| ENSG00000120647.9 | CCDC77 | rs10774170 | 0.000015 | ##### | Brain - Hippocampus | 440788 | -59212 | 940788 | Yes |
| ENSG00000120647.9 | CCDC77 | rs28850111 | 0.000015 | ##### | Brain - Cerebellar Hemisphere | 440788 | -59212 | 940788 | Yes |
| ENSG00000120647.9 | CCDC77 | rs2369280 | 0.000015 | ##### | Brain - Substantia nigra | 440788 | -59212 | 940788 | Yes |
| ENSG00000120647.9 | CCDC77 | rs11062934 | 0.000016 | ##### | Brain - Cerebellar Hemisphere | 440788 | -59212 | 940788 | Yes |
| ENSG00000120647.9 | CCDC77 | rs11062357 | 0.000016 | ##### | Brain - Hypothalamus | 440788 | -59212 | 940788 | Yes |
| ENSG00000120647.9 | CCDC77 | rs11062945 | 0.000016 | ##### | Brain - Substantia nigra | 440788 | -59212 | 940788 | Yes |
| ENSG00000120647.9 | CCDC77 | rs2369281 | 0.000016 | ##### | Brain - Putamen (basal gangli | 440788 | -59212 | 940788 | Yes |
| ENSG00000120647.9 | CCDC77 | rs28850111 | 0.000016 | ##### | Brain - Substantia nigra | 440788 | -59212 | 940788 | Yes |
| ENSG00000120647.9 | CCDC77 | rs2080402 | 0.000017 | ##### | Brain - Cortex | 440788 | -59212 | 940788 | Yes |
| ENSG00000120647.9 | CCDC77 | rs4980892 | 0.000017 | ##### | Brain - Substantia nigra | 440788 | -59212 | 940788 | Yes |
| ENSG00000120647.9 | CCDC77 | rs10848961 | 0.000017 | ##### | Brain - Substantia nigra | 440788 | -59212 | 940788 | Yes |
| ENSG00000120647.9 | CCDC77 | rs2080403 | 0.000017 | ##### | Brain - Cerebellum | 440788 | -59212 | 940788 | Yes |
| ENSG00000120647.9 | CCDC77 | rs2289957 | 0.000017 | ##### | Brain - Cerebellum | 440788 | -59212 | 940788 | Yes |
| ENSG00000120647.9 | CCDC77 | rs11062943 | 0.000018 | ##### | Brain - Cortex | 440788 | -59212 | 940788 | Yes |
| ENSG00000120647.9 | CCDC77 | rs4980819 | 0.000018 | ##### | Brain - Cortex | 440788 | -59212 | 940788 | Yes |
| ENSG00000120647.9 | CCDC77 | rs55657100 | 0.000018 | ##### | Brain - Anterior cingulate cort | 440788 | -59212 | 940788 | Yes |
| ENSG00000120647.9 | CCDC77 | rs2284332 | 0.000018 | ##### | Brain - Hippocampus | 440788 | -59212 | 940788 | Yes |
| ENSG00000120647.9 | CCDC77 | rs2229351 | 0.000018 | ##### | Brain - Hippocampus | 440788 | -59212 | 940788 | Yes |

| | | | | | | | | | |
|---|---|---|---|---|---|---|---|---|---|
| ENSG00000120647.9 | CCDC77 | rs6489477 | 0.000018 | ##### | Brain - Hippocampus | 440788 | -59212 | 940788 | Yes |
| ENSG00000120647.9 | CCDC77 | rs6489478 | 0.000018 | ##### | Brain - Hippocampus | 440788 | -59212 | 940788 | Yes |
| ENSG00000120647.9 | CCDC77 | rs487252 | 0.000018 | ##### | Brain - Cortex | 440788 | -59212 | 940788 | Yes |
| ENSG00000120647.9 | CCDC77 | rs10848650 | 0.000018 | ##### | Brain - Cortex | 440788 | -59212 | 940788 | Yes |
| ENSG00000120647.9 | CCDC77 | rs56042953 | 0.000019 | ##### | Brain - Nucleus accumbens (b | 440788 | -59212 | 940788 | Yes |
| ENSG00000120647.9 | CCDC77 | rs61907026 | 0.000019 | ##### | Brain - Anterior cingulate cort | 440788 | -59212 | 940788 | Yes |
| ENSG00000120647.9 | CCDC77 | rs79839498 | 0.000019 | ##### | Brain - Anterior cingulate cort | 440788 | -59212 | 940788 | Yes |
| ENSG00000120647.9 | CCDC77 | rs61907030 | 0.000019 | ##### | Brain - Anterior cingulate cort | 440788 | -59212 | 940788 | Yes |
| ENSG00000120647.9 | CCDC77 | rs61907031 | 0.000019 | ##### | Brain - Anterior cingulate cort | 440788 | -59212 | 940788 | Yes |
| ENSG00000120647.9 | CCDC77 | rs11615564 | 0.000019 | ##### | Brain - Anterior cingulate cort | 440788 | -59212 | 940788 | Yes |
| ENSG00000120647.9 | CCDC77 | rs11062813 | 0.000019 | ##### | Brain - Amygdala | 440788 | -59212 | 940788 | Yes |
| ENSG00000120647.9 | CCDC77 | rs3753156 | 0.000019 | ##### | Brain - Anterior cingulate cort | 440788 | -59212 | 940788 | Yes |
| ENSG00000120647.9 | CCDC77 | rs12581260 | 0.000019 | ##### | Brain - Anterior cingulate cort | 440788 | -59212 | 940788 | Yes |
| ENSG00000120647.9 | CCDC77 | rs16929783 | 0.000019 | ##### | Brain - Anterior cingulate cort | 440788 | -59212 | 940788 | Yes |
| ENSG00000120647.9 | CCDC77 | rs7307754 | 0.000019 | ##### | Brain - Cerebellar Hemisphere | 440788 | -59212 | 940788 | Yes |
| ENSG00000120647.9 | CCDC77 | rs1048466 | 0.000021 | ##### | Brain - Cortex | 440788 | -59212 | 940788 | Yes |
| ENSG00000120647.9 | CCDC77 | rs2253565 | 0.000021 | ##### | Brain - Hippocampus | 440788 | -59212 | 940788 | Yes |
| ENSG00000120647.9 | CCDC77 | rs12370433 | 0.000021 | ##### | Brain - Cerebellar Hemisphere | 440788 | -59212 | 940788 | Yes |
| ENSG00000120647.9 | CCDC77 | rs11063005 | 0.000021 | ##### | Brain - Cortex | 440788 | -59212 | 940788 | Yes |
| ENSG00000120647.9 | CCDC77 | rs11063010 | 0.000021 | ##### | Brain - Cortex | 440788 | -59212 | 940788 | Yes |
| ENSG00000120647.9 | CCDC77 | rs1860361 | 0.000021 | ##### | Brain - Hypothalamus | 440788 | -59212 | 940788 | Yes |
| ENSG00000120647.9 | CCDC77 | rs2286606 | 0.000021 | ##### | Brain - Cortex | 440788 | -59212 | 940788 | Yes |
| ENSG00000120647.9 | CCDC77 | rs576994 | 0.000021 | ##### | Brain - Hippocampus | 440788 | -59212 | 940788 | Yes |
| ENSG00000120647.9 | CCDC77 | rs797759 | 0.000021 | ##### | Brain - Hippocampus | 440788 | -59212 | 940788 | Yes |
| ENSG00000120647.9 | CCDC77 | rs479836 | 0.000021 | ##### | Brain - Hippocampus | 440788 | -59212 | 940788 | Yes |
| ENSG00000120647.9 | CCDC77 | rs4980810 | 0.000021 | ##### | Brain - Hippocampus | 440788 | -59212 | 940788 | Yes |
| ENSG00000120647.9 | CCDC77 | rs11062302 | 0.000021 | ##### | Brain - Hippocampus | 440788 | -59212 | 940788 | Yes |
| ENSG00000120647.9 | CCDC77 | rs12314497 | 0.000021 | ##### | Brain - Hippocampus | 440788 | -59212 | 940788 | Yes |
| ENSG00000120647.9 | CCDC77 | rs534925791 | 0.000021 | ##### | Brain - Hippocampus | 440788 | -59212 | 940788 | Yes |
| ENSG00000120647.9 | CCDC77 | rs144729439 | 0.000021 | ##### | Brain - Hippocampus | 440788 | -59212 | 940788 | Yes |
| ENSG00000120647.9 | CCDC77 | rs11062385 | 0.000021 | ##### | Brain - Hippocampus | 440788 | -59212 | 940788 | Yes |
| ENSG00000120647.9 | CCDC77 | rs11611810 | 0.000021 | ##### | Brain - Hippocampus | 440788 | -59212 | 940788 | Yes |

| | | | | | | | | | |
|---|---|---|---|---|---|---|---|---|---|
| ENSG00000120647.9 | CCDC77 | rs10848715 | 0.000021 | ##### | Brain - Hippocampus | 440788 | -59212 | 940788 | Yes |
| ENSG00000120647.9 | CCDC77 | rs10848716 | 0.000021 | ##### | Brain - Hippocampus | 440788 | -59212 | 940788 | Yes |
| ENSG00000120647.9 | CCDC77 | rs61907027 | 0.000021 | ##### | Brain - Hippocampus | 440788 | -59212 | 940788 | Yes |
| ENSG00000120647.9 | CCDC77 | rs7301422 | 0.000021 | ##### | Brain - Hippocampus | 440788 | -59212 | 940788 | Yes |
| ENSG00000120647.9 | CCDC77 | rs11062512 | 0.000021 | ##### | Brain - Hippocampus | 440788 | -59212 | 940788 | Yes |
| ENSG00000120647.9 | CCDC77 | rs3759371 | 0.000021 | ##### | Brain - Hippocampus | 440788 | -59212 | 940788 | Yes |
| ENSG00000120647.9 | CCDC77 | rs6489448 | 0.000021 | ##### | Brain - Hippocampus | 440788 | -59212 | 940788 | Yes |
| ENSG00000120647.9 | CCDC77 | rs7316660 | 0.000021 | ##### | Brain - Hippocampus | 440788 | -59212 | 940788 | Yes |
| ENSG00000120647.9 | CCDC77 | rs6489449 | 0.000021 | ##### | Brain - Hippocampus | 440788 | -59212 | 940788 | Yes |
| ENSG00000120647.9 | CCDC77 | rs10735021 | 0.000021 | ##### | Brain - Hippocampus | 440788 | -59212 | 940788 | Yes |
| ENSG00000120647.9 | CCDC77 | rs10848845 | 0.000021 | ##### | Brain - Hippocampus | 440788 | -59212 | 940788 | Yes |
| ENSG00000120647.9 | CCDC77 | rs10774137 | 0.000021 | ##### | Brain - Hippocampus | 440788 | -59212 | 940788 | Yes |
| ENSG00000120647.9 | CCDC77 | rs11835325 | 0.000021 | ##### | Brain - Hippocampus | 440788 | -59212 | 940788 | Yes |
| ENSG00000120647.9 | CCDC77 | rs10848849 | 0.000021 | ##### | Brain - Hippocampus | 440788 | -59212 | 940788 | Yes |
| ENSG00000120647.9 | CCDC77 | rs6489454 | 0.000021 | ##### | Brain - Hippocampus | 440788 | -59212 | 940788 | Yes |
| ENSG00000120647.9 | CCDC77 | rs7134608 | 0.000021 | ##### | Brain - Hippocampus | 440788 | -59212 | 940788 | Yes |
| ENSG00000120647.9 | CCDC77 | rs7138890 | 0.000021 | ##### | Brain - Hippocampus | 440788 | -59212 | 940788 | Yes |
| ENSG00000120647.9 | CCDC77 | rs10431355 | 0.000021 | ##### | Brain - Hippocampus | 440788 | -59212 | 940788 | Yes |
| ENSG00000120647.9 | CCDC77 | rs6489458 | 0.000021 | ##### | Brain - Hippocampus | 440788 | -59212 | 940788 | Yes |
| ENSG00000120647.9 | CCDC77 | rs2300134 | 0.000021 | ##### | Brain - Hippocampus | 440788 | -59212 | 940788 | Yes |
| ENSG00000120647.9 | CCDC77 | rs3782852 | 0.000021 | ##### | Brain - Hippocampus | 440788 | -59212 | 940788 | Yes |
| ENSG00000120647.9 | CCDC77 | rs201124968 | 0.000021 | ##### | Brain - Hippocampus | 440788 | -59212 | 940788 | Yes |
| ENSG00000120647.9 | CCDC77 | rs201946814 | 0.000021 | ##### | Brain - Hippocampus | 440788 | -59212 | 940788 | Yes |
| ENSG00000120647.9 | CCDC77 | rs6489463 | 0.000021 | ##### | Brain - Hippocampus | 440788 | -59212 | 940788 | Yes |
| ENSG00000120647.9 | CCDC77 | rs2079785 | 0.000021 | ##### | Brain - Hippocampus | 440788 | -59212 | 940788 | Yes |
| ENSG00000120647.9 | CCDC77 | rs7963937 | 0.000021 | ##### | Brain - Hippocampus | 440788 | -59212 | 940788 | Yes |
| ENSG00000120647.9 | CCDC77 | rs2369276 | 0.000021 | ##### | Brain - Hippocampus | 440788 | -59212 | 940788 | Yes |
| ENSG00000120647.9 | CCDC77 | rs7953137 | 0.000021 | ##### | Brain - Hippocampus | 440788 | -59212 | 940788 | Yes |
| ENSG00000120647.9 | CCDC77 | rs6489471 | 0.000021 | ##### | Brain - Hippocampus | 440788 | -59212 | 940788 | Yes |
| ENSG00000120647.9 | CCDC77 | rs7312065 | 0.000021 | ##### | Brain - Hippocampus | 440788 | -59212 | 940788 | Yes |
| ENSG00000120647.9 | CCDC77 | rs34380840 | 0.000021 | ##### | Brain - Hippocampus | 440788 | -59212 | 940788 | Yes |
| ENSG00000120647.9 | CCDC77 | rs10744561 | 0.000021 | ##### | Brain - Cortex | 440788 | -59212 | 940788 | Yes |

| | | | | | | | | | |
|---|---|---|---|---|---|---|---|---|---|
| ENSG00000120647.9 | CCDC77 | rs7311740 | 0.000021 | ##### | Brain - Putamen (basal gangli | 440788 | -59212 | 940788 | Yes |
| ENSG00000120647.9 | CCDC77 | rs7295011 | 0.000022 | ##### | Brain - Putamen (basal gangli | 440788 | -59212 | 940788 | Yes |
| ENSG00000120647.9 | CCDC77 | rs3213751 | 0.000022 | ##### | Brain - Hippocampus | 440788 | -59212 | 940788 | Yes |
| ENSG00000120647.9 | CCDC77 | rs12370433 | 0.000022 | ##### | Brain - Cortex | 440788 | -59212 | 940788 | Yes |
| ENSG00000120647.9 | CCDC77 | rs10848987 | 0.000022 | ##### | Brain - Putamen (basal gangli | 440788 | -59212 | 940788 | Yes |
| ENSG00000120647.9 | CCDC77 | rs16930719 | 0.000022 | ##### | Brain - Cortex | 440788 | -59212 | 940788 | Yes |
| ENSG00000120647.9 | CCDC77 | rs12298843 | 0.000023 | ##### | Brain - Hippocampus | 440788 | -59212 | 940788 | Yes |
| ENSG00000120647.9 | CCDC77 | rs11062934 | 0.000024 | ##### | Brain - Cortex | 440788 | -59212 | 940788 | Yes |
| ENSG00000120647.9 | CCDC77 | rs3815558 | 0.000024 | ##### | Brain - Hippocampus | 440788 | -59212 | 940788 | Yes |
| ENSG00000120647.9 | CCDC77 | rs78998485 | 0.000024 | ##### | Brain - Hippocampus | 440788 | -59212 | 940788 | Yes |
| ENSG00000120647.9 | CCDC77 | rs4980884 | 0.000024 | ##### | Brain - Hippocampus | 440788 | -59212 | 940788 | Yes |
| ENSG00000120647.9 | CCDC77 | rs7135768 | 0.000024 | ##### | Brain - Hippocampus | 440788 | -59212 | 940788 | Yes |
| ENSG00000120647.9 | CCDC77 | rs6489450 | 0.000024 | ##### | Brain - Hippocampus | 440788 | -59212 | 940788 | Yes |
| ENSG00000120647.9 | CCDC77 | rs4980906 | 0.000024 | ##### | Brain - Frontal Cortex (BA9) | 440788 | -59212 | 940788 | Yes |
| ENSG00000120647.9 | CCDC77 | rs16930719 | 0.000025 | ##### | Brain - Nucleus accumbens (b | 440788 | -59212 | 940788 | Yes |
| ENSG00000120647.9 | CCDC77 | rs580420 | 0.000026 | ##### | Brain - Hypothalamus | 440788 | -59212 | 940788 | Yes |
| ENSG00000120647.9 | CCDC77 | rs505211 | 0.000026 | ##### | Brain - Hypothalamus | 440788 | -59212 | 940788 | Yes |
| ENSG00000120647.9 | CCDC77 | rs532463 | 0.000026 | ##### | Brain - Hypothalamus | 440788 | -59212 | 940788 | Yes |
| ENSG00000120647.9 | CCDC77 | rs505574 | 0.000026 | ##### | Brain - Hypothalamus | 440788 | -59212 | 940788 | Yes |
| ENSG00000120647.9 | CCDC77 | rs1088324 | 0.000026 | ##### | Brain - Hypothalamus | 440788 | -59212 | 940788 | Yes |
| ENSG00000120647.9 | CCDC77 | rs12229673 | 0.000026 | ##### | Brain - Cortex | 440788 | -59212 | 940788 | Yes |
| ENSG00000120647.9 | CCDC77 | rs11062986 | 0.000026 | ##### | Brain - Frontal Cortex (BA9) | 440788 | -59212 | 940788 | Yes |
| ENSG00000120647.9 | CCDC77 | rs75611255 | 0.000026 | ##### | Brain - Hypothalamus | 440788 | -59212 | 940788 | Yes |
| ENSG00000120647.9 | CCDC77 | rs10848647 | 0.000026 | ##### | Brain - Cortex | 440788 | -59212 | 940788 | Yes |
| ENSG00000120647.9 | CCDC77 | rs11062945 | 0.000026 | ##### | Brain - Spinal cord (cervical c- | 440788 | -59212 | 940788 | Yes |
| ENSG00000120647.9 | CCDC77 | rs3782860 | 0.000026 | ##### | Brain - Cortex | 440788 | -59212 | 940788 | Yes |
| ENSG00000120647.9 | CCDC77 | rs3213750 | 0.000027 | ##### | Brain - Anterior cingulate cort | 440788 | -59212 | 940788 | Yes |
| ENSG00000120647.9 | CCDC77 | rs4980881 | 0.000027 | ##### | Brain - Anterior cingulate cort | 440788 | -59212 | 940788 | Yes |
| ENSG00000120647.9 | CCDC77 | rs17800720 | 0.000028 | ##### | Brain - Cortex | 440788 | -59212 | 940788 | Yes |
| ENSG00000120647.9 | CCDC77 | rs3837515 | 0.000028 | ##### | Brain - Cortex | 440788 | -59212 | 940788 | Yes |
| ENSG00000120647.9 | CCDC77 | rs495316 | 0.000028 | ##### | Brain - Hippocampus | 440788 | -59212 | 940788 | Yes |
| ENSG00000120647.9 | CCDC77 | rs4980908 | 0.000028 | ##### | Brain - Cerebellar Hemisphere | 440788 | -59212 | 940788 | Yes |

| | | | | | | | | | |
|---|---|---|---|---|---|---|---|---|---|
| ENSG00000120647.9 | CCDC77 | rs2369281 | 0.000029 | ##### | Brain - Cerebellar Hemisphere | 440788 | -59212 | 940788 | Yes |
| ENSG00000120647.9 | CCDC77 | rs141522933 | 0.000029 | ##### | Brain - Caudate (basal ganglia | 440788 | -59212 | 940788 | Yes |
| ENSG00000120647.9 | CCDC77 | rs476646 | 0.000029 | ##### | Brain - Caudate (basal ganglia | 440788 | -59212 | 940788 | Yes |
| ENSG00000120647.9 | CCDC77 | rs497740 | 0.000029 | ##### | Brain - Cerebellar Hemisphere | 440788 | -59212 | 940788 | Yes |
| ENSG00000120647.9 | CCDC77 | rs12231564 | 0.000031 | ##### | Brain - Cortex | 440788 | -59212 | 940788 | Yes |
| ENSG00000120647.9 | CCDC77 | rs1548904 | 0.000031 | ##### | Brain - Cortex | 440788 | -59212 | 940788 | Yes |
| ENSG00000120647.9 | CCDC77 | rs1972582 | 0.000031 | ##### | Brain - Cortex | 440788 | -59212 | 940788 | Yes |
| ENSG00000120647.9 | CCDC77 | rs10774038 | 0.000031 | ##### | Brain - Cortex | 440788 | -59212 | 940788 | Yes |
| ENSG00000120647.9 | CCDC77 | rs476646 | 0.000031 | ##### | Brain - Cerebellar Hemisphere | 440788 | -59212 | 940788 | Yes |
| ENSG00000120647.9 | CCDC77 | rs3753158 | 0.000031 | ##### | Brain - Anterior cingulate cort | 440788 | -59212 | 940788 | Yes |
| ENSG00000120647.9 | CCDC77 | rs10161265 | 0.000033 | ##### | Brain - Nucleus accumbens (b | 440788 | -59212 | 940788 | Yes |
| ENSG00000120647.9 | CCDC77 | rs4980897 | 0.000033 | ##### | Brain - Nucleus accumbens (b | 440788 | -59212 | 940788 | Yes |
| ENSG00000120647.9 | CCDC77 | rs12833509 | 0.000033 | ##### | Brain - Frontal Cortex (BA9) | 440788 | -59212 | 940788 | Yes |
| ENSG00000120647.9 | CCDC77 | rs11063005 | 0.000033 | ##### | Brain - Nucleus accumbens (b | 440788 | -59212 | 940788 | Yes |
| ENSG00000120647.9 | CCDC77 | rs11063010 | 0.000033 | ##### | Brain - Nucleus accumbens (b | 440788 | -59212 | 940788 | Yes |
| ENSG00000120647.9 | CCDC77 | rs11062167 | 0.000034 | ##### | Brain - Cerebellum | 440788 | -59212 | 940788 | Yes |
| ENSG00000120647.9 | CCDC77 | rs55990424 | 0.000034 | ##### | Brain - Cerebellum | 440788 | -59212 | 940788 | Yes |
| ENSG00000120647.9 | CCDC77 | rs10848987 | 0.000034 | ##### | Brain - Cortex | 440788 | -59212 | 940788 | Yes |
| ENSG00000120647.9 | CCDC77 | rs12301816 | 0.000035 | ##### | Brain - Anterior cingulate cort | 440788 | -59212 | 940788 | Yes |
| ENSG00000120647.9 | CCDC77 | rs797765 | 0.000036 | ##### | Brain - Anterior cingulate cort | 440788 | -59212 | 940788 | Yes |
| ENSG00000120647.9 | CCDC77 | rs7136957 | 0.000036 | ##### | Brain - Hippocampus | 440788 | -59212 | 940788 | Yes |
| ENSG00000120647.9 | CCDC77 | rs1207324 | 0.000036 | ##### | Brain - Hippocampus | 440788 | -59212 | 940788 | Yes |
| ENSG00000120647.9 | CCDC77 | rs11615492 | 0.000037 | ##### | Brain - Caudate (basal ganglia | 440788 | -59212 | 940788 | Yes |
| ENSG00000120647.9 | CCDC77 | rs10848987 | 0.000037 | ##### | Brain - Cerebellum | 440788 | -59212 | 940788 | Yes |
| ENSG00000120647.9 | CCDC77 | rs56042953 | 0.000038 | ##### | Brain - Anterior cingulate cort | 440788 | -59212 | 940788 | Yes |
| ENSG00000120647.9 | CCDC77 | rs10774192 | 0.000038 | ##### | Brain - Cortex | 440788 | -59212 | 940788 | Yes |
| ENSG00000120647.9 | CCDC77 | rs7969761 | 0.000038 | ##### | Brain - Cortex | 440788 | -59212 | 940788 | Yes |
| ENSG00000120647.9 | CCDC77 | rs530500 | 0.000039 | ##### | Brain - Hypothalamus | 440788 | -59212 | 940788 | Yes |
| ENSG00000120647.9 | CCDC77 | rs4980896 | 0.00004 | ##### | Brain - Cerebellum | 440788 | -59212 | 940788 | Yes |
| ENSG00000120647.9 | CCDC77 | rs11062102 | 0.00004 | ##### | Brain - Frontal Cortex (BA9) | 440788 | -59212 | 940788 | Yes |
| ENSG00000120647.9 | CCDC77 | rs10774020 | 0.00004 | ##### | Brain - Frontal Cortex (BA9) | 440788 | -59212 | 940788 | Yes |
| ENSG00000120647.9 | CCDC77 | | 0.000041 | ##### | Brain - Hypothalamus | 440788 | -59212 | 940788 | Yes |

| | | | | | | | | | |
|---|---|---|---|---|---|---|---|---|---|
| ENSG00000120647.9 | CCDC77 | rs56042953 | 0.000042 | ##### | Brain - Cortex | 440788 | -59212 | 940788 | Yes |
| ENSG00000120647.9 | CCDC77 | rs10848644 | 0.000042 | ##### | Brain - Cerebellum | 440788 | -59212 | 940788 | Yes |
| ENSG00000120647.9 | CCDC77 | rs12229673 | 0.000043 | ##### | Brain - Frontal Cortex (BA9) | 440788 | -59212 | 940788 | Yes |
| ENSG00000120647.9 | CCDC77 | rs12231564 | 0.000043 | ##### | Brain - Frontal Cortex (BA9) | 440788 | -59212 | 940788 | Yes |
| ENSG00000120647.9 | CCDC77 | rs10848639 | 0.000044 | ##### | Brain - Cerebellar Hemisphere | 440788 | -59212 | 940788 | Yes |
| ENSG00000120647.9 | CCDC77 | rs35222334 | 0.000044 | ##### | Brain - Caudate (basal ganglia | 440788 | -59212 | 940788 | Yes |
| ENSG00000120647.9 | CCDC77 | rs11062071 | 0.000044 | ##### | Brain - Caudate (basal ganglia | 440788 | -59212 | 940788 | Yes |
| ENSG00000120647.9 | CCDC77 | rs11062085 | 0.000044 | ##### | Brain - Caudate (basal ganglia | 440788 | -59212 | 940788 | Yes |
| ENSG00000120647.9 | CCDC77 | rs12580917 | 0.000044 | ##### | Brain - Caudate (basal ganglia | 440788 | -59212 | 940788 | Yes |
| ENSG00000120647.9 | CCDC77 | rs11062163 | 0.000046 | ##### | Brain - Caudate (basal ganglia | 440788 | -59212 | 940788 | Yes |
| ENSG00000120647.9 | CCDC77 | rs487252 | 0.000046 | ##### | Brain - Nucleus accumbens (b | 440788 | -59212 | 940788 | Yes |
| ENSG00000120647.9 | CCDC77 | rs12830750 | 0.000047 | ##### | Brain - Caudate (basal ganglia | 440788 | -59212 | 940788 | Yes |
| ENSG00000120647.9 | CCDC77 | rs11062072 | 0.000047 | ##### | Brain - Caudate (basal ganglia | 440788 | -59212 | 940788 | Yes |
| ENSG00000120647.9 | CCDC77 | rs7969059 | 0.000047 | ##### | Brain - Caudate (basal ganglia | 440788 | -59212 | 940788 | Yes |
| ENSG00000120647.9 | CCDC77 | rs199903042 | 0.000047 | ##### | Brain - Caudate (basal ganglia | 440788 | -59212 | 940788 | Yes |
| ENSG00000120647.9 | CCDC77 | rs12579445 | 0.000047 | ##### | Brain - Caudate (basal ganglia | 440788 | -59212 | 940788 | Yes |
| ENSG00000120647.9 | CCDC77 | rs73029312 | 0.000048 | ##### | Brain - Cerebellum | 440788 | -59212 | 940788 | No |
| ENSG00000120647.9 | CCDC77 | rs149376548 | 0.000048 | ##### | Brain - Cerebellum | 440788 | -59212 | 940788 | No |
| ENSG00000120647.9 | CCDC77 | rs1048466 | 0.000049 | ##### | Brain - Nucleus accumbens (b | 440788 | -59212 | 940788 | Yes |
| ENSG00000120647.9 | CCDC77 | rs67523949 | 0.000049 | ##### | Brain - Cerebellum | 440788 | -59212 | 940788 | Yes |
| ENSG00000120647.9 | CCDC77 | rs10848987 | 0.00005 | ##### | Brain - Nucleus accumbens (b | 440788 | -59212 | 940788 | Yes |
| ENSG00000120647.9 | CCDC77 | rs12372234 | 0.00005 | ##### | Brain - Putamen (basal gangli | 440788 | -59212 | 940788 | Yes |
| ENSG00000120647.9 | CCDC77 | rs11613331 | 0.000051 | ##### | Brain - Cortex | 440788 | -59212 | 940788 | Yes |
| ENSG00000120647.9 | CCDC77 | rs496206 | 0.000051 | ##### | Brain - Caudate (basal ganglia | 440788 | -59212 | 940788 | Yes |
| ENSG00000120647.9 | CCDC77 | rs11610539 | 0.000052 | ##### | Brain - Caudate (basal ganglia | 440788 | -59212 | 940788 | Yes |
| ENSG00000120647.9 | CCDC77 | rs2041298 | 0.000052 | ##### | Brain - Putamen (basal gangli | 440788 | -59212 | 940788 | Yes |
| ENSG00000120647.9 | CCDC77 | rs11062945 | 0.000054 | ##### | Brain - Putamen (basal gangli | 440788 | -59212 | 940788 | Yes |
| ENSG00000120647.9 | CCDC77 | rs525631 | 0.00006 | ##### | Brain - Caudate (basal ganglia | 440788 | -59212 | 940788 | Yes |
| ENSG00000120647.9 | CCDC77 | rs55657100 | 0.000063 | ##### | Brain - Nucleus accumbens (b | 440788 | -59212 | 940788 | Yes |
| ENSG00000120647.9 | CCDC77 | rs7136116 | 0.000063 | ##### | Brain - Cortex | 440788 | -59212 | 940788 | Yes |
| ENSG00000120647.9 | CCDC77 | rs11610457 | 0.000063 | ##### | Brain - Cortex | 440788 | -59212 | 940788 | Yes |
| ENSG00000120647.9 | CCDC77 | rs11062929 | 0.000064 | ##### | Brain - Cortex | 440788 | -59212 | 940788 | Yes |

| | | | | | | | | | |
|---|---|---|---|---|---|---|---|---|---|
| ENSG00000120647.9 | CCDC77 | rs11062986 | 0.000066 | ##### | Brain - Cortex | 440788 | -59212 | 940788 | Yes |
| ENSG00000120647.9 | CCDC77 | rs12826998 | 0.000068 | ##### | Brain - Cerebellar Hemisphere | 440788 | -59212 | 940788 | Yes |
| ENSG00000120647.9 | CCDC77 | rs10082788 | 0.000074 | ##### | Brain - Cerebellum | 440788 | -59212 | 940788 | Yes |
| ENSG00000120647.9 | CCDC77 | rs11062930 | 0.000075 | ##### | Brain - Cortex | 440788 | -59212 | 940788 | Yes |
| ENSG00000120647.9 | CCDC77 | rs72647390 | 0.000078 | ##### | Brain - Cortex | 440788 | -59212 | 940788 | Yes |
| ENSG00000120647.9 | CCDC77 | rs117399790 | 0.000078 | ##### | Brain - Cortex | 440788 | -59212 | 940788 | Yes |
| ENSG00000120647.9 | CCDC77 | rs72650727 | 0.000078 | ##### | Brain - Cortex | 440788 | -59212 | 940788 | Yes |
| ENSG00000117640.17 | MTFR1L | rs1127818 | 1.30E-11 | ##### | Brain - Cerebellum | 25826774 | 25326774 | 26326774 | Yes |
| ENSG00000117640.17 | MTFR1L | rs213622 | 2.30E-11 | ##### | Brain - Cerebellum | 25826774 | 25326774 | 26326774 | Yes |
| ENSG00000117640.17 | MTFR1L | rs11247764 | 4.40E-11 | ##### | Brain - Cerebellum | 25826774 | 25326774 | 26326774 | Yes |
| ENSG00000117640.17 | MTFR1L | rs213623 | 5.30E-11 | ##### | Brain - Cerebellum | 25826774 | 25326774 | 26326774 | Yes |
| ENSG00000117640.17 | MTFR1L | rs2782824 | 9.20E-11 | ##### | Brain - Cerebellum | 25826774 | 25326774 | 26326774 | Yes |
| ENSG00000117640.17 | MTFR1L | rs213638 | 9.80E-11 | ##### | Brain - Cerebellum | 25826774 | 25326774 | 26326774 | Yes |
| ENSG00000117640.17 | MTFR1L | rs4659394 | 1.00E-10 | ##### | Brain - Cerebellum | 25826774 | 25326774 | 26326774 | Yes |
| ENSG00000117640.17 | MTFR1L | rs3767876 | 1.10E-10 | ##### | Brain - Cerebellum | 25826774 | 25326774 | 26326774 | Yes |
| ENSG00000117640.17 | MTFR1L | rs1835308 | 1.20E-10 | ##### | Brain - Cerebellum | 25826774 | 25326774 | 26326774 | Yes |
| ENSG00000117640.17 | MTFR1L | rs159523 | 2.40E-10 | ##### | Brain - Cerebellum | 25826774 | 25326774 | 26326774 | Yes |
| ENSG00000117640.17 | MTFR1L | rs807070 | 3.10E-10 | ##### | Brain - Cerebellum | 25826774 | 25326774 | 26326774 | Yes |
| ENSG00000117640.17 | MTFR1L | rs159528 | 3.10E-10 | ##### | Brain - Cerebellum | 25826774 | 25326774 | 26326774 | Yes |
| ENSG00000117640.17 | MTFR1L | rs807053 | 3.10E-10 | ##### | Brain - Cerebellum | 25826774 | 25326774 | 26326774 | Yes |
| ENSG00000117640.17 | MTFR1L | rs159525 | 3.10E-10 | ##### | Brain - Cerebellum | 25826774 | 25326774 | 26326774 | Yes |
| ENSG00000117640.17 | MTFR1L | rs190043 | 4.70E-10 | ##### | Brain - Cerebellum | 25826774 | 25326774 | 26326774 | Yes |
| ENSG00000117640.17 | MTFR1L | rs807067 | 8.70E-10 | ##### | Brain - Cerebellum | 25826774 | 25326774 | 26326774 | Yes |
| ENSG00000117640.17 | MTFR1L | rs213637 | 9.00E-10 | ##### | Brain - Cerebellum | 25826774 | 25326774 | 26326774 | Yes |
| ENSG00000117640.17 | MTFR1L | rs213624 | 1.10E-09 | ##### | Brain - Cerebellum | 25826774 | 25326774 | 26326774 | Yes |
| ENSG00000117640.17 | MTFR1L | rs807069 | 1.90E-09 | ##### | Brain - Cerebellum | 25826774 | 25326774 | 26326774 | Yes |
| ENSG00000117640.17 | MTFR1L | rs4659393 | 1.90E-09 | ##### | Brain - Cerebellum | 25826774 | 25326774 | 26326774 | Yes |
| ENSG00000117640.17 | MTFR1L | rs11247824 | 1.90E-09 | ##### | Brain - Cerebellum | 25826774 | 25326774 | 26326774 | Yes |
| ENSG00000117640.17 | MTFR1L | rs159530 | 1.90E-09 | ##### | Brain - Cerebellum | 25826774 | 25326774 | 26326774 | Yes |
| ENSG00000117640.17 | MTFR1L | rs159527 | 1.90E-09 | ##### | Brain - Cerebellum | 25826774 | 25326774 | 26326774 | Yes |
| ENSG00000117640.17 | MTFR1L | rs2095426 | 1.90E-09 | ##### | Brain - Cerebellum | 25826774 | 25326774 | 26326774 | Yes |
| ENSG00000117640.17 | MTFR1L | rs3129533 | 1.90E-09 | ##### | Brain - Cerebellum | 25826774 | 25326774 | 26326774 | Yes |

| | | | | | | | | | |
|---|---|---|---|---|---|---|---|---|---|
| ENSG00000117640.17 | MTFR1L | rs1537682 | 1.90E-09 | ##### | Brain - Cerebellum | 25826774 | 25326774 | 26326774 | Yes |
| ENSG00000117640.17 | MTFR1L | rs159522 | 1.90E-09 | ##### | Brain - Cerebellum | 25826774 | 25326774 | 26326774 | Yes |
| ENSG00000117640.17 | MTFR1L | rs159521 | 1.90E-09 | ##### | Brain - Cerebellum | 25826774 | 25326774 | 26326774 | Yes |
| ENSG00000117640.17 | MTFR1L | rs6872 | 2.10E-09 | ##### | Brain - Cortex | 25826774 | 25326774 | 26326774 | Yes |
| ENSG00000117640.17 | MTFR1L | rs10449775 | 2.50E-09 | ##### | Brain - Cerebellum | 25826774 | 25326774 | 26326774 | Yes |
| ENSG00000117640.17 | MTFR1L | rs213621 | 3.20E-09 | ##### | Brain - Cerebellum | 25826774 | 25326774 | 26326774 | Yes |
| ENSG00000117640.17 | MTFR1L | rs2992073 | 9.30E-09 | ##### | Brain - Cortex | 25826774 | 25326774 | 26326774 | Yes |
| ENSG00000117640.17 | MTFR1L | rs1127818 | 1.10E-08 | ##### | Brain - Cerebellar Hemisphere | 25826774 | 25326774 | 26326774 | Yes |
| ENSG00000117640.17 | MTFR1L | rs11247764 | 1.30E-08 | ##### | Brain - Cerebellar Hemisphere | 25826774 | 25326774 | 26326774 | Yes |
| ENSG00000117640.17 | MTFR1L | rs213623 | 2.10E-08 | ##### | Brain - Cerebellar Hemisphere | 25826774 | 25326774 | 26326774 | Yes |
| ENSG00000117640.17 | MTFR1L | rs4659394 | 2.40E-08 | ##### | Brain - Cerebellar Hemisphere | 25826774 | 25326774 | 26326774 | Yes |
| ENSG00000117640.17 | MTFR1L | rs213620 | 2.50E-08 | ##### | Brain - Cerebellum | 25826774 | 25326774 | 26326774 | Yes |
| ENSG00000117640.17 | MTFR1L | rs3767876 | 2.50E-08 | ##### | Brain - Cerebellar Hemisphere | 25826774 | 25326774 | 26326774 | Yes |
| ENSG00000117640.17 | MTFR1L | rs1835308 | 2.50E-08 | ##### | Brain - Cerebellar Hemisphere | 25826774 | 25326774 | 26326774 | Yes |
| ENSG00000117640.17 | MTFR1L | rs2782824 | 3.20E-08 | ##### | Brain - Cerebellar Hemisphere | 25826774 | 25326774 | 26326774 | Yes |
| ENSG00000117640.17 | MTFR1L | rs159526 | 3.20E-08 | ##### | Brain - Cortex | 25826774 | 25326774 | 26326774 | Yes |
| ENSG00000117640.17 | MTFR1L | rs807070 | 3.30E-08 | ##### | Brain - Cerebellar Hemisphere | 25826774 | 25326774 | 26326774 | Yes |
| ENSG00000117640.17 | MTFR1L | rs159528 | 3.30E-08 | ##### | Brain - Cerebellar Hemisphere | 25826774 | 25326774 | 26326774 | Yes |
| ENSG00000117640.17 | MTFR1L | rs807053 | 3.30E-08 | ##### | Brain - Cerebellar Hemisphere | 25826774 | 25326774 | 26326774 | Yes |
| ENSG00000117640.17 | MTFR1L | rs159525 | 3.30E-08 | ##### | Brain - Cerebellar Hemisphere | 25826774 | 25326774 | 26326774 | Yes |
| ENSG00000117640.17 | MTFR1L | rs159523 | 3.30E-08 | ##### | Brain - Cerebellar Hemisphere | 25826774 | 25326774 | 26326774 | Yes |
| ENSG00000117640.17 | MTFR1L | rs159532 | 3.50E-08 | ##### | Brain - Cerebellum | 25826774 | 25326774 | 26326774 | Yes |
| ENSG00000117640.17 | MTFR1L | rs807057 | 3.50E-08 | ##### | Brain - Cerebellum | 25826774 | 25326774 | 26326774 | Yes |
| ENSG00000117640.17 | MTFR1L | rs807056 | 3.50E-08 | ##### | Brain - Cerebellum | 25826774 | 25326774 | 26326774 | Yes |
| ENSG00000117640.17 | MTFR1L | rs213638 | 4.70E-08 | ##### | Brain - Cerebellar Hemisphere | 25826774 | 25326774 | 26326774 | Yes |
| ENSG00000117640.17 | MTFR1L | rs3738450 | 5.00E-08 | ##### | Brain - Cortex | 25826774 | 25326774 | 26326774 | Yes |
| ENSG00000117640.17 | MTFR1L | rs213622 | 5.10E-08 | ##### | Brain - Cerebellar Hemisphere | 25826774 | 25326774 | 26326774 | Yes |
| ENSG00000117640.17 | MTFR1L | rs298440 | 7.20E-08 | ##### | Brain - Cortex | 25826774 | 25326774 | 26326774 | Yes |
| ENSG00000117640.17 | MTFR1L | rs159531 | 7.30E-08 | ##### | Brain - Cortex | 25826774 | 25326774 | 26326774 | Yes |
| ENSG00000117640.17 | MTFR1L | rs2294226 | 7.60E-08 | ##### | Brain - Cortex | 25826774 | 25326774 | 26326774 | Yes |
| ENSG00000117640.17 | MTFR1L | rs807062 | 1.20E-07 | ##### | Brain - Cortex | 25826774 | 25326774 | 26326774 | Yes |
| ENSG00000117640.17 | MTFR1L | rs1961658 | 1.20E-07 | ##### | Brain - Cortex | 25826774 | 25326774 | 26326774 | Yes |

| | | | | | | | | | |
|---|---|---|---|---|---|---|---|---|---|
| ENSG00000117640.17 | MTFR1L | rs3120160 | 1.60E-07 | ##### | Brain - Cerebellum | 25826774 | 25326774 | 26326774 | Yes |
| ENSG00000117640.17 | MTFR1L | rs5773140 | 1.60E-07 | ##### | Brain - Cortex | 25826774 | 25326774 | 26326774 | Yes |
| ENSG00000117640.17 | MTFR1L | rs1257163 | 1.60E-07 | ##### | Brain - Cortex | 25826774 | 25326774 | 26326774 | Yes |
| ENSG00000117640.17 | MTFR1L | rs11376635 | 1.60E-07 | ##### | Brain - Cerebellum | 25826774 | 25326774 | 26326774 | Yes |
| ENSG00000117640.17 | MTFR1L | rs182455 | 1.60E-07 | ##### | Brain - Cerebellum | 25826774 | 25326774 | 26326774 | Yes |
| ENSG00000117640.17 | MTFR1L | rs807055 | 1.60E-07 | ##### | Brain - Cerebellum | 25826774 | 25326774 | 26326774 | Yes |
| ENSG00000117640.17 | MTFR1L | rs2782823 | 2.00E-07 | ##### | Brain - Cortex | 25826774 | 25326774 | 26326774 | Yes |
| ENSG00000117640.17 | MTFR1L | rs159533 | 2.10E-07 | ##### | Brain - Cerebellum | 25826774 | 25326774 | 26326774 | Yes |
| ENSG00000117640.17 | MTFR1L | rs3120159 | 2.80E-07 | ##### | Brain - Cerebellum | 25826774 | 25326774 | 26326774 | Yes |
| ENSG00000117640.17 | MTFR1L | rs34239512 | 2.90E-07 | ##### | Brain - Cortex | 25826774 | 25326774 | 26326774 | Yes |
| ENSG00000117640.17 | MTFR1L | rs17163302 | 2.90E-07 | ##### | Brain - Cortex | 25826774 | 25326774 | 26326774 | Yes |
| ENSG00000117640.17 | MTFR1L | rs213629 | 2.90E-07 | ##### | Brain - Cortex | 25826774 | 25326774 | 26326774 | Yes |
| ENSG00000117640.17 | MTFR1L | rs2565032 | 2.90E-07 | ##### | Brain - Cortex | 25826774 | 25326774 | 26326774 | Yes |
| ENSG00000117640.17 | MTFR1L | rs213648 | 2.90E-07 | ##### | Brain - Cortex | 25826774 | 25326774 | 26326774 | Yes |
| ENSG00000117640.17 | MTFR1L | rs7549097 | 3.50E-07 | ##### | Brain - Cerebellum | 25826774 | 25326774 | 26326774 | Yes |
| ENSG00000117640.17 | MTFR1L | rs6872 | 3.50E-07 | ##### | Brain - Caudate (basal ganglia | 25826774 | 25326774 | 26326774 | Yes |
| ENSG00000117640.17 | MTFR1L | rs807061 | 3.60E-07 | ##### | Brain - Cortex | 25826774 | 25326774 | 26326774 | Yes |
| ENSG00000117640.17 | MTFR1L | rs807060 | 4.10E-07 | ##### | Brain - Cerebellum | 25826774 | 25326774 | 26326774 | Yes |
| ENSG00000117640.17 | MTFR1L | rs159519 | 4.30E-07 | ##### | Brain - Cerebellum | 25826774 | 25326774 | 26326774 | Yes |
| ENSG00000117640.17 | MTFR1L | rs213641 | 4.40E-07 | ##### | Brain - Cerebellum | 25826774 | 25326774 | 26326774 | Yes |
| ENSG00000117640.17 | MTFR1L | rs2280997 | 4.50E-07 | ##### | Brain - Cortex | 25826774 | 25326774 | 26326774 | Yes |
| ENSG00000117640.17 | MTFR1L | rs7513196 | 4.80E-07 | ##### | Brain - Cerebellum | 25826774 | 25326774 | 26326774 | Yes |
| ENSG00000117640.17 | MTFR1L | rs5773140 | 4.90E-07 | ##### | Brain - Cerebellar Hemisphere | 25826774 | 25326774 | 26326774 | Yes |
| ENSG00000117640.17 | MTFR1L | rs2992073 | 4.90E-07 | ##### | Brain - Cerebellar Hemisphere | 25826774 | 25326774 | 26326774 | Yes |
| ENSG00000117640.17 | MTFR1L | rs1257163 | 4.90E-07 | ##### | Brain - Cerebellar Hemisphere | 25826774 | 25326774 | 26326774 | Yes |
| ENSG00000117640.17 | MTFR1L | rs4659382 | 6.10E-07 | ##### | Brain - Cortex | 25826774 | 25326774 | 26326774 | Yes |
| ENSG00000117640.17 | MTFR1L | rs807057 | 6.60E-07 | ##### | Brain - Cerebellar Hemisphere | 25826774 | 25326774 | 26326774 | Yes |
| ENSG00000117640.17 | MTFR1L | rs807056 | 6.60E-07 | ##### | Brain - Cerebellar Hemisphere | 25826774 | 25326774 | 26326774 | Yes |
| ENSG00000117640.17 | MTFR1L | rs159532 | 6.60E-07 | ##### | Brain - Cerebellar Hemisphere | 25826774 | 25326774 | 26326774 | Yes |
| ENSG00000117640.17 | MTFR1L | rs718391 | 6.80E-07 | ##### | Brain - Cerebellum | 25826774 | 25326774 | 26326774 | Yes |
| ENSG00000117640.17 | MTFR1L | rs6659086 | 7.40E-07 | ##### | Brain - Cerebellum | 25826774 | 25326774 | 26326774 | Yes |
| ENSG00000117640.17 | MTFR1L | rs10719116 | 7.50E-07 | ##### | Brain - Cerebellum | 25826774 | 25326774 | 26326774 | Yes |

| | | | | | | | | | |
|---|---|---|---|---|---|---|---|---|---|
| ENSG00000117640.17 | MTFR1L | rs116609713 | 7.50E-07 | ##### | Brain - Cerebellum | 25826774 | 25326774 | 26326774 | Yes |
| ENSG00000117640.17 | MTFR1L | rs2294226 | 9.60E-07 | ##### | Brain - Caudate (basal ganglia | 25826774 | 25326774 | 26326774 | Yes |
| ENSG00000117640.17 | MTFR1L | rs7524771 | 0.000001 | ##### | Brain - Cerebellum | 25826774 | 25326774 | 26326774 | Yes |
| ENSG00000117640.17 | MTFR1L | rs11247710 | 0.0000012 | ##### | Brain - Cerebellum | 25826774 | 25326774 | 26326774 | Yes |
| ENSG00000117640.17 | MTFR1L | rs11589618 | 0.0000013 | ##### | Brain - Cerebellum | 25826774 | 25326774 | 26326774 | Yes |
| ENSG00000117640.17 | MTFR1L | rs7518451 | 0.0000013 | ##### | Brain - Cerebellum | 25826774 | 25326774 | 26326774 | Yes |
| ENSG00000117640.17 | MTFR1L | rs36024836 | 0.0000013 | ##### | Brain - Cerebellum | 25826774 | 25326774 | 26326774 | Yes |
| ENSG00000117640.17 | MTFR1L | rs7524154 | 0.0000013 | ##### | Brain - Cerebellum | 25826774 | 25326774 | 26326774 | Yes |
| ENSG00000117640.17 | MTFR1L | rs1127818 | 0.0000014 | ##### | Brain - Caudate (basal ganglia | 25826774 | 25326774 | 26326774 | Yes |
| ENSG00000117640.17 | MTFR1L | rs10902675 | 0.0000015 | ##### | Brain - Cerebellum | 25826774 | 25326774 | 26326774 | Yes |
| ENSG00000117640.17 | MTFR1L | rs190043 | 0.0000015 | ##### | Brain - Cerebellar Hemisphere | 25826774 | 25326774 | 26326774 | Yes |
| ENSG00000117640.17 | MTFR1L | rs807069 | 0.0000017 | ##### | Brain - Cerebellar Hemisphere | 25826774 | 25326774 | 26326774 | Yes |
| ENSG00000117640.17 | MTFR1L | rs4659393 | 0.0000017 | ##### | Brain - Cerebellar Hemisphere | 25826774 | 25326774 | 26326774 | Yes |
| ENSG00000117640.17 | MTFR1L | rs11247824 | 0.0000017 | ##### | Brain - Cerebellar Hemisphere | 25826774 | 25326774 | 26326774 | Yes |
| ENSG00000117640.17 | MTFR1L | rs159530 | 0.0000017 | ##### | Brain - Cerebellar Hemisphere | 25826774 | 25326774 | 26326774 | Yes |
| ENSG00000117640.17 | MTFR1L | rs159527 | 0.0000017 | ##### | Brain - Cerebellar Hemisphere | 25826774 | 25326774 | 26326774 | Yes |
| ENSG00000117640.17 | MTFR1L | rs2095426 | 0.0000017 | ##### | Brain - Cerebellar Hemisphere | 25826774 | 25326774 | 26326774 | Yes |
| ENSG00000117640.17 | MTFR1L | rs3129533 | 0.0000017 | ##### | Brain - Cerebellar Hemisphere | 25826774 | 25326774 | 26326774 | Yes |
| ENSG00000117640.17 | MTFR1L | rs1537682 | 0.0000017 | ##### | Brain - Cerebellar Hemisphere | 25826774 | 25326774 | 26326774 | Yes |
| ENSG00000117640.17 | MTFR1L | rs159522 | 0.0000017 | ##### | Brain - Cerebellar Hemisphere | 25826774 | 25326774 | 26326774 | Yes |
| ENSG00000117640.17 | MTFR1L | rs159521 | 0.0000017 | ##### | Brain - Cerebellar Hemisphere | 25826774 | 25326774 | 26326774 | Yes |
| ENSG00000117640.17 | MTFR1L | rs11585897 | 0.0000019 | ##### | Brain - Cortex | 25826774 | 25326774 | 26326774 | Yes |
| ENSG00000117640.17 | MTFR1L | rs807061 | 0.000002 | ##### | Brain - Cerebellar Hemisphere | 25826774 | 25326774 | 26326774 | Yes |
| ENSG00000117640.17 | MTFR1L | rs807062 | 0.000002 | ##### | Brain - Cerebellar Hemisphere | 25826774 | 25326774 | 26326774 | Yes |
| ENSG00000117640.17 | MTFR1L | rs1961658 | 0.000002 | ##### | Brain - Cerebellar Hemisphere | 25826774 | 25326774 | 26326774 | Yes |
| ENSG00000117640.17 | MTFR1L | rs11247716 | 0.0000022 | ##### | Brain - Cerebellum | 25826774 | 25326774 | 26326774 | Yes |
| ENSG00000117640.17 | MTFR1L | rs213624 | 0.0000023 | ##### | Brain - Cerebellar Hemisphere | 25826774 | 25326774 | 26326774 | Yes |
| ENSG00000117640.17 | MTFR1L | rs2782823 | 0.0000025 | ##### | Brain - Cerebellar Hemisphere | 25826774 | 25326774 | 26326774 | Yes |
| ENSG00000117640.17 | MTFR1L | rs11247709 | 0.0000025 | ##### | Brain - Cerebellum | 25826774 | 25326774 | 26326774 | Yes |
| ENSG00000117640.17 | MTFR1L | rs2786872 | 0.0000032 | ##### | Brain - Caudate (basal ganglia | 25826774 | 25326774 | 26326774 | Yes |
| ENSG00000117640.17 | MTFR1L | rs3738450 | 0.0000035 | ##### | Brain - Caudate (basal ganglia | 25826774 | 25326774 | 26326774 | Yes |
| ENSG00000117640.17 | MTFR1L | rs74378533 | 0.0000037 | ##### | Brain - Cerebellum | 25826774 | 25326774 | 26326774 | Yes |

| | | | | | | | | | |
|---|---|---|---|---|---|---|---|---|---|
| ENSG00000117640.17 | MTFR1L | rs298439 | 0.0000044 | ##### | Brain - Cortex | 25826774 | 25326774 | 26326774 | Yes |
| ENSG00000117640.17 | MTFR1L | rs718391 | 0.0000047 | ##### | Brain - Cerebellar Hemisphere | 25826774 | 25326774 | 26326774 | Yes |
| ENSG00000117640.17 | MTFR1L | rs12030759 | 0.0000047 | ##### | Brain - Cortex | 25826774 | 25326774 | 26326774 | Yes |
| ENSG00000117640.17 | MTFR1L | rs10902676 | 0.0000048 | ##### | Brain - Cerebellum | 25826774 | 25326774 | 26326774 | Yes |
| ENSG00000117640.17 | MTFR1L | rs34239512 | 0.0000048 | ##### | Brain - Caudate (basal ganglia | 25826774 | 25326774 | 26326774 | Yes |
| ENSG00000117640.17 | MTFR1L | rs17163302 | 0.0000048 | ##### | Brain - Caudate (basal ganglia | 25826774 | 25326774 | 26326774 | Yes |
| ENSG00000117640.17 | MTFR1L | rs213629 | 0.0000048 | ##### | Brain - Caudate (basal ganglia | 25826774 | 25326774 | 26326774 | Yes |
| ENSG00000117640.17 | MTFR1L | rs2565032 | 0.0000048 | ##### | Brain - Caudate (basal ganglia | 25826774 | 25326774 | 26326774 | Yes |
| ENSG00000117640.17 | MTFR1L | rs213648 | 0.0000048 | ##### | Brain - Caudate (basal ganglia | 25826774 | 25326774 | 26326774 | Yes |
| ENSG00000117640.17 | MTFR1L | rs3738450 | 0.0000049 | ##### | Brain - Cerebellar Hemisphere | 25826774 | 25326774 | 26326774 | Yes |
| ENSG00000117640.17 | MTFR1L | rs2280997 | 0.0000056 | ##### | Brain - Caudate (basal ganglia | 25826774 | 25326774 | 26326774 | Yes |
| ENSG00000117640.17 | MTFR1L | rs2246979 | 0.0000057 | ##### | Brain - Caudate (basal ganglia | 25826774 | 25326774 | 26326774 | Yes |
| ENSG00000117640.17 | MTFR1L | rs7513196 | 0.0000057 | ##### | Brain - Cerebellar Hemisphere | 25826774 | 25326774 | 26326774 | Yes |
| ENSG00000117640.17 | MTFR1L | rs2294226 | 0.0000057 | ##### | Brain - Cerebellar Hemisphere | 25826774 | 25326774 | 26326774 | Yes |
| ENSG00000117640.17 | MTFR1L | rs159531 | 0.0000062 | ##### | Brain - Cerebellar Hemisphere | 25826774 | 25326774 | 26326774 | Yes |
| ENSG00000117640.17 | MTFR1L | rs7549097 | 0.0000064 | ##### | Brain - Cerebellar Hemisphere | 25826774 | 25326774 | 26326774 | Yes |
| ENSG00000117640.17 | MTFR1L | rs6696298 | 0.0000064 | ##### | Brain - Cerebellum | 25826774 | 25326774 | 26326774 | Yes |
| ENSG00000117640.17 | MTFR1L | rs159532 | 0.000007 | ##### | Brain - Cortex | 25826774 | 25326774 | 26326774 | Yes |
| ENSG00000117640.17 | MTFR1L | rs11247735 | 0.0000078 | ##### | Brain - Cerebellum | 25826774 | 25326774 | 26326774 | Yes |
| ENSG00000117640.17 | MTFR1L | rs807061 | 0.0000079 | ##### | Brain - Caudate (basal ganglia | 25826774 | 25326774 | 26326774 | Yes |
| ENSG00000117640.17 | MTFR1L | rs807067 | 0.000008 | ##### | Brain - Cerebellar Hemisphere | 25826774 | 25326774 | 26326774 | Yes |
| ENSG00000117640.17 | MTFR1L | rs213620 | 0.0000084 | ##### | Brain - Cerebellar Hemisphere | 25826774 | 25326774 | 26326774 | Yes |
| ENSG00000117640.17 | MTFR1L | rs11585897 | 0.0000094 | ##### | Brain - Cerebellar Hemisphere | 25826774 | 25326774 | 26326774 | Yes |
| ENSG00000117640.17 | MTFR1L | rs34239512 | 0.0000099 | ##### | Brain - Cerebellar Hemisphere | 25826774 | 25326774 | 26326774 | Yes |
| ENSG00000117640.17 | MTFR1L | rs17163302 | 0.0000099 | ##### | Brain - Cerebellar Hemisphere | 25826774 | 25326774 | 26326774 | Yes |
| ENSG00000117640.17 | MTFR1L | rs213629 | 0.0000099 | ##### | Brain - Cerebellar Hemisphere | 25826774 | 25326774 | 26326774 | Yes |
| ENSG00000117640.17 | MTFR1L | rs2565032 | 0.0000099 | ##### | Brain - Cerebellar Hemisphere | 25826774 | 25326774 | 26326774 | Yes |
| ENSG00000117640.17 | MTFR1L | rs213648 | 0.0000099 | ##### | Brain - Cerebellar Hemisphere | 25826774 | 25326774 | 26326774 | Yes |
| ENSG00000117640.17 | MTFR1L | rs10449775 | 0.00001 | ##### | Brain - Cerebellar Hemisphere | 25826774 | 25326774 | 26326774 | Yes |
| ENSG00000117640.17 | MTFR1L | rs213621 | 0.00001 | ##### | Brain - Cerebellar Hemisphere | 25826774 | 25326774 | 26326774 | Yes |
| ENSG00000117640.17 | MTFR1L | rs2744772 | 0.000011 | ##### | Brain - Caudate (basal ganglia | 25826774 | 25326774 | 26326774 | Yes |
| ENSG00000117640.17 | MTFR1L | rs213637 | 0.000011 | ##### | Brain - Cerebellar Hemisphere | 25826774 | 25326774 | 26326774 | Yes |

| | | | | | | | | | |
|---|---|---|---|---|---|---|---|---|---|
| ENSG00000117640.17 | MTFR1L | rs11247710 | 0.000013 | ##### | Brain - Cerebellar Hemisphere | 25826774 | 25326774 | 26326774 | Yes |
| ENSG00000117640.17 | MTFR1L | rs11585897 | 0.000014 | ##### | Brain - Caudate (basal ganglia | 25826774 | 25326774 | 26326774 | Yes |
| ENSG00000117640.17 | MTFR1L | rs159526 | 0.000015 | ##### | Brain - Cerebellar Hemisphere | 25826774 | 25326774 | 26326774 | Yes |
| ENSG00000117640.17 | MTFR1L | rs807060 | 0.000015 | ##### | Brain - Cerebellar Hemisphere | 25826774 | 25326774 | 26326774 | Yes |
| ENSG00000117640.17 | MTFR1L | rs159533 | 0.000017 | ##### | Brain - Cerebellar Hemisphere | 25826774 | 25326774 | 26326774 | Yes |
| ENSG00000117640.17 | MTFR1L | rs5773140 | 0.000017 | ##### | Brain - Caudate (basal ganglia | 25826774 | 25326774 | 26326774 | Yes |
| ENSG00000117640.17 | MTFR1L | rs1257163 | 0.000017 | ##### | Brain - Caudate (basal ganglia | 25826774 | 25326774 | 26326774 | Yes |
| ENSG00000117640.17 | MTFR1L | rs12405225 | 0.000017 | ##### | Brain - Cortex | 25826774 | 25326774 | 26326774 | Yes |
| ENSG00000117640.17 | MTFR1L | rs159531 | 0.000017 | ##### | Brain - Caudate (basal ganglia | 25826774 | 25326774 | 26326774 | Yes |
| ENSG00000117640.17 | MTFR1L | rs298438 | 0.000019 | ##### | Brain - Caudate (basal ganglia | 25826774 | 25326774 | 26326774 | Yes |
| ENSG00000117640.17 | MTFR1L | rs11376635 | 0.000019 | ##### | Brain - Cerebellar Hemisphere | 25826774 | 25326774 | 26326774 | Yes |
| ENSG00000117640.17 | MTFR1L | rs182455 | 0.000019 | ##### | Brain - Cerebellar Hemisphere | 25826774 | 25326774 | 26326774 | Yes |
| ENSG00000117640.17 | MTFR1L | rs807055 | 0.000019 | ##### | Brain - Cerebellar Hemisphere | 25826774 | 25326774 | 26326774 | Yes |
| ENSG00000117640.17 | MTFR1L | rs3120160 | 0.000019 | ##### | Brain - Cerebellar Hemisphere | 25826774 | 25326774 | 26326774 | Yes |
| ENSG00000117640.17 | MTFR1L | rs11589618 | 0.000021 | ##### | Brain - Cerebellar Hemisphere | 25826774 | 25326774 | 26326774 | Yes |
| ENSG00000117640.17 | MTFR1L | rs7518451 | 0.000021 | ##### | Brain - Cerebellar Hemisphere | 25826774 | 25326774 | 26326774 | Yes |
| ENSG00000117640.17 | MTFR1L | rs36024836 | 0.000021 | ##### | Brain - Cerebellar Hemisphere | 25826774 | 25326774 | 26326774 | Yes |
| ENSG00000117640.17 | MTFR1L | rs7524154 | 0.000021 | ##### | Brain - Cerebellar Hemisphere | 25826774 | 25326774 | 26326774 | Yes |
| ENSG00000117640.17 | MTFR1L | rs10719116 | 0.000021 | ##### | Brain - Cerebellar Hemisphere | 25826774 | 25326774 | 26326774 | Yes |
| ENSG00000117640.17 | MTFR1L | rs116609713 | 0.000021 | ##### | Brain - Cerebellar Hemisphere | 25826774 | 25326774 | 26326774 | Yes |
| ENSG00000117640.17 | MTFR1L | rs2997447 | 0.000021 | ##### | Brain - Frontal Cortex (BA9) | 25826774 | 25326774 | 26326774 | Yes |
| ENSG00000117640.17 | MTFR1L | rs2280997 | 0.000021 | ##### | Brain - Cerebellum | 25826774 | 25326774 | 26326774 | Yes |
| ENSG00000117640.17 | MTFR1L | rs11247709 | 0.000021 | ##### | Brain - Cerebellar Hemisphere | 25826774 | 25326774 | 26326774 | Yes |
| ENSG00000117640.17 | MTFR1L | rs4659382 | 0.000021 | ##### | Brain - Cerebellum | 25826774 | 25326774 | 26326774 | Yes |
| ENSG00000117640.17 | MTFR1L | rs298440 | 0.000021 | ##### | Brain - Cerebellar Hemisphere | 25826774 | 25326774 | 26326774 | Yes |
| ENSG00000117640.17 | MTFR1L | rs298439 | 0.000022 | ##### | Brain - Caudate (basal ganglia | 25826774 | 25326774 | 26326774 | Yes |
| ENSG00000117640.17 | MTFR1L | rs3014690 | 0.000022 | ##### | Brain - Caudate (basal ganglia | 25826774 | 25326774 | 26326774 | Yes |
| ENSG00000117640.17 | MTFR1L | rs807057 | 0.000022 | ##### | Brain - Cortex | 25826774 | 25326774 | 26326774 | Yes |
| ENSG00000117640.17 | MTFR1L | rs807056 | 0.000022 | ##### | Brain - Cortex | 25826774 | 25326774 | 26326774 | Yes |
| ENSG00000117640.17 | MTFR1L | rs2992073 | 0.000023 | ##### | Brain - Caudate (basal ganglia | 25826774 | 25326774 | 26326774 | Yes |
| ENSG00000117640.17 | MTFR1L | rs1810985 | 0.000023 | ##### | Brain - Cortex | 25826774 | 25326774 | 26326774 | Yes |
| ENSG00000117640.17 | MTFR1L | rs11247833 | 0.000023 | ##### | Brain - Cortex | 25826774 | 25326774 | 26326774 | Yes |

| | | | | | | | | | |
|---|---|---|---|---|---|---|---|---|---|
| ENSG00000117640.17 | MTFR1L | rs2280997 | 0.000025 | ##### | Brain - Cerebellar Hemisphere | 25826774 | 25326774 | 26326774 | Yes |
| ENSG00000117640.17 | MTFR1L | rs213623 | 0.000027 | ##### | Brain - Caudate (basal ganglia | 25826774 | 25326774 | 26326774 | Yes |
| ENSG00000117640.17 | MTFR1L | rs7416351 | 0.000027 | ##### | Brain - Cortex | 25826774 | 25326774 | 26326774 | Yes |
| ENSG00000117640.17 | MTFR1L | rs298440 | 0.000028 | ##### | Brain - Caudate (basal ganglia | 25826774 | 25326774 | 26326774 | Yes |
| ENSG00000117640.17 | MTFR1L | rs10902710 | 0.000028 | ##### | Brain - Cortex | 25826774 | 25326774 | 26326774 | Yes |
| ENSG00000117640.17 | MTFR1L | rs10902675 | 0.000028 | ##### | Brain - Cerebellar Hemisphere | 25826774 | 25326774 | 26326774 | Yes |
| ENSG00000117640.17 | MTFR1L | rs807062 | 0.000029 | ##### | Brain - Caudate (basal ganglia | 25826774 | 25326774 | 26326774 | Yes |
| ENSG00000117640.17 | MTFR1L | rs1961658 | 0.000029 | ##### | Brain - Caudate (basal ganglia | 25826774 | 25326774 | 26326774 | Yes |
| ENSG00000117640.17 | MTFR1L | rs298438 | 0.000031 | ##### | Brain - Cortex | 25826774 | 25326774 | 26326774 | Yes |
| ENSG00000117640.17 | MTFR1L | rs1044183 | 0.000031 | ##### | Brain - Caudate (basal ganglia | 25826774 | 25326774 | 26326774 | Yes |
| ENSG00000117640.17 | MTFR1L | rs4659382 | 0.000032 | ##### | Brain - Caudate (basal ganglia | 25826774 | 25326774 | 26326774 | Yes |
| ENSG00000117640.17 | MTFR1L | rs10902709 | 0.000032 | ##### | Brain - Cortex | 25826774 | 25326774 | 26326774 | Yes |
| ENSG00000117640.17 | MTFR1L | rs3120160 | 0.000033 | ##### | Brain - Cortex | 25826774 | 25326774 | 26326774 | Yes |
| ENSG00000117640.17 | MTFR1L | rs2294226 | 0.000035 | ##### | Brain - Cerebellum | 25826774 | 25326774 | 26326774 | Yes |
| ENSG00000117640.17 | MTFR1L | rs159526 | 0.000037 | ##### | Brain - Caudate (basal ganglia | 25826774 | 25326774 | 26326774 | Yes |
| ENSG00000117640.17 | MTFR1L | rs3738450 | 0.000038 | ##### | Brain - Cerebellum | 25826774 | 25326774 | 26326774 | Yes |
| ENSG00000117640.17 | MTFR1L | rs6659086 | 0.000038 | ##### | Brain - Cerebellar Hemisphere | 25826774 | 25326774 | 26326774 | Yes |
| ENSG00000117640.17 | MTFR1L | rs4659394 | 0.000042 | ##### | Brain - Caudate (basal ganglia | 25826774 | 25326774 | 26326774 | Yes |
| ENSG00000117640.17 | MTFR1L | rs11247716 | 0.000044 | ##### | Brain - Cerebellar Hemisphere | 25826774 | 25326774 | 26326774 | Yes |
| ENSG00000117640.17 | MTFR1L | rs159519 | 0.000044 | ##### | Brain - Cerebellar Hemisphere | 25826774 | 25326774 | 26326774 | Yes |
| ENSG00000117640.17 | MTFR1L | rs6696298 | 0.000045 | ##### | Brain - Cerebellar Hemisphere | 25826774 | 25326774 | 26326774 | Yes |
| ENSG00000117640.17 | MTFR1L | rs10902711 | 0.000049 | ##### | Brain - Cortex | 25826774 | 25326774 | 26326774 | Yes |
| ENSG00000117640.17 | MTFR1L | rs149560263 | 0.000049 | ##### | Brain - Cortex | 25826774 | 25326774 | 26326774 | Yes |
| ENSG00000117640.17 | MTFR1L | rs807060 | 0.000053 | ##### | Brain - Cortex | 25826774 | 25326774 | 26326774 | Yes |
| ENSG00000117640.17 | MTFR1L | rs56361727 | 0.000054 | ##### | Brain - Cortex | 25826774 | 25326774 | 26326774 | Yes |
| ENSG00000117640.17 | MTFR1L | rs4659382 | 0.000056 | ##### | Brain - Cerebellar Hemisphere | 25826774 | 25326774 | 26326774 | Yes |
| ENSG00000117640.17 | MTFR1L | rs7524771 | 0.000058 | ##### | Brain - Cerebellar Hemisphere | 25826774 | 25326774 | 26326774 | Yes |
| ENSG00000117640.17 | MTFR1L | rs11247735 | 0.000059 | ##### | Brain - Cerebellar Hemisphere | 25826774 | 25326774 | 26326774 | Yes |
| ENSG00000117640.17 | MTFR1L | rs5773140 | 0.000062 | ##### | Brain - Cerebellum | 25826774 | 25326774 | 26326774 | Yes |
| ENSG00000117640.17 | MTFR1L | rs1257163 | 0.000062 | ##### | Brain - Cerebellum | 25826774 | 25326774 | 26326774 | Yes |
| ENSG00000117640.17 | MTFR1L | rs6872 | 0.000062 | ##### | Brain - Cerebellum | 25826774 | 25326774 | 26326774 | Yes |
| ENSG00000117640.17 | MTFR1L | rs159519 | 0.000063 | ##### | Brain - Cortex | 25826774 | 25326774 | 26326774 | Yes |

| | | | | | | | | | |
|---|---|---|---|---|---|---|---|---|---|
| ENSG00000117640.17 | MTFR1L | rs2782823 | 0.000065 | ##### | Brain - Caudate (basal ganglia | 25826774 | 25326774 | 26326774 | Yes |
| ENSG00000117640.17 | MTFR1L | rs10902676 | 0.000069 | ##### | Brain - Cerebellar Hemisphere | 25826774 | 25326774 | 26326774 | Yes |
| ENSG00000117640.17 | MTFR1L | rs3120159 | 0.000069 | ##### | Brain - Cortex | 25826774 | 25326774 | 26326774 | Yes |
| ENSG00000117640.17 | MTFR1L | rs11585897 | 0.000071 | ##### | Brain - Cerebellum | 25826774 | 25326774 | 26326774 | Yes |
| ENSG00000117640.17 | MTFR1L | rs2992065 | 0.000073 | ##### | Brain - Cortex | 25826774 | 25326774 | 26326774 | Yes |
| ENSG00000117640.17 | MTFR1L | rs2992073 | 0.000073 | ##### | Brain - Cerebellum | 25826774 | 25326774 | 26326774 | Yes |
| ENSG00000117640.17 | MTFR1L | rs807061 | 0.000074 | ##### | Brain - Cerebellum | 25826774 | 25326774 | 26326774 | Yes |
| ENSG00000117640.17 | MTFR1L | rs34239512 | 0.000083 | ##### | Brain - Cerebellum | 25826774 | 25326774 | 26326774 | Yes |
| ENSG00000117640.17 | MTFR1L | rs17163302 | 0.000083 | ##### | Brain - Cerebellum | 25826774 | 25326774 | 26326774 | Yes |
| ENSG00000117640.17 | MTFR1L | rs213629 | 0.000083 | ##### | Brain - Cerebellum | 25826774 | 25326774 | 26326774 | Yes |
| ENSG00000117640.17 | MTFR1L | rs2565032 | 0.000083 | ##### | Brain - Cerebellum | 25826774 | 25326774 | 26326774 | Yes |
| ENSG00000117640.17 | MTFR1L | rs213648 | 0.000083 | ##### | Brain - Cerebellum | 25826774 | 25326774 | 26326774 | Yes |
| ENSG00000117640.17 | MTFR1L | rs807062 | 0.000098 | ##### | Brain - Cerebellum | 25826774 | 25326774 | 26326774 | Yes |
| ENSG00000117640.17 | MTFR1L | rs6694897 | 0.000098 | ##### | Brain - Cerebellum | 25826774 | 25326774 | 26326774 | Yes |
| ENSG00000117640.17 | MTFR1L | rs3120159 | 0.0001 | ##### | Brain - Cerebellar Hemisphere | 25826774 | 25326774 | 26326774 | Yes |
