## Supplementary table 13 for "Exome-wide analysis reveals role of *LRP1* and additional novel loci in cognition"

**Suuplementary Table 13: eQTL variants controlling expression of *APOC1* and *LRP1* in the brain**

| **Header Descriptions** | Gencode ID, Gene Symbol for mapped gene: Gene ID and Symbol of genes identified to be associated with complex processing speed and visual attention from our gene-based tests<br>SNP Id: rsid of significant eQTLs<br>NES: Normalized effect size of association of the significant eQTLs with tissue expression<br>P-value: p-value of association |
|---|---|

| Gencode Id | Gene Symbo | Variant ID | SNP Id | P-Value | NES | Tissue |
|---|---|---|---|---|---|---|
| ENSG000001 | APOC1 | chr19_44382483_A_G_ | rs113265033 | 0.0000034 | 1.80E+00 | Brain - Amygdala |
| ENSG000001 | APOC1 | chr19_44946776_C_T_ | rs9304644 | 0.0000039 | 3.10E-01 | Brain - Spinal cord (cervical c-1) |
| ENSG000001 | APOC1 | chr19_44950979_G_A_ | rs7246900 | 0.0000053 | 3.10E-01 | Brain - Spinal cord (cervical c-1) |
| ENSG000001 | APOC1 | chr19_44951502_A_G_ | rs7247227 | 0.0000053 | 3.10E-01 | Brain - Spinal cord (cervical c-1) |
| ENSG000001 | APOC1 | chr19_44952201_G_A_ | rs892101 | 0.0000053 | 3.10E-01 | Brain - Spinal cord (cervical c-1) |
| ENSG000001 | APOC1 | chr19_44952844_G_A_ | rs4803777 | 0.0000053 | 3.10E-01 | Brain - Spinal cord (cervical c-1) |
| ENSG000001 | APOC1 | chr19_44953923_T_C_ | rs3760627 | 0.0000053 | 3.10E-01 | Brain - Spinal cord (cervical c-1) |
| ENSG000001 | APOC1 | chr19_44954036_C_T_ | rs66867801 | 0.0000053 | 3.10E-01 | Brain - Spinal cord (cervical c-1) |
| ENSG000001 | APOC1 | chr19_44954062_G_A_ | rs66771331 | 0.0000053 | 3.10E-01 | Brain - Spinal cord (cervical c-1) |
| ENSG000001 | APOC1 | chr19_44924096_C_G_ | rs4803770 | 0.000016 | -2.10E-01 | Brain - Nucleus accumbens (basal ganglia) |
| ENSG000001 | APOC1 | chr19_44918715_AG_/ | rs12721052 | 0.000017 | -2.10E-01 | Brain - Nucleus accumbens (basal ganglia) |
| ENSG000001 | APOC1 | chr19_44913221_A_G_ | rs584007 | 0.000019 | -2.40E-01 | Brain - Nucleus accumbens (basal ganglia) |
| ENSG000001 | APOC1 | chr19_44913034_C_T_ | rs59325138 | 0.000034 | -2.00E-01 | Brain - Nucleus accumbens (basal ganglia) |
| ENSG000001 | APOC1 | chr19_44915704_T_C_ | rs3826688 | 0.000036 | -2.30E-01 | Brain - Nucleus accumbens (basal ganglia) |
| ENSG000001 | APOC1 | chr19_44967656_C_T_ | rs10775543 | 0.000042 | 3.50E-01 | Brain - Nucleus accumbens (basal ganglia) |
| ENSG000001 | LRP1 | chr12_56963245_T_C_ | rs533784783 | 0.0000049 | 1.00E+00 | Brain - Putamen (basal ganglia) |
| ENSG000001 | LRP1 | chr12_56983563_C_T_ | rs4759042 | 0.0000086 | 1.70E-01 | Brain - Cerebellar Hemisphere |
| ENSG000001 | LRP1 | chr12_56972426_A_G_ | rs7132082 | 0.00001 | 1.70E-01 | Brain - Cerebellar Hemisphere |
| ENSG000001 | LRP1 | chr12_56929739_A_G_ | rs840161 | 0.000015 | -1.70E-01 | Brain - Cerebellar Hemisphere |
| ENSG000001 | LRP1 | chr12_56932208_C_C/ | rs35668586 | 0.000015 | -1.70E-01 | Brain - Cerebellar Hemisphere |
| ENSG000001 | LRP1 | chr12_56968638_G_A_ | rs1843314 | 0.000018 | 1.70E-01 | Brain - Cerebellar Hemisphere |
| ENSG000001 | LRP1 | chr12_56878830_C_T_ | rs4759036 | 0.000027 | 1.50E-01 | Brain - Cerebellar Hemisphere |

| | | | | | | |
|---|---|---|---|---|---|---|
| ENSG000001 | LRP1 | chr12_56879082_G_A | rs9668259 | 0.000027 | 1.50E-01 | Brain - Cerebellar Hemisphere |
| ENSG000001 | LRP1 | chr12_56879697_C_T | rs7961602 | 0.000027 | 1.50E-01 | Brain - Cerebellar Hemisphere |
| ENSG000001 | LRP1 | chr12_56881744_C_A | rs4477473 | 0.000027 | 1.50E-01 | Brain - Cerebellar Hemisphere |
| ENSG000001 | LRP1 | chr12_56882855_T_C | rs7960618 | 0.000027 | 1.50E-01 | Brain - Cerebellar Hemisphere |
| ENSG000001 | LRP1 | chr12_56883024_G_T | rs7955918 | 0.000027 | 1.50E-01 | Brain - Cerebellar Hemisphere |
| ENSG000001 | LRP1 | chr12_56884055_A_G | rs7959654 | 0.000027 | 1.50E-01 | Brain - Cerebellar Hemisphere |
| ENSG000001 | LRP1 | chr12_56884384_C_T | rs7960137 | 0.000027 | 1.50E-01 | Brain - Cerebellar Hemisphere |
| ENSG000001 | LRP1 | chr12_56885817_T_C | rs7979655 | 0.000027 | 1.50E-01 | Brain - Cerebellar Hemisphere |
| ENSG000001 | LRP1 | chr12_56886172_G_T | rs7967940 | 0.000027 | 1.50E-01 | Brain - Cerebellar Hemisphere |
| ENSG000001 | LRP1 | chr12_56887050_CAT | rs55904820 | 0.000027 | 1.50E-01 | Brain - Cerebellar Hemisphere |
| ENSG000001 | LRP1 | chr12_56887618_G_A | rs7972591 | 0.000027 | 1.50E-01 | Brain - Cerebellar Hemisphere |
| ENSG000001 | LRP1 | chr12_56889007_C_T | rs11172033 | 0.000027 | 1.50E-01 | Brain - Cerebellar Hemisphere |
| ENSG000001 | LRP1 | chr12_56889425_A_G | rs56023460 | 0.000027 | 1.50E-01 | Brain - Cerebellar Hemisphere |
| ENSG000001 | LRP1 | chr12_56891306_C_T | rs7959725 | 0.000027 | 1.50E-01 | Brain - Cerebellar Hemisphere |
| ENSG000001 | LRP1 | chr12_56893209_T_A | rs12425604 | 0.000027 | 1.50E-01 | Brain - Cerebellar Hemisphere |
| ENSG000001 | LRP1 | chr12_56893910_G_T | rs55812315 | 0.000027 | 1.50E-01 | Brain - Cerebellar Hemisphere |
| ENSG000001 | LRP1 | chr12_56897715_C_G | rs4759041 | 0.000027 | 1.50E-01 | Brain - Cerebellar Hemisphere |
| ENSG000001 | LRP1 | chr12_56897910_A_C | rs61939604 | 0.000027 | 1.50E-01 | Brain - Cerebellar Hemisphere |
| ENSG000001 | LRP1 | chr12_56902640_G_A | rs2888108 | 0.000027 | -1.50E-01 | Brain - Cerebellar Hemisphere |
| ENSG000001 | LRP1 | chr12_56907610_T_C | rs2888107 | 0.000027 | -1.50E-01 | Brain - Cerebellar Hemisphere |
| ENSG000001 | LRP1 | chr12_56907614_C_T | rs2888106 | 0.000027 | -1.50E-01 | Brain - Cerebellar Hemisphere |
| ENSG000001 | LRP1 | chr12_57149789_C_T | rs10876966 | 0.000036 | 2.50E-01 | Brain - Cerebellum |
| ENSG000001 | LRP1 | chr12_56890814_C_T | rs7956391 | 0.000038 | 1.50E-01 | Brain - Cerebellar Hemisphere |
| ENSG000001 | LRP1 | chr12_56880864_A_G | rs10876943 | 0.000039 | 1.50E-01 | Brain - Cerebellar Hemisphere |
| ENSG000001 | LRP1 | chr12_56890843_TTG | rs144095801 | 0.000039 | 1.50E-01 | Brain - Cerebellar Hemisphere |
| ENSG000001 | LRP1 | chr12_56890939_A_G | rs7959253 | 0.000039 | 1.50E-01 | Brain - Cerebellar Hemisphere |
| ENSG000001 | LRP1 | chr12_56892579_C_C | rs56073765 | 0.000039 | 1.50E-01 | Brain - Cerebellar Hemisphere |
| ENSG000001 | LRP1 | chr12_56892580_A_A | rs55851749 | 0.000039 | 1.50E-01 | Brain - Cerebellar Hemisphere |
| ENSG000001 | LRP1 | chr12_56892581_C_G | rs55889072 | 0.000039 | 1.50E-01 | Brain - Cerebellar Hemisphere |
| ENSG000001 | LRP1 | chr12_56853738_G_A | rs150248294 | 0.000075 | 8.20E-01 | Brain - Putamen (basal ganglia) |
| ENSG000001 | LRP1 | chr12_56879533_C_T | rs181457987 | 0.000075 | 8.20E-01 | Brain - Putamen (basal ganglia) |
| ENSG000001 | LRP1 | chr12_56901340_T_C | rs182925459 | 0.000075 | 8.20E-01 | Brain - Putamen (basal ganglia) |

| | | | | | | |
|---|---|---|---|---|---|---|
| ENSG000001: | LRP1 | chr12_56904636_T_C_ | rs151016514 | 0.000075 | 8.20E-01 | Brain - Putamen (basal ganglia) |
| ENSG000001: | LRP1 | chr12_56909015_A_C_ | rs1846398 | 0.000088 | -1.40E-01 | Brain - Cerebellar Hemisphere |
| ENSG000001: | LRP1 | chr12_56912893_C_T_ | rs10735871 | 0.000088 | -1.40E-01 | Brain - Cerebellar Hemisphere |
| ENSG000001: | LRP1 | chr12_56915152_G_A_ | rs1909328 | 0.000088 | -1.40E-01 | Brain - Cerebellar Hemisphere |
| ENSG000001: | LRP1 | chr12_57052061_A_C_ | rs324021 | 0.00011 | 1.50E-01 | Brain - Cerebellar Hemisphere |
| ENSG000001: | LRP1 | chr12_57056027_C_T_ | rs1044931 | 0.00011 | 1.50E-01 | Brain - Cerebellar Hemisphere |
