## Supplementary information for "Exome-wide analysis reveals role of *LRP1* and additional novel loci in cognition"

**Phenotype selection:**

We chose mean time to correctly identify matches (Field Id: 20023), fluid intelligence score (Field Id: 20016), maximum digits remembered correctly (Field Id: 20240), number of symbol digit matches made correctly (Field Id: 20159), alphanumeric path (Field Id: 20157) as measures of reaction time, fluid intelligence, digit recall and digit-symbol substitution, trail-making respectively. We derived proportion of incorrect matches in a round obtained as the ratio of number of incorrect matches in a row (Field Id:20132) to the sum of number of correct (Field Id: 20131) and incorrect matches in a row as a measure of pair-matching.

Apart from these, we also considered genetic sex (Field Id: 22001), age at recruitment (Field Id: 21022), educational qualification (Field Id: 6138) as covariates. We categorized educational qualification into three categories- i) College or university degree (code: 1), ii) A levels/ AS levels or equivalent (code: 2), and iii) Either of O-levels/ GCSEs or equivalent (code: 3), CSEs or equivalent (code: 4) or NVQ or HND or HNC equivalent (code:5) (where, GCSE: General Certificate of Secondary Education ,CSE: Certificate of Secondary Education, NVQ: National Vocation Qualification, HNC: Higher National Certificate; HND: Higher National Diploma).

**Genotype Quality Control:**

Each chromosome data was split across multiple blocks. However, all the blocks did not have variant information for all samples. We deemed the blocks which did not have information for all 200,624 samples as incompatible for merging, and excluded those blocks. Our resultant working file contained 13,567,285 variants on autosomes for 200,624 individuals. We filtered biallelic variants and set half-calls to missing. We excluded samples with a sample-call rate of < 99% and excluded variants with variant call-rate < 99%. We also excluded variants with a minor allele frequency (MAF) <0.001 in the entire dataset. Using these filters, we retained 187,154 individuals with 239,076 variants. To detect sample outliers based on their deviation from European ancestry, we extracted these 239,076 variants from five 1000 genomes superpopulations namely CEU (Utah residents (CEPH) with Northern and Western European ancestry), TSI (Toscani in Italia), GBR (British in England and Scotland), FIN (Finnish in Finland) and IBS (Iberian populations in Spain) (<http://ftp.1000genomes.ebi.ac.uk/vol1/ftp/>). We removed 119 duplicate variants and then we pruned the variants with a window size of 500kb and pairwise $r^{2}$>0.05 to get an independent set of variants. We built the kinship using GCTA model^1^ based on the independent set of variants so obtained. It is to be noted that these variants were pruned only to get the kinship matrix. Thereafter, we calculated top 20 principal components (PCs) on the pruned 1000 genomes data. We then mapped the partially filtered UK Biobank data onto these principal components and identified 22,063 samples as outliers which we consequently removed from the study. Then, we removed samples with heterozygosity rate exceeding four standard deviations from mean heterozygosity to exclude samples with low sample quality and high inbreeding. We also excluded variants which deviate from the Hardy Weinberg equilibrium with a p-value < ${10}^{-4}$. To remove cryptic relatedness beyond 2^nd^ degree, we used plink2’s ‘king-cutoff’ flag so that no two individuals have a kinship coefficient > 0.0884. We also removed individuals who had withdrawn from the UK Biobank study at this stage leading to a sample size of 157,160 with 211,134 variants.

We additionally carried out a few quality control steps to assess variant calling quality. We excluded 37 variants with a QUAL score < 30. We annotated the variants using GATK ‘VariantAnnotator’ tool. All remaining variants had quality by depth (QD) ≤ 20, median mapping quality (MMQ) ≥ 40 and inbreeding coefficient ≥-0.8. Per-sample genotyping-quality (GQ) was also ≥ 20 for all variants. We used a filtering criterion to exclude variants for which >1% of the samples had missing sample -level DP information, but no variants were excluded with this filter. We removed SNPs for which >10% samples had a DP < 7 and Indels for which >10% samples had DP < 10. We retained variants having no homozygous alternate allele carrier and no heterozygous alternate allele carrier with an allele balance (AB) ≥ 0.15 for SNPs and ≥ 0.20 for indels.^2,3^ We calculated allele balance for each sample as the ratio of per sample allele depth (AD) for alternative allele to the sum of per-sample allele depths for both reference and alternate allele. These steps were critically carried out to remove variants with poor genotyping quality so as to reduce false-positive findings in downstream analyses. After all quality checks, we retained 157,160 individuals with 211,012 variants (Supplementary Table 1).

**Heritability Analysis:**

Heritability is a measure of the amount of observable variation in a population that can be attributed to inherited individual genetic factors. Broad-sense heritability refers to the ratio of total genetic variance to total phenotypic variance whereas narrow sense heritability is the ratio of additive genetic variance to total phenotypic variance.

At first, we used the BLD-LDAK^4^ model to perform heritability analysis from summary statistics for each cognition phenotype. This BLD-LDAK model computes the expected heritability contributed by a SNP depending on its minor allele frequency (MAF), linkage disequilibrium (LD) and 64 functional annotations (6 LD-related annotations, 24 function indicators and 34 auxiliary annotations, detailed information in supplementary)^4^ which are largely indicators of buffer regions for the functional categories. We employed the following heritability model:

$$E\left[ h_{j}^{2} \right]=\left[ f_{j}\left( 1-f_{j} \right) \right]^{0.75}(\tau_{1}b_{1j}+\tau_{2}b_{2j}+\ldots+\tau_{64}b_{64j}+\tau_{65}w_{j}+\tau_{66})$$

Where$f_{j}$ represents the MAF of j^th^ variant, $b_{1}, b_{2},{\ldots,b}_{64}$are the annotations and $w_{j}$ represents the LDAK weighting which tends to be lower for regions in high LD and these weights are computed using only high-quality SNPs. These annotations are predefined using the genomic coordinates from the GRCh37/hg19 build. But UK Biobank data is in GRCh38/hg38. So, we used CrossMap liftover tool.^5^ 413 variants failed to get mapped to the target build. Of the remaining 210599 variants, 5593 were multi-character allelic variants. We thus performed heritability analyses with 205,006 SNPs. We generated summary statistics by classical linear regression model of phenotype on genotype adjusting for age, sex.

However, we observed that for some of the cognitive phenotypes, this model was over-complicated and gave incorrect estimates due to high number of predictors. Thus, we considered the LDAK model to conduct heritability analyses of our cognitive phenotypes.

The LDAK model^6^ computes the expected heritability contributed by a SNP depending on its minor allele frequency (MAF) and linkage disequilibrium (LD). We finally employed the heritability model:

$$E\left[ h_{j}^{2} \right]={\tau_{1}w_{j}\left[ f_{j}\left( 1-f_{j} \right) \right]}^{0.75}$$

wher$e f_{j}'s$ are the MAFs and $w_{j}$ *a*re the LDAK weights which tends to be lower for regions in high LD and vice-versa. These weights are computed using only high-quality SNPs. At first, the SNPs were chunked into sections with each section containing approximately 1000 SNPs (buffer of 100kb is kept on either side). Then LD pruning was carried out, so that no pair of SNPs within a region of 100kb had squared correlation greater than 0.98. To these SNPs, non-zero weights based on LD were assigned. Here again, we performed the heritability analyses with 205,006 SNPs with co-ordinates corresponding to hg19 build.

**Single Variant Association**

***Baseline Model:***

We regressed each cognitive phenotype on age, gender, educational qualification, top 10 principal components, and *APOE*-carrier status as described above, and then inverse-normalized the residuals before fitting it against genotype. The model we used was as follows:

$Y_{i}=\mu_{i}+\delta_{1i}{age}_{i}+\delta_{2i}{sex}_{i}+\delta_{3i}{Edu}_{i}+\delta_{4i}{APOE status}_{i}+\delta_{5i}{PC}_{1i}++\delta_{14i}{PC}_{10i}+\varepsilon_{i}$ …..[1]

$$Inverse\_Normal(e_{i})=\alpha_{j}+\beta_{j}X_{ij}+I.................................................[2]$$

where $Y_{i}$ is the phenotype value for i^th^ individual, $e_{i}$ is the residual for i^th^ individual from the first equation and $X_{\mathrm{ij}}$ is the number of alternate alleles in the genotype of i^th^ individual with respect to j^th^ genetic variant. $X_{\mathrm{ij}}$ can be 0, 1 or 2 and is assumed to follow a binomial distribution with alternate allele frequency as success probability. The null hypothesis for each of the j variants is $H_{0}: \beta_{j}=0$ was tested using Wald’s test statistic $T_{j}=\frac{\hat{\beta}_{j}^{2}}{var\left( \hat{\beta}_{j} \right)}\sim\chi_{1}^{2}$ under $H_{0}$ with p-value *P(*$\chi_{1}^{2}>$ $T_{j})$. The proportion of variation explained by the j^th^ variant was then calculated using $2\beta_{j}^{2}f_{j}\left( 1-f_{j} \right)$ where $f_{j}$ is the alternate allele frequency of the j^th^ variant. This served as the baseline model.

***Model with lipid/glycemic traits as covariates:***

Further, to control for age-related metabolic conditions that could adversely affect cognition, we added lipid levels (serum total cholesterol, triglycerides, HDL and LDL Direct) (referred to as Model 2 in main text), glucose and HbA1c levels (referred to as Model 3 in main text) separately as covariates to the baseline model (equation 1) to obtain the residuals for association testing.

We considered an exome-wide level of significance at 0.05/211012 =2.37*10^-7^ (Bonferroni adjusted). For each of these tests, if the genomic inflation factor was greater than 1.1, we applied genomic control by dividing each test statistic for all variants tested (say m) by the genomic inflation factor calculated as $\lambda=\frac{median(T_{1},T_{2},\ldots,T_{m})}{median(\chi_{1}^{2})}$ .

The single-variant association results based on whole exome obtained from five cognitive domains (there were no significant variants associated with the phenotype for working memory with genomic inflation factor *λ_GC_* < 1.1) have been given in Table 1 (summarized) and Supplementary Table 4 (detailed).

**Genes functioning along with *APOE* in pathways relevant to cognition:**

In the central nervous system, *APOE* is majorly expressed in astrocytes, microglia, vascular mural cells and choroid plexus cells.^7^ Post secretion from these cells, it is lipidated by cell surface ATP-binding cassette transporters to form lipoprotein particles and functions in transporting cholesterol and other lipids to neurons by binding to cell-surface receptors in an isoform-dependent manner.^8^ ApoE4, the isoform conferring maximum risk for Alzheimer’s and related dementia is associated with suppressed lipid metabolism and inefficient delivery of essential fatty acids (EFAs) like DHA, that are required in synaptic maintenance, to cerebral neurons resulting in inhibited functioning of glucose transporters like *GLUT* resulting in decreased glucose uptake in the brain, which is one of the earliest signs of AD.^9–12^ Reduced glucose levels in the brain lead to depletion of acetyl-Co A, resulting in reduced synthesis of acetylcholine^12^ which leads to impairment in neurotransmission. Cholesterol in the brain is majorly synthesized endogenously and is a complex process that starts with converting acetyl-Co A into a series of intermediate products to yield the final product as cholesterol.^13^ Changes in levels of acetyl-CoA lead to disruption of cholesterol homeostasis. Abnormally lipidated ApoE-containing high-density lipoprotein (HDL)-like lipoproteins binds to neuronal surface low-density lipoprotein receptors, encoded by *LDLR* and *LRP1* and is internalized via receptor-mediated endocytosis and degraded to release cholesterol that can be used for synaptic formation.^14^ Changes in cholesterol homeostasis leads to aberrant processing of APP protein leading to accumulation of Aβ and cell death^13^ ApoE enhances seeding and fibrillization of Aβ and further promotes Aβ deposition. The clearance of soluble Aβ occurs in an isoform-dependent manner, with ApoE4 being the least efficient. The lipidation status of *APOE* also plays an important role here.^15^ The cerebrovascular system is another major Aβ clearance pathway. Aβ binds to ApoE to form a complex which impedes its clearance across the blood-brain barrier via *LRP1* and *LRP2* and triglyceride-rich very low-density lipoprotein receptor (VLDR).^16^ ApoE (especially E4) impedes clearance of Aβ by binding competitively to Aβ receptors of the *LDLR* family that are present on the surface of the glial cells for their uptake and lysosomal degradation.^8^ It also mediates microglial response to amyloid plaques. ApoE also plays a role in neuroinflammatory pathways in an isoform dependent manner. ApoE-binding may interfere with other pathways of lipid metabolism that are routed through the same receptors. Figure 1 summarizes the putative pathways in which *APOE*-isoforms could potentially regulate neuronal dysfunction. Thus, for this analysis, we identify and retrieve genes^17^ by using search terms: APOE + amyloid, APOE + cholesterol, APOE + lipoprotein, APOE + cholesterol + amyloid beta, APOE + lipoprotein + amyloid-beta, APOE + glucose, APOE + glucose + amyloid-beta, APOE + triglycerides, APOE + inflammatory response. We get *TREM2, APP A4, LRP1, SORL1, PSEN1, LDLR , LRP8, SCARB1, LCAT, ABCA1, ANGPTL3, APOC1, APOBR, LRP10, MAPT, LGALS1 , NR1H4, PDCD4, HP, SNCA.*

**Gene-based association tests:**

To determine the influence of all variants on a gene to the phenotype of the i^th^ individual we used the following baseline model :

$y_{i}=\alpha_{0}+\alpha^{'}\boldsymbol{X}_{i}+\beta^{'}\boldsymbol{G}_{i}+\varepsilon_{i}$ where  *α=* [α_1_,…, α_m_]*'* correspond to the vector of regression coefficients for the m covariates pertaining to age, gender, PCs educational qualification, *APOE* carrier status and the lipid and glycaemic phenotypes in some cases, ***β*** *= [*β_1_,…,β_p_*]'* correspond to the vector of regression coefficients for the p observed genetic variants in the gene considered, and ɛ_i_ is an error term with a mean of zero and a variance of σ^2^. To test whether all these variants in the gene influence the phenotype, our null hypothesis is *H_0_:* ***β = 0***, that is, β_1_ = β_2_ = … = β_p_ = 0 . This test assumes that each β_j_ follows an arbitrary distribution with a mean of zero and a variance of w_j_τ, where τ is a variance component and w_j_ is the prespecified weight for variant j. Then H_0_*:* ***β = 0*** boils down to testing H_0_: τ = 0. The SKAT test statistic is as follows

$$\boldsymbol{Q}_{\boldsymbol{K}}={(\mathbf{y}\mathbf{-}\hat{\boldsymbol{\mu}})}^{'}\mathbf{K}(\mathbf{y}\mathbf{-}\hat{\boldsymbol{\mu}})$$

where $\hat{\boldsymbol{\mu}}$ is the predicted mean of phenotype y under H_0_, ***K*** is an n × n matrix called the kernel function with the (i, i')^th^ element equal to  $K(\boldsymbol{G}_{\boldsymbol{i}},\boldsymbol{G}_{\boldsymbol{i'}})=\sum_{j=1}^{p} w_{j}G_{ij}G_{i'j}$ which measures the genetic similarity between subjects i and i' with respect to the p variants in the gene considered. Each weight $w_{j}$signify the relative contribution of the j^th^ variant to the score statistic. We set $\sqrt{w_{j}}$ =Beta($\mathrm{MAF}_{j} ;a_{1}$=1 $a_{2}$=25), the beta distribution density function calculated at the sample minor-allele frequency (MAF) for the j^th^ variant. This choice of parameters $a_{1}$ and $a_{2}$ puts more weight on the contribution of rare variants with decent non-zero weight on common variants. Under the null hypothesis, Q_K_ follows a mixture of chi-square distributions which is approximated computationally using Davies method.^18,19^

The burden method ^20^ assumes each ${\beta_{\mathbf{j}}=w}_{j}^{'}\beta_{\mathbf{c}}$ with each w_j_ being a function of MAF of the j^th^ variant. The burden score statistic is:

$$Q_{B}=\left[ \sum_{i=1}^{n} \left( y_{i}-\hat{\mu}_{i} \right)\left( \sum_{j=1}^{p} w_{j}^{'}g_{\mathrm{ij}} \right) \right]^{2}$$

The SKAT-O test statistic is thus given by $Q=\rho Q_{B}+\left( 1-\rho\right)Q_{K};0\leq\rho\leq1$ which asymptotically follows a mixture of a chi-square distribution with one degree of freedom, and an approximated mixture of chi-square distributions with a proper adjustment.

To summarize, we performed both SKAT and SKAT-O tests on cognitive phenotypes with age, gender, educational status, *APOE*-carrier status and top 10 principal components as covariates at first. With this baseline model, we added lipid levels (cholesterol, triglycerides, HDL and LDL Direct), glucose and HbA1c levels separately as covariates. Thus, for each of the phenotypes, we conducted seven gene-based tests. Here again, association was based on inverse-normalized residuals for the phenotypes.

We set the level of significance for these tests at 0.0025 (Bonferroni adjusted) and obtained significant results from two cognitive domains (complex processing speed and visual attention) as summarized in (Supplementary Table 5).

**Pairwise interaction analyses:**

We performed our analysis using the following model for ${H_{0}:\beta}_{3}=0$, when

$$Y=\beta_{0}+ \beta_{1}X_{1}+\beta_{2}X_{2}+\beta_{3}X_{1}X_{2}+\varepsilon$$

where Y’s are the inverse-normalized residuals obtained as stated in main text, X_1_ and X_2_ are the genotypes of the two SNPs whose interaction is being tested , X_1_X_2_ is their genotypic combination with $\beta_{1},\beta_{2}$ being their main effects and $\beta_{3}$ signifying the interaction effect of the two SNPs. The test statistic used is $T=\frac{\hat{\beta}_{3}^{2}}{\mathrm{var}\left( \hat{\beta}_{3} \right)}\sim\chi_{1}^{2}$ under H_0_ (p < 0.05 considered for significance).

Our results from these test have been summarized in Tables 2 and 3.We used the nominal level of significance for each of these tests.

**Bivariate association tests:**

We jointly analysed each cognitive test with each of the lipid and glucose phenotypes. For the two traits (one cognitive and the other lipid/glucose phenotype), we had our marginal models as follows.

$$\mathbf{Y}_{\mathbf{k}}=\alpha_{\mathbf{k}}+\beta_{k}\mathbf{X}+\varepsilon,$$

k=1,2 where $\beta_{k}$ is the genetic effect on the k^th^ trait and $Y_{k}$ is the vector of covariate adjusted response for the k^th^ trait. The Wald statistic for the k^th^ hypothesis $H_{0,k}: \beta_{k}=0$ is given by $Z_{k}=\frac{\hat{\beta_{k}}}{\mathrm{se}\left( \hat{\beta}_{k} \right)}$

The global null hypothesis is $H_{0}: \beta_{1}=\beta_{2}=0$. Under $H_{0}$, $\mathbf{Z}=(Z_{1},Z_{2})'$ has an asymptotic bivariate normal distribution with mean **0** and covariance matrix R, where R represents the 2 × 2 correlation matrix of the two traits $Y_{1}$ and $Y_{2}$. The test statistic is given by:

$T_{metaMANOVA}=Z^{'}\hat{R}^{-1}Z$ which under $H_{0}$ asymptotically follows $\chi_{2}^{2}$ distribution. We considered 1.25*10-7 as level of significance and summarized our results in Supplementary Tables 6-10.

**Mediation Analysis:**

We have carried out single variant association with each lipid and glycemic test using our baseline covariates to find out if our variants identified from the single variant association test are also associated lipid and/or glycemic phenotypes. We have considered an exome-wide significance level of ~2.5*10^-7^ for this association and a suggestive significance level of 10^-5^. If the variants associated with lipid and /or glycemic phenotypes were associated to the respective cognition phenotypes, when additionally controlled for that lipid/glycemic trait, with a reduced absolute effect size, we considered suggestive mediation effect of that variant. For this analysis we considered p-value 10^-5^ as suggestive evidence of association.

**Supplementary Figures**

**Supplementary Figure 1: Manhattan plots for mean reaction time**

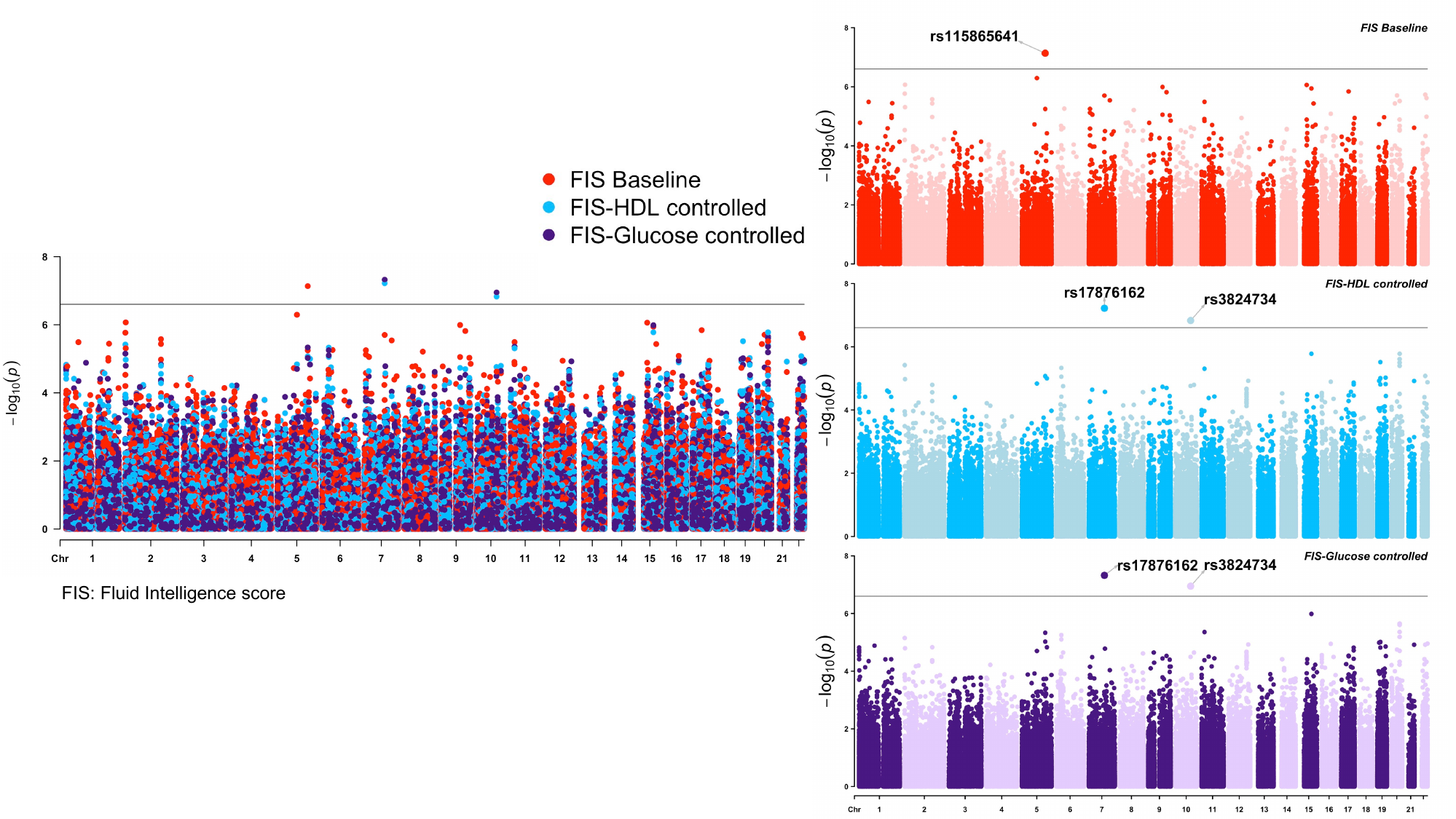
Manhattan plots showing significantly associated loci with fluid intelligence score when controlled for baseline covariates (red) and additionally with serum HDL (light blue) and serum glucose (violet) levels. The line indicates the exome-wide threshold.

**Supplementary Figure 2: Manhattan plots for mean reaction time**

Manhattan plots showing significantly associated loci with mean reaction time when controlled for baseline covariates (red), and additionally with serum HDL (light blue), serum LDL (blue), serum total cholesterol (green), serum triglyceride (pink), serum glucose (violet) and serum HbA1c (brown) levels. The line indicates the exome-wide threshold.

**Supplementary Figure 3: Manhattan plots for maximum symbol digit substitutions**

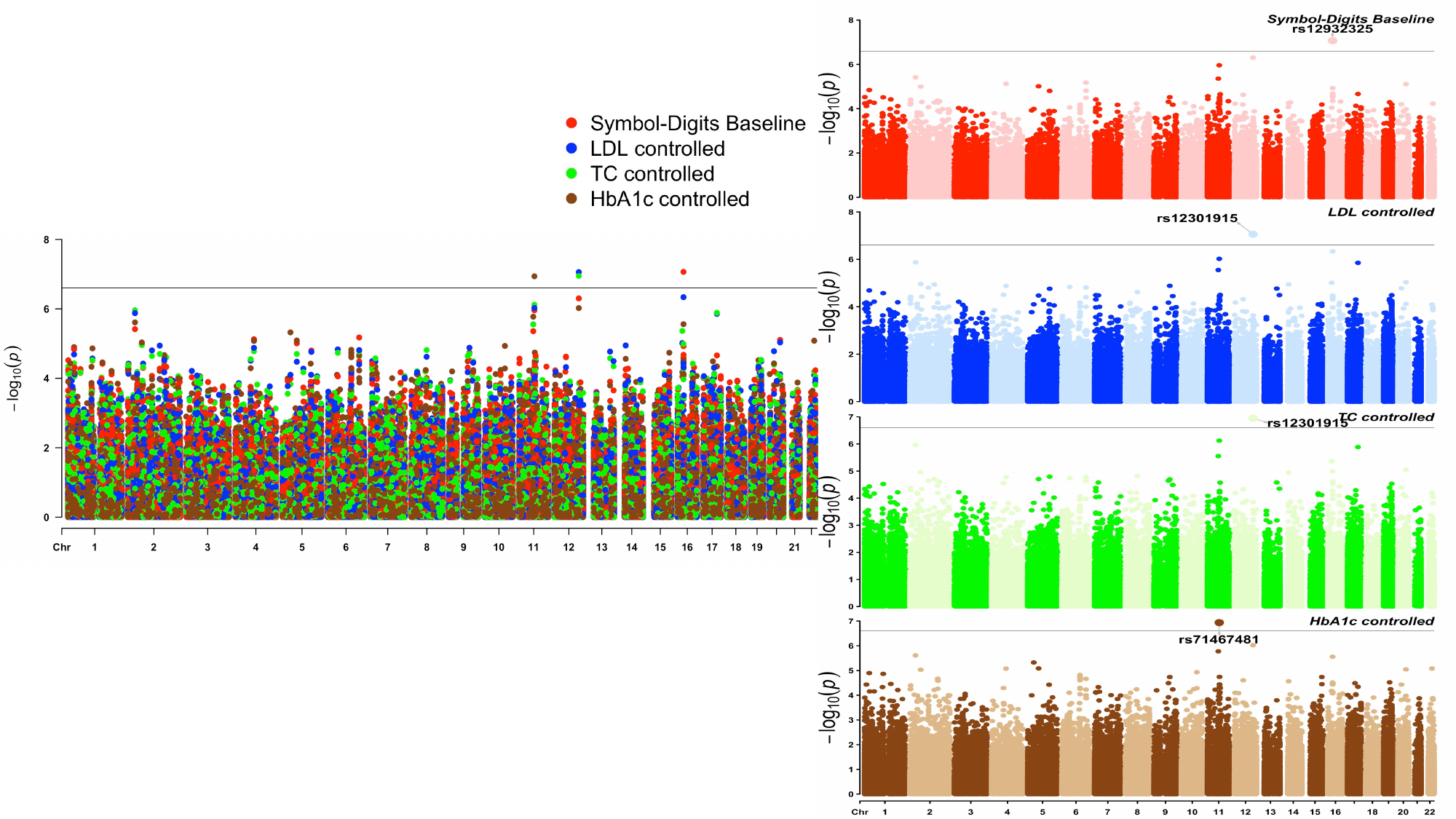

Manhattan plots showing significantly associated loci with maximum symbol digit substitutions when controlled for baseline covariates (red), and additionally with serum LDL (blue), serum total cholesterol (green), and serum HbA1c (brown) levels. The line indicates the exome-wide threshold.

**Supplementary Figure 4: Manhattan plots for proportion of incorrect matches**

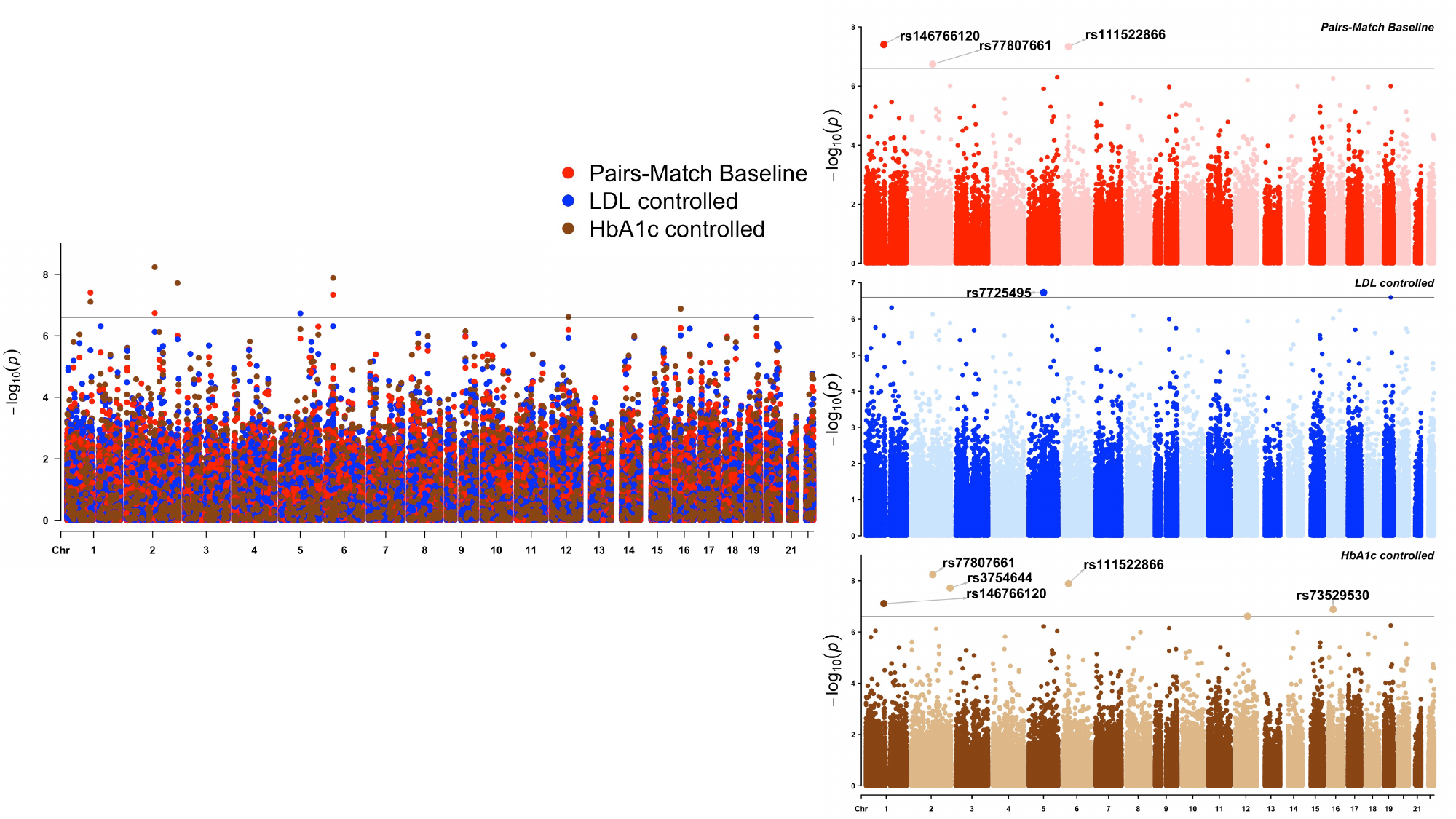

Manhattan plots showing significantly associated loci with proportion of incorrect matches when controlled for baseline covariates (red), and additionally with serum LDL (blue), and serum HbA1c (brown) levels. The line indicates the exome-wide threshold.

**Supplementary Figure 5: QQ-plots for fluid intelligence, mean reaction time, proportion of incorrect pair matches and maximum symbol-digits substitutions (from left to right) while controlling for covariates**

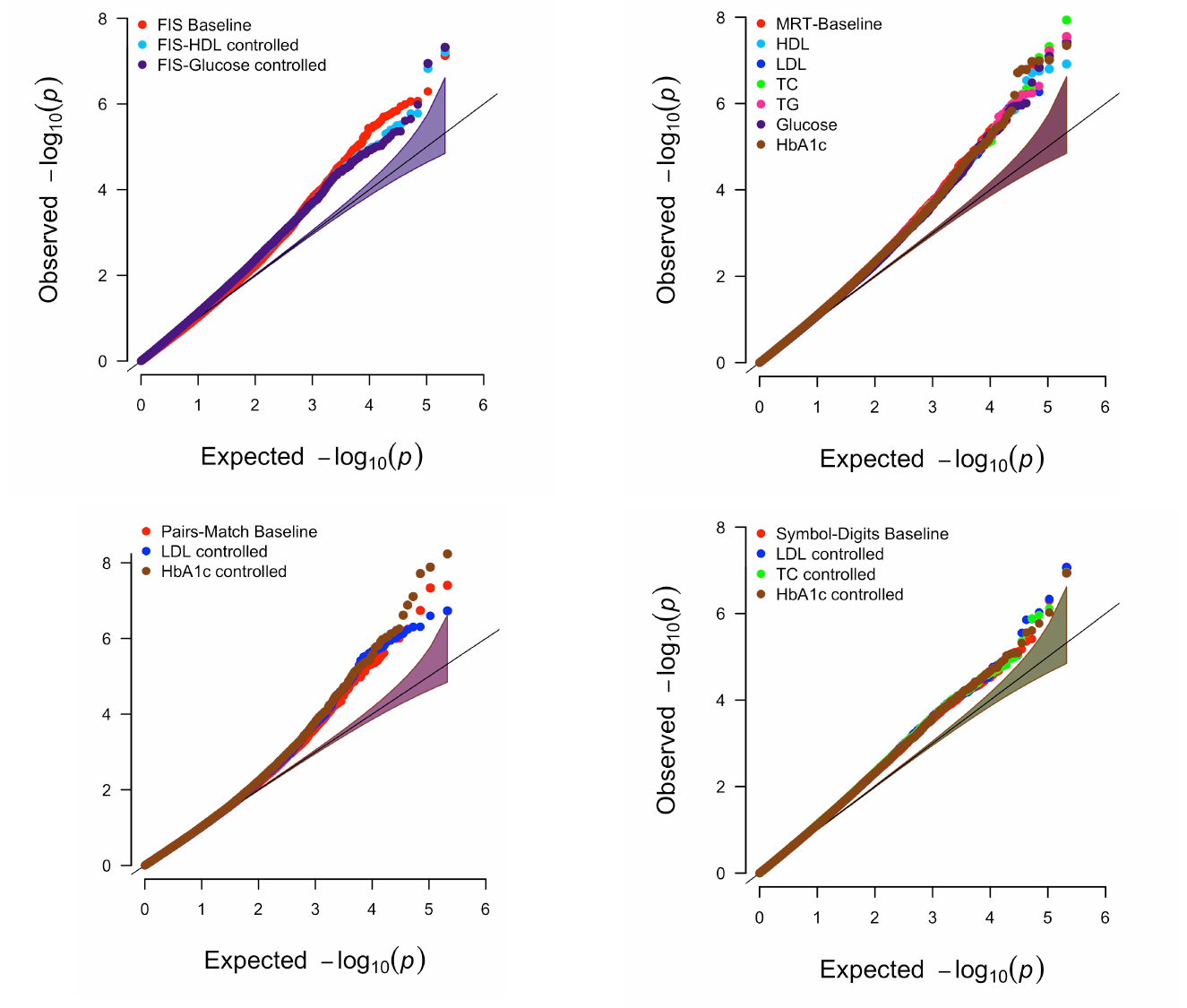

Adjustment with baseline covariates represented by red, additional metabolic covariates are represented as follows: HDL in sky blue, LDL in blue, total cholesterol (in green), triglyceride in pink, glucose in violet and HbA1c in brown.

**Supplementary Figure 6: QQ and Manhattan plots for alphanumeric trail duration while controlling for serum HDL levels**

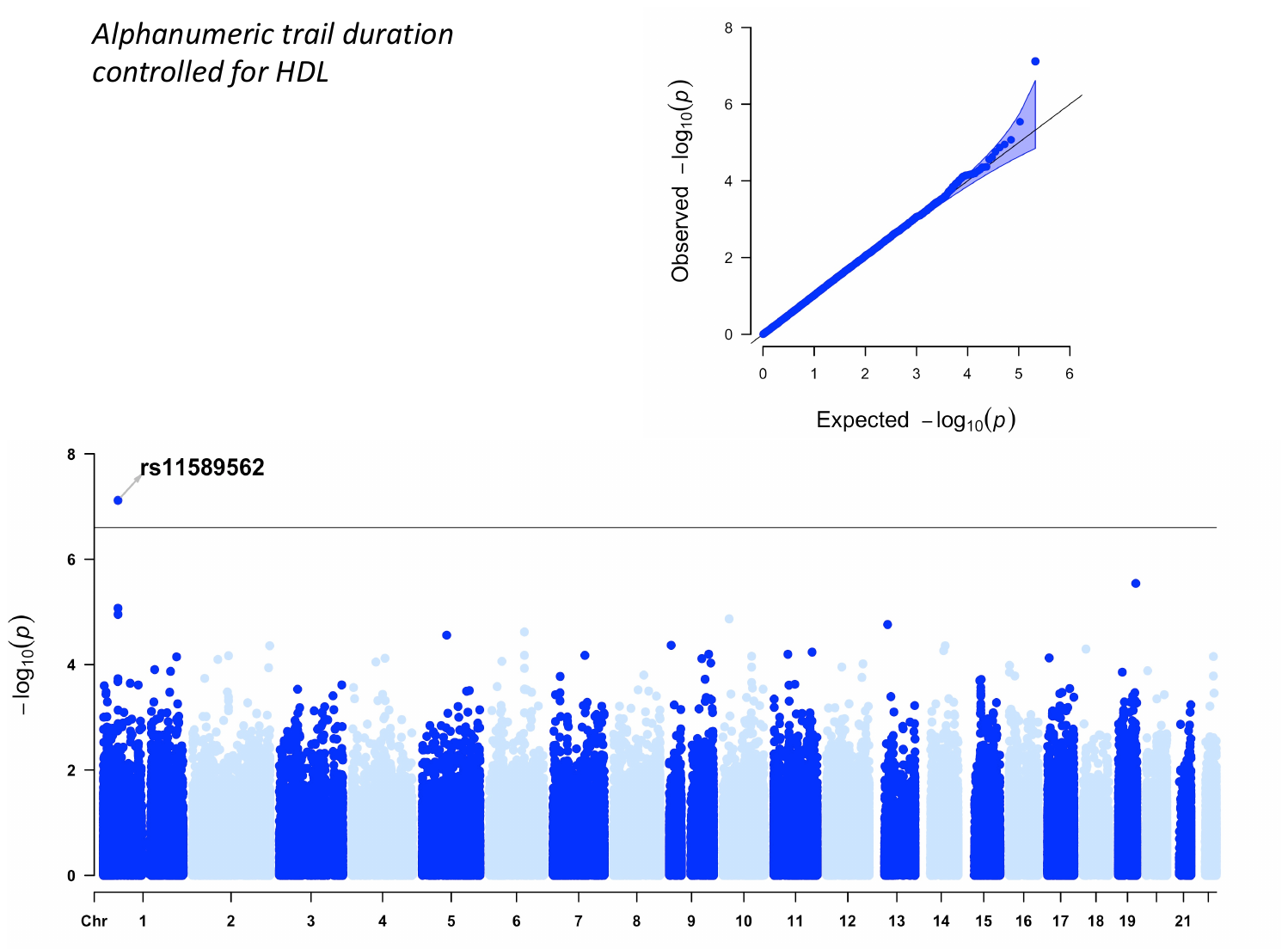

**Supplementary Figure 7 :Expression of *PON2* in the brain**

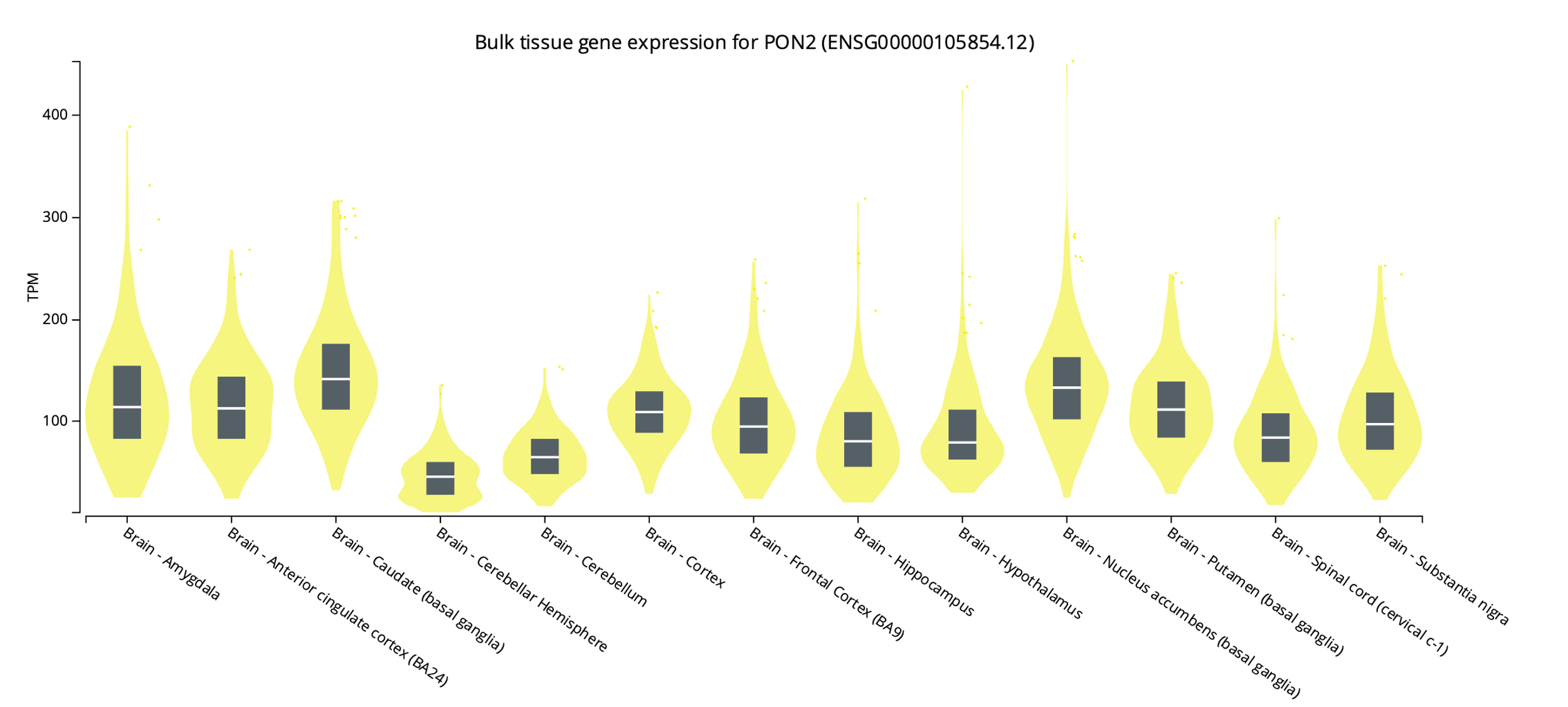

TPM stands for transcripts per million

**Supplementary Figure 8: Expression of *PPFIA1* in the brain**

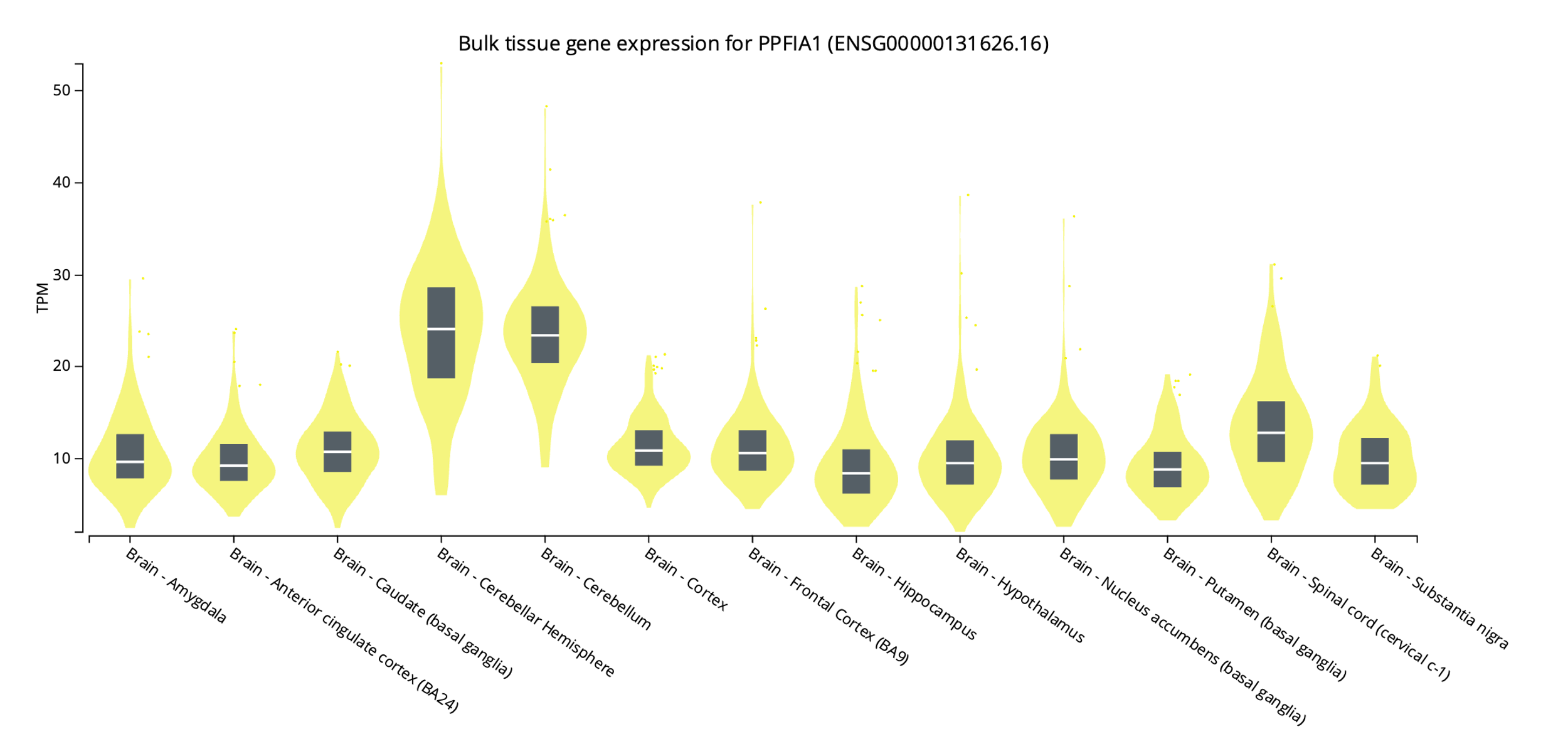

TPM stands for transcripts per million

**Supplementary Figure 9: Gene expression for *AMIGO1* in the brain**

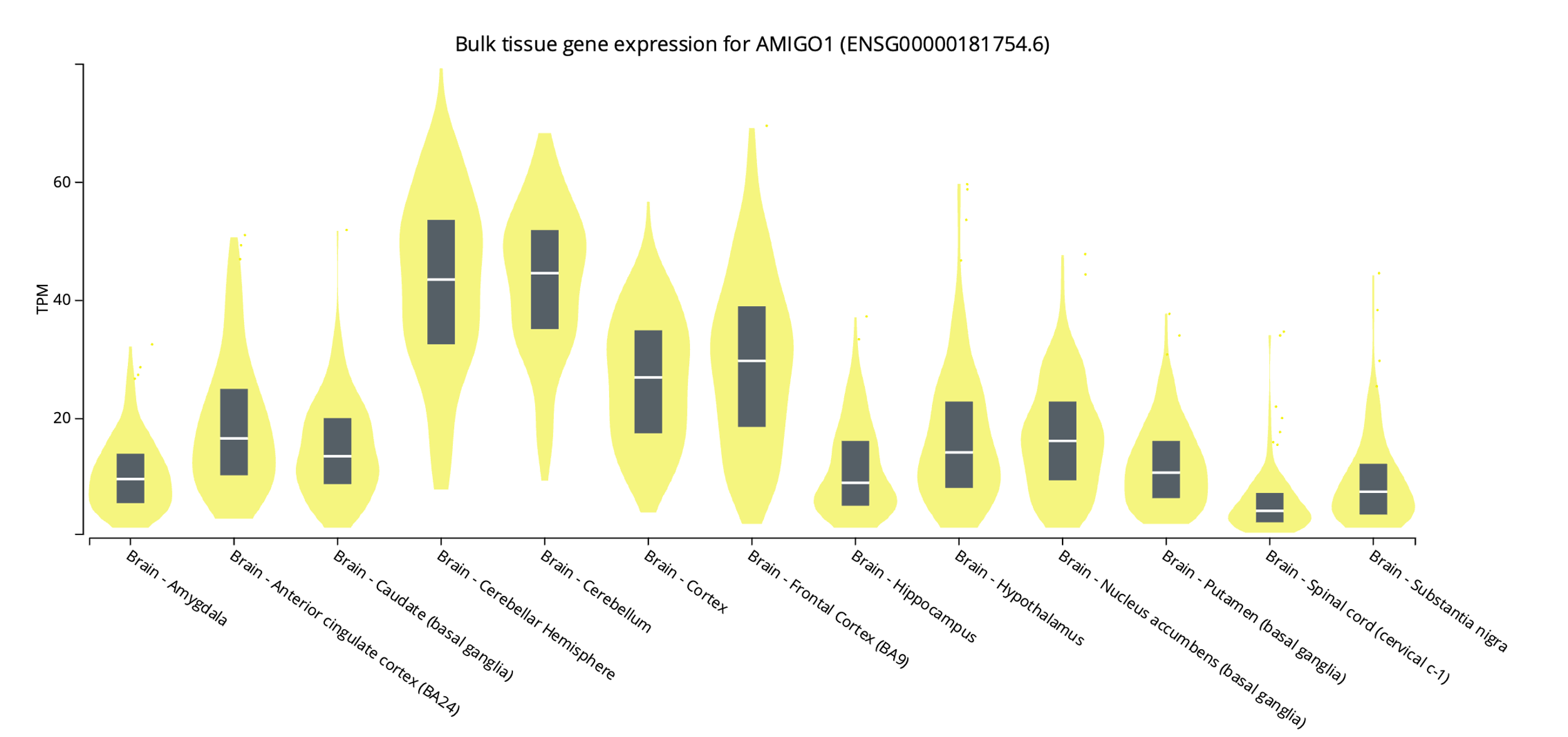
TPM stands for transcripts per million

**Supplementary Figure 10: Gene expression for *PTPN18* in the brain**

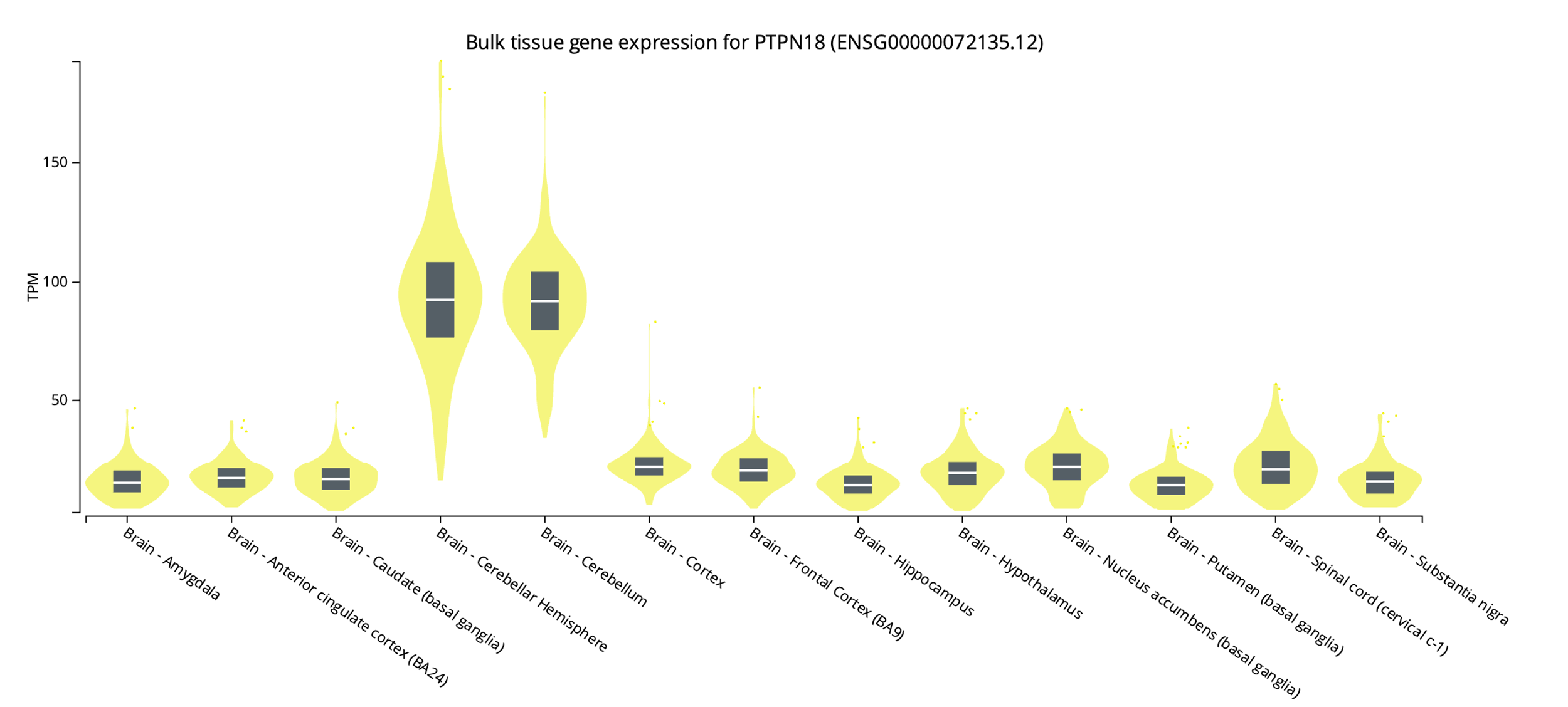

TPM stands for transcripts per million

**Supplementary Figure 11: Gene expression for *MAST2* in the brain**

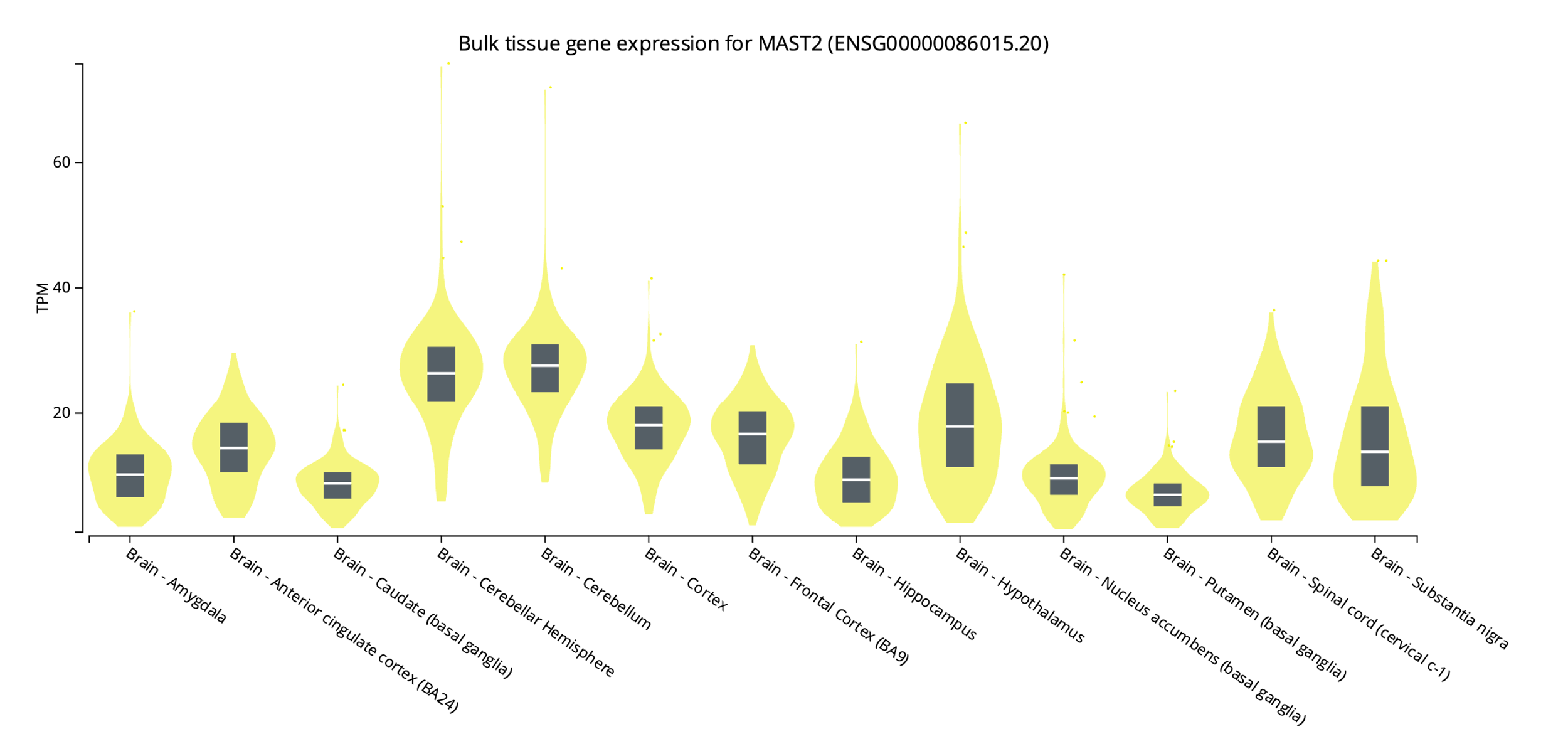

TPM stands for transcripts per million

**Supplementary Tables**

**Supplementary Table 1: Quality control steps for genotypic data**

| **Steps (thresholds for each step below detailed in methods)** | **Samples Excluded** | **Samples**  **Retained** | **Variants Excluded** | **Variants retained** |
| --- | --- | --- | --- | --- |
| 1. Incompatible blocks discarded-> Starting point |  | 200,624 |  | 13,567,285 |
| 2. Biallelic and sample half-calls set to missing |  | 200,624 | 1,426,145 | 12,141,140 |
| 3. Sample missingness | 13,050 | 187,574 | - | 12,141,140 |
| 4. Genotype missingness |  | 187,574 | 214,822 | 11,926,318 |
| 5. MAF |  | 187,574 | 11,687,242 | 239,076 |
| 6. LDAK (Outliers based on deviation from European Ancestry ) | 22,063 | 165,511 | - | 239,076 |
| 7. Participants who had withdrawn from the study | 2 | 165,509 |  | 239,076 |
| 8. Heterozygosity | 3,789 | 161,720 |  | 239,076 |
| 9. Hardy Weinberg Equilibrium |  | 161,720 | 27,942 | 211,134 |
| 10. Relatedness | 4,546 | 157,174 |  | 211,134 |
| **11.Variant Call Filters Exclusion Criteria** |  |  |  |  |
| Qual < 30 |  |  | 37 | 211,097 |
| QD > 20 |  | 157,174 | - | 211,097 |
| MMQ < 40 |  |  | - | 211,097 |
| Inbreeding Coeff < -0.8 |  |  | - | 211,097 |
| If >10% samples have SAMPLE DP<10, exclude the variant |  |  | - | 211,097 |
| retain only variants that meet either of the following criteria: (i) at least one homozygous variant carrier; or (ii) at least one heterozygous variant carrier with an allele balance (AB) greater than the cut-off (AB ≥ 0.15 for SNVs and AB ≥ 0.20 for indels). |  |  | 85 | 211,012 |
| 12. Further withdrawal | 14 | 157,160 | - | 211,012 |

**Supplementary Table 2: Comparison of SNP heritability estimates with previous studies**

| **Phenotype** | **Domain** | **Heritability estimates calculated controlling for**  **age and gender in our study** | **Heritability Estimates from previous family-based/twin studies** | [GWAS ATLAS estimates](https://atlas.ctglab.nl/)^21^ |
| --- | --- | --- | --- | --- |
| Fluid Intelligence Score | Fluid Intelligence | 42% | 58-60%*^22^; | 17.59%-20% |
| Mean Reaction Time | Simple Processing Speed | 6%*** | 38%**^23^; 35%**^24^ | 6.61% |
| Symbol Digit Matches | Complex Processing Speed | 25% | 50-60%^25^; 42%**^26^ | 11.09% |
| Proportion of Incorrect Matches | Episodic Memory | 7% | 30-60%*^27^ | 3.57% |
| Alphanumeric Trail Duration | Visual Attention | 23% | 39-65%**^26,28,29^; 22.4%**^28^ | 12.19% |
| Maximum Digits Remembered | Working Memory | 13% | 43.7%*^30^ | 7.3%-11.89% |

**Supplementary Table 3: *APOE* carrier status**

| ***APOE* Genotype** | **Number of Samples** | **APOE carrier status** |
| --- | --- | --- |
| E1/E1 | 0 |  |
| E1/E2 | 0 |  |
| E1/E3 | 3810 | Neutral |
| E1/E4 | 0 |  |
| E2/E2 | 946 | Protective/Beneficial |
| E2/E3 | 19103 | Protective/Beneficial |
| E2/E4 | 0 |  |
| E3/E3 | 92827 | Neutral |
| E3/E4 | 36810 | Risk |
| E4/E4 | 3571 | Risk |
| Missing | 93 |  |
| **Total** | **157160** |  |

**Supplementary Table 4: Single variant association analysis – detailed results**

| **Phenotype**  **/ Domain :** | **Fluid Intelligence Score (FIS)/ Reasoning, fluid intelligence cognitive domain** | | | | | | | | |
| --- | --- | --- | --- | --- | --- | --- | --- | --- | --- |
| **MODELS** | **Chr:Position ; rsid – REF/ALT**  **(ALT=Effect allele); Effect allele frequency** | **Informative Samples** | **Effect size: β (Effect Allele)** | **Standard Error (SE)** | **p-value** | **% of Variance Explained** | **Novelty for cognition** | **Genomic Annotation ;**  **Gene** | **CADD score (Phred scaled)**  **;**  **LofTool score** |
| **Baseline^a^** | chr5:141185287; rs115865641 –  G/A; 0.009 | 36457 | 0.036 | 0.006 | 7.33E-08 | 0.002% | Novel | 3' UTR variant ; *PCDHB16* | 3.05;  0.71 |
| **Baseline + HDL** | chr7:95406923; rs17876162 –  A/G; 0.001 | 31885 | 0.035 | 0.007 | 6.04E-08 | 3.16E-04% | Novel | Intron variant ;  *PON2* | 5.37;  0.82 |
|  | chr10:92240093; rs3824734 –  A/G; 0.594 | 31852 | -0.042 | 0.008 | 1.48E-07 | 0.086% | Novel | Synonymous variant ;  *CPEB3* | 11.74;  0.06 |
| **Baseline + Glucose** | chr7:95406923; rs17876162 –  A/G; 0.001 | 31852 | 0.036 | 0.007 | 4.75E-08 | 3.20E-04% | Novel | Intron variant;  *PON2s* | 5.37;  0.82 |
|  | chr10:92240093; rs3824734 –  A/G; 0.594 | 31852 | -0.043 | 0.008 | 1.13E-07 | 0.087% | Novel | Synonymous variant ;  *CPEB3* | 11.74;  0.06 |
| **Phenotype**  **/ Domain :** | **Mean time to identify matches / Simple processing speed cognitive domain** | | | | | | | | |
| **MODELS** | **Chr:Position ; rsid – REF/ALT**  **(ALT=Effect allele)** | **Informative Samples** | **Effect size: β (Effect Allele)** | **Standard Error (SE)** | **p-value** | **% of Variance Explained** | **Novelty for cognition** | **Genomic Annotation ;**  **Gene** | **CADD score (Phred scaled)**  **;**  **LofTool score** |
| **Baseline** | chr6:130365270; rs3813363 - C/T; 0.324 | 121127 | -0.024 | 0.004 | 5.91E-08 | 0.025% | - | 5' UTR variant;  *SAMD3* | 7.30;  0.94 |
|  | chr17:46171482 ; rs17662853 - G/A; 0.149 | 121127 | -0.025 | 0.005 | 1.04E-07 | 0.016% | - | Missense variant (p.Thr221Ile);  *KANSL1* | 23.90;  - |
|  | chr19:6732114 ; rs73922480 - C/T; 0.001 | 121127 | -0.584 | 0.106 | 3.66E-08 | 0.097% | Novel | Synonymous variant ;  *GPR108* | 4.16;  0.76 |
|  | chr19:6732194 ; rs77285514 - C/T; 0.002 | 121127 | -0.448 | 0.085 | 1.45E-07 | 0.066% | Novel | Intron variant ;  *GPR108* | 0.59;  0.76 |
| **Baseline + HDL** | chr6:130365270; rs3813363 - C/T; 0.324 | 105977 | -0.025 | 0.005 | 1.59E-07 | 0.026% | - | 5' UTR variant;  *SAMD3* | 7.30;  0.94 |
|  | chr12:440788 ; rs11062991 - G/T; 0.011 | 105977 | -0.109 | 0.021 | 1.78E-07 | 0.026% | Novel | Intron variant ;  *CCDC77* | 4.22;  1.00 |
|  | chr15:70891939; rs2959174 - G/T; 0.407 | 105977 | -0.022 | 0.004 | 1.96E-07 | 0.024% | Novel | Intron/Synonymous variant; *LRRC49/THAP10* | 2.75;  0.48 |
|  | chr17:46171482; rs17662853 - G/A; 0.149 | 105977 | -0.027 | 0.005 | 1.22E-07 | 0.018% | - | Missense variant (p.Thr221Ile) ; *KANSL1* | 23.90;  - |
| **Baseline + LDL** | chr6:130365270; rs3813363 - C/T; 0.324 | 115316 | -0.025 | 0.004 | 3.89E-08 | 0.027% | - | 5' UTR variant ;  *SAMD3* | 7.30;  0.94 |
|  | chr17:46171482 ; rs17662853 - G/A; 0.149 | 115316 | -0.026 | 0.005 | 9.70E-08 | 0.017% | - | Missense variant (p.Thr221Ile) ;  *KANSL1* | 23.90;  - |
| **Baseline + TC** | chr6:130365270; rs3813363 - C/T; 0.324 | 115520 | -0.024 | 0.004 | 4.79E-08 | 0.026% | - | 5' UTR variant ;  *SAMD3* | 7.30;  0.94 |
|  | chr15:70891939; rs2959174. - G/T; 0.407 | 115520 | -0.022 | 0.004 | 8.61E-08 | 0.024% | Novel | Intron/Synonymous variant ; *LRRC49/THAP10* | 2.75;  0.48 |
|  | chr17:46171482 ; rs17662853 - G/A; 0.149 | 115520 | -0.028 | 0.005 | 1.17E-08 | 0.019% | - | Missense variant (p.Thr221Ile) ;  *KANSL1* | 23.90;  - |
| **Baseline + TG** | chr6:130365270; rs3813363 - C/T; 0.324 | 115425 | -0.024 | 0.004 | 6.03E-08 | 0.026% | - | 5' UTR variant ;  *SAMD3* | 7.30;  0.94 |
|  | chr17:46171482 ; rs17662853 - G/A; 0.149 | 115425 | -0.027 | 0.005 | 2.83E-08 | 0.018% | - | Missense variant (p.Thr221Ile) ;  *KANSL1* | 23.90;  - |
| **Baseline + Glucose** | chr6:130365270; rs3813363 - C/T; 0.324 | 105898 | -0.025 | 0.005 | 8.10E-08 | 0.028% | - | 5' UTR variant ;  *SAMD3* | 7.30;  0.94 |
|  | chr12:440788 ; rs11062991 - G/T; 0.011 | 105898 | -0.110 | 0.021 | 1.46E-07 | 0.027% | Novel | Intron variant ;  *CCDC77* | 4.22;  1.00 |
|  | chr17:46171482 ; rs17662853 - G/A; 0.149 | 105898 | -0.028 | 0.005 | 4.27E-08 | 0.019% | - | Missense variant (p.Thr221Ile) ;  *KANSL1* | 23.90;  - |
| **Baseline + HbA1c** | chr1:25826774; rs201404149 - C/T; 0.001 | 115563 | -0.311 | 0.060 | 1.93E-07 | 0.027% | Novel | Synonymous variant ;  *MTFR1L* | 9.64;  - |
|  | chr6:130365270; rs3813363 - C/T; 0.324 | 115563 | -0.025 | 0.004 | 4.50E-08 | 0.026% | - | 5' UTR variant ;  *SAMD3* | 7.30;  0.94 |
|  | chr15:70832754; rs3825970 - G/A; 0.586 | 115563 | 0.022 | 0.004 | 1.64E-07 | 0.024% | Novel | Synonymous variant ;  *LARP6* | 0.895;  0.18 |
|  | chr15:70832865; rs1549317 - A/G; 0.589 | 115563 | 0.023 | 0.004 | 9.71E-08 | 0.025% | Novel | Synonymous variant ;  *LARP6* | 6.676;  0.18 |
|  | chr17:46171482 ; rs17662853 - G/A; 0.149 | 115563 | -0.025 | 0.005 | 1.64E-07 | 0.016% | - | Missense variant (p.Thr221Ile) ;  *KANSL1* | 23.90;  - |
|  | chr19:6732114 ; rs73922480 - C/T; 0.001 | 115563 | -0.595 | 0.112 | 1.03E-07 | 0.101% | Novel | Synonymous variant ;  *GPR108* | 4.16;  0.76 |
|  | chr19:6732194 ; rs77285514 - C/T; 0.002 | 115563 | -0.476 | 0.089 | 1.07E-07 | 0.074% | Novel | Intron variant ; *GPR108* | 0.59;  0.76 |
| **Phenotype**  **/ Domain :** | **Maximum correct symbol-digit substitutions / Complex processing speed cognitive domain** | | | | | | | | |
| **MODELS** | **Chr:Position ; rsid – REF/ALT**  **(ALT=Effect allele)** | **Informative Samples** | **Effect size: β (Effect Allele)** | **Standard Error (SE)** | **p-value** | **% of Variance Explained** | **Novelty for cognition** | **Genomic Annotation ;**  **Gene** | **CADD score (Phred scaled)**  **;**  **LofTool score** |
| **Baseline** | chr16:27462539; rs12932325 –  G/A; 0.142 | 35174 | -0.040 | 0.007 | 8.56E-08 | 0.039% | Novel | Intron variant;  *GTF3C1* | 0.88;  0.28 |
| **Baseline + LDL** | chr12:112482065; rs12301915-C/A; 0.013 | 33433 | 0.039 | 0.007 | 8.69E-08 | 0.004% | Novel | Intron variant;  *PTPN11* | 14.34;  0.05 |
| **Baseline + TC** | chr12:112482065; rs12301915-C/A; 0.013 | 33495 | 0.039 | 0.007 | 1.12E-07 | 0.004% | Novel | Intron variant;  *PTPN11* | 14.34;0.05 |
| **Baseline + HbA1c** | chr11:70324575;  rs71467481- G/A; 0.021 | 33503 | 0.030 | 0.006 | 1.15E-07 | 0.004% | Novel | Intron variant;  *PPFIA1* | 0.09;  0.23 |
| **Phenotype**  **/ Domain :** | **Proportion of incorrect pair matches / episodic memory domain** | | | | | | | | |
| **MODELS** | **Chr:Position ; rsid – REF/ALT**  **(ALT=Effect allele)** | **Informative Samples** | **Effect size: β (Effect Allele)** | **Standard Error (SE)** | **p-value** | **% of Variance Explained** | **Novelty for cognition** | **Genomic Annotation ;**  **Gene** | **CADD score (Phred scaled)**  **;**  **LofTool score** |
| **Baseline^a^** | chr1:109508840; rs146766120-C/T; 0.001 | 34120 | -1.080 | 0.180 | 3.93E-08 | 0.319% | Novel | Missense variant (p.Ala25Thr);  *AMIGO1* | 15.97;  - |
|  | chr2:130356125; rs77807661 - C/T; 0.002 | 34120 | -2.157 | 0.378 | 1.82E-07 | 1.711% | Novel | Synonymous variant;  *PTPN18* | 6.70;  0.41 |
|  | chr6:33688801; rs111522866-C/T; 0.002 | 34120 | -0.882 | 0.147 | 4.62E-08 | 0.243% | Novel | Intron variant;  *ITPR3* | 0.06;  0.05 |
| **Baseline + LDL^a^** | chr5:90502412; rs7725495 –  G/A; 0.002 | 32423 | -0.034 | 0.006 | 1.86E-07 | 3.74E-04% | Novel | Intron variant;  *POLR3G* | 0.59;  0.53 |
| **Baseline + HbA1c^a^** | chr1:109508840;rs146766120-C/T; 0.001 | 32494 | -1.091 | 0.186 | 7.80E-08 | 0.326% | Novel | Missense variant (p.Ala25Thr);  *AMIGO1* | 15.97;  - |
|  | chr2:130356125 ; rs77807661-C/T; 0.002 | 32494 | -2.598 | 0.408 | 5.81E-09 | 2.481% | Novel | Synonymous variant;  *PTPN18* | 6.70;  0.41 |
|  | chr2:236419198 ; rs3754644 –  T/C; 0.001 | 32494 | -0.971 | 0.158 | 1.92E-08 | 0.235% | Novel | Missense variant (p.Gln369Arg);  *IQCA1* | 7.61;  - |
|  | chr6:33688801; rs111522866 –  C/T; 0.002 | 32494 | -0.927 | 0.149 | 1.30E-08 | 0.269% | Novel | Intron variant;  *ITPR3* | 0.06;  0.05 |
|  | chr16:28904132; rs73529530-T/C; 0.003 | 32494 | -0.044 | 0.008 | 1.31E-07 | 0.001% | Novel | Intron variant;  *ATP2A1* | 0.08;  0.08 |
| **Phenotype**  **/ Domain :** | **Duration of alphanumeric trail / Visual Attention domain** | | | | | | | | |
| **MODELS** | **Chr:Position ; rsid – REF/ALT**  **(ALT=Effect allele)** | **Informative Samples** | **Effect size: β (Effect Allele)** | **Standard Error (SE)** | **p-value** | **% of Variance Explained** | **Novelty for cognition** | **Genomic Annotation ;**  **Gene** | **CADD score (Phred scaled)**  **;**  **LofTool score** |
| **Baseline + HDL^a^** | chr1:46001049; rs11589562-  T/C; 0.419 | 27214 | -0.050 | 0.009 | 7.64E-08 | 0.122% | Novel | Intron variant;  *MAST2* | 7.84;  0.84 |

a: GC corrected models

chr: Chromosome

β: Effect size of the association and SE: standard error of β

CADD score: Combined Annotation Dependent Depletion score evaluates the deleteriousness of the variants. Higher the CADD score, more deleterious is the variant.

Loftool score: A gene intolerance score based on loss of function variants. A lower score indicates more intolerance to functional variation.

We note that the effect allele given in our single variant association allele is not always the risk allele for cognitive decline. The direction of association has different implication on cognition for different phenotypes. For example, when the direction of association of the effect allele is negative for average trail duration, mean reaction time and proportion of incorrect matches, the effect allele is actually the protective allele for cognitive decline, whereas a positive association implies that the effect allele is the risk increasing allele for cognitive decline. Similarly, negative association of effect allele with fluid intelligence score as well as with maximum symbol-digit substitutions imply that the effect allele is in essence the risk allele affecting cognition in their corresponding domains and vice versa.

**Supplementary Table 5: Gene-based association results**

| **Domain** | **Phenotype** | **Covariate** | **Gene** | **N_INFO** | **SKATO-rho** | **P-value** | **Comment** |
| --- | --- | --- | --- | --- | --- | --- | --- |
| Complex Processing Speed | Maximum Symbol-digit substitutions | Baseline | *APOC1*^a^ | 35174 | 1 | 0.0013 | Burden |
|  |  | Baseline+HDL | *-* | - | - | - | - |
|  |  | Baseline+LDL | *APOC1*^a^ | 33433 | 1 | 0.0013 | Burden |
|  |  | Baseline+TC | *APOC1*^a^ | 33495 | 1 | 0.0011 | Burden |
|  |  | Baseline+TG | *APOC1*^a^ | 33468 | 1 | 0.0015 | Burden |
|  |  | Baseline+HbA1c | *APOC1* ^a^ | 33503 | 1 | 0.0027 | Burden |
| Visual Attention | Alphanumeric Trail Duration | Baseline | *APOC1* ^a^ | 31200 | 1 | 0.0021 | Burden |
|  |  | Baseline+HDL | *LRP1*^b^ | 27214 | 0 | 0.0018 | SKAT |
|  |  | Baseline+LDL | *APOC1* ^a^ | 29660 | 1 | 0.0013 | Burden |
|  |  | Baseline+TC | *APOC1* ^a^ | 29715 | 1 | 0.0011 | Burden |
|  |  | Baseline+TG | *APOC1* ^a^ | 29692 | 1 | 0.0018 | Burden |
|  |  | Baseline+HbA1c | APOC1 ^a^ | 29730 | 1 | 0.0013 | Burden |

‘a’:SKAT-O

‘b’:SKAT

N_INFO: Number of informative samples

SKATO-rho: Εstimate of ρ used in SKATO test statistic; ρ=0 implies burden test has been performed and ρ=1 implies SKAT test has been performed.

**Supplementary Table 6: Bivariate association and mediation for single variant hits for fluid intelligence**

| **DOMAIN:** | **Fluid Intelligence** | | | |
| --- | --- | --- | --- | --- |
|  | **Chr:Pos;**  **rsid**  **(Mapped Gene)** | **chr5:141185287;**  **rs115865641**  **(*PCDHB16*)** | **chr7:95406923;**  **rs17876162**  **(*PON2*)** | **chr10:92240093;**  **rs3824734 (*CPEB3*)** |
| **Covariates controlled** | Baseline | β = 0.036;  p = 7.33E-08 | β = 0.03  p = 1.99E-06^a^ | - |
|  | Baseline + HDL | β = 0.029;  p = 8.58E-06^a^ | β = 0.035  p = 6.04E-08 | β = -0.042  p = 1.48E-07 |
|  | Baseline + LDL | - | - | - |
|  | Baseline + TC | - | - | - |
|  | Baseline + TG | - | - | - |
|  | Baseline + Glucose | β = 0.029;  p = 9.44E-06^a^ | β = 0.036  p = 4.75E-08 | β = -0.043  p = 1.13E-07 |
|  | Baseline + HbA1c | - | - | - |
| **Associated to** | HDL | β = 0.026;  p = 1.78E-11 | - | - |
|  | LDL | - | - | - |
|  | TC | - | - | - |
|  | TG | - | - | - |
|  | Glucose | β = 0.042;  p = 2.11E-27 | - | - |
|  | HbA1c | - | - | - |
| **Bivariate association of cognitive phenotype with** | HDL | p = 2.53E-15 | - | - |
|  | LDL | p = 5.26E-11 | p = 6.58E-08 | - |
|  | TC | p = 3.43E-09 | - | - |
|  | TG | p = 3.38E-12 | - | - |
|  | Glucose | p = 7.63E-30 | p = 1.16E-08 | - |
|  | HbA1c | p = 1.73E-08 | - | - |

‘-‘:Insignificant association; ‘a’: Nearly significant (of order E-06); ‘b’: Suggestive (order E-05)

β : Effect size of the respective associations; p : p-value

The “Covariates controlled” section contains repeated information from supplementary table 4 for clarity. And lipid, glucose associations are appearing only in this table under “Associated to” section. Results from our bivariate analysis have also been represented here.

**Supplementary Table 7: Bivariate association and mediation for single variant hits for simple processing speed**

| **DOMAIN:** | **Simple processing speed** | | | | | |
| --- | --- | --- | --- | --- | --- | --- |
|  | **Chr:Pos;**  **rsid**  **(Mapped Gene)** | **chr1:25826774;**  **rs201404149 (*MTFR1L*)** | **chr6:13036527;**  **rs3813363 (*SAMD3*)** | **chr12:440788;**  **rs11062991 (*CCDC77*)** | **chr15:70832754;**  **rs3825970 (*LARP6*)** | **chr15:70832865;**  **rs1549317 (*LARP6*)** |
| **Covariates controlled** | Baseline | β = -0.289  p = 7.69E-07^a^ | β = -0.024  p = 5.91E-08 | - | - | - |
|  | Baseline+ HDL | - | β = -0.025  p = 1.59E-07 | β = -0.109  p = 1.78E-07 | - | - |
|  | Baseline+ LDL | - | β = -0.025  p = 3.89E-08 | - | - | - |
|  | Baseline+ TC | - | β = -0.024  p = 4.79E-08 | - | - | - |
|  | Baseline+ TG | - | β = -0.024  p = 6.03E-08 | - | - | - |
|  | Baseline+  Glucose | β = -0.251  p = 5.13E-05^b^ | β = -0.025  p = 8.10E-08 | β = -0.110  p = 1.46E-07 | - | - |
|  | Baseline+  HbA1c | β = -0.311  p = 1.93E-07 | β = -0.025  p = 4.50E-08 | - | β = 0.022  p = 1.64E-07 | β = 0.023  p = 9.71E-08 |
| **Associated to** | HDL | - | - | - | - | - |
|  | LDL | - | - | - | - | - |
|  | TC | - | - | - | - | - |
|  | TG | - | - | - | - | - |
|  | Glucose | β = -0.279  p = 4.77E-06 | - | - | - | - |
|  | HbA1c | - | - | - | - | - |
| **Bivariate association of cognitive phenotype with** | HDL | - | - | - | - | - |
|  | LDL | - | p = 6.11E-08 | - | - | - |
|  | TC | - | p = 7.47E-08 | - | - | - |
|  | TG | - | p = 1.32E-08 | - | - | - |
|  | Glucose | p = 4.91E-10 | p = 3.98E-08 | - | p = 7.67E-08 | p = 5.99E-08 |
|  | HbA1c | - | - | - | - | - |

‘-‘:Insignificant association; ‘a’: Nearly significant (of order E-06); ‘b’: Suggestive (order E-05)

β : Effect size of the respective associations; p : p-value

The “Covariates controlled” section contains repeated information from Supplementary Table 4 for clarity. And lipid, glucose associations are appearing only in this table under “Associated to” section. Results from our bivariate analysis have also been represented here.

**Supplementary Table 8: Bivariate association and mediation for single variant hits for simple processing speed (continued)**

| **DOMAIN:** | **Simple Processing Speed** | | | | |
| --- | --- | --- | --- | --- | --- |
|  | **Chr:Pos;**  **rsid**  **(Mapped Gene)** | **chr15:70891939;**  **rs2959174 (*LRRC49/***  ***THAP10*)** | **chr17:46171482;**  **rs17662853 (*KANSL1*)** | **chr19:6732114;**  **rs73922480 (*GPR108*)** | **chr19:6732194;**  **rs77285514**  **(*GPR108*)** |
| **Covariates controlled** | Baseline | - | β = -0.025  p = 1.04E-07 | β = -0.584  p = 3.66E-08 | β = -0.448  p = 1.45E-07 |
|  | Baseline+ HDL | β = -0.022  p = 1.96E-07 | β = -0.027  p = 1.22E-07 | - | - |
|  | Baseline+ LDL | - | β = -0.026  p = 9.70E-08 | - | - |
|  | Baseline+ TC | β = -0.022  p = 8.61E-08 | β = -0.028  p = 1.17E-08 | - | - |
|  | Baseline+ TG | - | β = -0.027  p = 2.83E-08 | - | - |
|  | Baseline+  Glucose | - | β = -0.028  p = 4.27E-08 | - | - |
|  | Baseline+  HbA1c | - | β = -0.025  p = 1.64E-07 | β = -0.595  p = 1.03E-07 | β = -0.476  p = 1.07E-07 |
| **Associated to** | HDL | - | - | - | - |
|  | LDL | - | - | - | - |
|  | TC | - | - | - | - |
|  | TG | - | - | - | - |
|  | Glucose | - | - | - | - |
|  | HbA1c | - | - | - | - |
| **Bivariate association of cognitive phenotype with** | HDL | - | - | p = 1.01E-07 | - |
|  | LDL | - | - | - | - |
|  | TC | - | - | - | - |
|  | TG | - | - | p = 1.11E-07 | - |
|  | Glucose | - | - | - | - |
|  | HbA1c | - | p = 1.63E-10 | p = 3.20E-08 | - |

‘-‘:Insignificant association; ‘a’: Nearly significant (of order E-06); ‘b’: Suggestive (order E-05)

β : Effect size of the respective associations; p : p-value

The “Covariates controlled” section contains repeated information from Supplementary Table 4 for clarity. And lipid, glucose associations are appearing only in this table under “Associated to” section. Results from our bivariate analysis have also been represented here.

**Supplementary Table 9: Bivariate association and mediation for single variant hits for complex processing speed**

| **DOMAIN:** | **Complex Processing Speed** | | | |
| --- | --- | --- | --- | --- |
|  | **Variant**  **(Chr:Pos; rsid (Mapped Gene)** | **chr11:70324575;**  **rs71467481 (*PPFIA1*)** | **chr12:112482065;**  **rs12301915 (*PTPN11*)** | **chr16:27462539;**  **rs12932325 (*GTF3C1*)** |
| **Covariates controlled** | Baseline | β = 0.027;  p = 1.10E-06^a^ | β = 0.036;  p = 5.00E-07^a^ | β = -0.040  p = 8.56E-08 |
|  | Baseline + HDL | - | β = 0.038;  p = 5.77E-07^a^ | - |
|  | Baseline + LDL | - | β = 0.039  p = 8.69E-08 | - |
|  | Baseline + TC | - | β = 0.039  p = 1.12E-07 | - |
|  | Baseline + TG | - | - | - |
|  | Baseline + Glucose | β = 0.026;  p = 7.65E-06^a^ | β = 0.037;  p = 1.08E-06^a^ | - |
|  | Baseline + HbA1c | β = 0.030  p = 1.15E-07 | - | - |
| **Associated to** | HDL | β = 0.013  9.38E-05^b^ | β = -0.018  p = 1.26E-07 | - |
|  | LDL | - | β = -0.018  3.43E-05^b^ | - |
|  | TC | - | - | - |
|  | TG | - | β = -0.019  p = 8.01E-06^a^ | - |
|  | Glucose | β = 0.025  p = 7.26E-15 | β = 0.042  p = 1.73E-20 | - |
|  | HbA1c | - | - | - |
| **Bivariate association of cognitive phenotype with** | HDL | p = 3.86E-08 | p = 8.46E-11 | - |
|  | LDL | p = 3.71E-08 | p = 3.84E-09 | - |
|  | TC | - | - | - |
|  | TG | - | p = 2.38E-09 | - |
|  | Glucose | p = 1.90E-17 | p = 5.55E-23 | - |
|  | HbA1c | - | - | - |

‘-‘:Insignificant association; ‘a’: Nearly significant (of order E-06) ;‘b’: Suggestive (order E-05)

β : Effect size of the respective associations; p : p-value

The “Covariates controlled” section contains repeated information from Supplementary Table 4 for clarity. And lipid, glucose associations are appearing only in this table under “Associated to” section. Results from our bivariate analysis have also been represented here.

**Supplementary Table 10: Bivariate association and mediation for single variant hits for episodic memory**

| **DOMAIN:** | **Episodic Memory** | | | | | | |
| --- | --- | --- | --- | --- | --- | --- | --- |
|  | **Variant**  **(Chr:Pos; rsid (Mapped Gene)** | **chr1:**  **109508840;**  **;rs146766120 (*AMIGO1*)** | **chr2:**  **130356125; rs77807661 (*PTPN18*)** | **chr2:**  **236419198**  **; rs3754644 (*IQCA1)*** | **chr5:**  **90502412**  **; rs7725495**  **(*POLR3G*)** | **chr6:**  **33688801;**  **rs111522866 (*ITPR3*)** | **chr16:**  **28904132;**  **rs73529530 (*ATP2A1*)** |
| **Covariates controlled** | Baseline | β = -1.080  p = 3.93E-08 | β = -2.157  p = 1.82E-07 | - | β = -0.031  p = 1.24E-06^a^ | β = -0.882  p = 4.62E-08 | β = -0.040;  p = 5.59E-07^a^ |
|  | Baseline+ HDL | - | - | - | β = -0.031  p = 3.72E-06^a^ | - | β = -0.038;  p =1.31E-07^a^ |
|  | Baseline+ LDL | - | - | - | β = -0.034  p = 1.86E-07 | - | - |
|  | Baseline+ TC | - | - | - | - | - | - |
|  | Baseline+ TG | - | - | - | β = -0.032  p = 7.44E-07^a^ | - | - |
|  | Baseline+  Glucose | - | - | - | β = -0.031  p = 4.20E-06^a^ | β = -0.886;  p = 8.68E-07^a^ | β = -0.037;  p = 1.40E-05^b^ |
|  | Baseline+  HbA1c | β = -1.091  p = 7.80E-08 | β = -2.598  p = 5.81E-09 | β = -0.971  p =1.92E-08 | - | β = -0.927  p = 1.30E-08 | β = -0.044  p = 1.31E-07 |
| **Associated to** | HDL | - | - | - | β = 0.019  p = 6.67E-08 | - | β = 0.022  p = 2.03E-06^a^ |
|  | LDL | - | - | - | β= -0.015  p = 7.72E-06^a^ | - | - |
|  | TC | - | - | - | - | - | - |
|  | TG | - | - | - | β = -0.016  p = 2.18E-06^a^ | - | - |
|  | Glucose | - | - | - | β = 0.038  p = 3.32E-27 | β = -0.296  p = 1.11E-05^b^ | β = 0.039  p =2.38E-17 |
|  | HbA1c | - | - | - | - | - | - |
| **Bivariate association of cognitive phenotype with** | HDL | p = 1.35E-08 | p = 1.17E-08 | - | p = 2.18E-11 | p = 1.13E-08 | p = 1.60E-10 |
|  | LDL | p = 1.32E-08 | p = 7.65E-08 | - | p = 4.46E-10 | p = 1.29E-08 | p = 4.67E-09 |
|  | TC | p = 1.30E-08 | p = 8.35E-08 | - | p = 3.13E-08 | p = 1.46E-08 | p = 6.93E-08 |
|  | TG | p = 1.41E-08 | p = 6.64E-08 | - | p = 3.81E-10 | p = 2.61E-09 | p = 4.96E-08 |
|  | Glucose | p = 2.05E-11 | p = 2.30E-10 | p = 1.47E-08 | p = 7.36E-31 | p = 2.21E-13 | p = 9.55E-22 |
|  | HbA1c | p = 7.97E-11 | p = 3.72E-08 | p = 4.72E-08 | - | p = 1.62E-09 | - |

‘-‘:Insignificant association ;‘a’: Nearly significant (of order E-06) ; ‘b’: Suggestive (order E-05); β : Effect size of the respective associations; p : p-value

The “Covariates controlled” section contains repeated information from Supplementary Table 4 for clarity. And lipid, glucose associations are appearing only in this table under “Associated to” section. Results from our bivariate analysis have also been represented here.

**References:**

1.	Yang J, Lee SH, Goddard ME, Visscher PM. GCTA: a tool for genome-wide complex trait analysis. *Am J Hum Genet*. 2011;88(1):76-82. doi:10.1016/j.ajhg.2010.11.011

2.	Backman JD, Li AH, Marcketta A, et al. Exome sequencing and analysis of 454,787 UK Biobank participants. *Nature*. 2021;599(7886):628-634. doi:10.1038/s41586-021-04103-z

3.	Zhao Y, Stankovic S, Koprulu M, et al. GIGYF1 loss of function is associated with clonal mosaicism and adverse metabolic health. *Nat Commun*. 2021;12(1):4178. doi:10.1038/s41467-021-24504-y

4.	Speed D, Holmes J, Balding DJ. Evaluating and improving heritability models using summary statistics. *Nat Genet*. 2020;52(4):458-462. doi:10.1038/s41588-020-0600-y

5.	Zhao H, Sun Z, Wang J, Huang H, Kocher JP, Wang L. CrossMap: a versatile tool for coordinate conversion between genome assemblies. *Bioinformatics*. 2014;30(7):1006-1007. doi:10.1093/bioinformatics/btt730

6.	Speed D, Hemani G, Johnson MR, Balding DJ. Improved Heritability Estimation from Genome-wide SNPs. *The American Journal of Human Genetics*. 2012;91(6):1011-1021. doi:10.1016/j.ajhg.2012.10.010

7.	Troutwine BR, Hamid L, Lysaker CR, Strope TA, Wilkins HM. Apolipoprotein E and Alzheimer’s disease. *Acta Pharm Sin B*. 2022;12(2):496-510. doi:10.1016/j.apsb.2021.10.002

8.	Long JM, Holtzman DM. Alzheimer Disease: An Update on Pathobiology and Treatment Strategies. *Cell*. 2019;179(2):312-339. doi:https://doi.org/10.1016/j.cell.2019.09.001

9.	Lane RM, Farlow MR. Lipid homeostasis and apolipoprotein E in the development and progression of Alzheimer’s disease. *J Lipid Res*. 2005;46(5):949-968. doi:10.1194/jlr.M400486-JLR200

10.	Schaefer EJ, Bongard V, Beiser AS, et al. Plasma Phosphatidylcholine Docosahexaenoic Acid Content and Risk of Dementia and Alzheimer Disease: The Framingham Heart Study. *Arch Neurol*. 2006;63(11):1545-1550. doi:10.1001/archneur.63.11.1545

11.	An Y, Varma VR, Varma S, et al. Evidence for brain glucose dysregulation in Alzheimer’s disease. *Alzheimers Dement*. 2018;14(3):318-329. doi:10.1016/j.jalz.2017.09.011

12.	Henderson ST. High carbohydrate diets and Alzheimer’s disease. *Med Hypotheses*. 2004;62(5):689-700. doi:https://doi.org/10.1016/j.mehy.2003.11.028

13.	Zhang J, Liu Q. Cholesterol metabolism and homeostasis in the brain. *Protein Cell*. 2015;6(4):254-264. doi:10.1007/s13238-014-0131-3

14.	Leduc V, Jasmin-Bélanger S, Poirier J. APOE and cholesterol homeostasis in Alzheimer’s disease. *Trends Mol Med*. 2010;16(10):469-477. doi:https://doi.org/10.1016/j.molmed.2010.07.008

15.	Yamazaki Y, Zhao N, Caulfield TR, Liu CC, Bu G. Apolipoprotein E and Alzheimer disease: pathobiology and targeting strategies. *Nat Rev Neurol*. 2019;15(9):501-518. doi:10.1038/s41582-019-0228-7

16.	Marques F, Sousa JC, Sousa N, Palha JA. Blood–brain-barriers in aging and in Alzheimer’s disease. *Mol Neurodegener*. 2013;8(1):38. doi:10.1186/1750-1326-8-38

17.	Consortium TU. UniProt: the universal protein knowledgebase in 2021. *Nucleic Acids Res*. 2021;49(D1):D480-D489. doi:10.1093/nar/gkaa1100

18.	Wu MC, Lee S, Cai T, Li Y, Boehnke M, Lin X. Rare-variant association testing for sequencing data with the sequence kernel association test. *Am J Hum Genet*. 2011;89(1):82-93. doi:10.1016/j.ajhg.2011.05.029

19.	Davies RB. Algorithm AS 155: The Distribution of a Linear Combination of χ 2 Random Variables. *Appl Stat*. 1980;29(3):323. doi:10.2307/2346911

20.	Lee S, Emond MJ, Bamshad MJ, et al. Optimal unified approach for rare-variant association testing with application to small-sample case-control whole-exome sequencing studies. *Am J Hum Genet*. 2012;91(2):224-237. doi:10.1016/j.ajhg.2012.06.007

21.	Watanabe K, Stringer S, Frei O, et al. A global overview of pleiotropy and genetic architecture in complex traits. *Nat Genet*. 2019;51(9):1339-1348. doi:10.1038/s41588-019-0481-0

22.	Cattell RB. The heritability of fluid, gf, and crystallised, gc, intelligence, estimated by a least squares use of the MAVA method. *British Journal of Educational Psychology*. 1980;50(3):253-265. doi:10.1111/j.2044-8279.1980.tb00809.x

23.	KUNTSI J, ROGERS H, SWINARD G, et al. Reaction time, inhibition, working memory and ‘delay aversion’ performance: genetic influences and their interpretation. *Psychol Med*. 2006;36(11):1613-1624. doi:10.1017/S0033291706008580

24.	Finkel D, Pedersen NL. Genetic and environmental contributions to the associations between intraindividual variability in reaction time and cognitive function. *Aging, Neuropsychology, and Cognition*. 2014;21(6):746-764. doi:10.1080/13825585.2013.874523

25.	Pahlen S, Hamdi NR, Dahl Aslan AK, et al. Age-moderation of genetic and environmental contributions to cognitive functioning in mid- and late-life for specific cognitive abilities. *Intelligence*. 2018;68:70-81. doi:10.1016/j.intell.2017.12.004

26.	Knowles EEM, Carless MA, de Almeida MAA, et al. Genome-wide significant localization for working and spatial memory: Identifying genes for psychosis using models of cognition. *American Journal of Medical Genetics Part B: Neuropsychiatric Genetics*. 2014;165(1):84-95. doi:10.1002/ajmg.b.32211

27.	Papassotiropoulos A, de Quervain DJF. Genetics of human episodic memory: dealing with complexity. *Trends Cogn Sci*. 2011;15(9):381-387. doi:10.1016/j.tics.2011.07.005

28.	Hagenaars SP, Cox SR, Hill WD, et al. Genetic contributions to Trail Making Test performance in UK Biobank. *Mol Psychiatry*. 2018;23(7):1575-1583. doi:10.1038/mp.2017.189

29.	Vasilopoulos T, Franz CE, Panizzon MS, et al. Genetic architecture of the Delis-Kaplan executive function system Trail Making Test: Evidence for distinct genetic influences on executive function. *Neuropsychology*. 2012;26(2):238-250. doi:10.1037/a0026768

30.	Jensen AR, Marisi DQ. A note on the heritability of memory span. *Behav Genet*. 1979;9(5):379-387. doi:10.1007/BF01066976
